# Deep quantitative glycoproteomics reveals gut microbiome induced remodeling of the brain glycoproteome

**DOI:** 10.1101/2023.09.13.557529

**Authors:** Clément M Potel, Mira Lea Burtscher, Martin Garrido-Rodriguez, Amber Brauer-Nikonow, Isabelle Becher, Athanasios Typas, Michael Zimmermann, Mikhail M Savitski

**Affiliations:** Genome Biology Unit, European Molecular Biology Laboratory, Heidelberg, Germany; Faculty of Biosciences, Heidelberg University, Heidelberg, Germany; Heidelberg University, Faculty of Medicine, and Heidelberg University Hospital, Institute for Computational Biomedicine, Bioquant, Heidelberg, Germany; Structural and Computational Biology Unit, European Molecular Biology Laboratory, Heidelberg, Germany

## Abstract

**Highlights:**

- High throughput glycoproteomics method with multiplexed quantification
- 25-fold improvement of the mouse brain glycoproteome coverage
- Structural features dictate level of glycosite micro-heterogeneity
- Gut microbiome composition extensively impacts the brain glycoproteome
- Modulation of glycosylation is site-specific

Protein glycosylation is a highly diverse post-translational modification, modulating key cellular processes such as cell signaling, adhesion and cell-cell interactions. Its deregulation has been associated with various pathologies, including cancer and neurological diseases. Methods capable of quantifying glycosylation dynamics are essential to start unraveling the biological functions of protein glycosylation. Here we present Deep Quantitative Glycoprofiling (DQGlyco), a method that combines high-throughput sample preparation, high-sensitivity detection, and precise multiplexed quantification of protein glycosylation. We used DQGlyco to profile the mouse brain glycoproteome, in which we identify 158,972 and 15,056 unique N- and O-glycopeptides localized on 3,199 and 2,365 glycoproteins, respectively - this amounts to 25-fold more glycopeptides identified compared to previous studies. We observed extensive heterogeneity of glycoforms and determined their functional and structural preferences. The presence of a defined gut microbiota resulted in extensive remodeling of the brain glycoproteome when compared to that of germ-free animals, exemplifying how the gut microbiome may affect brain protein functions.

## Introduction

Protein glycosylation is one of the most common protein modifications, consisting of a covalent linkage of sugar molecules to amino acid side chains. About one third of human proteins are going through the secretory pathway during which the vast majority of proteins are glycosylated on specific asparagine side chains (N-glycosylation) upon translocation to the endoplasmic reticulum (Helenius and Aebi, 2001; Schjoldager et al., 2020). Furthermore, glycans are extensively modified in the Golgi apparatus before reaching their final destination, which can be the extracellular space, the plasma membrane or other membrane-bound organelles (Helenius and Aebi, 2001). In contrast, O-glycosylation happens mainly on serine and threonine residues, in the Golgi or the cytoplasm (Schjoldager et al., 2020; Spiro, 2002).

Protein glycosylation is the most diverse posttranslational modification, as around 200 different glycosylation enzymes can act sequentially to create glycans of highly heterogeneous composition and structure (Schjoldager et al., 2020). In addition to glycosylation macroheterogeneity, which refers to the fact that proteins can be glycosylated at different sites, there is also glycosylation microheterogeneity, which accounts for the variety of glycans that can be attached to the same site. Although the regulatory roles of glycosylation microheterogeneity is a relatively new concept, it is now accepted that the alteration of protein glycoforms can impact protein functions by changing protein-protein or protein-ligand interactions (Varki, 2017). In general, protein glycosylation is involved in the regulation of key biological processes including cell signaling, cell adhesion or endocytosis (Helenius and Aebi, 2001; Schjoldager et al., 2020; Spiro, 2002; Varki, 2017) and deregulation of protein glycosylation has been associated with the onset of many pathologies, including cancer and neuronal disorders (Pinho and Reis, 2015; Reily et al., 2019). Particularly important and prevalent in the brain, glycosylation is known to regulate receptor trafficking, surface mobility and ligand binding affinity, and is generally involved in the maintainance of synapse physiology (Scott and Panin, 2014a). While essential to understand the role of glycosylation in health and disease and the molecular mechanisms underlying the modulation of glycoprotein functions, the comprehensive mapping and quantification of glycosylation macro- and microheterogeneity is currently difficult. The main challenge lies in the lack of highly selective enrichment strategies of the glycoproteome and high-throughput quantitative approaches to measure glycosylation.

The substoichiometric nature of glycosylation - due to the presence of multiple glycoforms on the same site - and lower ionization efficiency when compared to non-modified peptides makes glycopeptide enrichment strategies mandatory to achieve high sensitivity (Riley et al., 2021; Stavenhagen et al., 2013). Several methods have been developed to enrich glycopeptides from protein digests (Riley et al., 2021), including lectin affinity chromatography (LAC) (Sharon and Lis, 2004) and hydrophilic interaction chromatography (HILIC) (Hägglund et al., 2004). While highly useful, these methods have technical limitations, such as biases towards certain types of glycan compositions or structures and low enrichment specificities due to the inability to perform stringent washes to remove non-glycosylated peptides while preserving lectin protein structure for LAC and the co-enrichment of hydrophilic non-modified peptides for HILIC.

Chemical coupling of glycopeptides to beads functionalized with reactive groups generally leverages the reversible reaction between phenyl-boronic acid (PBA) derivatives and 1,2- or 1,3-diols present in sugar molecules, resulting in the formation of a covalent bond between functionalized beads and glycopeptides at high pH. In theory, such an approach has two key advantages: the glycopeptide-bead covalent linkage enables stringent washes to increase glycopeptides enrichment specificity, and glycopeptides can be enriched in an unbiased manner, as nearly all contain reactive diols groups. Nevertheless, chemical coupling historically identified less glycopeptides than LAC or HILIC enrichments, and often requires in-house preparation of functionalized beads. The use of amine-free binding buffers was recently shown to improve the capture of glycopeptides by beads functionalized by a PBA derivative (Morgenstern et al., 2022), opening the door for the broad use of PBA derivatives in glycoproteomics.

Here we optimized an enrichment and prefractionation strategy that substantially increases the coverage of the glycoproteome, and is coupled to a multiplexed and affordable quantification strategy. We quantified more than 50,000 unique glycopeptides per experiment in brain tissues, and identified in total 158,972 unique glycopeptides across all experiments. This amounts to a 25-fold improvement when compared to previous state-of-the-art glycoproteomics studies, and enabled us to study the structural and sequence dependent preferences of the different glycoforms. We also detected extensive changes in protein glycoform abundance in the brain of mice colonized with defined gut microbiomes compared to that of germfree mice, gaining further insights into the poorly characterized glycosylation regulation processes.

## Results

### Sensitive and specific enrichment of protein glycosylation

We developed a glycoproteomics workflow (Figure 1A) that uses commercially available and cost-effective silica beads functionalized with PBA to selectively enrich intact glycopeptides from human cells or mouse brain tissue protein digests, while eliminating contaminants. All steps of the workflow are performed in 96-well (filter)-plates to increase samples processing throughput. Besides glycopeptides, other sugar-containing biomolecules can also bind to PBA beads, such as glycolipids or RNA molecules (via the terminal nucleosides), and thus interfere with the detection of glycopeptides. While glycolipids are easily removed during protein precipitation, nucleic acids co-precipitate with proteins, as illustrated by the high abundance of an RNA marker ion (330.06 m/z) (Potel et al., 2018) in glycopeptides enriched samples after 2% SDS lysis followed by protein precipitation (Figure S1A). To address this issue, we lyzed the samples in a buffer composed of a high concentration of chaotropic salts and 40% acetonitrile, inducing nucleic acid precipitation while proteins remain in solution. Nucleic acid aggregates were then filtered out on 96-well filter plates before increasing acetonitrile concentration to precipitate proteins prior to tryptic enzymatic digestion. This fast sample preparation workflow (1h for 96 samples) enabled efficient removal of RNA molecules (Figure S1A), increasing the number of unique N-glycopeptides identification by 60% (Figure S1B, Supplementary Data 1). Next, we adjusted the mass over charge (m/z) MS1 scan range in order to preferentially select glycopeptides ions which are higher in mass compared to non-modified peptides ions that non-specifically bind to PBA beads (median molecular mass > 3,500 Da vs 1,300 Da in this study, respectively, Figure S1C). This increased the number of identified unique N-glycopeptides by 20% as well as the enrichment specificity of N-glycopeptides by 13% compared to a commonly used mass over charge scan range (Figures S1D and S1E, Supplementary Data 1). In total, using MSFragger (Polasky et al., 2020) as a search engine, we were able to identify on average 11,072 unique glycopeptides, 1,724 glycosites and 794 glycoproteins in human cell lines (HeLa and HEK293T) per single shot replicate. These numbers increased to 16,644 unique glycopeptides, 2,530 glycosites and 1,077 glycoproteins in mouse brain samples (Figure 1B).

**Figure 1:**
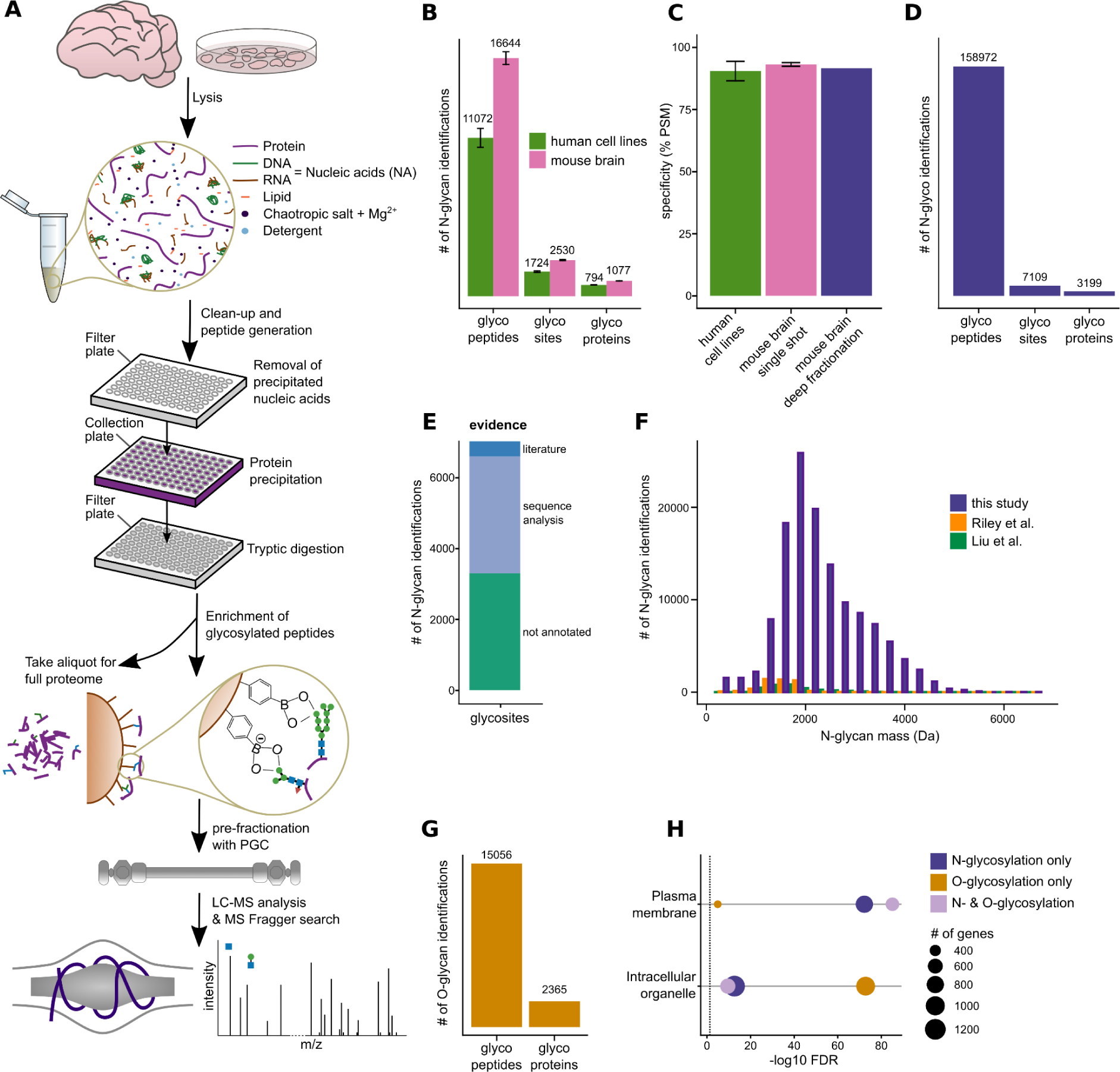
Deep profiling of protein glycosylation in human cell lines and mouse brain. **A** Deep Quantitative Glycoprofiling workflow, (DQGlyco). **B** Average numbers of unique glycopeptides, glycosites, and glycoproteins identified from LC-MS/MS replicates of unfractionated samples from human cell lines (HelaK and HEK293T cells) and mouse brain. All samples were analyzed in triplicates. **C** Glycopeptides enrichment specificity, calculated as the ratio of glycopeptide spectrum matches to all peptide spectrum matches (i.e. glycopeptides + non-modified peptides spectrum matches) for unfractionated samples from human cell lines (HeLaK and HEK293T) and mouse brain, as well as from Porous Graphitic Carbon (PGC) fractionated mouse brain samples. **D** Total number of identified N-glycopeptides (unique sequence and glycan composition), N-glycosites and N-glycoproteins from PGC fractionated mouse brain samples. **E** Uniprot annotation of glycosites identified in mouse brain samples. Literature evidence corresponds to experimental evidence while the majority of sites is annotated based on sequence analysis, i.e. prediction of glycosite based on the presence of the N-X-S/T, X≠P, glycosylation sequons. **F** Total number of unique glycopeptides of the PGC fractionated mouse brain sample per binned m/z range compared to recent large-scale glycoproteomic studies(Liu et al., 2017; Riley et al., 2019). **G** Total number of O-glycopeptides and O-glycoproteins identified in the PGC fractionated mouse brain sample. **H** Gene Ontology enrichment analysis (stringdb) for proteins found to be only N-glycosylated, only O-glycosylated or N- and O-glycosylated for two selected terms (intracellular organelle and plasma membrane).

Overall, DQGlyco enables high-throughput sample preparation and enrichment of hundreds of samples per day, as all steps are performed in 96-well plates. Moreover, DQGlyco significantly improves the detected glycoproteome coverage when compared to previous studies (Figure S2A), without the need for fractionation. Remarkably, the enrichment selectivity was greater than 90% for all samples (Figure 1C, Supplementary Data 1).

### Porous Graphitic Carbon off-line fractionation enables deep mouse brain glycoproteome analysis

As protein glycosylation is abundant and known to have important biological roles in the brain, and because brain tissue samples present a higher complexity when compared to cell lines derived samples, we sought to further improve the glycoproteome coverage in mouse brain samples. To do so, we used Porous Graphitic Carbon (PGC) as a first dimension of chromatographic separation for glycopeptides prior to a second, classically used C18-based reversed-phase separation coupled to the mass spectrometer. Due to a mixed mode retention mechanism (West et al., 2010), PGC can more efficiently resolve different glycan species than traditional reversed-phase chromatography (Jensen et al., 2012; Melmer et al., 2011) and presents a good orthogonality with C18 for the separation of non-modified peptides (Zhao et al., 2014). We thus inferred that a first dimension of separation using PGC would facilitate a better capture of glycosylation microheterogeneity and improve the glycoproteome coverage.

In total, we were able to identify 158,972 unique N-glycopeptides and 7,109 N-glycosites located on 3,199 N-glycoproteins (Figure 1D, Supplementary Data 2) after PGC fractionation. In terms of unique N-glycopeptides, this represents a greater than 25-fold improvement compared to current state-of-the-art N-glycoproteomics studies using LAC (Riley et al., 2019) or HILIC based enrichment strategies (Liu et al., 2017) on mouse brain samples (Figure S2A). Our dataset includes 60% of proteins which have been previously identified in diverse regions of the mouse brain (Giansanti et al., 2022) and annotated as glycosylated in Uniprot (Figure S2B). Overall, around half of the glycosites and glycoproteins identified in this study were not reported as glycosylated in Uniprot, while only 5% were previously experimentally validated (Figures 1E and S2C). Thus, this study substantially expands the known mouse brain N-glycoproteome. The proteins reported as glycosylated in Uniprot are predominantly annotated as part of the plasma membrane while previously unreported glycosylated proteins are mainly annotated as part of subcellular organelles (Figure S2D). The fact that close to 300 proteins found to be N-glycosylated in our dataset were not identified in the mouse proteome atlas (Giansanti et al., 2022) study (Figure S2E) highlights the sensitivity of our method as these proteins are likely expressed at low level. Typically, glycopeptide enrichment methods have biases towards certain glycan compositions. Therefore, we compared the different glycan compositions or masses identified by alternative enrichment strategies and found that N-glycans identified by our PBA-based approach encompass N-glycans previously identified by different approaches (Figure 1F), while greatly improving the glycoproteome coverage, suggesting a low enrichment bias towards specific glycopeptides.

It is estimated that N-glycopeptides are around 10-fold more abundant than O-glycopeptides in mammalian cells, hence O-glycopeptides identification usually necessitates dedicated analytical strategies (Riley et al., 2021). For this reason, most O-glycoproteomics studies usually focus on specific protein classes (e.g. mucins (Malaker et al., 2022)), have a bias towards specific types of glycans (e.g. small glycans (Woo et al., 2018; Yang et al., 2018)) or do not allow the characterization of intact O-glycopeptides. Here, we reasoned that the deep glycoproteome coverage obtained by DQGlyco could enable the simultaneous identification of both N- and O-glycopeptides. In total, we identified 15,056 O-glycopeptides on 2,365 O-glycoproteins in mouse brain samples (Figure 1G, Supplementary Data 2), constituting to our knowledge the most extensive characterization of intact O-glycosylation in any mammalian tissue. It has to be noted that as higher-energy collisional dissociation (HCD) fragmentation was used in this study, we could not confidently assign the site of glycosylation of a given glycopeptide (Riley et al., 2020). Still we could identify over 400 unique O-glycan compositions, with a molecular weight distribution shifted to lower masses when compared to N-glycans (Figure S2F). We found about 1,400 proteins to be only O- and not N-glycosylated. Proteins that were solely O-glycosylated were significantly enriched in intracellular membrane-bound organelles, corroborating that N- and O-glycosylations serve different functions (Figure 1H). In accordance with previous reports, O-GlcNac modification (addition of a single sugar molecule) was enriched in intracellular organelles, whereas no difference in glycan elongation was observed between proteins belonging to the plasma membrane and to intracellular organelles (Figure S2F).

### Deep glycoproteomics profiling reveals extensive microheterogeneity

The high number of glycoforms identified in this study represents a rich resource for studying glycoform microheterogeneity. Although half of the sites were singly or doubly glycosylated, many sites were modified by multiple glycoforms (mean = 18, Figure 2A) and some sites exhibited extreme microheterogeneity, as for example the N205 site on the excitatory amino acid transporter 2 protein (EAAT2, also referred to as SLC1A2), on which 759 glycoforms were identified. Notably, we did not observe any correlation between the detected number of glycosites per protein or number of glycoforms per site and the abundance of the respective protein (Figures S3A,B). This suggests that the glycosylation heterogeneity observed is not biased towards easily detected/abundant proteins.

**Figure 2:**
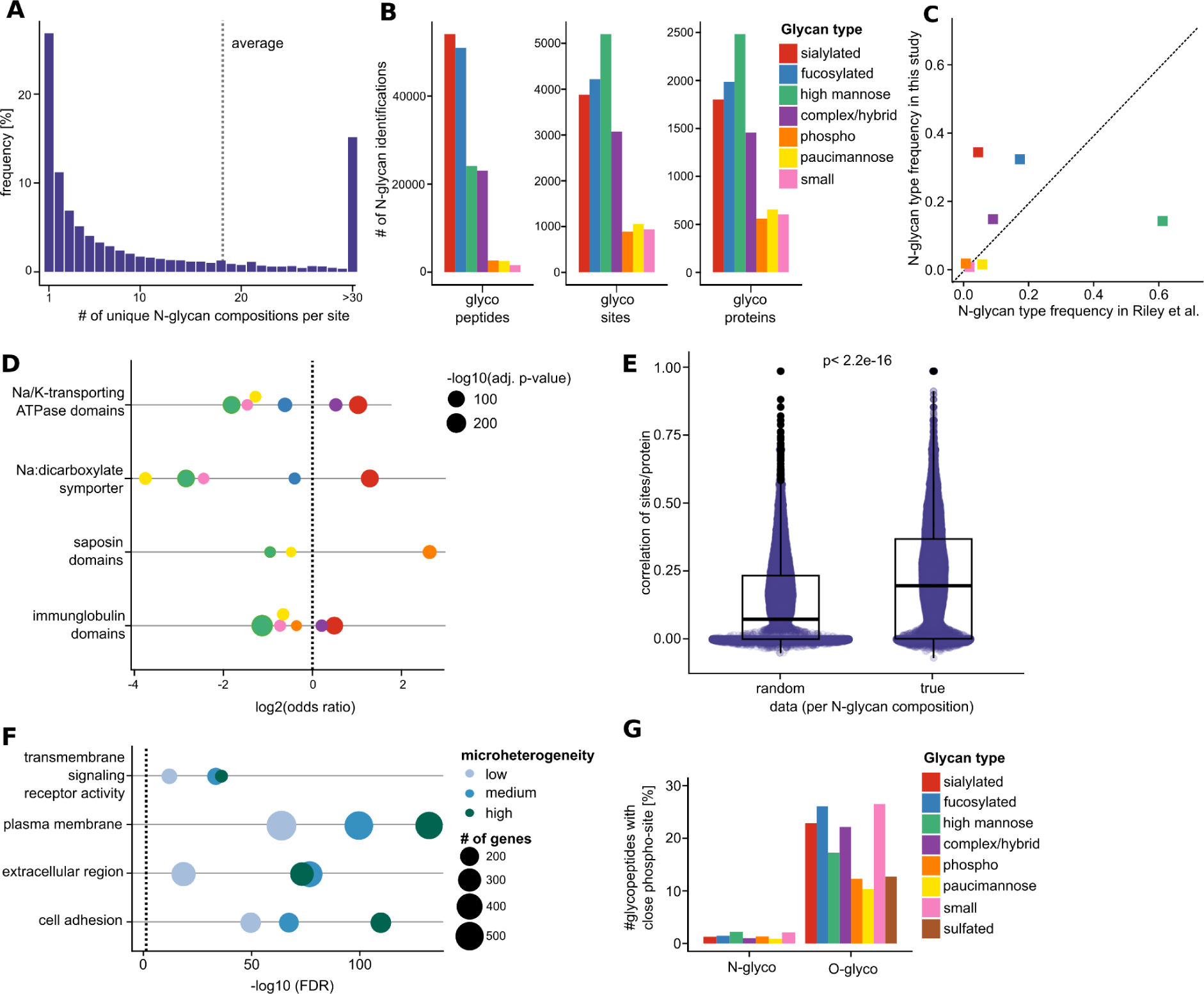
Protein glycosylation composition and microheterogeneity in the mouse brain. **A** Frequency of the number of unique glycan compositions per site in the PGC fractionated mouse brain N-glycosylation data (on average, 18 glycan compositions were identified per site). **B** Total number of unique glycopeptides, glycosites and glycoproteins per glycan class for the PGC fractionated mouse brain sample. **C** Frequencies of glycan classes identified in this study and in a recent glycoproteomic study using lectin-based enrichment(Riley et al., 2019). **D** Distinct glycan classes are significantly over- or underrepresented on specific functional protein domains from the InterPro database (log2 odds-ratio >/< 0, Fisher’s exact test, adjusted p-value < 1e-3, examples shown). **E** Glycosylation profiles of sites belonging to the same protein are significantly more correlated compared to when glycan compositions are shuffled across the whole glycoproteome (Kendall rank correlation coefficient, Wilcoxon signed rank test, p-value < 2.2e-16). **F** Results of the Gene Ontology enrichment analysis (stringdb) of proteins with sites displaying low, medium or high microheterogeneity (N = 1-2, 3-11, >11 unique glycan compositions per site, respectively) for selected significant terms (false-discovery rate < 0.05). **G** Frequency of glycosites or glycopeptides with a close known phosphosite (+/− 5 residues) per glycan class for O- and N-glycosylation data.

To facilitate data visualization, the different glycan compositions were classified into 8 classes based on their chemical nature. It is known that different classes of glycans can have different impacts on protein-protein or protein-ligand interactions through steric or electrostatic effects and serve as epitopes for specific lectin receptors (Helenius and Aebi, 2001; Schjoldager et al., 2020; Spiro, 2002; Varki, 2017). N-glycans are added in the ER as high-mannose and can be processed in the Golgi into glycans classified as high-mannose, small, paucimannose, phosphorylated, fucosylated, sialylated, sulfated (here only for O-glycosylation), while the remaining compositions were considered as complex/hybrid (Figure 2B, Figure S3C) (see methods section for details). Compared to the Riley et al. dataset (Riley et al., 2019), we identified significantly less high mannose N-glycosylation, which are often regarded as immature glycans as they can constitute intermediate species (less than 15% of all glycopeptides identified vs 60%), confirming that the commonly used conA lectin-based enrichment is biased towards high mannose glycosylation and indicating that deep coverage is needed to achieve comprehensive characterization of mature glycoforms (Figure 2C). In this study, close to half of the monoglycosylated sites were found to be modified by high-mannose, implying that most of the detected monoglycosylated sites are indeed subjected to a low number of processing events (Figure S3D).

The distribution of different classes of N-glycosylation differs between the two human cell lines and the mouse brain (Figure S3E), likely reflecting differences in the regulation of glycosylation processes. Little difference in enriched Gene Ontology terms was observed between the different glycan classes (Figure S3F), meaning that different types of glycosylation in general do not target proteins with specific functions. However, we observed significant differences between the enriched protein domains for the different N-glycosylation classes (Figure 2D). Sodium/Potassium transporting ATPase domains were for example strongly enriched in sialylated and paucimannose glycans, while saposin domains - specific to lysosomal proteins - were enriched for phosphoglycans. Phosphoglycans were in general strongly enriched on lysosomal proteins (Figure S3G), in line with the fact that mannose-6-phosphate serves as a tag to direct enzymes to the lysosome(Coutinho et al., 2012).

As the different sites of a given N-glycoprotein should be exposed to the same glycoenzymes, we investigated if microheterogeneity across different sites within the same protein was more similar than across the glycoproteome. We observed no clear pattern of co-occurrence of different classes of N-glycosylation either at the glycosite or at the glycoprotein level (Figures S4A,B). However, when correlating the exact N-glycoform compositions between sites localized on the same protein before and after random reshuffling of all glycoforms across all identified sites, we concluded that while there is a lot of variation in microheterogeneity between different sites on the same protein, this variation is significantly lower than across the glycoproteome as a whole (Figure 2E). Interestingly, significant differences were observed when classifying N-glycoproteins according to the level of site microheterogeneity, independently of glycan compositions: sites with high heterogeneity were enriched on extracellular proteins, involved in processes such as cell surface receptor signaling pathways and cell adhesion, both known to be modulated by glycosylation (Schjoldager et al., 2020) (Figure 2F).

No specific protein annotation was associated with a class of O-glycosylation, with the intracellular or plasma membrane O-glycoproteins having similar glycosylation patterns (Figure S4C). Proteins exhibiting only one O-glycoform were enriched in intracellular organelle proteins while proteins with higher number of identified O-glycoforms were enriched in membrane proteins involved in signaling (Figure S4D). It has been reported that O-GlcNAcylation (addition of a single sugar) and phosphorylation have a propensity to co-occur on the same protein regions, resulting in a PTM crosstalk (Hart et al., 2011). Here we show that around 25% of O-glycosylated peptides were also previously reported to be phosphorylated, independently of the glycan mass and composition, hinting at the possibility that the existence of a crosstalk can be extended to all glycoforms on O-glycosites. In comparison, less than 2% of N-glycosites were in close proximity to reported phosphorylation sites (Figure 2G).

### Glycosylation in a structural context

Next we investigated structural features influencing (i) which sites are glycosylated and (ii) the composition and number of different glycoforms at a given site. To do so, we compared the glycosylated asparagines to non-glycosylated but glycosylatable asparagines (i.e. part of glycosylation sequons N-X-S/T, X≠P, noted ngN) belonging to identified glycoproteins, and separated the different glycans according to the aforementioned classification or the number of glycoforms identified on a given site. We observed that when compared to ngN, glycosites are enriched in regions annotated as extracellular and lumenal while depleted in cytoplasmic parts of the proteins, suggesting that many potential glycosites for which we do not detect glycoforms are indeed not glycosylated in the cell (Figure 3A). Likewise, O-glycosylation is more prevalent in extracellular and lumenal regions (Figure S5A).

**Figure 3:**
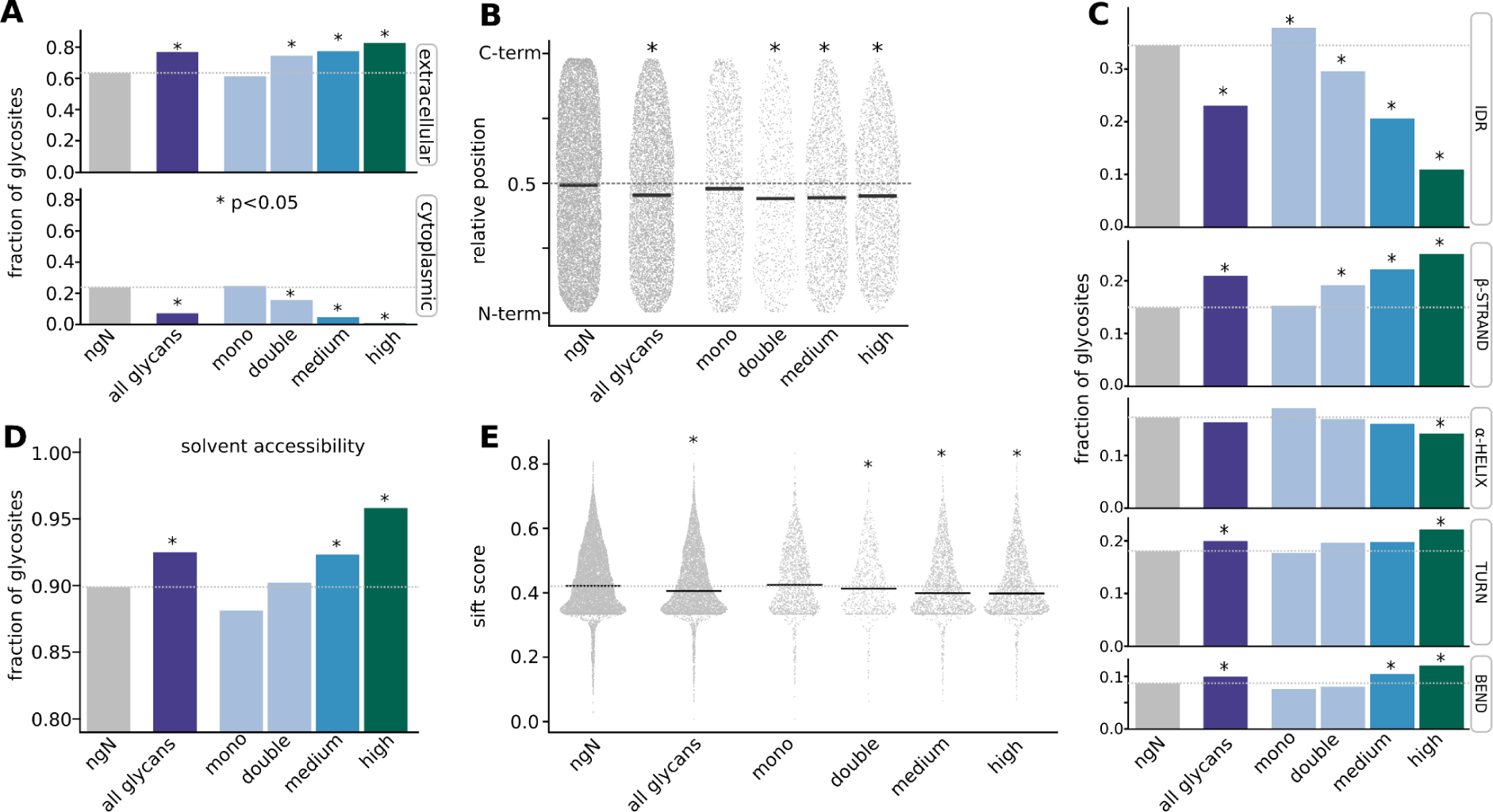
Glycosylation in a structural context. **A** Glycosites exhibiting high microheterogeneity are significantly enriched in extracellular and depleted in cytoplasmic regions of glycoproteins (N = 1, 2, 3-11 and >11 unique glycan compositions per site referred to as monoglycosylated, doubly glycosylated, medium and high microheterogeneity, respectively). Barplots indicate the fraction of glycosites inside the region of interest. Asterisks indicate Fisher’s exact test p-value < 0.05 using non-glycosylated but glycosylatable asparagines as background (noted ngN). **B** Glycosites are localized closer to the protein N-terminus when compared to ngN. The dotplot represents, for each site, its relative position within the protein. The solid lines indicate each group’s median relative position, and the asterisk indicates a Wilcoxon signed rank test p-value < 0.05 for each group compared to ngN. **C** Frequency of glycosites on defined structural motifs varies depending on the degree of site microheterogeneity. **D** Degree of glycosite microheterogeneity is associated with the site’s solvent accessibility. **E** Glycosites presenting higher microheterogeneity are more conserved when compared to monoglycosylated sites. Average SIFT scores across all potential amino-acid substitutions were used as a proxy for site conservation.

The N-glycosylated sites are also shifted significantly closer to the N-terminus compared to ngN, (Figure 3B), confirming a previous observation made on a few proteins that N-glycosylation efficiency is lower closer to the C-terminus of the protein (Bañó-Polo et al., 2011). We used AlphaFold (Jumper et al., 2021) and its metric (pPSE: prediction-aware part-sphere exposure) to assess structural disorderness of proteins (Bludau et al., 2022) to study glycosylation in a structural context. We found that glycosites were significantly depleted in disordered regions of proteins when compared to ngN, and in accordance with previous findings (Zielinska et al., 2010), N-glycosylation is enriched on beta strands, turns and bends (Figure 3C).

Only a few significant differences with small effect size were observed between the different glycosylation classes in the above analysis (Figures S5B-E). However, significant differences with larger effect size were observed when classifying glycosites based on their level of microheterogeneity. When compared to monoglycosylated sites, multiglycosylated sites are located more often on extracellular domains and are closer to the N-terminus of the protein (Figure 3A,B). Additionally, we observed a strong trend for sites with a higher number of glycoforms to be localized on more structurally ordered regions (Figure 3C), in line with the fact that extracellular regions are on average more structurally ordered than cytoplasmic regions (Figure S5F). When compared to sites presenting low microheterogeneity, highly modified sites are enriched on beta strands and depleted on alpha helices, as well as enriched at points of change in protein secondary structure (turns and bends, Figure 3C). This suggests that modulation of glycosite microheterogeneity could regulate protein conformation.

Interestingly, sites with low microheterogeneity, indicating few glycan processing events, were found to be localized on less ordered regions and exhibit less solvent accessibility when compared to multiglycosylated sites (Figure 3D). This suggests that specific ordered structures combined with high site solvent accessibility are needed for extensive glycan processing by glycosyltransferases. Finally, the sites with high numbers of glycoforms were also more conserved (lower SIFT score (Ng and Henikoff, 2003)) than monoglycosylated sites (Figure 3E). Taken together our data uncovers significant structural differences between glycosites with low or high microheterogeneity.

### Tandem mass tag based quantification of glycoforms

In order to understand mechanisms of glycosylation regulation and the functional consequences of glycosylation microheterogeneity robust and high throughput quantification methods are required. Up to now, most of the few existing quantitative glycoproteomics studies have relied on label-free approaches, limiting quantitative analyses. Indeed, label-free quantification is not compatible with the different pre-fractionation strategies that improve analytical depth.

Alternatively, the use of Tandem Mass Tags (TMT) labeling, which enables sample multiplexing and is compatible with prefractionation, has been described for glycoproteomics (Fang et al., 2020; Saraswat et al., 2022; Stadlmann et al., 2017). However, the high cost of TMT reagents prevents its routine use, as pre-enrichment labeling - where both the glycopeptides and non-modified peptides are labeled together - is recommended to ensure precise quantification. Here we demonstrate that it is possible to decrease the amount of TMT reagents used 200-fold when performing post-enrichment labeling, as only the substoichiometric glycopeptides need to be labeled. This affordable quantification method enables multiplexing of up to 18 conditions with high quantitative reproducibility as shown in Figures 4A,B and Figure S6A,B, Supplementary Data 3 (average correlation > 0,984 between biological replicates of different mouse brain samples at the glycopeptide level). This illustrates the high reproducibility of our sample preparation workflow and enrichment platform. In addition, this shows that the brain glycoproteome is tightly regulated and conserved between biological replicates, (Figures 4A,B and Figure S6A,B), in contrast to other tissues, such as plasma, for which high interindividual differences were reported (Knežević et al., 2009). Next we decided to investigate how glycosylation patterns change in vivo upon perturbations.

**Figure 4:**
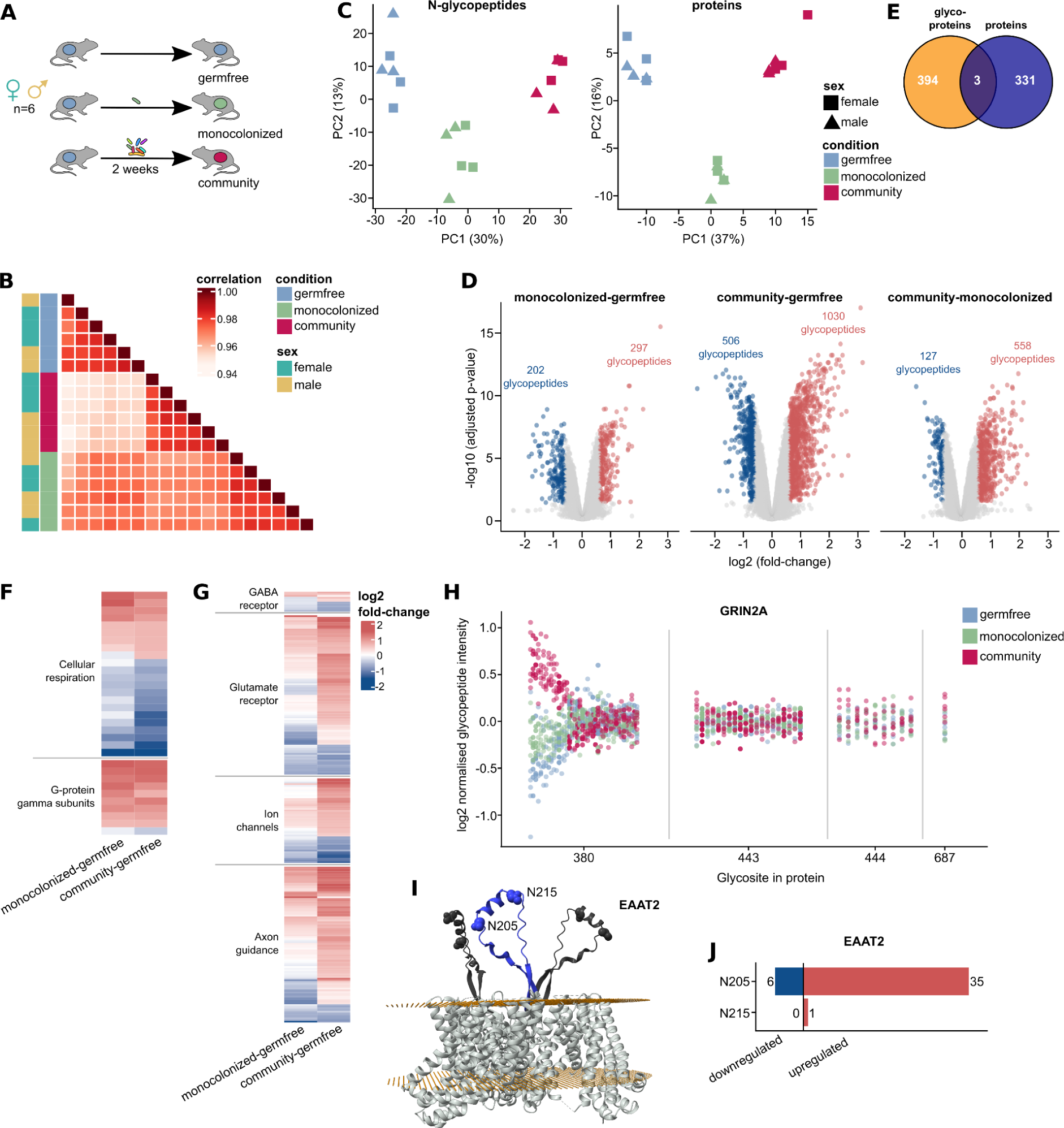
Impact of the gut microbiome on the mouse brain glycoproteome. **A** Experimental design: 3 groups of 6 adult germ-free mice (3 males + 3 females) were either monocolonized by *B. uniformis*, colonized by a defined 8-members community of human gut commensal or remained germ-free for 2 weeks. **B** Spearman correlation of glycopeptides intensities between biological replicates. **C** Principal Component Analysis (PCA) of the normalised reporter intensities of the 18 brain samples demonstrates gut microbiome dependent remodeling of the brain proteome and N-glycoproteome. **D** N-glycopeptide regulation in the mouse brain depending on the gut microbiome composition. **E** Proteins mostly exhibited significant regulation at either the glycoproteome or the proteome level specifically, with only three proteins being significantly regulated at both levels. **F** Mitochondrial proteins involved in cellular respiration as well as G-protein gamma subunits were shown to be regulated at the protein abundance level. **G** Amongst the proteins exhibiting significant regulation at the N-glycosylation level are many proteins for which glycosylation has been reported to impact protein function: neurotransmitter receptors (GABA and glutamate receptors), as well as ion channels and proteins involved in axon guidance. **H** Example of site-specific regulation of N-glycosylation on the GRIN2A glutamate receptor. Only one site exhibits significant regulation of defined glycan compositions (N380) while other sites belonging to the same protein remain unchanged. Each vertical line represents one glycopeptide while each dot represents a replicate (3 conditions x 6 biological replicates). **I** EAAT2 is a membrane-bound transporter which is heavily glycosylated at the N205 and N215 sites located on an extracellular loop (blue) protruding from the scaffolding domain of the protein. Here, we show the human structure combined with AlphaFold predictions for the extracellular region, as murine structure is not available (sequence homology > 95%). The light gray structure represents the template homotrimer (PDB accession: 7VR8), serving as a reference for protein architecture. Yellow planes depict the membrane orientation prediction. Dark-blue and dark-gray regions correspond to predicted structures aligned to the template structure. Spheres indicate the glycosites of interest (N205 and N215). **J** The glycoforms on the two glycosylation sites of the EAAT2 protein are differentially modulated, with only one glycoform being upregulated on N215, while extensive glycoform regulation is observed on the N205 site upon gut microbiota colonization.

### Impact of gut microbiome on the mouse brain glycoproteome

Alteration of the gut microbiome composition has been associated with behavioral changes and has been shown to impact brain development and function via nervous or chemical signaling along the gut-brain axis (Morais et al., 2021). However, underlying molecular mechanisms remain poorly characterized. Given the known importance of glycosylation for neuronal functions, we decided to assess the impact of different gut microbiome compositions on both the brain proteome and glycoproteome. Three groups of 6 adult germ-free C57BL/6 mice (3 males and 3 females) were either monocolonized for 2 weeks with *Bacteroides uniformis*, one of the most prevalent human gut microbes, by a defined 8-member community composed of human commensal gut bacteria (*B. uniformis*, *C. sporogenes*, *P. distasonis*, *L. gasseri*, *E. rectale*, *A. muciniphila*, *R. gnavus* and *P. copri*) or were kept germ-free (Figure 4A). Changes at both protein abundance and glycoform levels were quantified, with high reproducibility between biological replicates (Figure 4B,C, Figure S7A). In total, 334 out of 9,744 quantified proteins - in the unenriched sample - and 1,700 out of 56,171 quantified glycopeptides on 394 glycoproteins - in the enriched sample - were identified to significantly change at the proteome and glycoproteome levels, respectively (Figure 4D, Figure S7B), with changes at the glycoform level predominantly occurring on sites exhibiting high microheterogeneity (Figure S7C). Notably, only three proteins (ATP6V0A2, CNTNAP5A and ABHD14A) exhibited significant changes at both abundance and glycosylation levels (Figure 4E, Supplementary Data 3). A clear separation was observed between the 3 mice groups after principal component analysis, indicating gut microbiome-specific remodeling of the brain proteome and glycoproteome at the adult stage (Figure 4C).

Proteins significantly changing in abundance were enriched in mitochondrial proteins, in line with previous reports on gut microbiome-mitochondria crosstalk (Saint-Georges-Chaumet and Edeas, 2016), likely via the secretion of small molecules (Figure 4F). This could constitute an interesting perspective to investigate further in the context of the gut microbiome-brain axis, as mitochondrial function deregulation has been associated with neurological pathologies (McFarland et al., 2010). In addition, we observed abundance changes for 9 of the 12 G-proteins gamma subunits, with the 8 moderate/slow translocating G-proteins gamma identified being upregulated and the fast translocating Gng11 being downregulated (Figure 4F). G-proteins associate with transmembrane receptors to modulate and relay signals to the inside of the cell, and G-proteins gamma translocation kinetics has been associated with cell sensitivity and adaptation to extracellular signals (Kankanamge et al., 2022), hinting at a change in receptor activity.

N-glycosylation has been shown to play roles in the modulation of neurotransmission via regulation of neurotransmitter receptors ligand affinity binding, oligomerization states, trafficking or cell surface retention (Scott and Panin, 2014b). Amongst proteins exhibiting changes at the glycosylation levels, 13 glutamate and 6 GABA G-protein coupled receptor proteins were identified (Figure 4G). Glutamate and GABA are the main excitatory and inhibitory neurotransmitters, respectively. We additionally identified changes of glycoform abundance on several other ion channel proteins (Figure 4G). Here again, N-glycosylation has been shown to regulate neurotransmission by modulating membrane excitability via the regulation of ion channel proteins (Scott and Panin, 2014b). Interestingly, such proteins were found to exhibit glycosylation changes upon *in vitro* stimulation by synaptosome depolarization (Boll et al., 2020), suggesting that observed alterations of glycosylation are linked to changes in neuronal activity. Finally, proteins involved in axon guidance showed significant changes of glycoforms levels (Figure 4G). Notably, the absence of maternal microbiome has previously been linked to defects in fetal neurodevelopment, with reduced axonogenesis and downregulation of genes involved in axon guidance (Vuong et al., 2020). As glycosylation has been shown to be critical for axon guidance (Mutalik and Gupton, 2021) due to its importance in the regulation of cell adhesion and migration processes, our data provides another layer of evidence that the gut microbiome could impact axogenesis.

It is generally accepted that changes in protein glycosylation occur mainly via modulation of glycoenzymes abundance, localization or activity. In such cases, similar trends would be expected at the glycoproteome level and for glycosites within the same protein (e.g. general increase/decrease of sialylation via modulation of sialyltransferases activity or abundance). However, only one glycosyltransferase enzyme changed significantly in abundance in this study (ALG12, glycosyltransferase localized in the ER, which cannot be responsible for the vast changes in microheterogeneity observed). Also, no trend in specific regulation of one class of glycoform was observed (i.e. all glycan classes followed the same trend, Figure S7D) and glycopeptides belonging to the same glycan class localized on the same site were not co-regulated (Figure S7E,F). On the other hand, regulated glycoforms belonging to the same site, regardless of the glycan composition, tend to change in abundance in the same direction (Figure S7G,H), indicating a regulation of the overall level of glycosylation of specific glycosites. Furthermore, the deep glycoproteome coverage combined with precise quantification revealed extensive site-specific regulation of glycoforms - i.e. only specific glycoforms on specific glycosites within a protein are regulated while others remain unchanged (Supplementary Data 5) - as shown for the glutamate receptor ionotropic NMDA 2A (GRIN2A, Figure 4H). These observations suggest complex posttranslational mechanisms of regulation in order to control site specific glycosylation dynamics.

A striking case in point are the two glycosites identified on the excitatory amino acid transporter 2 (EAAT2, Figure 4I), on which a similar number of glycoforms were quantified (458 on N205, 460 on N215). While only 10 amino acids apart, the N205 site exhibits extensive upregulation of glycoforms in the brain of colonized mice, while only one glycoform was found to be significantly changing on the N215 site (Figure 4J). This family of transporters plays an essential role in the brain by maintaining low levels of extracellular excitatory neurotransmitter glutamate in the synaptic cleft to avoid excitotoxicity, and their deregulation has been associated with various neurological diseases (Malik and Willnow, 2019). Interestingly, a significant general decrease of glycosylation level on EAAT2 has been observed in brain samples of schizophrenic patients (Bauer et al., 2010), illustrating how the gut microbiome could impact brain physiology via the modulation of protein glycosylation.

## Discussion

Understanding the biological significance and regulation of the high microheterogeneity of protein glycosylation presents a formidable challenge. We believe our method, DQGlyco, which enables deep identification and precise quantification of glycoforms, paves the way for a better comprehension of protein glycosylation dynamics and its regulatory roles. It should be noted that in this study we distinguished glycoforms based on glycan composition. Taking into account the glycan structures will add another layer of complexity in the future.

Deep profiling of protein glycosylation in the mouse brain characterized site microheterogeneity to an unprecedented extent. While we observed that different classes of glycans have preferential localization on specific protein domains, there is an even stronger pattern of sites exhibiting different degrees of glycoform microheterogeneity to localize on different protein regions or on proteins of specific functions. This is particularly pronounced on cell surface receptors and in agreement with a previous study showing that a high extent of N-glycosylation is a feature that can be used to regulate cell surface exposition of receptors in response to metabolic changes (Lau et al., 2007). Taken together this confirms that high microheterogeneity is in general more likely to be associated with the regulation of receptor functions.

AlphaFold predictions (Jumper et al., 2021) enabled us to define site-specific structural features such as the fact that sites with higher degree of microheterogeneity are more likely to be located on highly ordered structures and to present high solvent accessibility. The latter is in line with previous work showing that solvent accessibility is an important parameter for further processing of high-mannose glycans to create mature glycan forms (Lee et al., 2014; Suga et al., 2018). We foresee that our extensive dataset will be a useful resource to further understand and predict the impact of protein structures on glycosylation and vice versa.

The affordable workflow for multiplexed quantitative analysis of protein glycoforms presented here makes it possible to systematically quantify protein glycosylation levels across many samples. This could prove especially useful to identify disease specific glycosylation patterns, which could lead to the development of targeted therapies (Mereiter et al., 2019) and more generally improve our understanding of the association between deregulation of glycosylation and disease states. While previous systematic studies quantified glycosylation changes in cell lines after perturbations directly targeting glycosyltransferases (for example inhibition (Fang et al., 2020) or gene knockout (Stadlmann et al., 2017)), here we observed significant remodeling of the brain glycoproteome *in vivo* upon changes of the gut microbiome composition, suggesting high plasticity of glycosylation in response to a broad range of stimuli. While the exact impact of the observed glycosylation changes on protein functions remain to be elucidated, the fact that many of these changes occur on neurotransmitter receptors or proteins involved in axon guidance, for which glycosylation has been previously shown to be essential for protein activity (Mutalik and Gupton, 2021; Scott and Panin, 2014b), sheds light on a new link between the gut microbiome and brain physiology.

Previous work has coined the term metaheterogeneity to reflect the fact that different sites on the same protein can be very different from each other in glycan content (Čaval et al., 2021). In our data we can see how perturbations can give rise to quantitative changes in site specific glycoform compositions, exemplifying how metaheterogeneity can be modulated in response to perturbations. Understanding the mechanisms behind these site specific modulations opens avenues for exciting future studies.

In summary, the presented workflow constitutes a framework to address many unanswered questions, such as the kinetics of glycosylation changes in response to perturbations, or the impact of glycosylation microheterogeneity modulation on protein function. The latter point represents a particularly difficult challenge as site mutagenesis, commonly used to assess functionality of other modifications (e.g. phosphorylation (Viéitez et al., 2022)), cannot address this issue, and no alternative strategy to alter glycosylation on a specific site has so far been described. A promising avenue for deciphering glycosylation functionality will be linking glycosylation changes to cellular phenotypes by subjecting cells to multiple perturbations, a strategy that has been successfully applied to protein phosphorylation (Leutert et al., 2022). Future efforts can also focus on measuring the effect of glycosylation changes on the biophysical properties of proteins, such as thermal stability (Potel et al., 2021; Savitski et al., 2014) which can inform on changes in ligand binding properties (Savitski et al., 2014) as well as protein interactions (Tan et al., 2018).

## Methods

### Cell culture

All cell lines used in this study were verified to be negative for mycoplasma contamination. HeLa Kyoto cells (a kind gift from the Ellenberg group, EMBL) and HEK293T cells were cultured in DMEM (Sigma-Aldrich, D5648) containing 4.5 mg ml−1 glucose, 10% (vol/vol) FBS (Gibco, 10270) and 1 mM L-glutamine (Gibco, 25030081) at 37 °C with 5% CO2. HeLaK and HEK293T cells (0.5 million) were seeded in 150-mm dishes and grown for 2 d. The cells were washed with ice-cold PBS (2.67 mM KCl, 1.5 mM KH2PO4, 137 mM NaCl and 8.1 mM NaH2PO4, pH 7.4) and collected by scraping. The cells were pelleted by centrifugation at 300g for 3 min. The cell pellets were flash frozen in liquid N2 and stored at −80 °C.

### Gnotobiotic animal experiments

All mouse experiments were performed using approved protocols by the EMBL ethics committee (licence 21-002_HD_MZ). Germfree (GF) C57BL/6 mice were maintained and bred in gnotobiotic isolators (CbC) with a 12-hour light/dark cycle. GF status was monitored by PCR (using 16S primer pair, Supplementary Data 3) and culture-based methods. Mice were provided with standard, autoclaved chow (1318 P FORTI, Altromin) ad libitum. Animals of mixed gender at an age between 8 and 17 weeks were used for all experiments. Colonization was performed by oral gavage of 200 μL of cryo-preserved bacteria from overnight cultures in modified Gifu Anaerobic Medium (mGAM, Nissui Pharmaceutical Co. Ltd.) of *B. uniformis* DSM6597 or with a mixed *in vitro* culture containing 8 bacterial species mixed in equal ratios, based on OD600 measurements. All inocula were made in advance and stored at −80°C until use. 14 days after colonization, the animals were euthanized and brains and cecal samples were collected.

### Determination of gut bacterial community composition

The relative cecal abundance of each species was determined from purified DNA by qPCR, (Supplementary Data 3). Hereto, DNA was extracted using a commercially available gDNA extraction kit (96 DNA kit, ZymoBiomics) following the manufacturer’s instructions. Species-specific primers were designed against intergenic regions of the published genome of each species (Supplementary Data 3). All primers were tested *in silico* and by qPCR for their reactivity to non-specific sites within the genomes of the other strains in the community. For each qPCR assay, a standard curve was calculated from genomic DNA (gDNA) isolated from each species. To this aim, gDNA was diluted to 0.5 ng/μL (as quantified by Quant-iT dsDNA BR assay kit and a Qubit fluorometer (Invitrogen). Ten-point standard curves were prepared from two-fold serial dilutions of this 0.5 ng/μL starting point. qPCR reactions were run in 20 μL volumes, containing 10 μL SYBR™ Green PCR Master Mix (ThermoFisher), 6.8 μL nuclease-free water, 1.2 μL of primers at 333nM final concentration each, and 2.0 μL template DNA. Amplification was performed using StepOnePlus™ Real-Time PCR System (Applied Biosystems™) machine running StepOne™ Software (version 2.3), using the manufacturer’s recommended protocol: 95°C for 10 minutes, followed by 40 cycles of 15 seconds 95°C and 1 minute at 60°C. Melting curve analysis followed after amplification, to ensure a single product in each reaction. qPCR estimate of the amount of DNA from each strain was estimated by normalization of the CT values by the estimated molecular weight of each strain’s genome, thereby converting the CT values to genome equivalents. Relative abundance was calculated as the proportion of a single strain genome equivalent to the total genome equivalents in the sample.

### Sample preparation

Samples were lysed in a buffer constituted of 4 M guanidinium isothiocyanate, 50 mM 2-[4-(2-hydroxyethyl)piperazin-1-yl]ethanesulfonic acid (HEPES), 10 mM tris(2-carboxyethyl)phosphine (TCEP), 1% N-lauroylsarcosine, 5% isoamyl alcohol and 40% acetonitrile adjusted to pH 8.5 with 10 M sodium hydroxide. The volume of lysis buffer used corresponded to approximately 5 times the sample volume. Human cell pellets were pipetted up and down while whole mouse brains were homogenized using a glass douncer. For the SDS lysis, samples were lysed in a buffer composed of 2% Sodium Dodecyl Sulfate, 50 mM HEPES and 10 mM TCEP. After 10 minutes of incubation at room temperature in a shaker at 1,000 rpm, samples were centrifuged at 14,000 rpm at room temperature for 5 minutes or filtered in multiscreenHTS-HV 0.45 µm 96 well filter plates with PVDF membranes (Merck Millipore) to remove cell debris and nucleic acid aggregates. Protein concentrations were determined using tryptophan fluorescence assay as described in Wisńiewski et al. (Wiśniewski and Gaugaz, 2015) and the samples were diluted to a maximum concentration of 25 µg/µL, if needed. Volumes of 80 µL of lysate per well were transferred into a multiscreenHTS-HV 0.45 µm 96 well filter plate, and 220 uL of ice-cold acetonitrile were added to induce protein precipitation. After 10 minutes, the plate was centrifuged to remove solution and protein precipitates were washed two times with 200 µL 80% acetonitrile and two times with 200 µL 70% ethanol (1,000 x g for 2 minutes for each step). Then, the digestion buffer composed of 100 mM HEPES pH 8.5, 5 mM TCEP, 20 mM chloroacetamide and trypsin (TPCK treated trypsin, Thermo Fisher Scientific) was added to the protein precipitates in the filter plate. The ratio of trypsin:protein was fixed to 1:25 w/w and the maximum final protein concentration to 10 µg/µL. Trypsin was reconstituted at a stock concentration of 10 µg/µL and added to the ice-cold digestion buffer just prior to the addition to the precipitates. Tryptic digestion was carried on overnight at room temperature under mild shaking (600 rpm). After digestion, samples were acidified to 1% TFA and desalted using Sep-pak tC18 columns (Waters), eluted using 0.1% TFA in 40% acetonitrile and dried using a vacuum concentrator.

### Glycopeptides enrichment

Dried peptides were resuspended in the glycopeptide enrichment buffer constituted of 50 mM carbonate buffer, pH 10.5 in 50% acetonitrile. The optimal sample concentration was determined to be 5 mg/mL. Silica beads functionalized with phenyl-boronic acid (Bondesil-PBA 40µm, Agilent) were used for glycopeptides enrichment at an optimized ratio of 1:2.5 sample:PBA beads w/w. Prior to addition to the samples, beads were washed 3 times with the glycopeptide enrichment buffer. The samples/beads mixtures were incubated in eppendorf tubes or 96 well plates at room temperature for 1h under sufficient shaking to prevent beads sedimentation. After incubation, the samples/beads mixtures were transferred to a multiscreenHTS-HV 0.45 µm 96 well filter plate, non-bound peptides were filtered off and beads were washed 7 times with the glycopeptide enrichment buffer, each time at a speed of 100 x g for one minute. Glycopeptides were eluted two times with 50 µL 1%TFA in 50% acetonitrile for 15 minutes at room temperature under shaking at 800 rpm before being dried in the 96 well elution plate or glass inserts using a vacuum concentrator.

### Tandem Mass Tag (TMT) labeling

Because of the presence of residual acidic salts, glycopeptides were resuspended for 5 minutes in 10 µL of 400 mM HEPES pH 10 prior to the addition of the TMTpro reagents (Thermo Fisher Scientific). To quantify the full proteome, 15 µg aliquots were taken after sample desalting and before glycopeptides enrichment, dried and resuspended in 10 µL of 100 mM HEPES pH 8.5 prior to TMT labeling. For all samples, 4 µL of TMTpro reagents at a concentration of 20 µg/µL in acetonitrile were added in each well. The labeling reaction was carried on for one hour at room temperature under mild shaking before being quenched by 5 µL of 5% hydroxylamine for 15 minutes. Samples belonging to the same TMT experiment were then pooled and dried using a vacuum concentrator. Prior to fractionation, the full proteome samples were desalted using Sep-pak tC18 columns (Waters), while the labeled glycopeptides were dissolved into 100 µL of 10% TFA and desalted using C18 stagetips (Rappsilber et al., 2007) made in-house and packed with 1 mg of C18 bulk material (ReproSil-Pur 120 C18-AQ 5 µm, Dr. Maisch) on top of the C18 resin disk (AttractSPE disks bio - C18, Affinisep).

### Porous Graphitic Carbon (PGC) fractionation

Samples were reconstituted in 18 uL buffer A (0.05% TFA in MS grade water supplemented by 2% acetonitrile) and the injection volume was fixed to 16 uL. The glycopeptides were separated on a Hypercarb column (100mm, 1.0mm ID, 3µm particle size, Thermo Fisher Scientific) at a temperature of 50°C and a flow rate of 75 µL/min using an Ultimate 3000 Liquid Chromatography system (Thermo Fisher Scientific). The linear separation gradient started 1 minute after injection and increased from 13% buffer B (0.05% TFA in acetonitrile) to 42% buffer B after 95 minutes, before increasing to 80% buffer B in 5 minutes. The column was washed with 80% buffer B for 5 minutes before being re-equilibrated for 5 minutes with 100% buffer A. Fractions were collected from 4.5 minutes to 100.5 minutes with a 2 minutes collection period, resulting in 48 fractions which were subsequently pooled into 24 fractions, each n fraction being pooled with the n + 24 fraction. Samples were dried using a vacuum concentrator prior to LC-MS/MS analysis.

### LC-MS/MS analysis

All samples were resuspended in a loading buffer containing 1% TFA, 50 mM citric acid and 2% acetonitrile in MS grade water. Peptides were separated using an UltiMate 3000 RSLCnano system (Thermo Fisher Scientific) equipped with a trapping cartridge (Precolumn; C18 PepMap 100, 5 μm, 300-μm i.d. × 5 mm, 100 Å) and an analytical column (Waters nanoEase HSS C18 T3, 75 μm × 25 cm, 1.8 μm, 100 Å). Solvent A was 0.1% formic acid in LC–MS-grade water, and solvent B was 0.1% formic acid in LC–MS-grade acetonitrile. Peptides were loaded onto the trapping cartridge (30 μl min^−1^ solvent A for 3 min) and eluted with a constant flow of 300 nL/min. Peptides were separated using a linear gradient of 8–28%, 5-25% and 6-26% B for 156 minutes for the full proteome, label free glycopeptides and TMT labeled glycopeptides samples, respectively, followed by an increase to 40% B within 4 minutes before washing at 85% B for 4 minutes and re-equilibration to initial conditions. The LC system was coupled to a Fusion Lumos Tribrid mass spectrometer (Thermo Fisher Scientific) operated in positive ion mode with a spray voltage of 2.4 kV and a capillary temperature of 275 °C. Full-scan MS spectra were acquired in profile mode in the Orbitrap using a resolution of 120,000, with a mass range of 375–1,500 m/z, 700-2,000 m/z and 800-2,000 m/z in the case of the full proteome, label-free glycopeptides and labeled glycopeptides, respectively. The maximum injection time was 50 ms and automatic gain control (AGC) was set to 4 × 10^5^ charges. The mass spectrometer was operated in data-dependent acquisition mode and precursors with charge states 2–7 and a minimum intensity of 2 × 10^5^ were selected for subsequent HCD fragmentation with a maximum duty cycle time of 3 seconds. Peptides were isolated using the quadrupole with an isolation window of 0.7 m/z and 1.4 m/z in the case of TMT labeled samples and label free samples, respectively. Precursors were fragmented with a normalized collision energy of 34%, 38% and stepped collision energy of 25, 36 and 45% for the full proteome, label-free glycopeptides and TMT labeled glycopeptides, respectively. A dynamic exclusion window of 45 seconds was used for the full proteome sample and 30 seconds for glycopeptides samples. MS/MS spectra were acquired in profile mode with a resolution of 30,000 in the Orbitrap. The maximum injection time was set to 100 ms for the full proteome and label free glycopeptides and 200ms for TMT labeled glycopeptides. AGC target was set to 1 × 10^5^ charges for full proteome and label free glycopeptides samples and to 2.5 × 10^5^ charges for TMT labeled glycopeptides.

### Data analysis

All raw files were converted to mzmL format using MSConvert from Proteowizard (Chambers et al., 2012), using peak picking from the vendor algorithm and keeping the 300 most intense peaks. Files were then searched using MSFragger v3.7 in Fragpipe v19.1 against the Swissprot *Homo sapiens* database (20,443 entries) in the case of human cells samples, the *Mus musculus* reference proteome (55,402 entries) in the case of label-free mouse brain samples and the *Mus musculus* reference proteome with one protein per gene sequence (22,084 entries) in the case of TMT labeled mouse brain samples. Label-free samples were searched against the large reference proteome, as were raw files from the mouse proteome atlas or the Riley et al. study in order to benchmark our method. TMT labeled samples were searched against a smaller database in order to reduce non-unique peptides, which were filtered off. Overall, total numbers of identification was similar with the two databases.

The default glyco-N-HCD, glyco-O-HCD and glyco-N-TMT workflows were used for the label-free N-glyco, label-free O-glyco and TMT labeled N-glyco database searches, respectively. In the case of N-glycosylation, a custom database containing 1799 unique glycan compositions (Supplementary Data 1) was used while the default MSFragger database was used for O-glycosylation. Glycan compositions were assigned using PTM-Shepherd. In the case of TMT labeled samples, TMTpro modification was fixed on lysine and variable on peptide N-terminus and channel normalization was disabled. The psm.tsv and protein.tsv output files were used for subsequent data analysis. In the case of O-glycosylation, all glycopeptides containing the N-glycosylation sequon (N-X-S/T) were removed to prevent misassignment. Peptides with a unique amino acid sequence combined with a unique glycan composition were considered as unique glycopeptides.

For downstream analysis, we extracted glycopeptides from the psm.tsv files and categorized them into seven or eight glycan classes for N- and O-glycosylation, respectively. The categories were defined as following: (i) phospho, containing a phosphate group, (ii) sialylated, containing a NeuAc sugar, (iii) fucosylated, containing fucose, (iv) high mannose, combining a HexNAc(2) core with between four and 12 hexose molecules, (v) paucimannose, combining a HexNAc(2) core with between one and three hexose molecules, (vi) small, composed of only a core with less than three HexNAc molecules and (vii) complex/hybrid, the rest. For O-glycosylation data, an additional sulfated class was defined for glycosylations containing a sulfation. Evidence annotation of known glycosites was downloaded from UniProt.

All included enrichment analyses were performed using the STRINGdb R package (Szklarczyk et al., 2021), version 11.5, with the full *Mus musculus* or *Homo sapiens* network. The default hypergeometric test and p-value adjustment procedure was used. Significant pathways were extracted by using a FDR cutoff of 5% (Supplementary Data 4). For the domain enrichment analysis, we performed an over-representation analysis on the contingency matrix of InterPro domains (Paysan-Lafosse et al., 2023) and glycan classes using Fisher’s exact test (Supplementary Data 4). p-values were adjusted for multiple testing using the Benjamini-Hochberg method. Significant domains were extracted by using an adjusted p-value cutoff of 0.1%.

To compare the similarity of glycosylation profiles of sites on one protein, we first prefiltered for proteins with more than one site. We then calculated the pairwise correlation of the number of all possible observed modifications between all sites of a protein using the Kendall rank correlation coefficient. To generate random correlations, we shuffled all glycosites to randomly assign glycan compositions to them. To test whether the observed difference of random and true correlations of sites is significant we performed a Wilcoxon signed rank test.

To investigate the distance between phosphosites and glycosites for different glycan types, we downloaded all known *Mus musculus* phosphosites from Uniprot. We defined a phosphosite as close if it is located +/− 5 residues around the glycosite in case of N-glycosylation and +/− 5 residues around the peptide start position in the case of O-glycosylation.

For the quantitative proteomics and glycoproteomics data of brains of mice exposed to different microbiome species, we first normalized the TMT reporter intensities for glycopeptides and proteins using the normalizeVSN function of the limma R package(Ritchie et al., 2015). To check the reproducibility of quantification, the pairwise Spearman correlation of all conditions and replicates was calculated. We corrected glycopeptide intensities per protein by removing abundance-derived intensity using a linear regression for all glycopeptides with a matched total protein intensity. The corrected and normalized glycopeptide intensities as well as the normalized protein intensities were used to generate PCA plots and the differential analysis using the limma R package (Ritchie et al., 2015). The intensity matrices were filtered for only features with complete quantification across all conditions (i.e. TMT reporter ion intensity different from 0 for all channels), resulting in 56171 glycopeptides and 9744 proteins in total. For the design, we considered the microbiome group as well as the sex of the mice. Contrasts were set for the comparison of the different microbiome groups. A significant change was assigned for glycopeptides or proteins with an absolute log2 fold-change > log2(1.5) and an adjusted p-value < 0.05.

### Topological, structural and conservation analysis of identified glycosites

To analyze the N-glycosylation data, we compared asparagines that were identified as being glycosylated to those that were not but had the potential to be glycosylated on the same proteins (i.e., N that are part of the glycosylation sequons N-X-S/T, X≠P, referred to as ngN). As O-glycosylation sites could not be confidently localized by HCD fragmentation, we assigned a putative glycosite by arbitrarily selecting the most central serine or threonine residue in the peptide. In cases where there was a tie, we used the ceiling. All serine and threonine residues identified in the glycoproteins were then used as a background for the O-glycosylation over-representation analysis.

We retrieved topological domain annotations for the proteins identified in the N-glycosylation and O-glycosylation mouse brain data from UniProt. We filtered these annotations to include only the two classes of interest, which were “cytoplasmic” and “extracellular”. Out of the 3,199 proteins identified in the N-glycosylation data and 2,365 in the O-glycosylation data, we found at least one annotation for 1,100 and 845 proteins, respectively. We used these subsets of proteins to conduct an over-representation analysis of glycosites that are located inside or outside the topological domains.

We used the structuremap python package (https://github.com/MannLabs/structuremap) to generate all of the structural annotations. Initially, we retrieved predicted structures for all proteins identified in the N-glycosylation data from the AlphaFold Protein Structure Database (Varadi et al., 2022) (2,807 structures out of the 3,199 glycoproteins identified). Residues were then annotated using the prediction-aware part-sphere exposure (pPSE) metric (Bludau et al., 2022), which scores disorder for each residue based on the number of residues in their structural proximity, taking into account the AlphaFold prediction error. As in the original manuscript (Bludau et al., 2022), we used a threshold of 34.27 on the smoothed pPSE (180°, 24Å) to determine if a protein site was part of an intrinsically disordered region. For the solvent accessibility analysis, we only considered sites in structured regions (not intrinsically disordered), and we used a cutoff of 5 amino acids in the directional, shorter pPSE (70°, 12Å) to determine if a protein site was solvent-accessible.

For the conservation analysis, we used SIFT scores (Ng and Henikoff, 2003). Initially, we retrieved precomputed SIFT scores at the genome level from the SIFT database (https://sift.bii.a-star.edu.sg/sift4g/). Since SIFT scores are calculated based on nucleotide substitutions, we obtained protein-level SIFT scores by averaging the SIFT scores for every potential amino acid substitution.

We used Fisher’s exact test to conduct the site-level over-representation analysis. For each region of interest, we created a contingency matrix formed by the sites located inside and outside the regions. When comparing average relative position and SIFT scores between glycosite groups, we employed the Wilcoxon Rank Sum Test.

To visualize the glycosites of interest in the excitatory amino acid transporter 2, and given that the mouse protein structure has not yet been resolved, we obtained the resolved structure for its human ortholog (sequence similarity > 95%). We first retrieved the structure and its membrane orientation prediction from the OPM server (Kato et al., 2022; Lomize et al., 2012) (accession number 9537). Since the structure of the TM4b-TM4c extracellular loop, where both glycosites are located, has not been resolved, we utilized AlphaFold2 predictions to visualize this specific region. Initially, we retrieved the predicted protein structure from AlphaFoldDB (Jumper et al., 2021; Varadi et al., 2022). Subsequently, we performed a structural alignment of the template and predicted structure for each monomer within the sequence window 190-235 using the MatchMaker algorithm with default parameters as implemented in the ChimeraX software (Meng et al., 2006).

## Code availability

All code used to perform the computational analyses described and to reproduce the figures is available at https://github.com/miralea/DQGlyco.

## Data Availability

All raw files, search parameters and search outputs were deposited to the ProteomeXchange Consortium via the PRIDE partner repository with the dataset identifier PXD042237 and will be made accessible upon publication.

## Acknowledgments

We thank A. Mateus and P. Cossart for insightful discussions and feedback on the manuscript. This work was supported by the European Molecular Biology Laboratory. C.M.P. was supported by a fellowship from the EMBL Interdisciplinary Postdoc (EI3POD) programme under Marie Skłodowska-Curie Actions COFUND (grant number 664726). A.T. is supported by an ERC consolidator grant, uCARE. M.M.S. is supported by the Allen Distinguished Investigator award through the Paul G. Allen Frontiers Group. We would like to thank all members of the Savitski, Zimmermann, and Typas groups for helpful discussions, the Proteomics Core Facility at the EMBL for expert help and George Maftei for assistance with animal experiments.

## Author contributions

C.M.P. and M.M.S. designed the study. C.M.P. developed the methodology and performed mass spectrometry experiments with help from M.L.B. and I.B.. C.M.P., M.L.B., and M.G-R. developed data analysis strategies and analyzed the data. A.B-N. performed mouse experiments and was supervised by M.Z.. M.Z. and A.T. provided scientific input on study design and data interpretation. C.M.P. and M.M.S. drafted the manuscript with contribution from M.L.B., M.G-R., I.B., M.Z., and A.T. which was reviewed and edited by all authors. C.M.P. and M.M.S. supervised the study.

---

**Figure S1:**
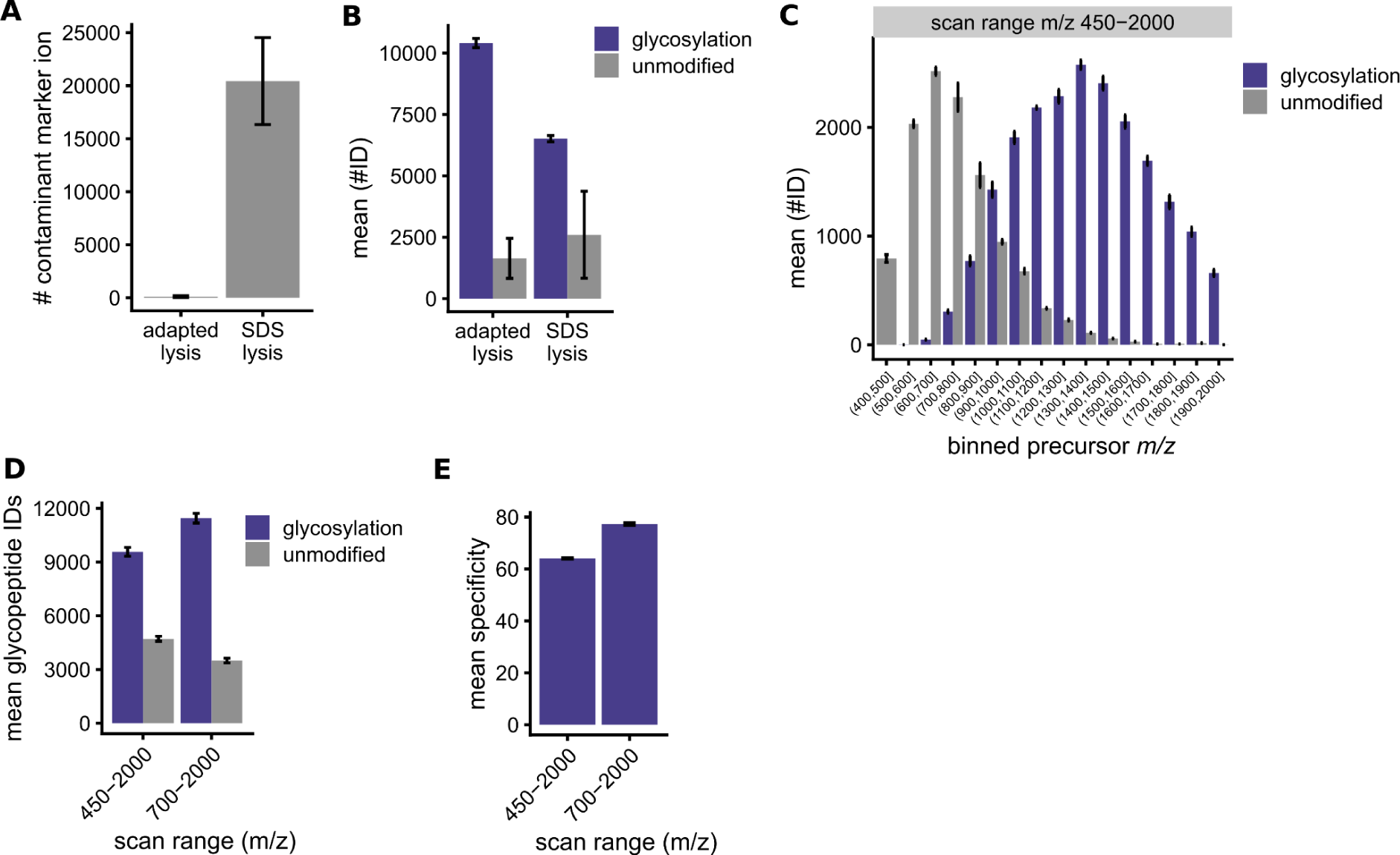
Optimization of DQGlyco. **A** Mean number of contaminant ions identified using a standard SDS lysis in comparison to an adapted optimized lysis in HEK293T cells. All samples were analyzed in duplicates. **B** Mean number of unique glycopeptides and unmodified peptides identified using a standard SDS lysis in comparison to an adapted optimized lysis in HEK293T cells. All samples were analyzed in triplicates. **C** Mean number of glycopeptides and unmodified peptides identified in HEK293T per scan range bin from 400-2000 m/z. All samples were analyzed in triplicates. **D** Mean number of glycopeptides and unmodified peptides identified for measurements with two different scan ranges in HEK293T cells. All samples were analyzed in triplicates. **E** Mean specificity (number of glycopeptide spectrum matches divided by all peptide spectrum matches) for measurements with two different scan ranges in HEK293T cells.

**Figure S2:**
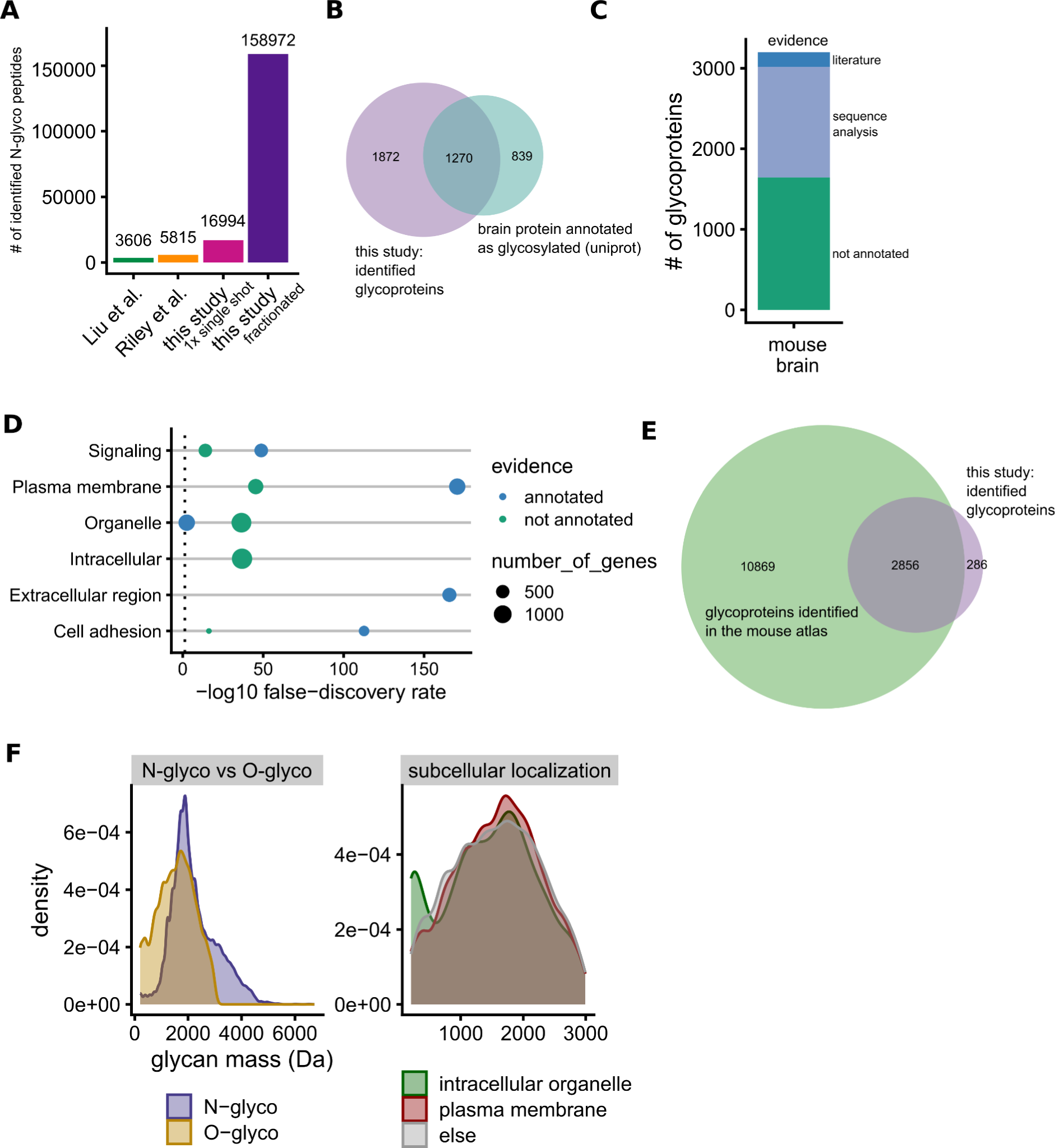
In-depth characterization of N- and O-glycosylation in the mouse brain. **A** Total number of unique glycopeptides (unique sequence and glycan composition) identified in this study and two recent N-glycoproteomic studies using off-line fractionation. **B** Overlap of glycoproteins identified in this study and proteins identified in the mouse brain which are annotated as glycosylated in the Uniprot database. **C** Uniprot annotation of glycoproteins identified in mouse brain samples. Literature evidence corresponds to experimental evidence while the majority of sites is annotated based on sequence analysis, i.e. prediction of glycosite based on the presence of the N-X-S/T, X≠P, glycosylation sequence. **D** Results of the Gene Ontology enrichment analysis (stringdb) of N-glycoproteins with and without annotation in the uniprot database. **E** Overlap of all proteins identified in the mouse brain of the mouse atlas and glycoproteins identified in this study. **F** Frequency of (i) O- and N-glycopeptides per glycan mass, (ii) O-glycopeptides of proteins assigned to a specific Gene Ontology term per glycan mass.

**Figure S3:**
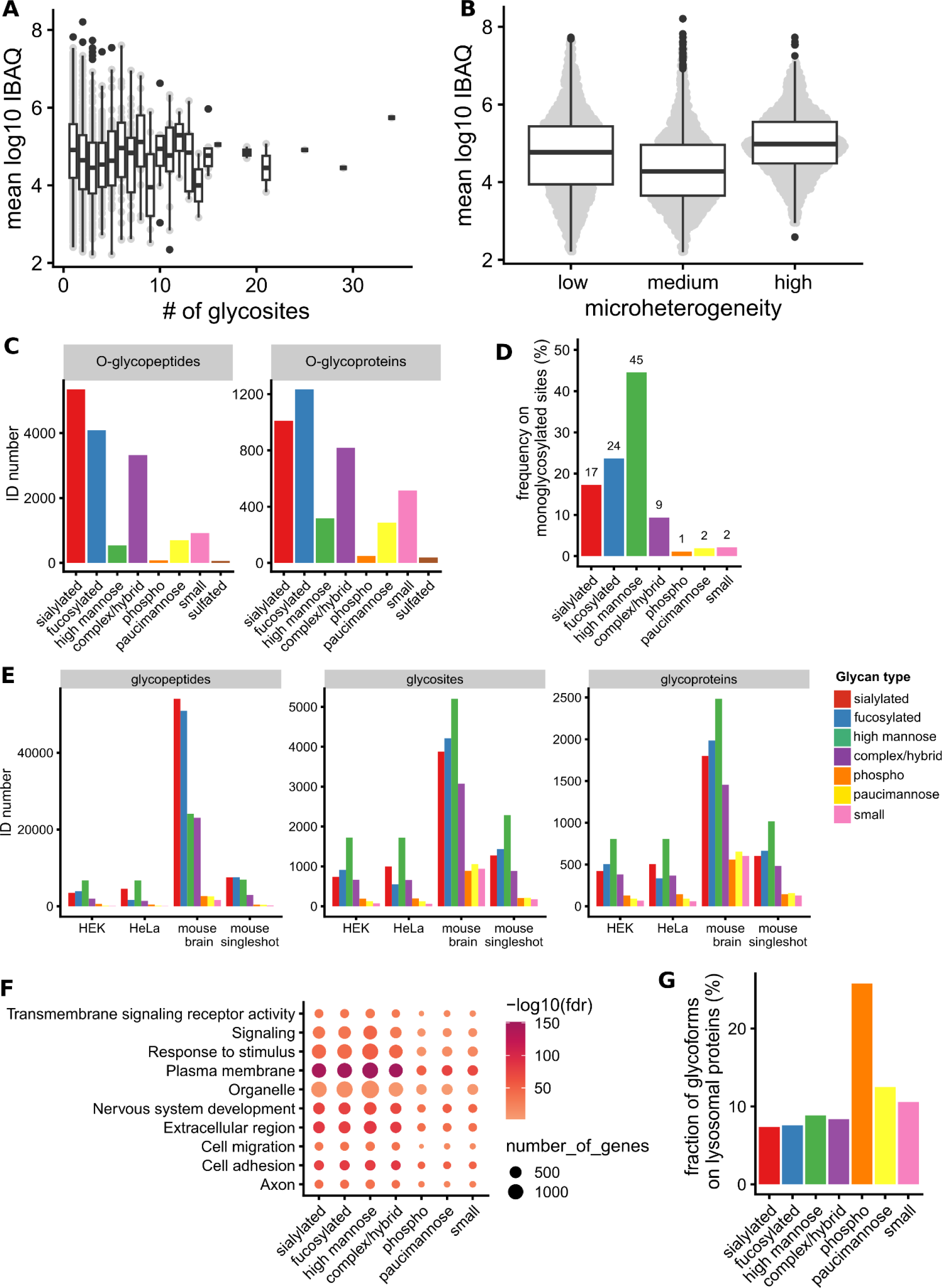
**A** Estimated abundances (IBAQ values) of glycoproteins quantified in the mouse brain compared with the number of sites per protein identified in this study. **B** Microheterogeneity of sites compared to the mean estimated abundance of the corresponding protein quantified in the mouse brain. **C** Total number of unique O-glycopeptides and O-glycoproteins per glycan class for the PGC fractionated mouse brain sample. **D** Frequency of N-glycosites for which only one glycan composition was identified in the PGC fractionated mouse brain sample per glycan class. **E** Total number of unique N-glycopeptides, N-glycosites and N-glycoproteins per glycan class for HEK293T cells single shot, HeLaK cells single shot, mouse brain single shot and the PGC fractionated mouse brain samples. **F** Results of the Gene Ontology enrichment analysis of N-glycoproteins with sites that are modified with different glycan classes. **G** Frequency of O-glycopeptides that map to lysosomal proteins per glycan class.

**Figure S4:**
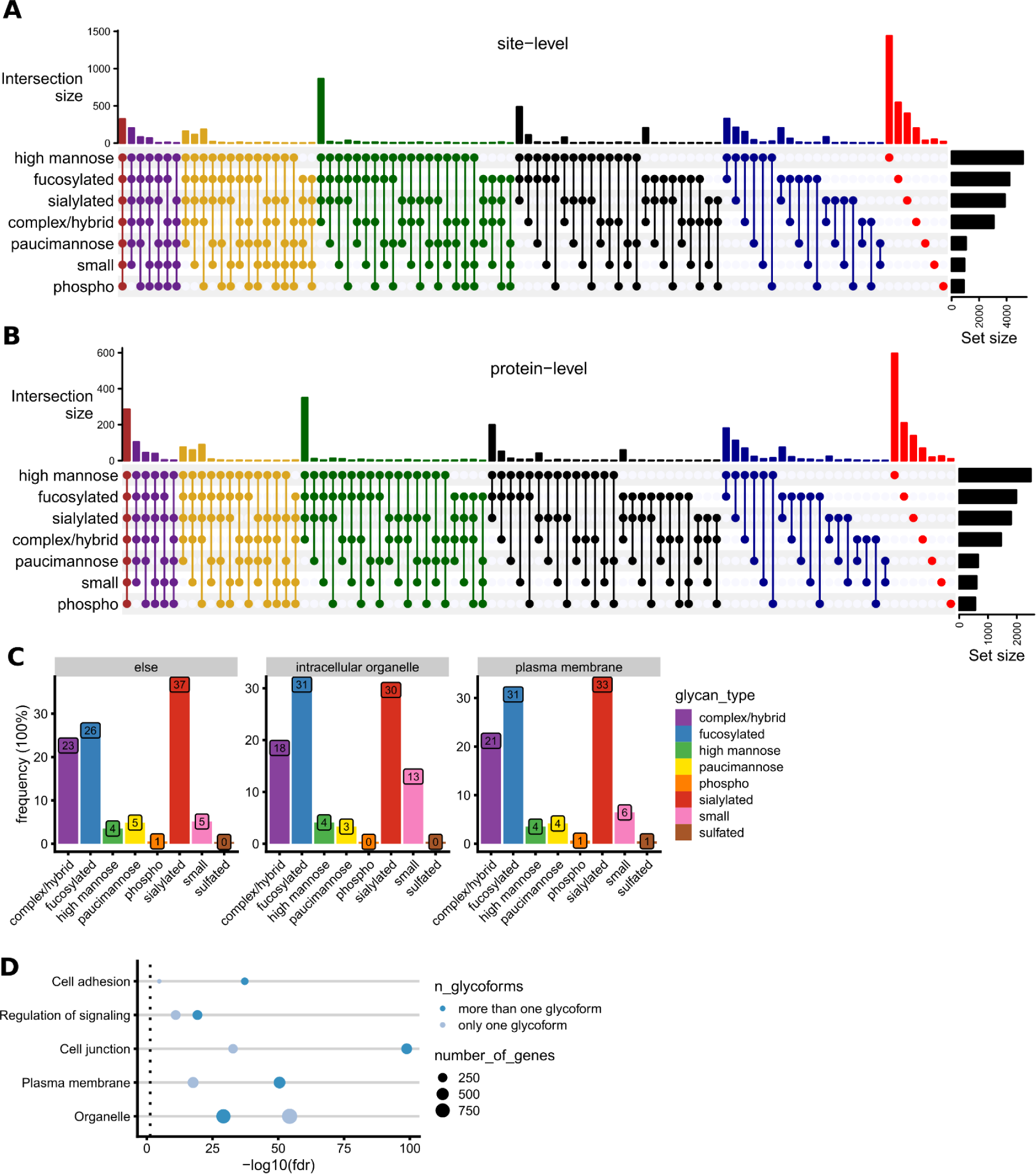
**A** Co-occurrence of glycan classes on N-glycosites displayed as an Upsetplot. **B** Co-occurrence of glycan classes on N-glycoproteins displayed as an Upsetplot. **C** Frequencies of glycan classes of O-glycoproteins for selected Gene Ontology terms. **D** Gene Ontology enrichment results (stringdb) for O-glycoproteins with sites that are either modified with only one glycan composition or with more than one glycan composition.

**Figure S5:**
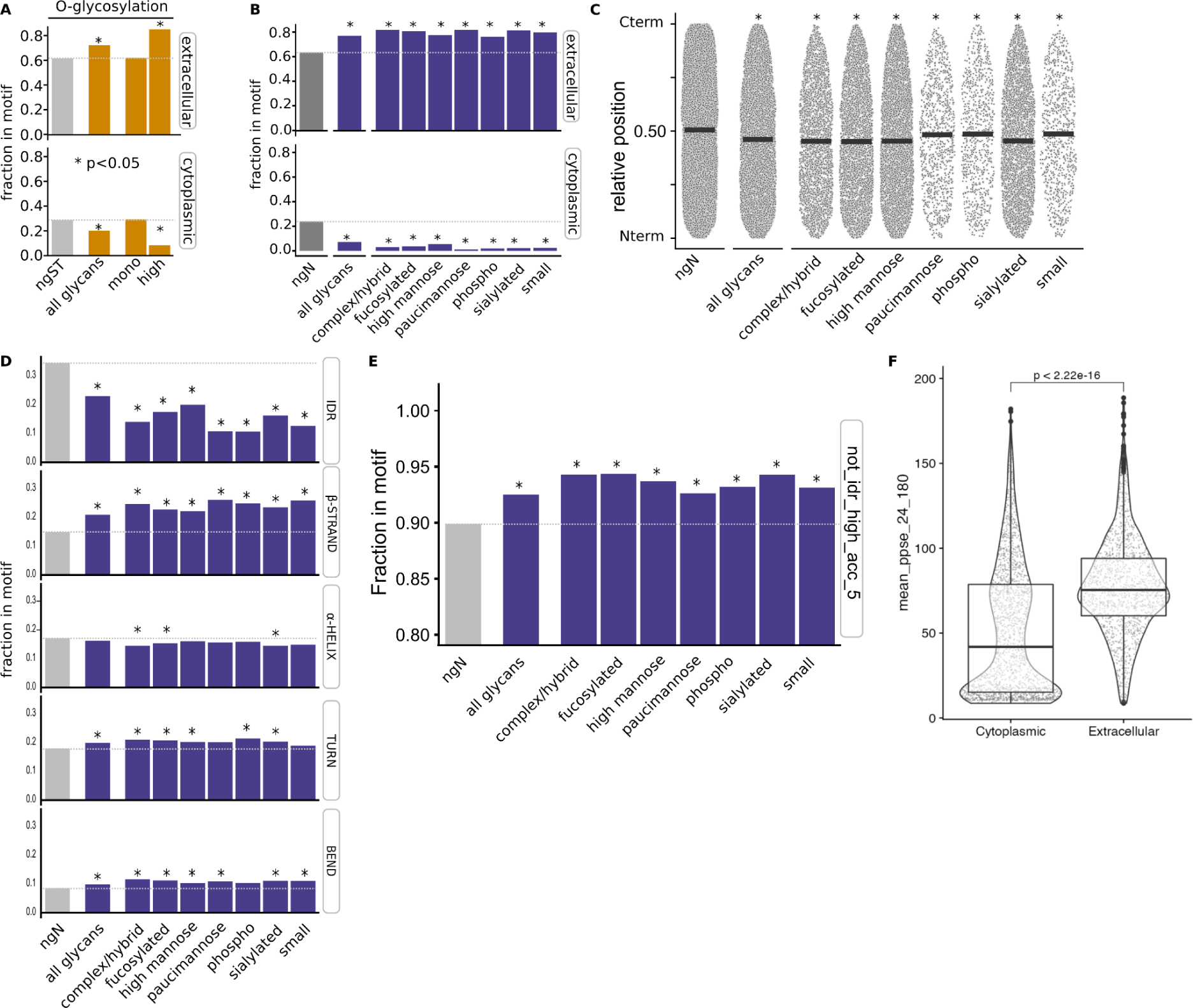
**A** Putative O-glycosylated sites exhibiting high microheterogeneity are significantly enriched in extracellular and depleted in cytoplasmic regions of glycoproteins (N = 1, and N > 1 referred to as monoglycosylated and high microheterogeneity, respectively). Barplots indicate the fraction of putative O-glycosites inside the region of interest. Asterisks indicate Fisher’s exact test p-value < 0.05 using all non-glycosylated Serines and Threonines residues in the proteins as background. **B** Enrichment of glycosites divided by glycan type in extracellular and cytoplasmic regions. **C** Relative protein position of glycosites divided by glycan type. **D** Frequency of glycosites divided by glycan type on defined structural motifs. **E** Enrichment of glycosites divided by glycan type in solvent-accessible regions. **F** Extracellular topological domains are found on more structured regions of proteins. Each point represents the average smoothed pPSE (180°, 24Å) across all the positions in a given topological domain. Displayed P value corresponds to the results of Wilcoxon Rank Sum Test used to compare means between both groups.

**Figure S6:**
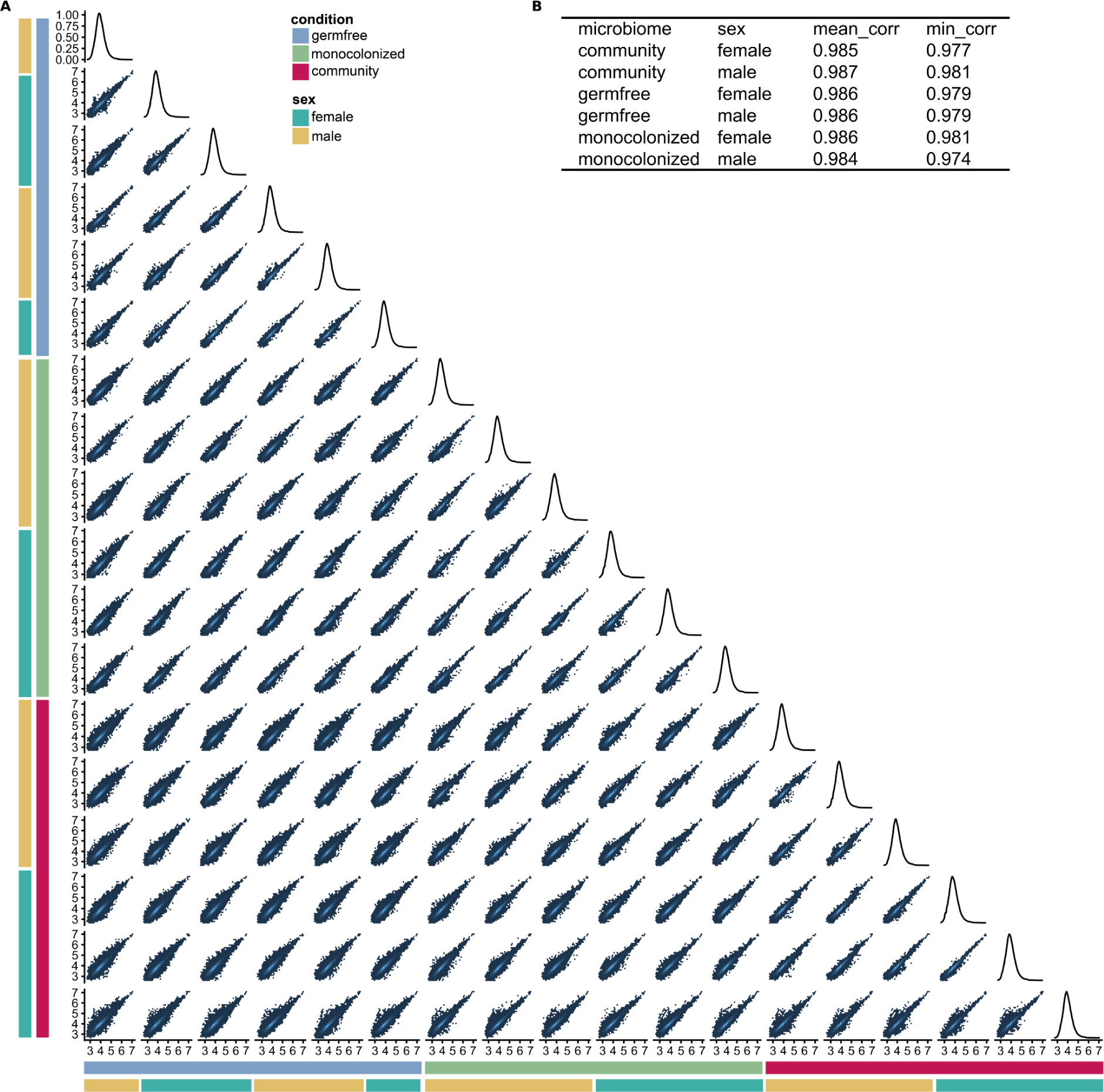
Reproducibility of quantitative N-glycosylation data of mouse brains for different microbiome groups. **A** Pairwise comparison of the reporter intensities of all replicates of all microbiome groups. **B** The mean and minimum spearman correlation of all biological replicates of the different conditions.

**Figure S7:**
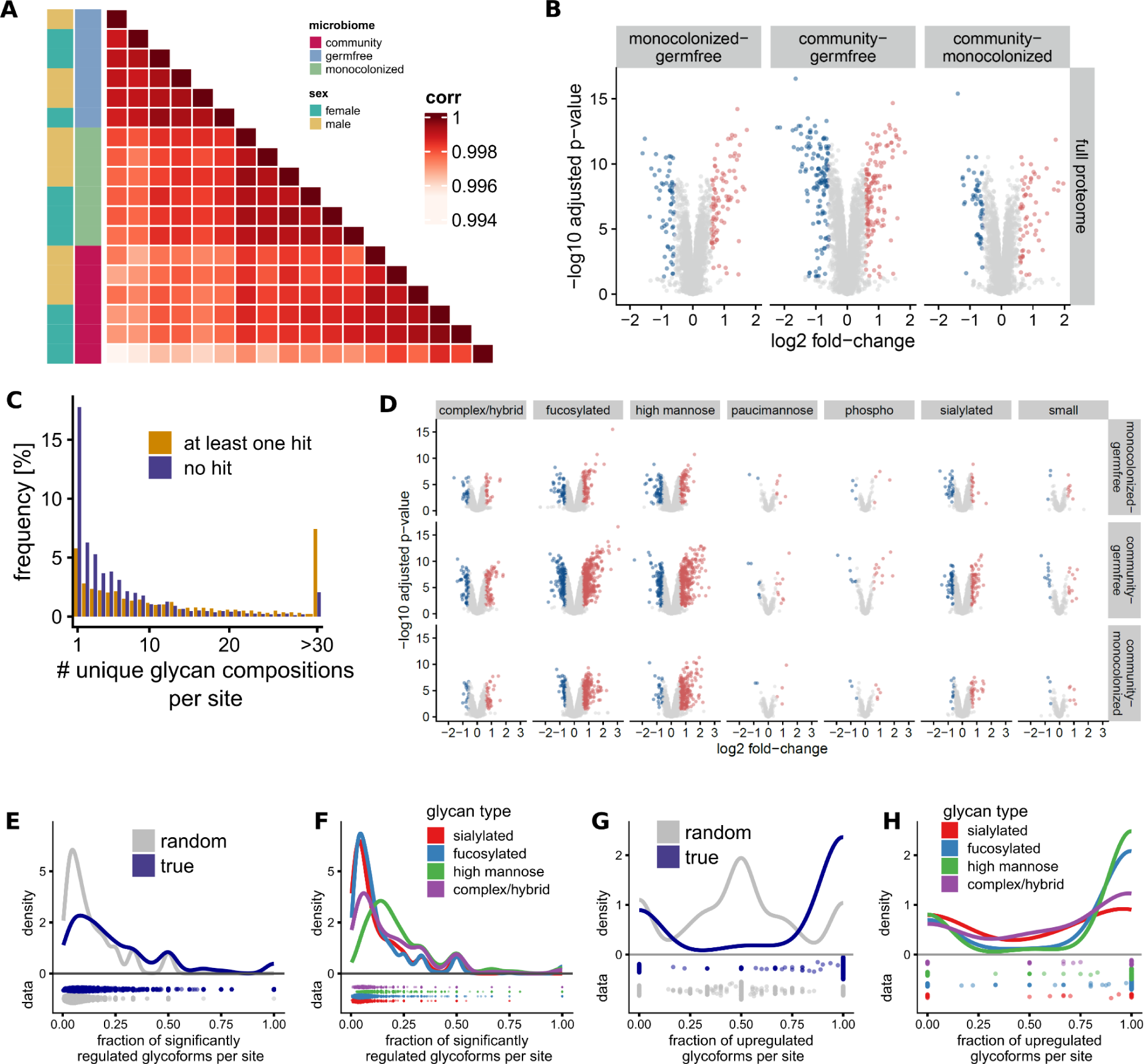
Quantitative profiling of gut microbiome dependent changes in the mouse brain proteome and N-glycoproteome. **A** Spearman correlation of protein intensities between biological replicates. **B** Protein-level regulation in the mouse brain depending on the gut microbiome composition. **C** Number of unique glycan compositions per site for sites exhibiting at least one significantly regulated glycoform or no glycoform regulated **D** Regulation of glycopeptides, with glycopeptides being grouped by glycan class. **E** Fraction of glycoforms which are regulated significantly per site with at least two glycoforms (blue) compared to randomly assigned regulation annotation (grey). On sites with multiple glycoforms detected only a small fraction of glycoforms changes significantly depending on the gut microbiome composition, suggesting that there is no general regulation at the glycosite level. **F** The fraction of glycoforms which are regulated significantly per site and glycan class with at least two glycoforms per class indicates that glycosite level regulation does not depend on a certain glycan class (i.e all fucosylated forms on one site are significantly upregulated). Only glycan types with enough data points shown. **G** Fraction of glycoforms which are upregulated out of all significantly regulated glycoforms per site compared to randomly assigned regulation direction (grey). For glycosites with at least one significantly regulated glycoform, most regulated glycoforms tended in the same direction of regulation, as opposed to random direction of regulation. **H** This trend for the direction of regulation can also be observed for different glycan classes. Only glycan types with enough data points shown.

