## Supplementary Data 5 for "Deep quantitative glycoproteomics reveals gut microbiome induced remodeling of the brain glycoproteome"

### Intercellular adhesion molecule 5

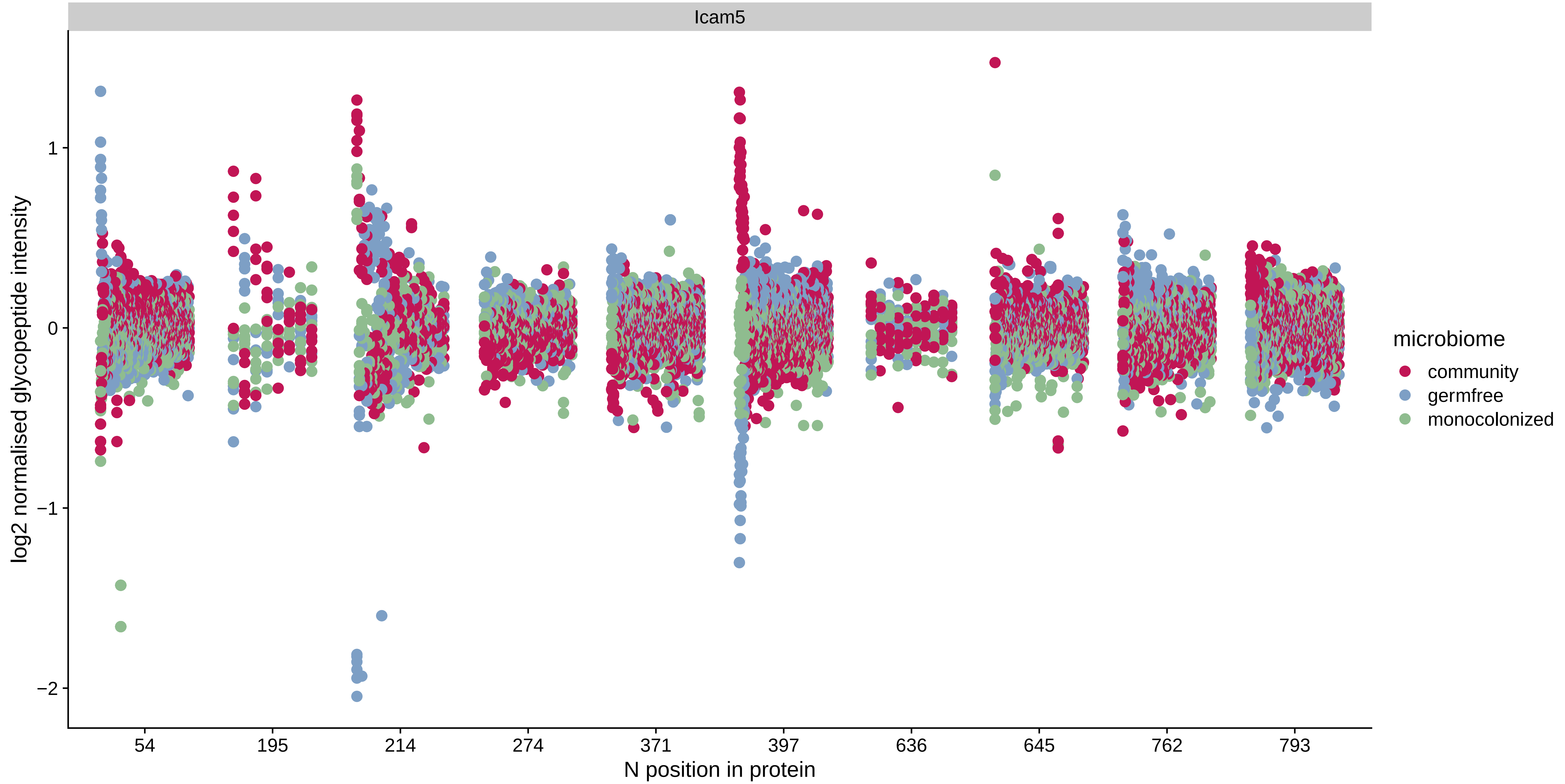

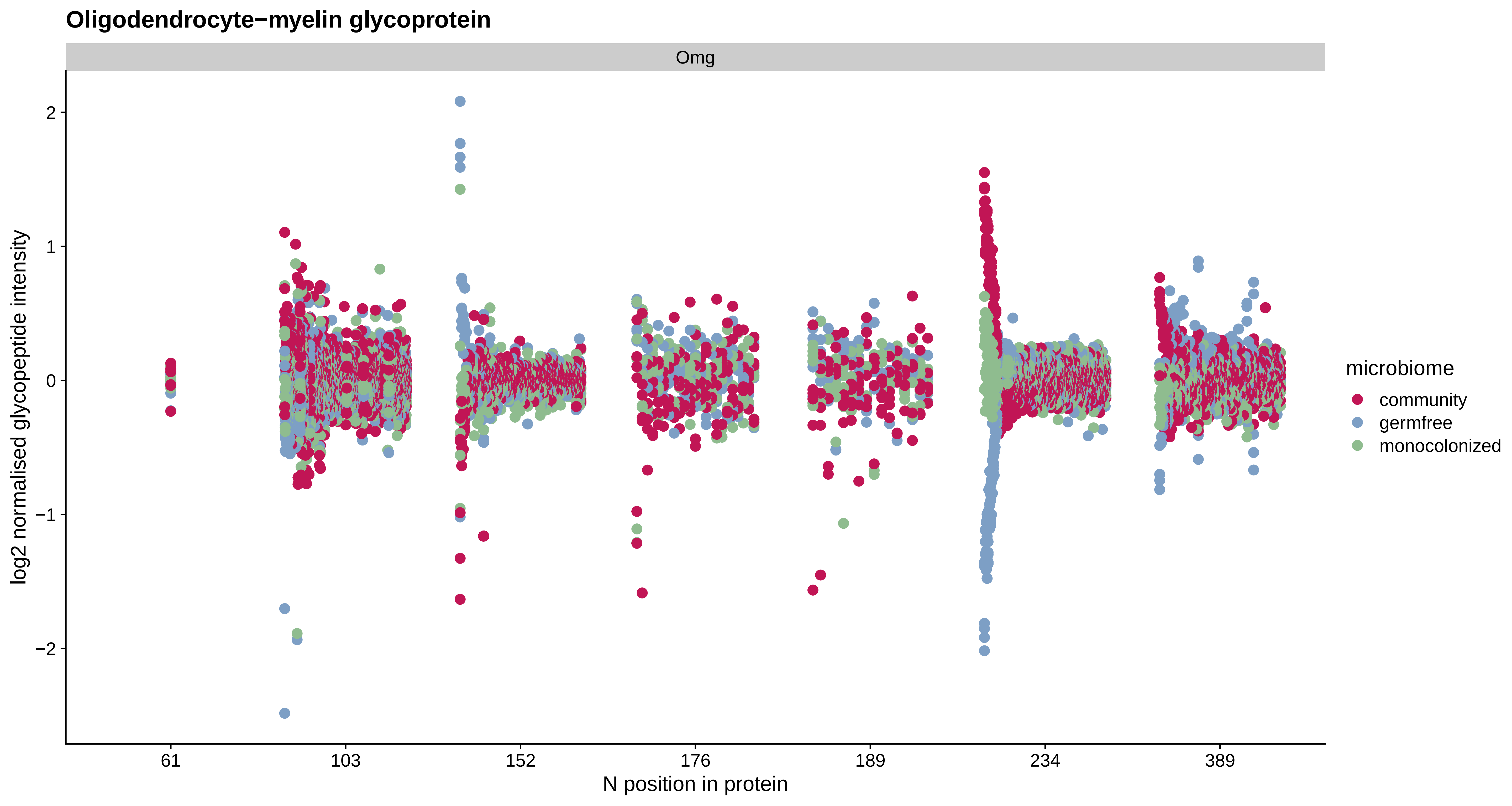

Oligodendrocyte−myelin glycoprotein
Omg
log2 normalised glycopeptide intensity
2
1
0
−1
−2
61
103
152
176
189
234
389
N position in protein
microbiome
community
germfree
monocolonized

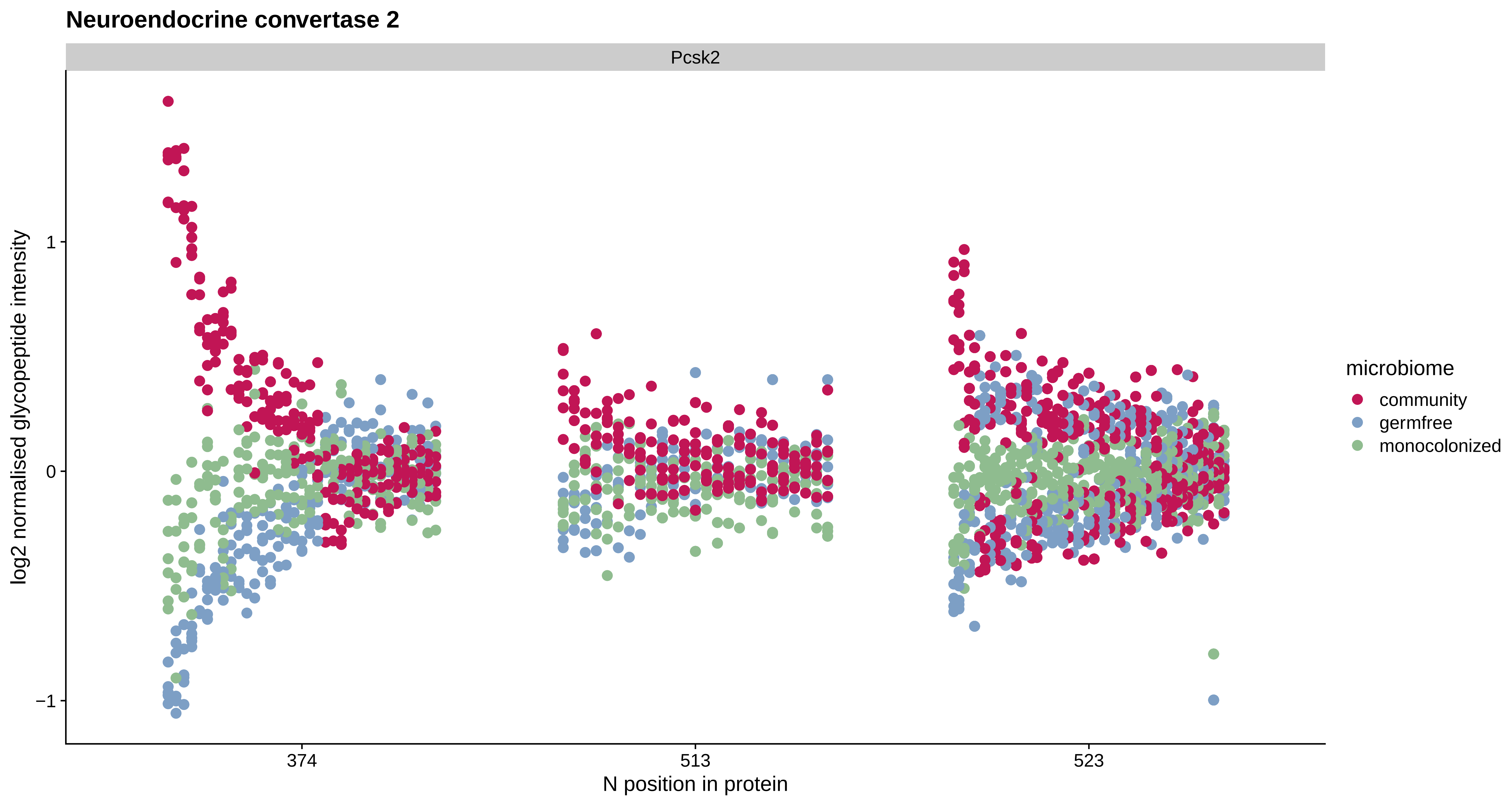

Neuroendocrine convertase 2
Pcsk2
log2 normalised glycopeptide intensity
1
0
−1
374
513
523
N position in protein
microbiome
community
germfree
monocolonized

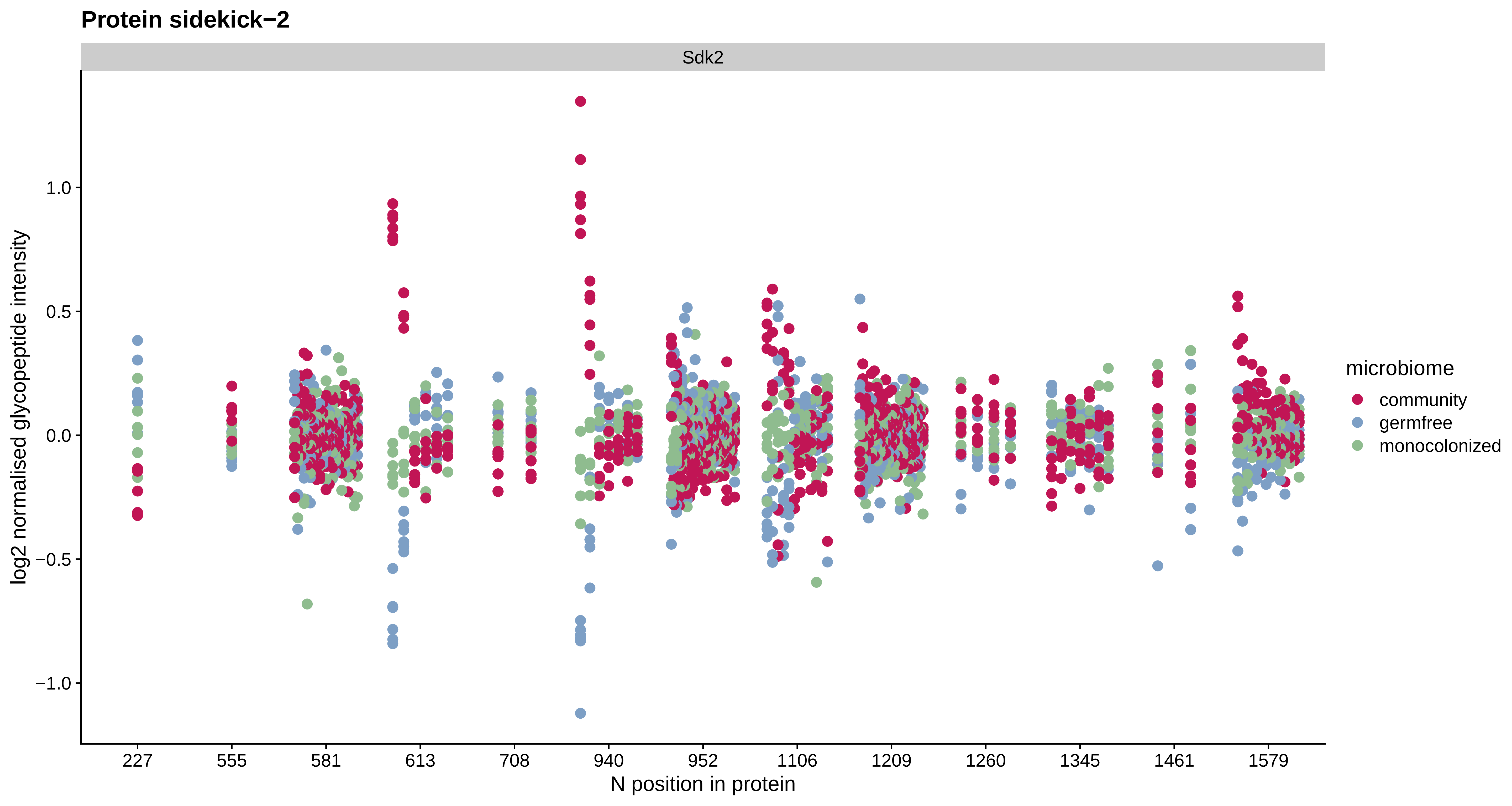

Protein sidekick−2
Sdk2
log2 normalised glycopeptide intensity
1.0
0.5
0.0
−0.5
−1.0
227
555
581
613
708
940
952
1106
1209
1260
1345
1461
1579
N position in protein
microbiome
community
germfree
monocolonized

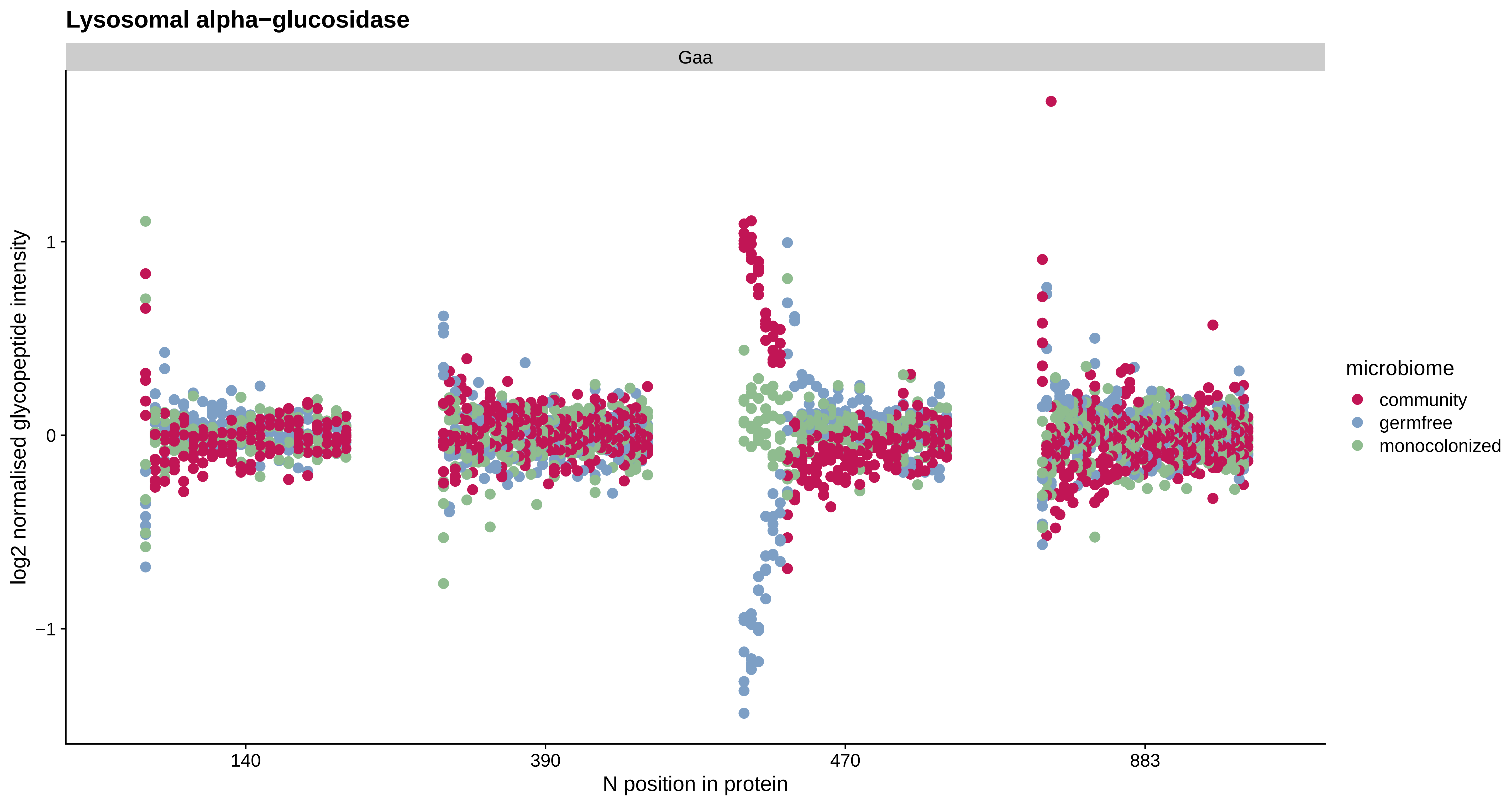
Lysosomal alpha−glucosidase
Gaa
log2 normalised glycopeptide intensity
1
0
−1
140
390
470
883
N position in protein
microbiome
community
germfree
monocolonized

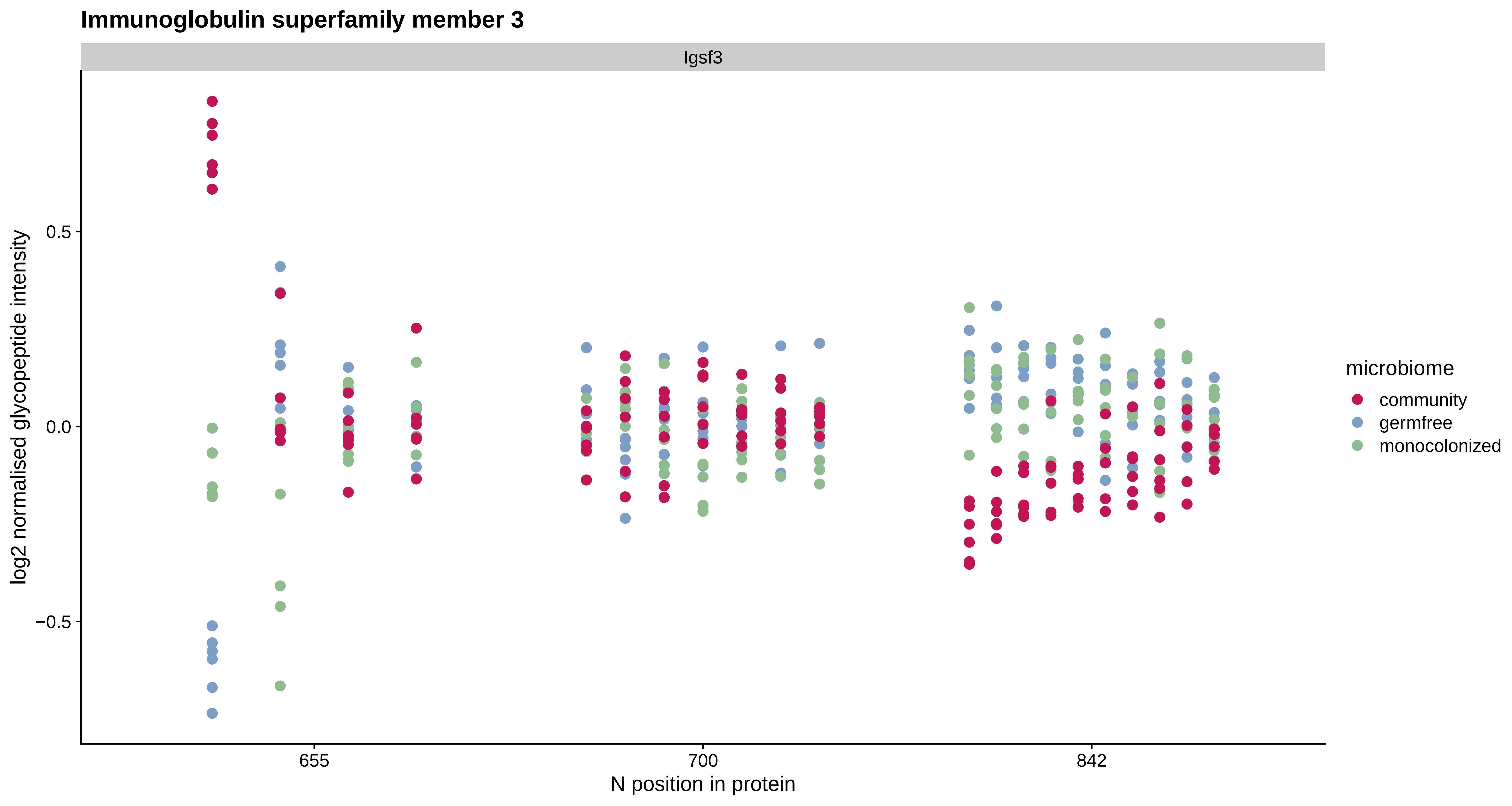
Immunoglobulin superfamily member 3
Igsf3
log2 normalised glycopeptide intensity
0.5
0.0
−0.5
655
700
842
N position in protein
microbiome
community
germfree
monocolonized

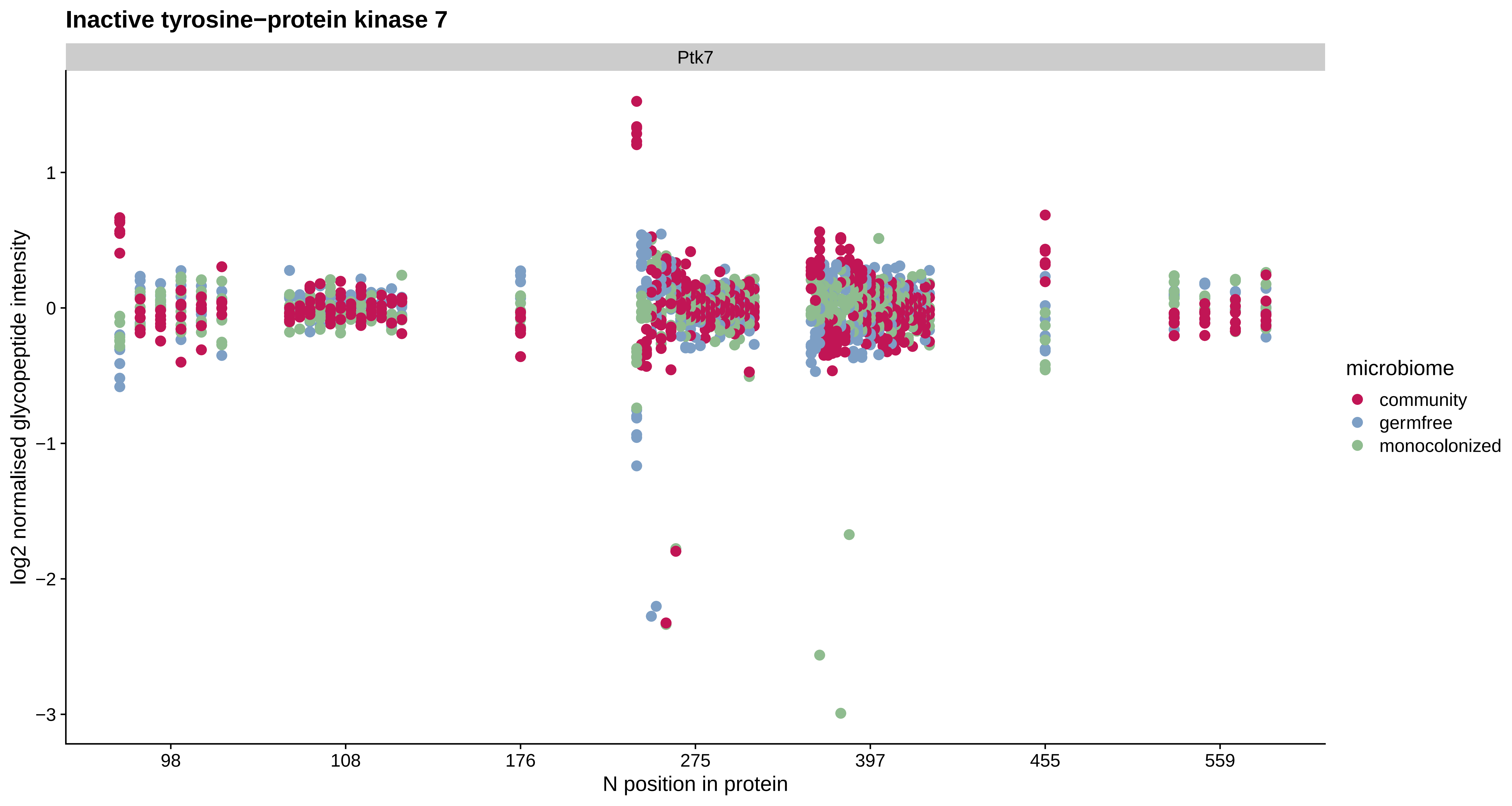

Inactive tyrosine−protein kinase 7
Ptk7
log2 normalised glycopeptide intensity
1
0
−1
−2
−3
98
108
176
275
397
455
559
N position in protein
microbiome
community
germfree
monocolonized

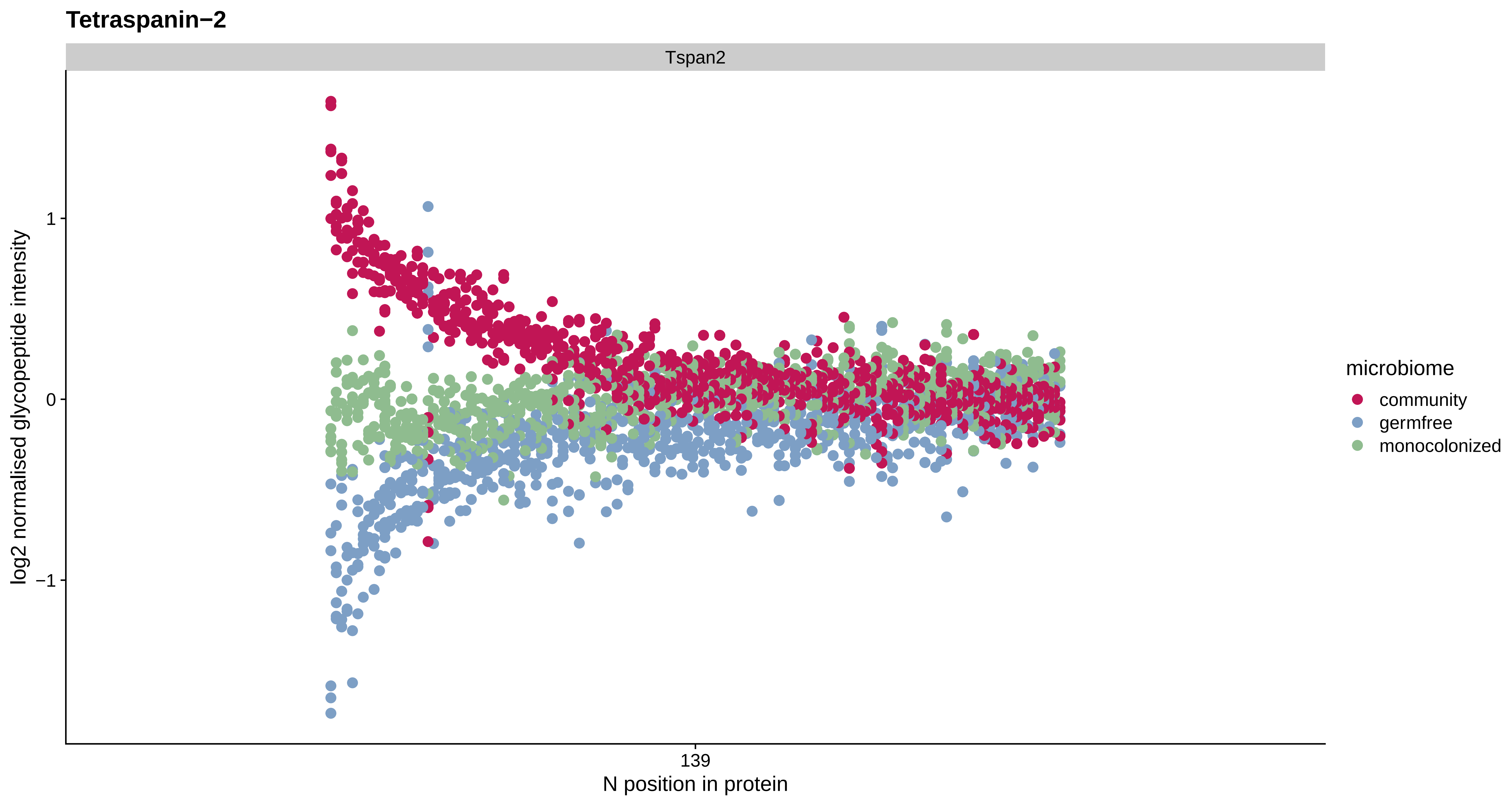
Tetraspanin−2
Tspan2
log2 normalised glycopeptide intensity
1
0
−1
139
N position in protein
microbiome
community
germfree
monocolonized

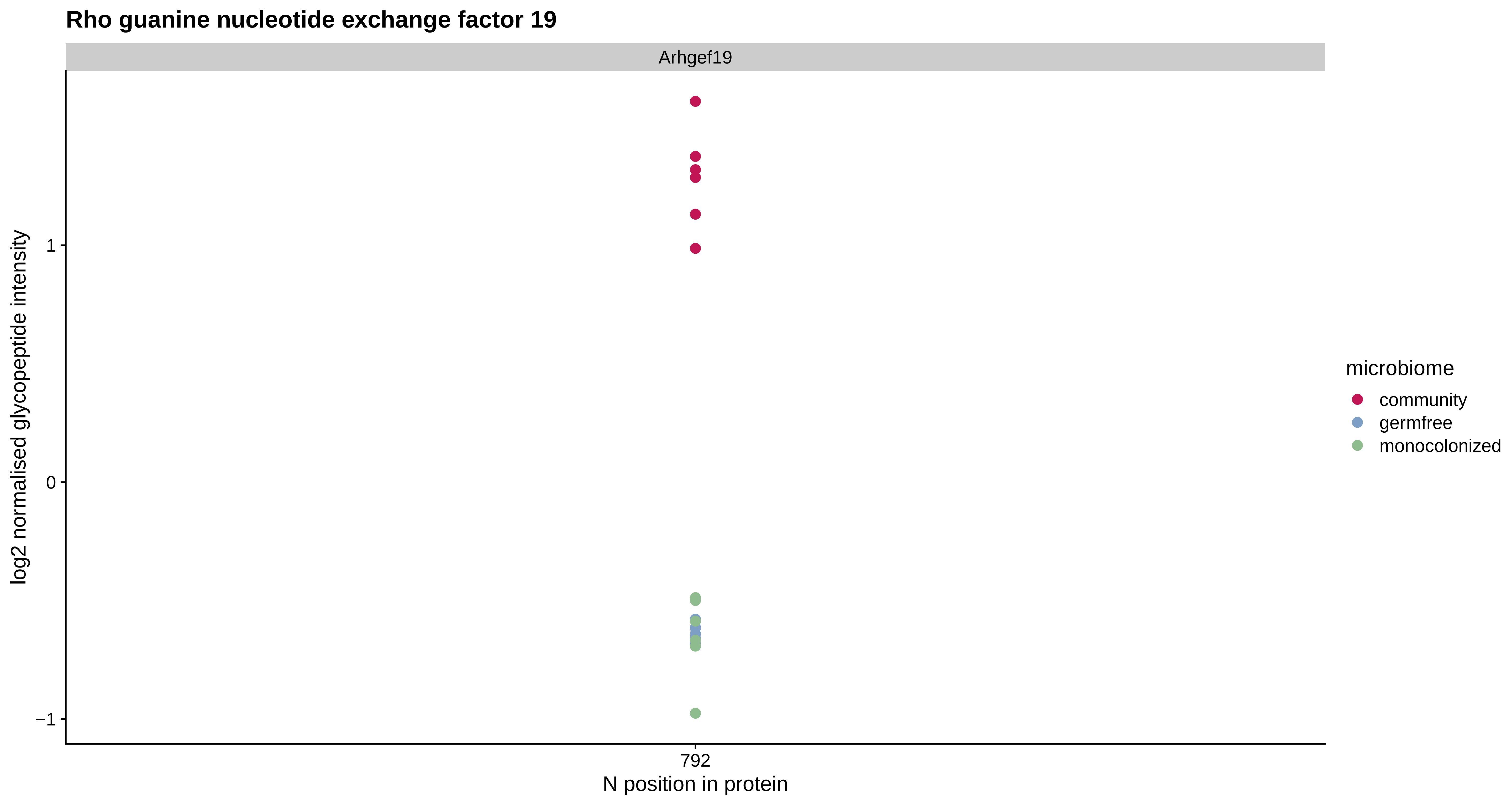

Rho guanine nucleotide exchange factor 19
Arhgef19
log2 normalised glycopeptide intensity
1
0
−1
792
N position in protein
microbiome
community
germfree
monocolonized

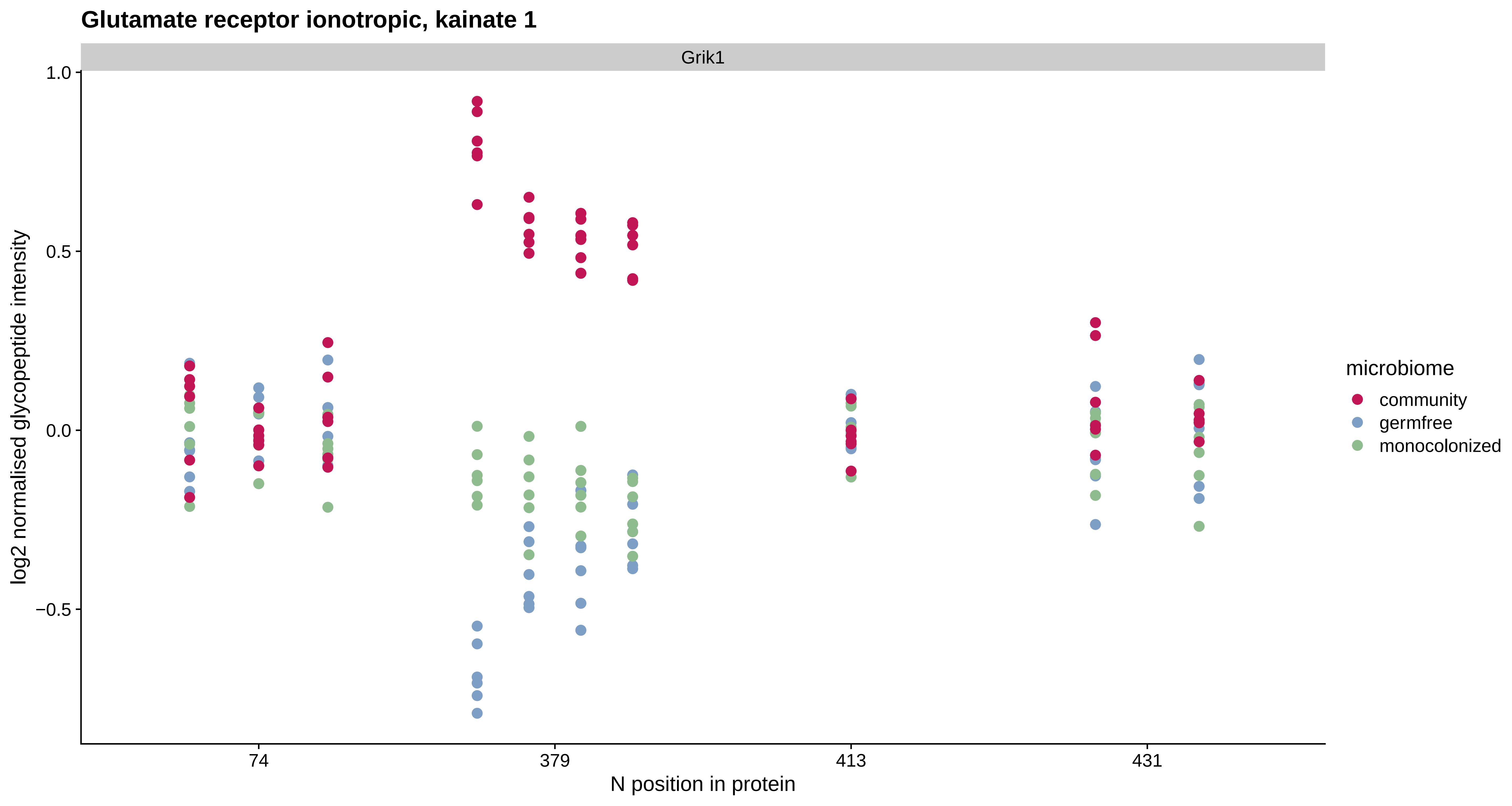
Glutamate receptor ionotropic, kainate 1
Grik1
1.0
0.5
0.0
−0.5
log2 normalised glycopeptide intensity
74
379
413
431
N position in protein
microbiome
community
germfree
monocolonized

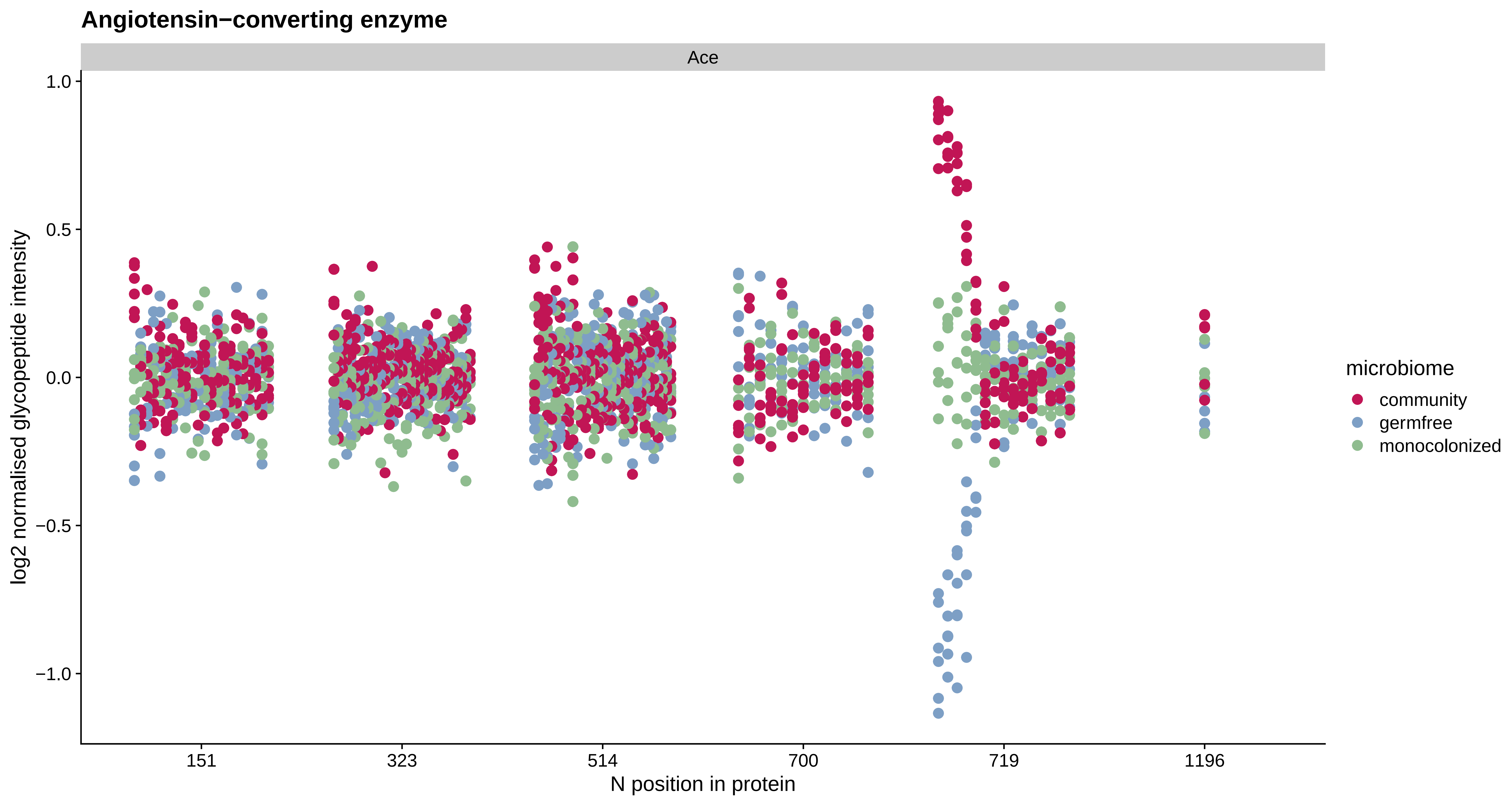

Angiotensin−converting enzyme
Ace
1.0
0.5
0.0
−0.5
−1.0
log2 normalised glycopeptide intensity
151
323
514
700
719
1196
N position in protein
microbiome
community
germfree
monocolonized

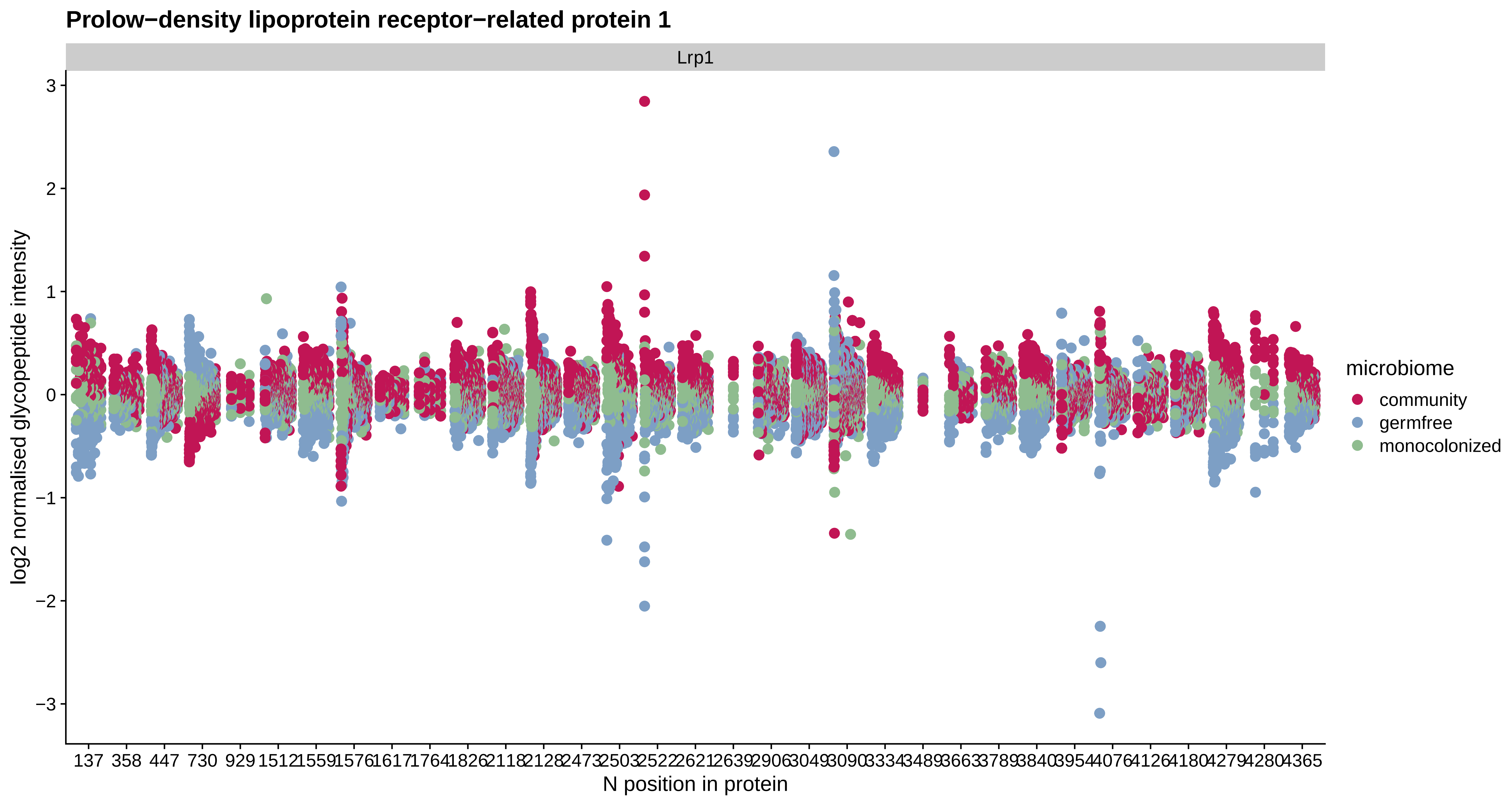
Prolow−density lipoprotein receptor−related protein 1
Lrp1
log2 normalised glycopeptide intensity
3
2
1
0
−1
−2
−3
137 358 447 730 929 1512 1559 1576 1617 1764 1826 2118 2128 2473 2503 2522 2621 2639 2906 3049 3090 3334 3489 3663 3789 3840 3954 4076 4126 4180 4279 4280 4365
N position in protein
microbiome
community
germfree
monocolonized

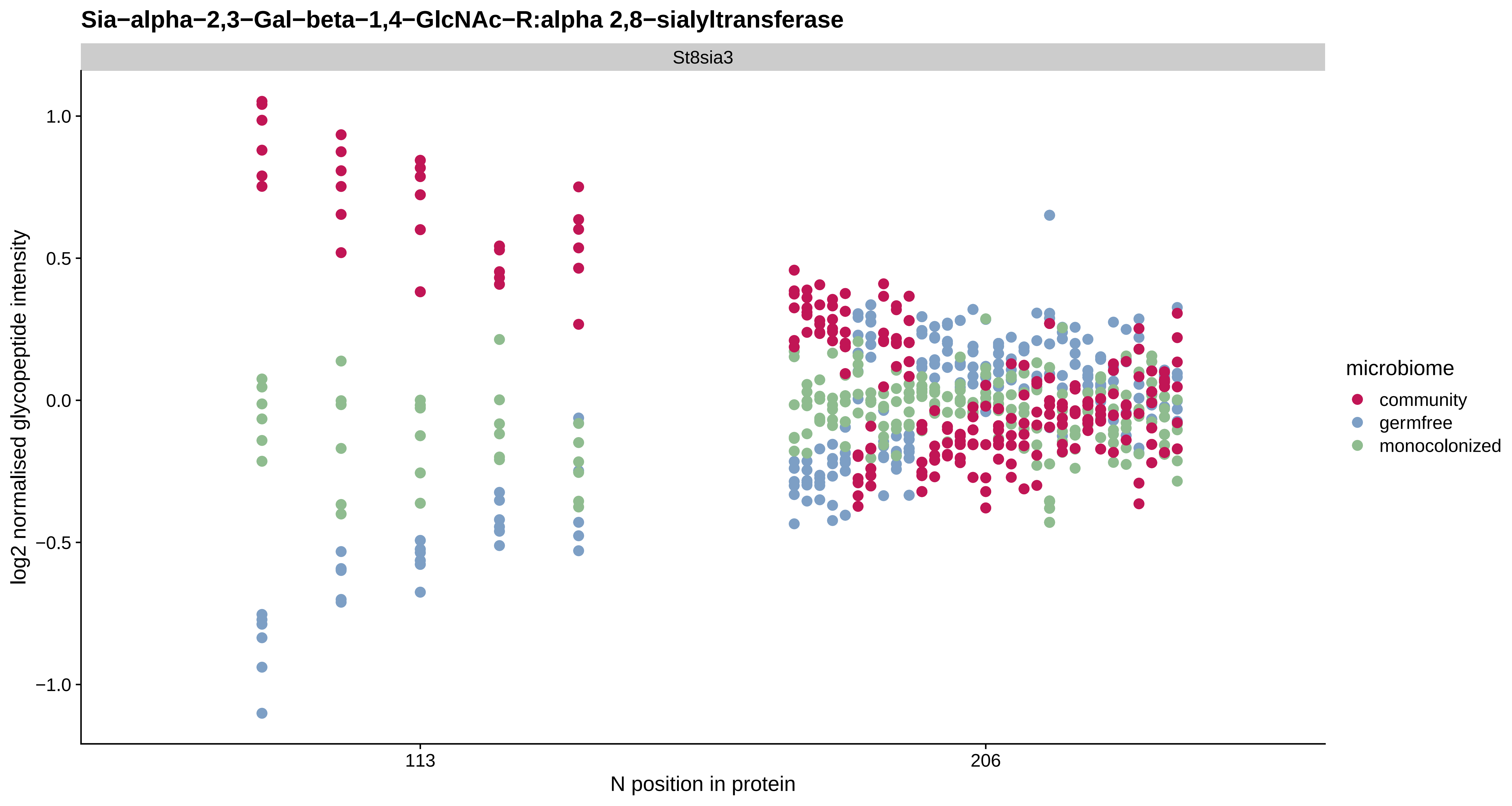

Sia−alpha−2,3−Gal−beta−1,4−GlcNAc−R:alpha 2,8−sialyltransferase
St8sia3
log2 normalised glycopeptide intensity
1.0
0.5
0.0
−0.5
−1.0
113
206
N position in protein
microbiome
community
germfree
monocolonized

#### Ankyrin repeat domain−containing protein 17

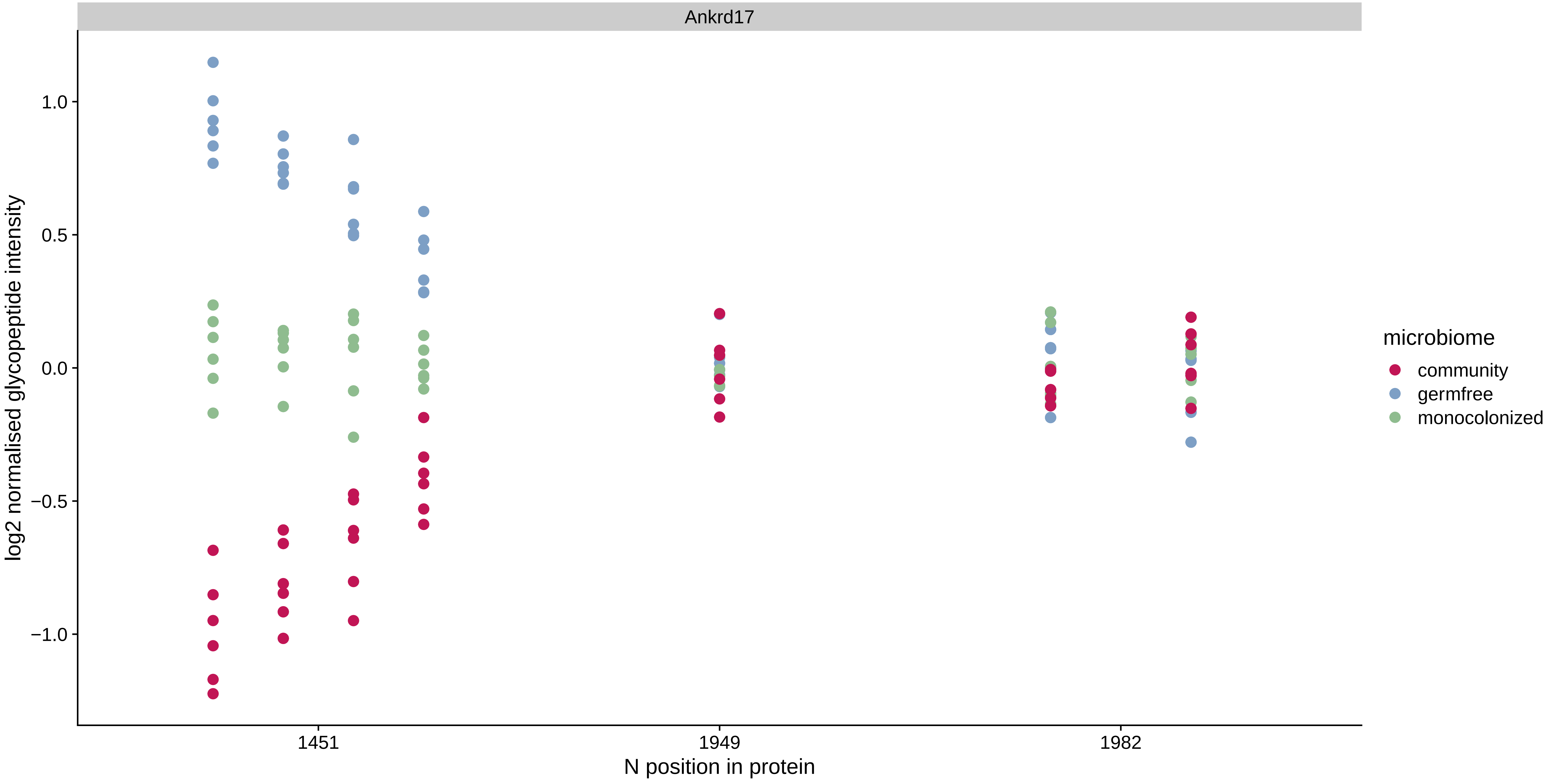

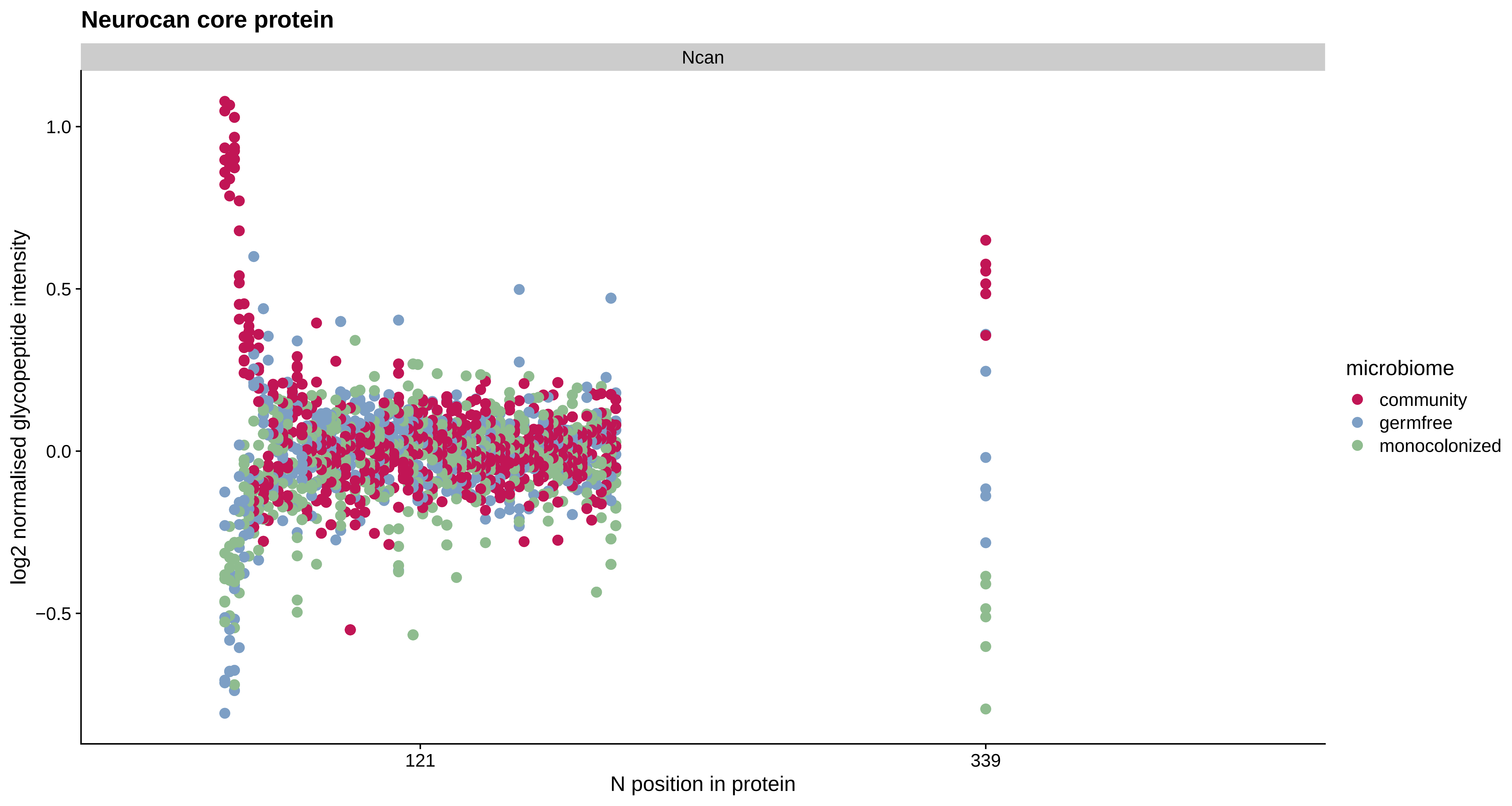
Neurocan core protein
Ncan
1.0
0.5
0.0
−0.5
log2 normalised glycopeptide intensity
121
339
N position in protein
microbiome
community
germfree
monocolonized

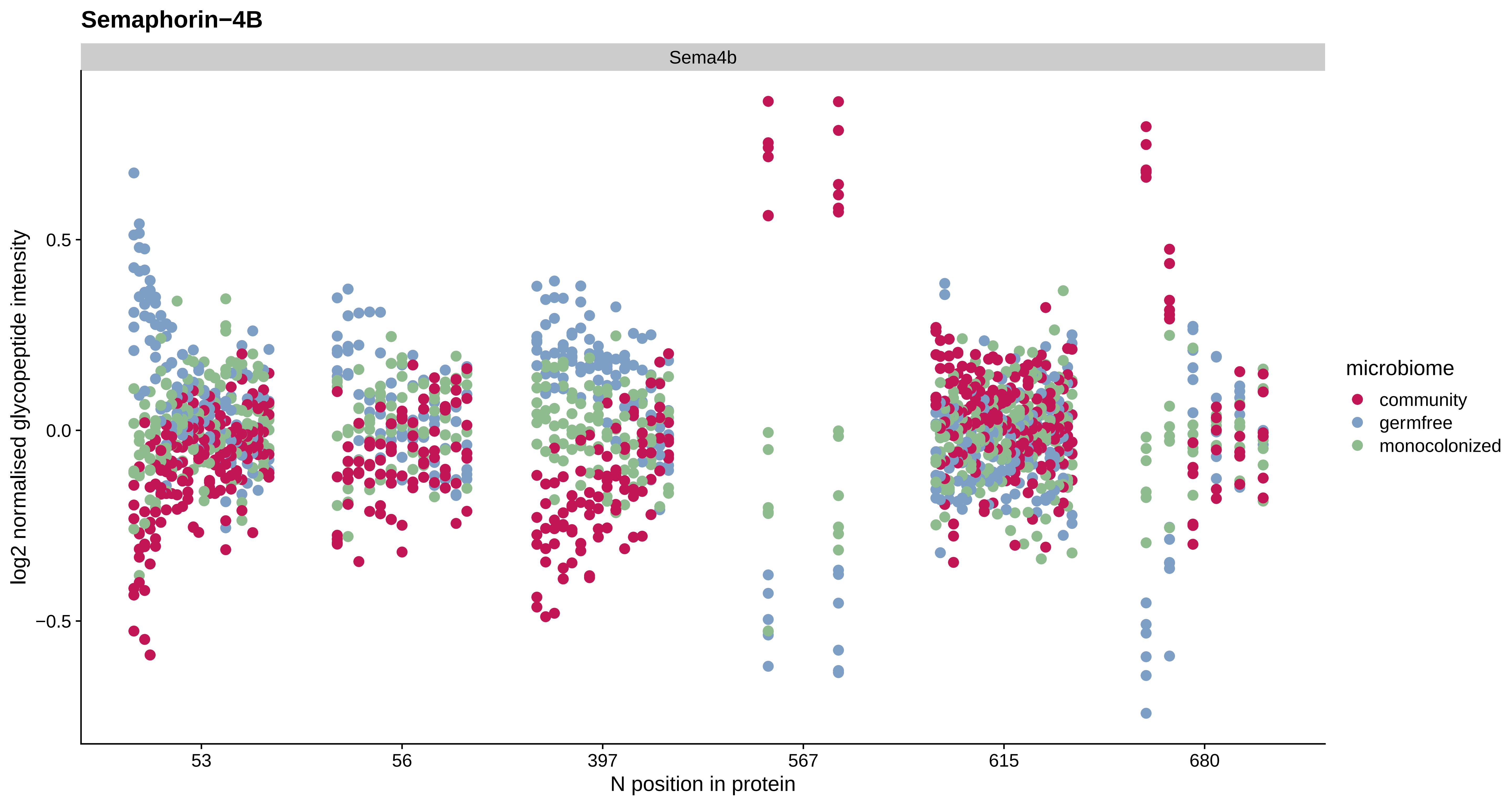
Semaphorin−4B
Sema4b
log2 normalised glycopeptide intensity
0.5
0.0
−0.5
53
56
397
567
615
680
N position in protein
microbiome
community
germfree
monocolonized

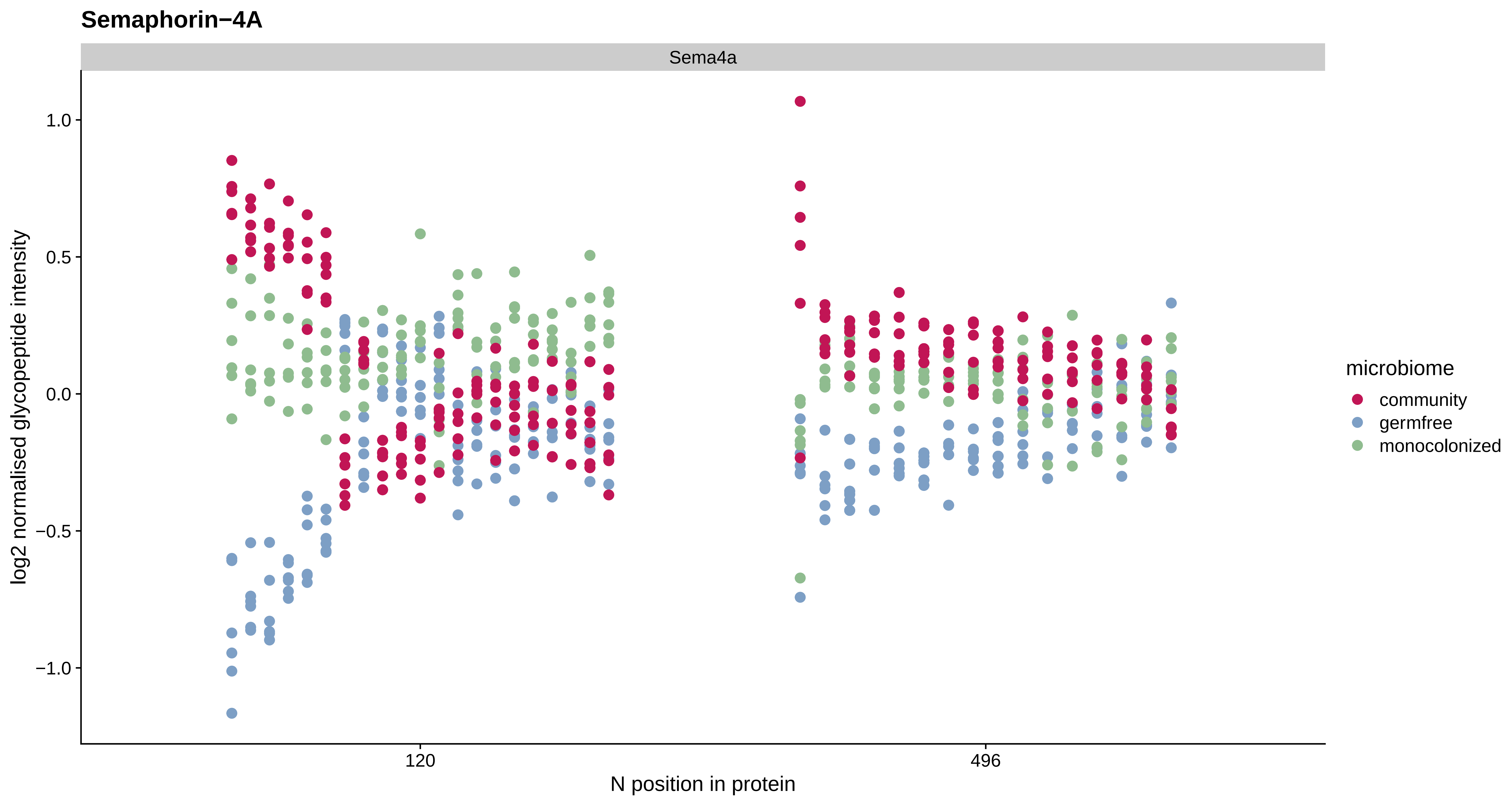

Semaphorin−4A
Sema4a
log2 normalised glycopeptide intensity
1.0
0.5
0.0
−0.5
−1.0
120
496
N position in protein
microbiome
community
germfree
monocolonized

#### Peroxidasin homolog

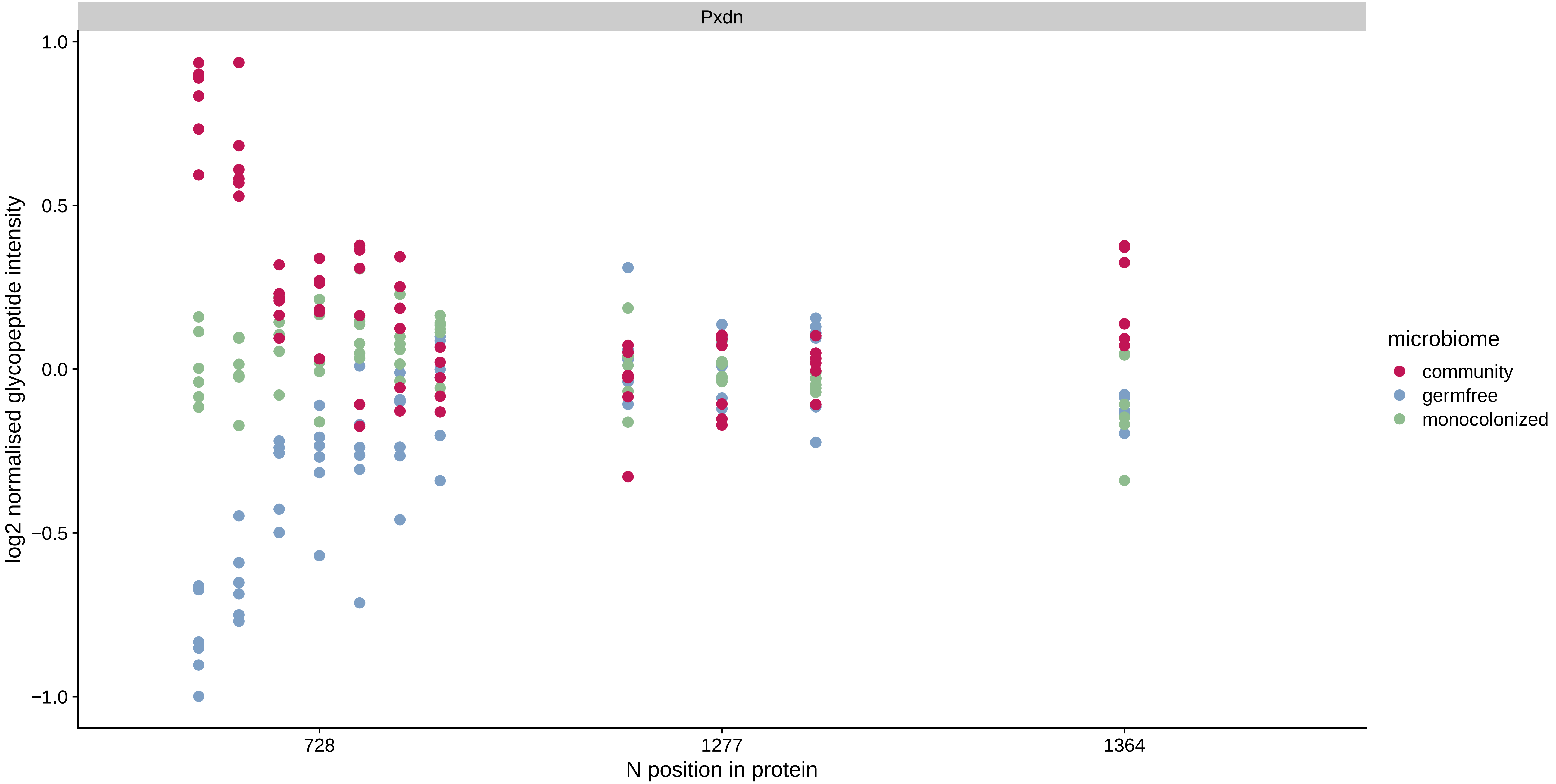

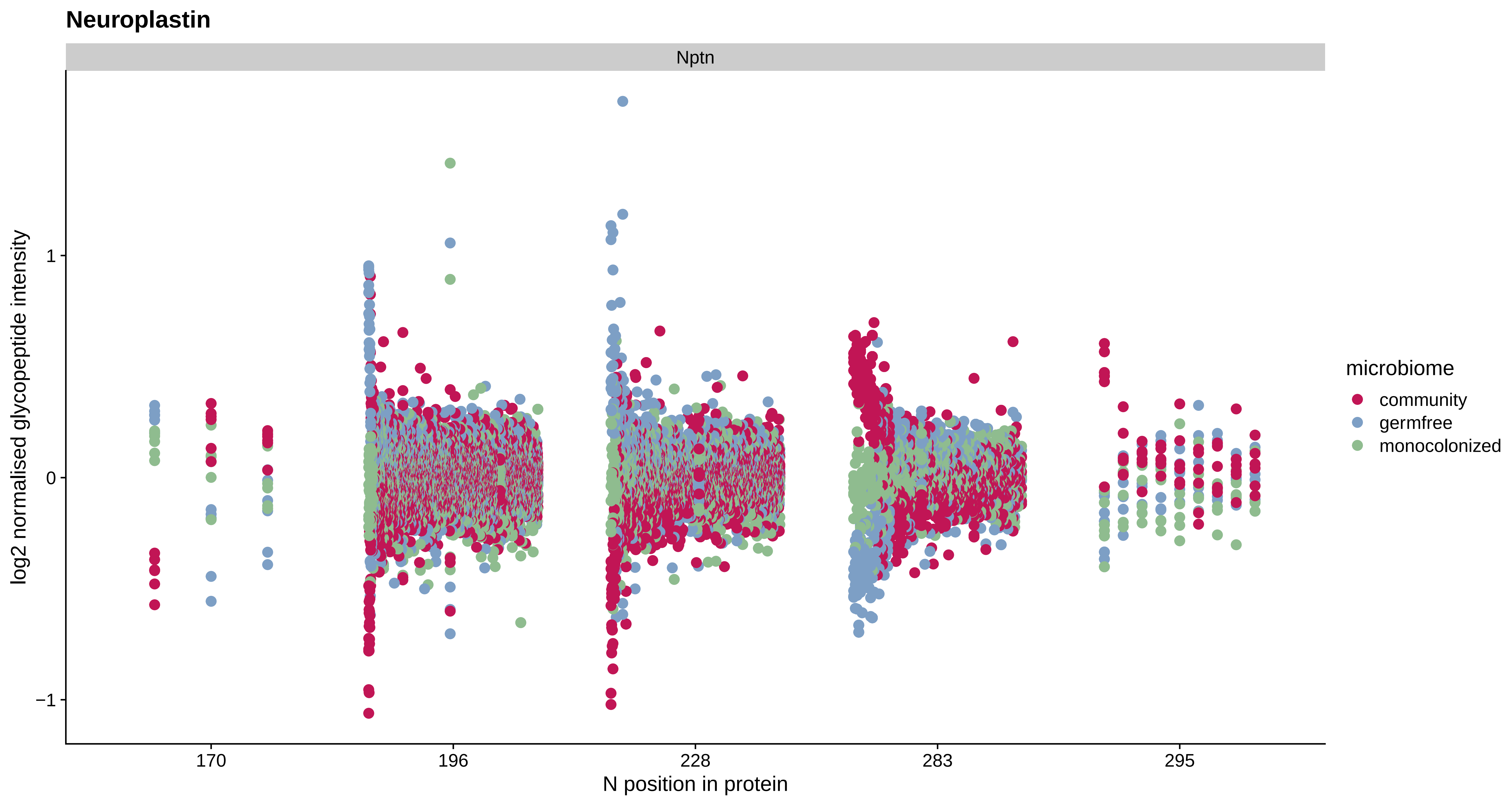

Neuroplastin
Nptn
log2 normalised glycopeptide intensity
1
0
−1
170
196
228
283
295
N position in protein
microbiome
community
germfree
monocolonized

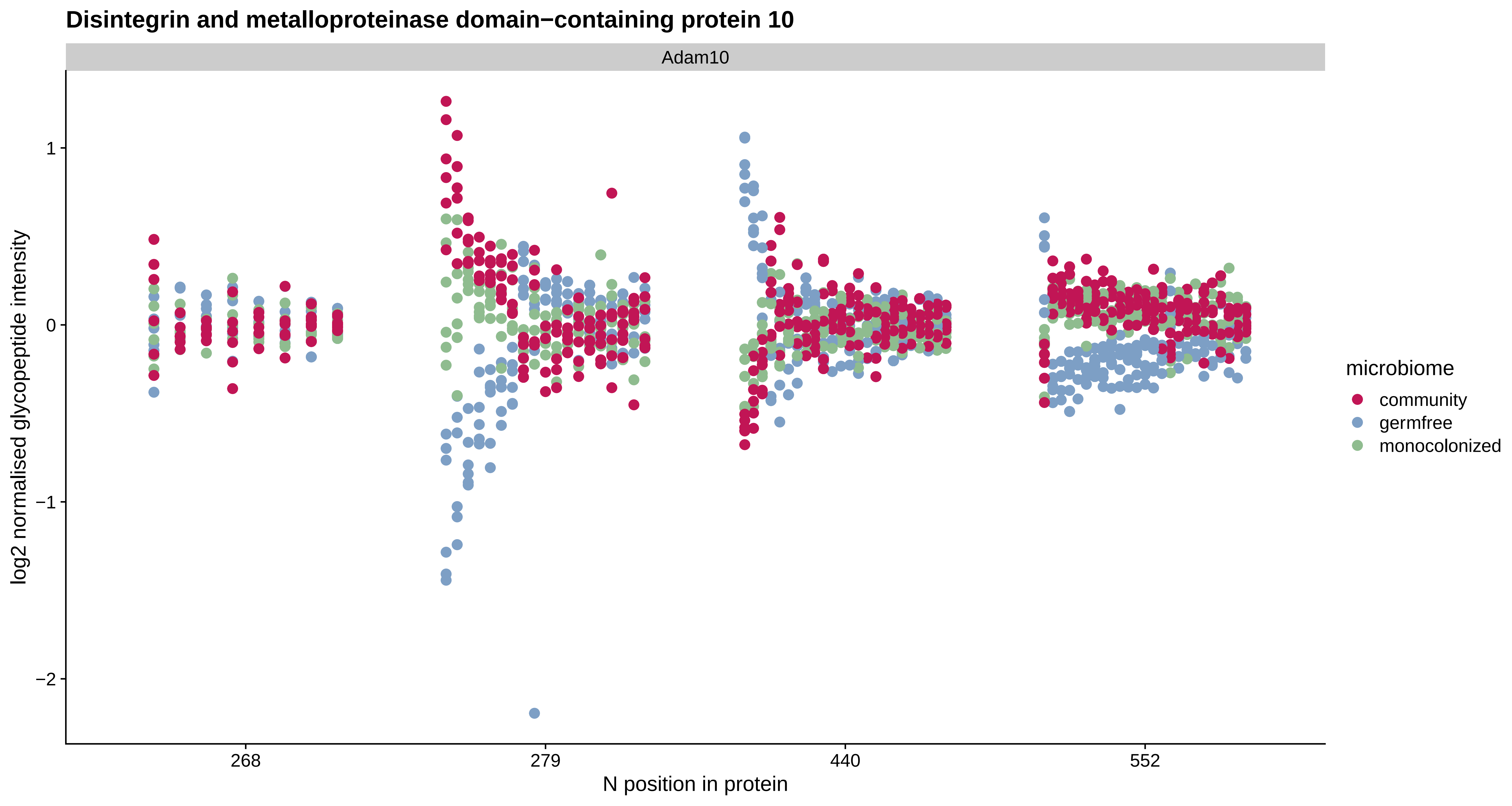
Disintegrin and metalloproteinase domain−containing protein 10
Adam10
log2 normalised glycopeptide intensity
1
0
−1
−2
268
279
440
552
N position in protein
microbiome
community
germfree
monocolonized

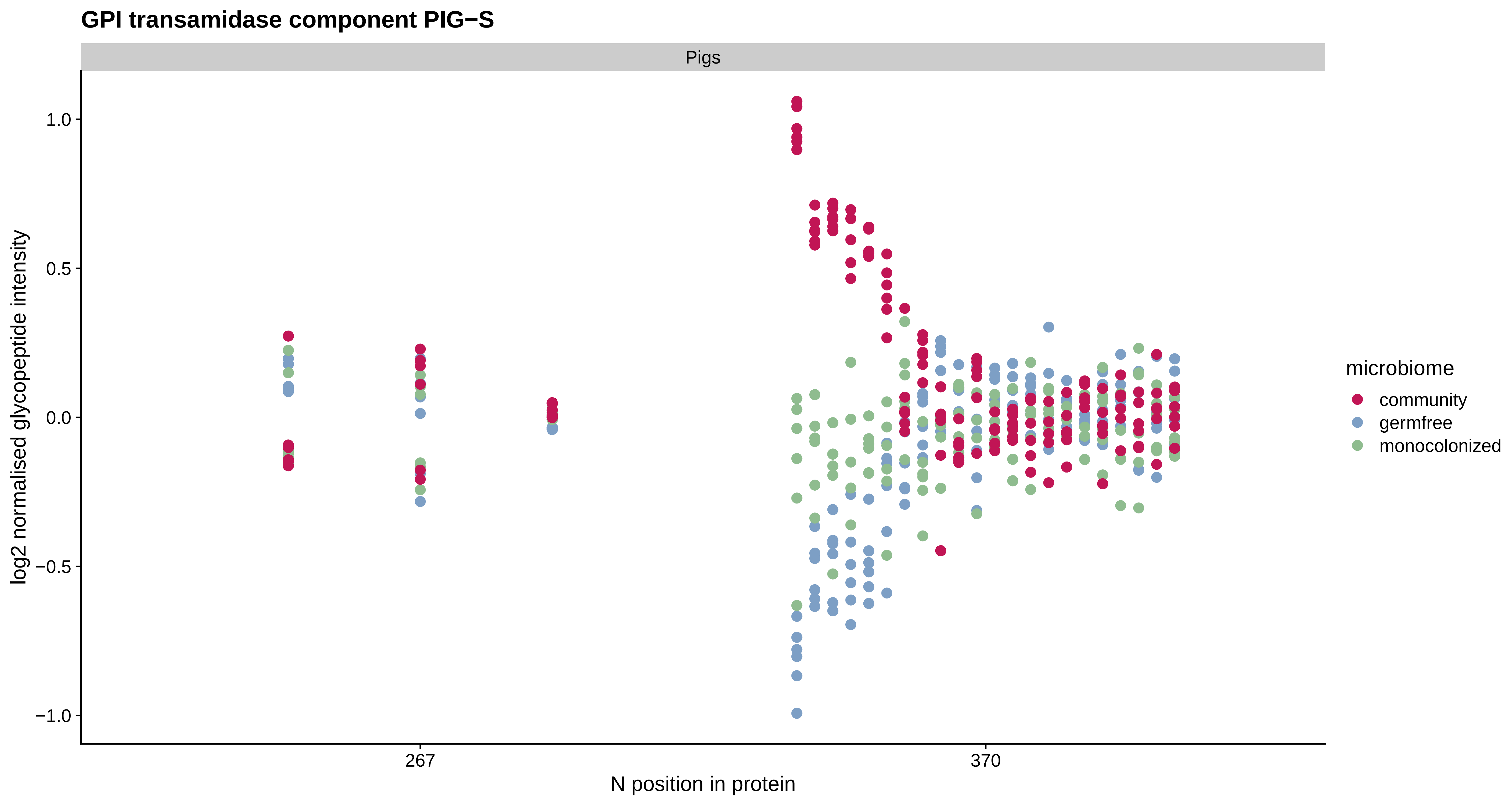

GPI transamidase component PIG−S
Pigs
1.0
0.5
0.0
−0.5
−1.0
log2 normalised glycopeptide intensity
267
370
N position in protein
microbiome
community
germfree
monocolonized

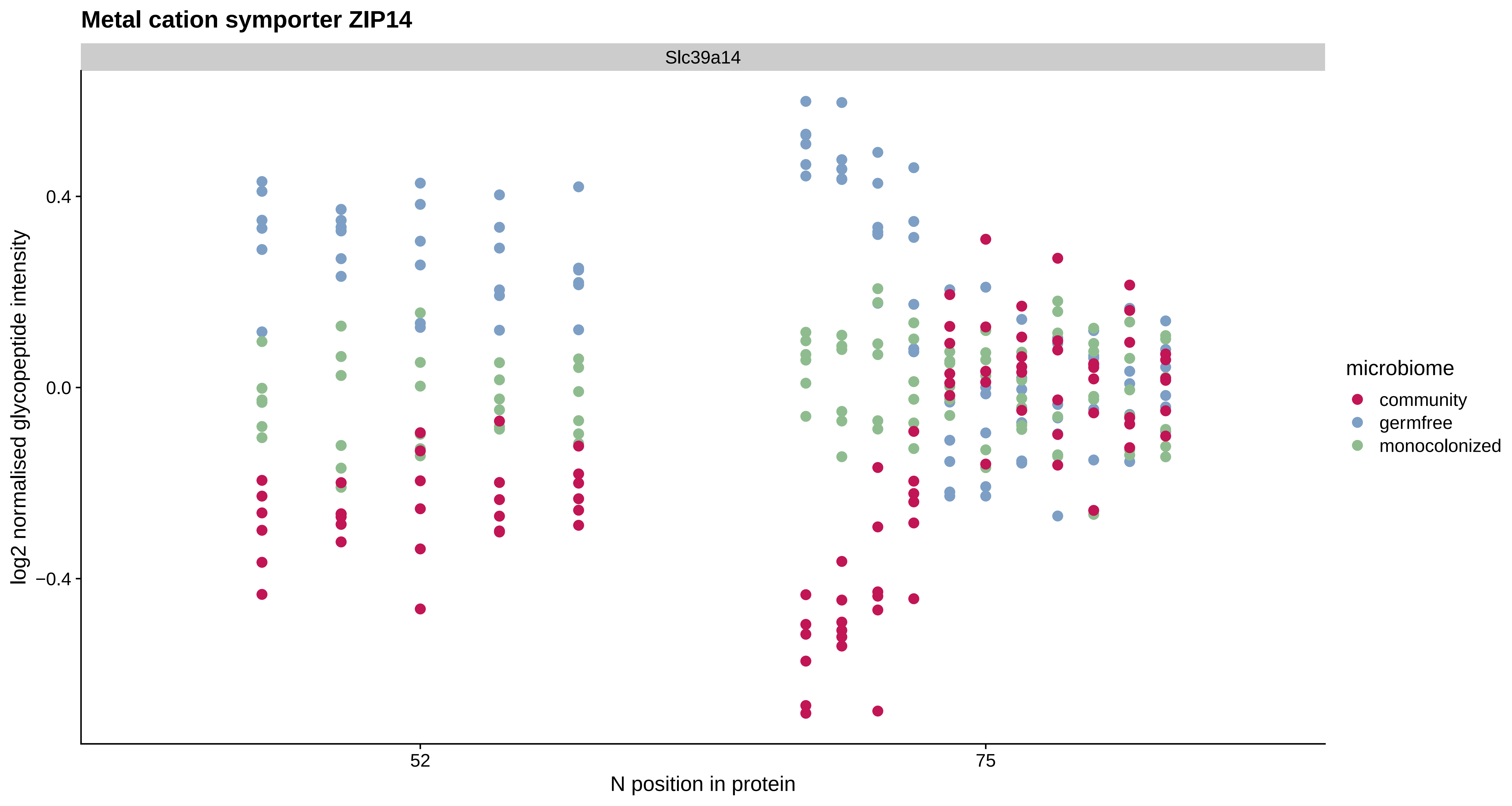

Metal cation symporter ZIP14
Slc39a14
log2 normalised glycopeptide intensity
0.4
0.0
−0.4
52
75
N position in protein
microbiome
community
germfree
monocolonized

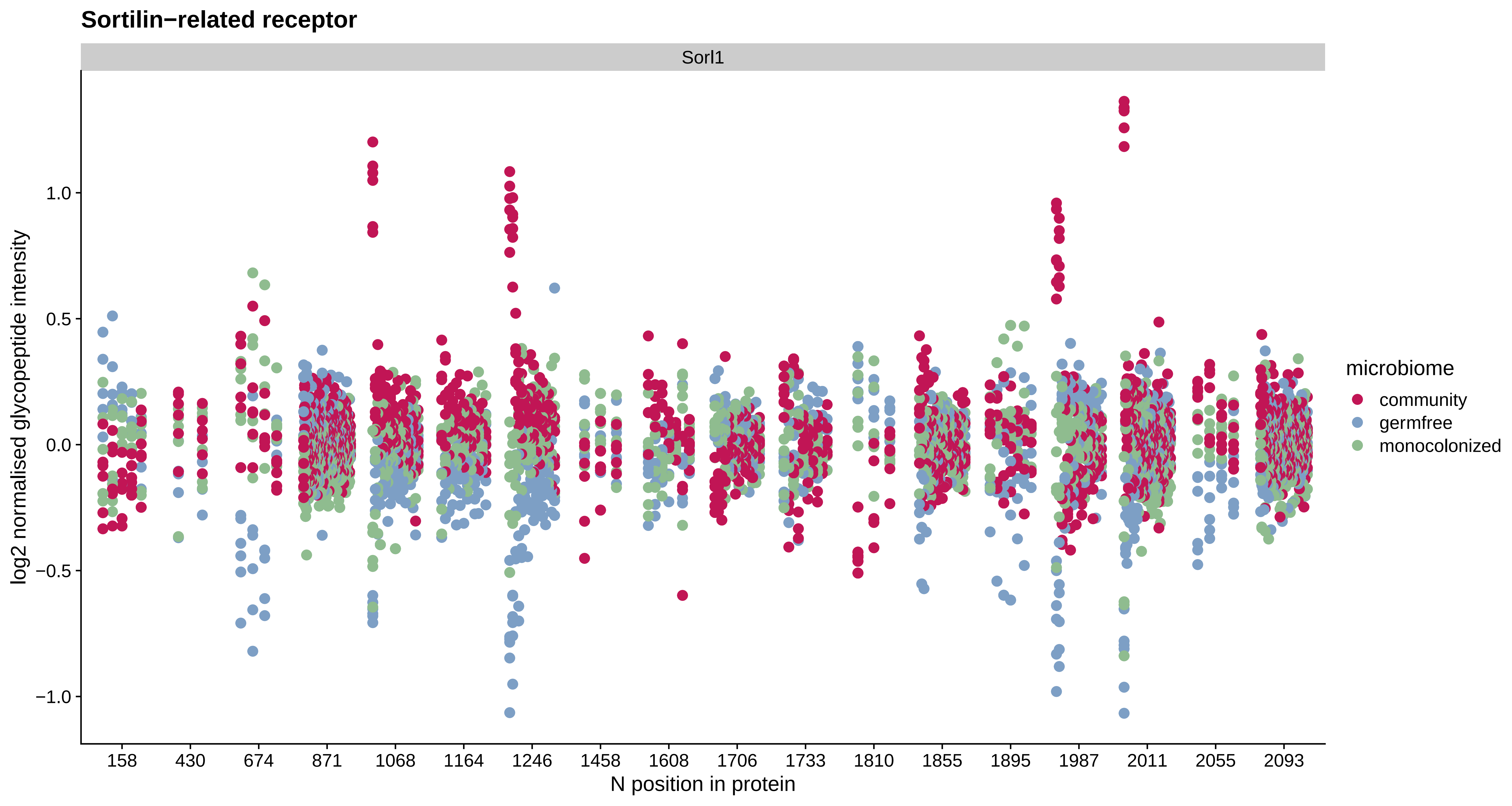

Sortilin−related receptor
Sorl1
log2 normalised glycopeptide intensity
1.0
0.5
0.0
−0.5
−1.0
158
430
674
871
1068
1164
1246
1458
1608
1706
1733
1810
1855
1895
1987
2011
2055
2093
N position in protein
microbiome
community
germfree
monocolonized

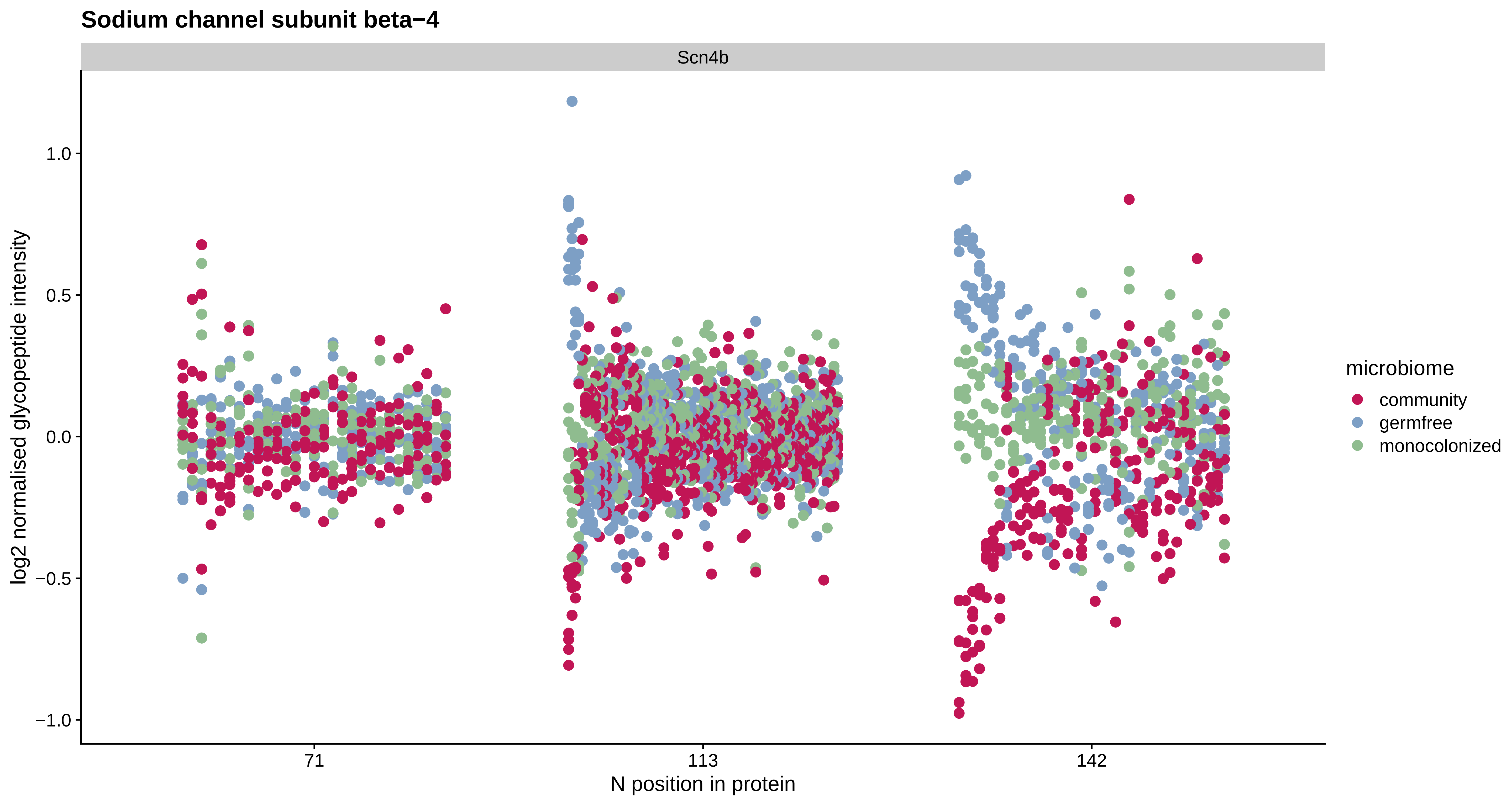

Sodium channel subunit beta−4
Scn4b
1.0
0.5
0.0
−0.5
−1.0
log2 normalised glycopeptide intensity
71
113
142
N position in protein
microbiome
community
germfree
monocolonized

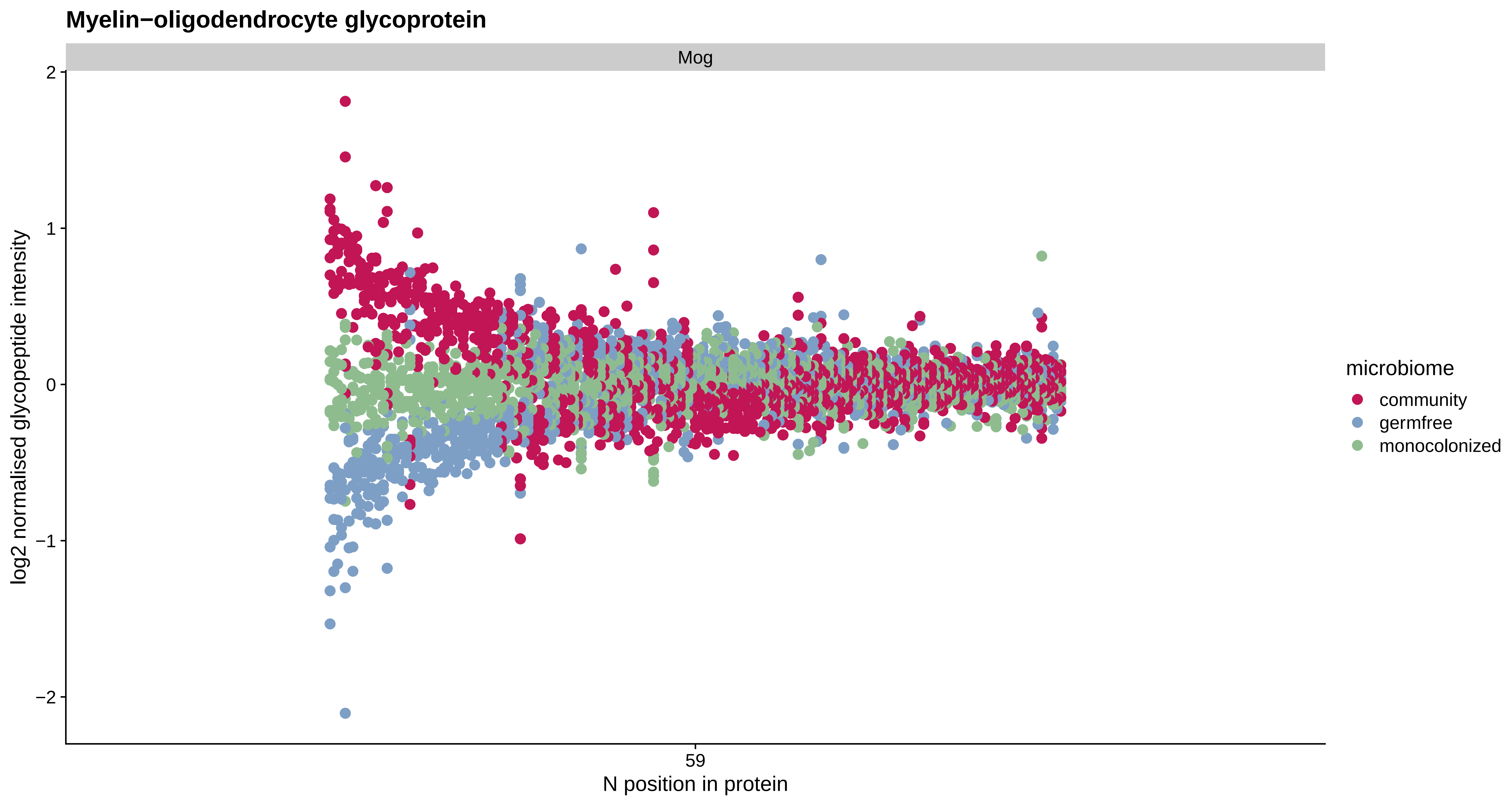
Myelin−oligodendrocyte glycoprotein
Mog
log2 normalised glycopeptide intensity
2
1
0
−1
−2
59
N position in protein
microbiome
community
germfree
monocolonized

#### Glutamate receptor 3

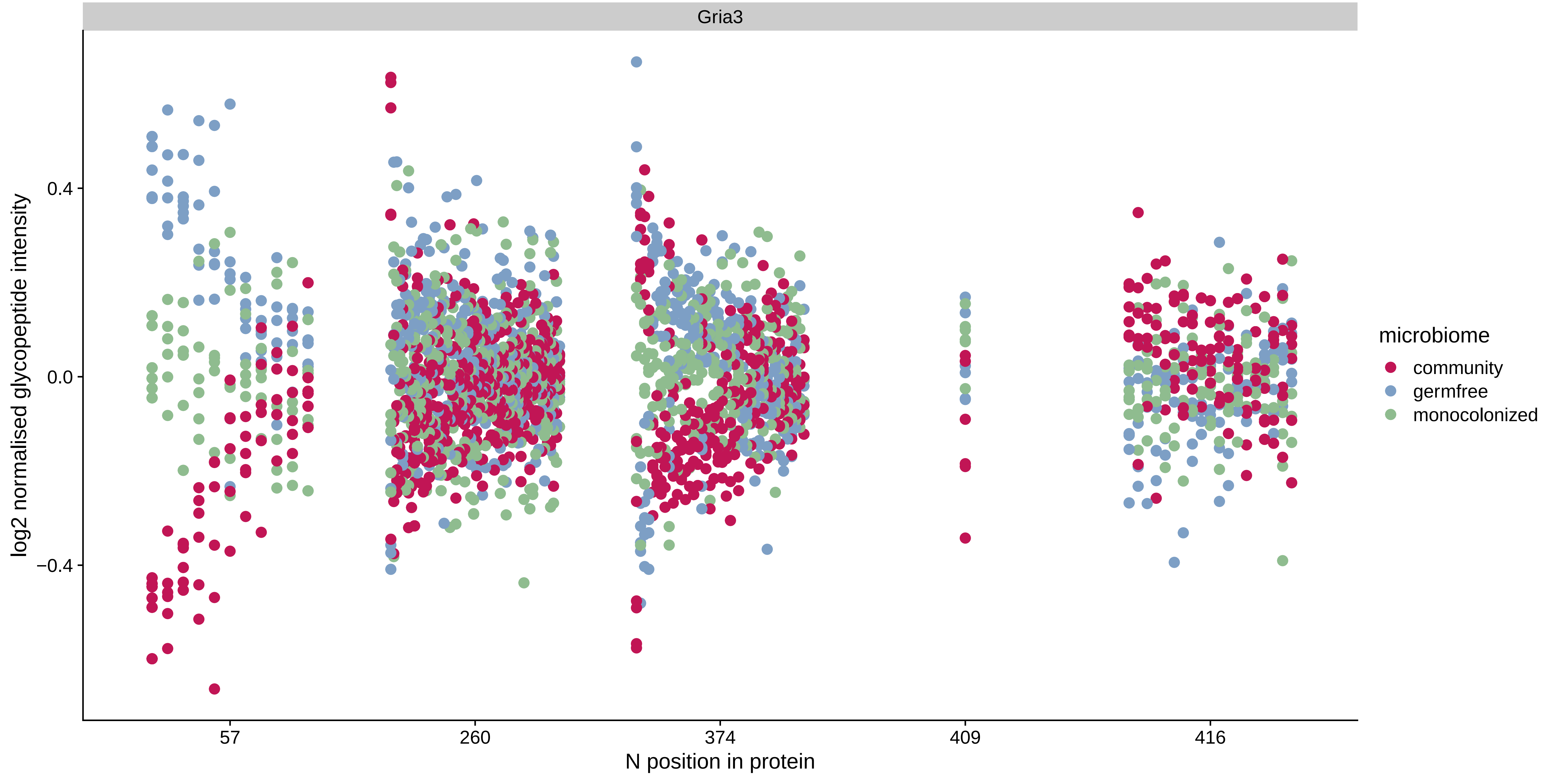

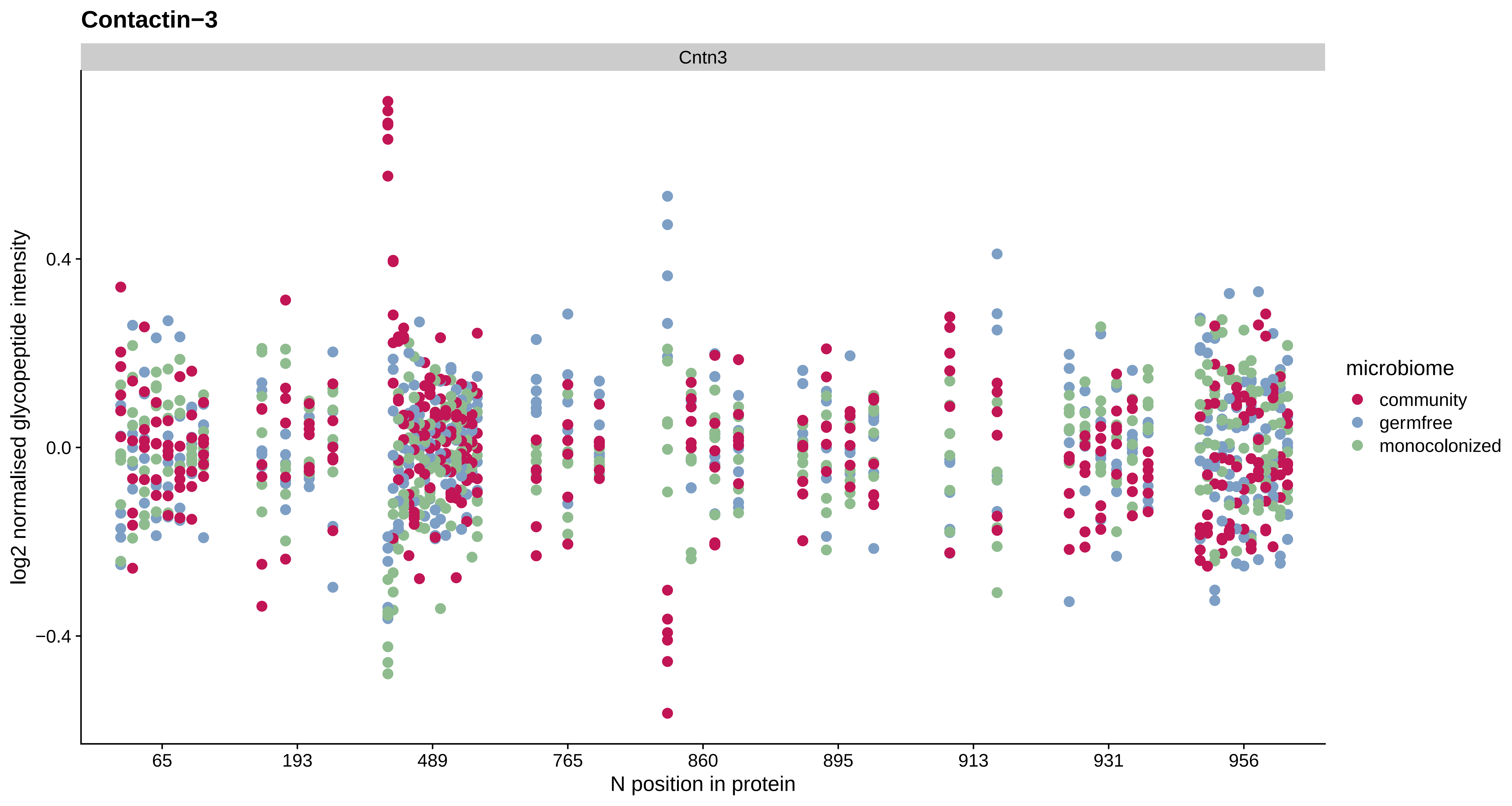
Contactin−3
Cntn3
log2 normalised glycopeptide intensity
0.4
0.0
−0.4
65
193
489
765
860
895
913
931
956
N position in protein
microbiome
community
germfree
monocolonized

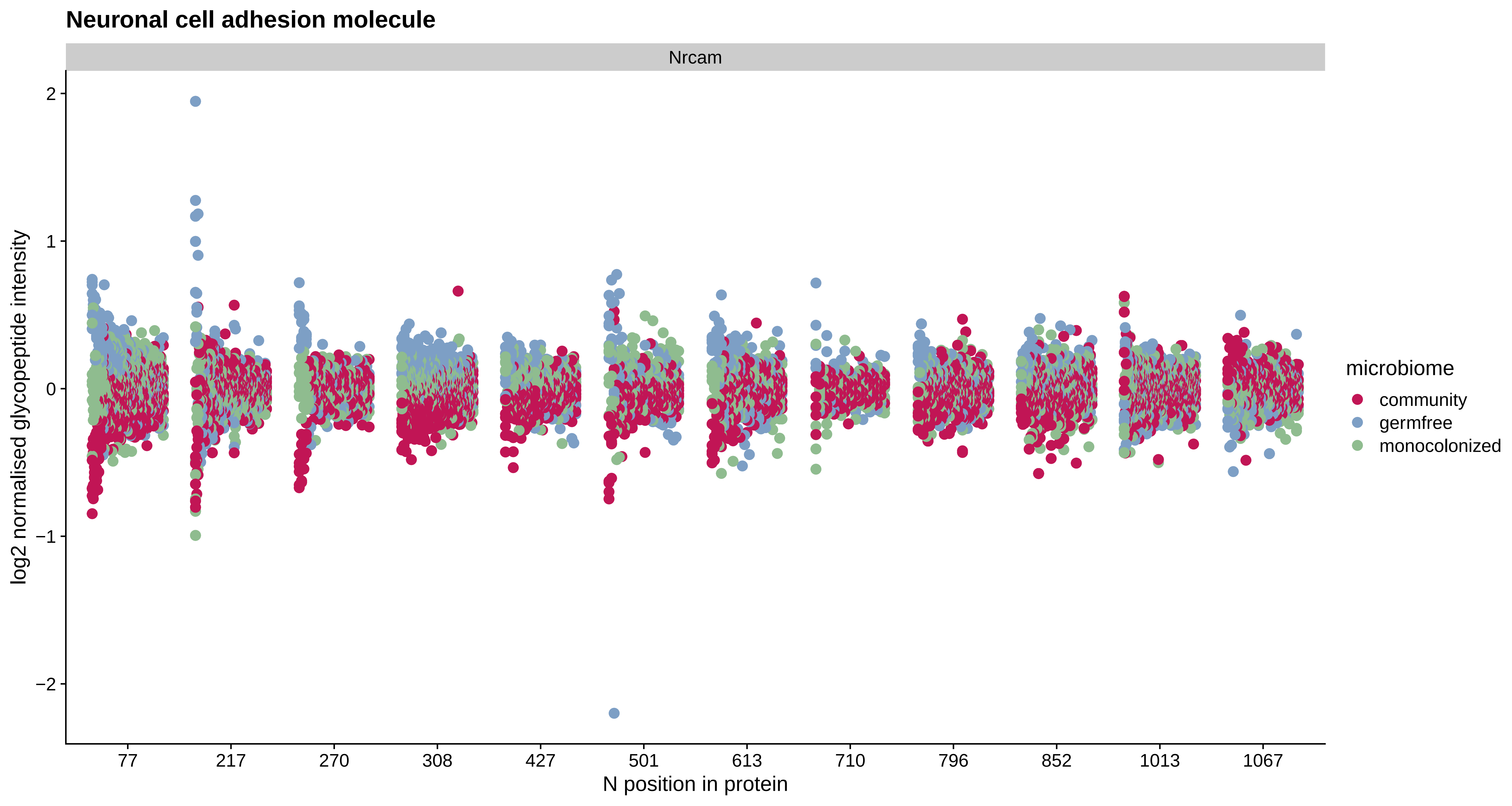
Neuronal cell adhesion molecule
Nrcam
log2 normalised glycopeptide intensity
2
1
0
−1
−2
77
217
270
308
427
501
613
710
796
852
1013
1067
N position in protein
microbiome
community
germfree
monocolonized

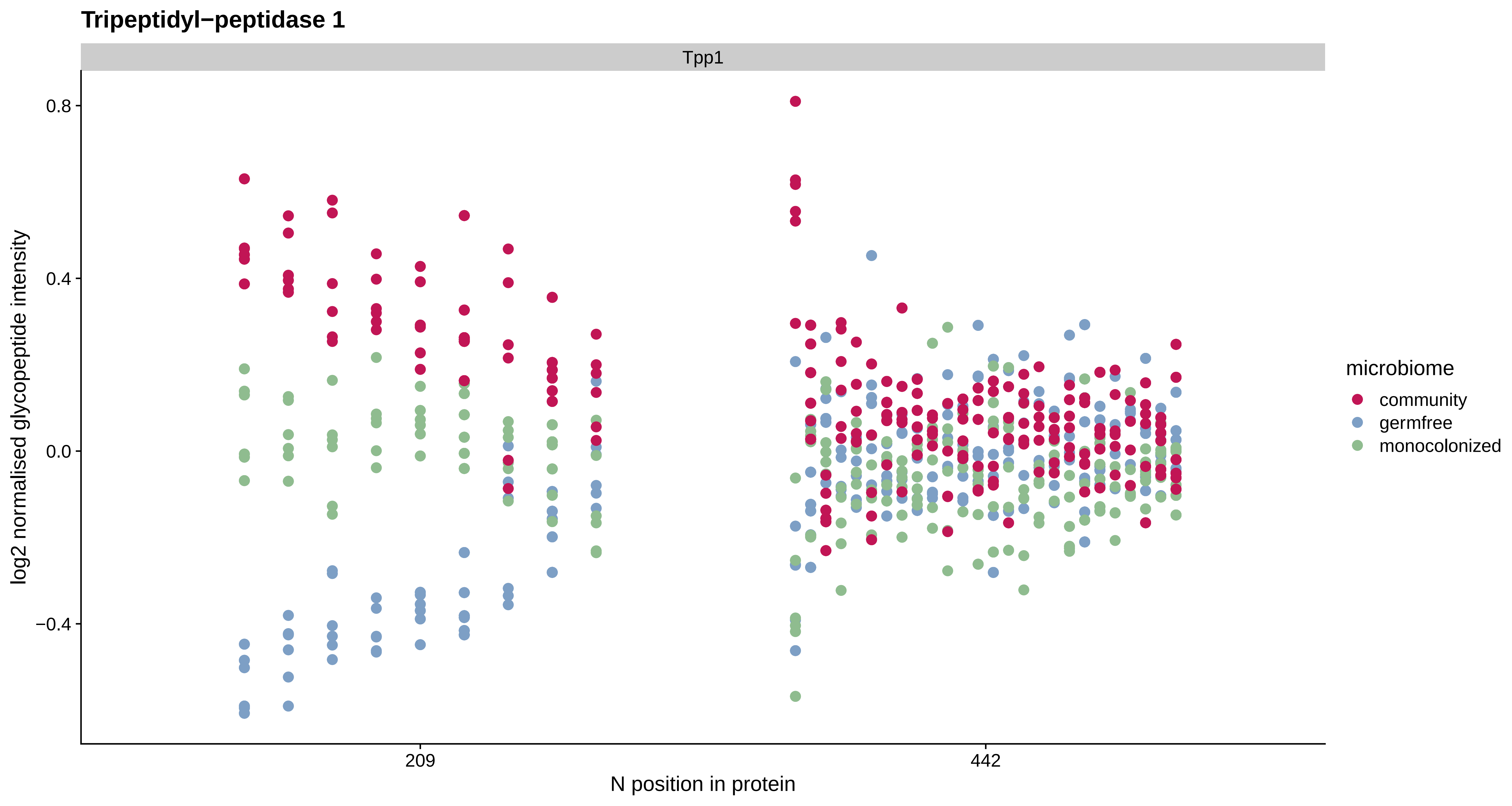
Tripeptidyl−peptidase 1
Tpp1
0.8
0.4
0.0
−0.4
log2 normalised glycopeptide intensity
209
442
N position in protein
microbiome
community
germfree
monocolonized

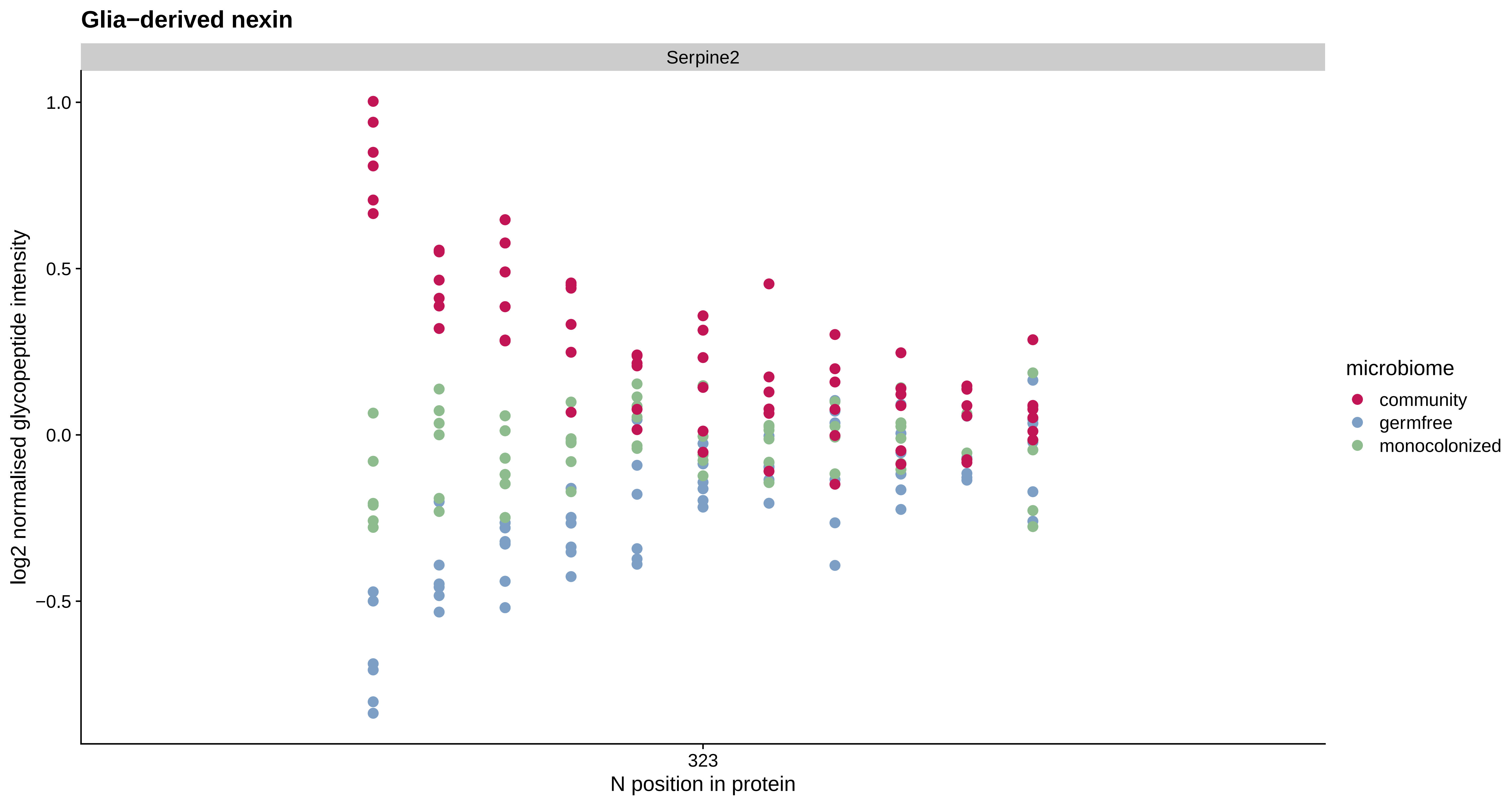

Glia−derived nexin
Serpine2
1.0
0.5
0.0
−0.5
log2 normalised glycopeptide intensity
323
N position in protein
microbiome
community
germfree
monocolonized

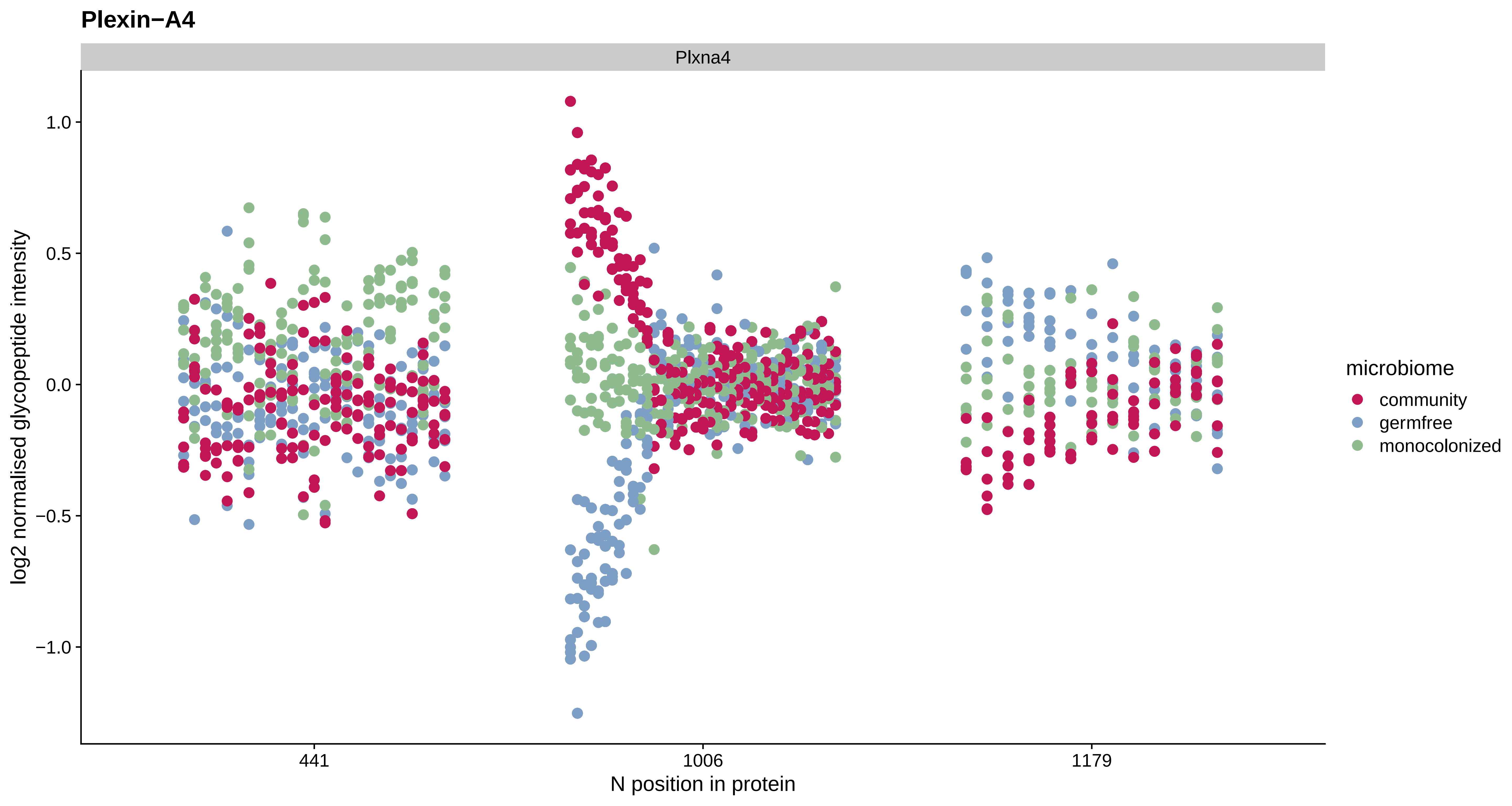
Plexin−A4
Plxna4
log2 normalised glycopeptide intensity
1.0
0.5
0.0
−0.5
−1.0
441
1006
1179
N position in protein
microbiome
community
germfree
monocolonized

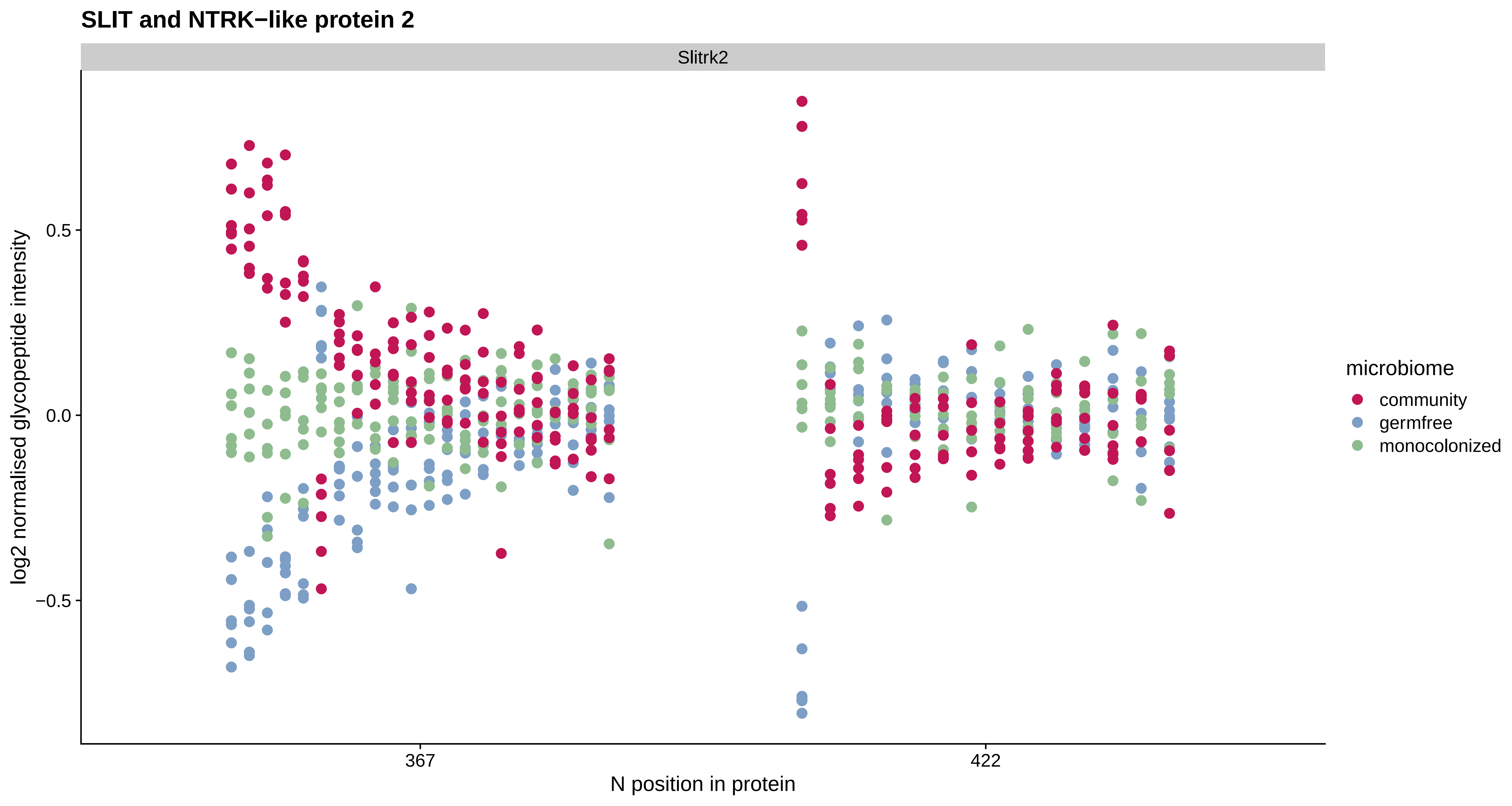
SLIT and NTRK−like protein 2
Slitrk2
log2 normalised glycopeptide intensity
0.5
0.0
−0.5
367
422
N position in protein
microbiome
community
germfree
monocolonized

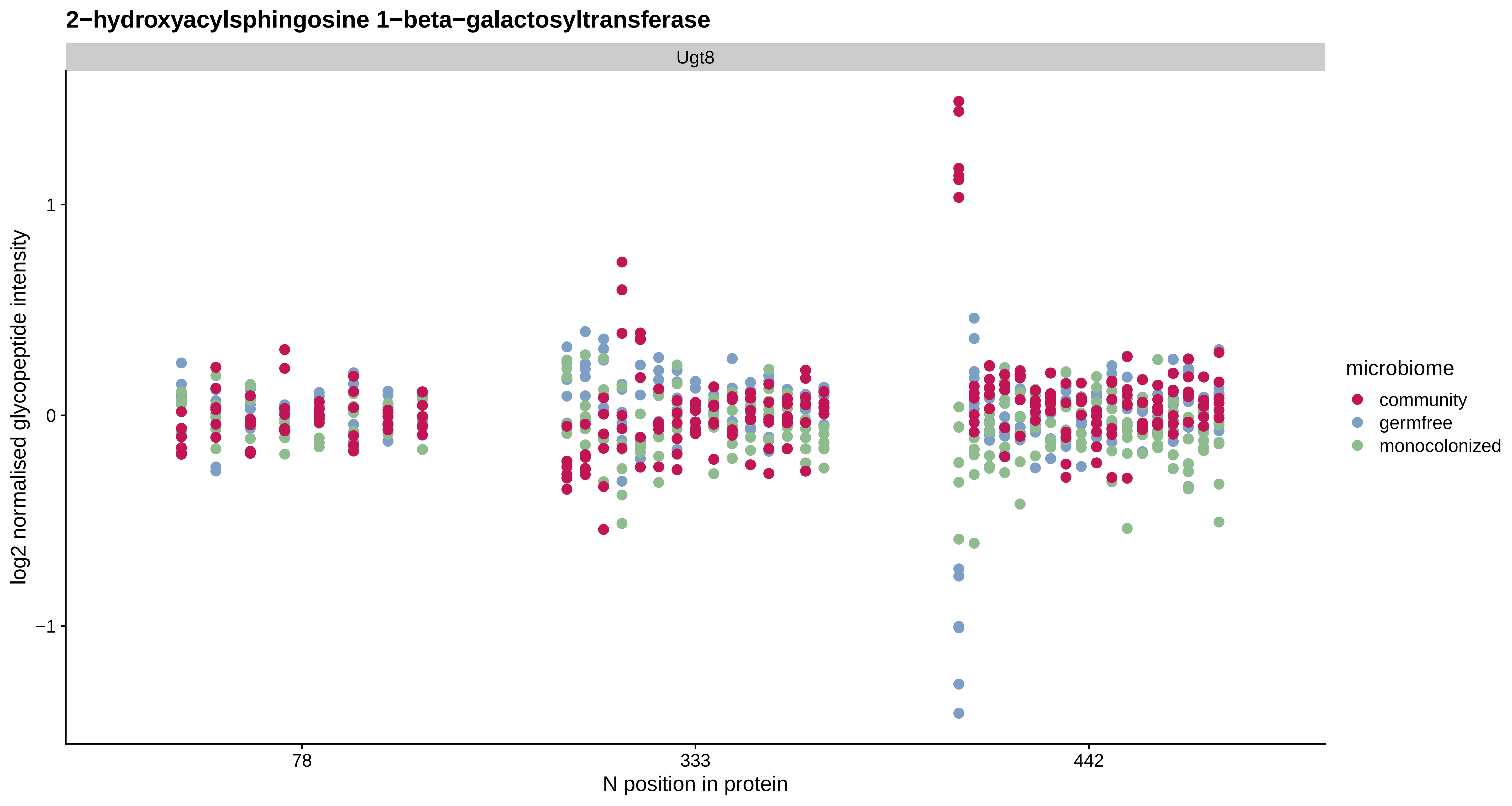
2−hydroxyacylsphingosine 1−beta−galactosyltransferase
Ugt8
log2 normalised glycopeptide intensity
1
0
−1
78
333
442
N position in protein
microbiome
community
germfree
monocolonized

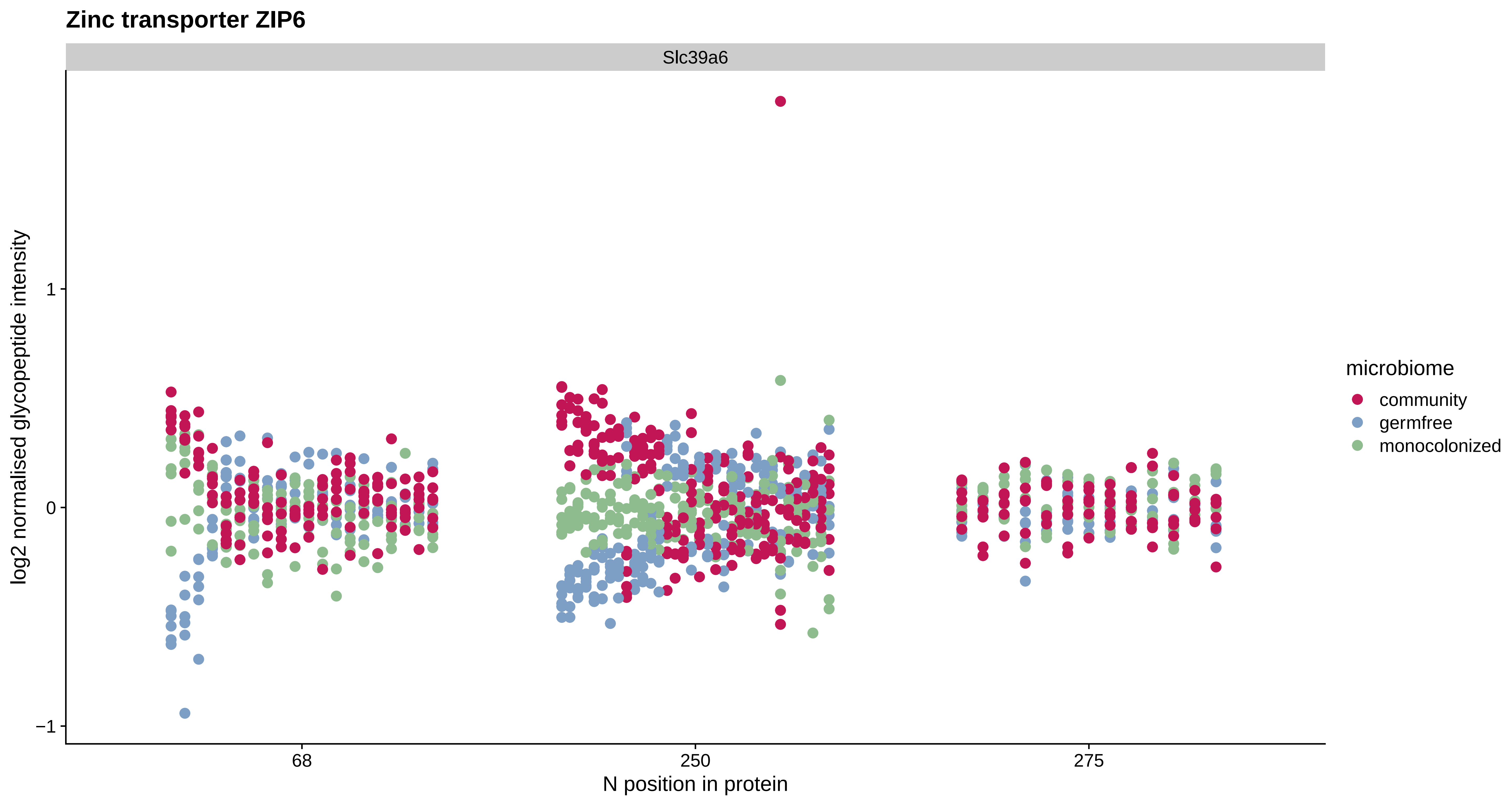

Zinc transporter ZIP6
Slc39a6
log2 normalised glycopeptide intensity
1
0
−1
68
250
275
N position in protein
microbiome
community
germfree
monocolonized

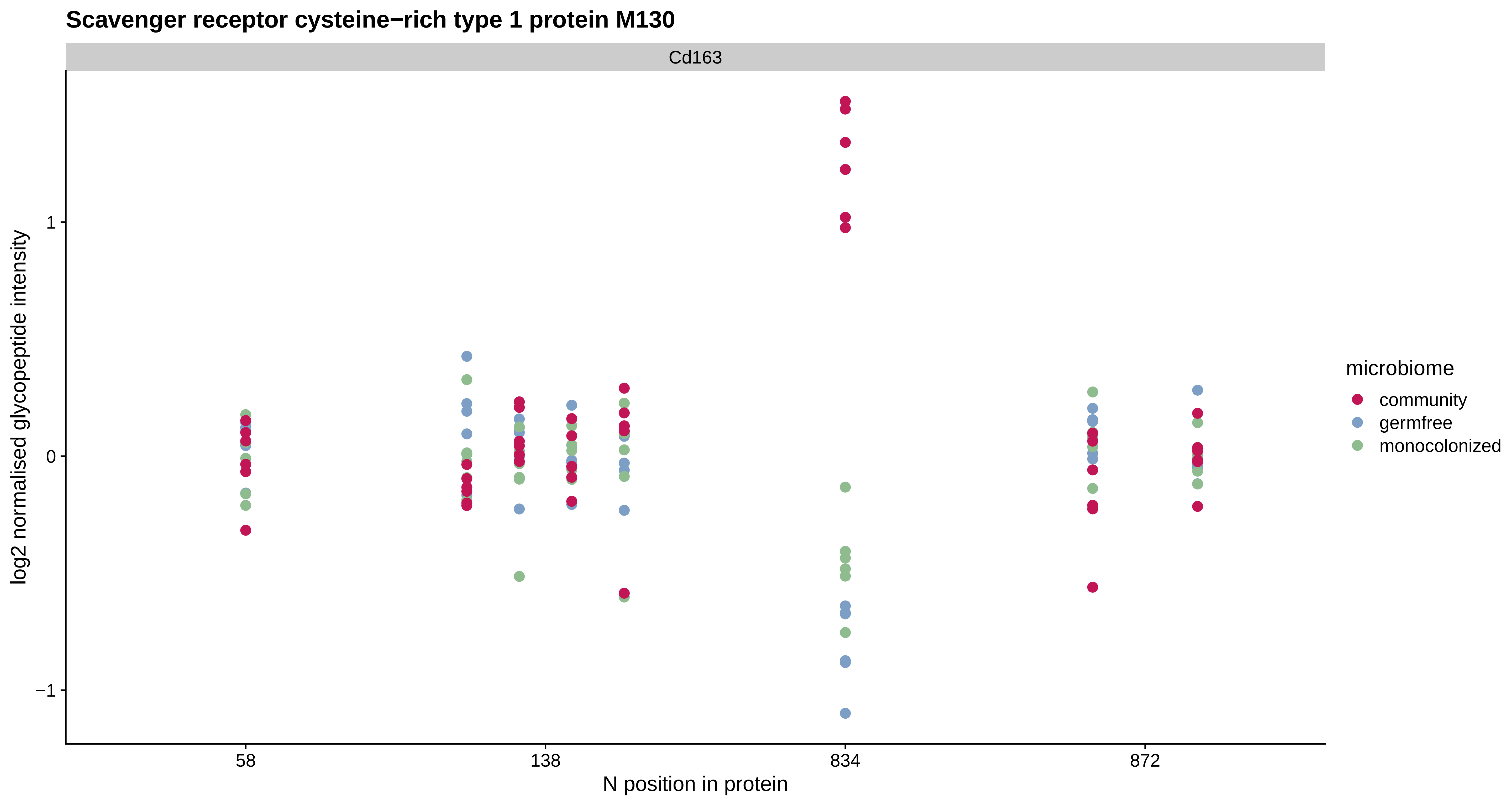
Scavenger receptor cysteine−rich type 1 protein M130
Cd163
log2 normalised glycopeptide intensity
1
0
−1
58
138
834
872
N position in protein
microbiome
community
germfree
monocolonized

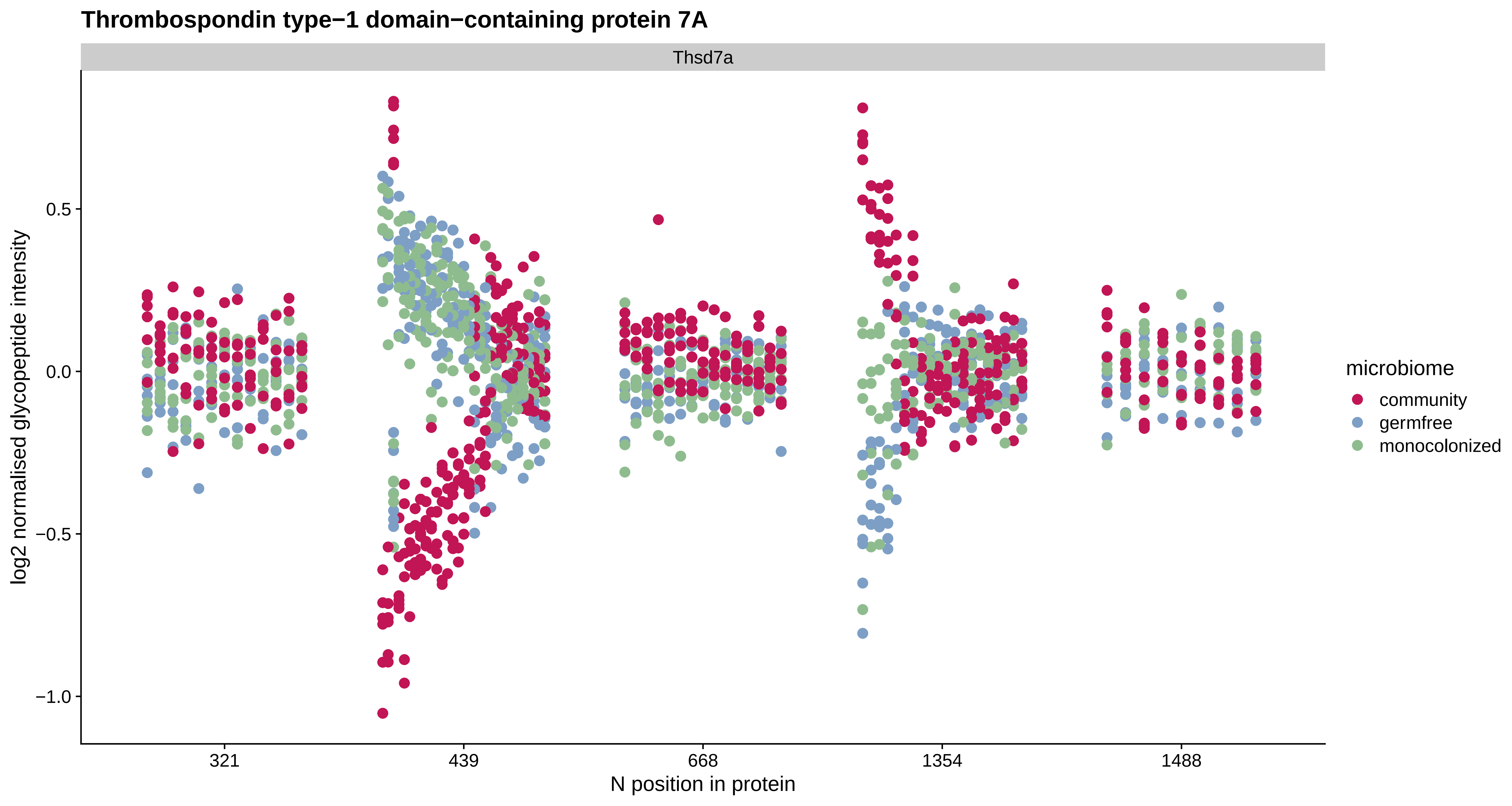

Thrombospondin type−1 domain−containing protein 7A
Thsd7a
log2 normalised glycopeptide intensity
0.5
0.0
−0.5
−1.0
321
439
668
1354
1488
N position in protein
microbiome
community
germfree
monocolonized

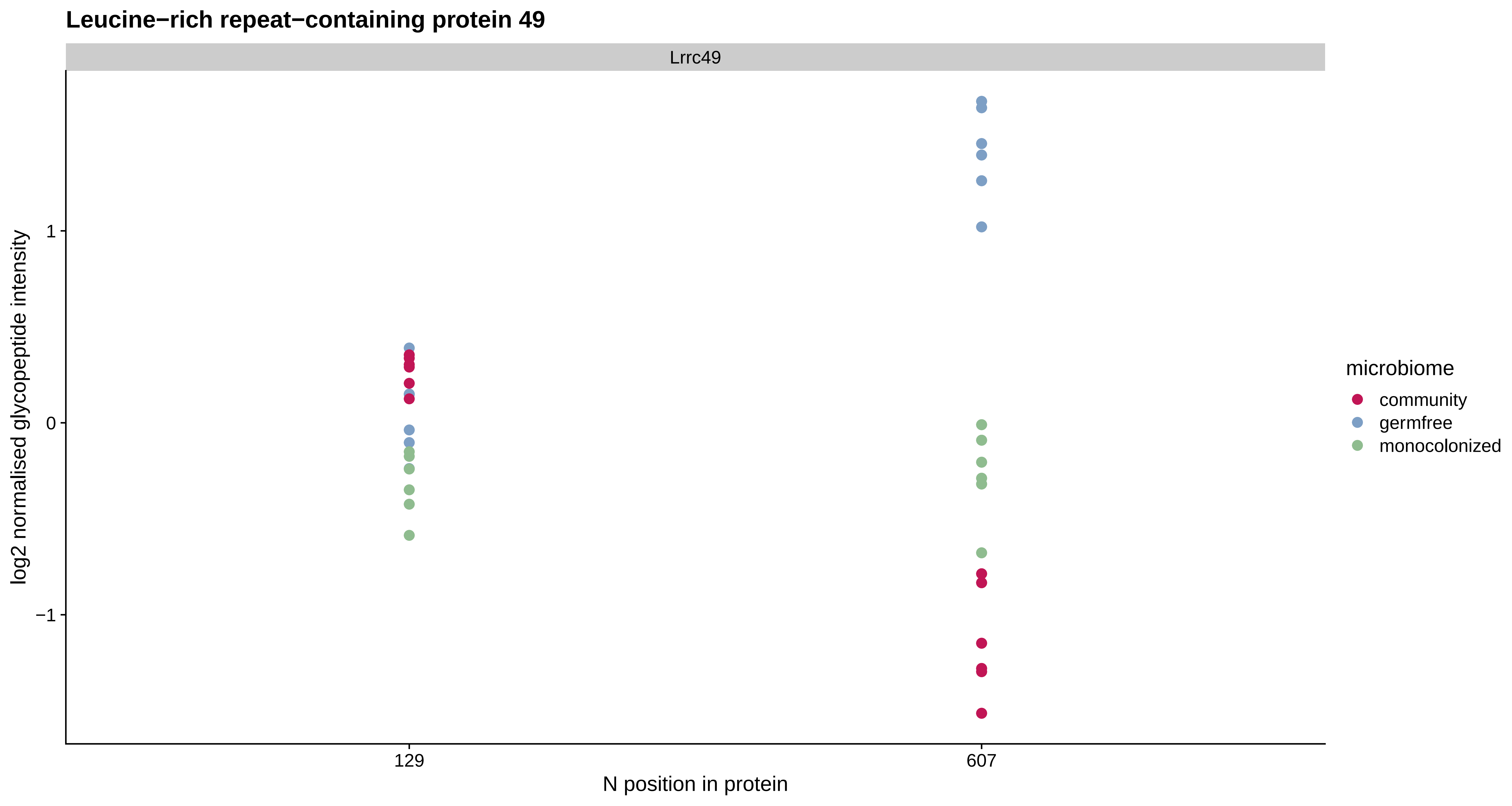

Leucine−rich repeat−containing protein 49
Lrrc49
log2 normalised glycopeptide intensity
1
0
−1
129
607
N position in protein
microbiome
community
germfree
monocolonized

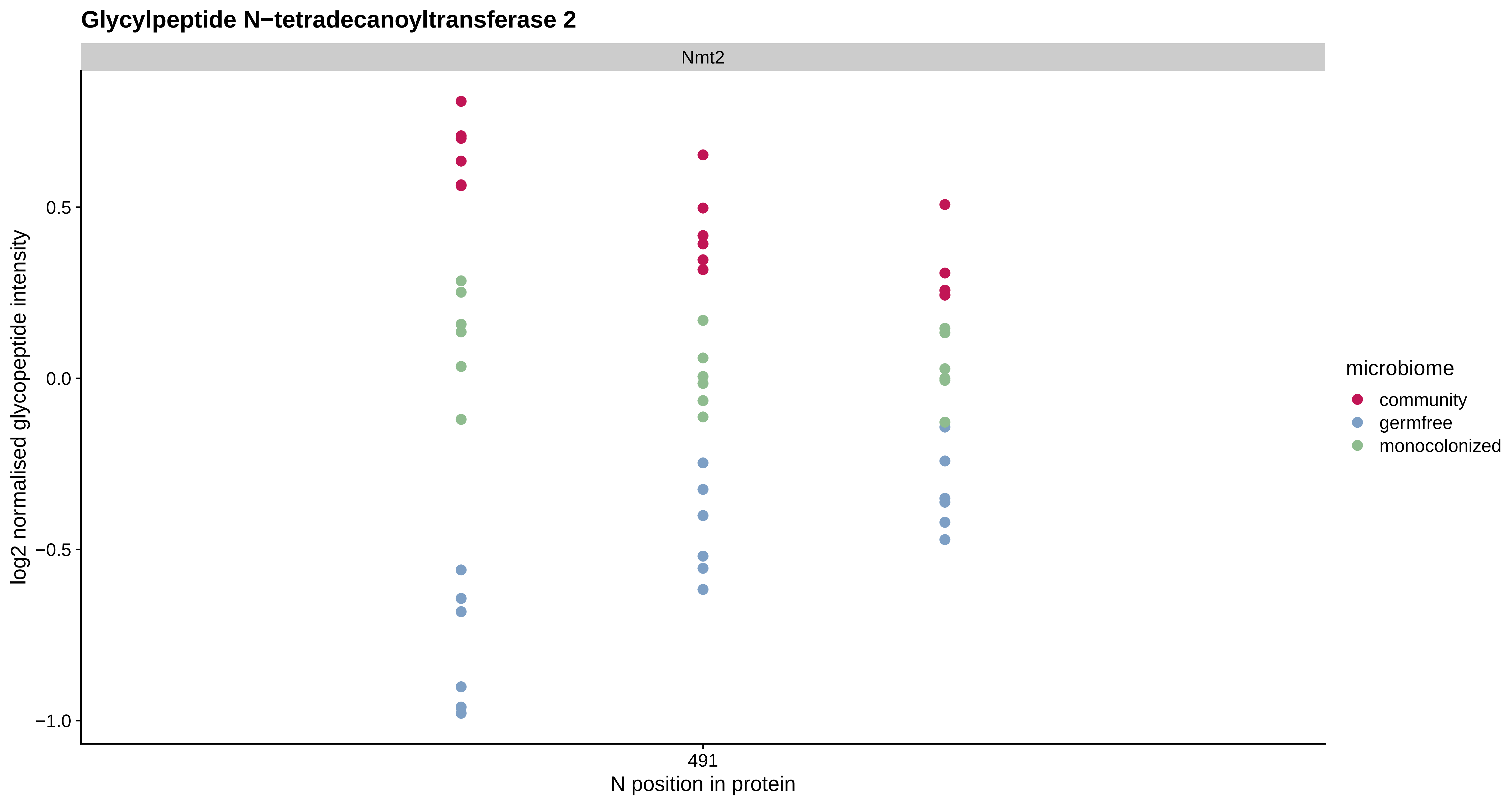

Glycylpeptide N−tetradecanoyltransferase 2
Nmt2
log2 normalised glycopeptide intensity
0.5
0.0
−0.5
−1.0
491
N position in protein
microbiome
community
germfree
monocolonized

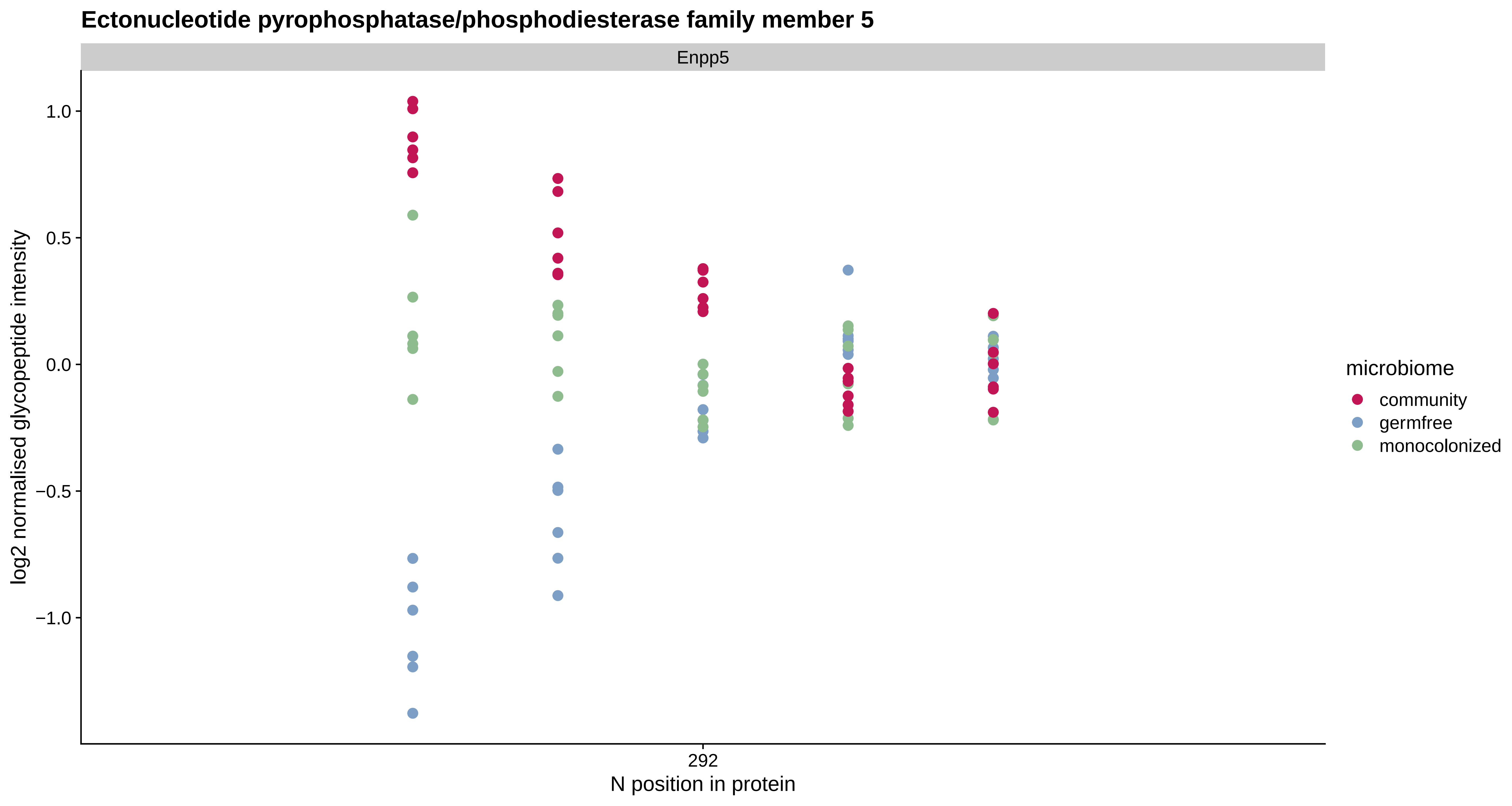

Ectonucleotide pyrophosphatase/phosphodiesterase family member 5
Enpp5
1.0
0.5
0.0
−0.5
−1.0
log2 normalised glycopeptide intensity
292
N position in protein
microbiome
community
germfree
monocolonized

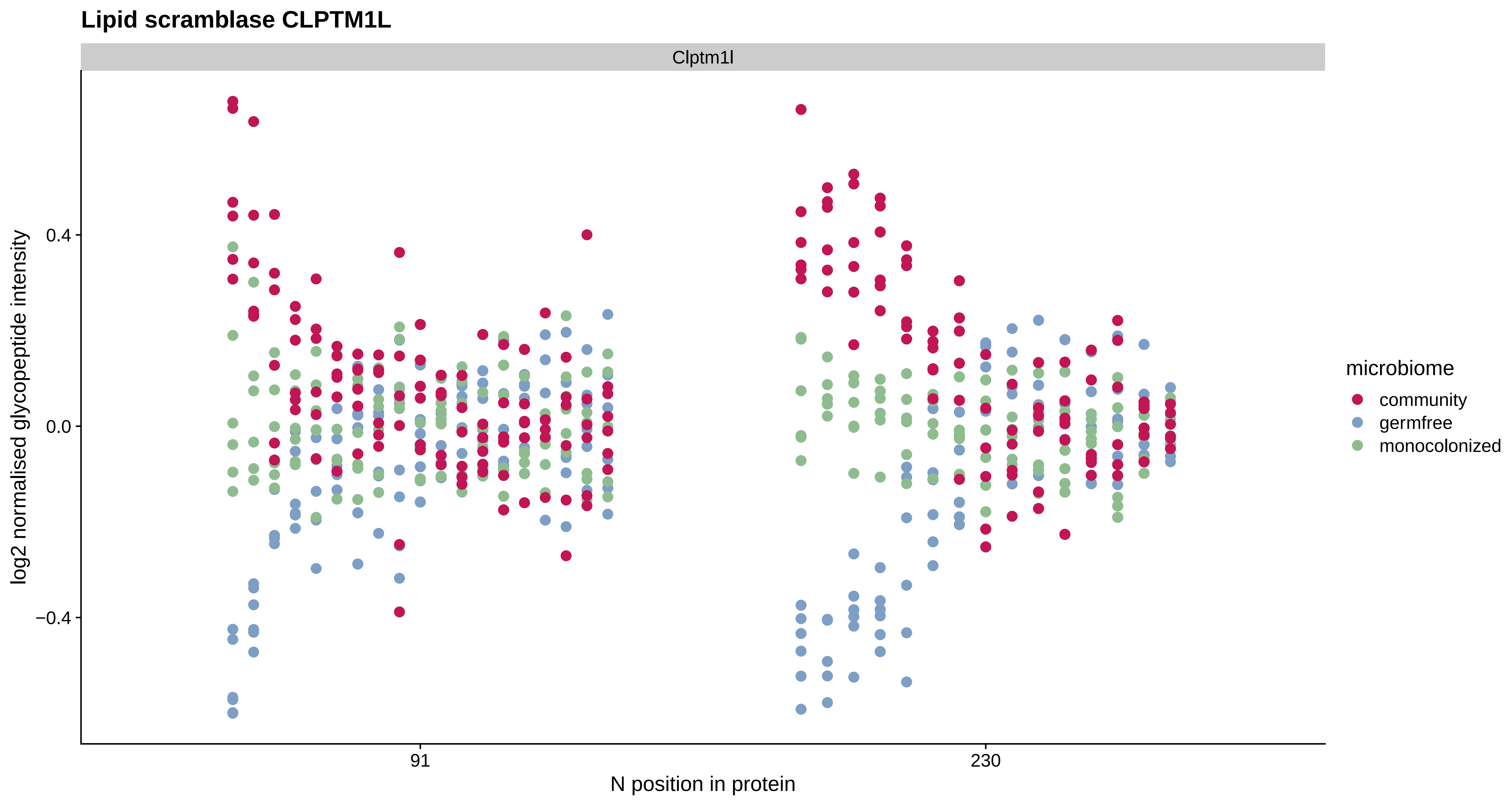

Lipid scramblase CLPTM1L
Clptm1l
0.4
0.0
−0.4
log2 normalised glycopeptide intensity
91
230
N position in protein
microbiome
community
germfree
monocolonized

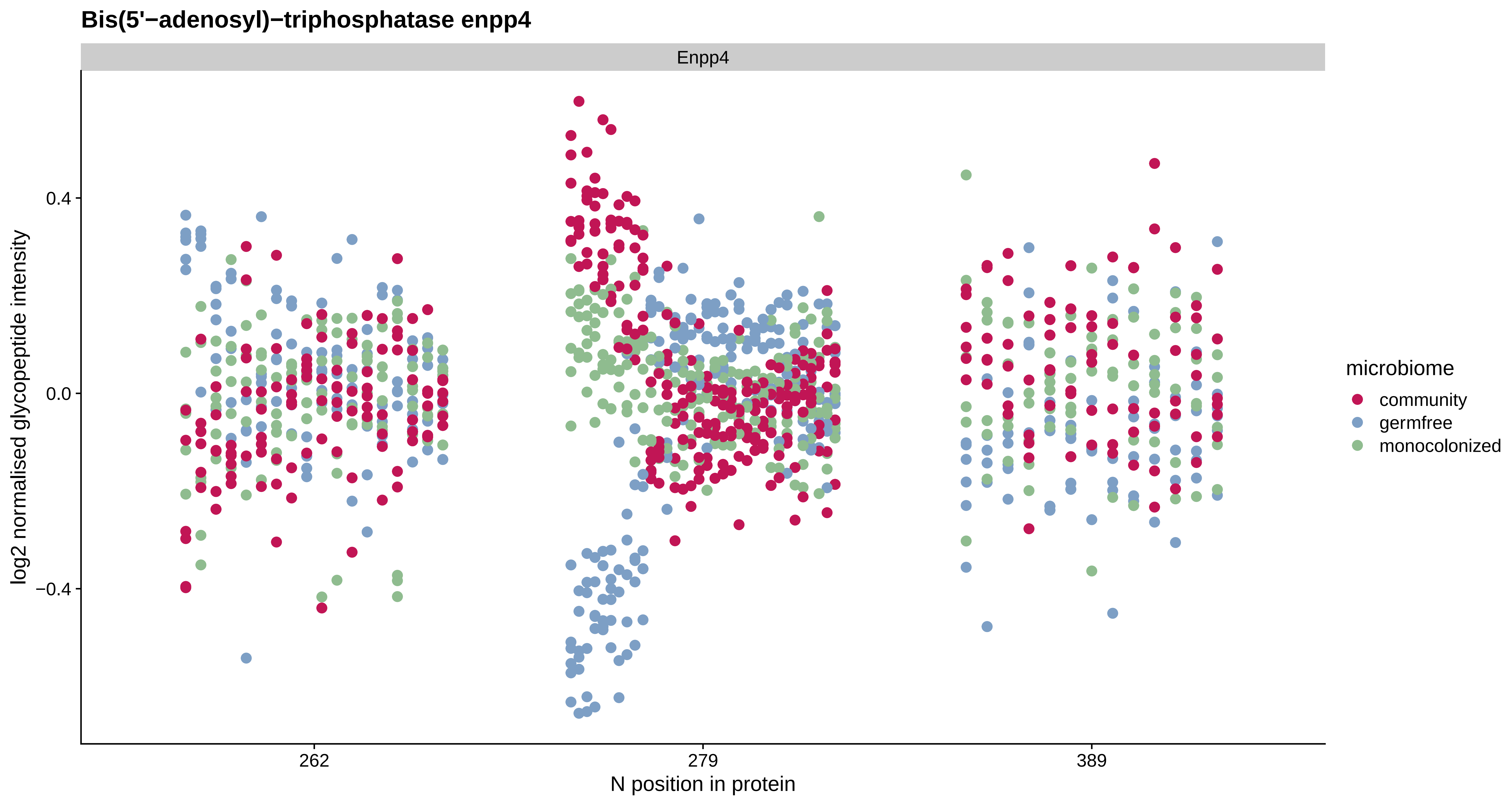

Bis(5'−adenosyl)−triphosphatase enpp4
Enpp4
0.4
0.0
−0.4
log2 normalised glycopeptide intensity
262
279
389
N position in protein
microbiome
community
germfree
monocolonized

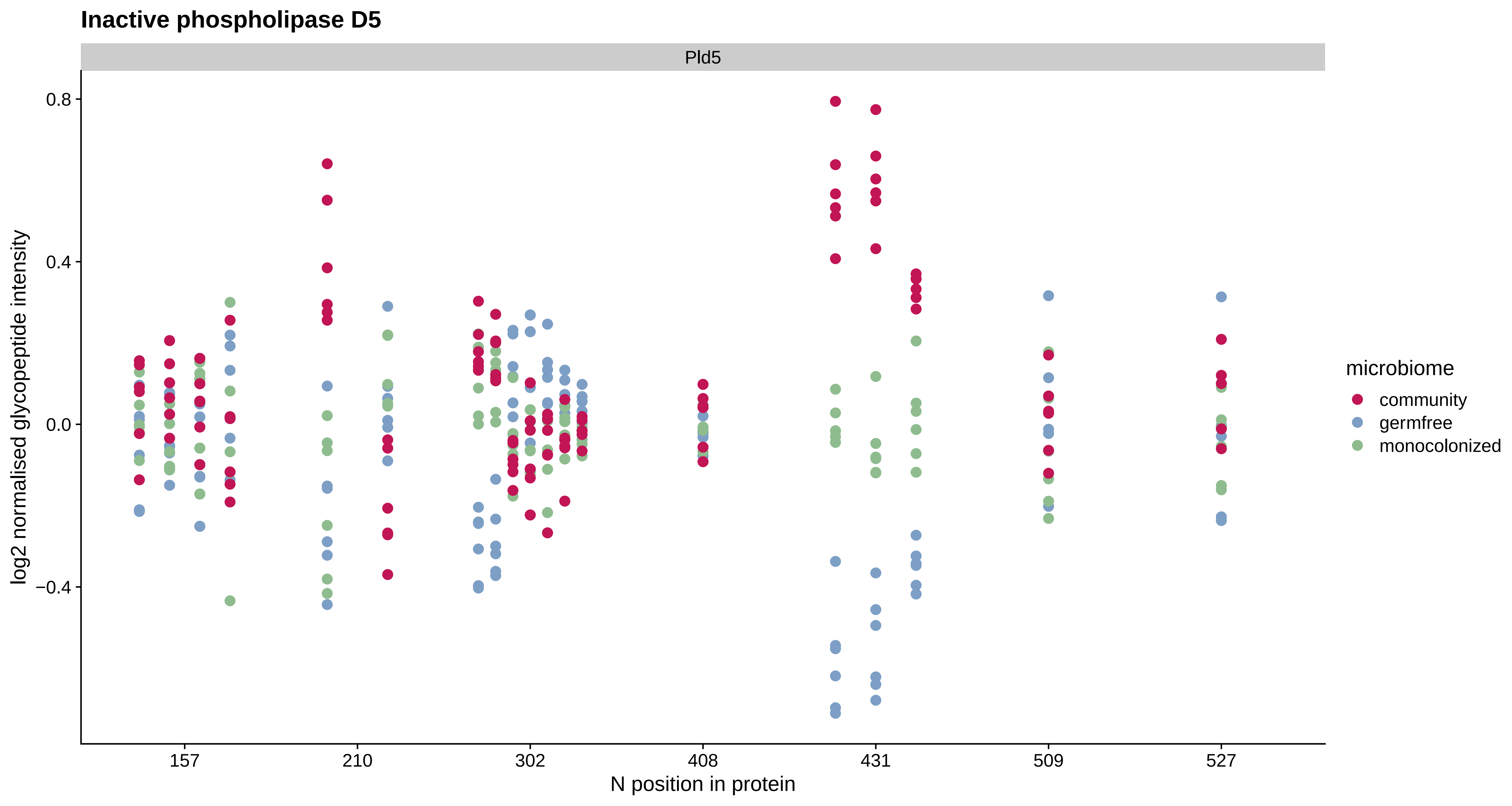

Inactive phospholipase D5
Pld5
log2 normalised glycopeptide intensity
0.8
0.4
0.0
−0.4
157
210
302
408
431
509
527
N position in protein
microbiome
community
germfree
monocolonized

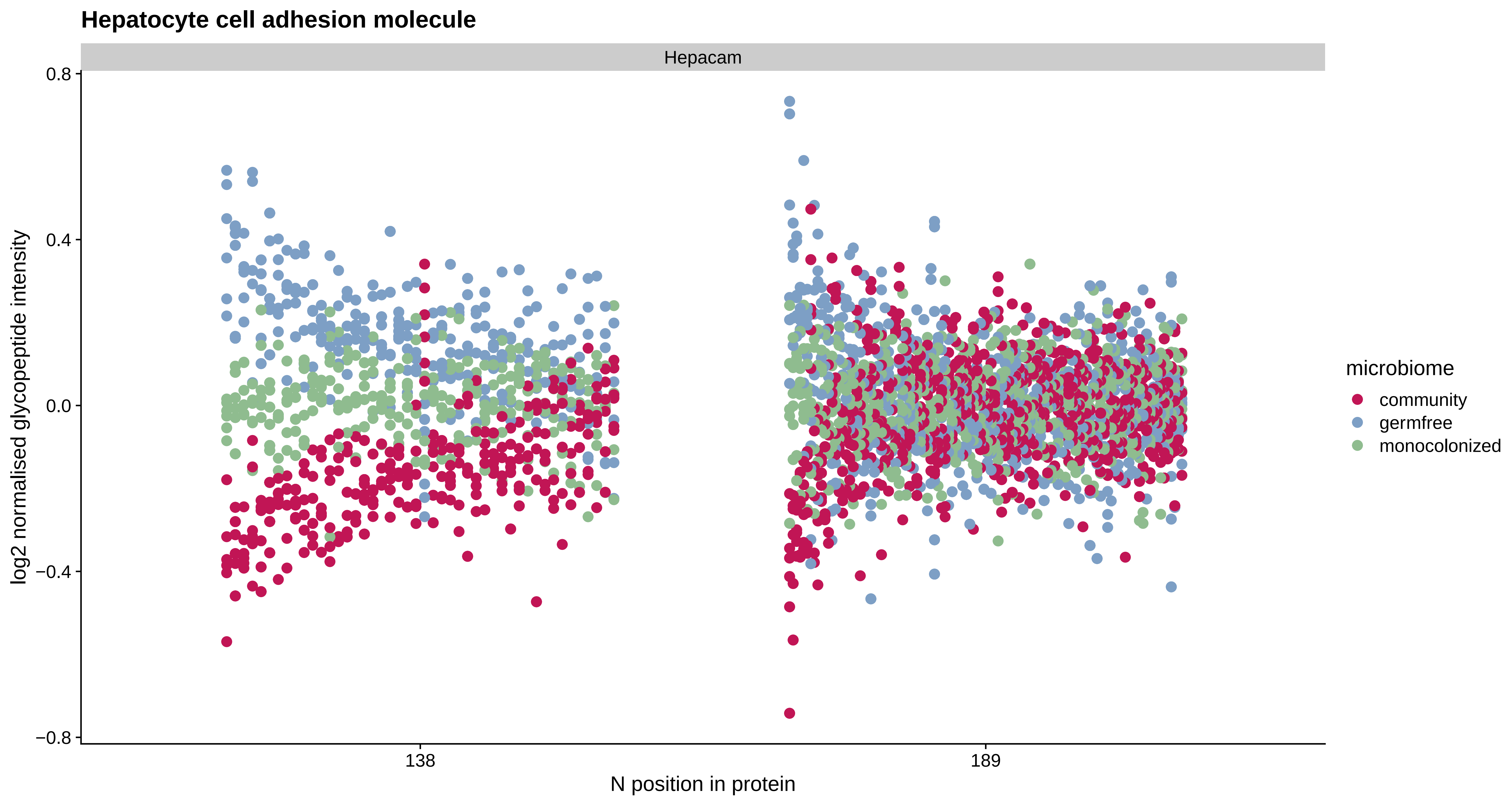
Hepatocyte cell adhesion molecule
Hepacam
0.8
0.4
0.0
−0.4
−0.8
log2 normalised glycopeptide intensity
138
189
N position in protein
microbiome
community
germfree
monocolonized

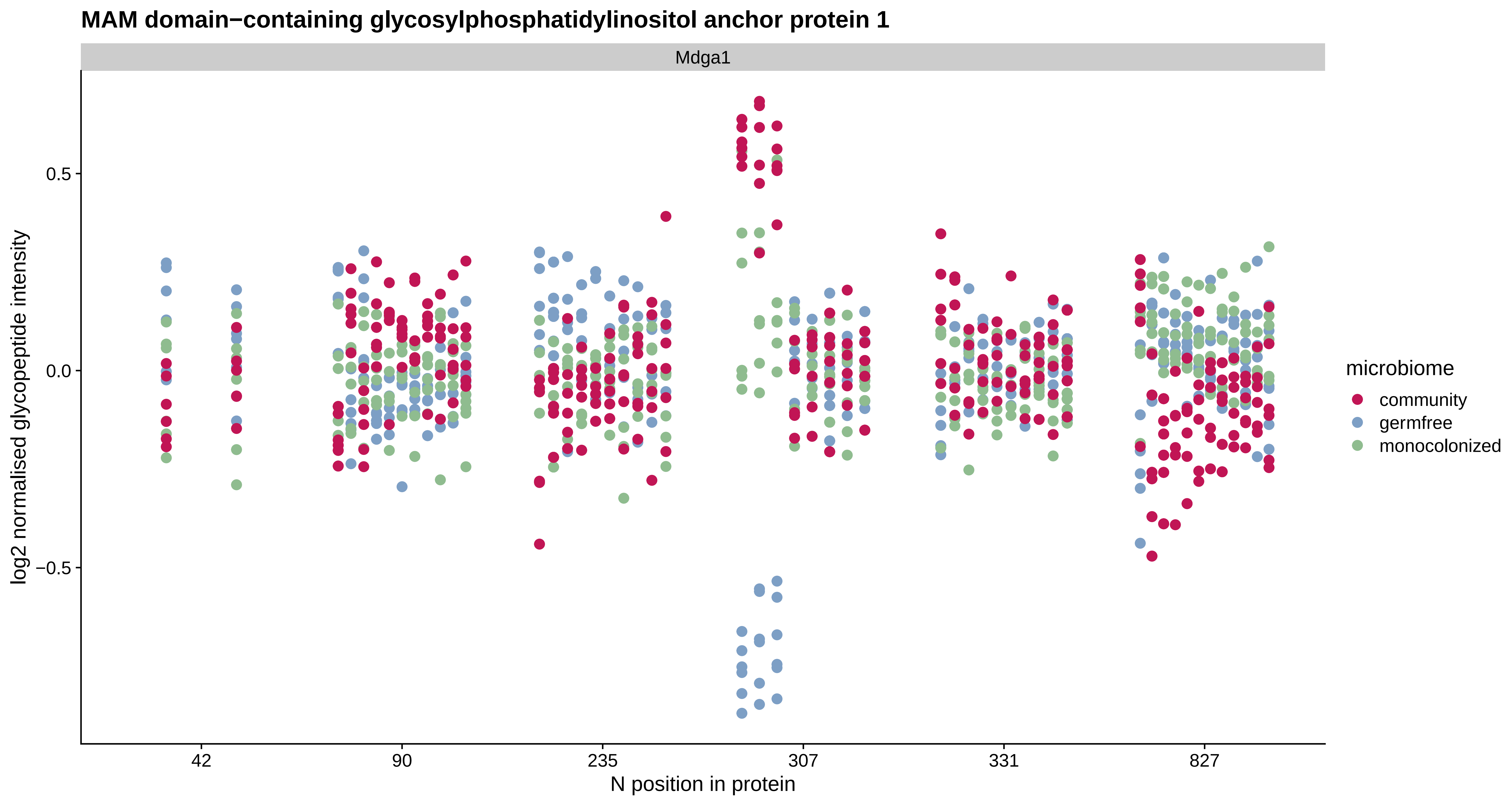
MAM domain−containing glycosylphosphatidylinositol anchor protein 1
Mdga1
log2 normalised glycopeptide intensity
0.5
0.0
−0.5
42
90
235
307
331
827
N position in protein
microbiome
community
germfree
monocolonized

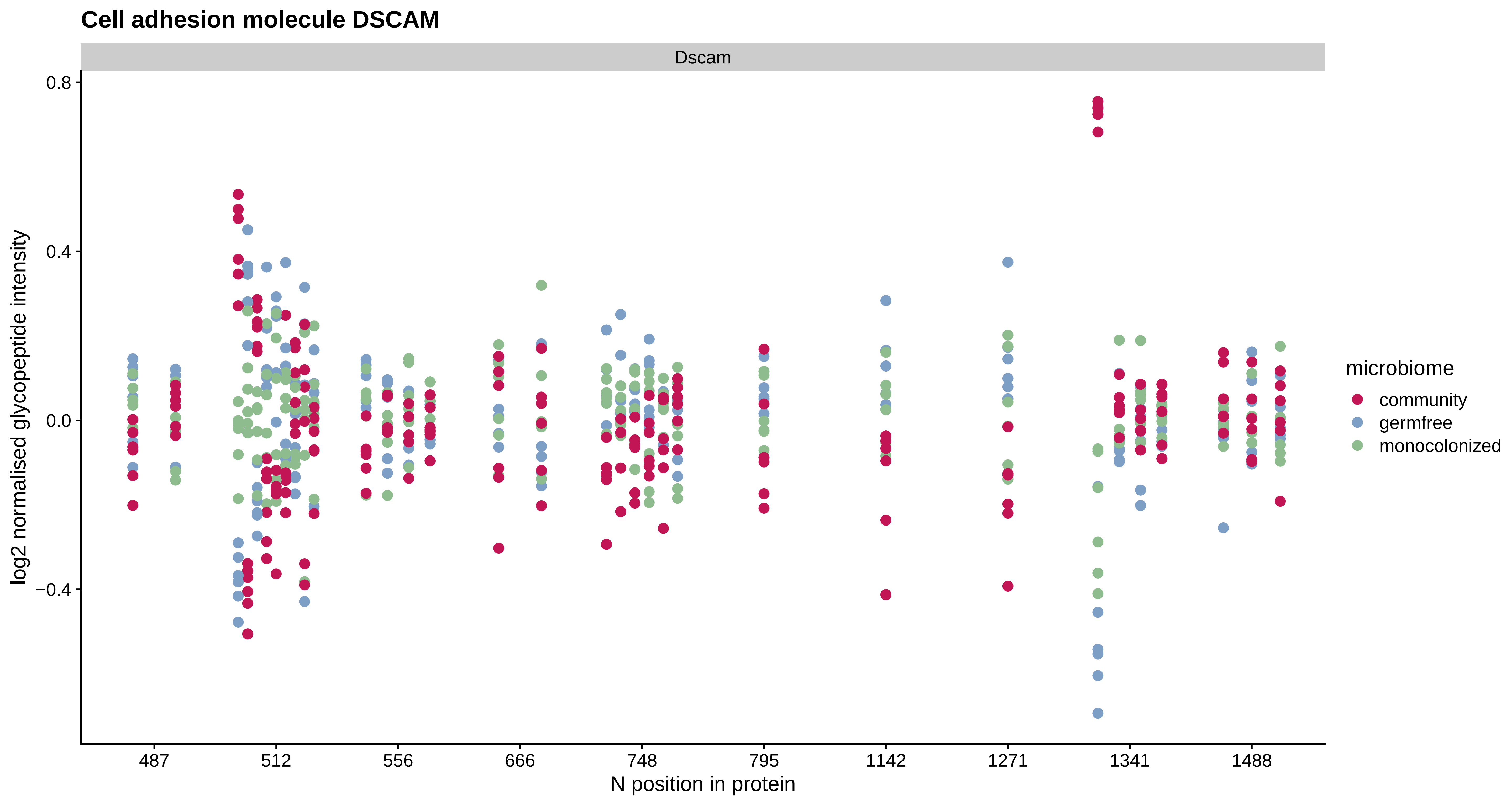

Cell adhesion molecule DSCAM
Dscam
0.8
0.4
0.0
−0.4
log2 normalised glycopeptide intensity
487
512
556
666
748
795
1142
1271
1341
1488
N position in protein
microbiome
community
germfree
monocolonized

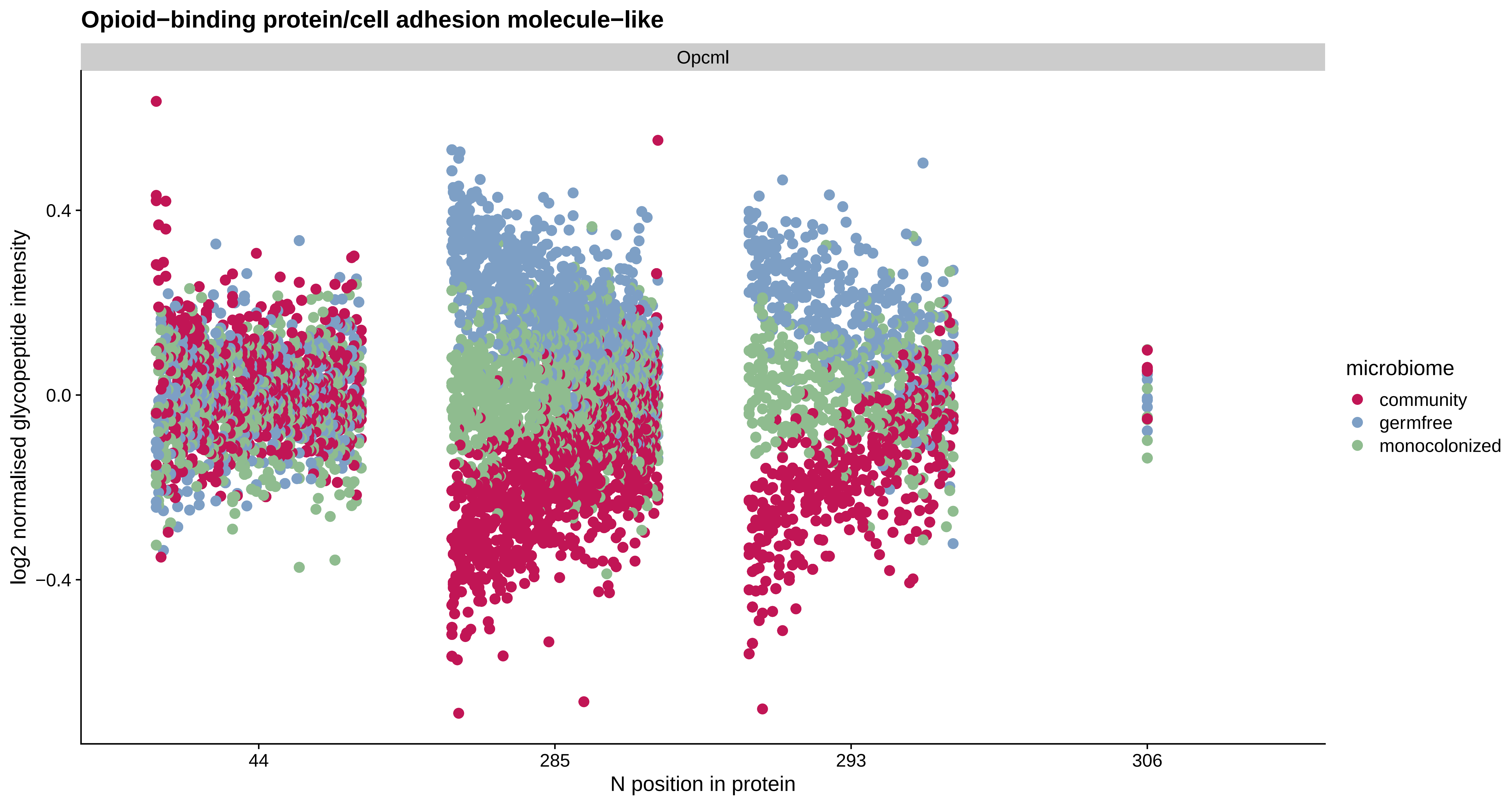
Opioid−binding protein/cell adhesion molecule−like
Opcml
log2 normalised glycopeptide intensity
0.4
0.0
−0.4
44
285
293
306
N position in protein
microbiome
community
germfree
monocolonized

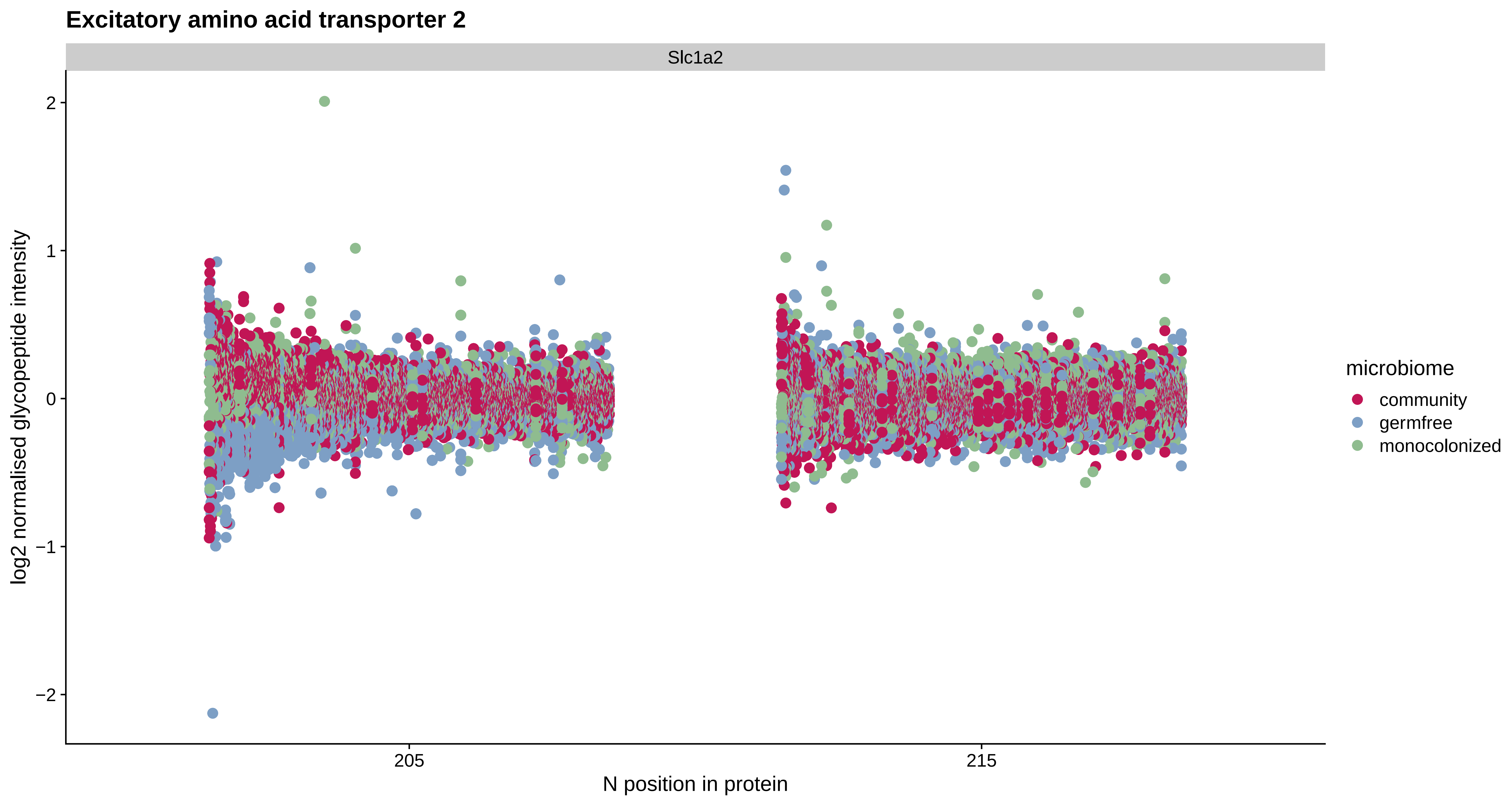
Excitatory amino acid transporter 2
Slc1a2
log2 normalised glycopeptide intensity
2
1
0
−1
−2
205
215
N position in protein
microbiome
community
germfree
monocolonized

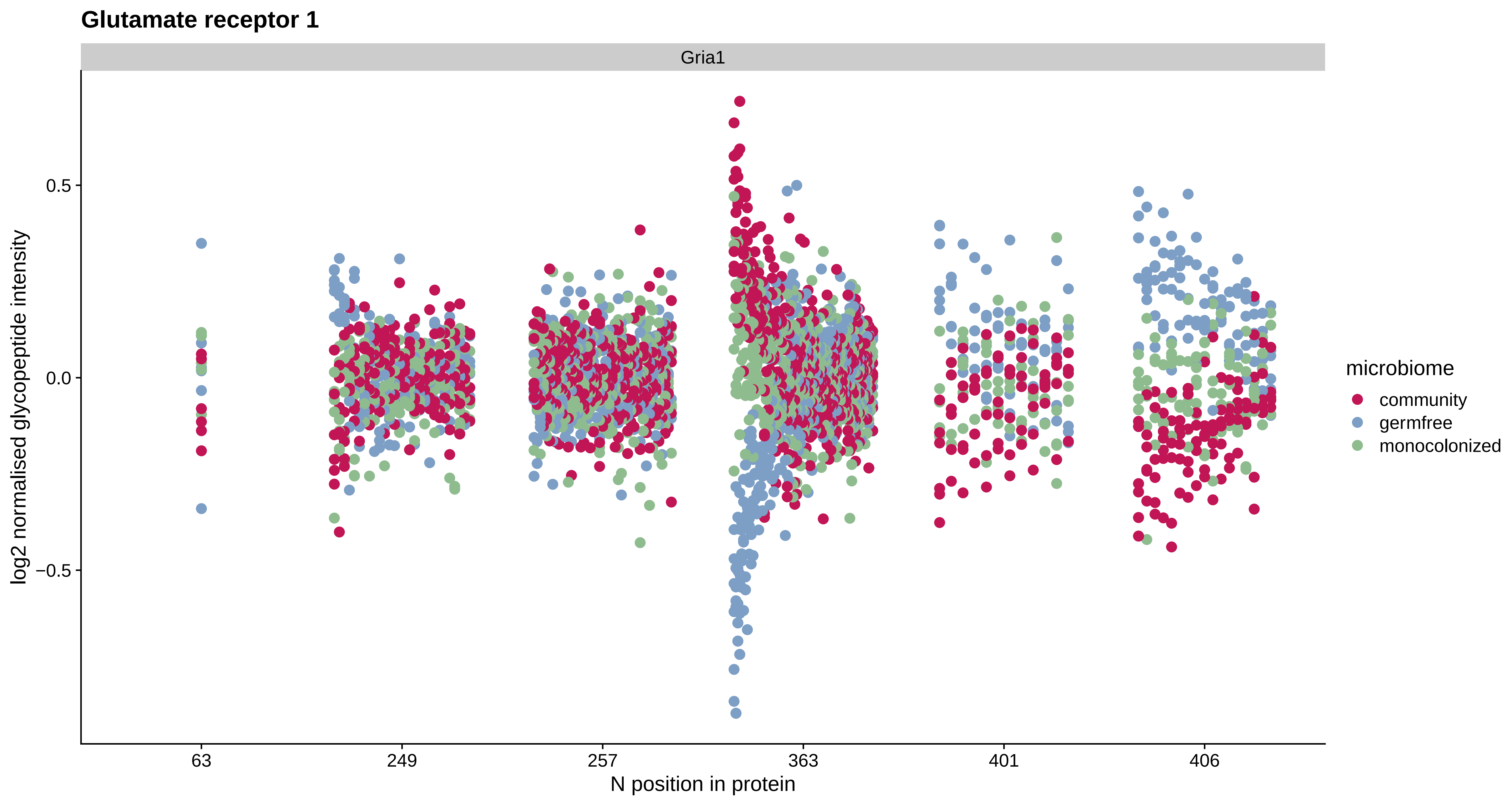

Glutamate receptor 1
Gria1
0.5
0.0
−0.5
log2 normalised glycopeptide intensity
63
249
257
363
401
406
N position in protein
microbiome
community
germfree
monocolonized

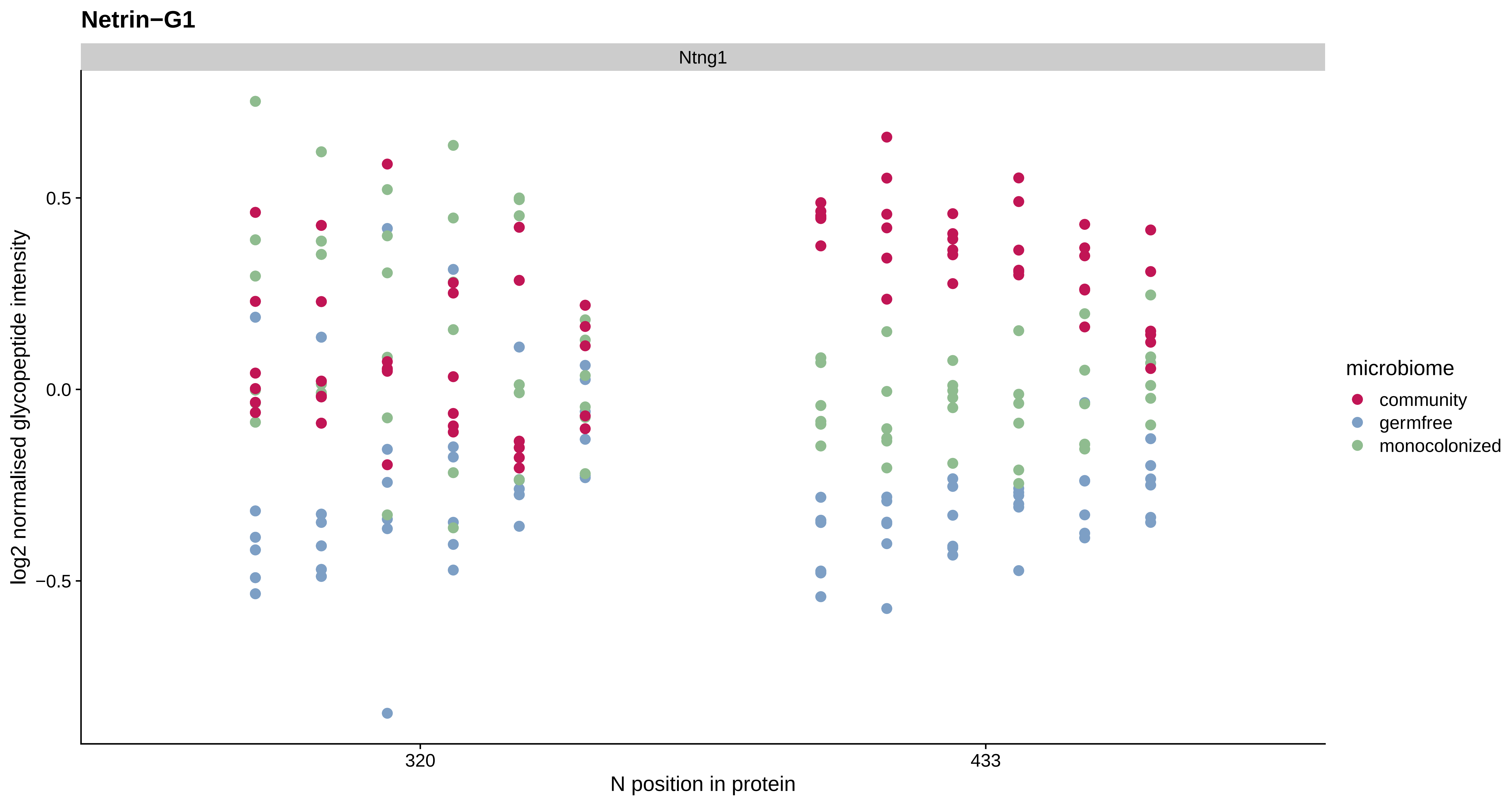
Netrin−G1
Ntng1
log2 normalised glycopeptide intensity
0.5
0.0
−0.5
320
433
N position in protein
microbiome
community
germfree
monocolonized

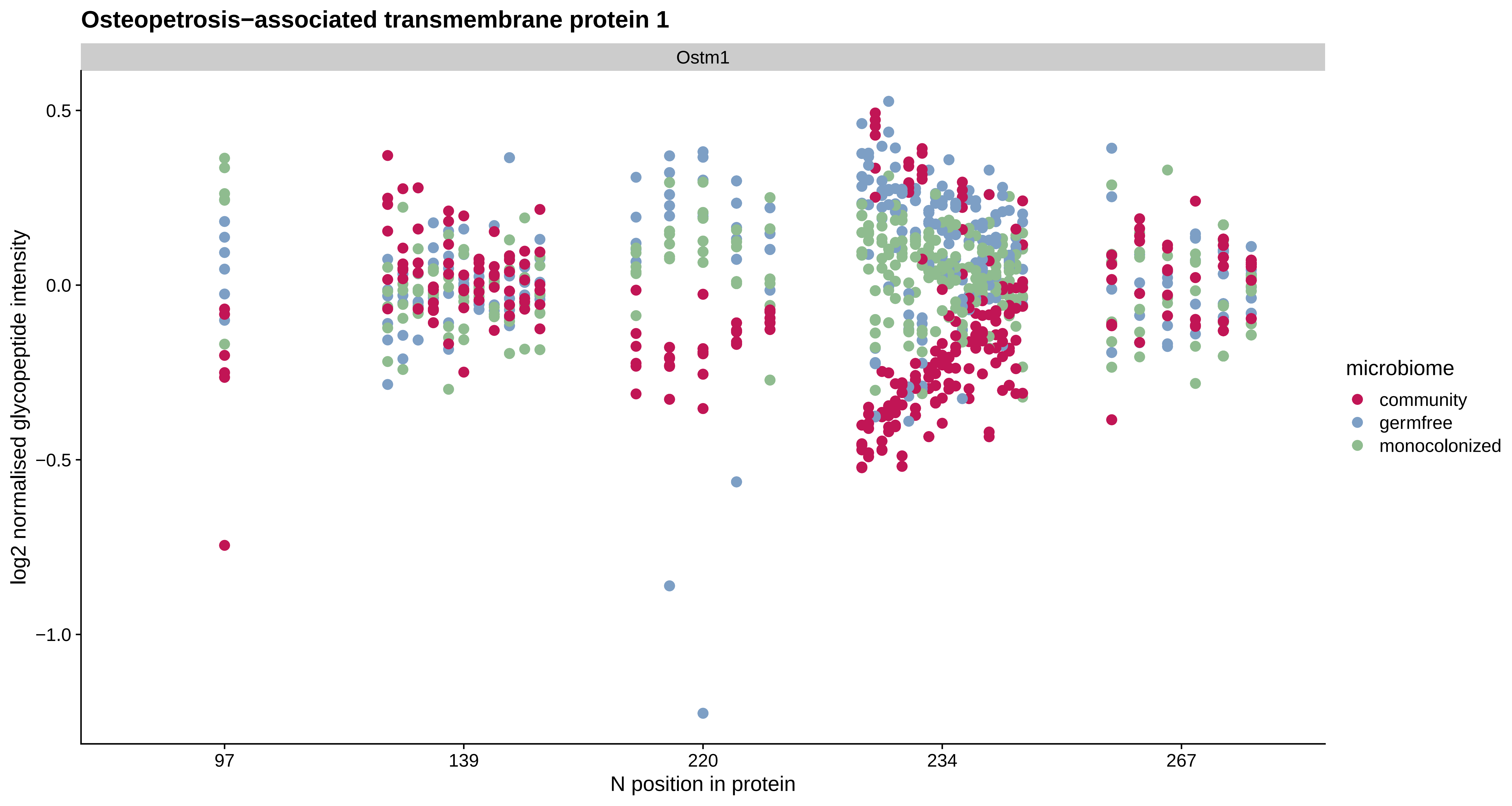
Osteopetrosis−associated transmembrane protein 1
Ostm1
0.5
0.0
−0.5
−1.0
log2 normalised glycopeptide intensity
97
139
220
234
267
N position in protein
microbiome
community
germfree
monocolonized

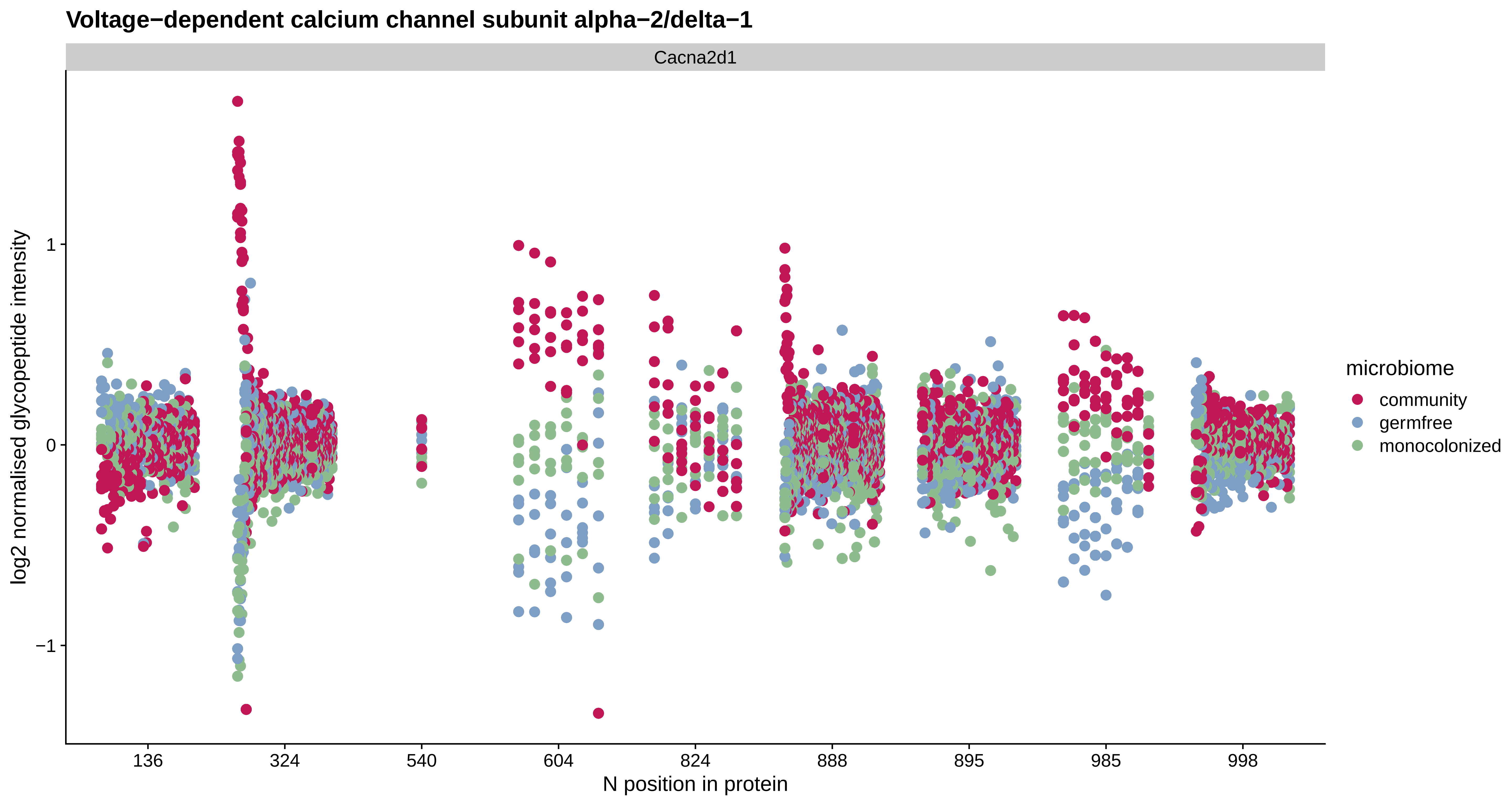

Voltage−dependent calcium channel subunit alpha−2/delta−1
Cacna2d1
log2 normalised glycopeptide intensity
1
0
−1
136
324
540
604
824
888
895
985
998
N position in protein
microbiome
community
germfree
monocolonized

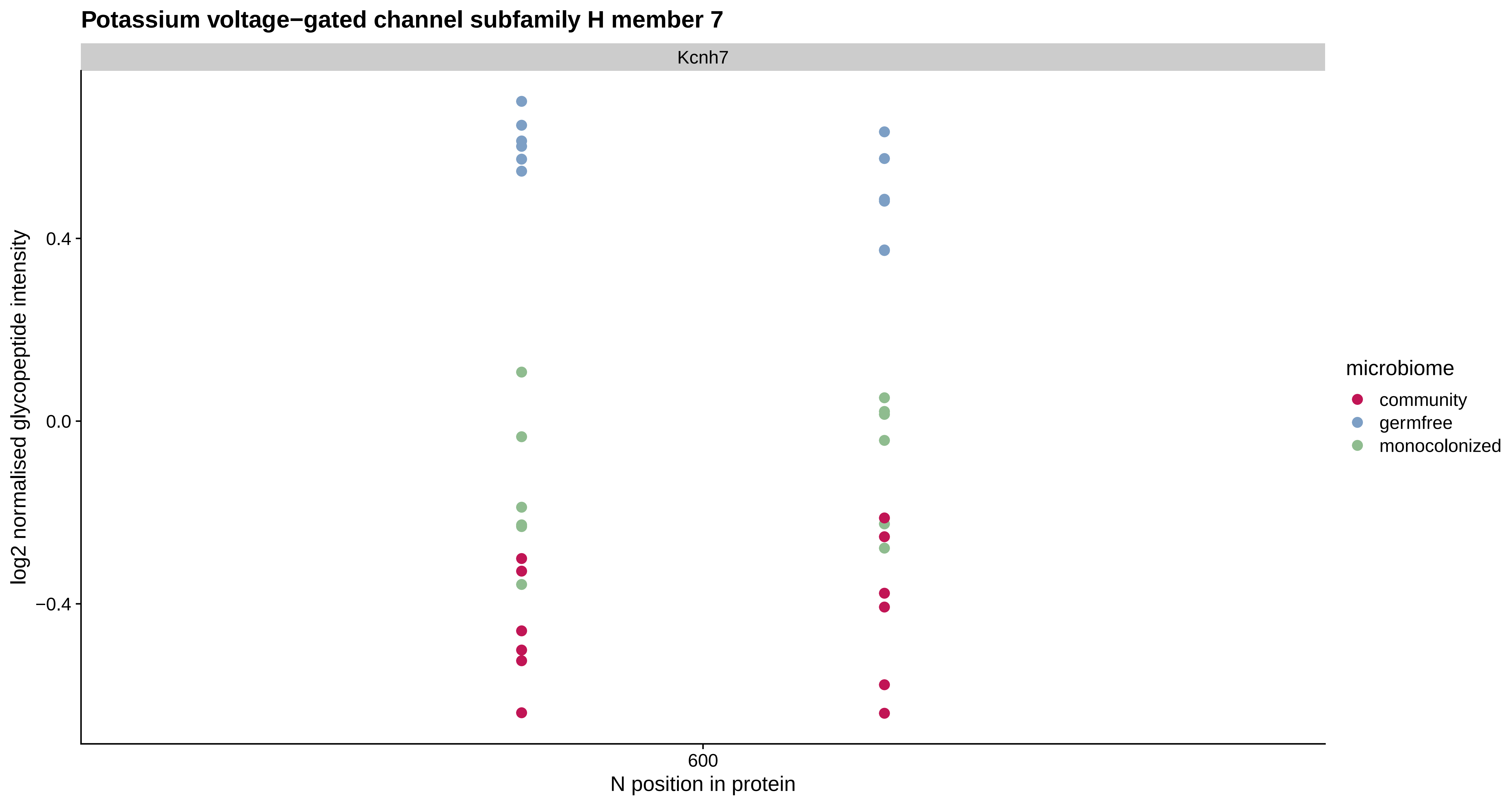

Potassium voltage−gated channel subfamily H member 7
Kcnh7
log2 normalised glycopeptide intensity
0.4
0.0
−0.4
600
N position in protein
microbiome
community
germfree
monocolonized

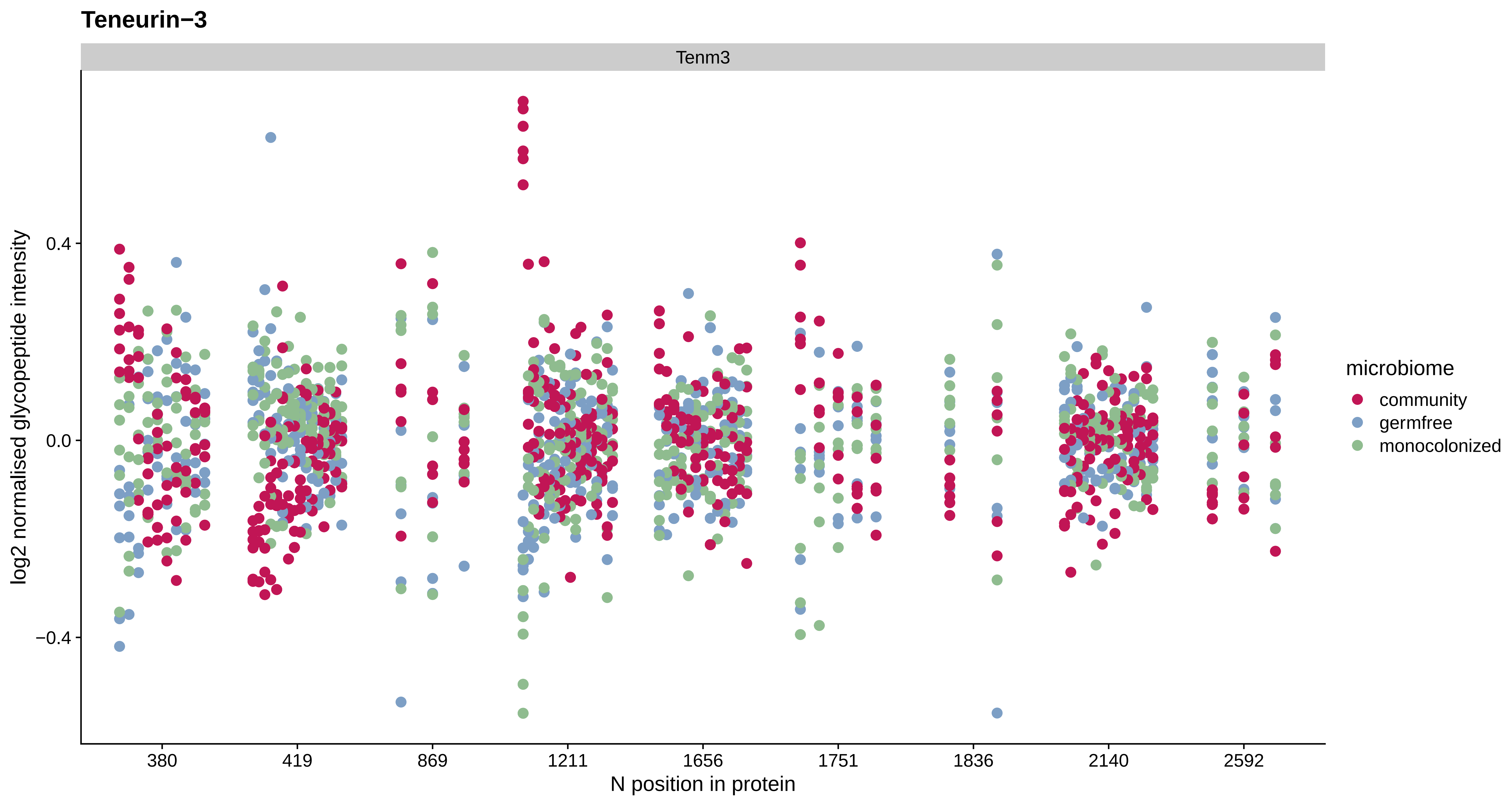
Teneurin−3
Tenm3
log2 normalised glycopeptide intensity
0.4
0.0
−0.4
380
419
869
1211
1656
1751
1836
2140
2592
N position in protein
microbiome
community
germfree
monocolonized

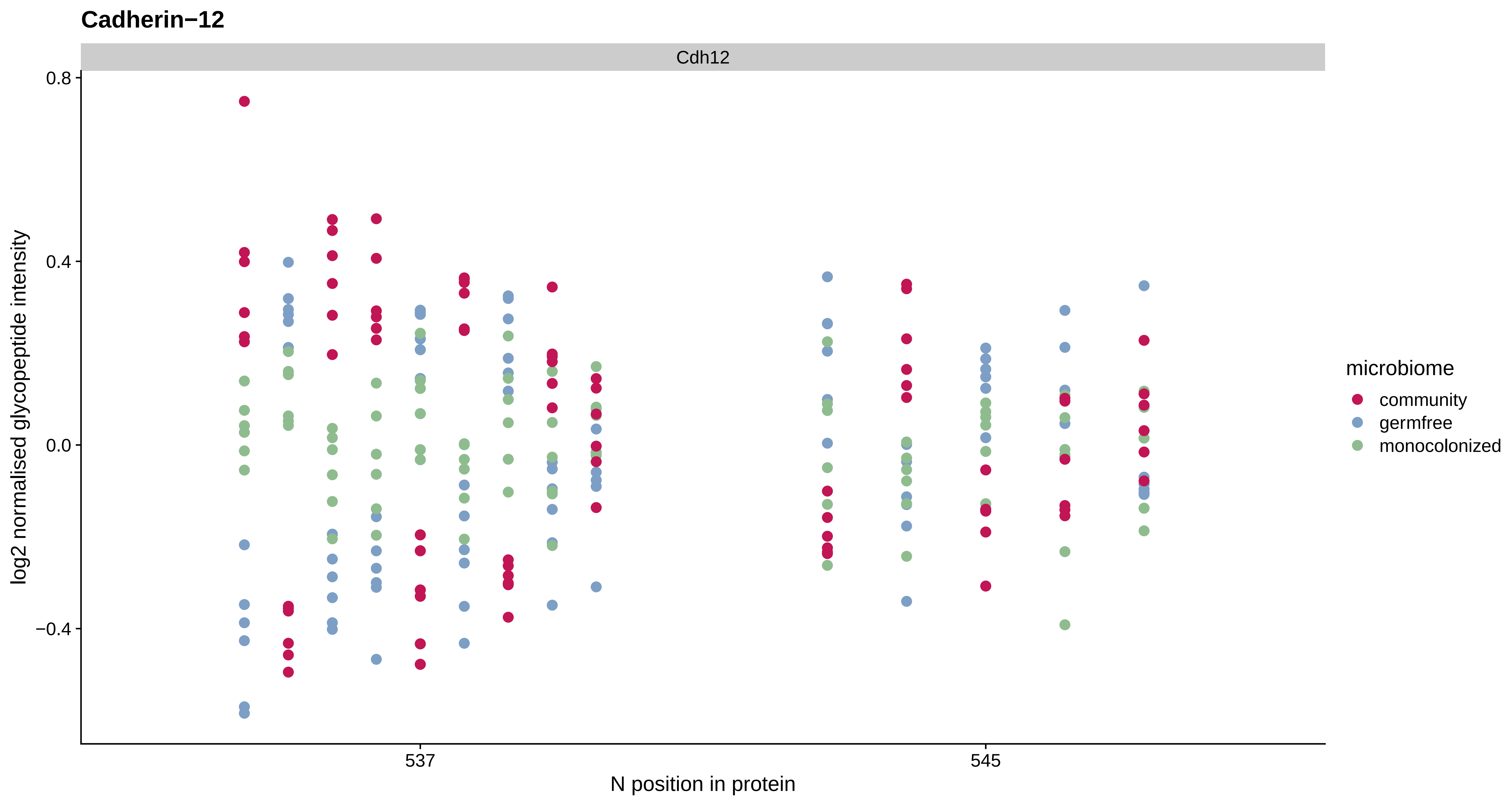

Cadherin−12
Cdh12
0.8
0.4
0.0
−0.4
log2 normalised glycopeptide intensity
537
545
N position in protein
microbiome
community
germfree
monocolonized

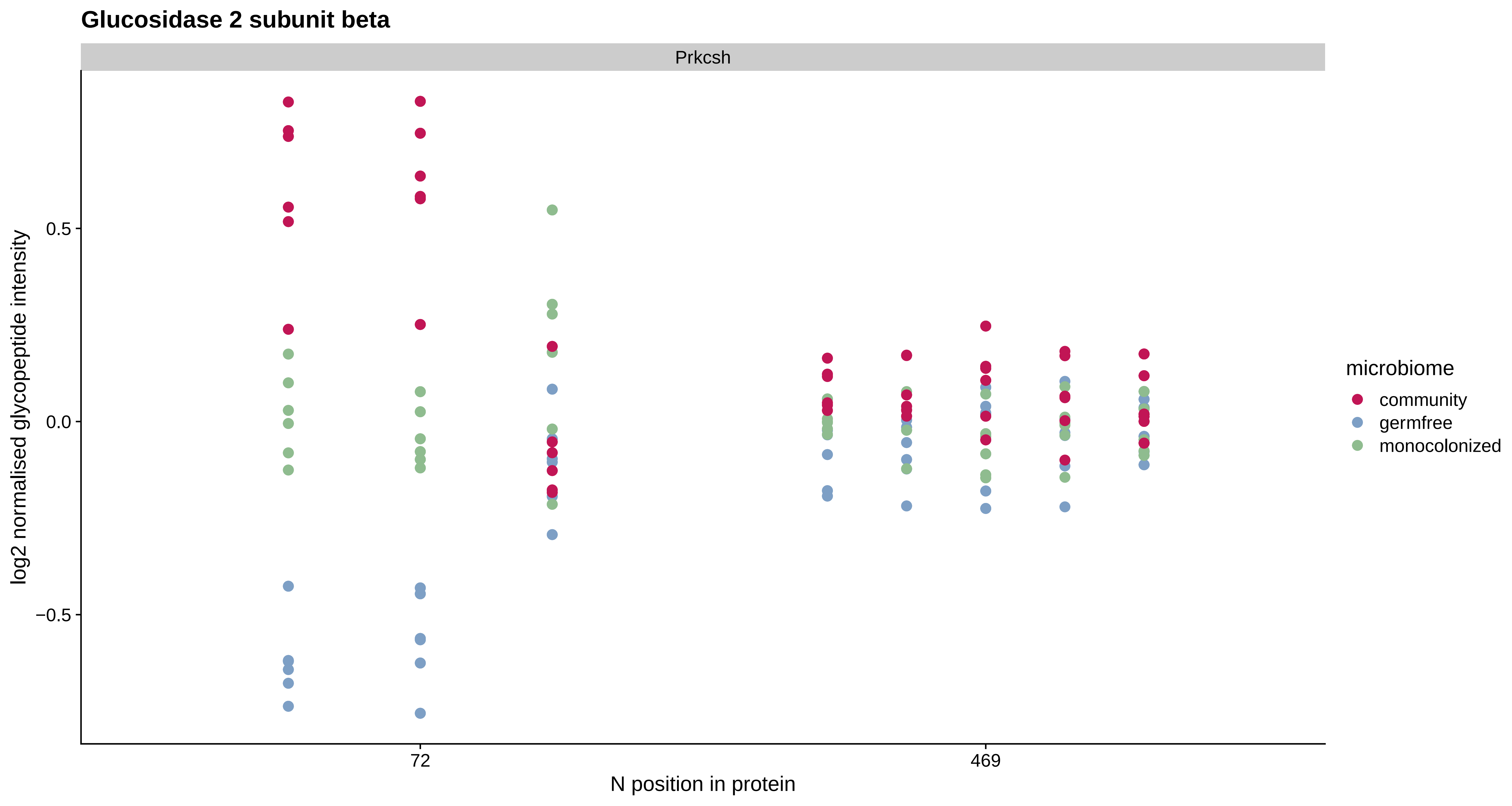

Glucosidase 2 subunit beta
Prkcsh
log2 normalised glycopeptide intensity
0.5
0.0
−0.5
72
469
N position in protein
microbiome
community
germfree
monocolonized

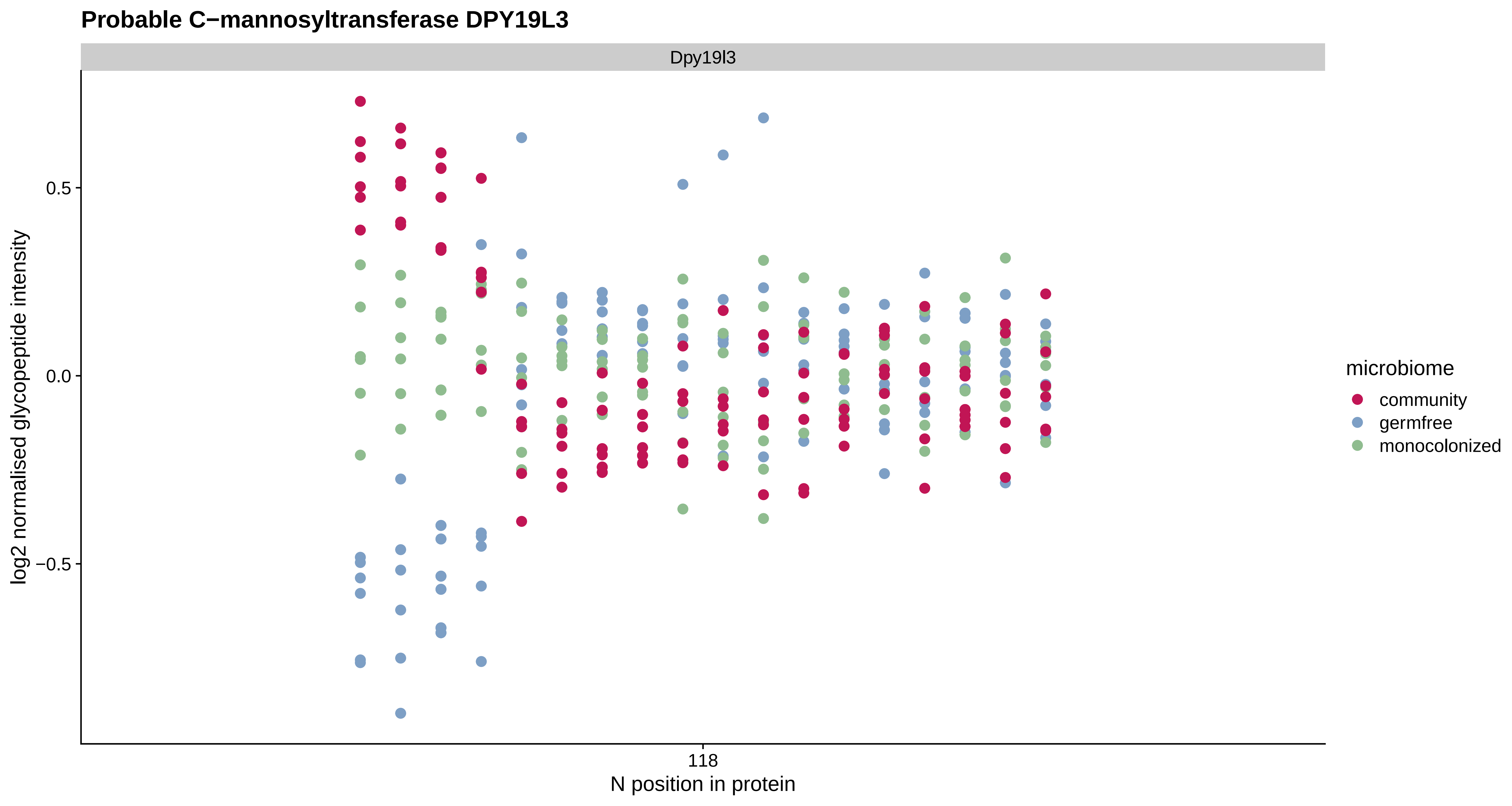

Probable C−mannosyltransferase DPY19L3
Dpy19l3
log2 normalised glycopeptide intensity
0.5
0.0
−0.5
118
N position in protein
microbiome
community
germfree
monocolonized

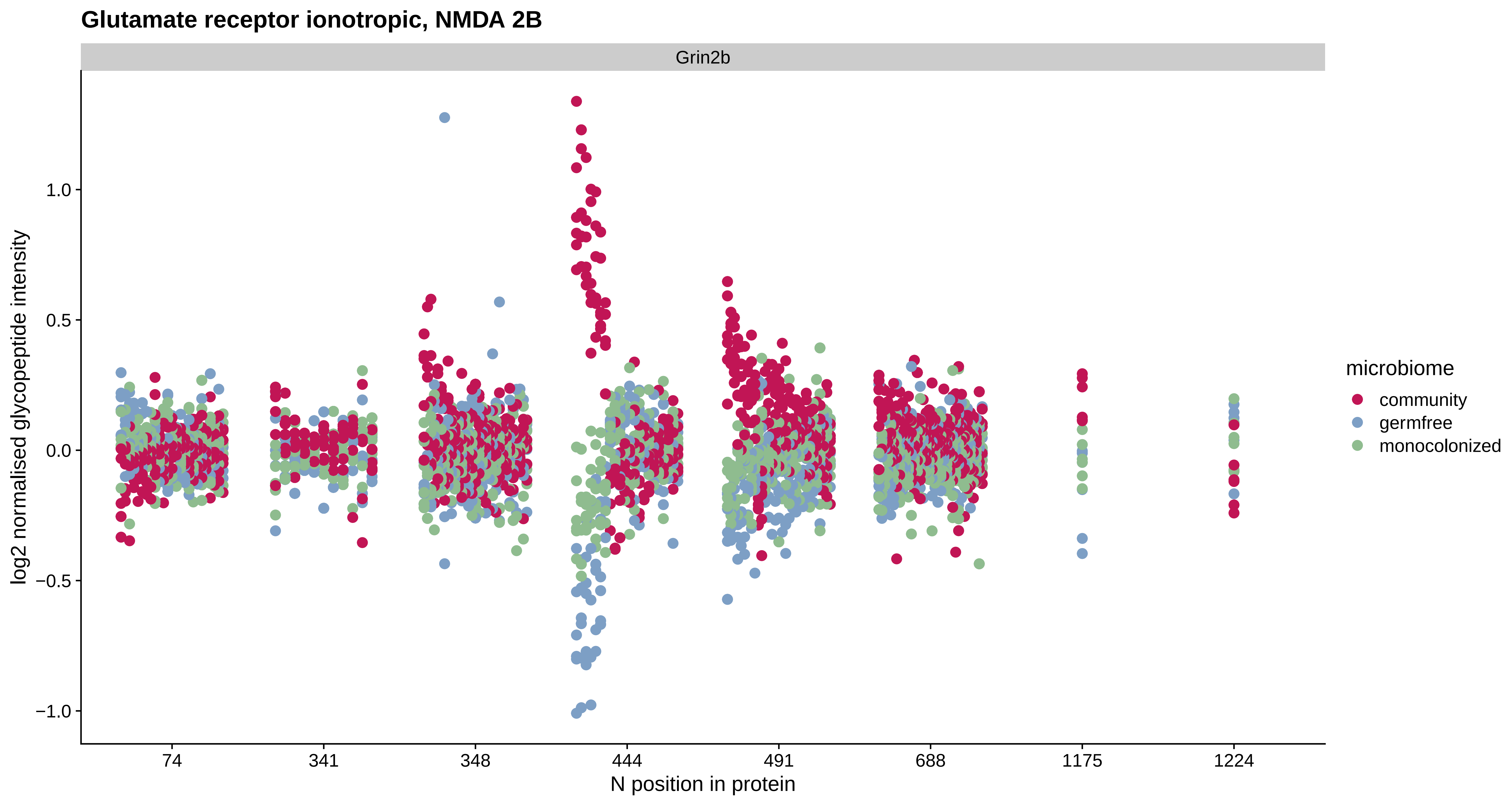
Glutamate receptor ionotropic, NMDA 2B
Grin2b
log2 normalised glycopeptide intensity
1.0
0.5
0.0
−0.5
−1.0
74
341
348
444
491
688
1175
1224
N position in protein
microbiome
community
germfree
monocolonized

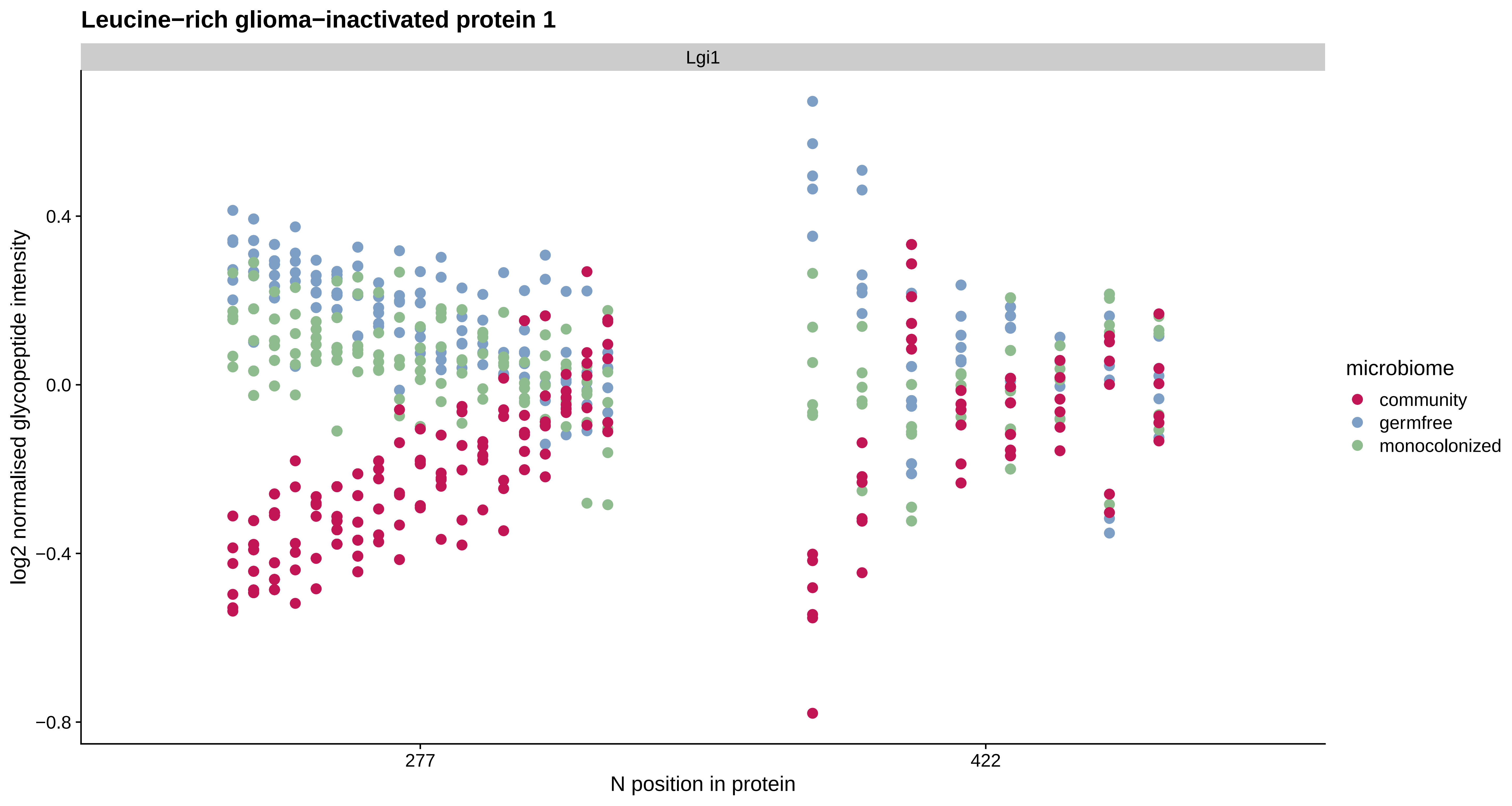
Leucine−rich glioma−inactivated protein 1
Lgi1
log2 normalised glycopeptide intensity
0.4
0.0
−0.4
−0.8
277
422
N position in protein
microbiome
community
germfree
monocolonized

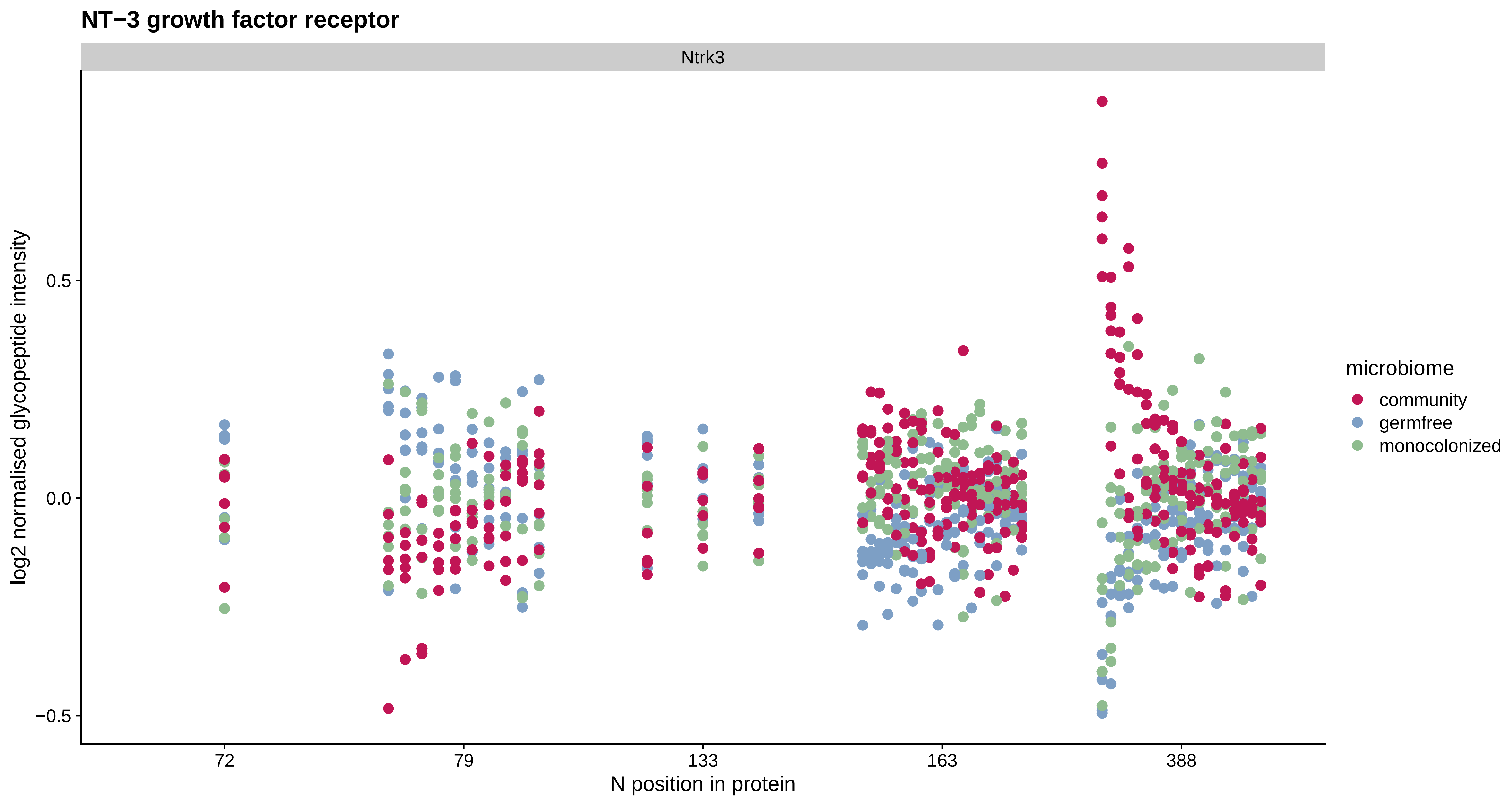
NT−3 growth factor receptor
Ntrk3
log2 normalised glycopeptide intensity
0.5
0.0
−0.5
72
79
133
163
388
N position in protein
microbiome
community
germfree
monocolonized

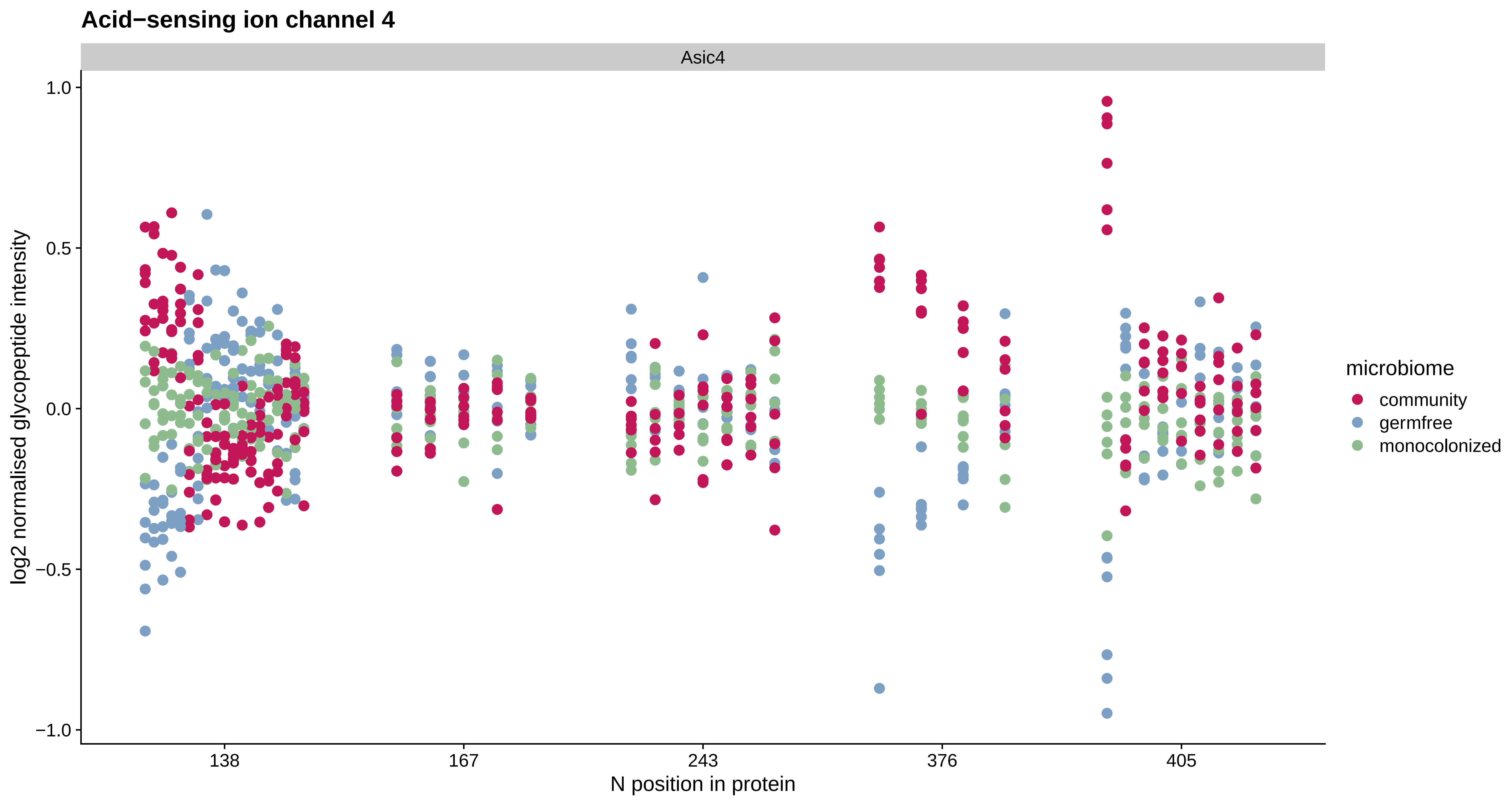
Acid−sensing ion channel 4
Asic4
log2 normalised glycopeptide intensity
1.0
0.5
0.0
−0.5
−1.0
138
167
243
376
405
N position in protein
microbiome
community
germfree
monocolonized

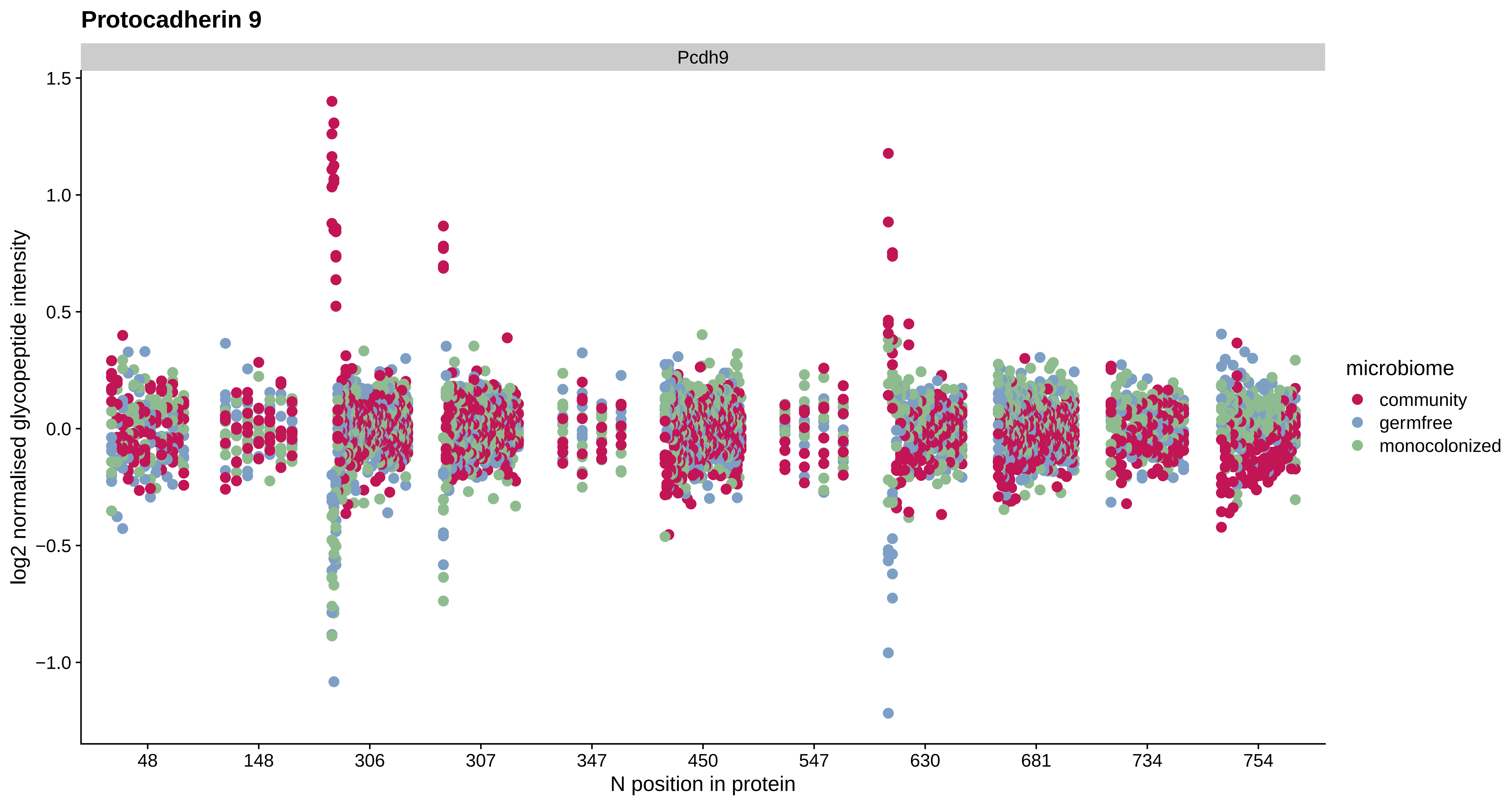
Protocadherin 9
Pcdh9
log2 normalised glycopeptide intensity
1.5
1.0
0.5
0.0
−0.5
−1.0
48
148
306
307
347
450
547
630
681
734
754
N position in protein
microbiome
community
germfree
monocolonized

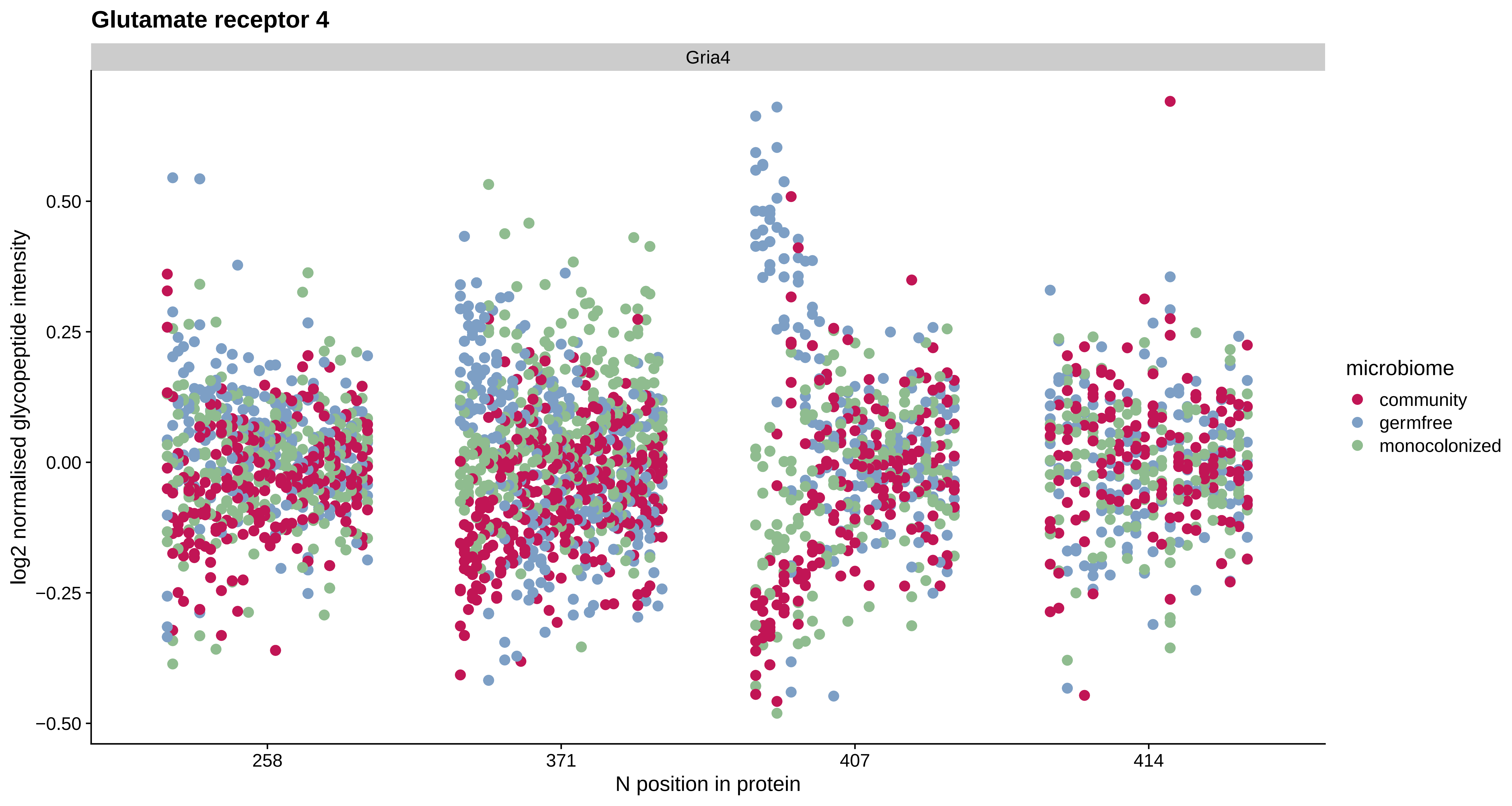

Glutamate receptor 4
Gria4
log2 normalised glycopeptide intensity
0.50
0.25
0.00
−0.25
−0.50
258
371
407
414
N position in protein
microbiome
community
germfree
monocolonized

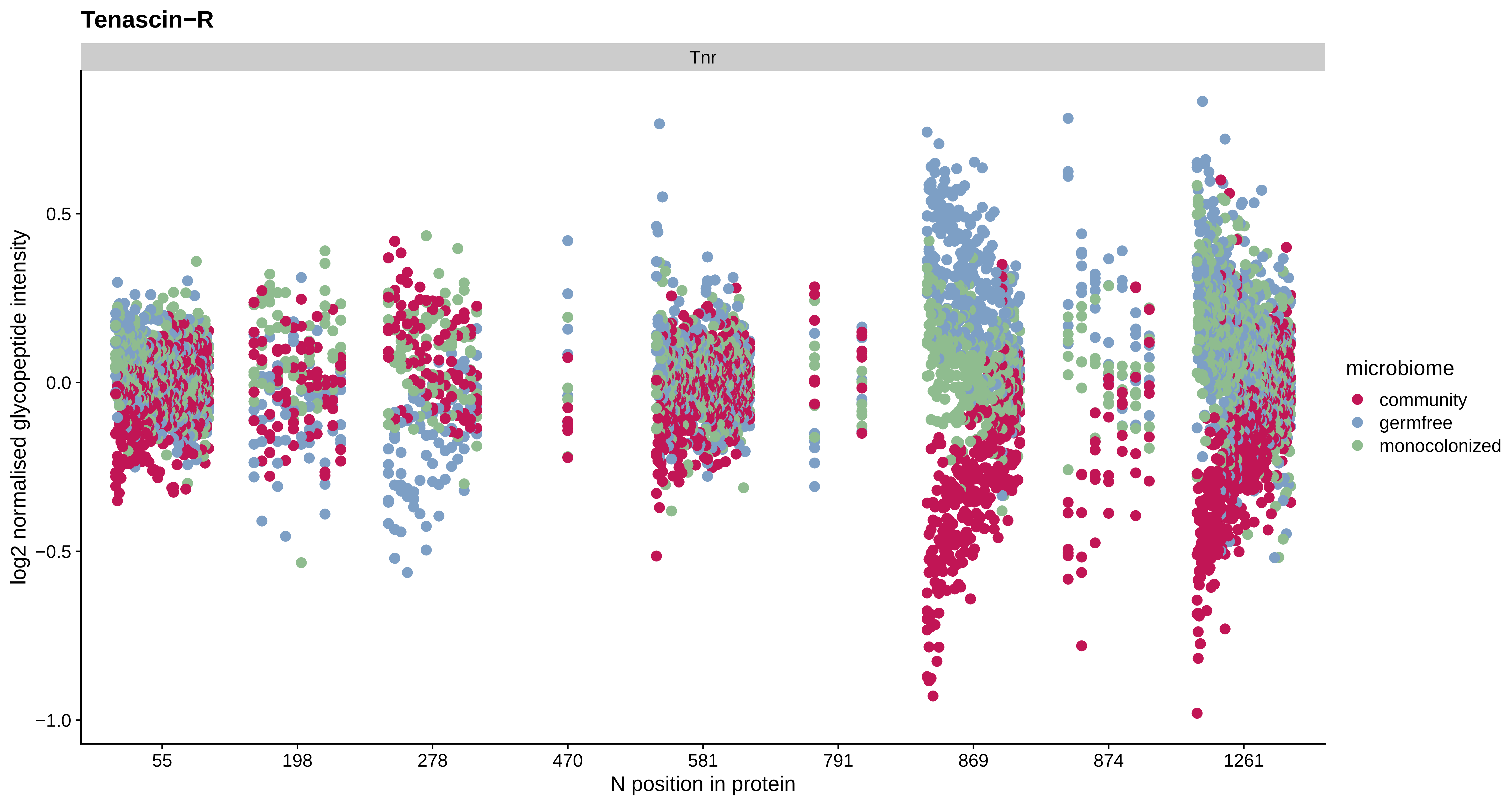

Tenascin−R
Tnr
log2 normalised glycopeptide intensity
0.5
0.0
−0.5
−1.0
55
198
278
470
581
791
869
874
1261
N position in protein
microbiome
community
germfree
monocolonized

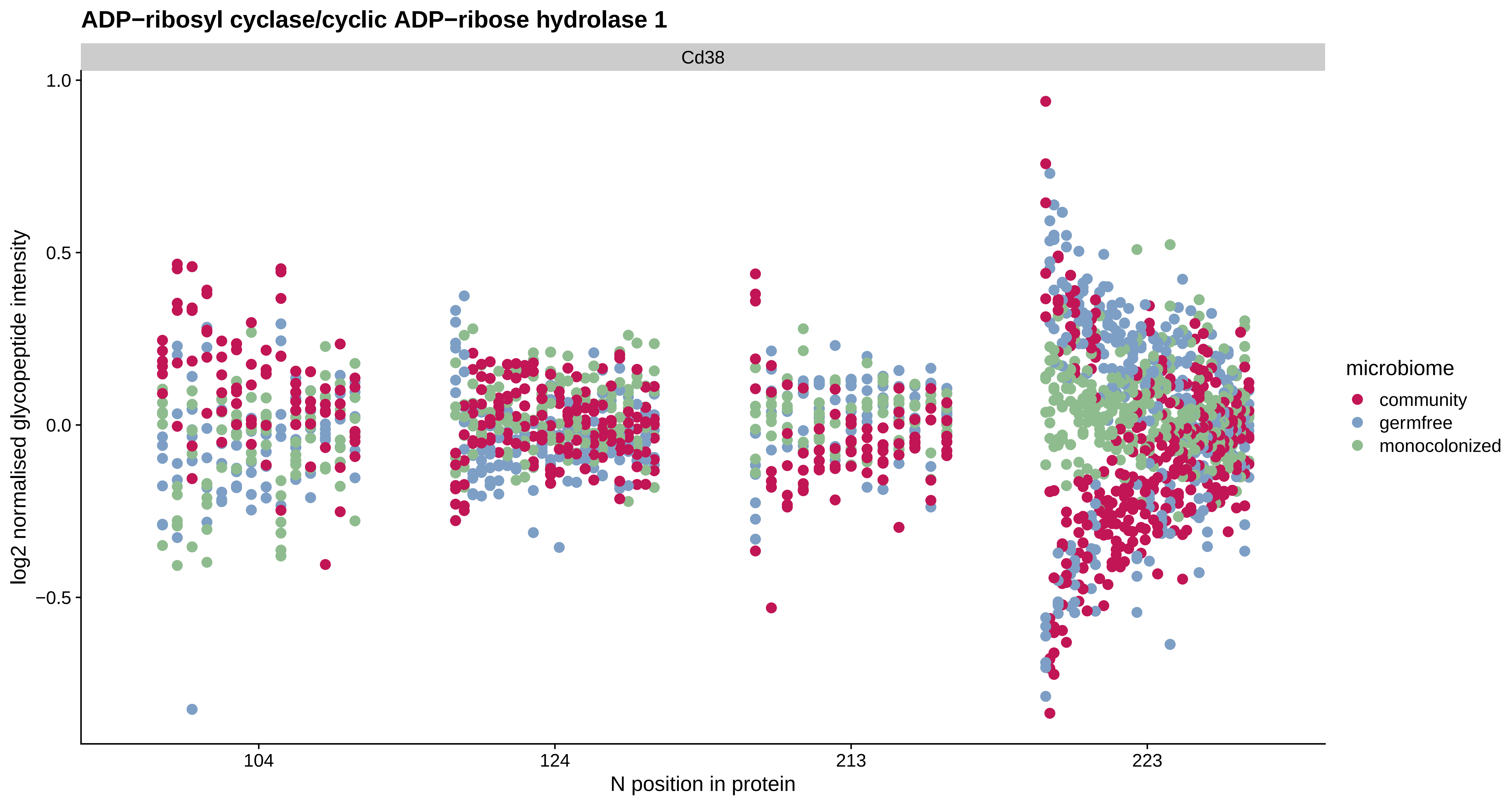

ADP−ribosyl cyclase/cyclic ADP−ribose hydrolase 1
Cd38
1.0
0.5
0.0
−0.5
log2 normalised glycopeptide intensity
104
124
213
223
N position in protein
microbiome
community
germfree
monocolonized

#### Attractin

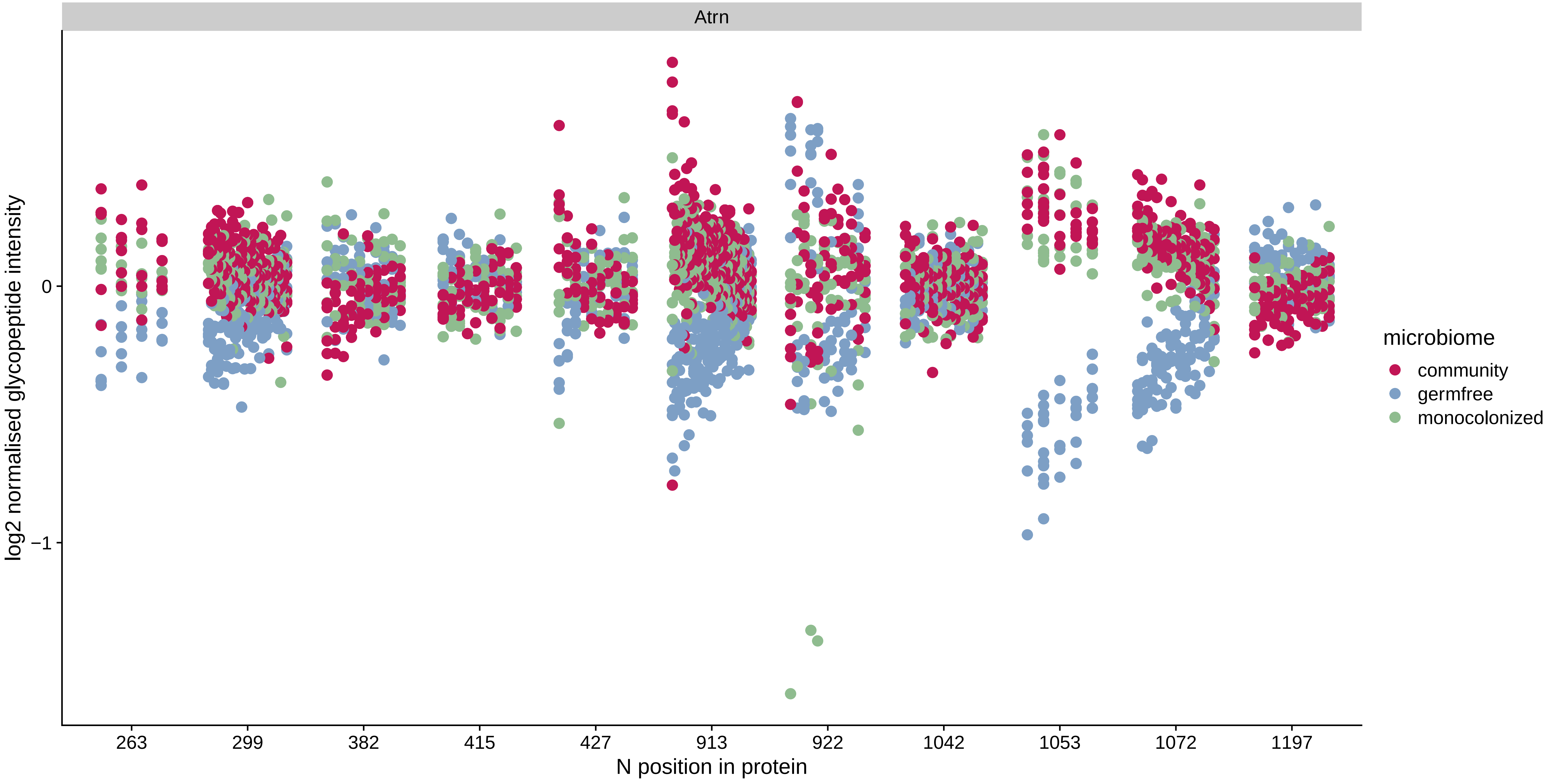

Atrn
log2 normalised glycopeptide intensity
0
−1
263
299
382
415
427
913
922
1042
1053
1072
1197
N position in protein
microbiome
community
germfree
monocolonized

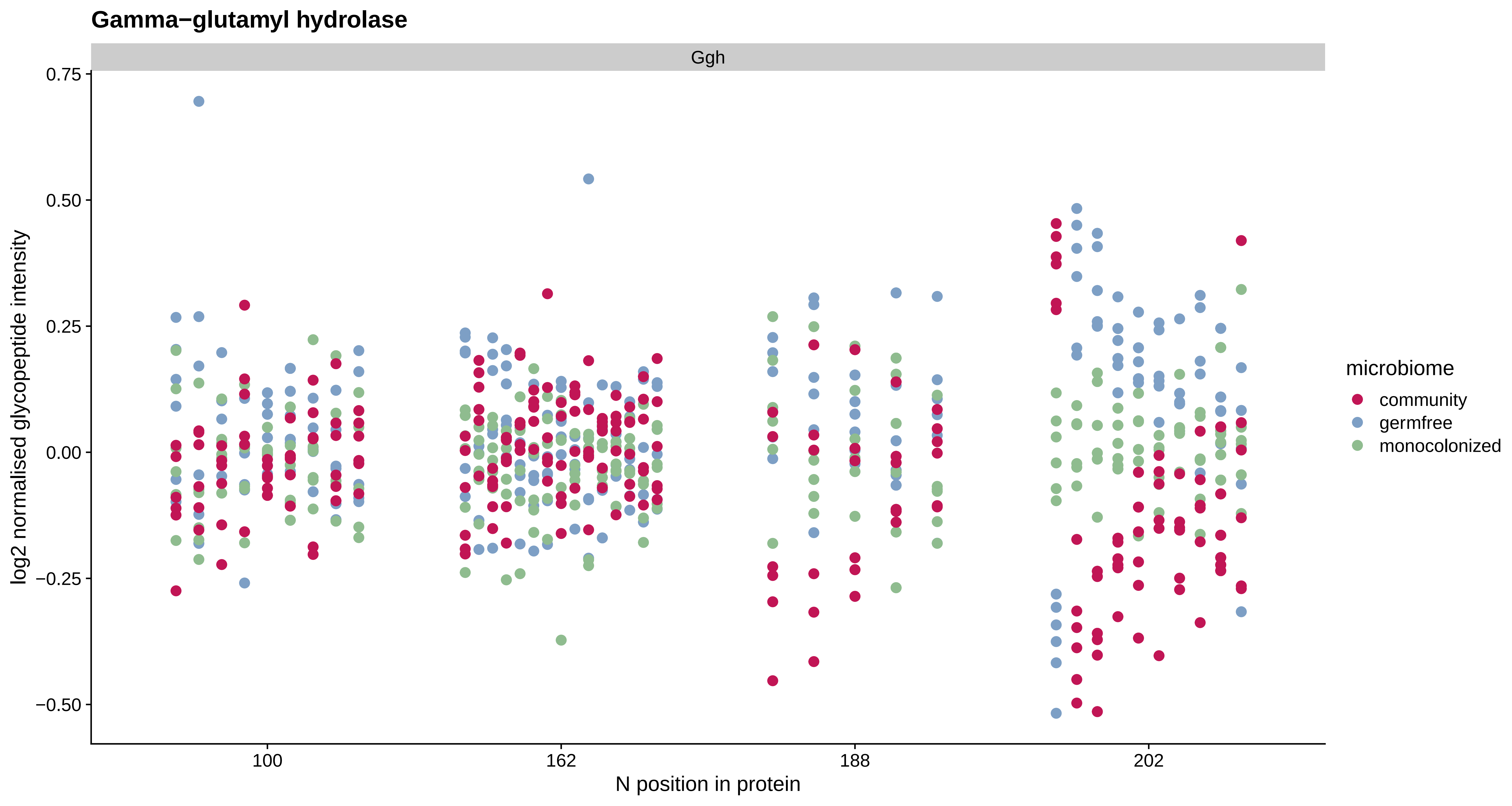

Gamma−glutamyl hydrolase
Ggh
0.75
0.50
0.25
0.00
−0.25
−0.50
log2 normalised glycopeptide intensity
100
162
188
202
N position in protein
microbiome
community
germfree
monocolonized

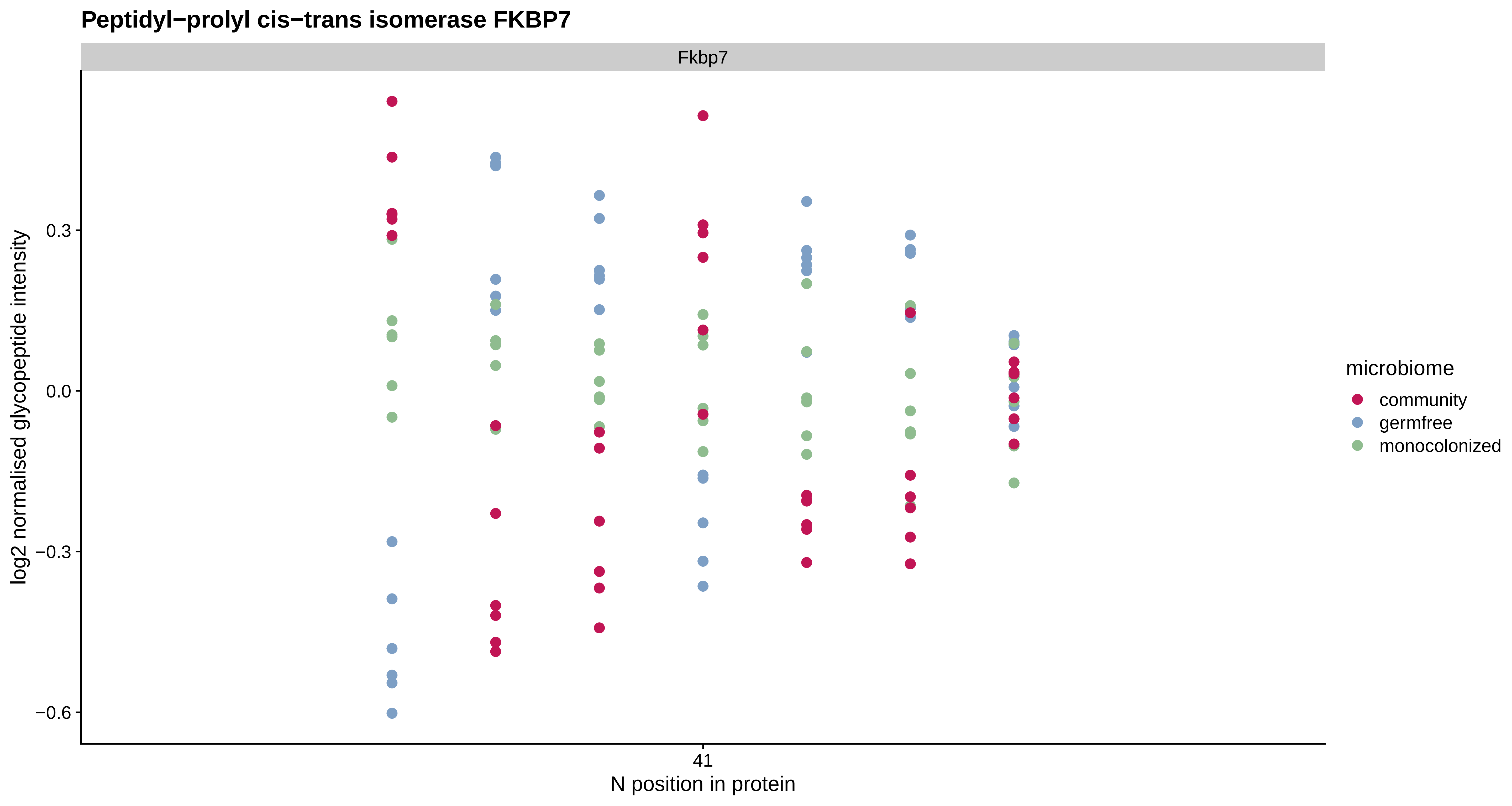
Peptidyl−prolyl cis−trans isomerase FKBP7
Fkbp7
log2 normalised glycopeptide intensity
0.3
0.0
−0.3
−0.6
41
N position in protein
microbiome
community
germfree
monocolonized

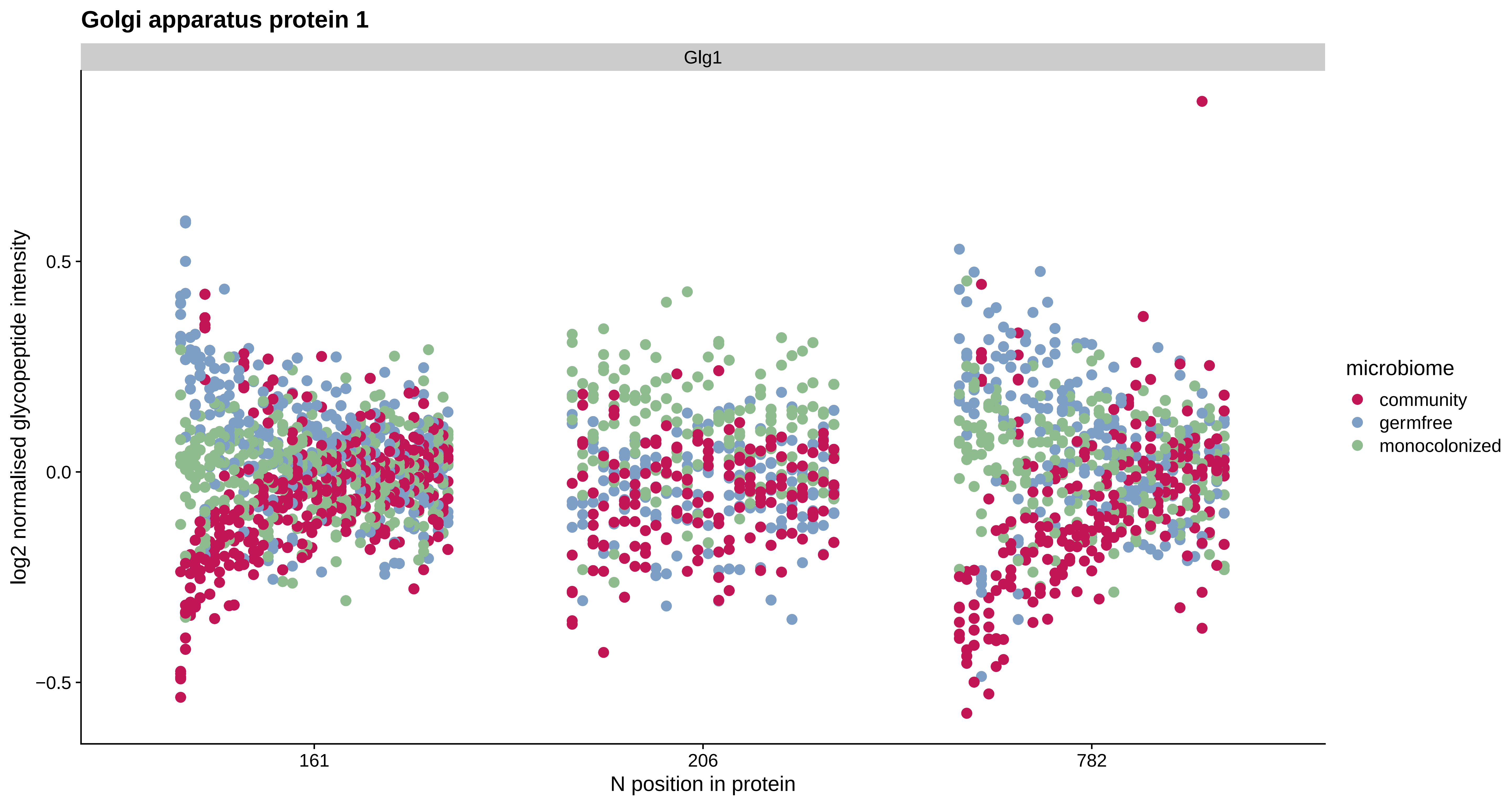

Golgi apparatus protein 1
Glg1
log2 normalised glycopeptide intensity
0.5
0.0
−0.5
161
206
782
N position in protein
microbiome
community
germfree
monocolonized

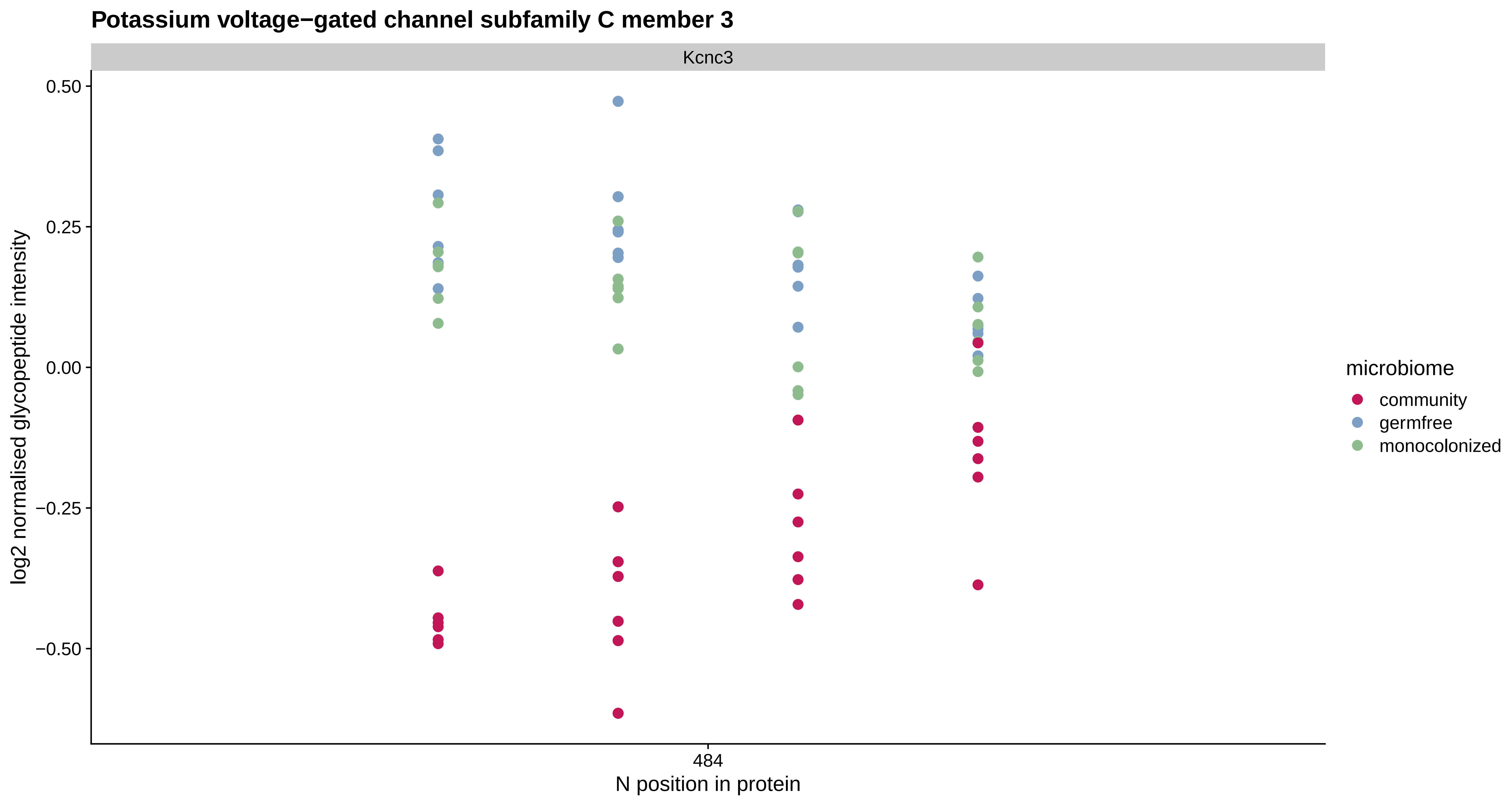
Potassium voltage−gated channel subfamily C member 3
Kcnc3
log2 normalised glycopeptide intensity
0.50
0.25
0.00
−0.25
−0.50
484
N position in protein
microbiome
community
germfree
monocolonized

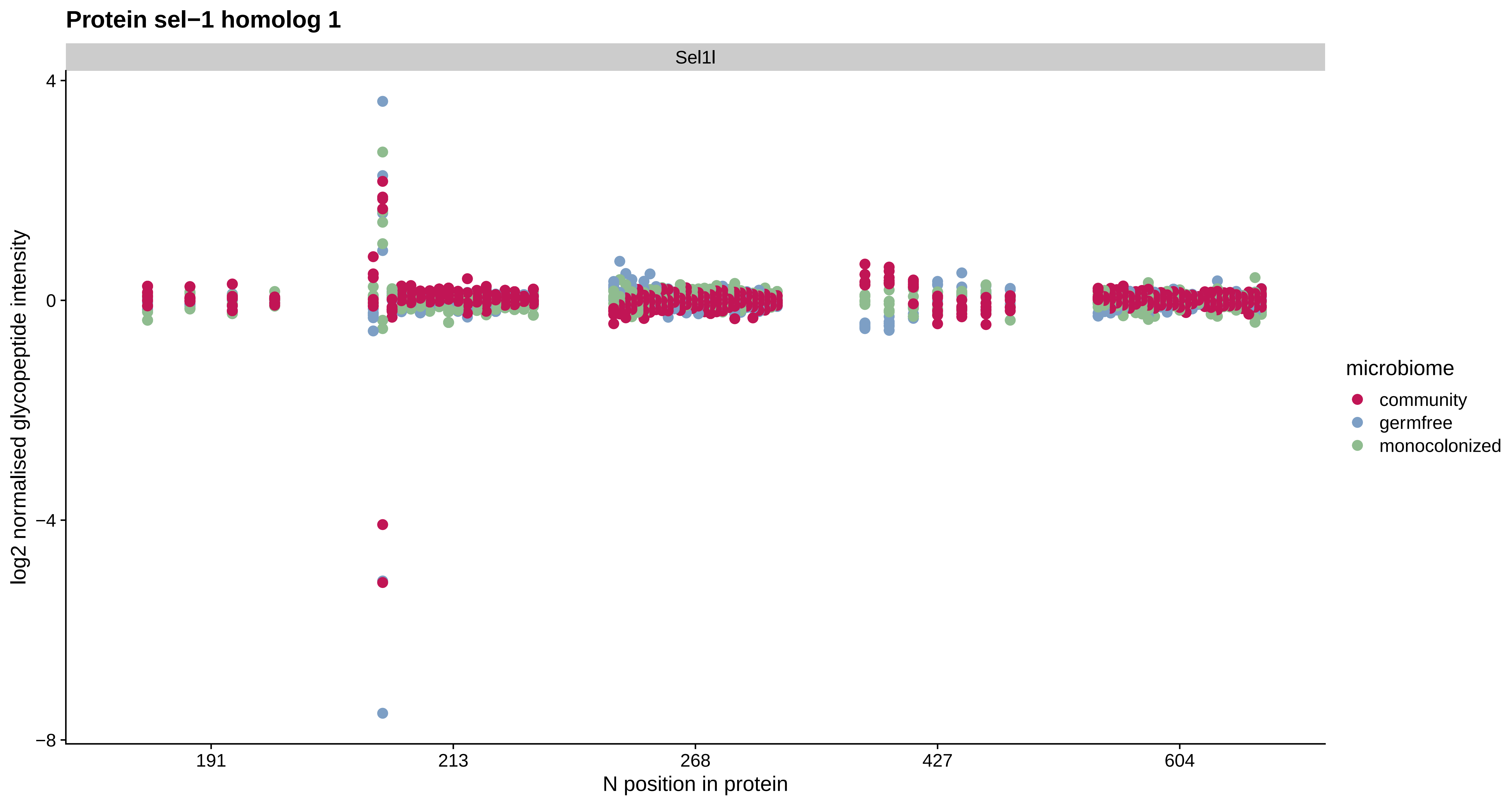

Protein sel−1 homolog 1
Sel1l
4
0
−4
−8
log2 normalised glycopeptide intensity
191
213
268
427
604
N position in protein
microbiome
community
germfree
monocolonized

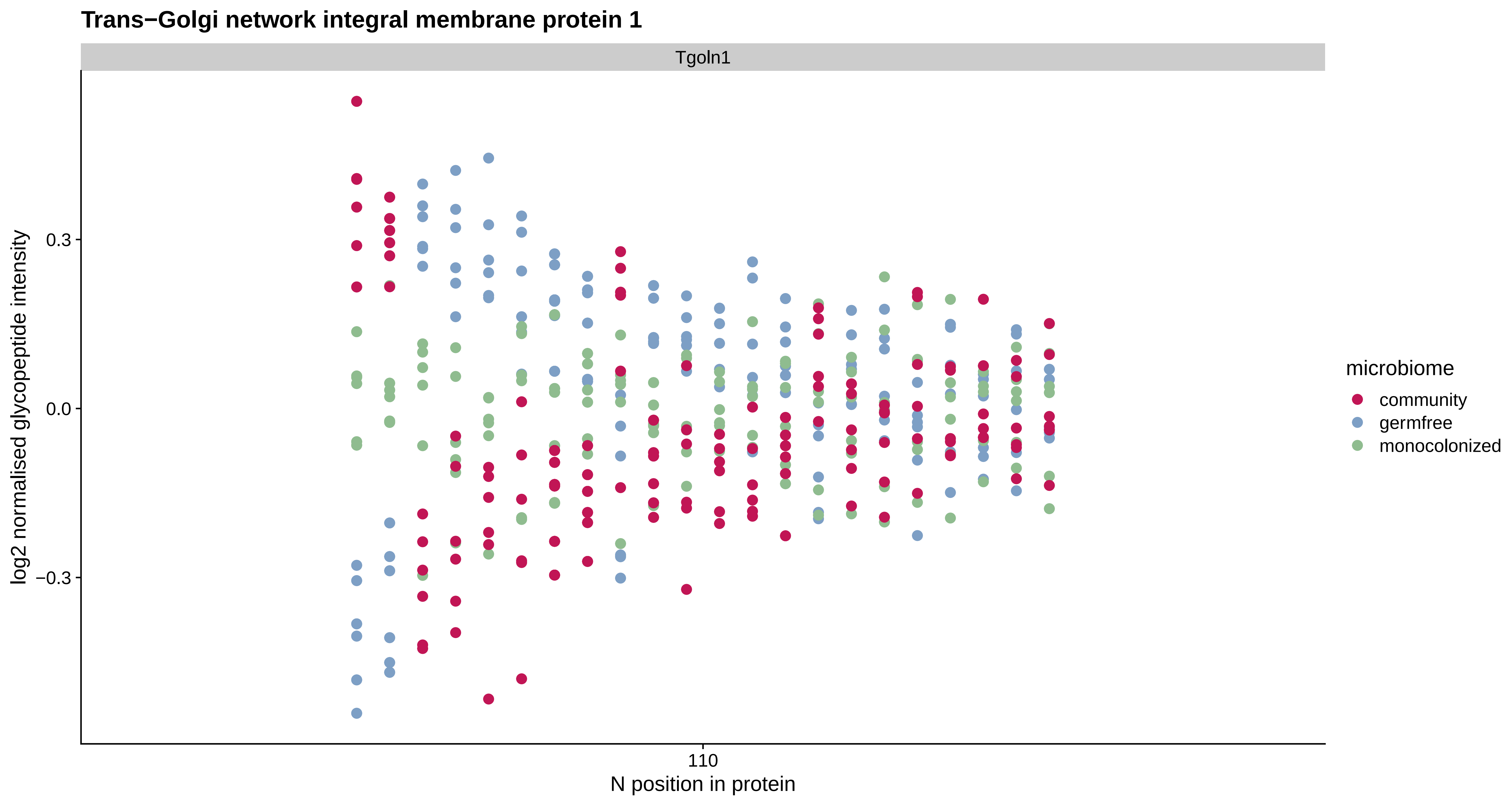

Trans−Golgi network integral membrane protein 1
Tgoln1
log2 normalised glycopeptide intensity
0.3
0.0
−0.3
110
N position in protein
microbiome
community
germfree
monocolonized

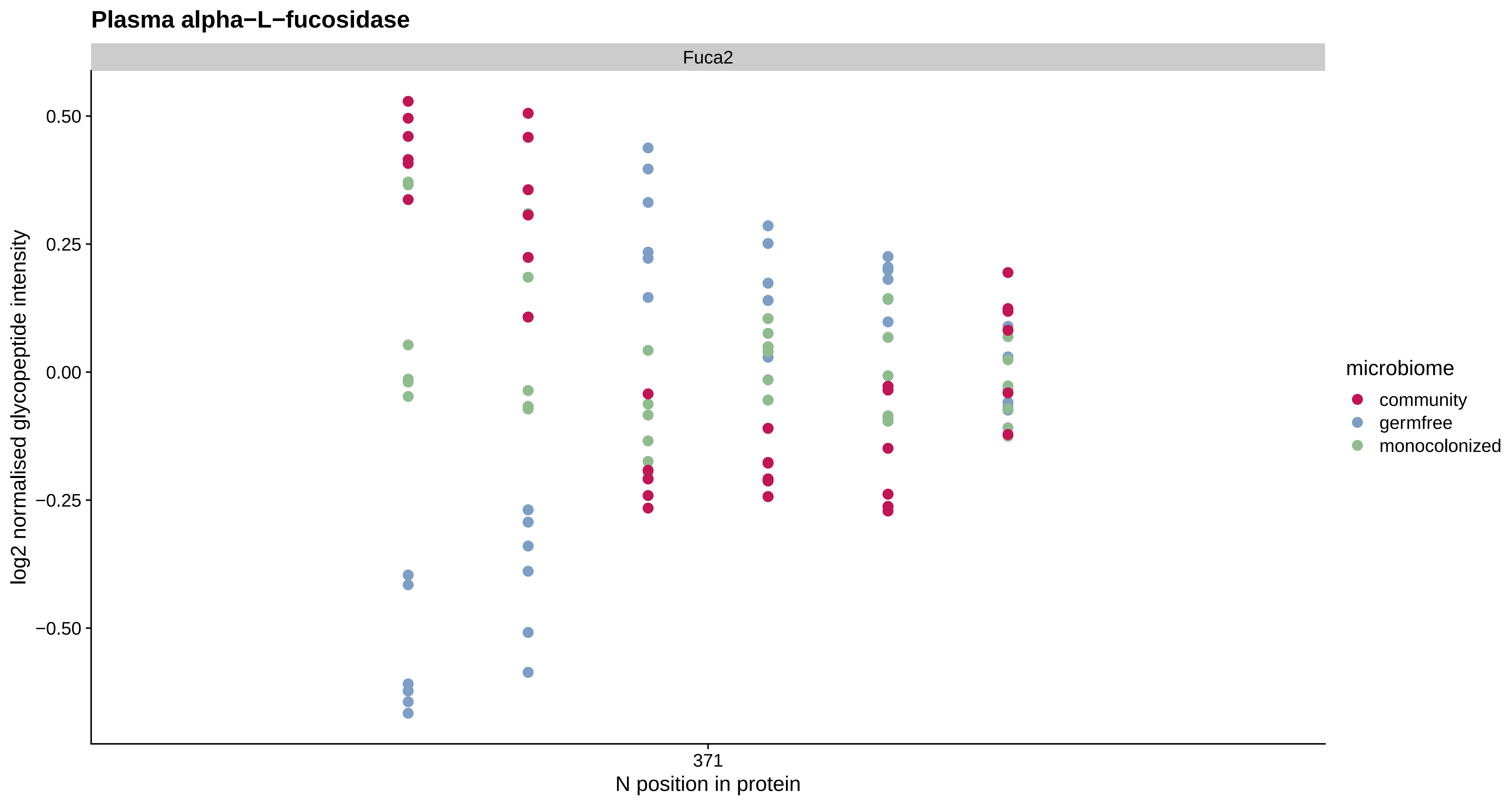

Plasma alpha−L−fucosidase
Fuca2
0.50
0.25
0.00
−0.25
−0.50
log2 normalised glycopeptide intensity
371
N position in protein
microbiome
community
germfree
monocolonized

**Protein HEG homolog 1**

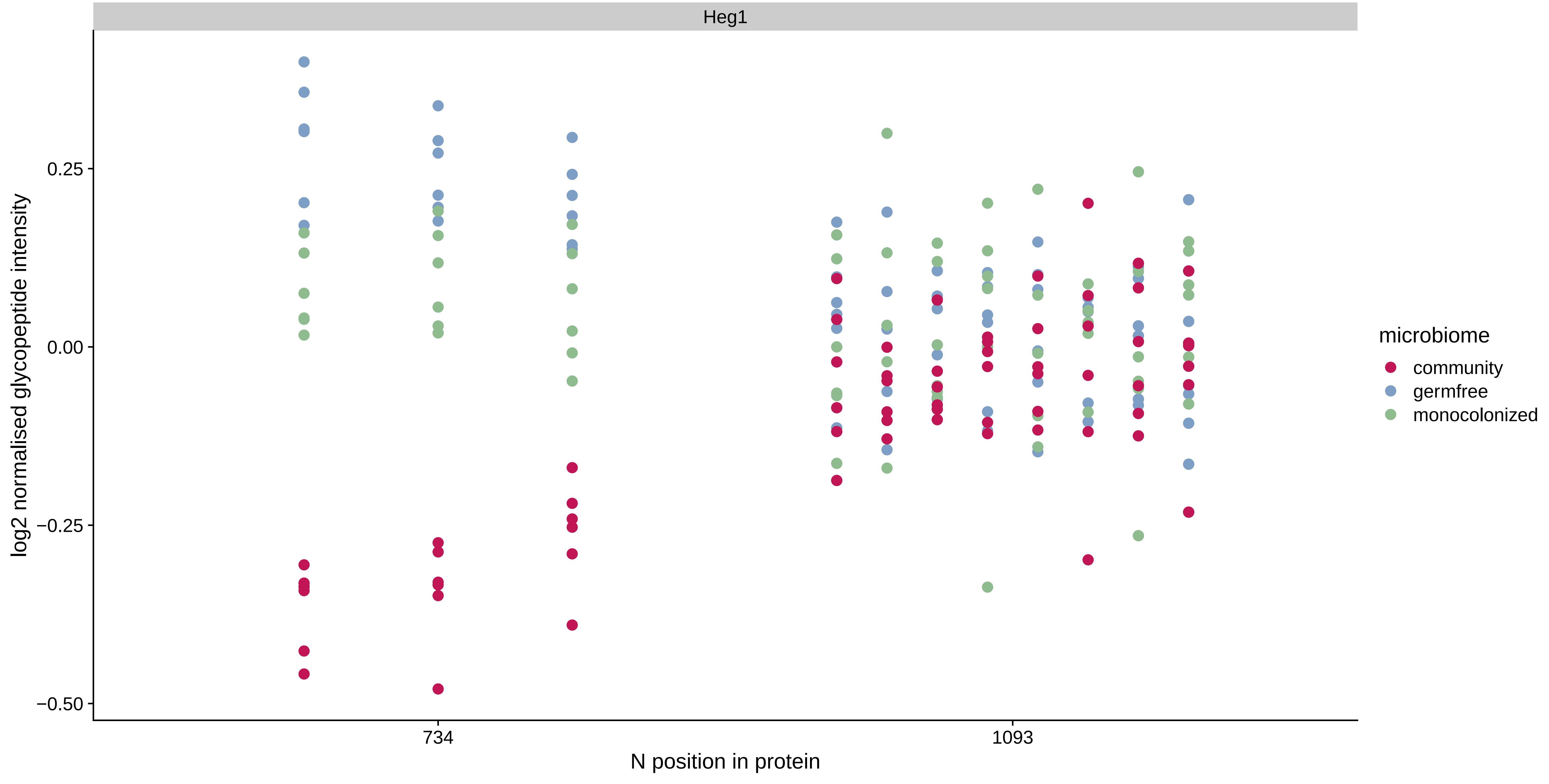

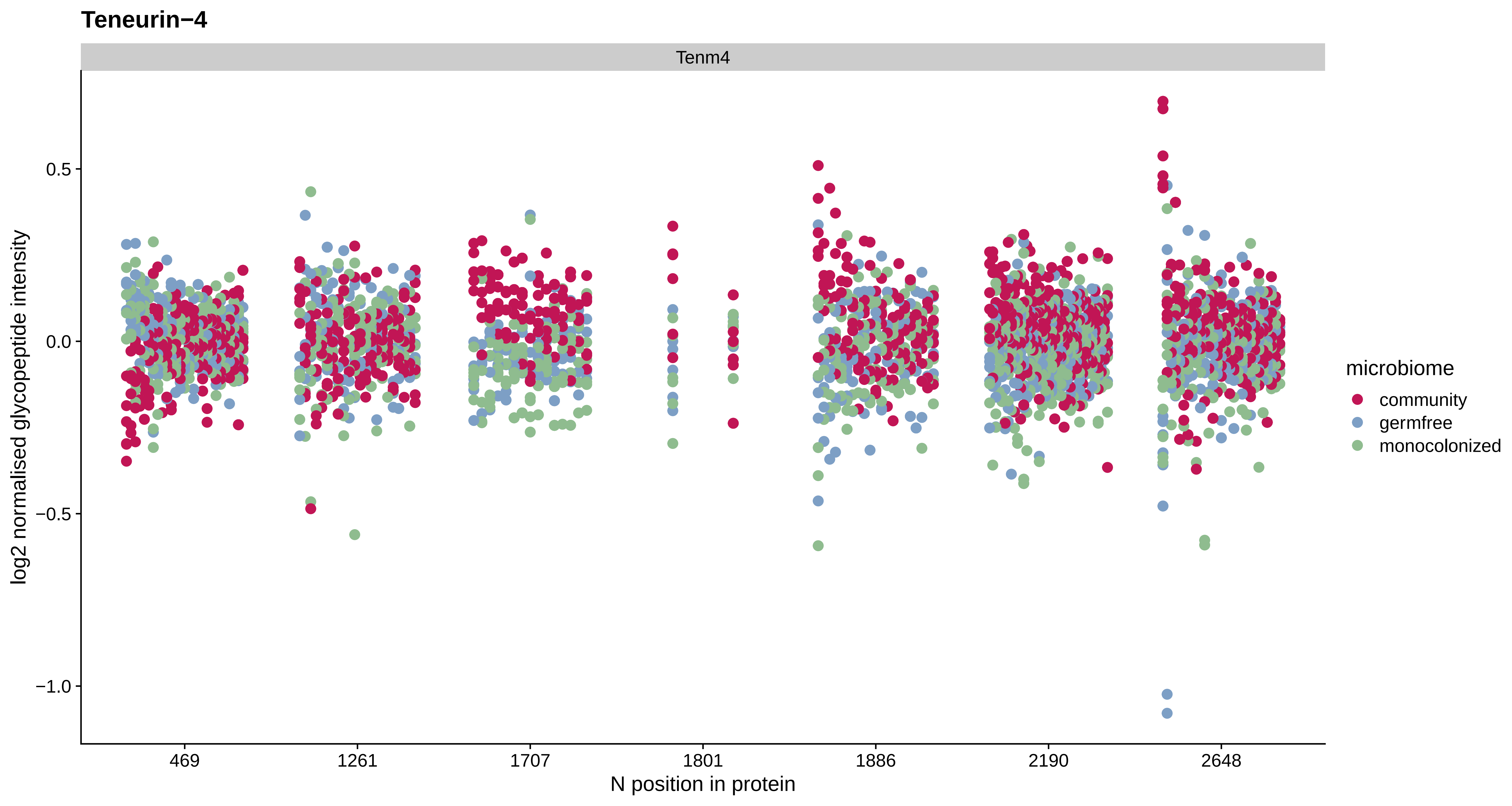

Teneurin−4
Tenm4
0.5
0.0
−0.5
−1.0
log2 normalised glycopeptide intensity
469
1261
1707
1801
1886
2190
2648
N position in protein
microbiome
community
germfree
monocolonized

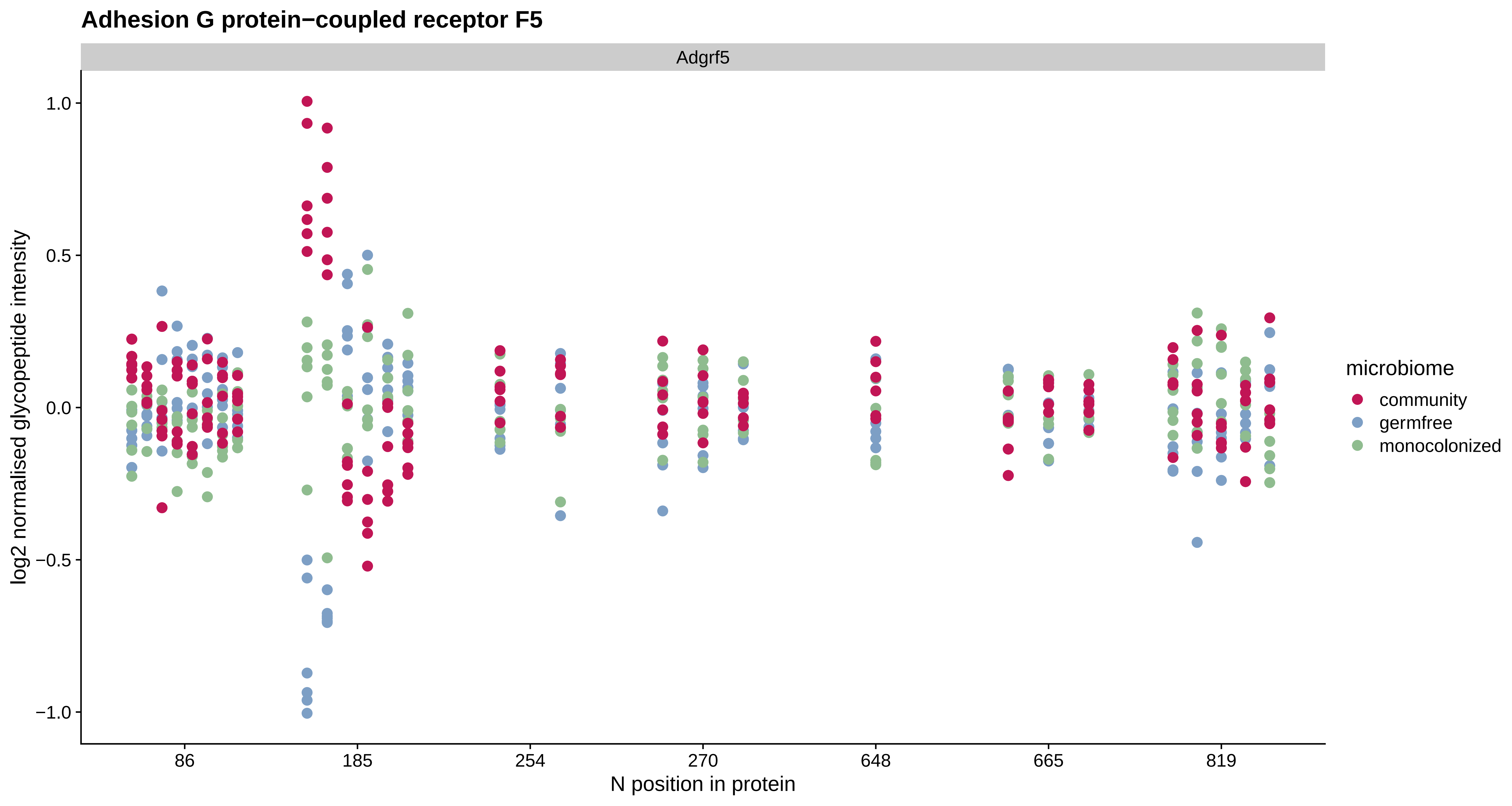
Adhesion G protein−coupled receptor F5
Adgrf5
log2 normalised glycopeptide intensity
1.0
0.5
0.0
−0.5
−1.0
86
185
254
270
648
665
819
N position in protein
microbiome
community
germfree
monocolonized

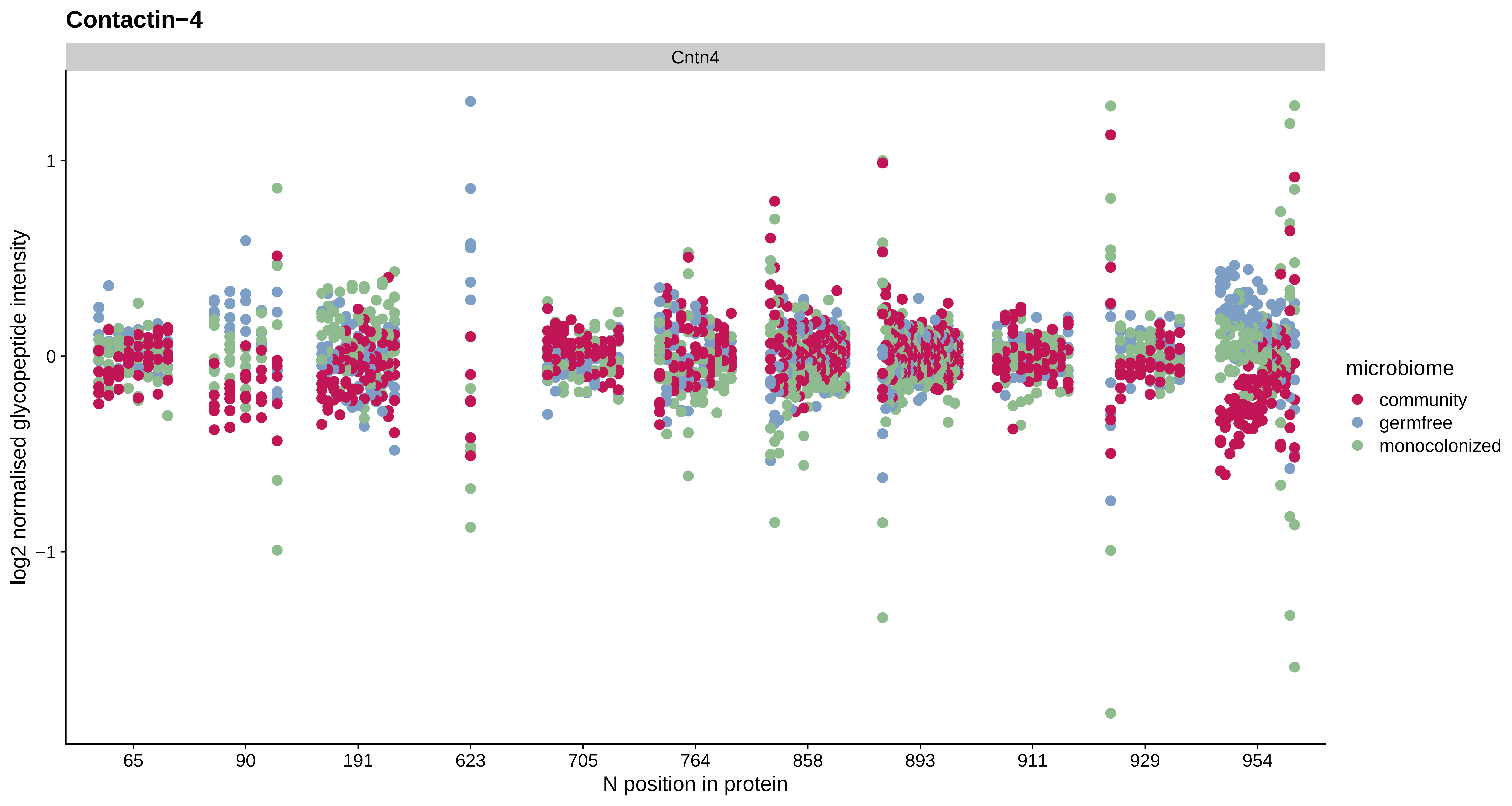
Contactin−4
Cntn4
log2 normalised glycopeptide intensity
1
0
−1
65
90
191
623
705
764
858
893
911
929
954
N position in protein
microbiome
community
germfree
monocolonized

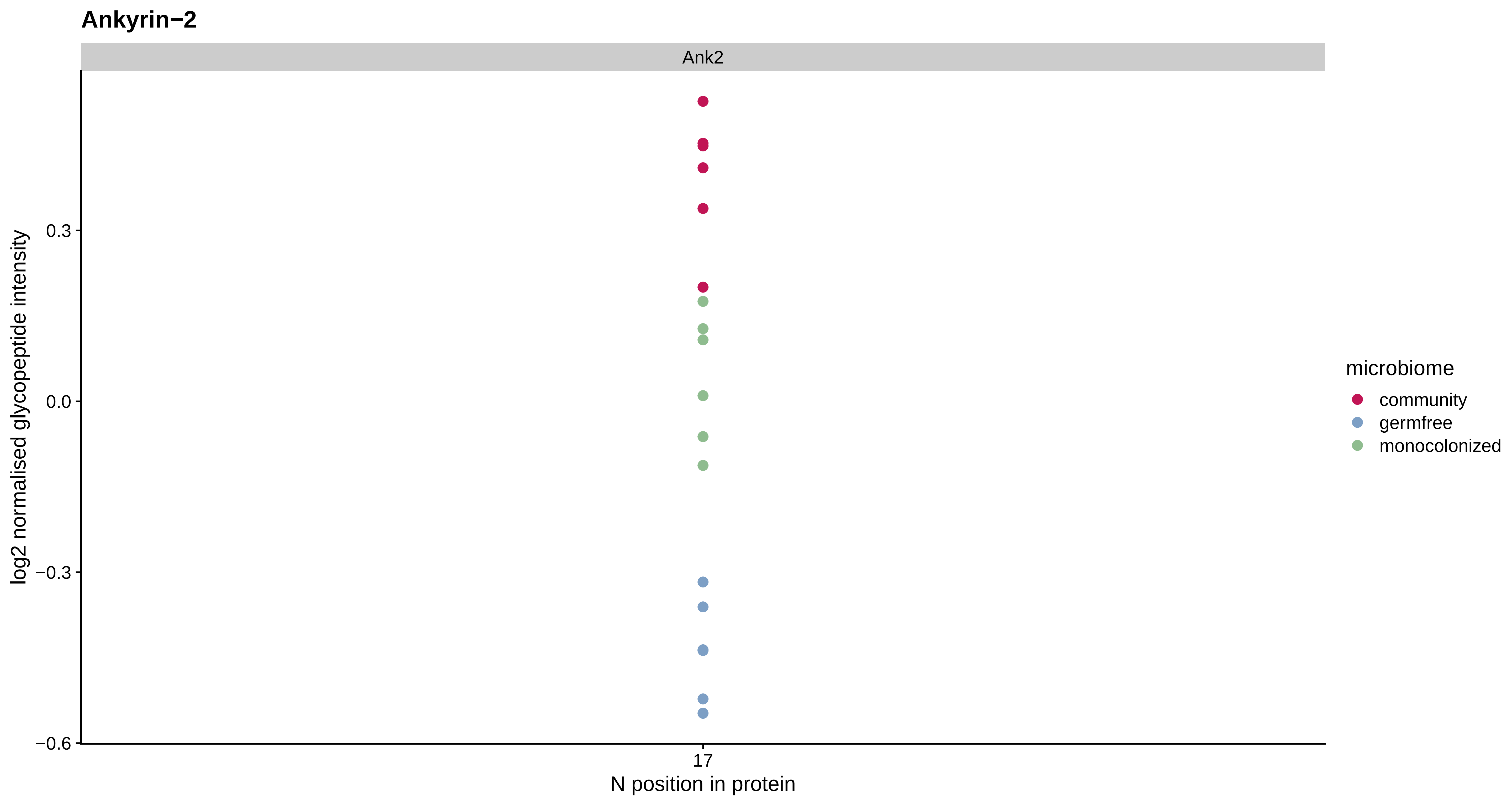
Ankyrin−2
Ank2
log2 normalised glycopeptide intensity
0.3
0.0
−0.3
−0.6
17
N position in protein
microbiome
community
germfree
monocolonized

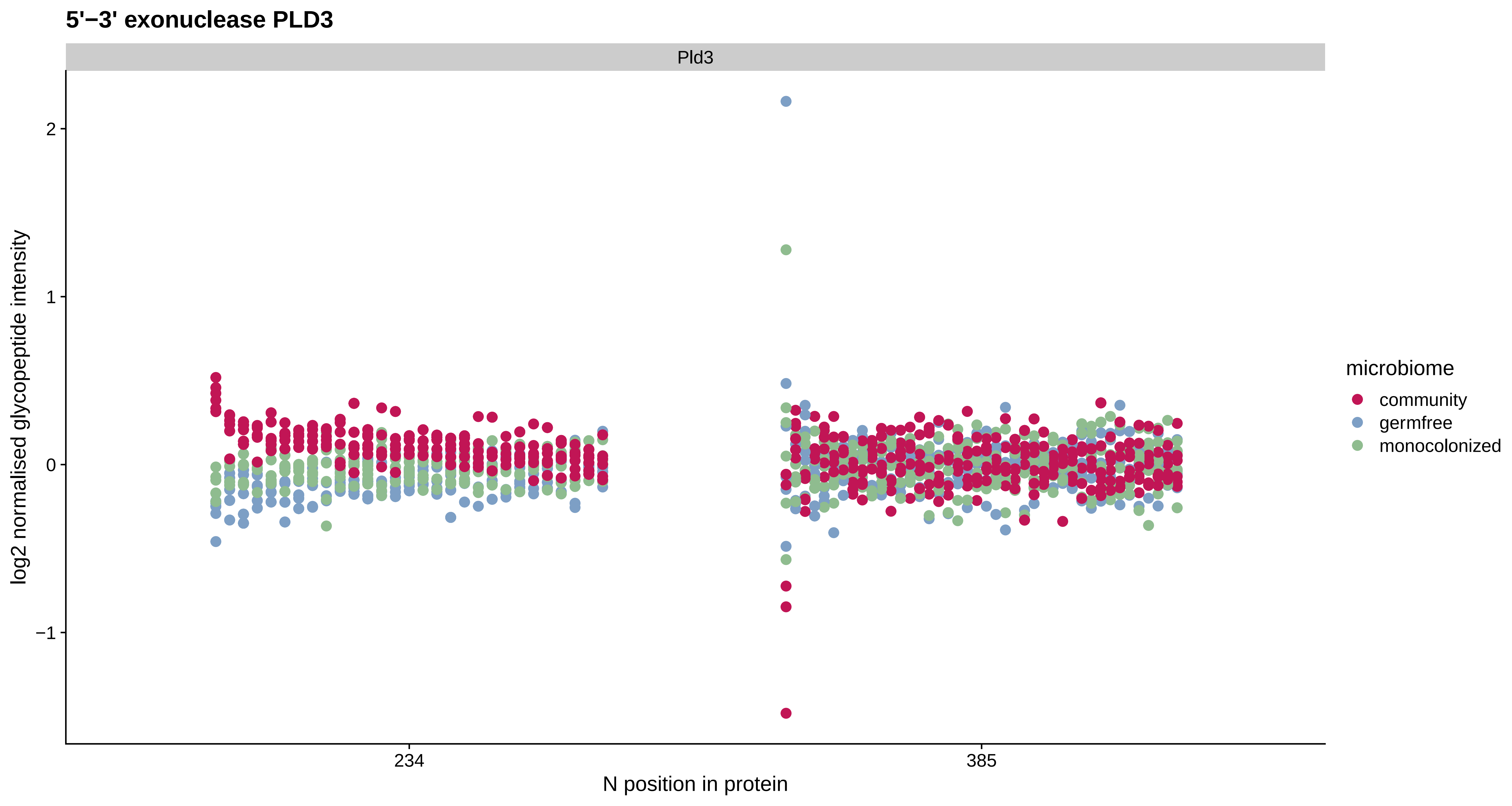
5'−3' exonuclease PLD3
Pld3
log2 normalised glycopeptide intensity
2
1
0
−1
234
385
N position in protein
microbiome
community
germfree
monocolonized

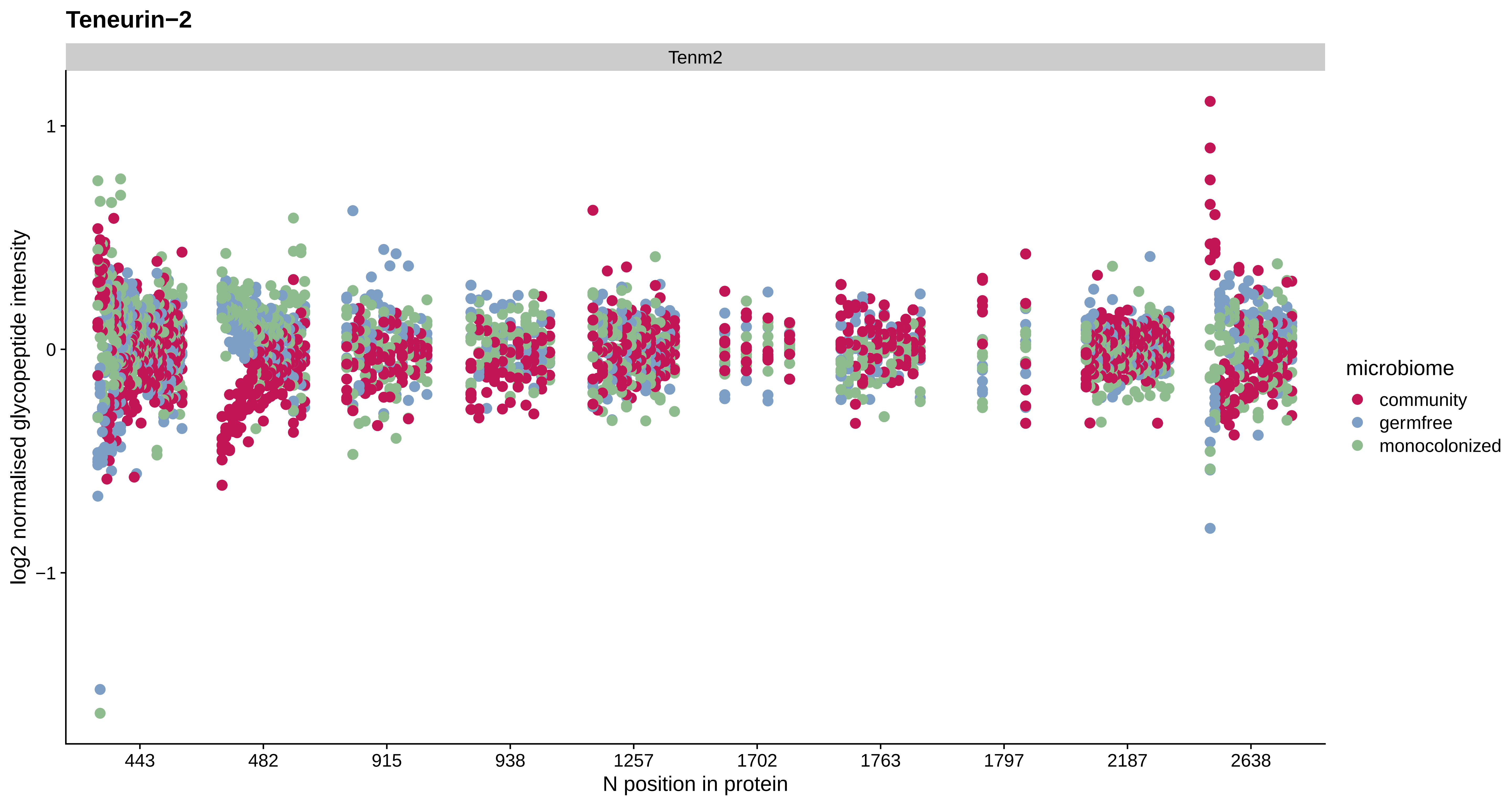
Teneurin−2
Tenm2
log2 normalised glycopeptide intensity
1
0
−1
443
482
915
938
1257
1702
1763
1797
2187
2638
N position in protein
microbiome
community
germfree
monocolonized

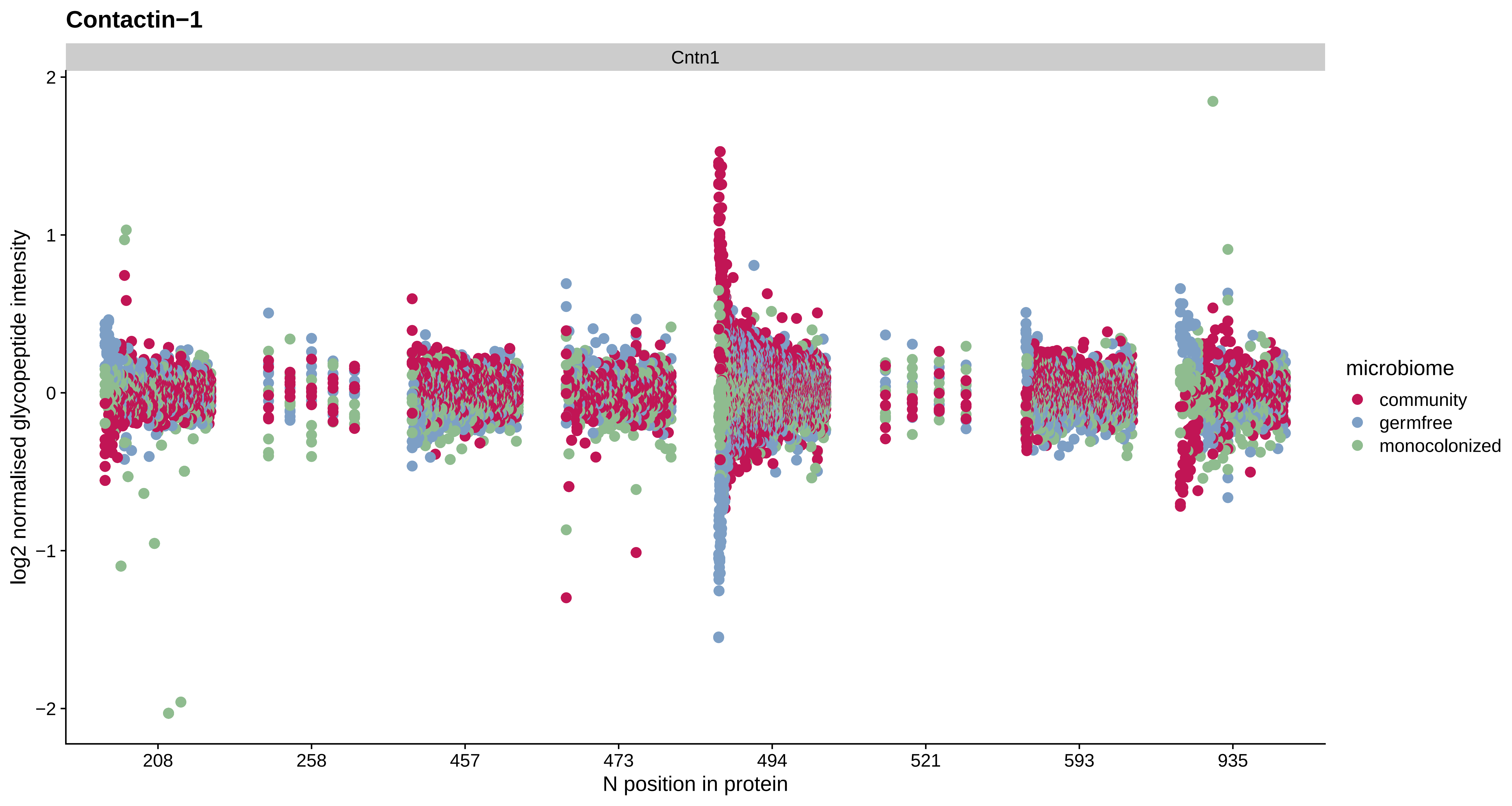
Contactin−1
Cntn1
log2 normalised glycopeptide intensity
2
1
0
−1
−2
208
258
457
473
494
521
593
935
N position in protein
microbiome
community
germfree
monocolonized

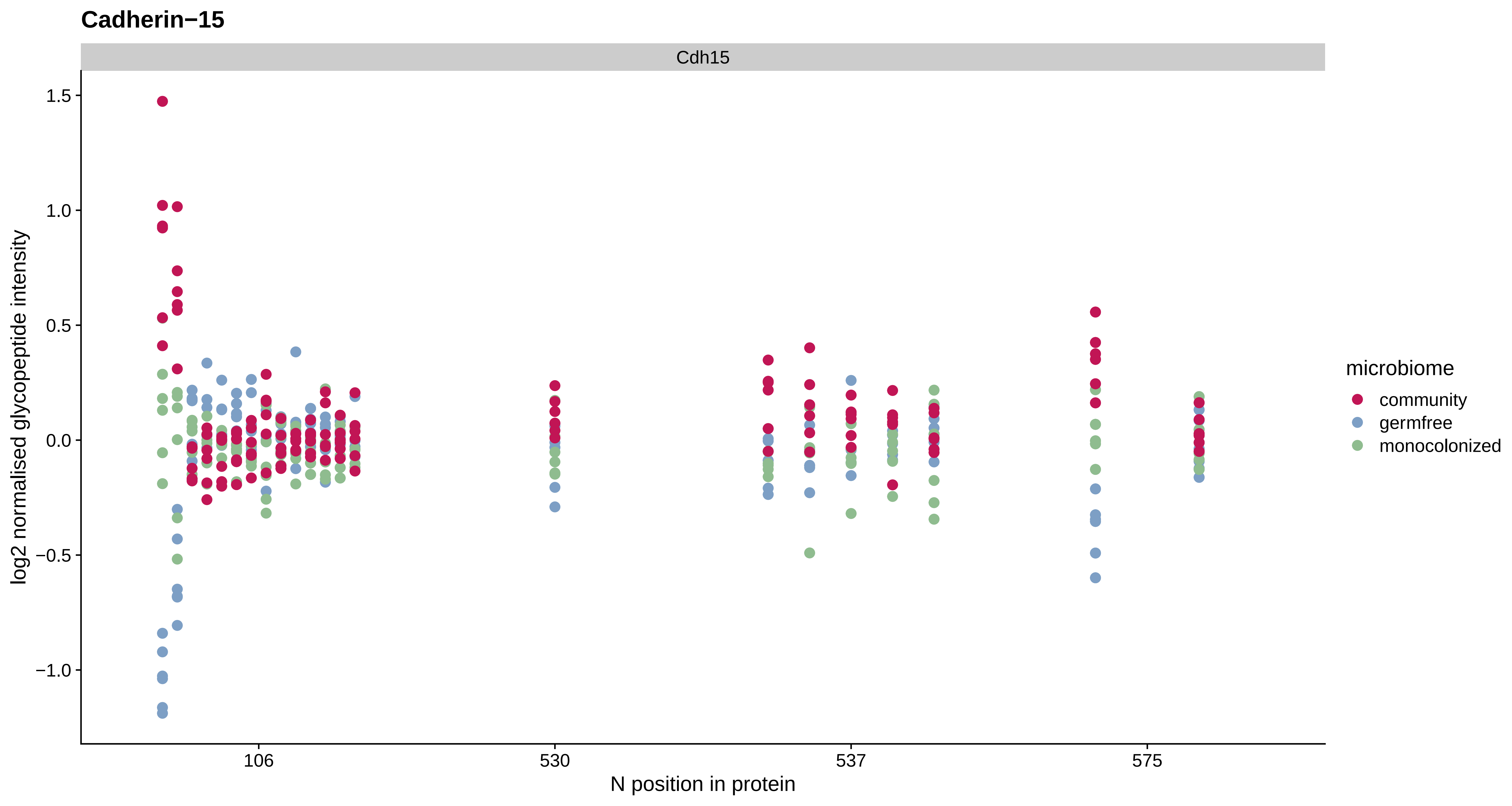
Cadherin−15
Cdh15
log2 normalised glycopeptide intensity
1.5
1.0
0.5
0.0
−0.5
−1.0
106
530
537
575
N position in protein
microbiome
community
germfree
monocolonized

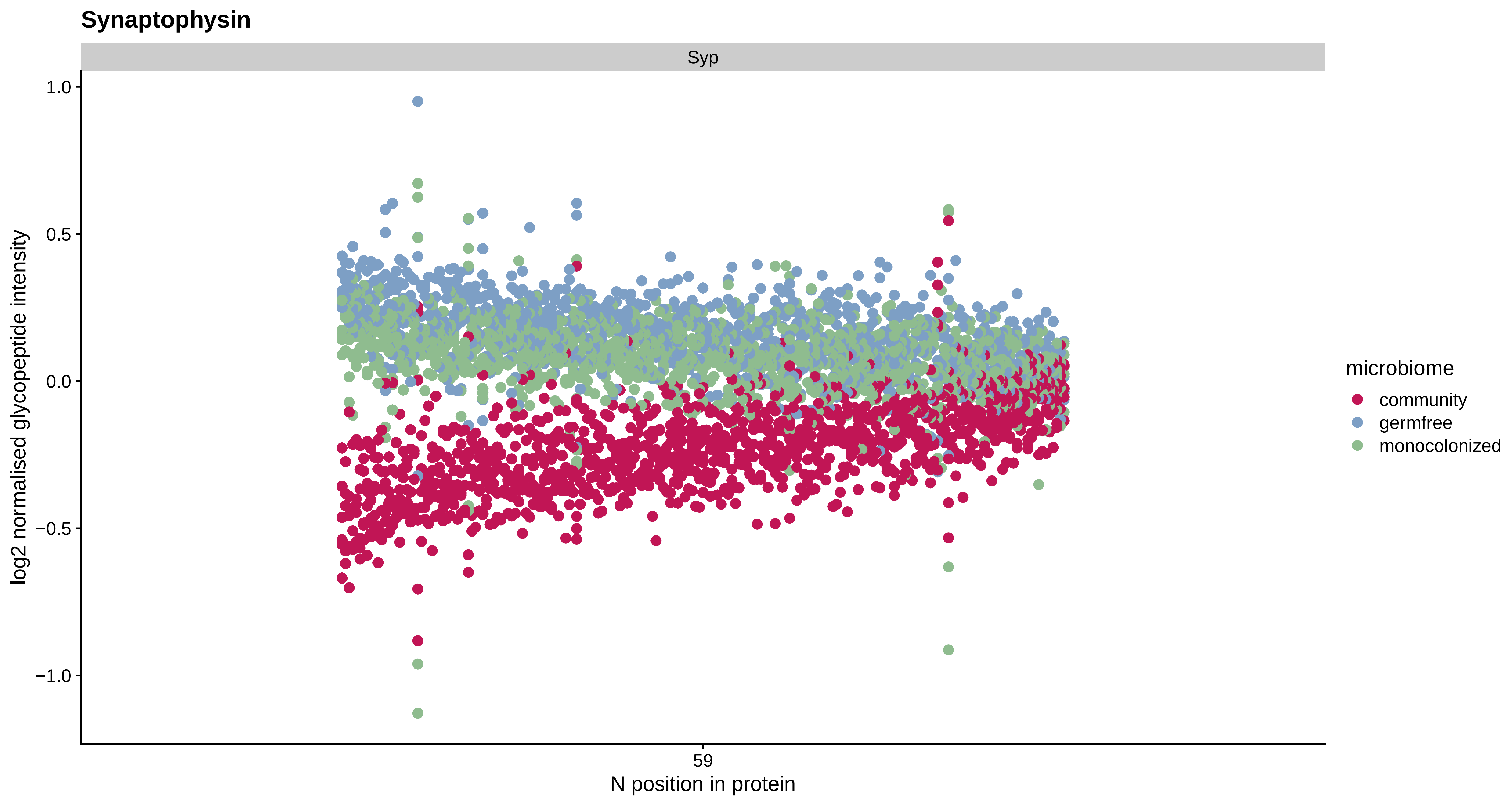

Synaptophysin
Syp
1.0
0.5
0.0
−0.5
−1.0
log2 normalised glycopeptide intensity
59
N position in protein
microbiome
community
germfree
monocolonized

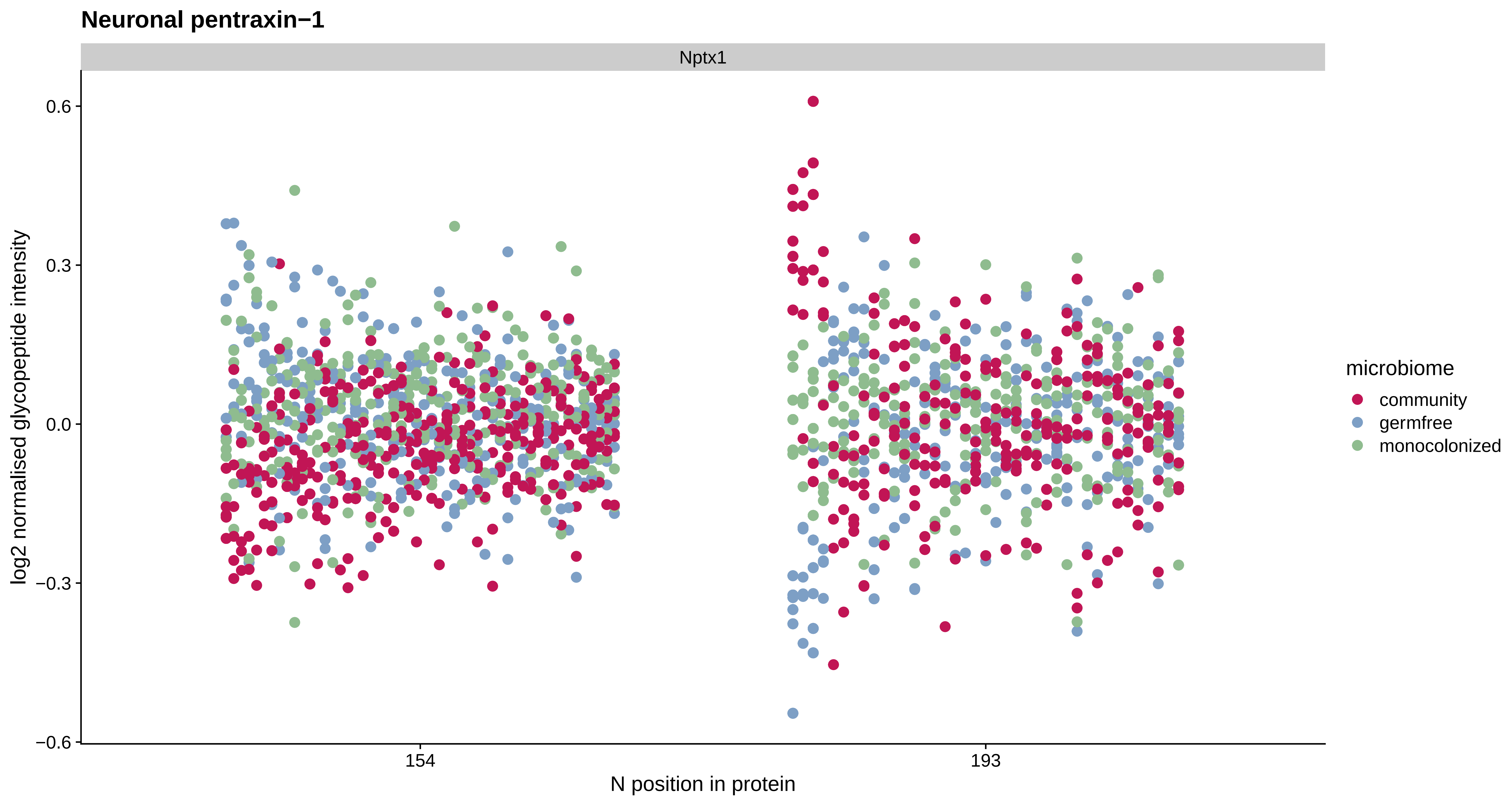

Neuronal pentraxin−1
Nptx1
log2 normalised glycopeptide intensity
0.6
0.3
0.0
−0.3
−0.6
154
193
N position in protein
microbiome
community
germfree
monocolonized

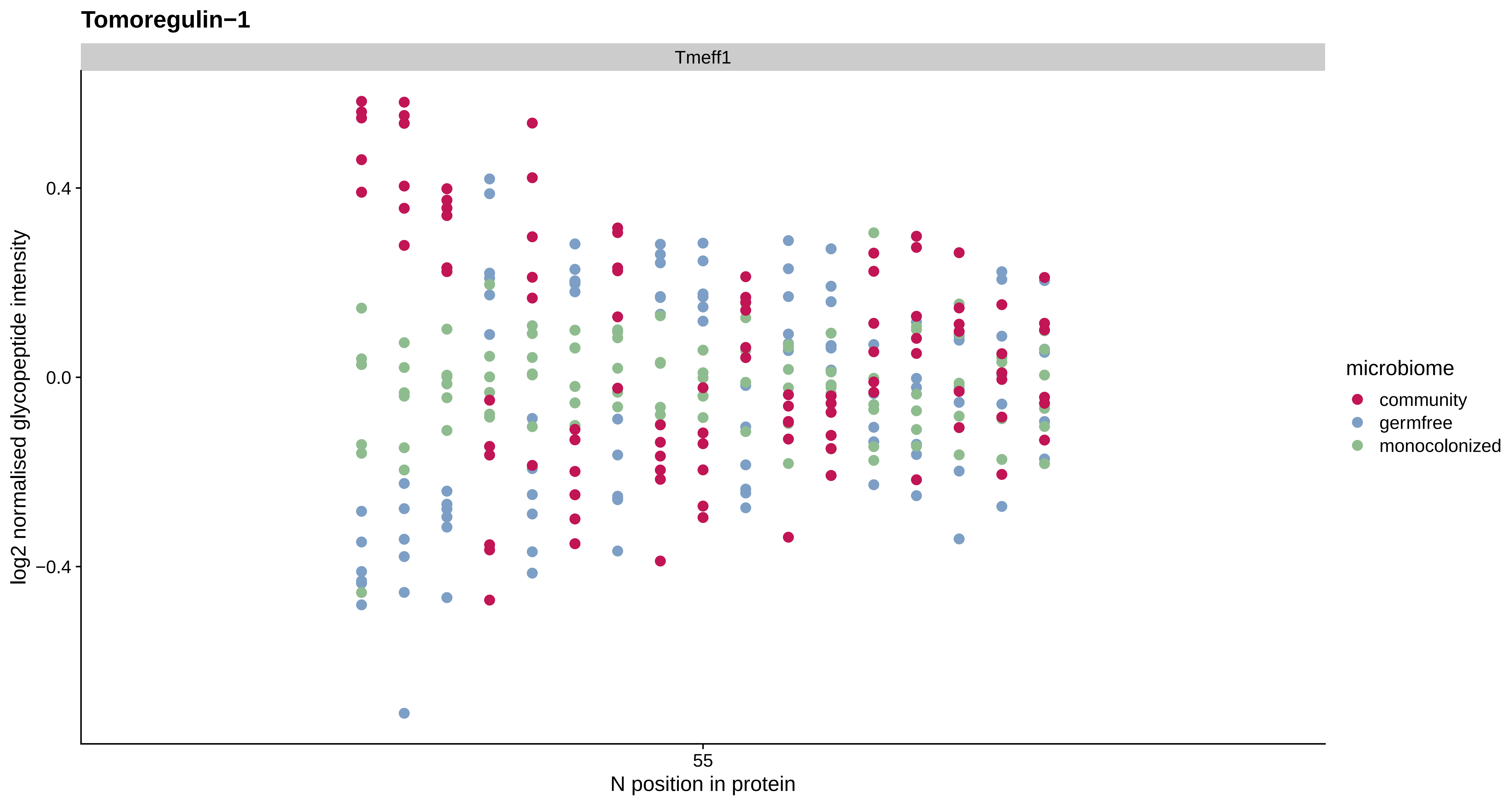
Tomoregulin−1
Tmeff1
log2 normalised glycopeptide intensity
0.4
0.0
−0.4
55
N position in protein
microbiome
community
germfree
monocolonized

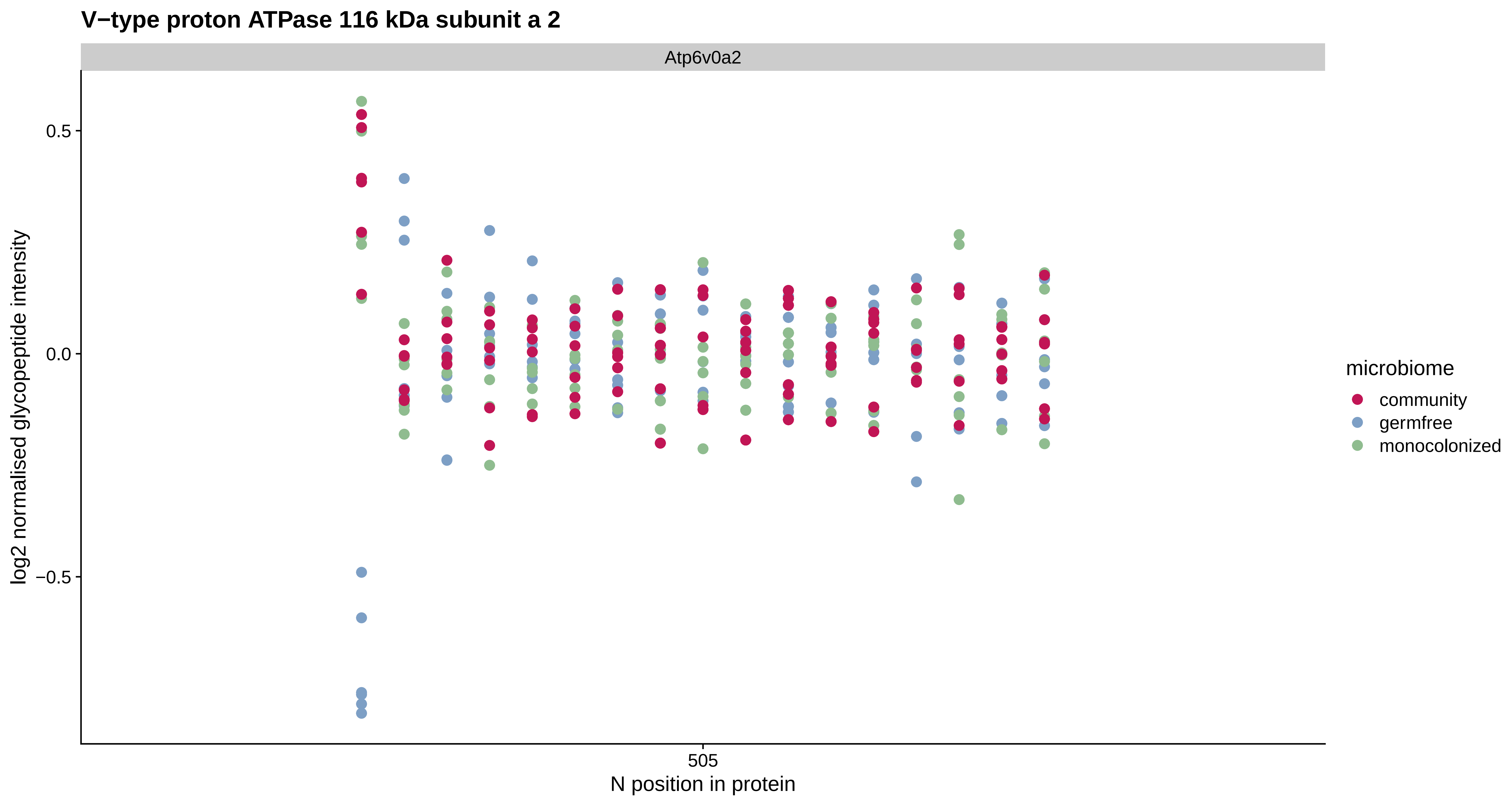
V−type proton ATPase 116 kDa subunit a 2
Atp6v0a2
log2 normalised glycopeptide intensity
0.5
0.0
−0.5
505
N position in protein
microbiome
community
germfree
monocolonized

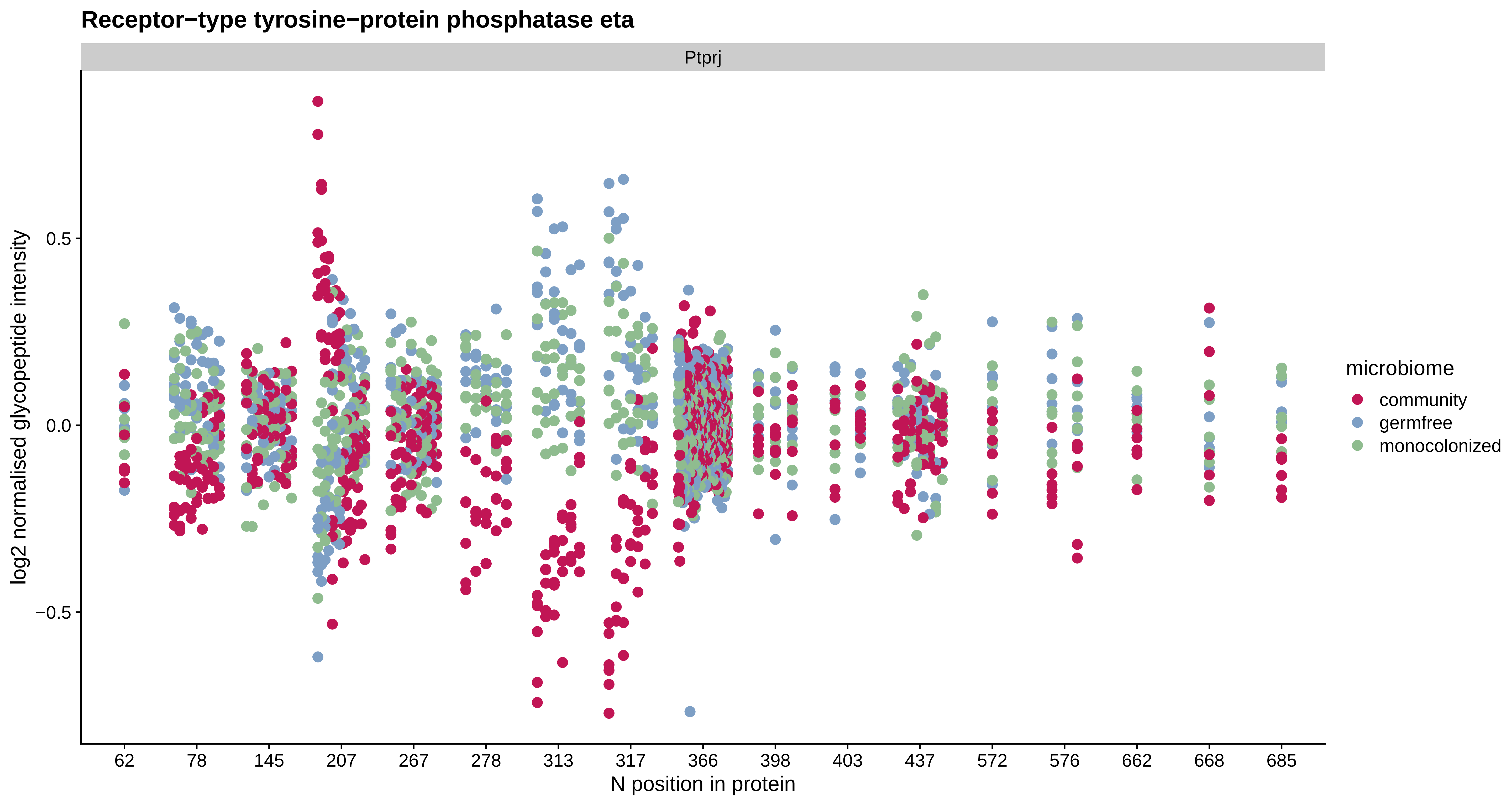
Receptor−type tyrosine−protein phosphatase eta
Ptprj
log2 normalised glycopeptide intensity
0.5
0.0
−0.5
62
78
145
207
267
278
313
317
366
398
403
437
572
576
662
668
685
N position in protein
microbiome
community
germfree
monocolonized

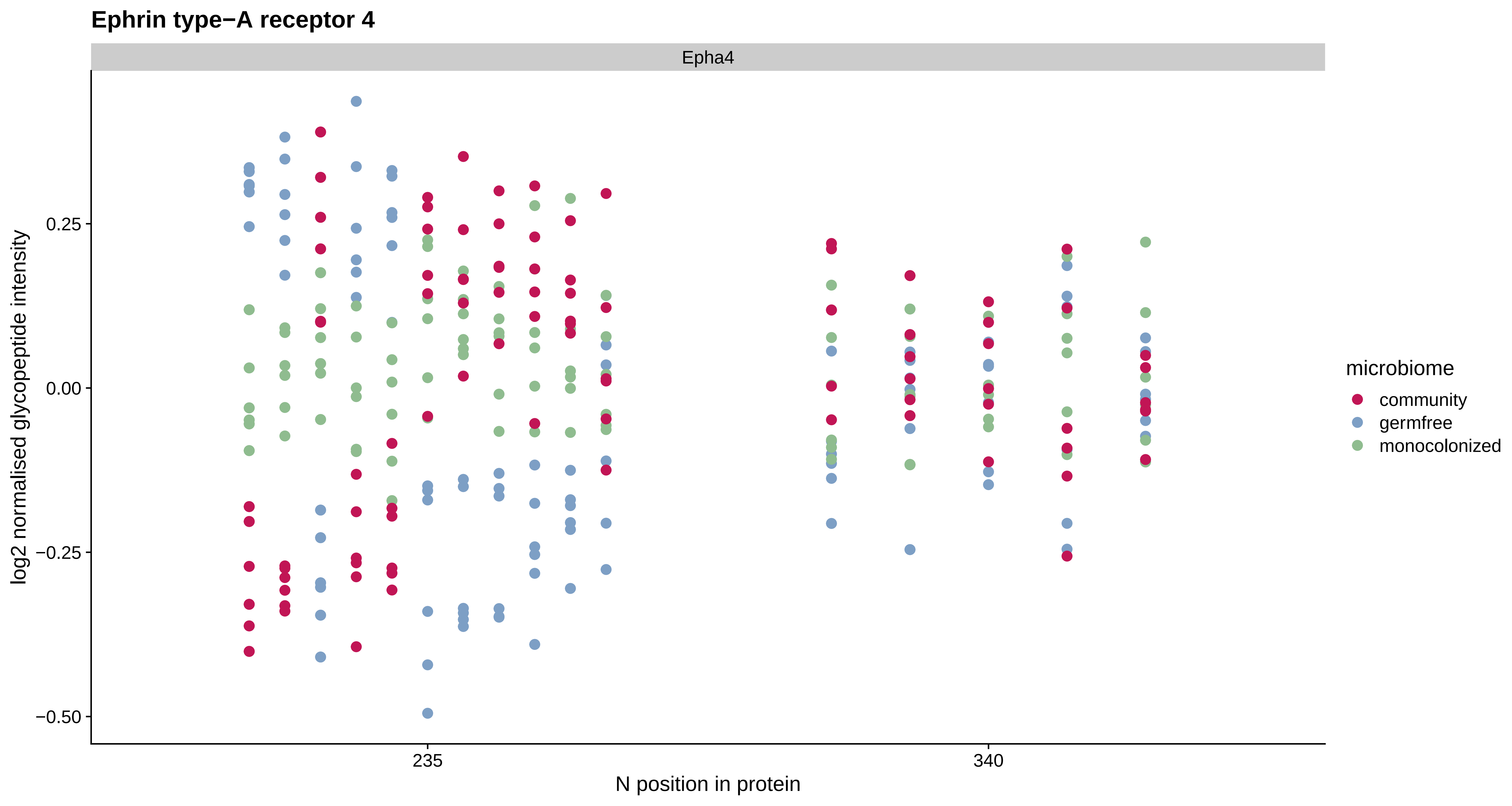
Ephrin type−A receptor 4
Epha4
log2 normalised glycopeptide intensity
0.25
0.00
−0.25
−0.50
235
340
N position in protein
microbiome
community
germfree
monocolonized

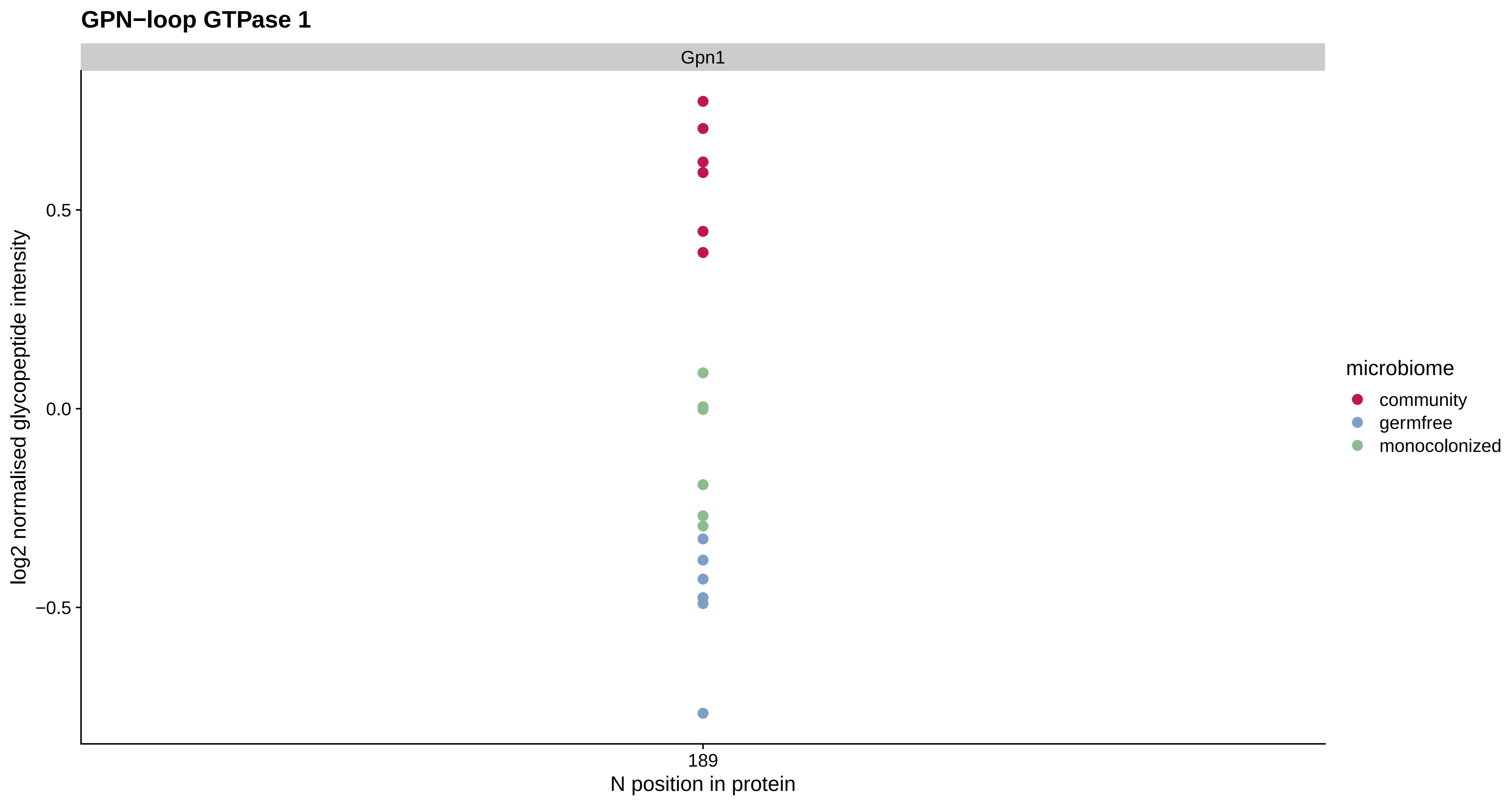
GPN−loop GTPase 1
Gpn1
0.5
0.0
−0.5
log2 normalised glycopeptide intensity
189
N position in protein
microbiome
community
germfree
monocolonized

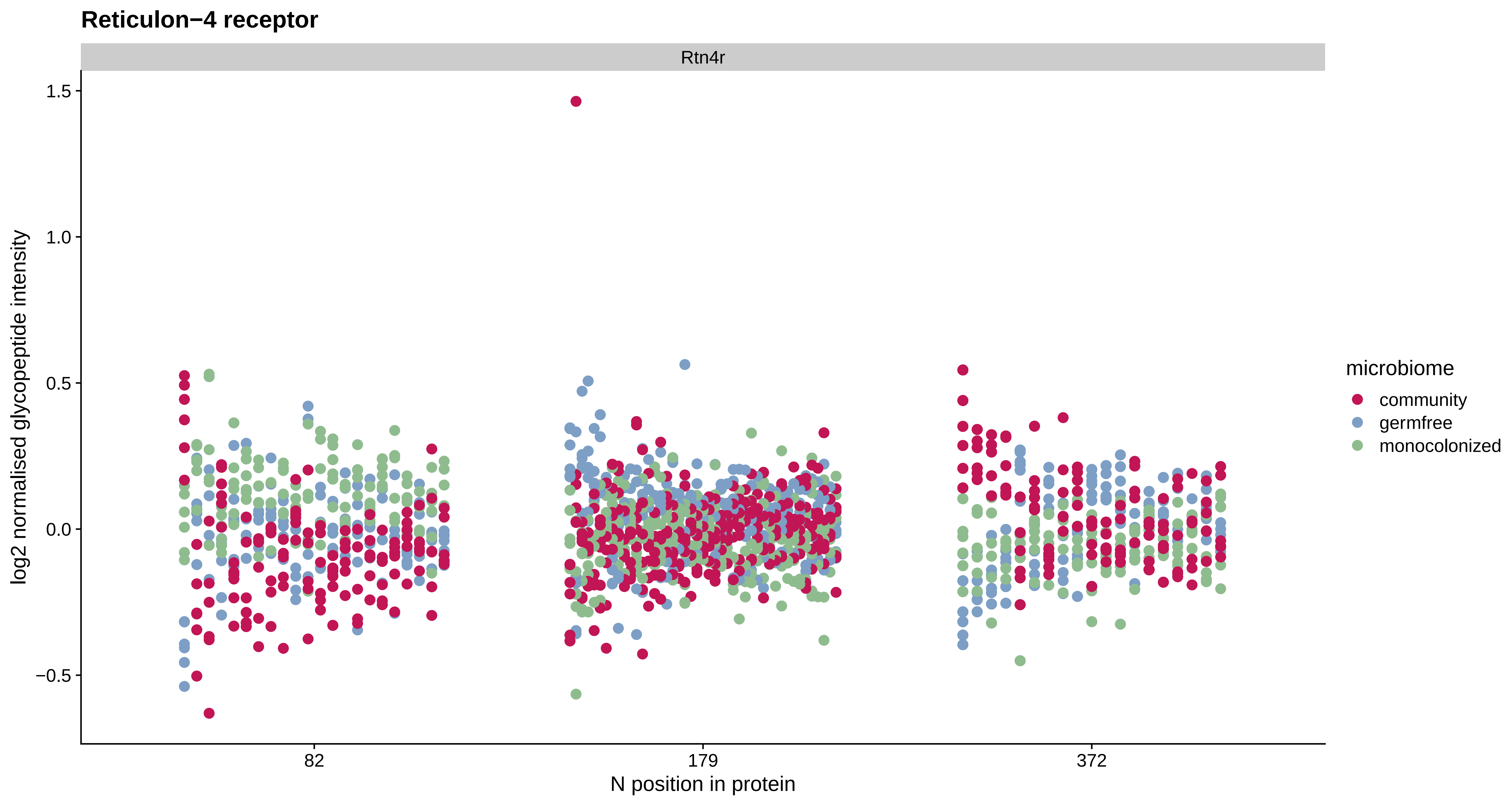
Reticulon−4 receptor
Rtn4r
1.5
1.0
0.5
0.0
−0.5
log2 normalised glycopeptide intensity
82
179
372
N position in protein
microbiome
community
germfree
monocolonized

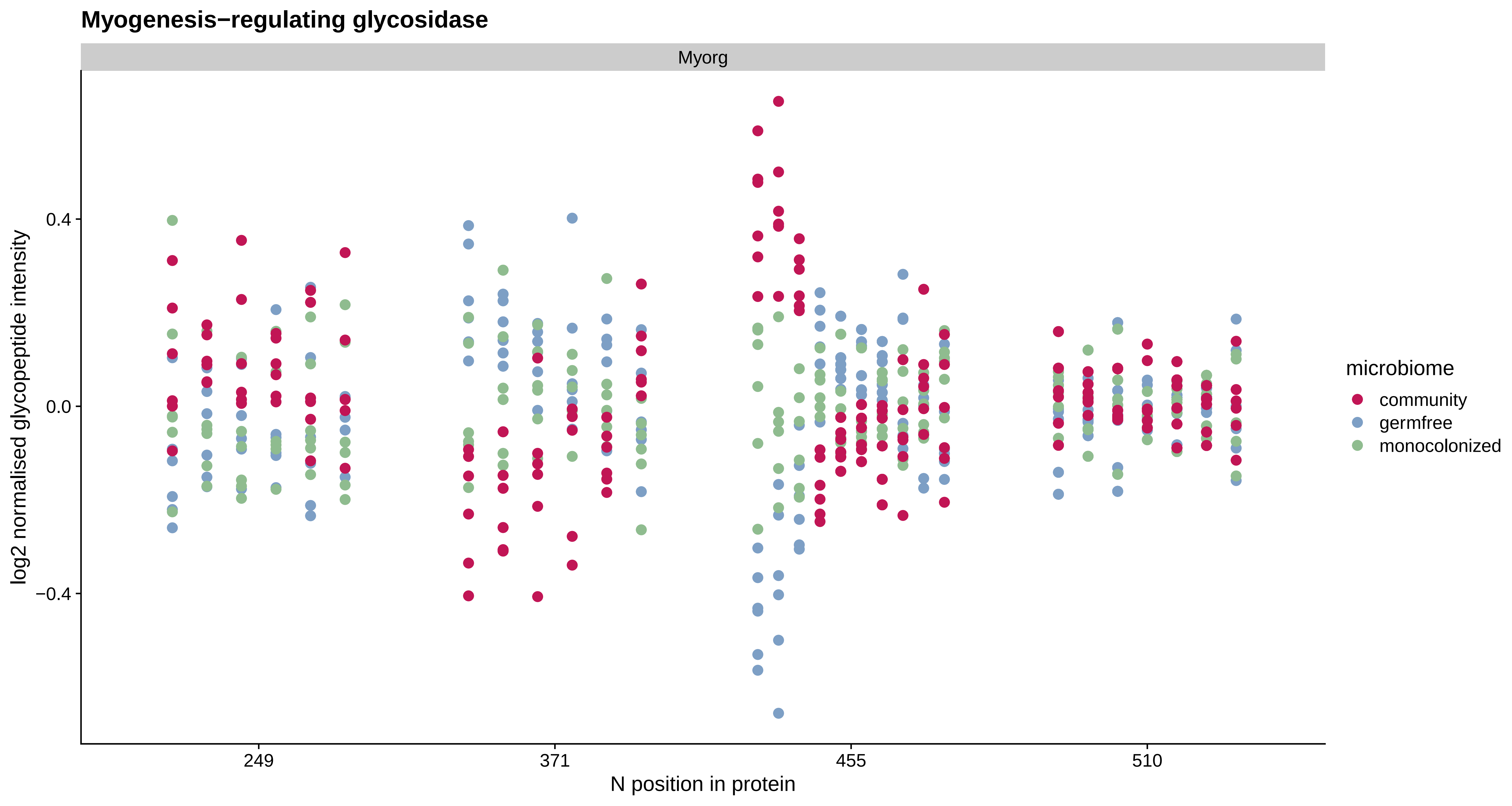
Myogenesis−regulating glycosidase
Myorg
0.4
0.0
−0.4
log2 normalised glycopeptide intensity
249
371
455
510
N position in protein
microbiome
community
germfree
monocolonized

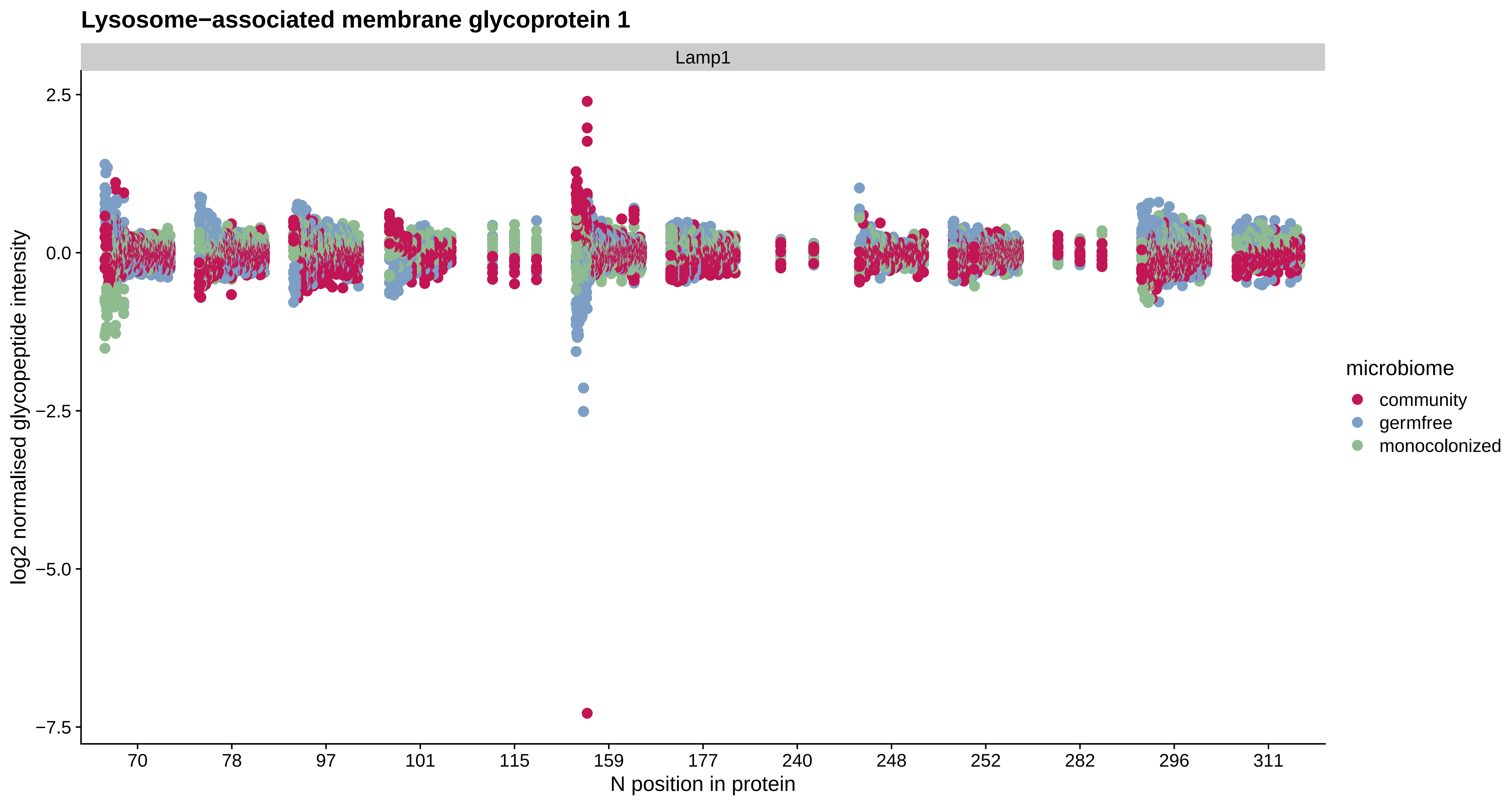
Lysosome−associated membrane glycoprotein 1
Lamp1
log2 normalised glycopeptide intensity
2.5
0.0
−2.5
−5.0
−7.5
70
78
97
101
115
159
177
240
248
252
282
296
311
N position in protein
microbiome
community
germfree
monocolonized

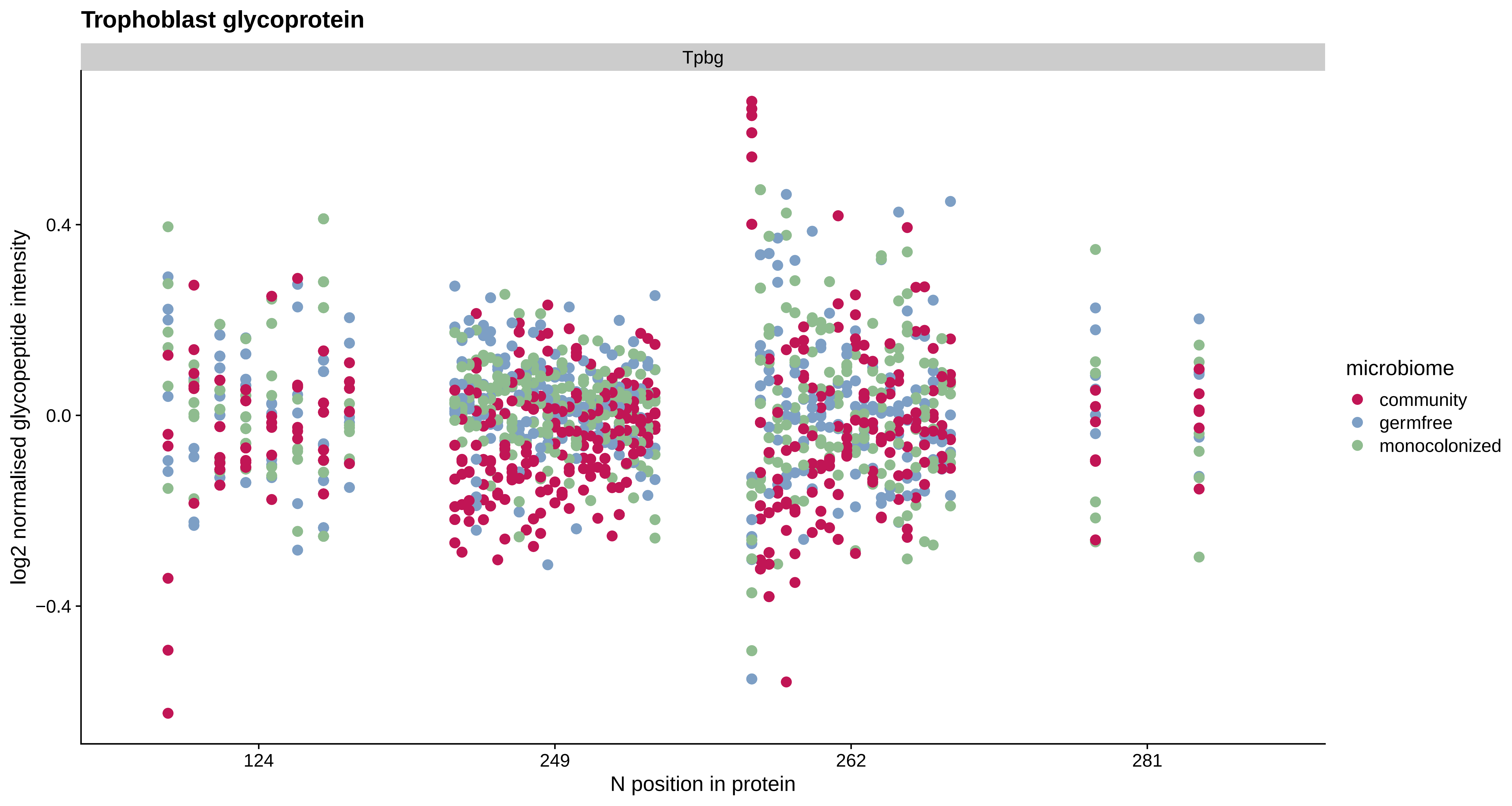
Trophoblast glycoprotein
Tpbg
log2 normalised glycopeptide intensity
0.4
0.0
−0.4
124
249
262
281
N position in protein
microbiome
community
germfree
monocolonized

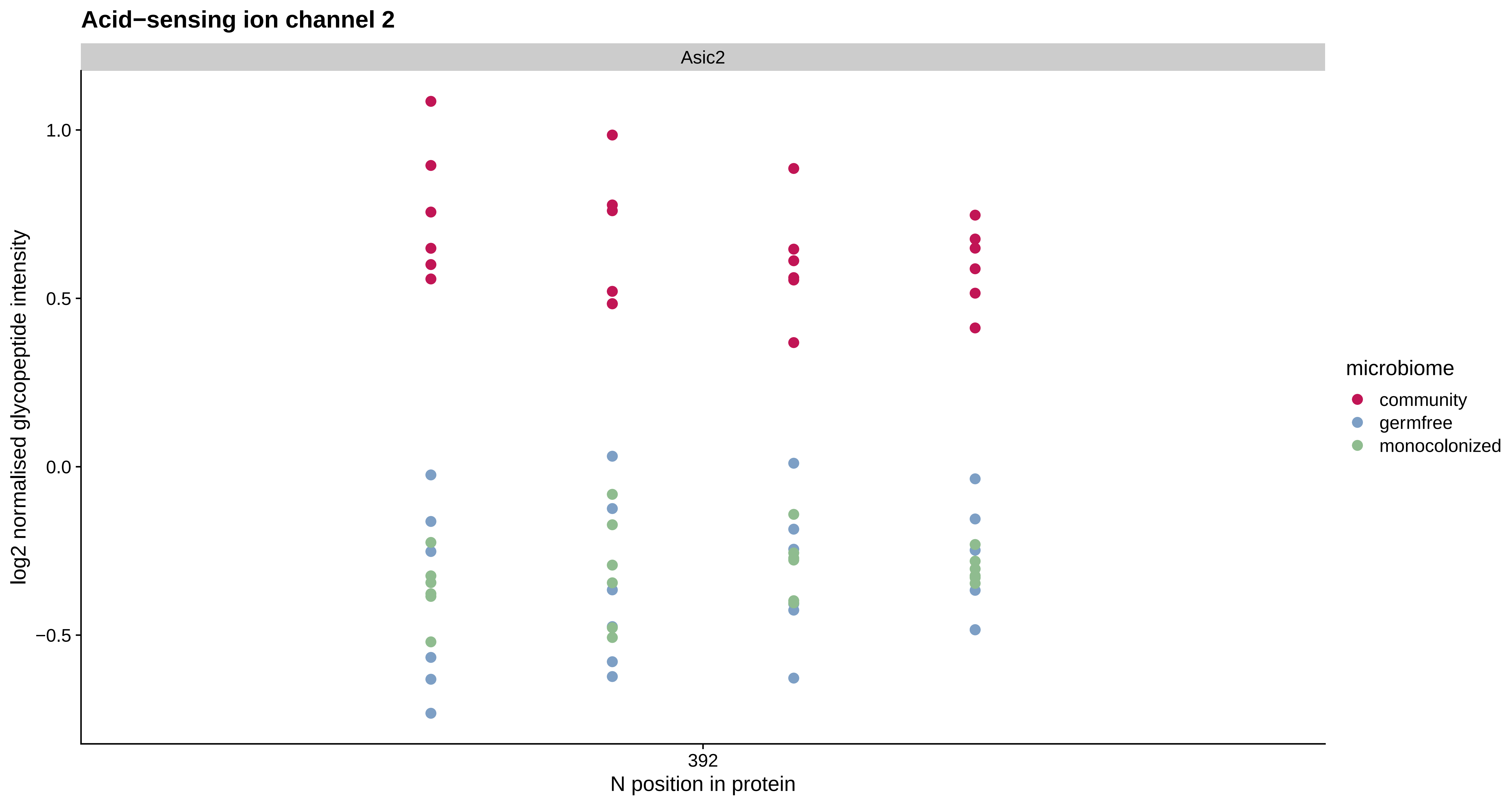

Acid−sensing ion channel 2
Asic2
log2 normalised glycopeptide intensity
1.0
0.5
0.0
−0.5
392
N position in protein
microbiome
community
germfree
monocolonized

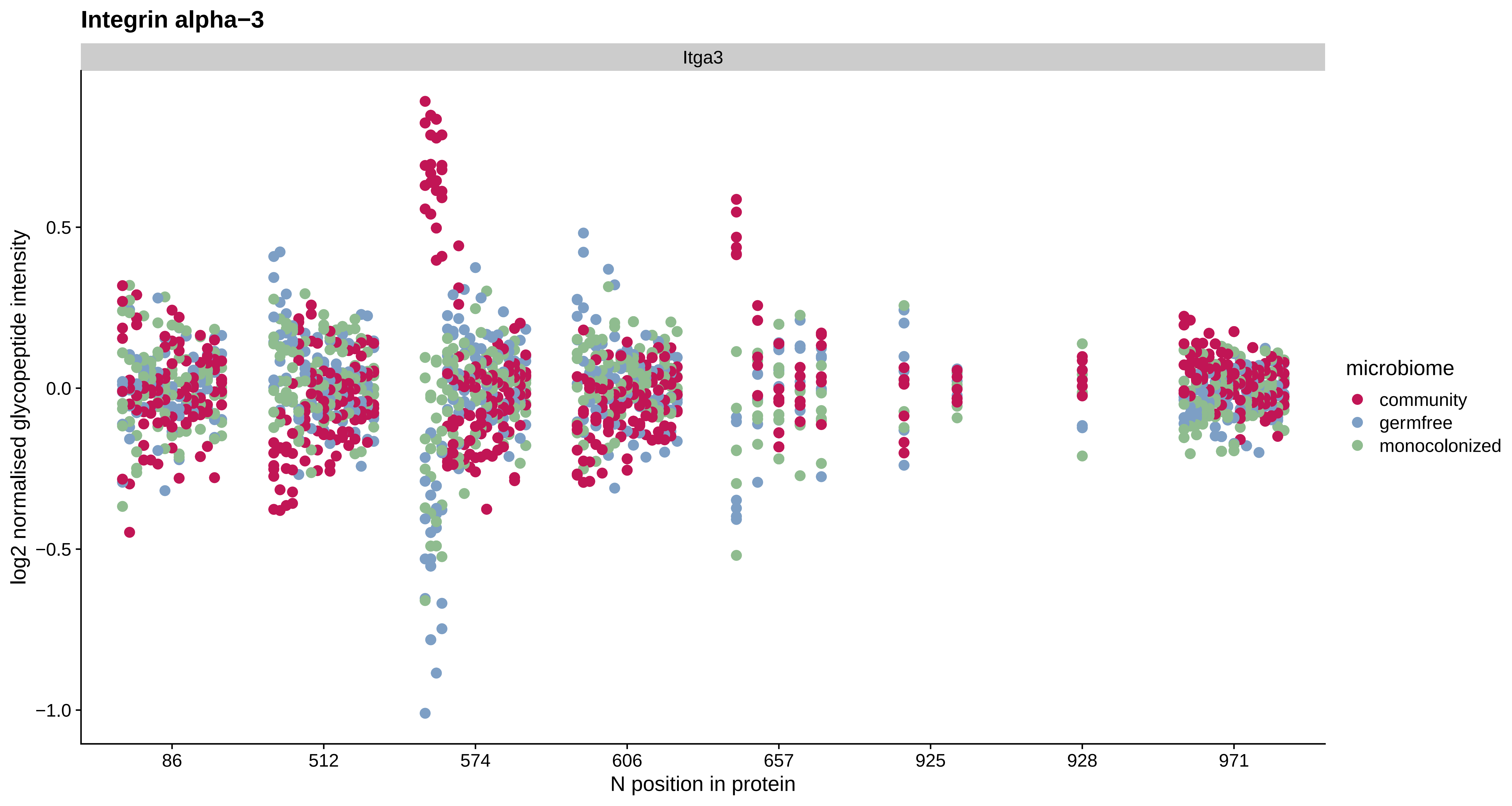
Integrin alpha−3
Itga3
log2 normalised glycopeptide intensity
0.5
0.0
−0.5
−1.0
86
512
574
606
657
925
928
971
N position in protein
microbiome
community
germfree
monocolonized

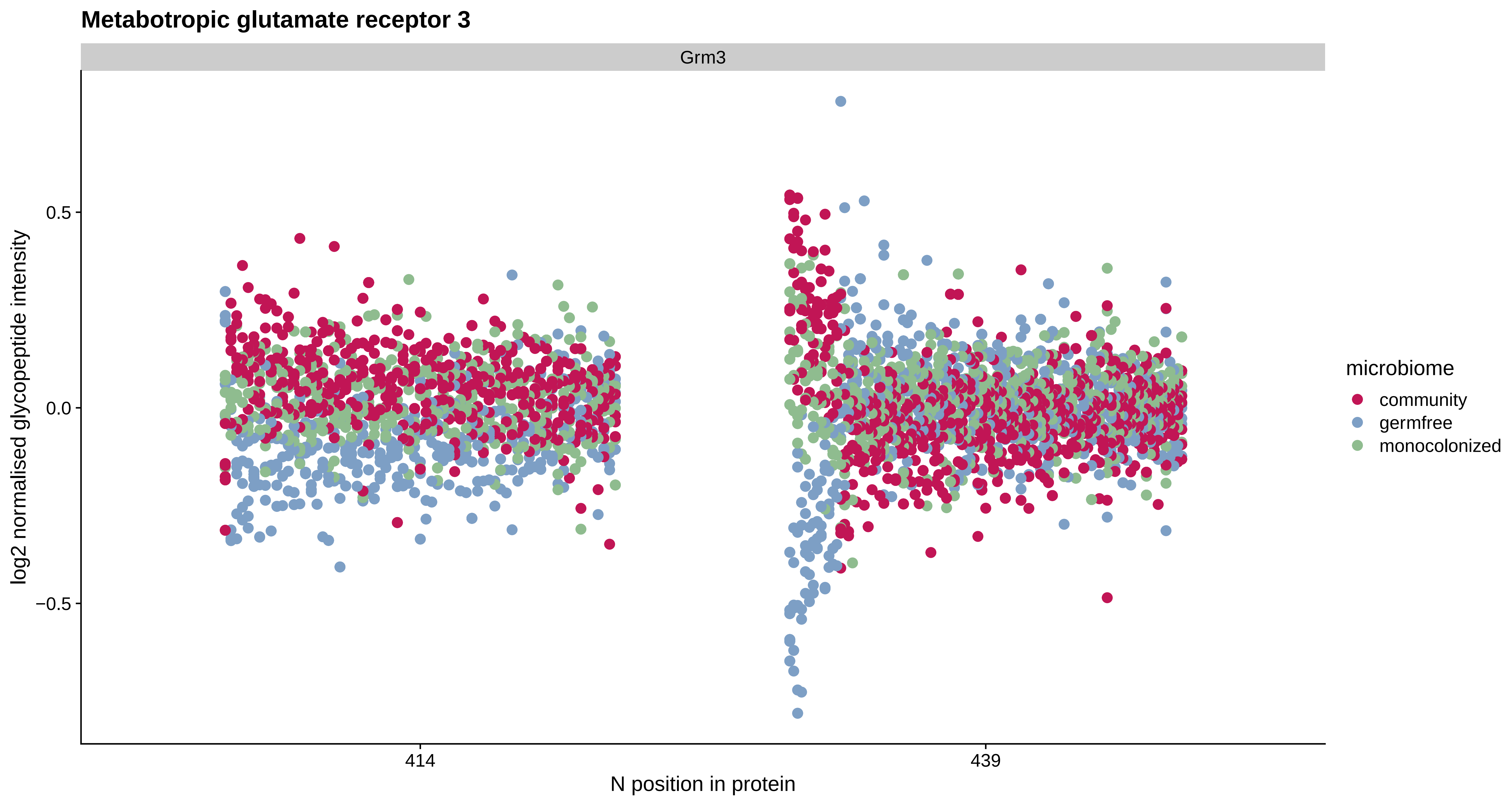

Metabotropic glutamate receptor 3
Grm3
log2 normalised glycopeptide intensity
0.5
0.0
−0.5
414
439
N position in protein
microbiome
community
germfree
monocolonized

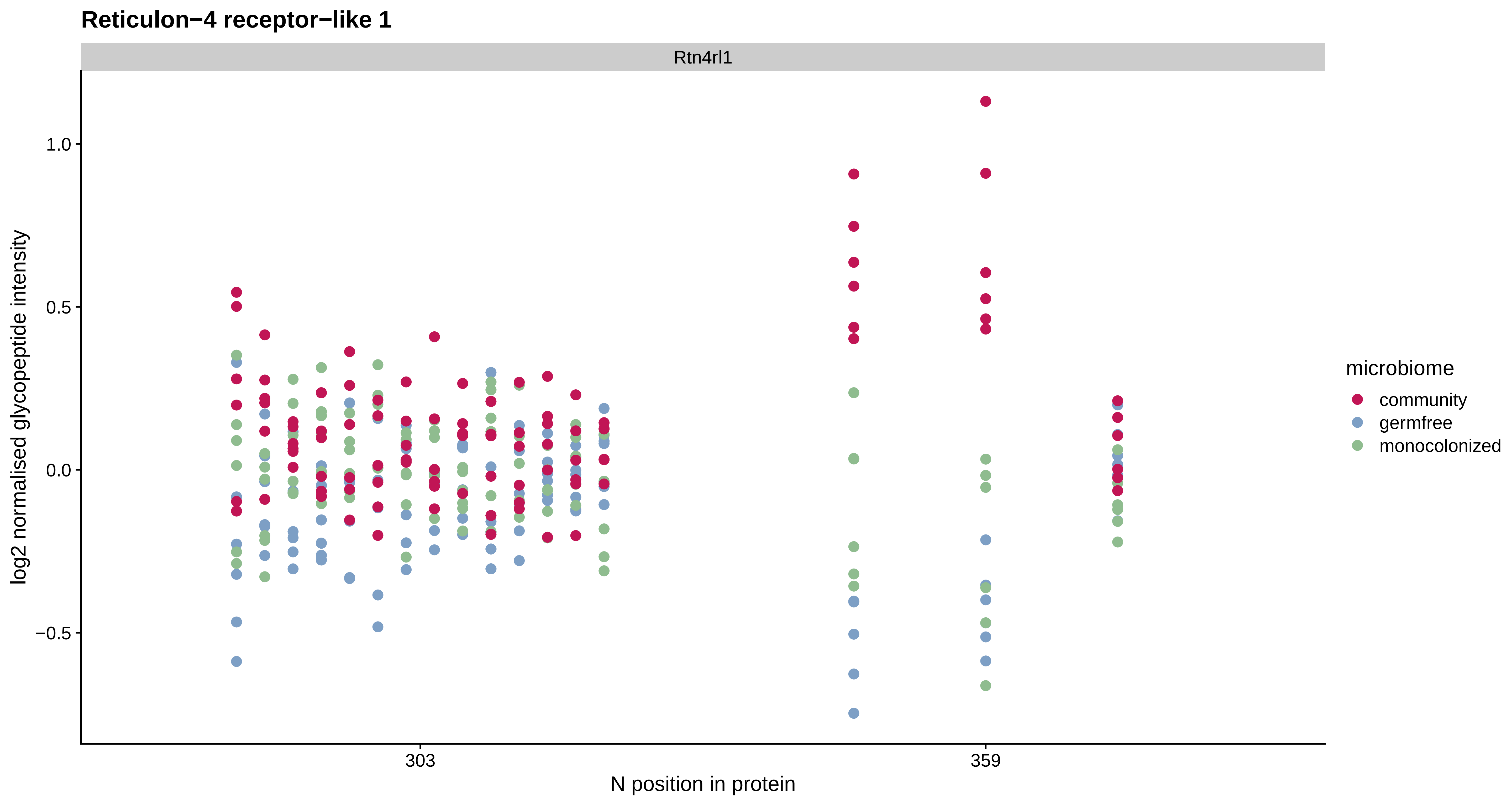
Reticulon−4 receptor−like 1
Rtn4rl1
log2 normalised glycopeptide intensity
1.0
0.5
0.0
−0.5
303
359
N position in protein
microbiome
community
germfree
monocolonized

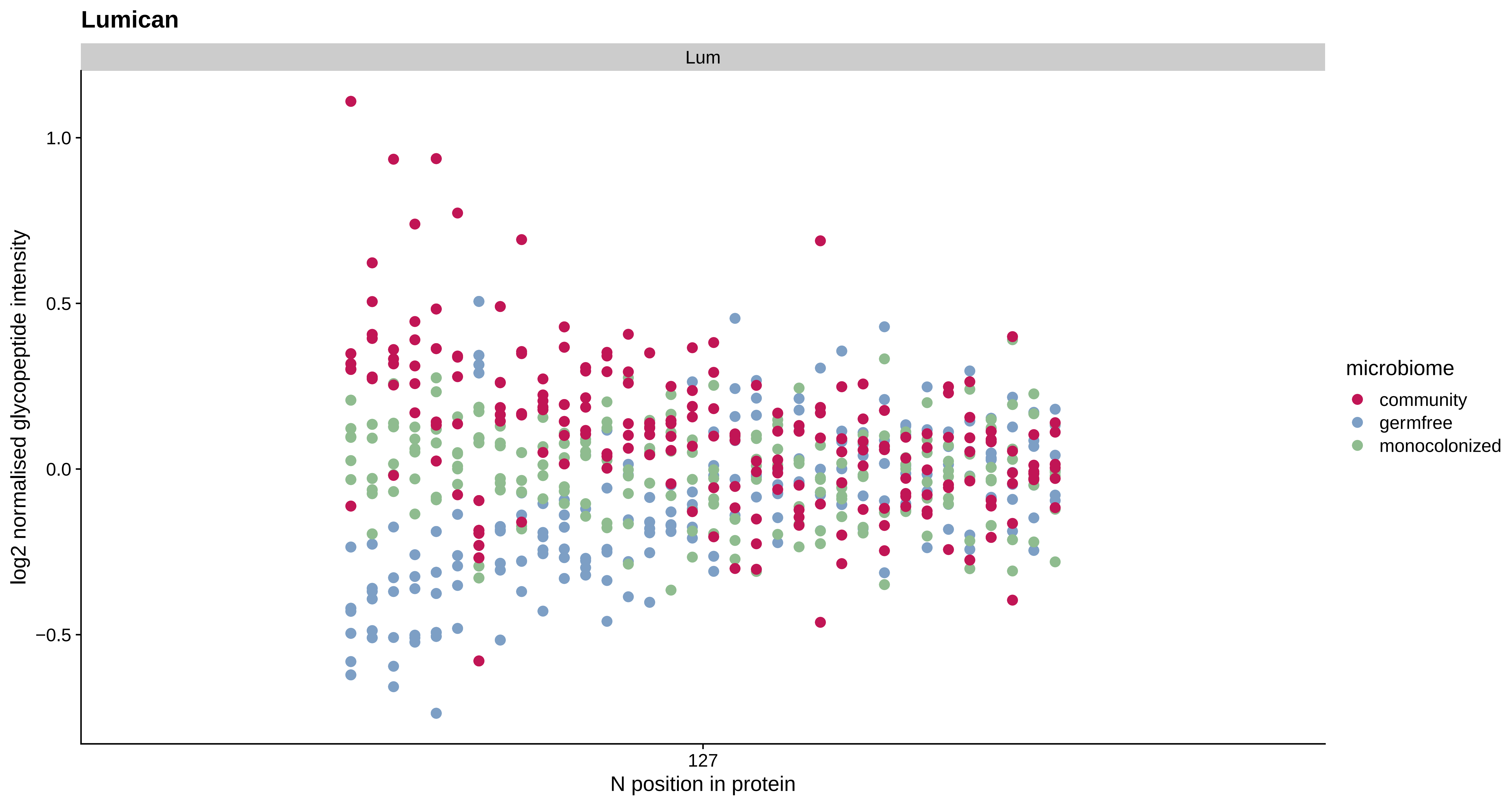
Lumican
Lum
1.0
0.5
0.0
−0.5
log2 normalised glycopeptide intensity
127
N position in protein
microbiome
community
germfree
monocolonized

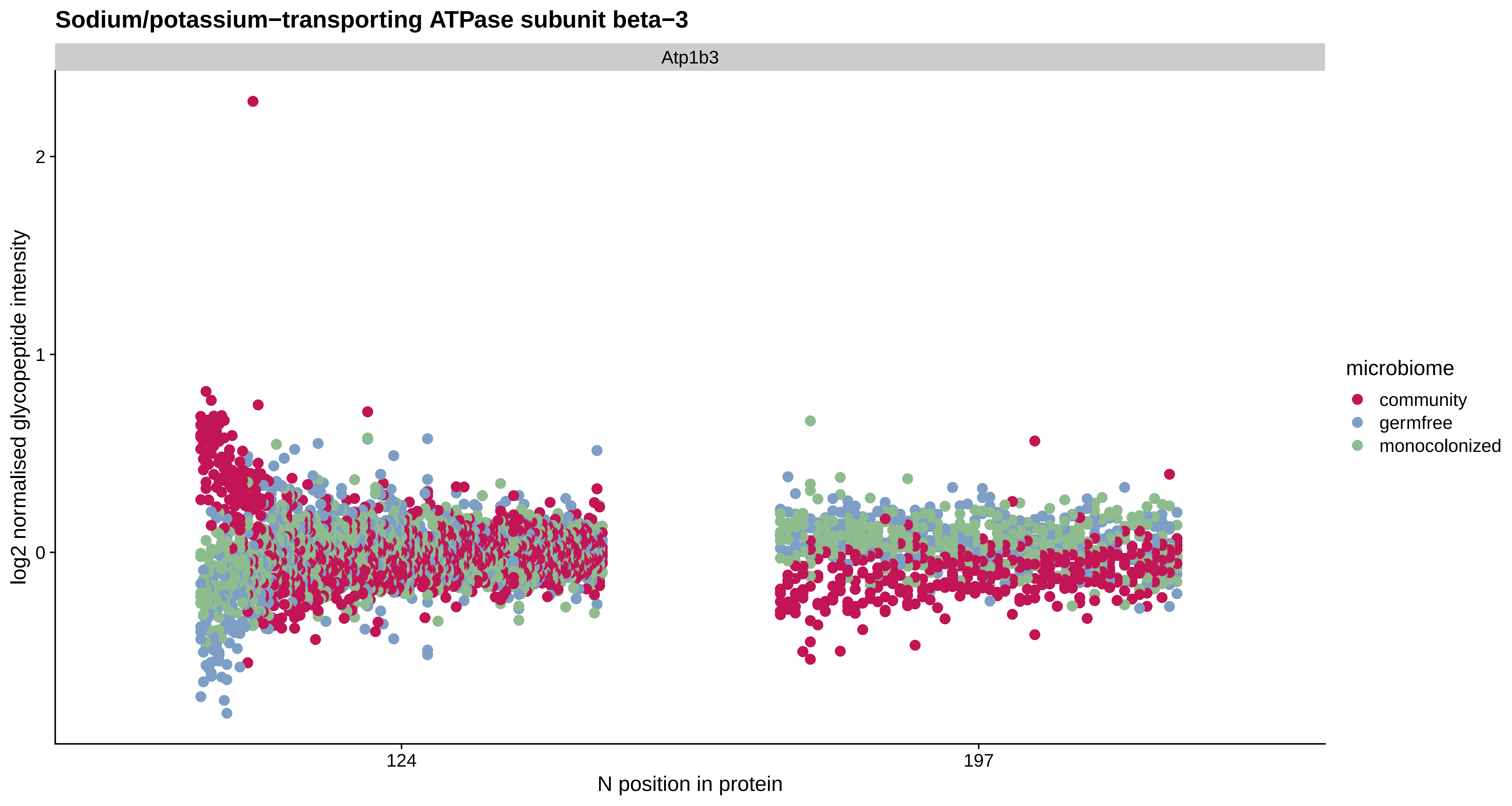

Sodium/potassium−transporting ATPase subunit beta−3
Atp1b3
log2 normalised glycopeptide intensity
2
1
0
124
197
N position in protein
microbiome
community
germfree
monocolonized

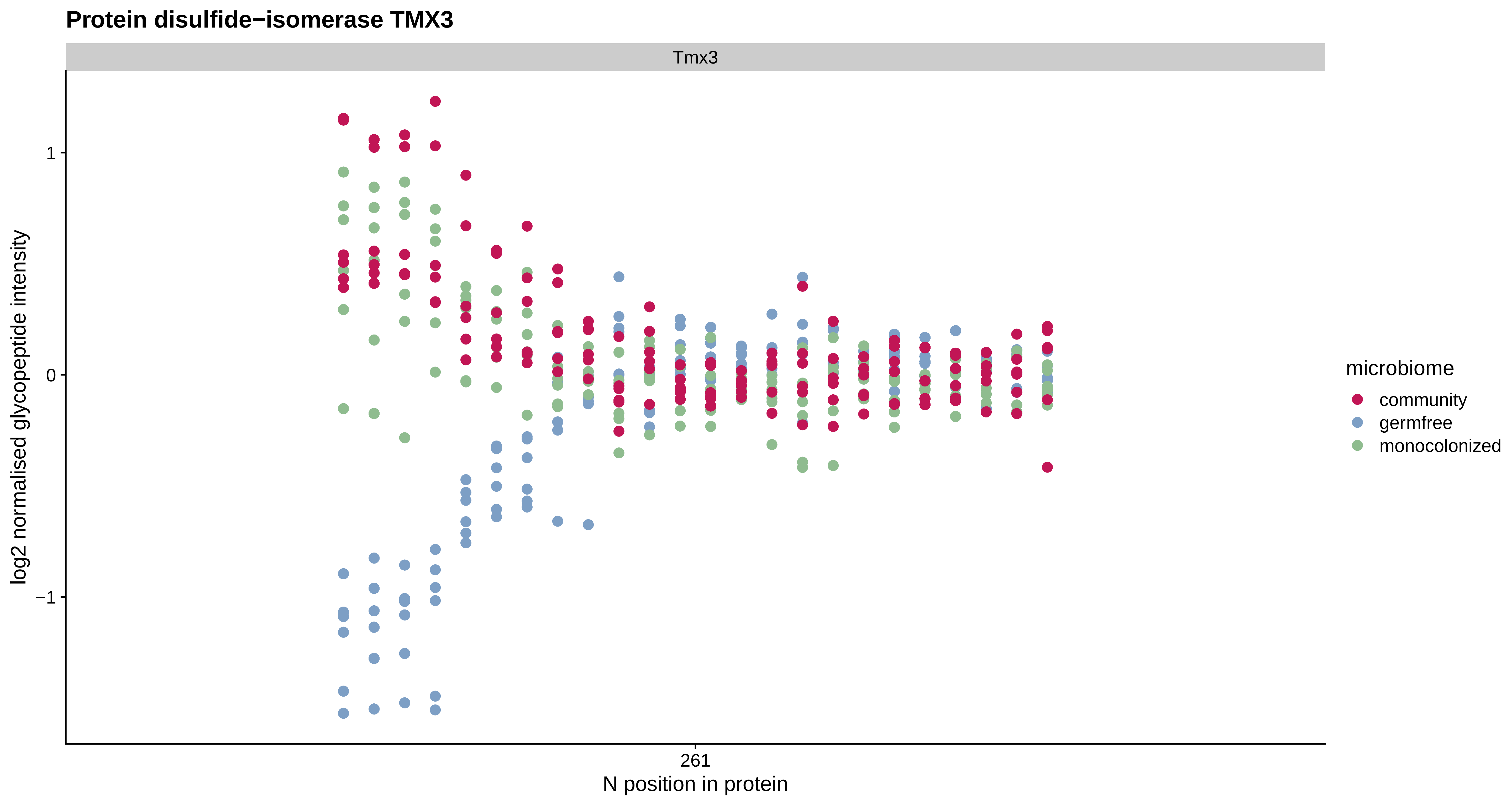
Protein disulfide−isomerase TMX3
Tmx3
log2 normalised glycopeptide intensity
1
0
−1
261
N position in protein
microbiome
community
germfree
monocolonized

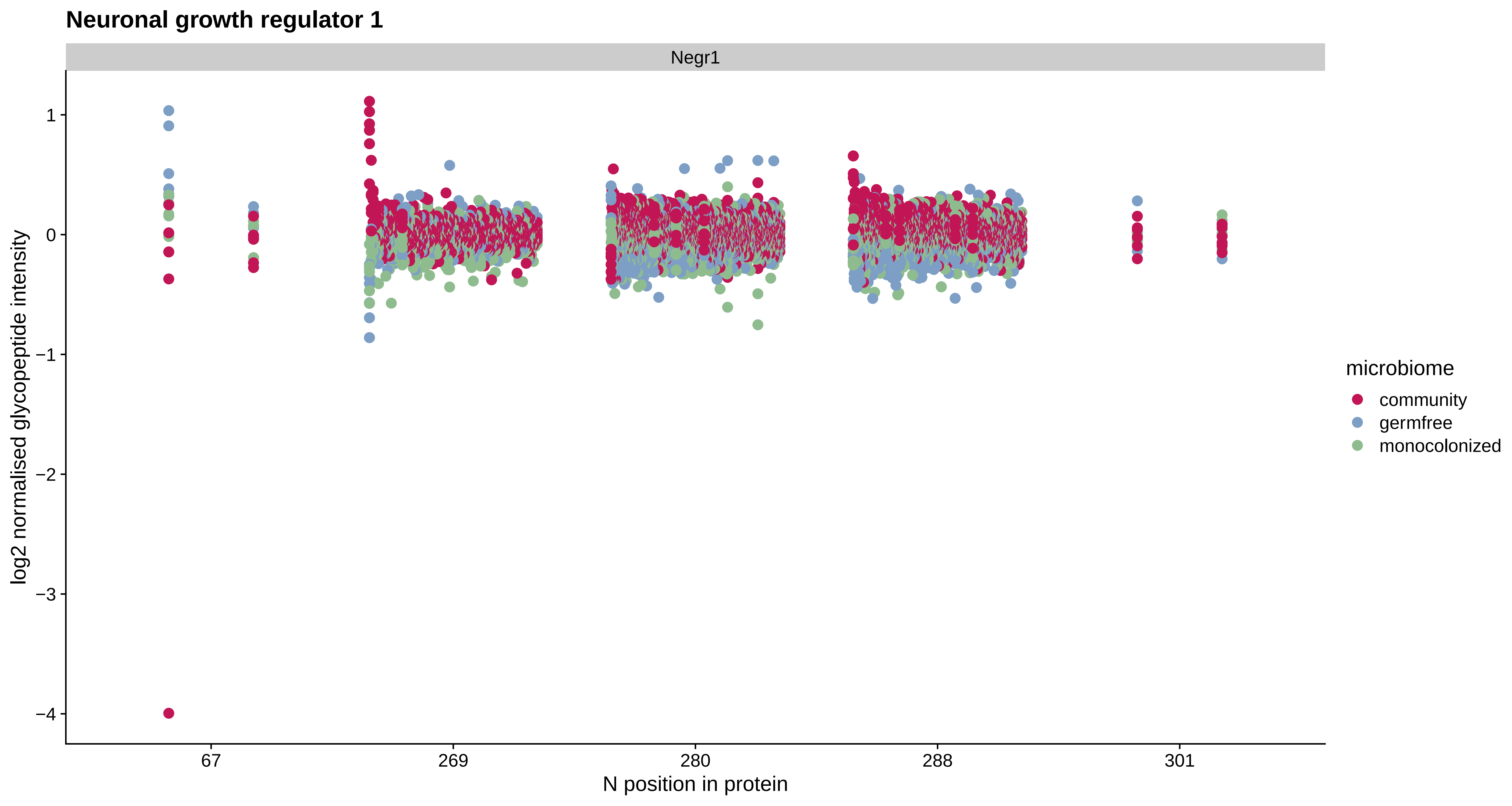
Neuronal growth regulator 1
Negr1
log2 normalised glycopeptide intensity
1
0
−1
−2
−3
−4
67
269
280
288
301
N position in protein
microbiome
community
germfree
monocolonized

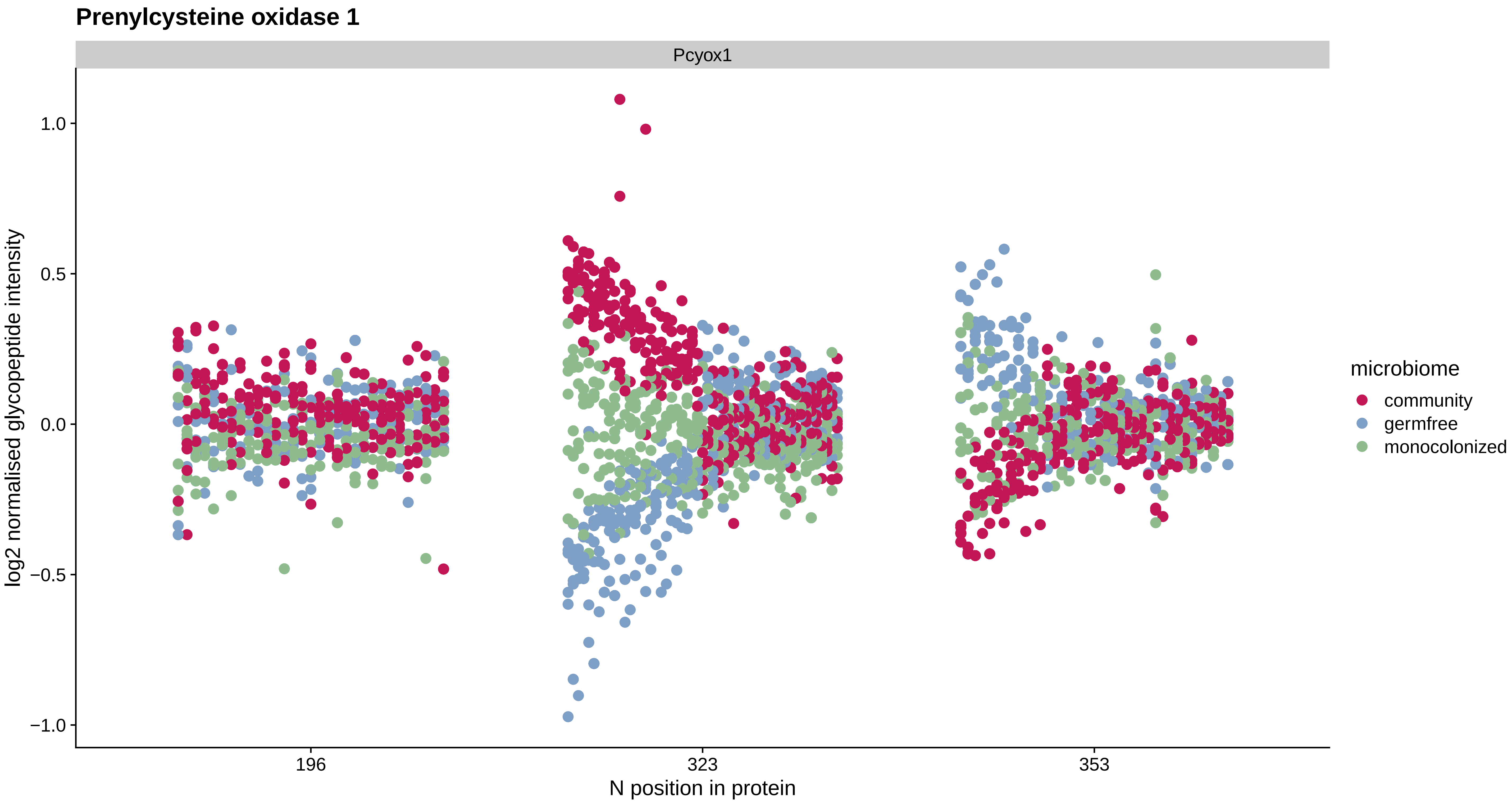

Prenylcysteine oxidase 1
Pcyox1
log2 normalised glycopeptide intensity
1.0
0.5
0.0
−0.5
−1.0
196
323
353
N position in protein
microbiome
community
germfree
monocolonized

#### Protocadherin gamma B6

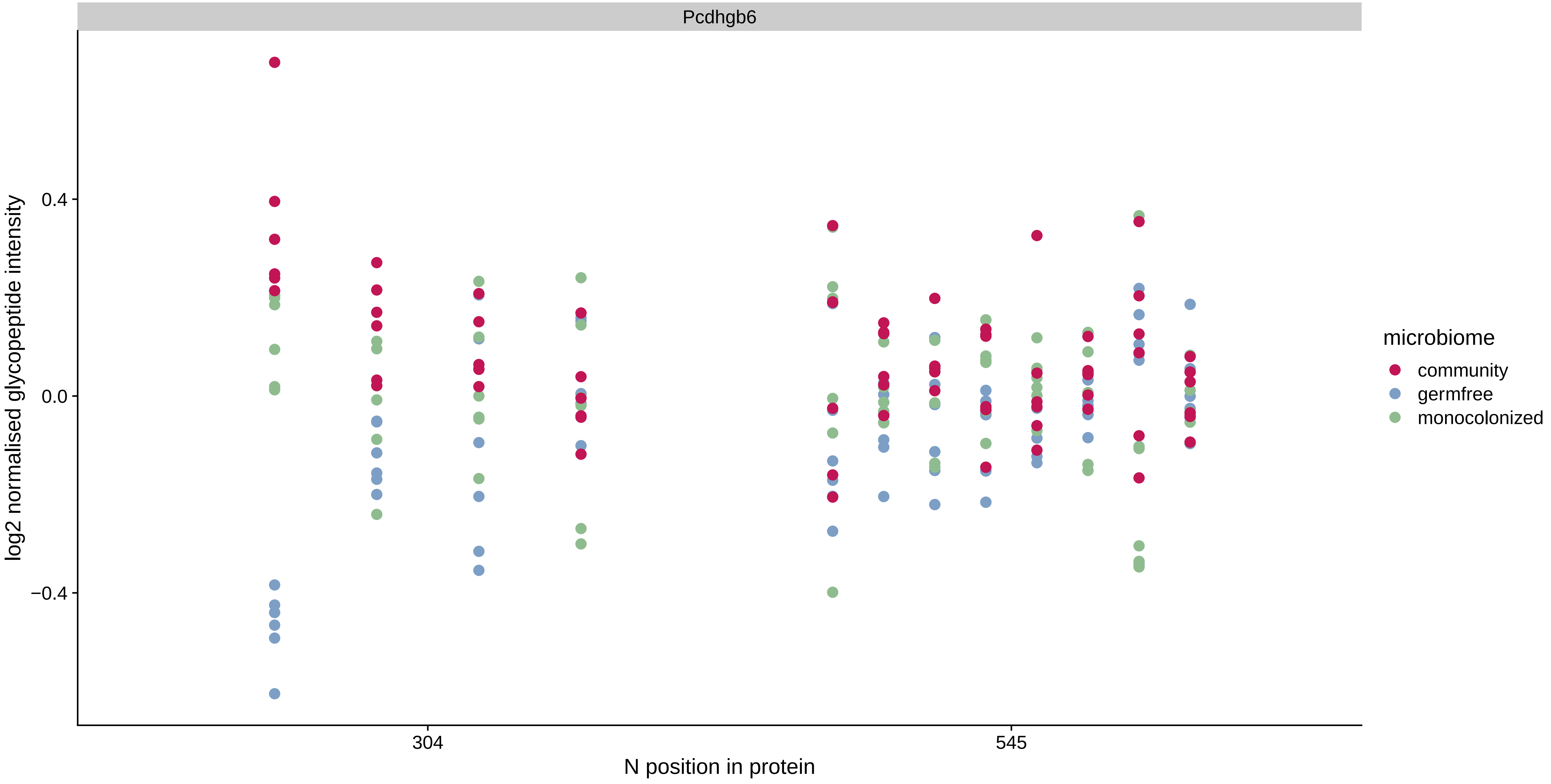

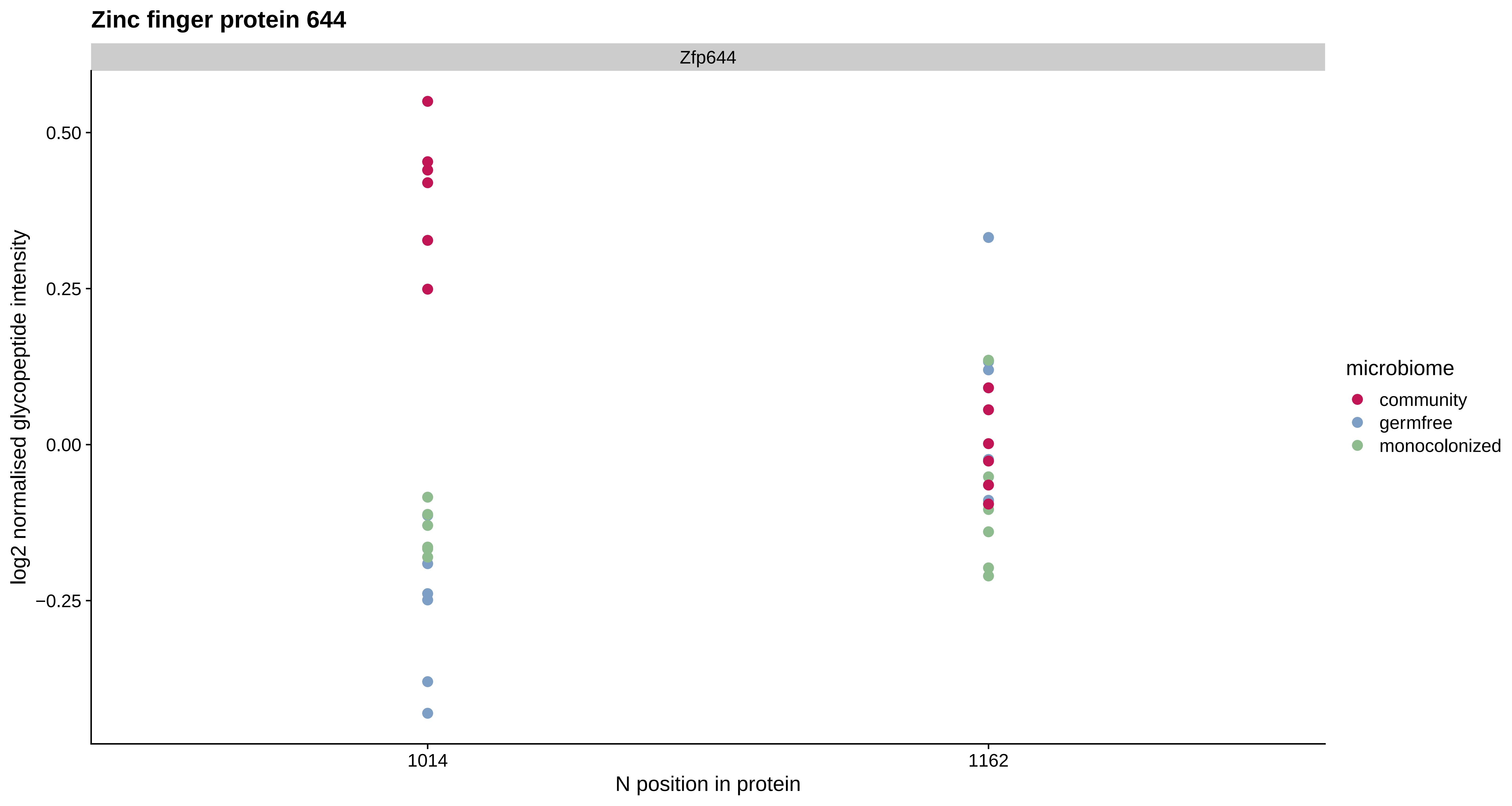
Zinc finger protein 644
Zfp644
0.50
0.25
0.00
−0.25
log2 normalised glycopeptide intensity
1014
1162
N position in protein
microbiome
community
germfree
monocolonized

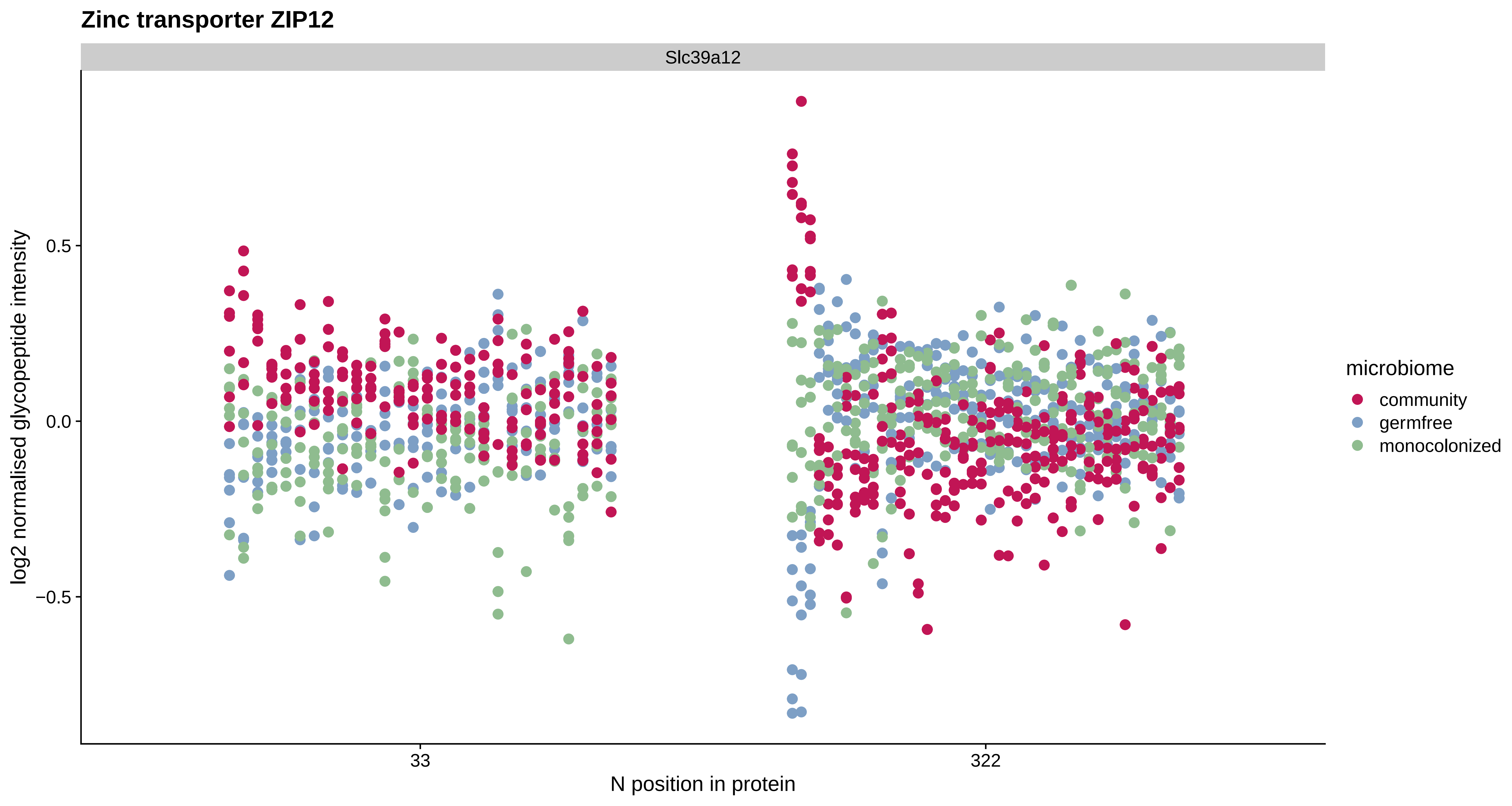

Zinc transporter ZIP12
Slc39a12
log2 normalised glycopeptide intensity
0.5
0.0
−0.5
33
322
N position in protein
microbiome
community
germfree
monocolonized

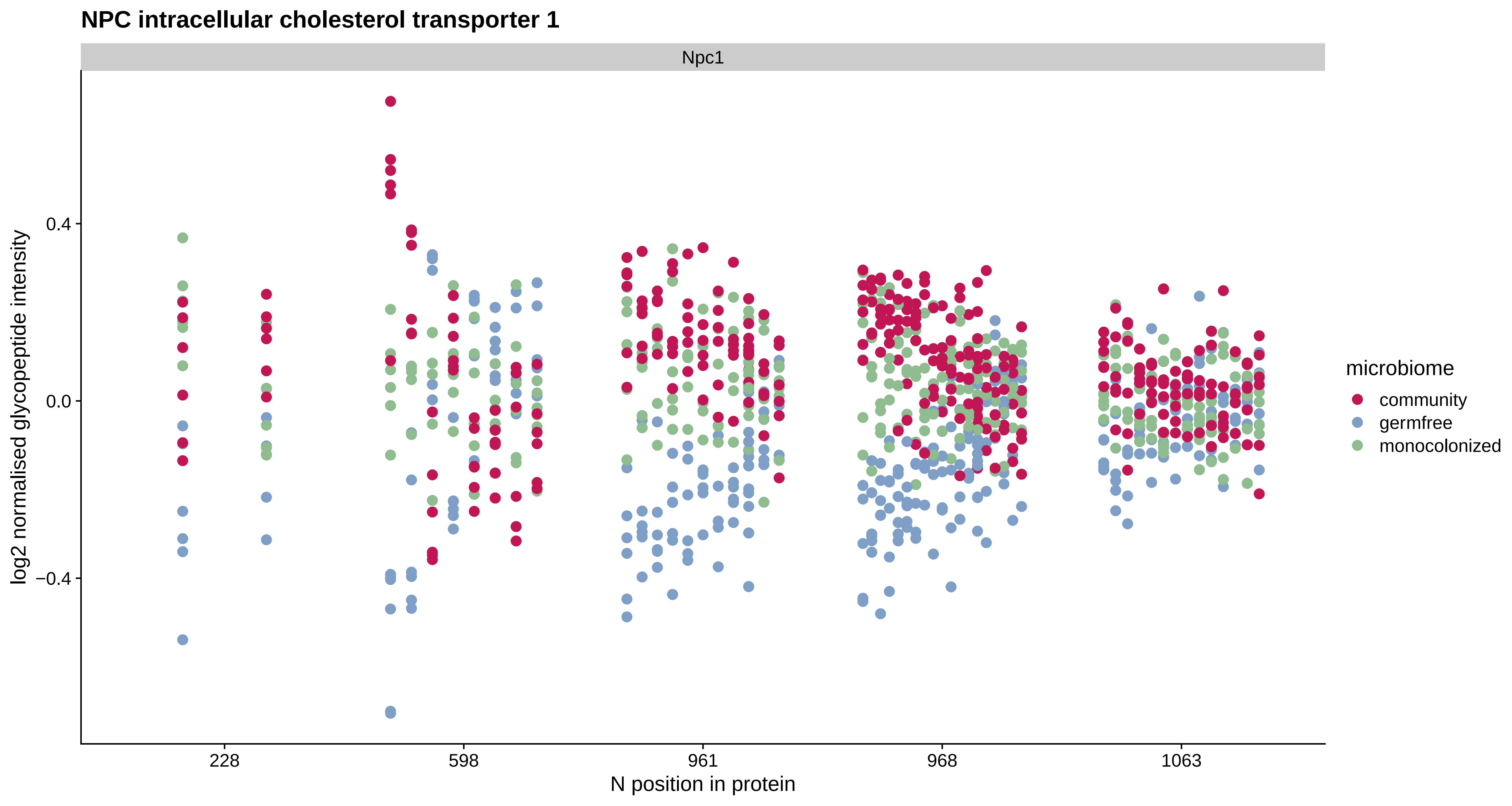
NPC intracellular cholesterol transporter 1
Npc1
log2 normalised glycopeptide intensity
0.4
0.0
−0.4
228
598
961
968
1063
N position in protein
microbiome
community
germfree
monocolonized

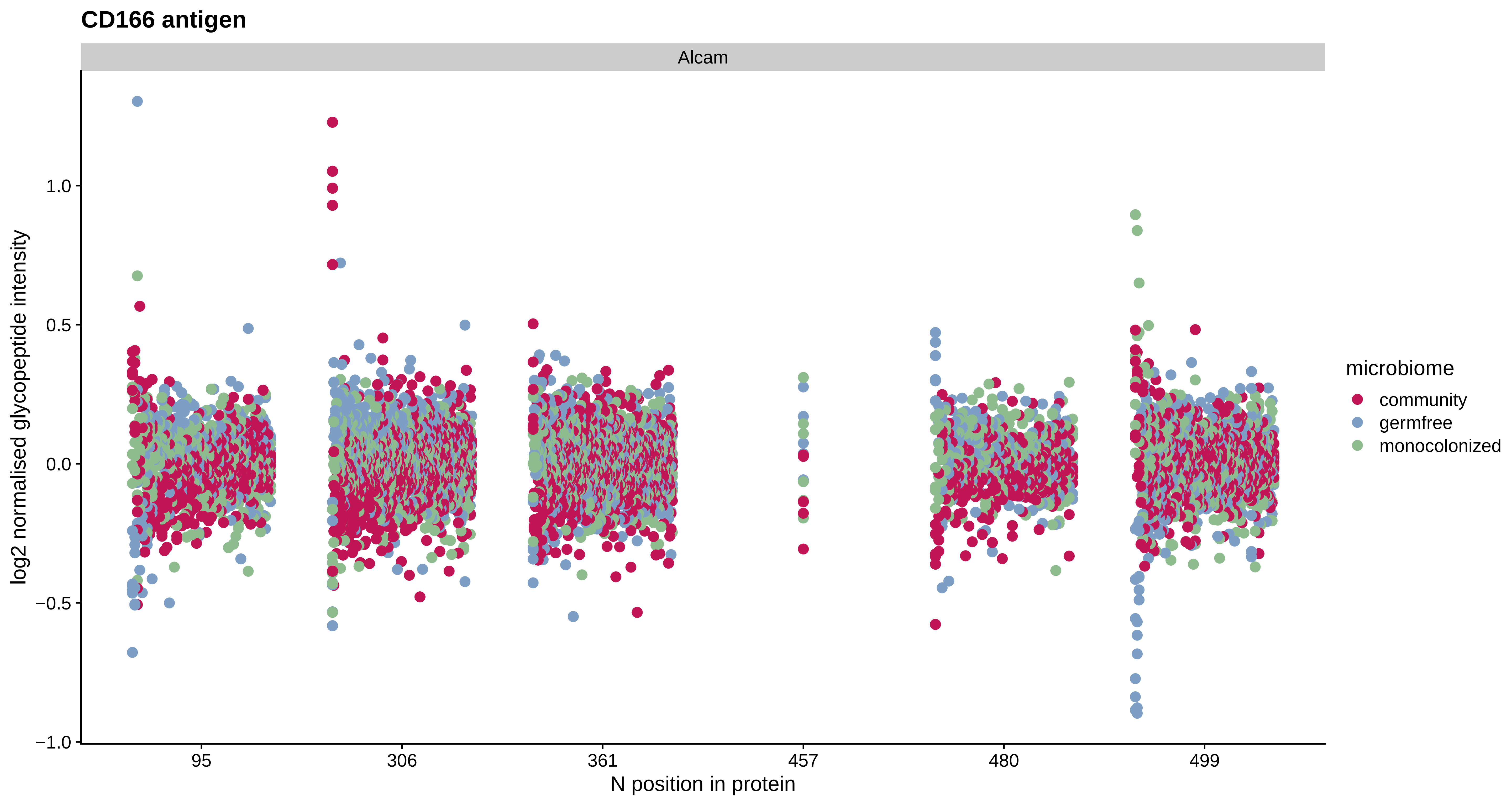
CD166 antigen
Alcam
log2 normalised glycopeptide intensity
1.0
0.5
0.0
−0.5
−1.0
95
306
361
457
480
499
N position in protein
microbiome
community
germfree
monocolonized

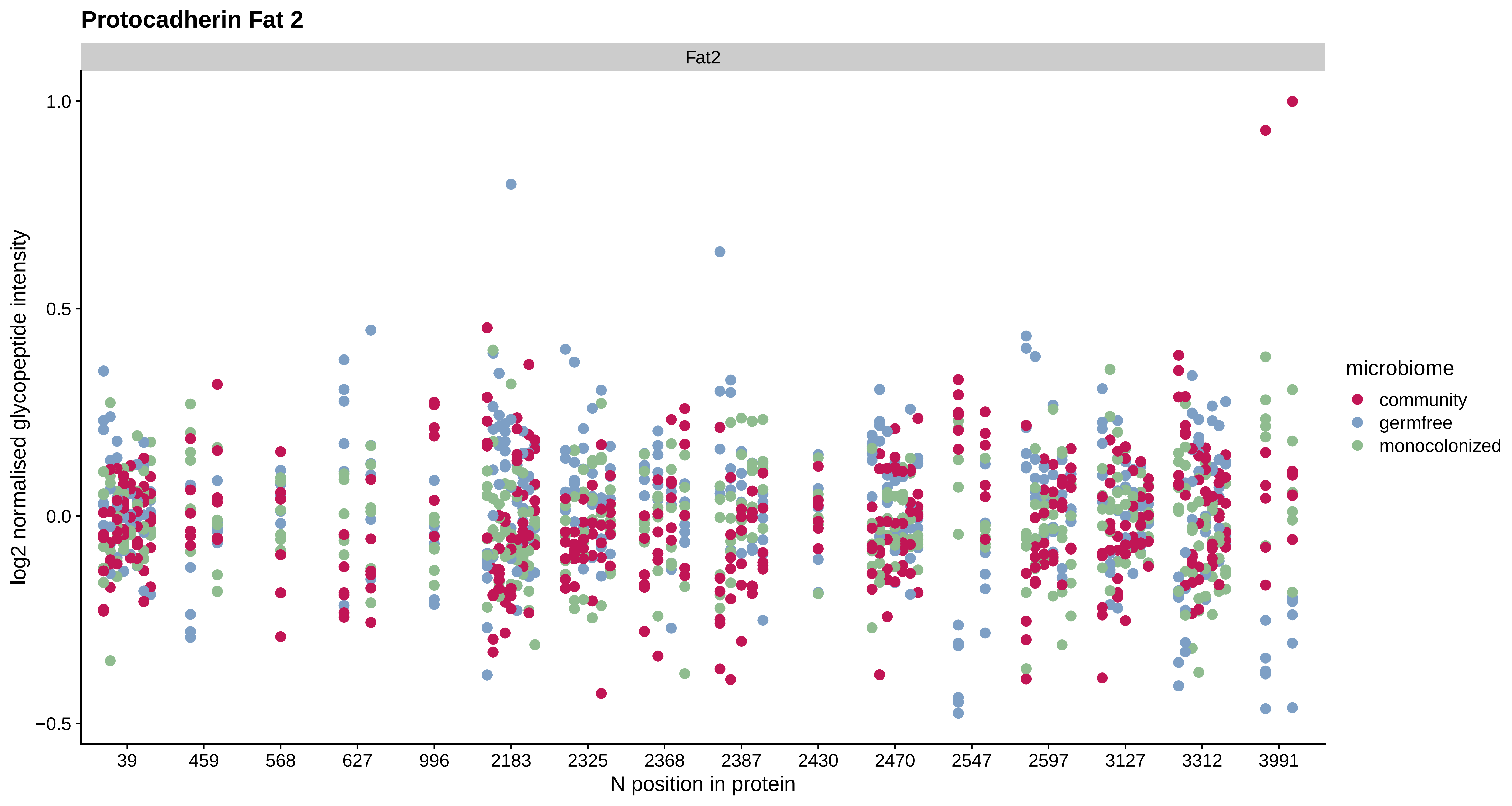
Protocadherin Fat 2
Fat2
log2 normalised glycopeptide intensity
1.0
0.5
0.0
−0.5
39
459
568
627
996
2183
2325
2368
2387
2430
2470
2547
2597
3127
3312
3991
N position in protein
microbiome
community
germfree
monocolonized

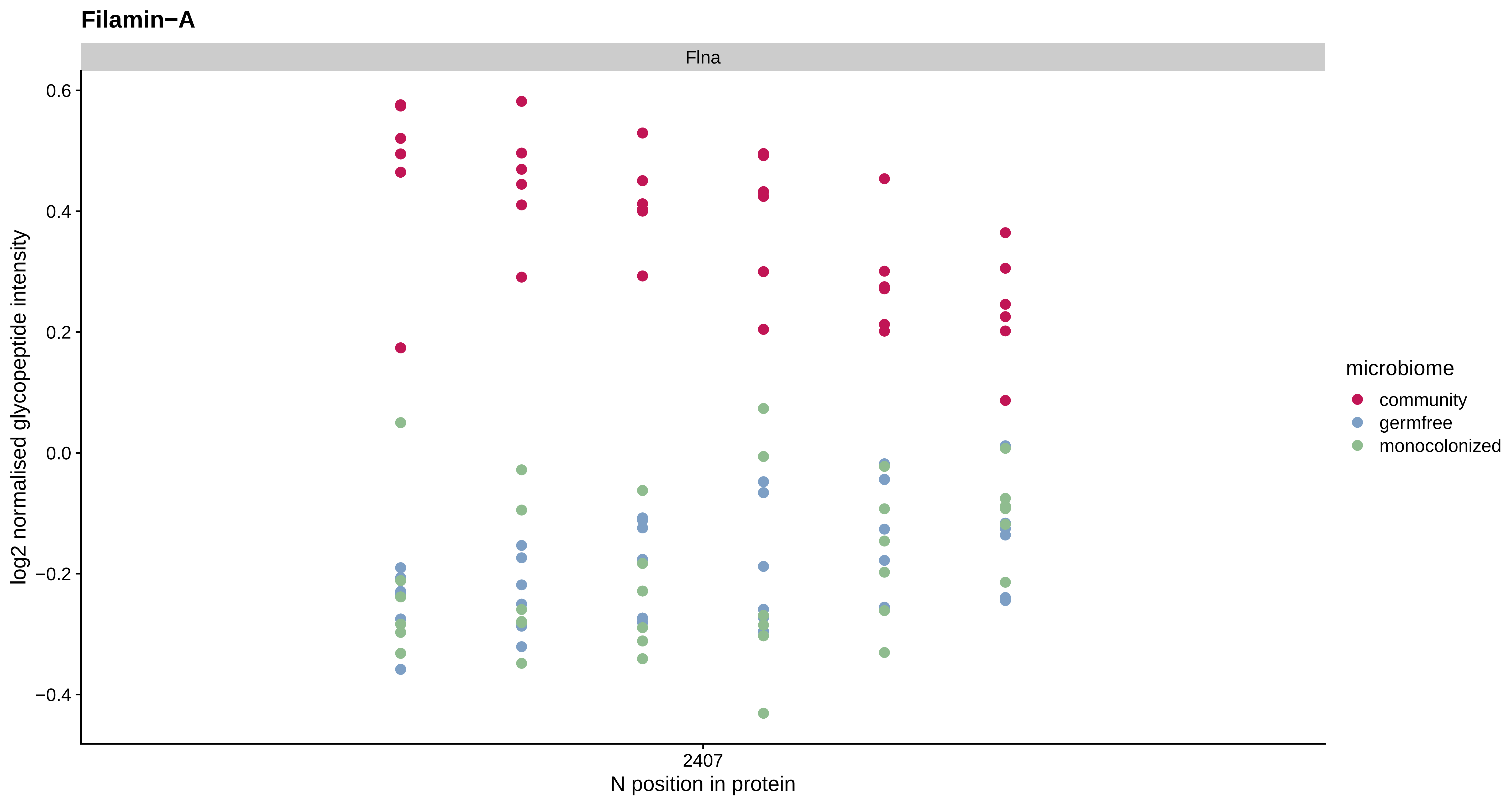

Filamin−A
Flna
0.6
0.4
0.2
0.0
−0.2
−0.4
log2 normalised glycopeptide intensity
2407
N position in protein
microbiome
community
germfree
monocolonized

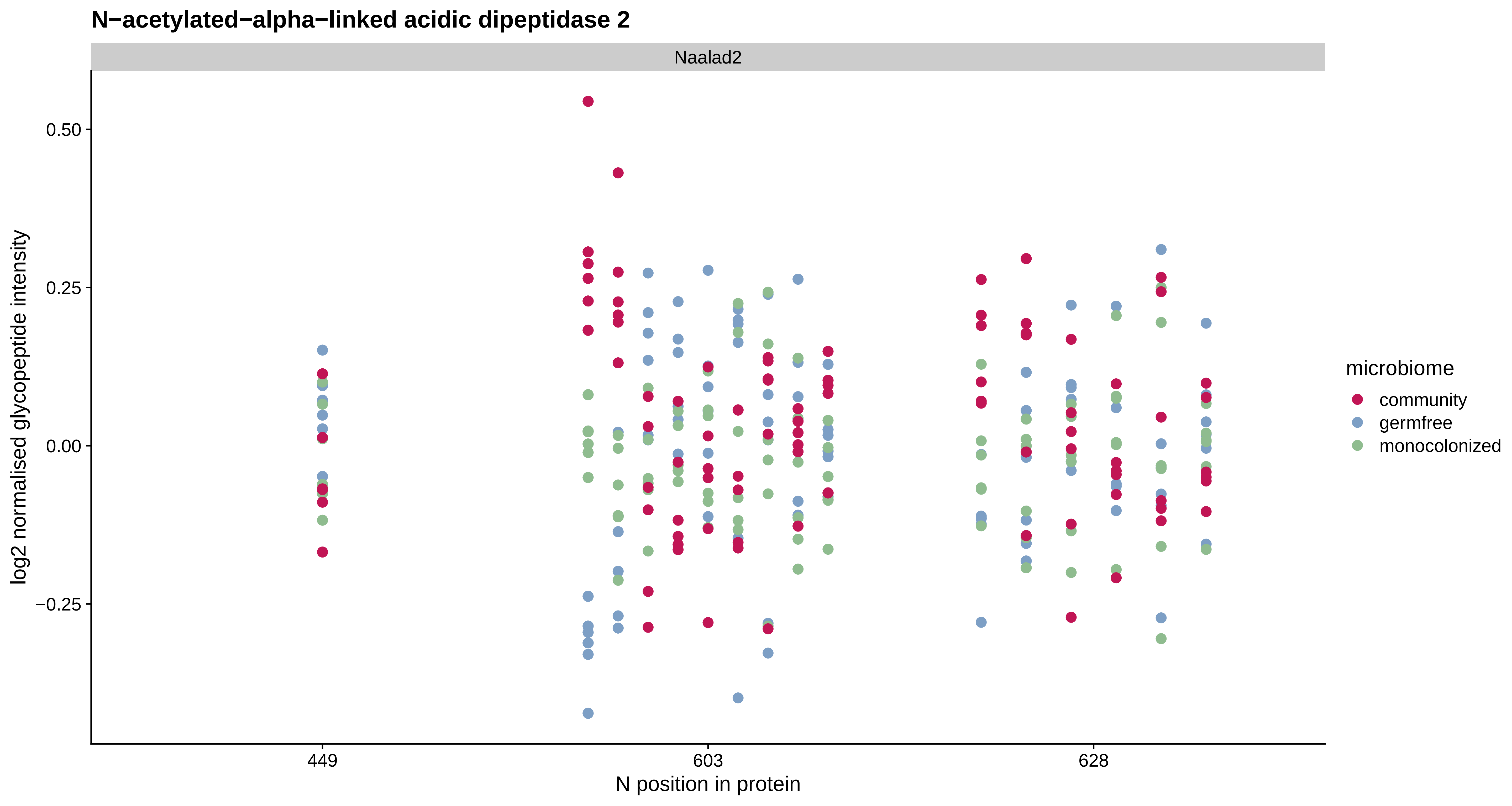

N−acetylated−alpha−linked acidic dipeptidase 2
Naalad2
log2 normalised glycopeptide intensity
0.50
0.25
0.00
−0.25
449
603
628
N position in protein
microbiome
community
germfree
monocolonized

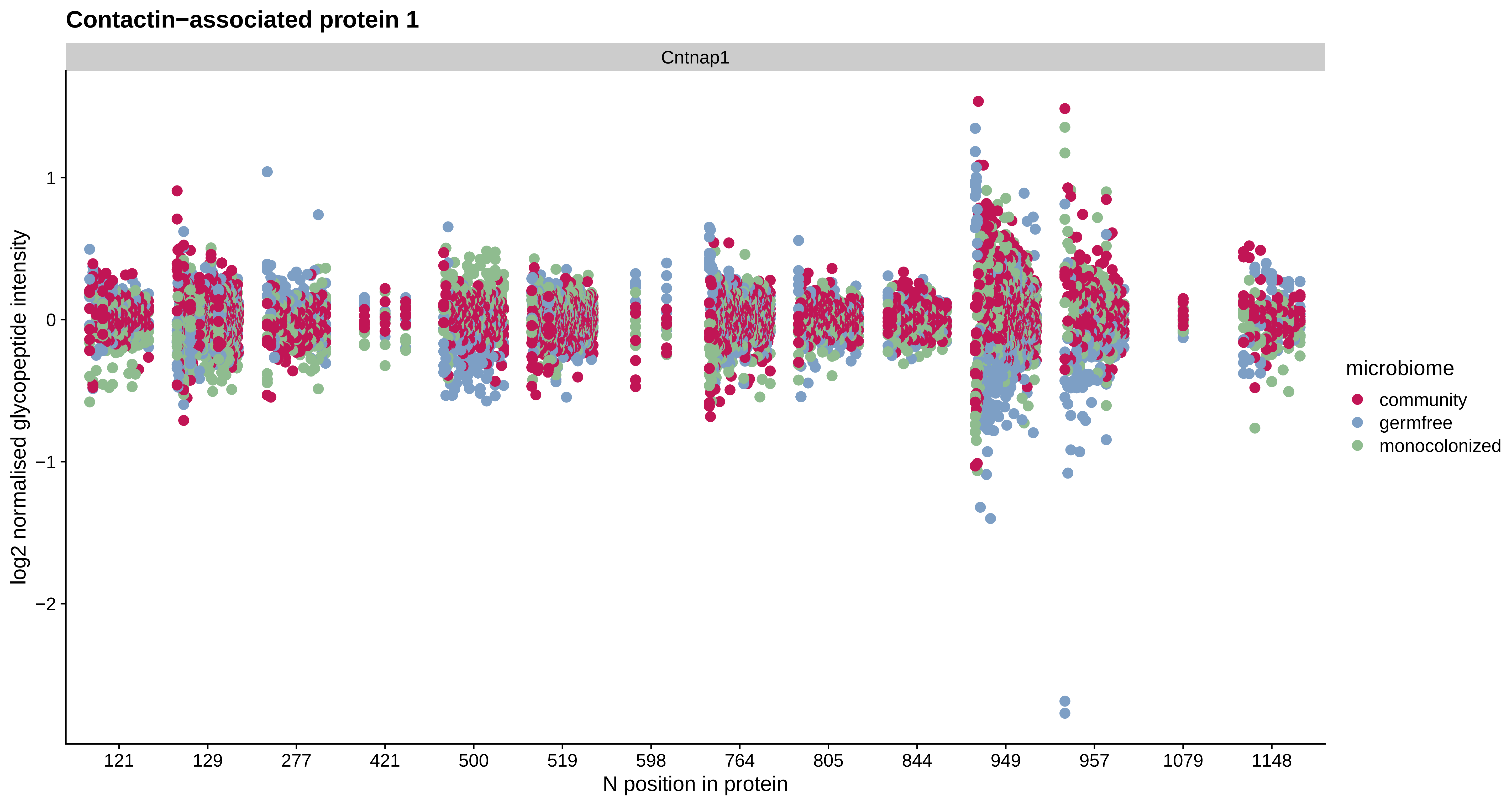

Contactin−associated protein 1
Cntnap1
log2 normalised glycopeptide intensity
1
0
−1
−2
121
129
277
421
500
519
598
764
805
844
949
957
1079
1148
N position in protein
microbiome
community
germfree
monocolonized

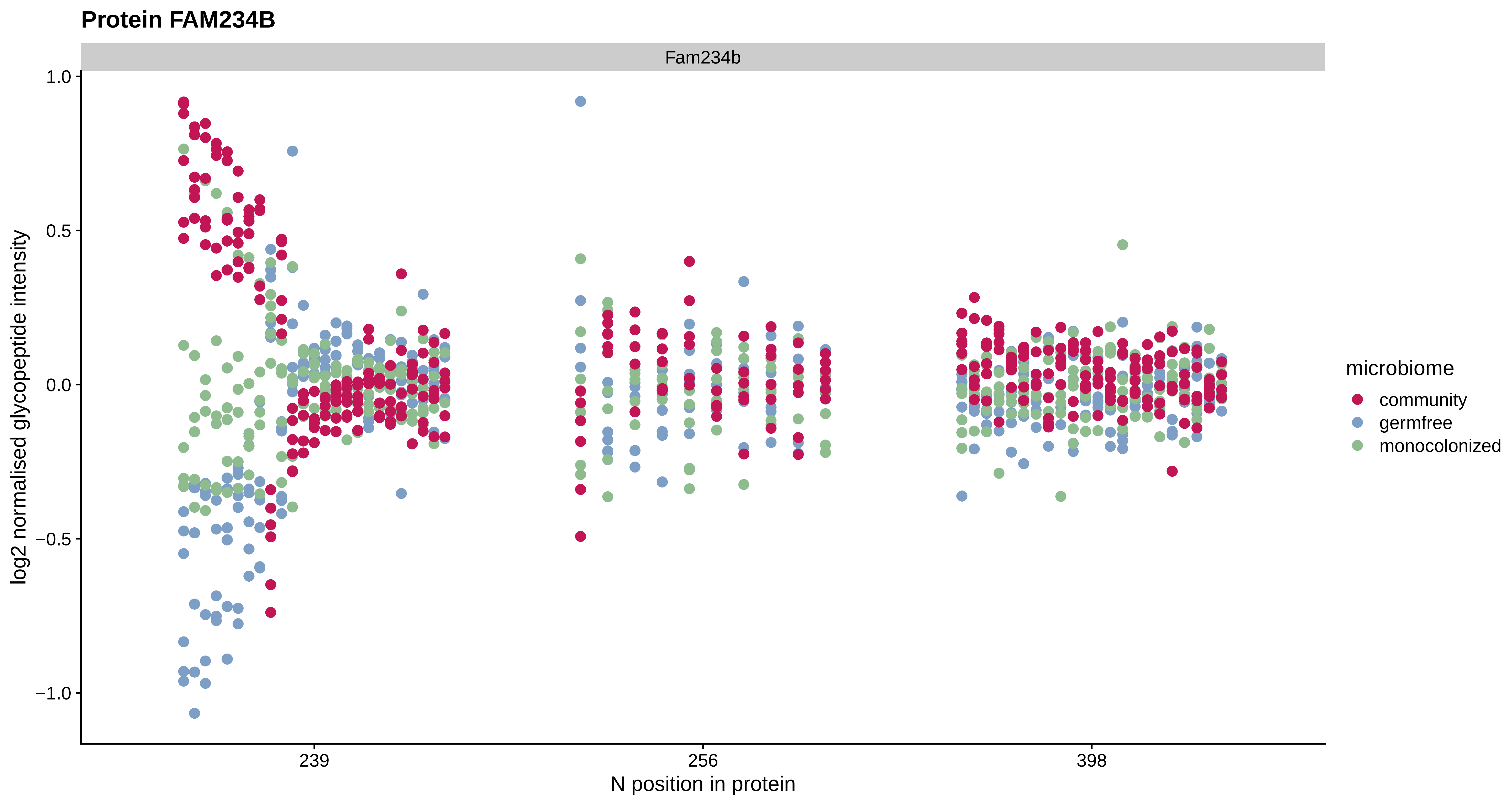

Protein FAM234B
Fam234b
1.0
0.5
0.0
−0.5
−1.0
log2 normalised glycopeptide intensity
239
256
398
N position in protein
microbiome
community
germfree
monocolonized

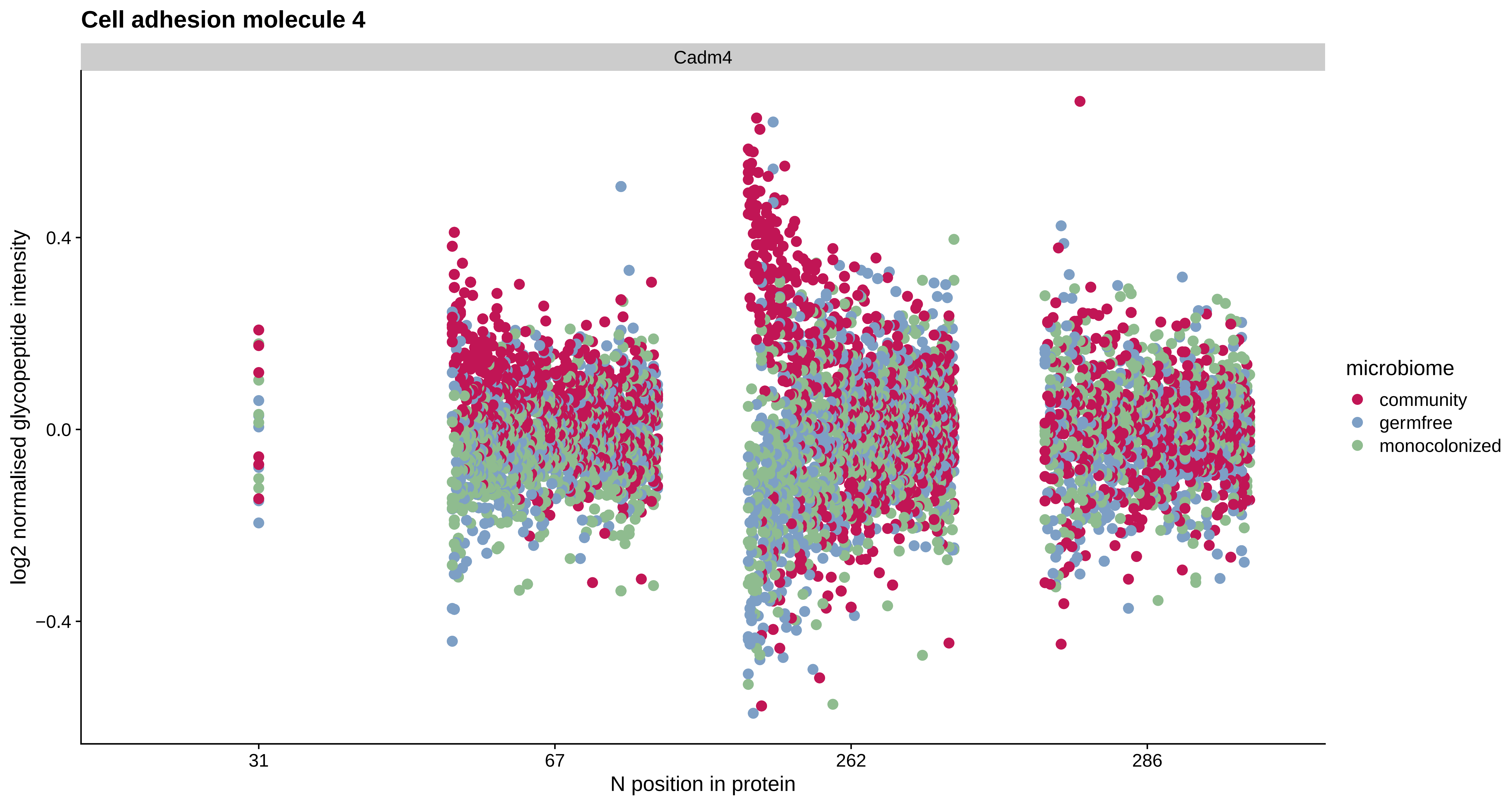

Cell adhesion molecule 4
Cadm4
log2 normalised glycopeptide intensity
0.4
0.0
−0.4
31
67
262
286
N position in protein
microbiome
community
germfree
monocolonized

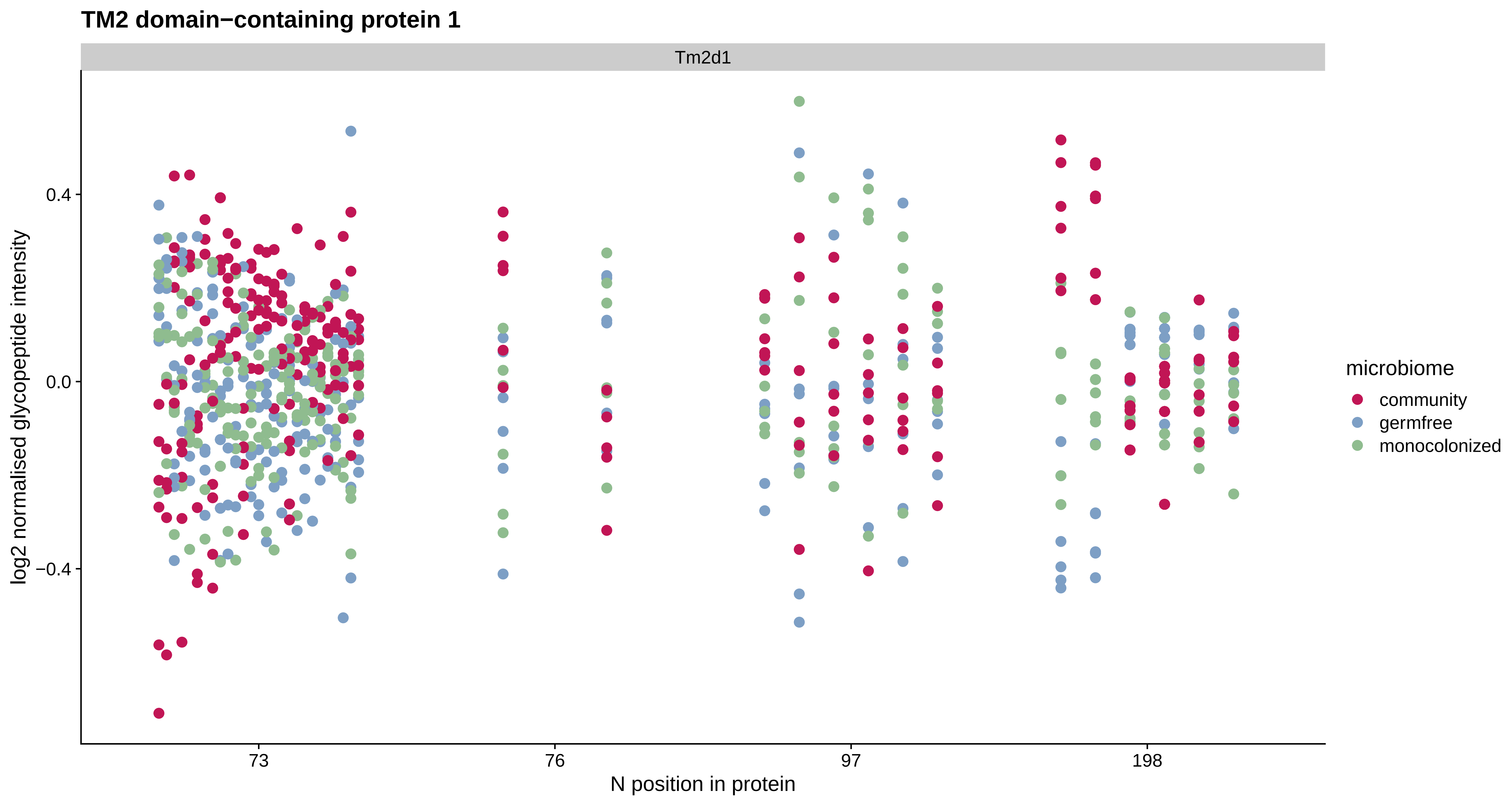
TM2 domain−containing protein 1
Tm2d1
log2 normalised glycopeptide intensity
0.4
0.0
−0.4
73
76
97
198
N position in protein
microbiome
community
germfree
monocolonized

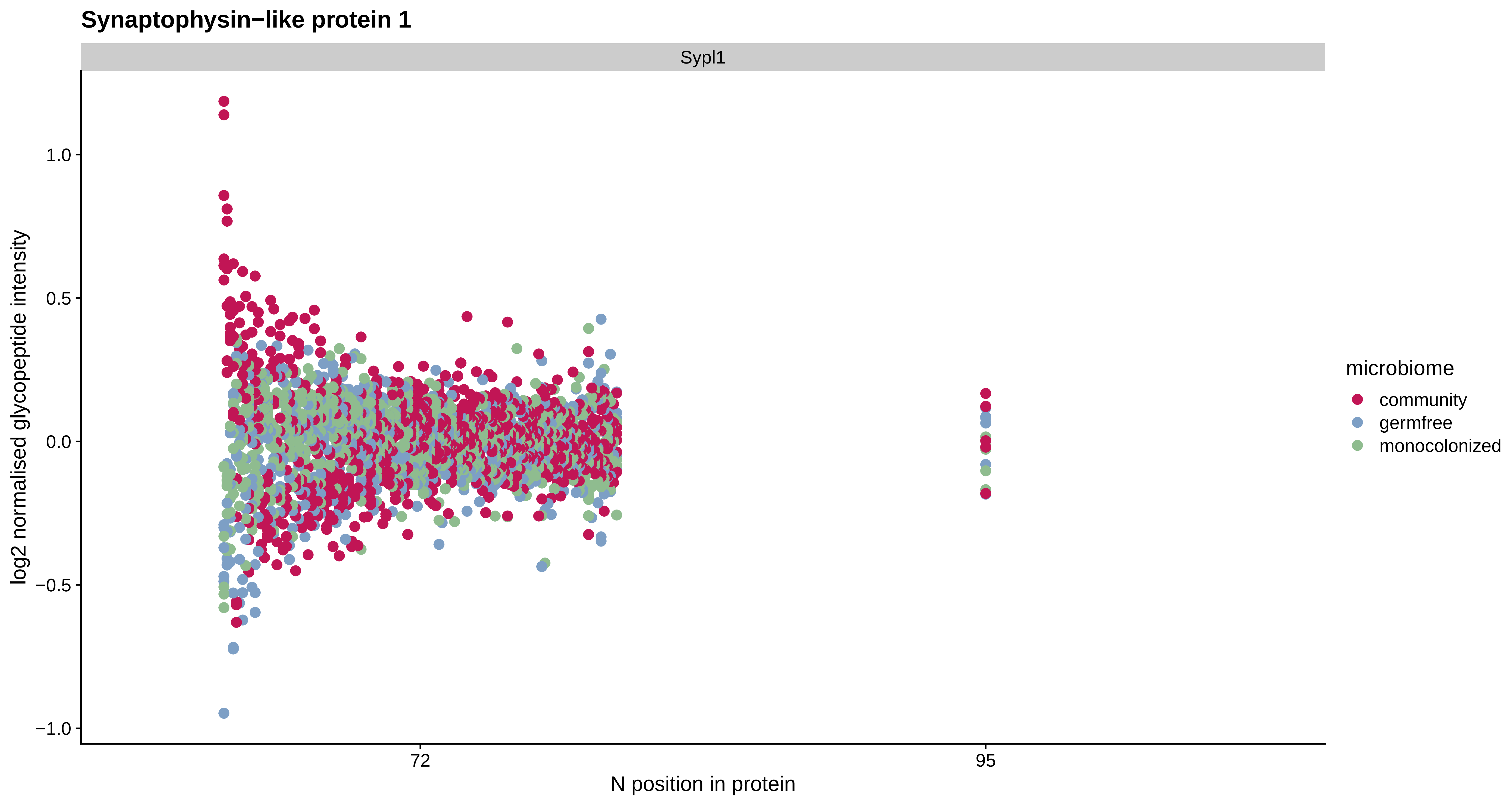
Synaptophysin−like protein 1
Sypl1
log2 normalised glycopeptide intensity
1.0
0.5
0.0
−0.5
−1.0
72
95
N position in protein
microbiome
community
germfree
monocolonized

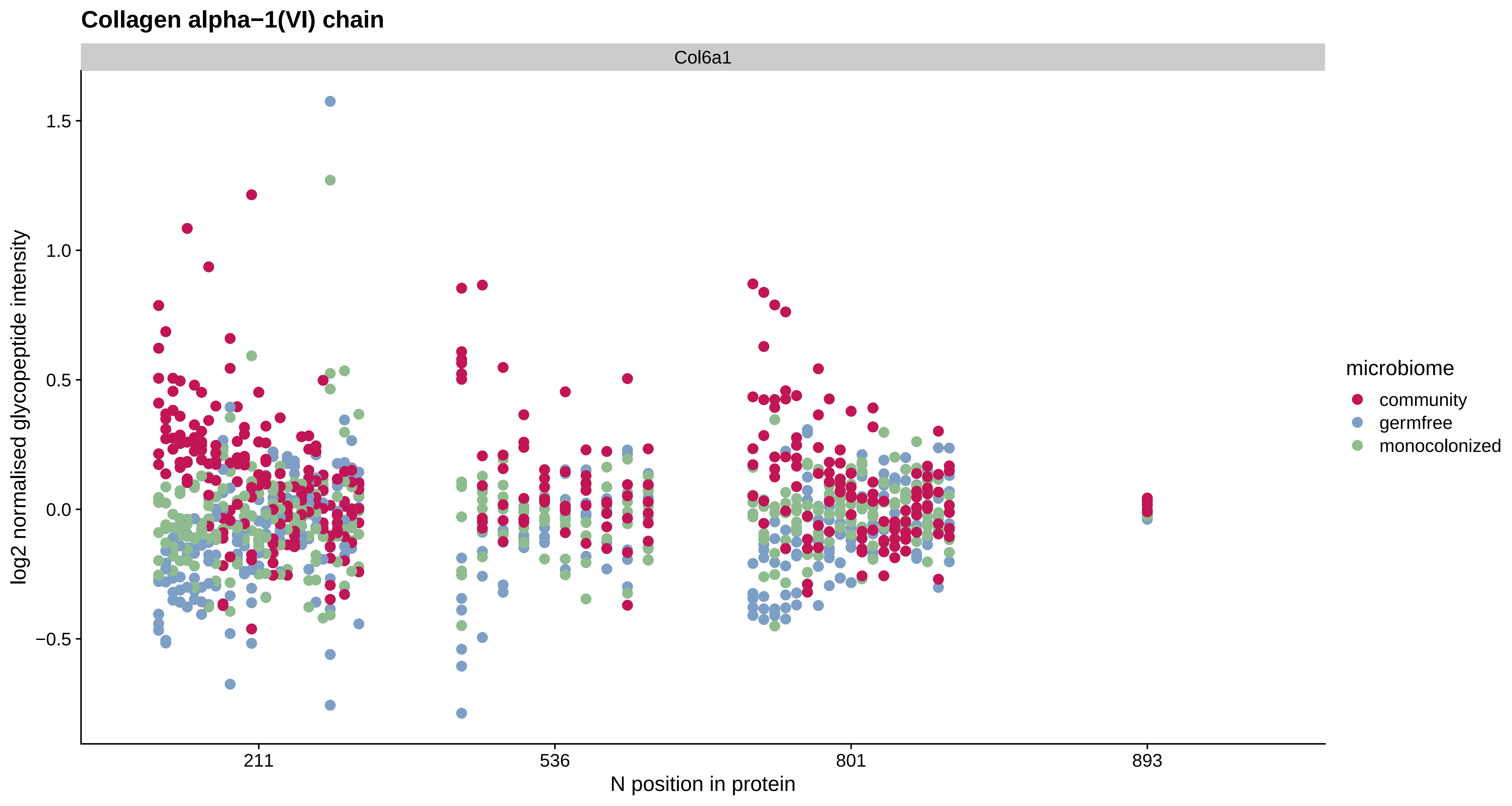

Collagen alpha−1(VI) chain
Col6a1
log2 normalised glycopeptide intensity
1.5
1.0
0.5
0.0
−0.5
211
536
801
893
N position in protein
microbiome
community
germfree
monocolonized

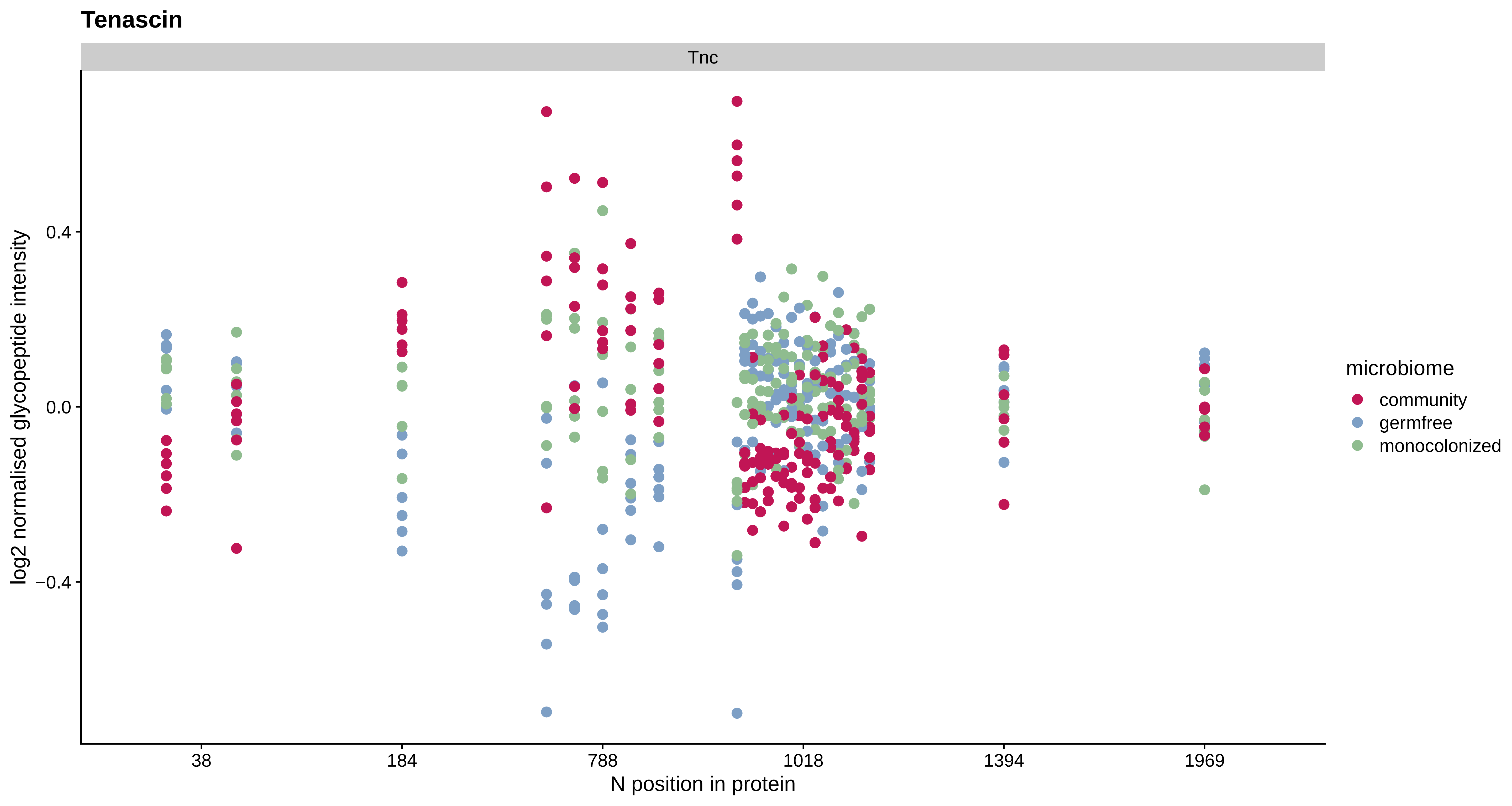

Tenascin
Tnc
log2 normalised glycopeptide intensity
0.4
0.0
−0.4
38
184
788
1018
1394
1969
N position in protein
microbiome
community
germfree
monocolonized

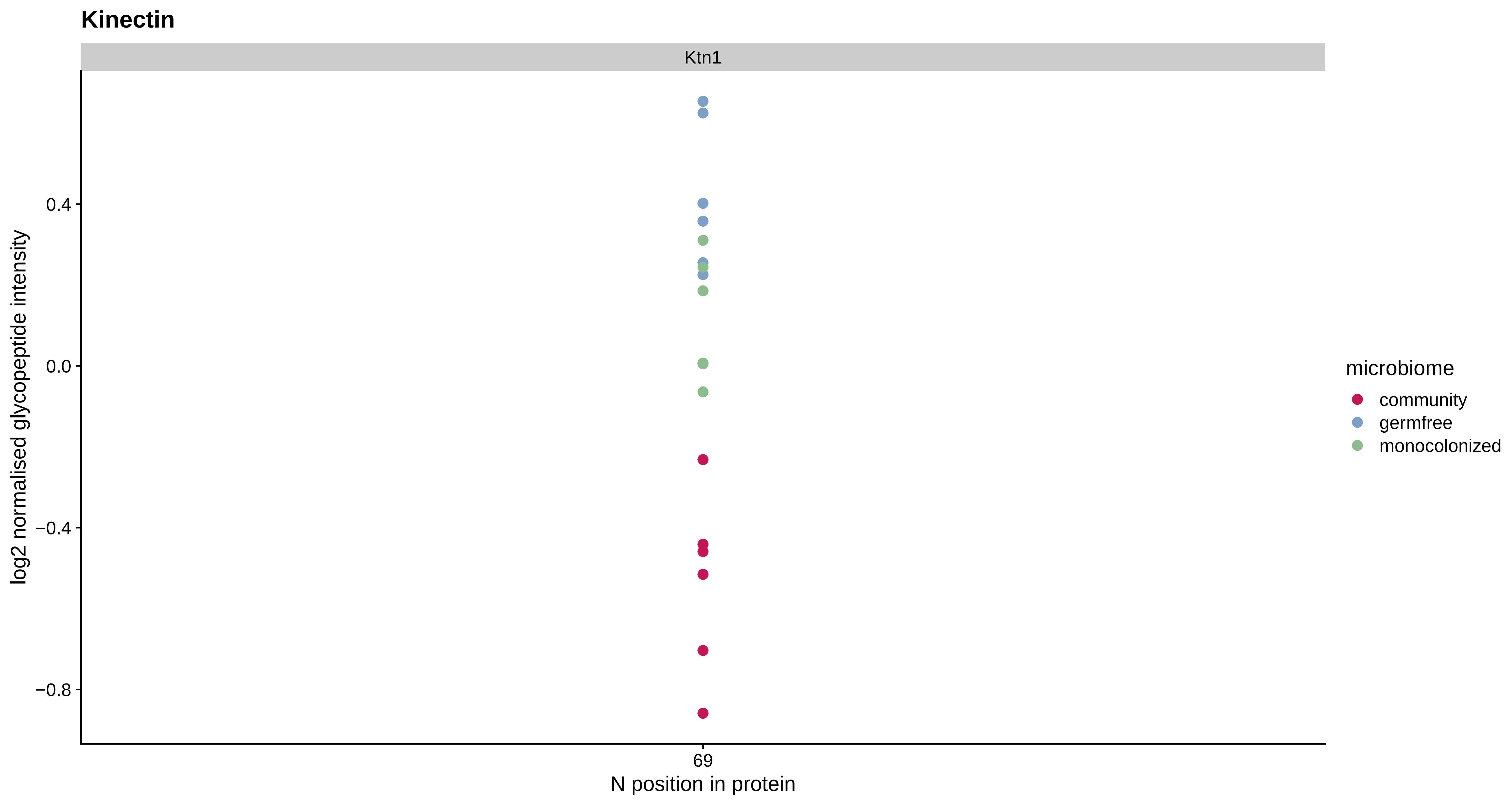

Kinectin
Ktn1
0.4
0.0
−0.4
−0.8
log2 normalised glycopeptide intensity
69
N position in protein
microbiome
community
germfree
monocolonized

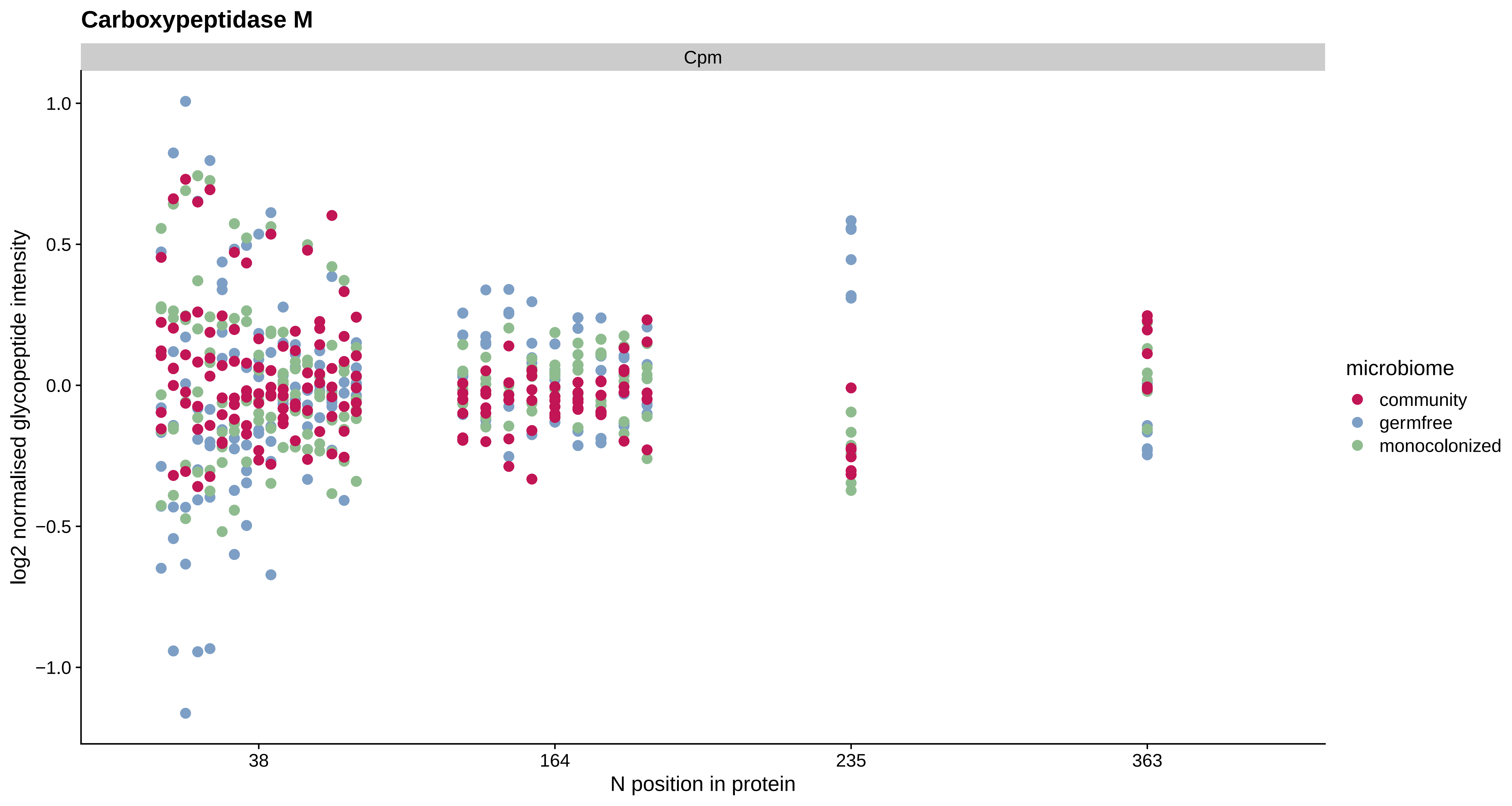
Carboxypeptidase M
Cpm
1.0
0.5
0.0
−0.5
−1.0
log2 normalised glycopeptide intensity
38
164
235
363
N position in protein
microbiome
community
germfree
monocolonized

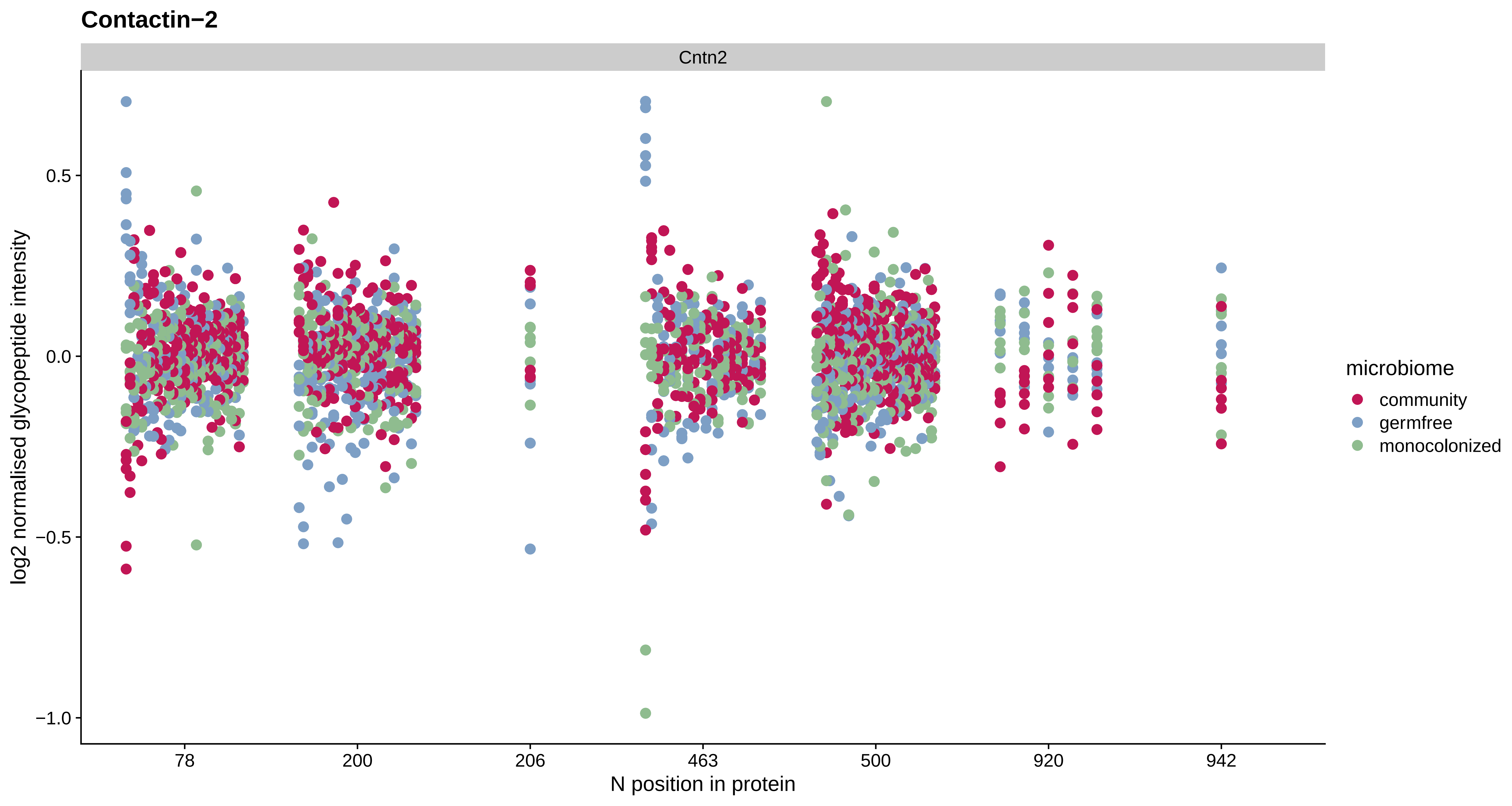
Contactin−2
Cntn2
log2 normalised glycopeptide intensity
0.5
0.0
−0.5
−1.0
78
200
206
463
500
920
942
N position in protein
microbiome
community
germfree
monocolonized

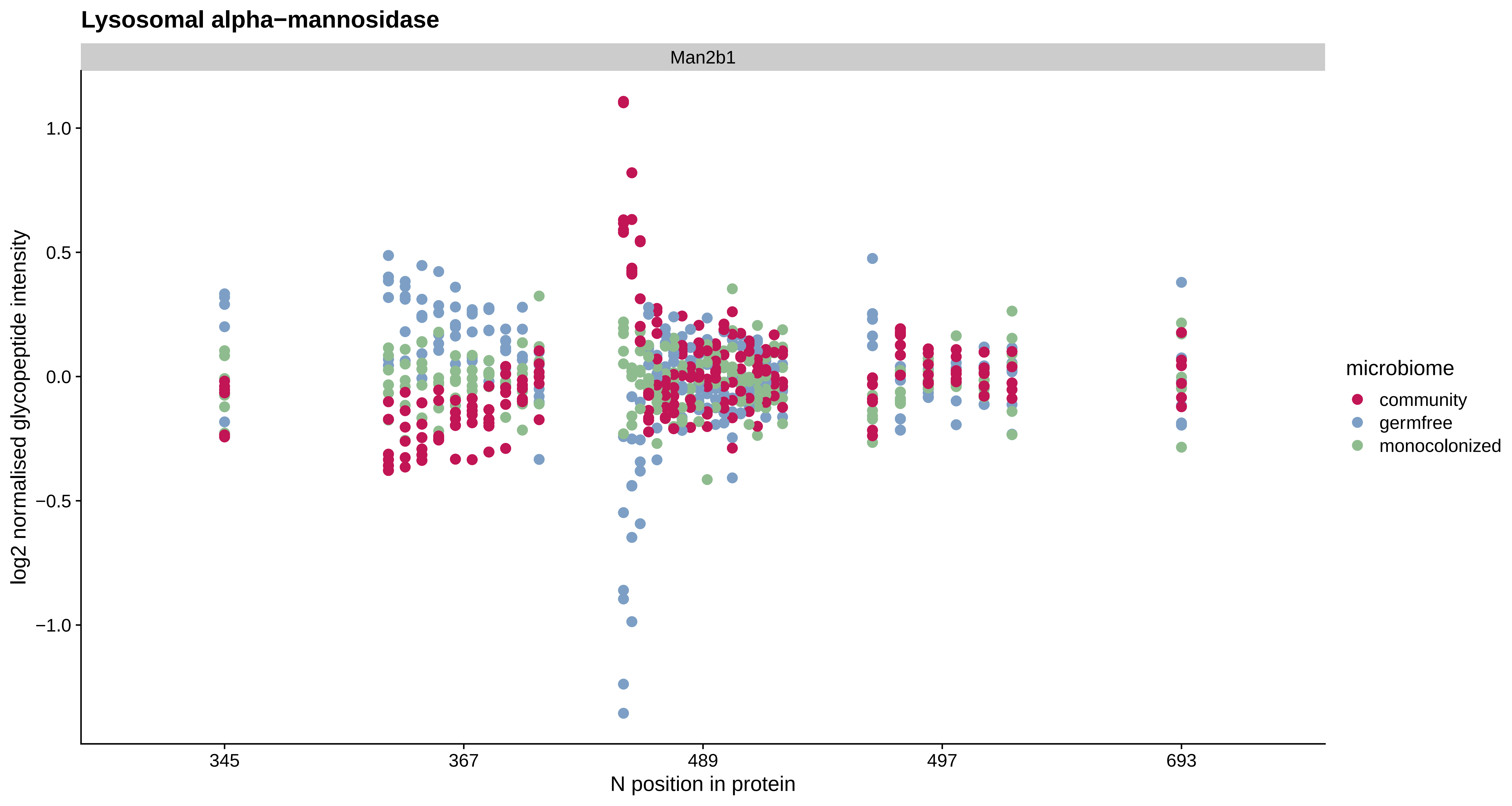

Lysosomal alpha−mannosidase
Man2b1
log2 normalised glycopeptide intensity
1.0
0.5
0.0
−0.5
−1.0
345
367
489
497
693
N position in protein
microbiome
community
germfree
monocolonized

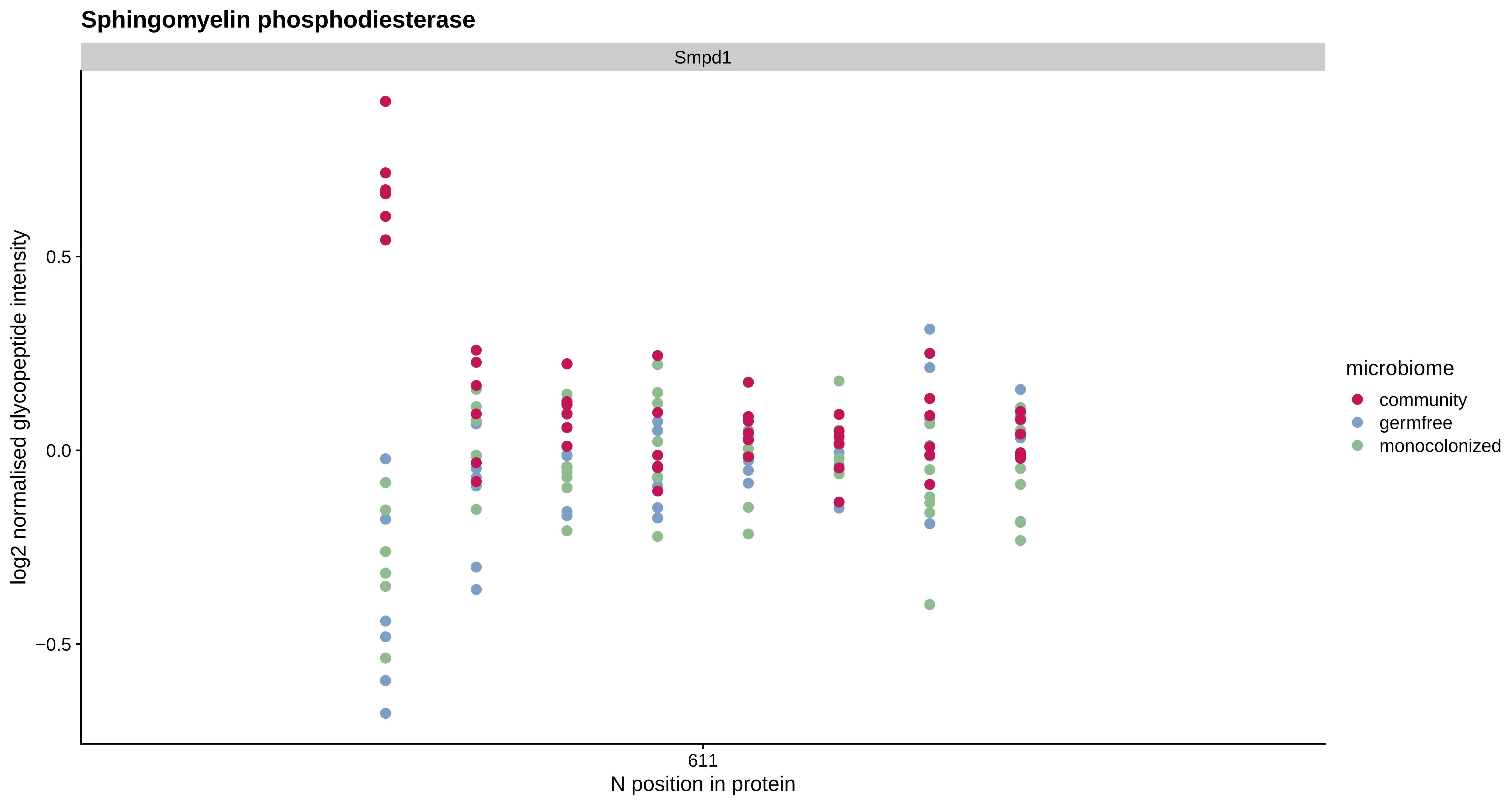

Sphingomyelin phosphodiesterase
Smpd1
log2 normalised glycopeptide intensity
0.5
0.0
−0.5
611
N position in protein
microbiome
community
germfree
monocolonized

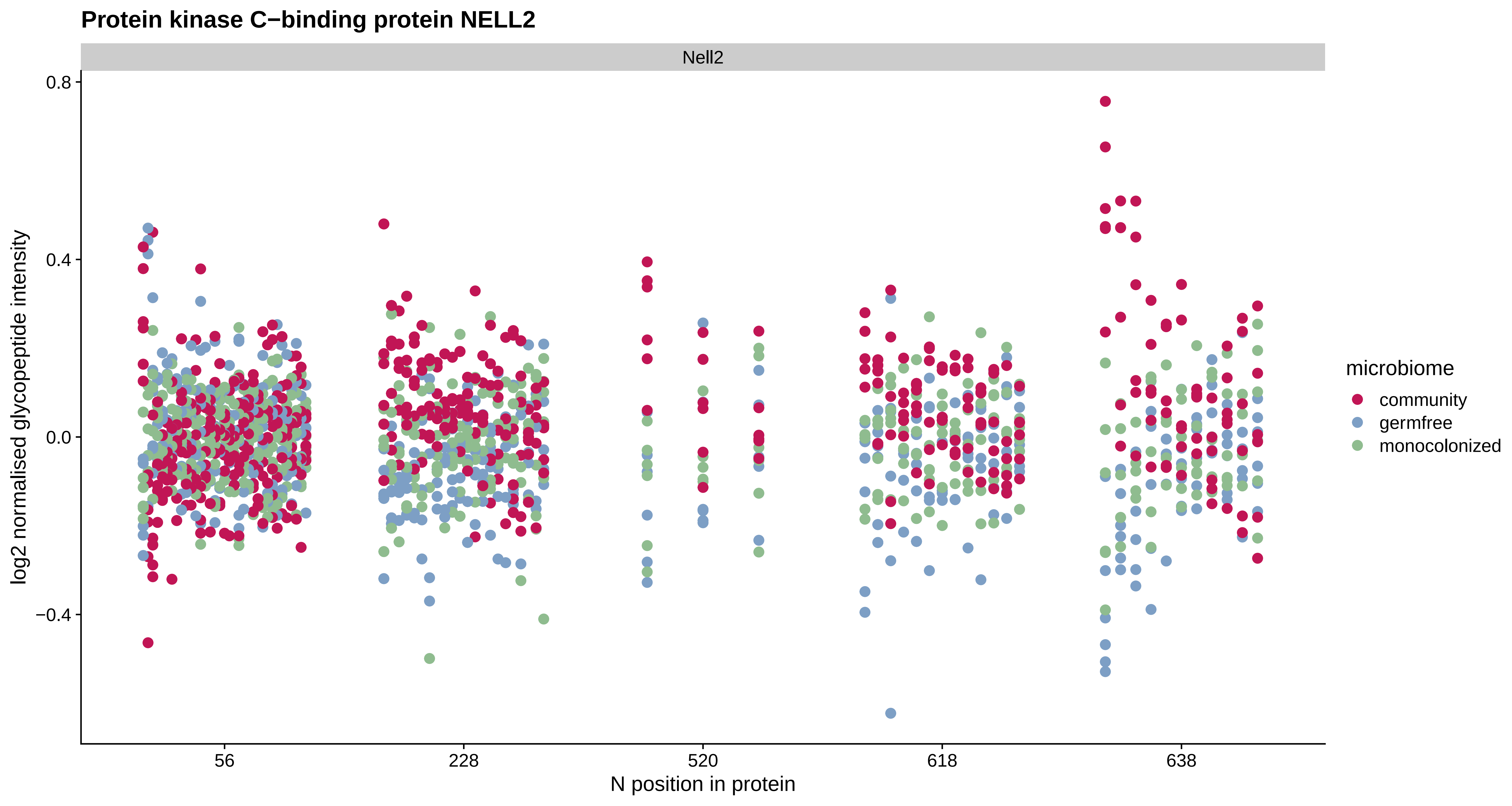

Protein kinase C−binding protein NELL2
Nell2
0.8
0.4
0.0
−0.4
log2 normalised glycopeptide intensity
56
228
520
618
638
N position in protein
microbiome
community
germfree
monocolonized

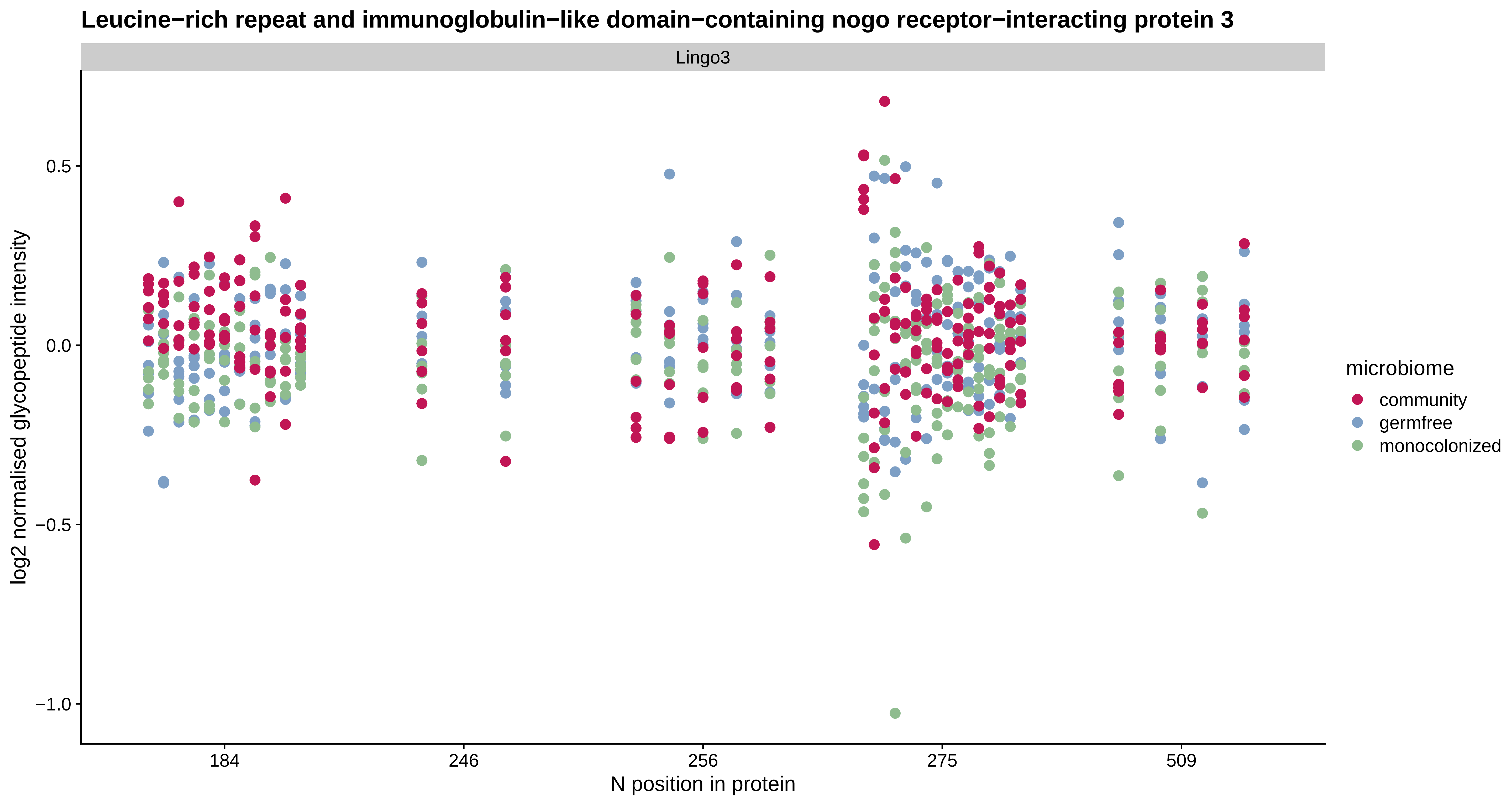

Leucine−rich repeat and immunoglobulin−like domain−containing nogo receptor−interacting protein 3
Lingo3
log2 normalised glycopeptide intensity
0.5
0.0
−0.5
−1.0
184
246
256
275
509
N position in protein
microbiome
community
germfree
monocolonized

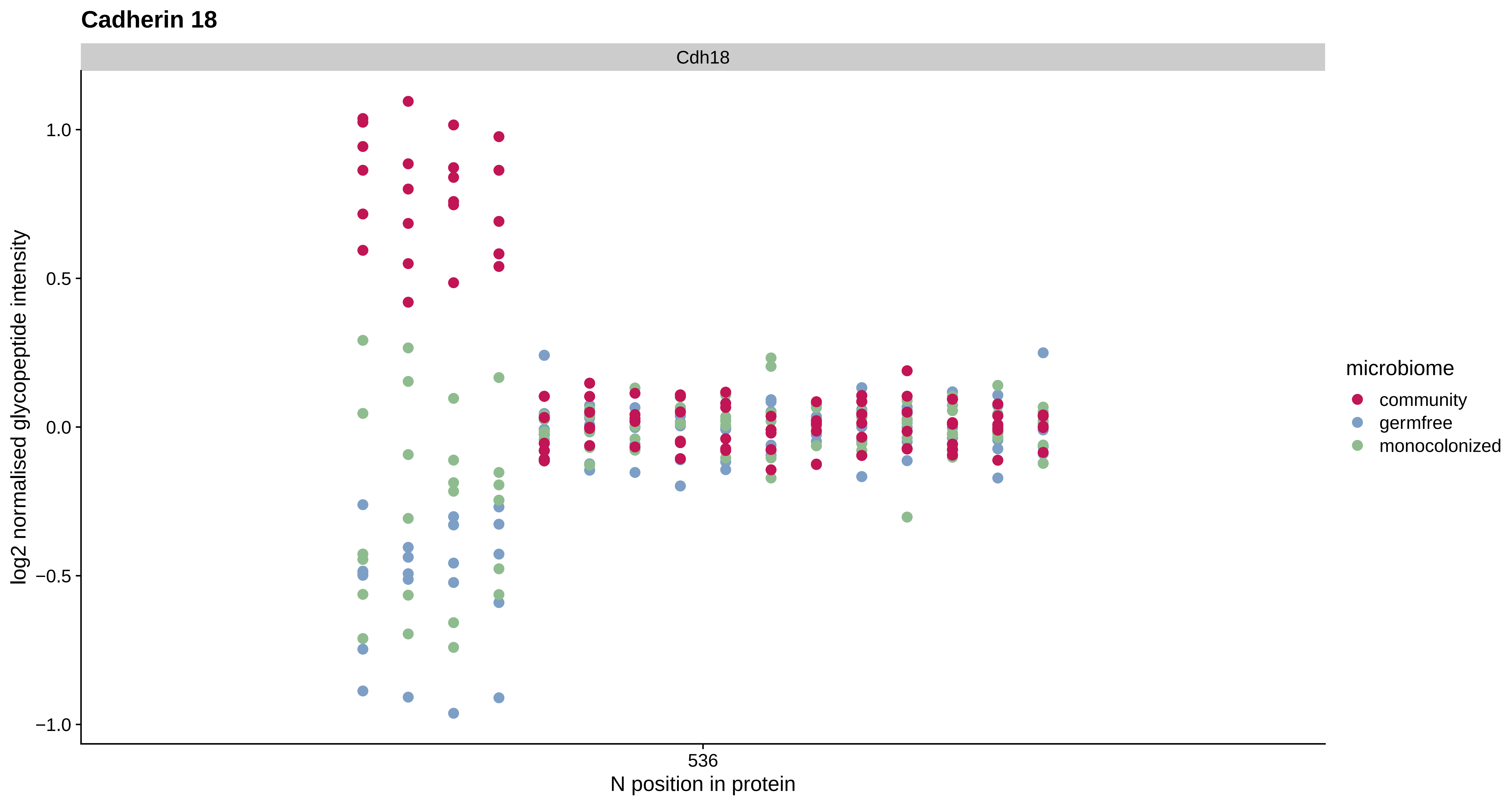

Cadherin 18
Cdh18
log2 normalised glycopeptide intensity
1.0
0.5
0.0
−0.5
−1.0
536
N position in protein
microbiome
community
germfree
monocolonized

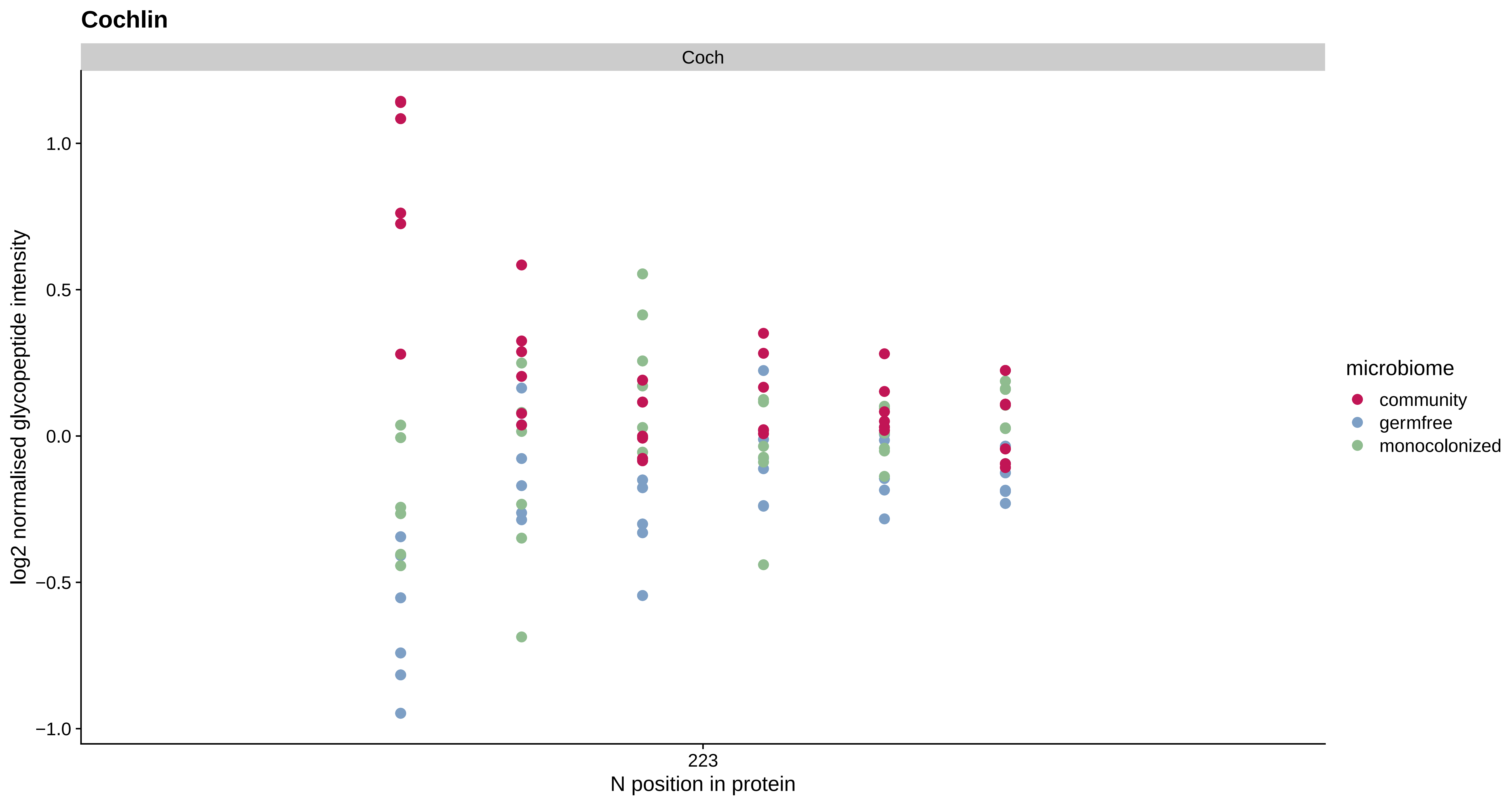

Cochlin
Coch
log2 normalised glycopeptide intensity
1.0
0.5
0.0
−0.5
−1.0
223
N position in protein
microbiome
community
germfree
monocolonized

#### Rap guanine nucleotide exchange factor (GEF) 6

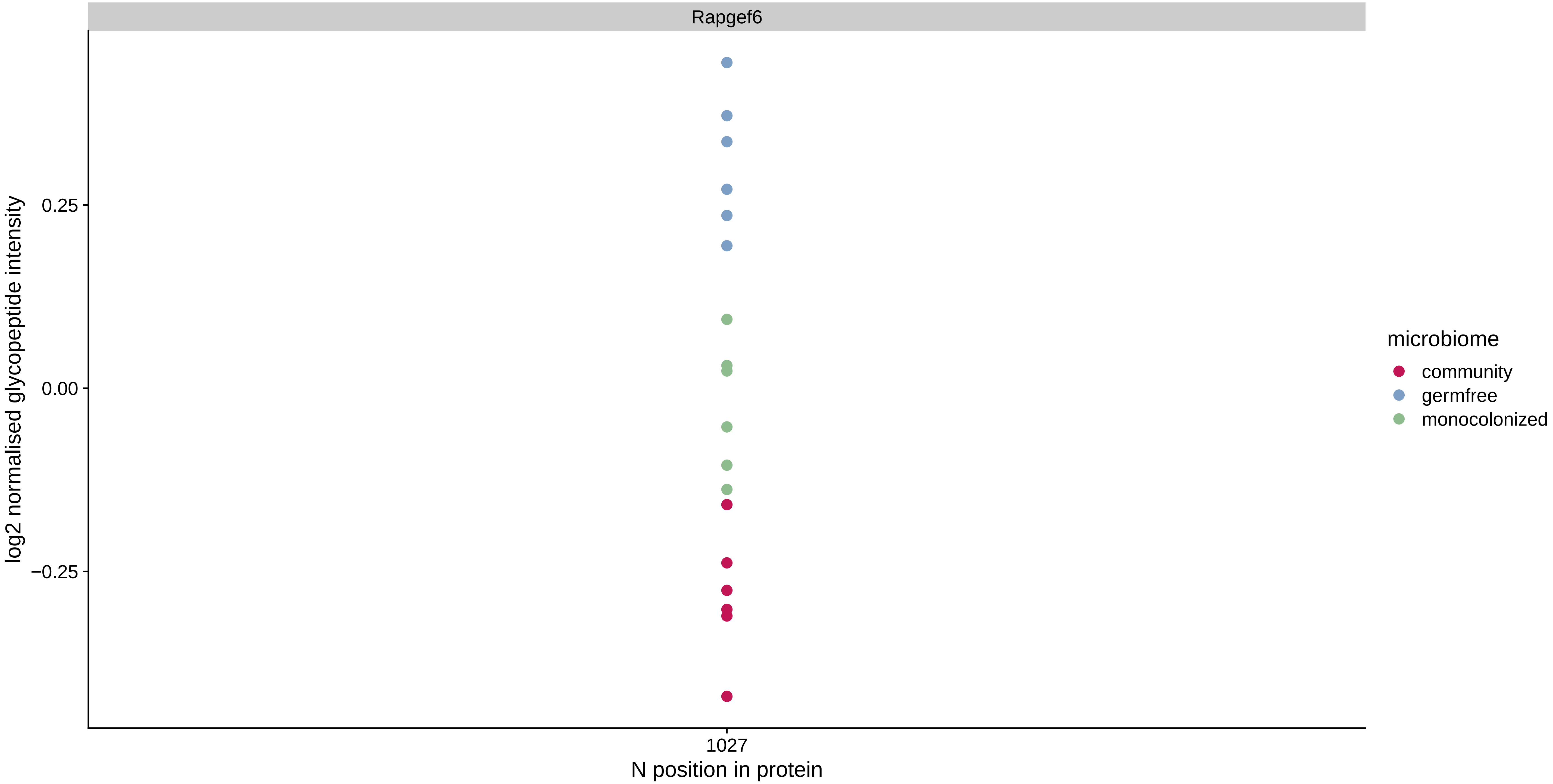

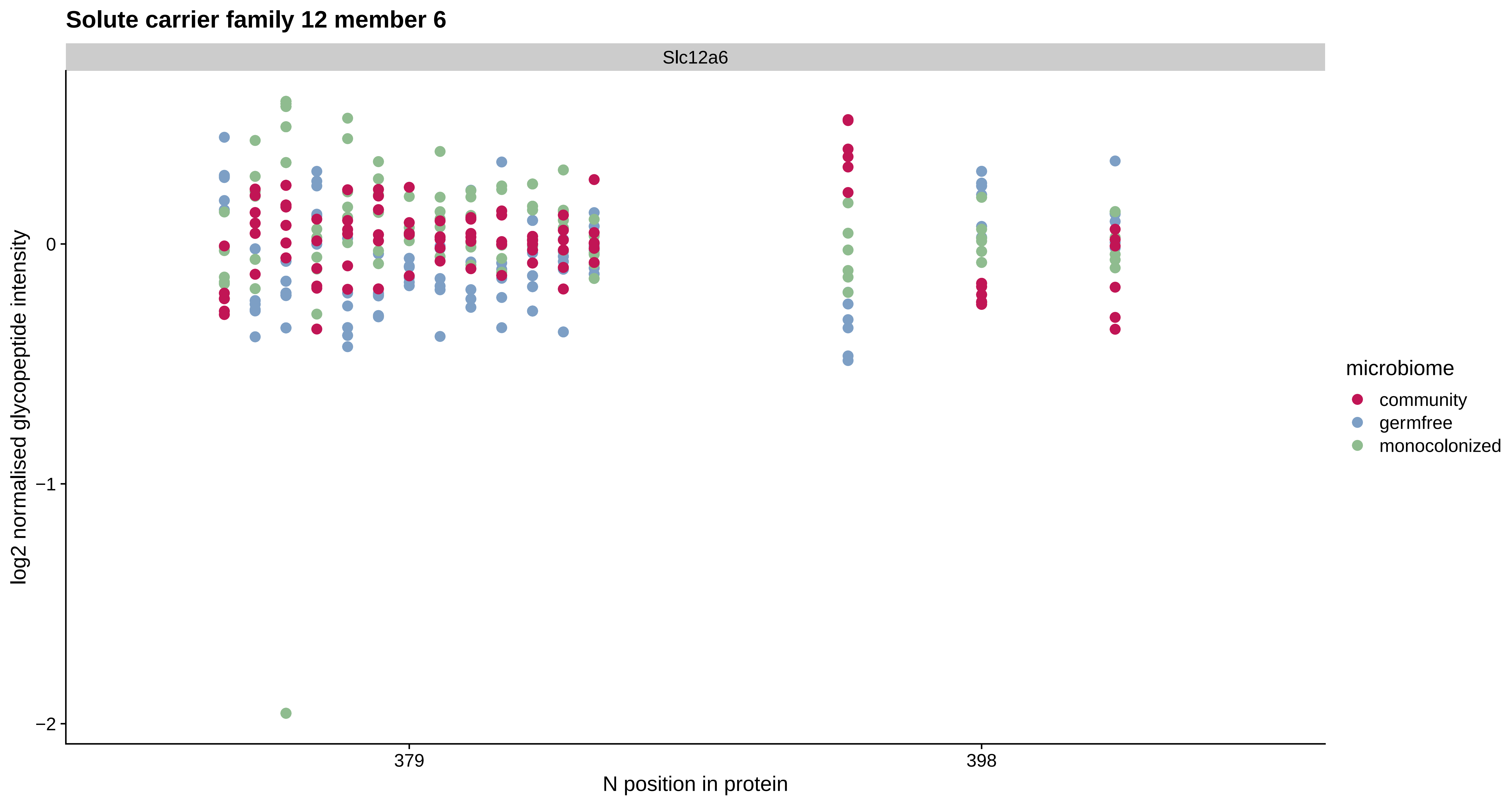

Solute carrier family 12 member 6
Slc12a6
log2 normalised glycopeptide intensity
0
−1
−2
379
398
N position in protein
microbiome
community
germfree
monocolonized

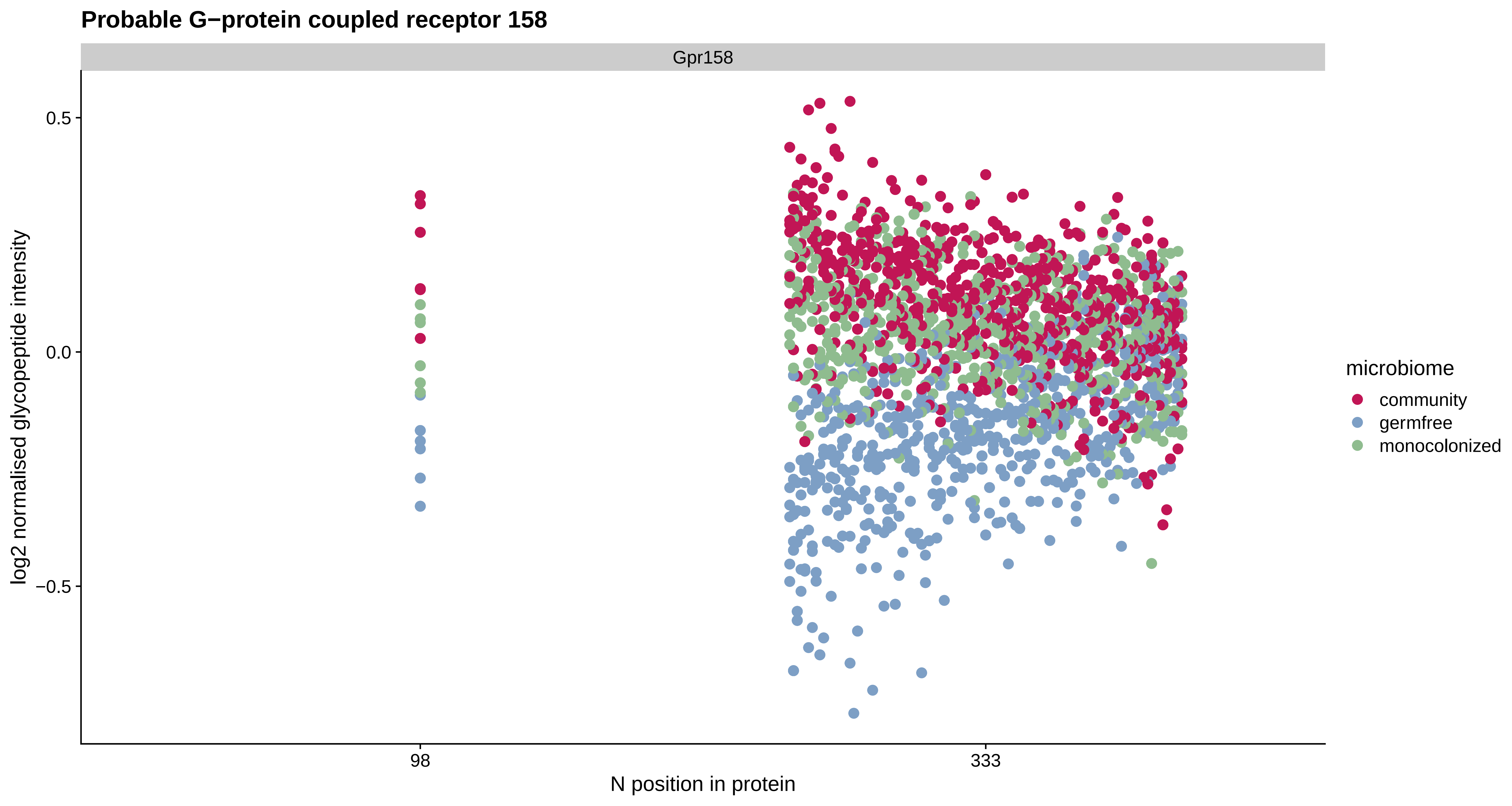

Probable G−protein coupled receptor 158
Gpr158
0.5
0.0
−0.5
log2 normalised glycopeptide intensity
98
333
N position in protein
microbiome
community
germfree
monocolonized

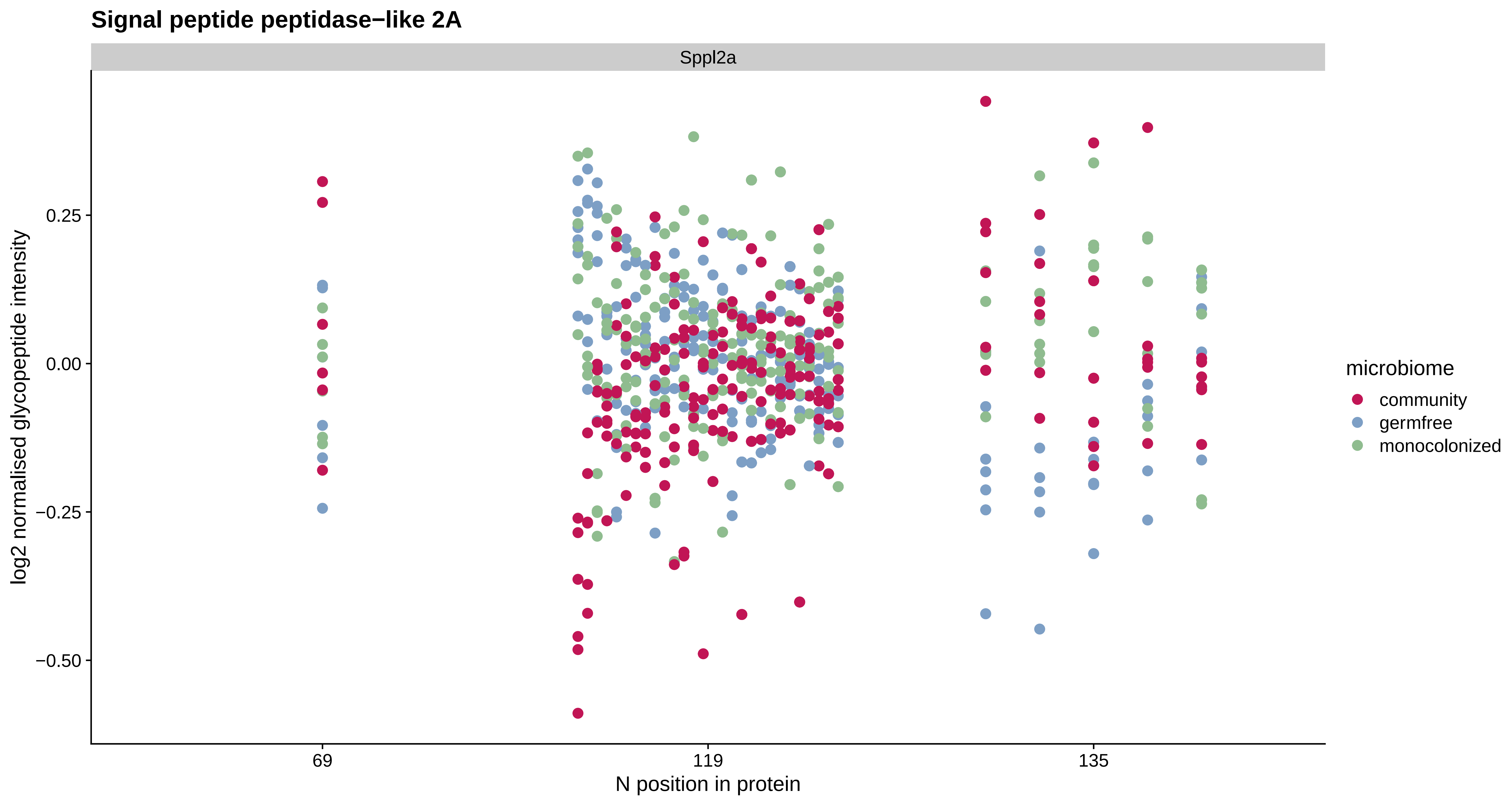

Signal peptide peptidase−like 2A
Sppl2a
log2 normalised glycopeptide intensity
0.25
0.00
−0.25
−0.50
69
119
135
N position in protein
microbiome
community
germfree
monocolonized

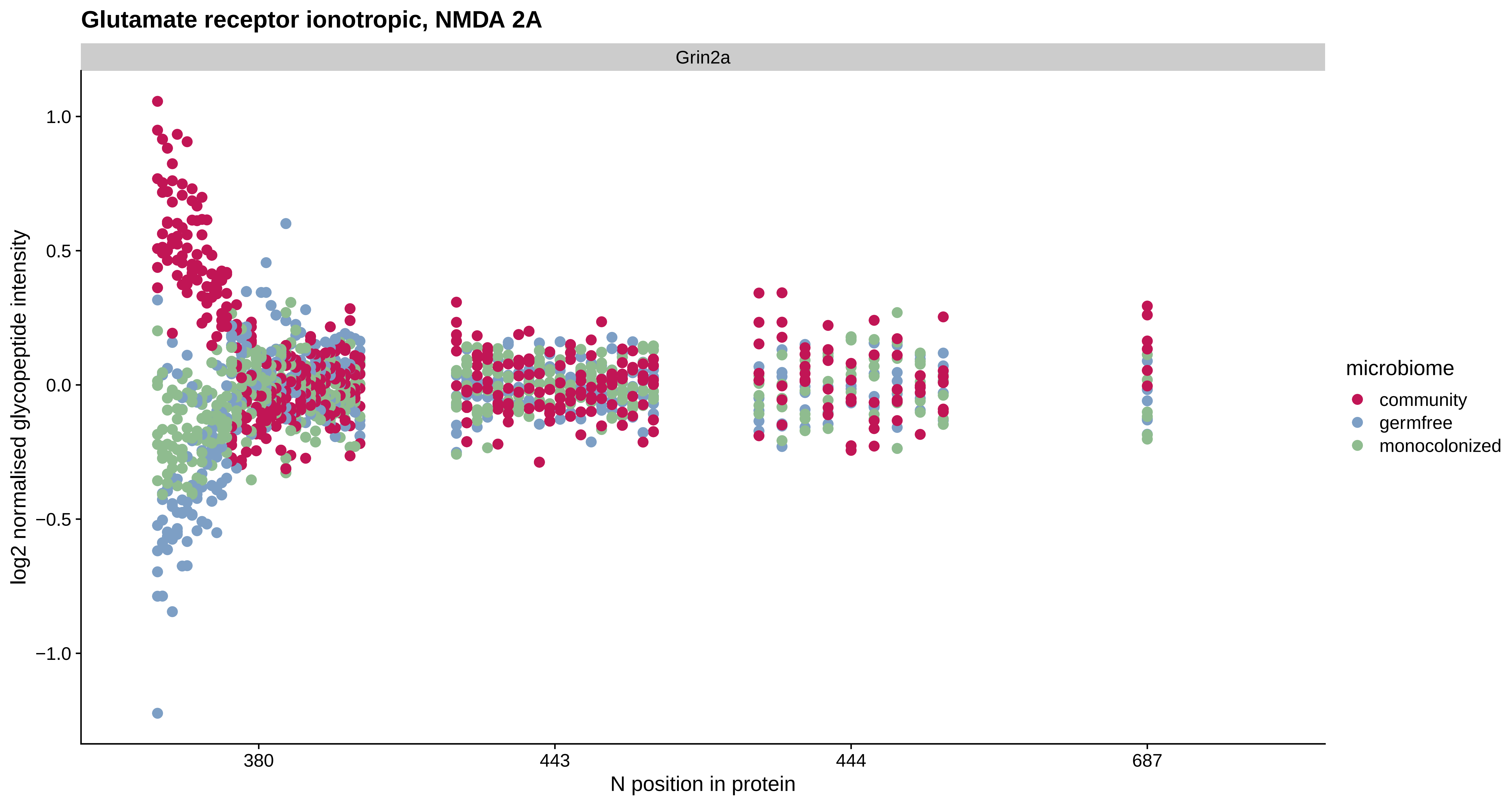
Glutamate receptor ionotropic, NMDA 2A
Grin2a
log2 normalised glycopeptide intensity
1.0
0.5
0.0
−0.5
−1.0
380
443
444
687
N position in protein
microbiome
community
germfree
monocolonized

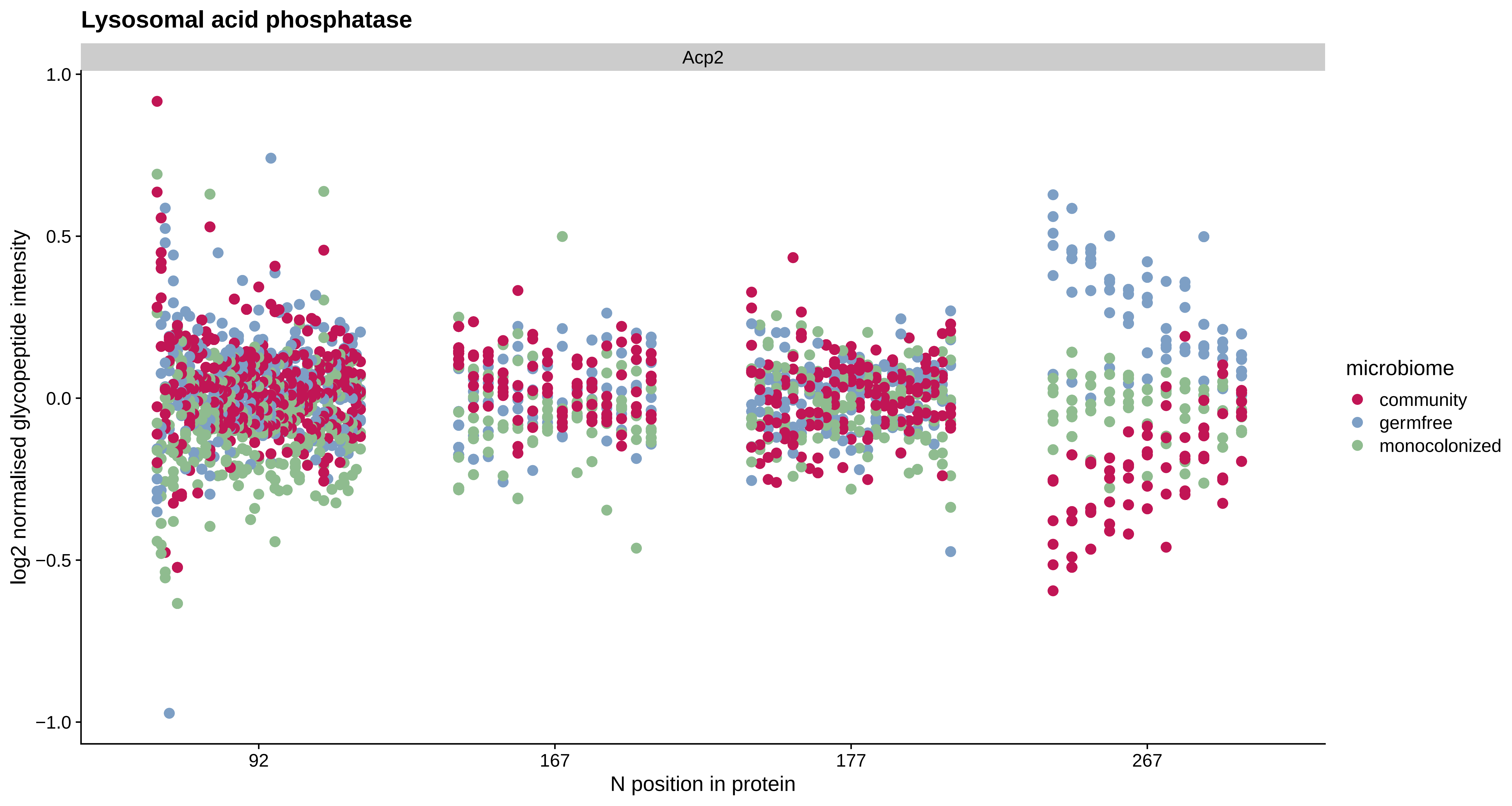

Lysosomal acid phosphatase
Acp2
1.0
0.5
0.0
−0.5
−1.0
log2 normalised glycopeptide intensity
92
167
177
267
N position in protein
microbiome
community
germfree
monocolonized

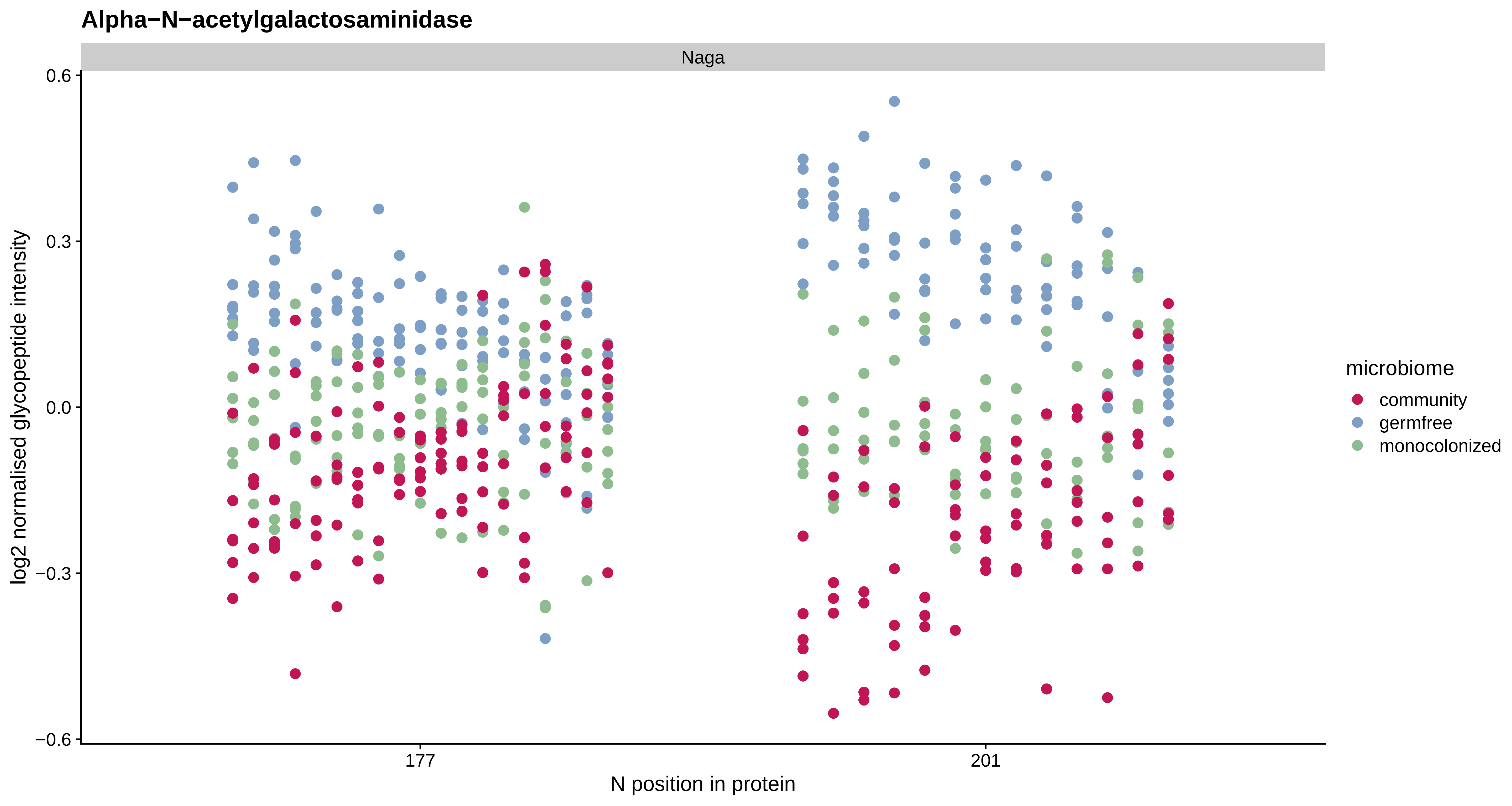

Alpha−N−acetylgalactosaminidase
Naga
log2 normalised glycopeptide intensity
0.6
0.3
0.0
−0.3
−0.6
177
201
N position in protein
microbiome
community
germfree
monocolonized

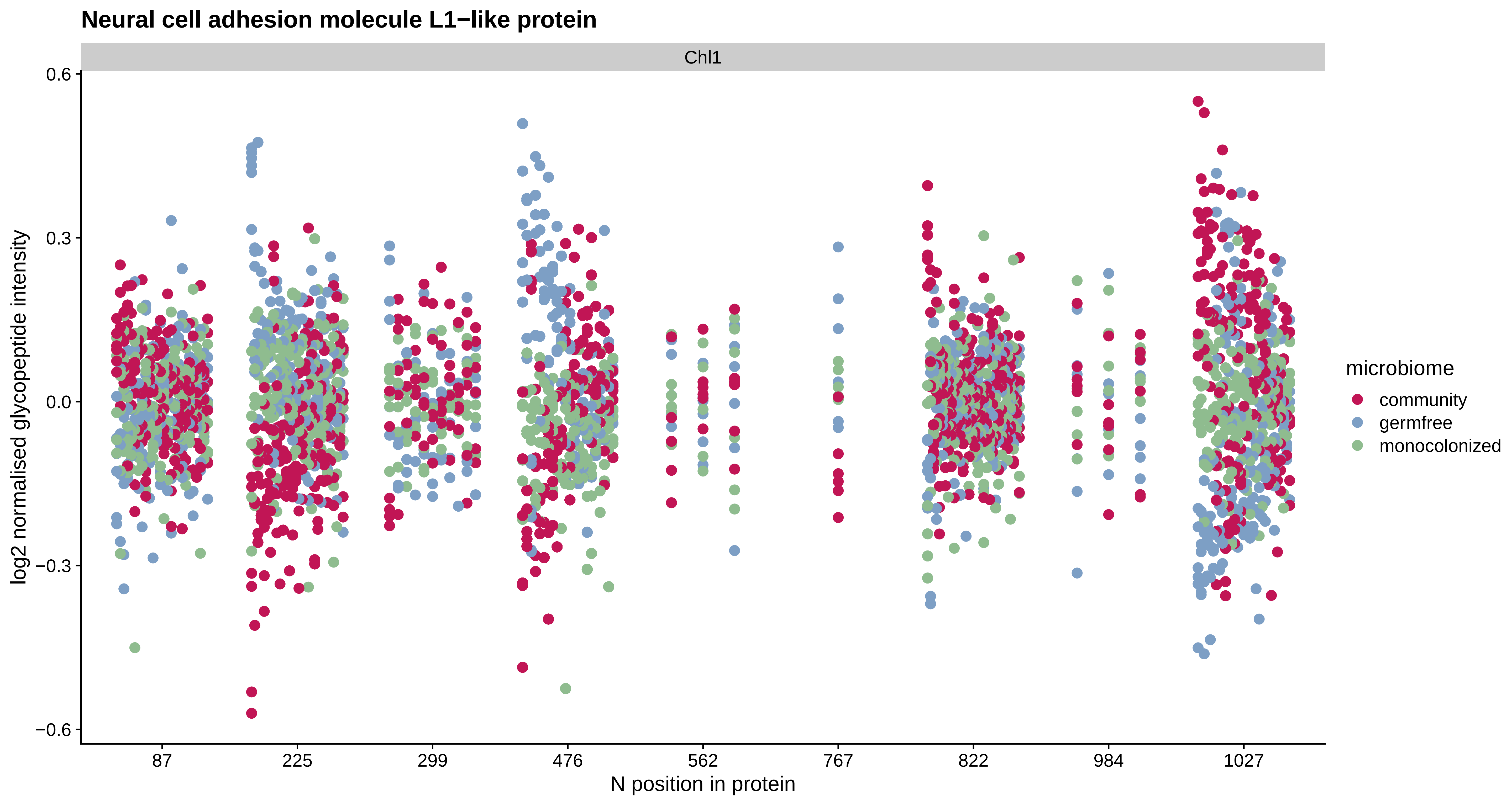
Neural cell adhesion molecule L1−like protein
Chl1
log2 normalised glycopeptide intensity
0.6
0.3
0.0
−0.3
−0.6
87
225
299
476
562
767
822
984
1027
N position in protein
microbiome
community
germfree
monocolonized

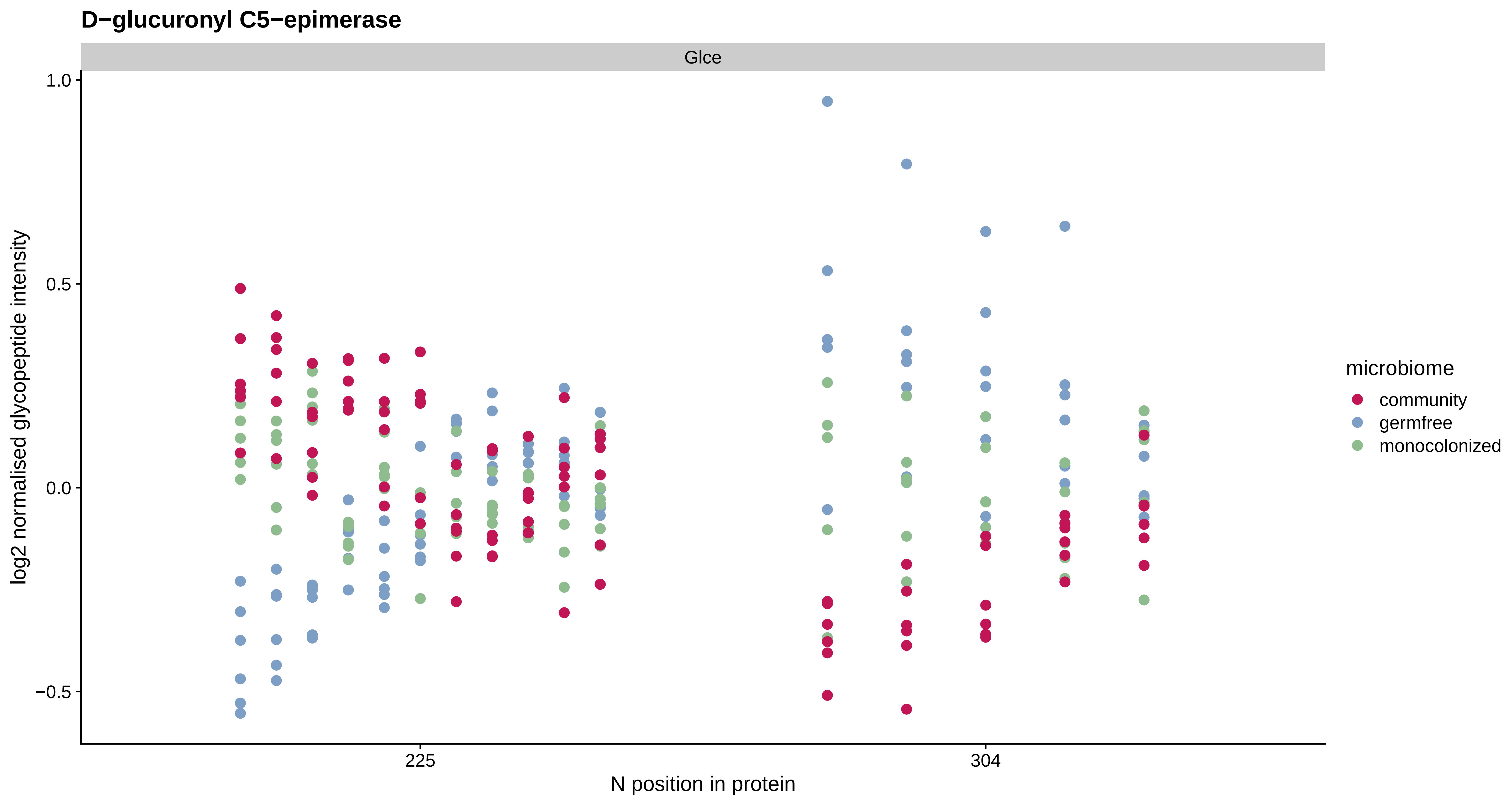
D−glucuronyl C5−epimerase
Glce
1.0
0.5
0.0
−0.5
log2 normalised glycopeptide intensity
225
304
N position in protein
microbiome
community
germfree
monocolonized

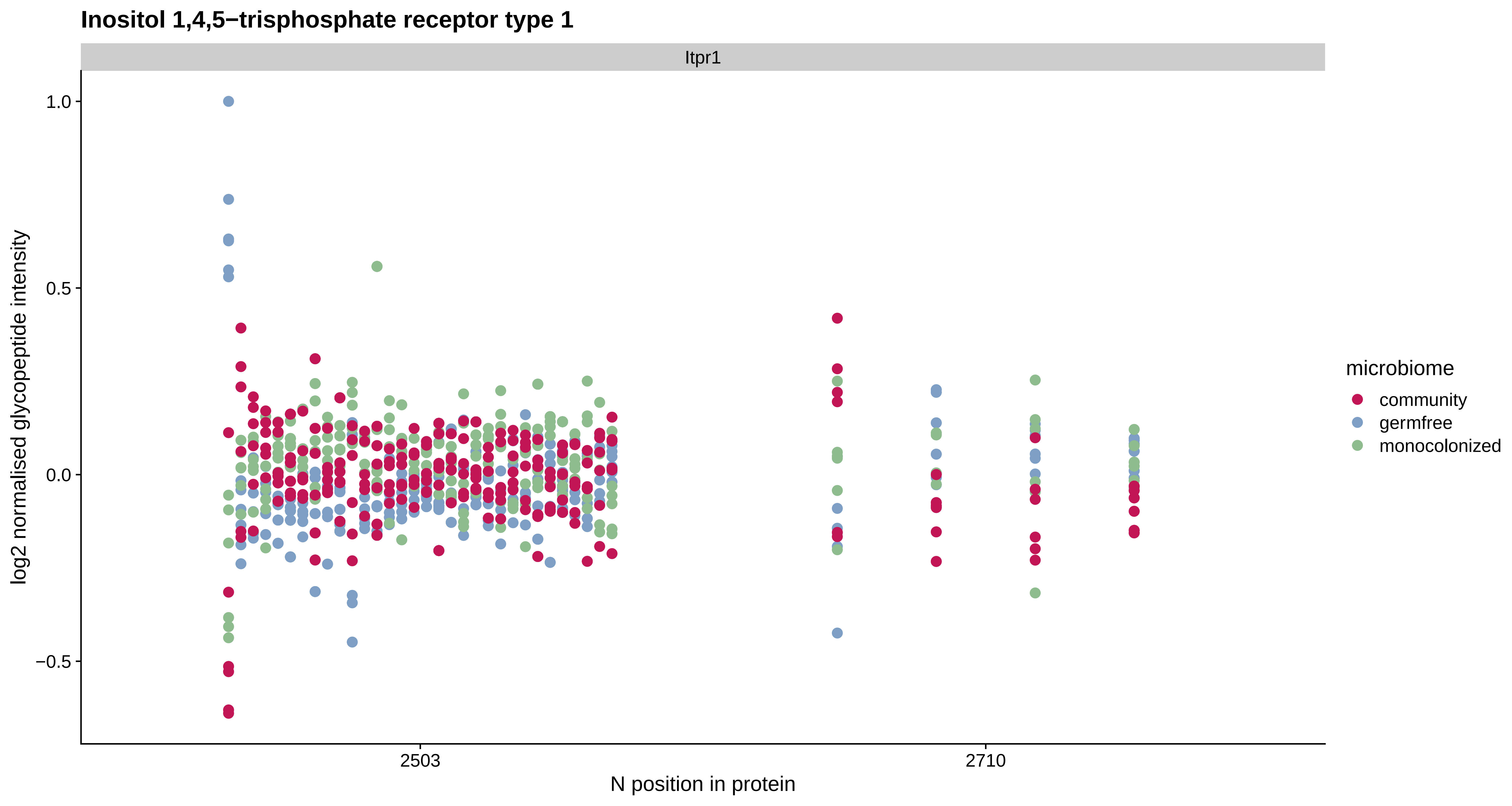
Inositol 1,4,5−trisphosphate receptor type 1
Itpr1
log2 normalised glycopeptide intensity
1.0
0.5
0.0
−0.5
2503
2710
N position in protein
microbiome
community
germfree
monocolonized

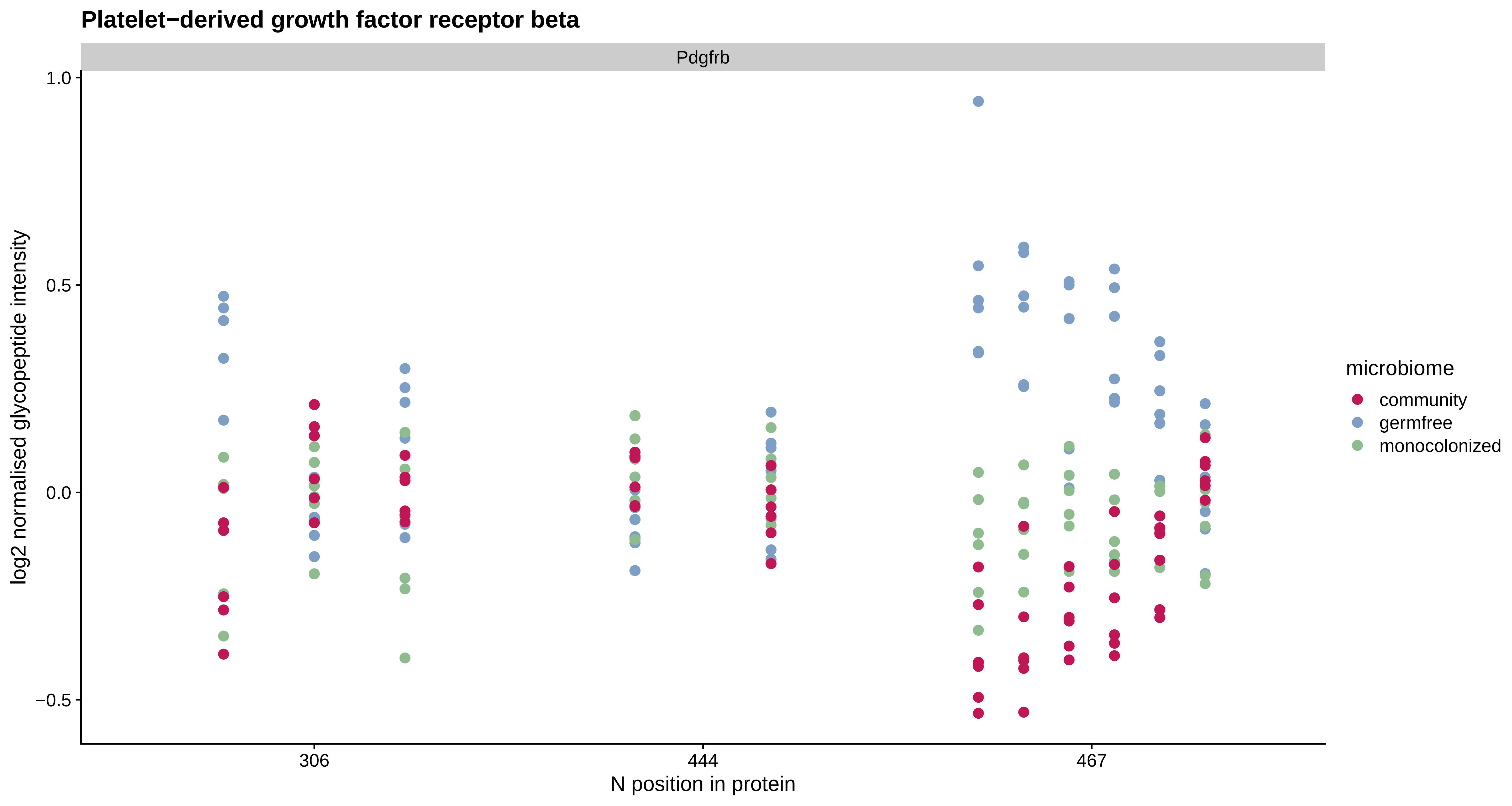

Platelet−derived growth factor receptor beta
Pdgfrb
1.0
0.5
0.0
−0.5
log2 normalised glycopeptide intensity
306
444
467
N position in protein
microbiome
community
germfree
monocolonized

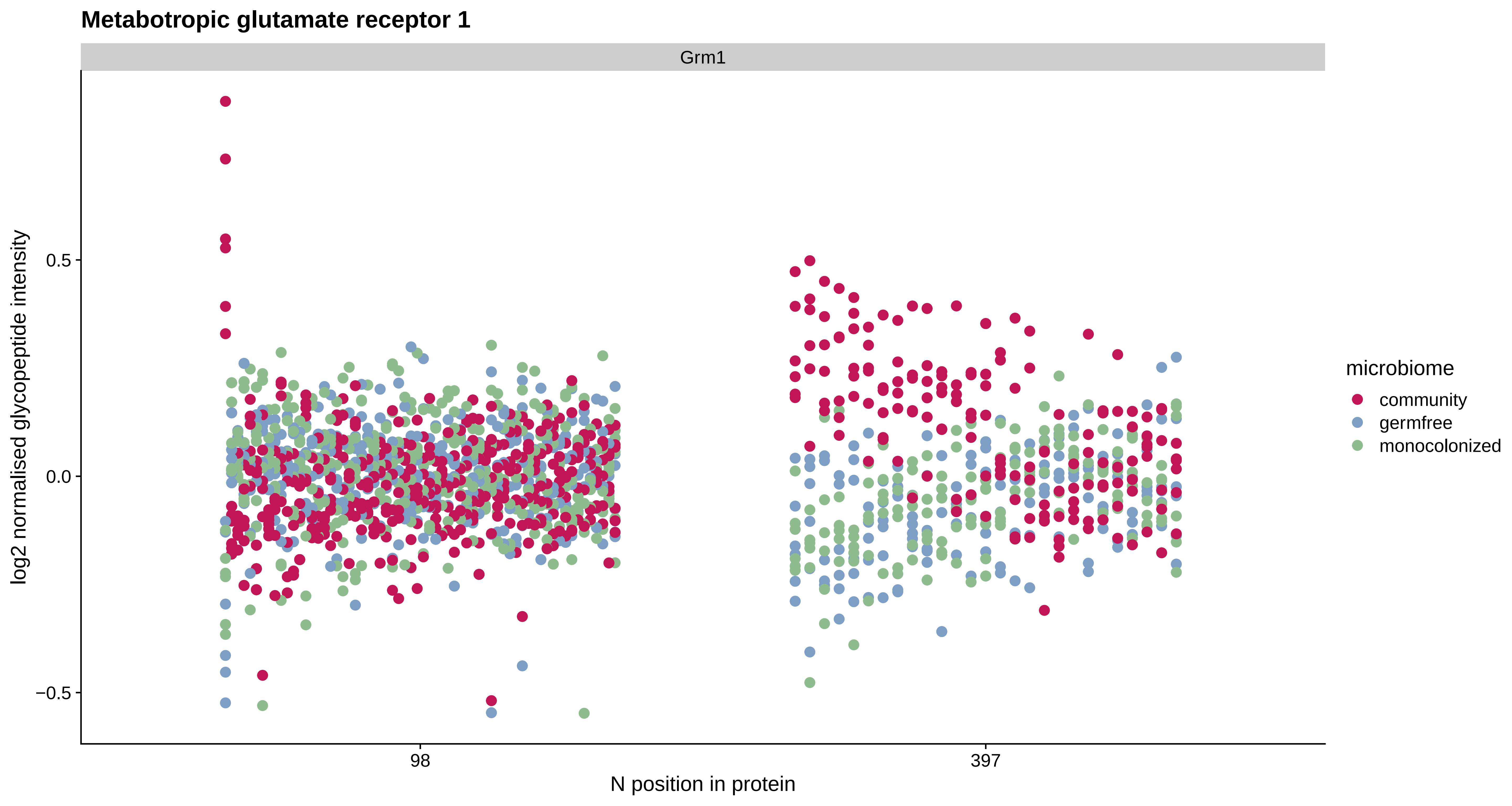

Metabotropic glutamate receptor 1
Grm1
log2 normalised glycopeptide intensity
0.5
0.0
−0.5
98
397
N position in protein
microbiome
community
germfree
monocolonized

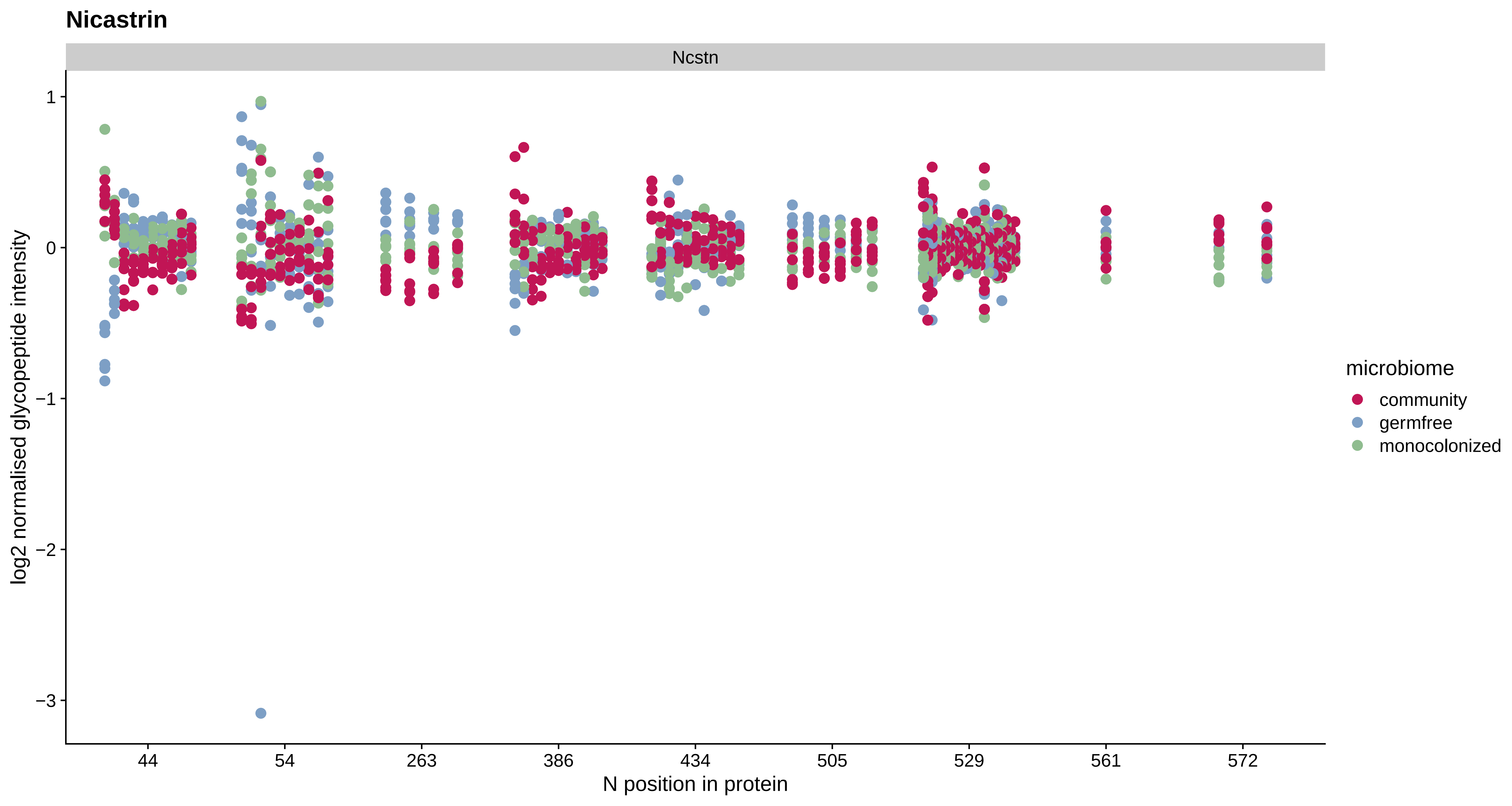
Nicastrin
Ncstn
1
0
−1
−2
−3
log2 normalised glycopeptide intensity
44
54
263
386
434
505
529
561
572
N position in protein
microbiome
community
germfree
monocolonized

#### Multiple inositol polyphosphate phosphatase 1

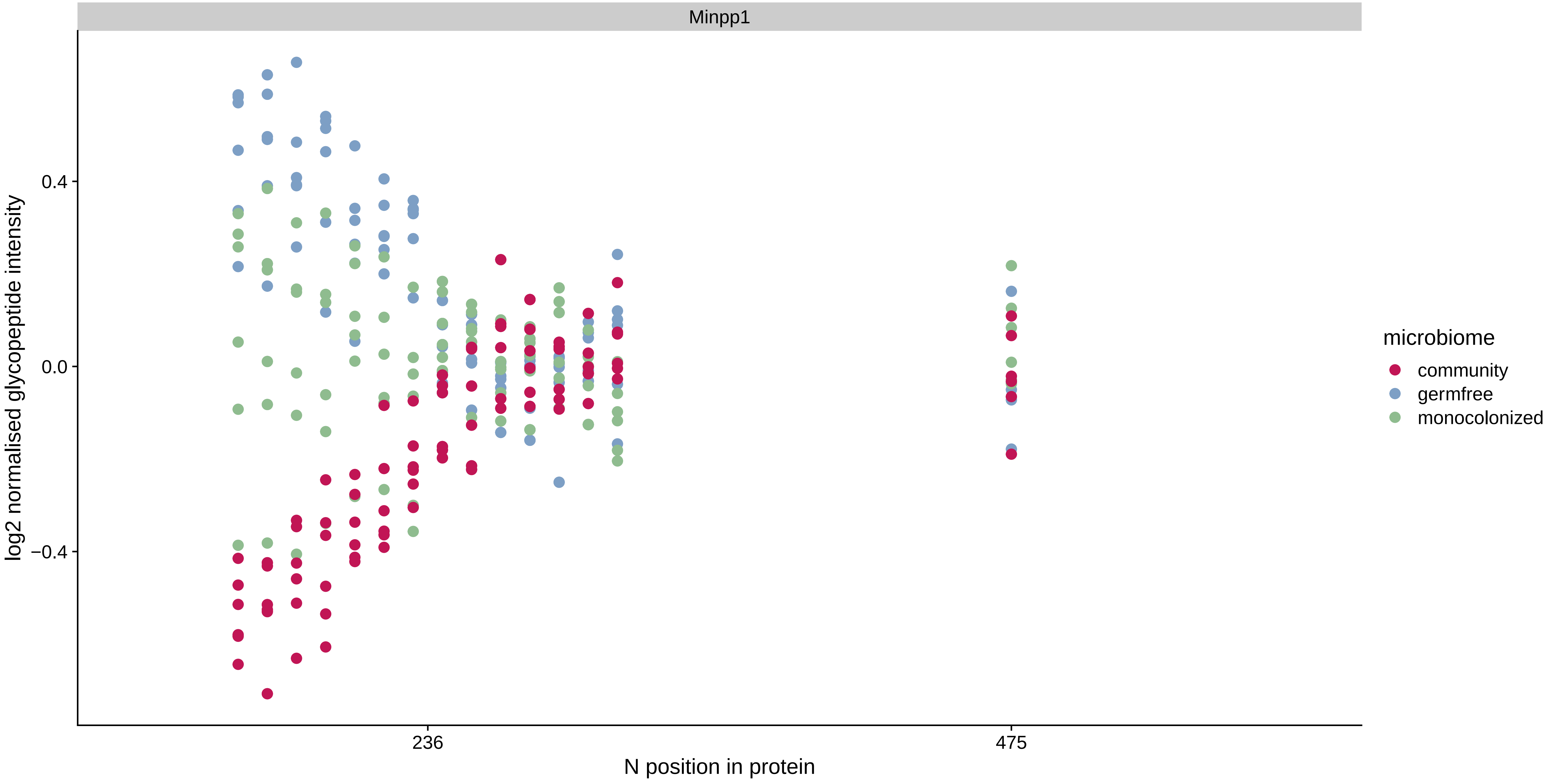

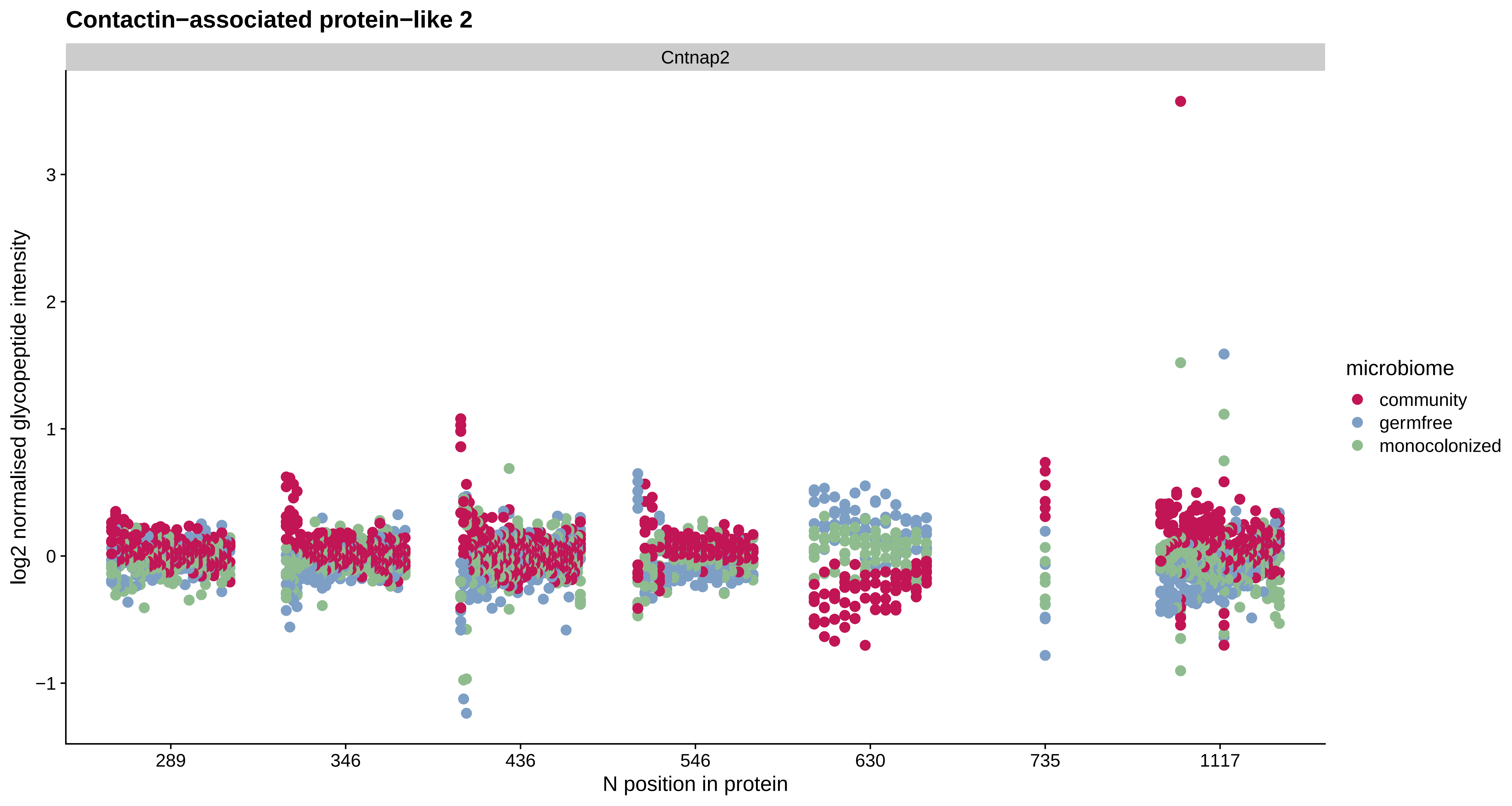
Contactin−associated protein−like 2
Cntnap2
log2 normalised glycopeptide intensity
3
2
1
0
−1
289
346
436
546
630
735
1117
N position in protein
microbiome
community
germfree
monocolonized

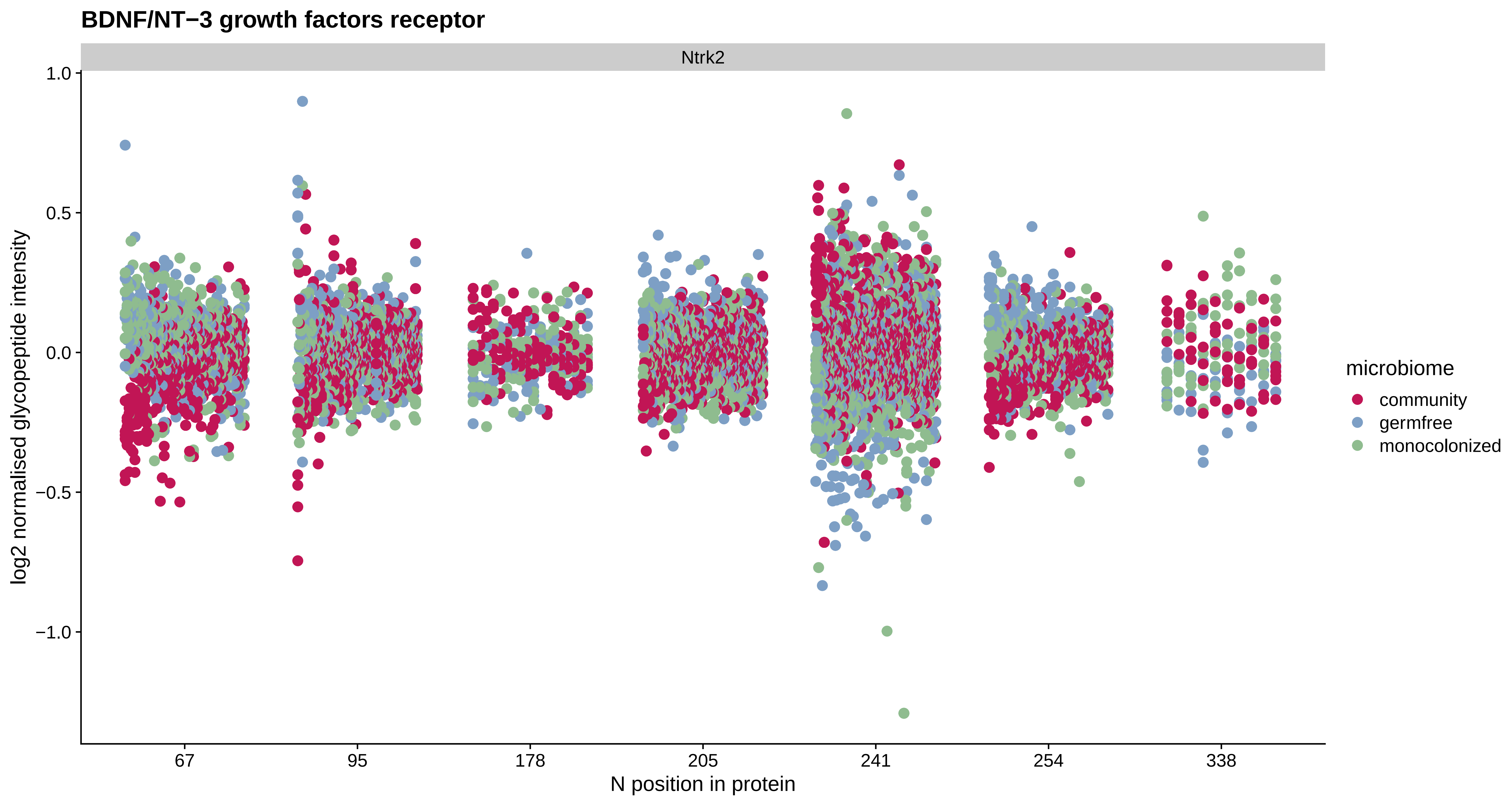

BDNF/NT−3 growth factors receptor
Ntrk2
log2 normalised glycopeptide intensity
1.0
0.5
0.0
−0.5
−1.0
67
95
178
205
241
254
338
N position in protein
microbiome
community
germfree
monocolonized

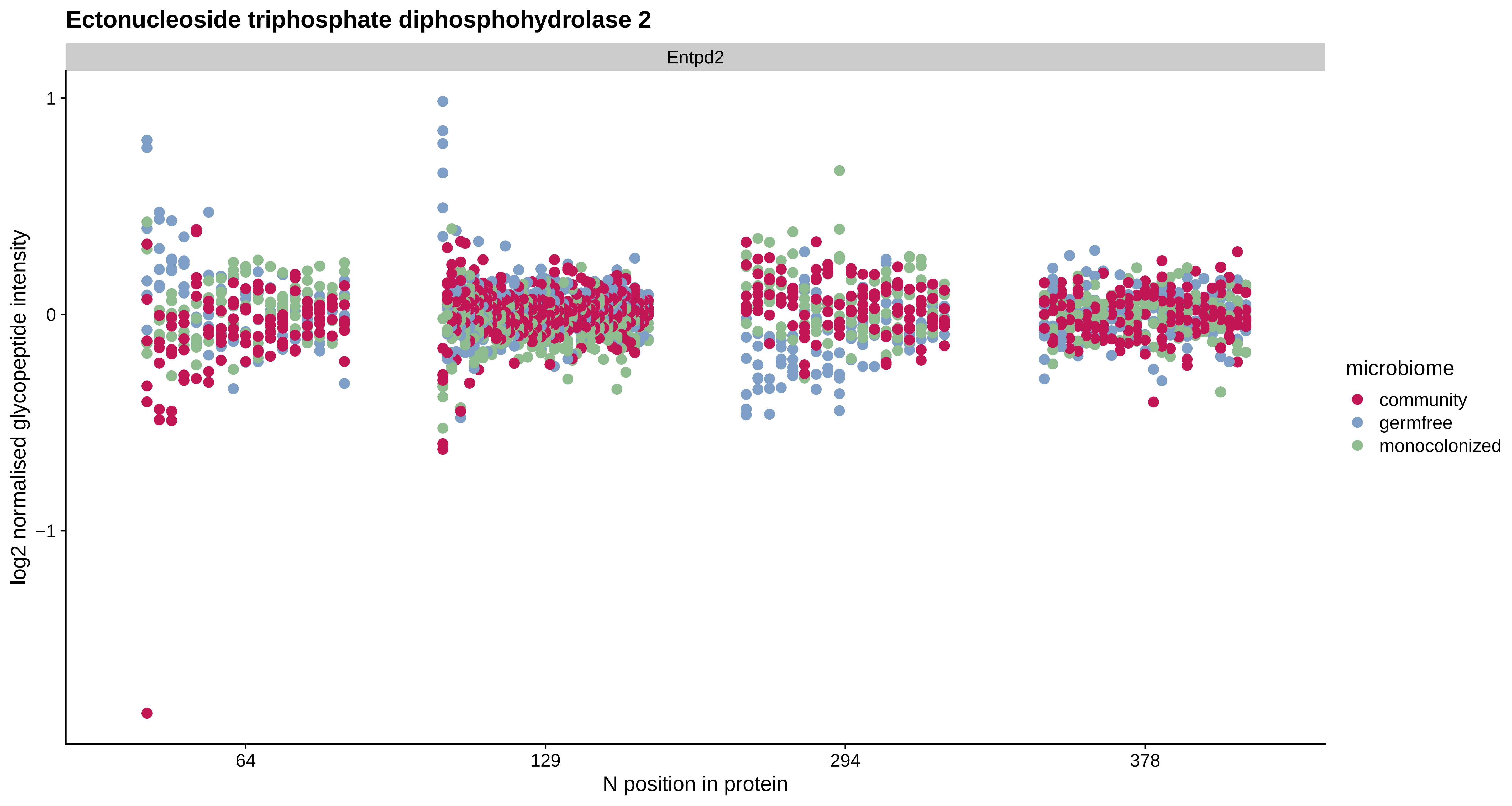

Ectonucleoside triphosphate diphosphohydrolase 2
Entpd2
log2 normalised glycopeptide intensity
1
0
−1
64
129
294
378
N position in protein
microbiome
community
germfree
monocolonized

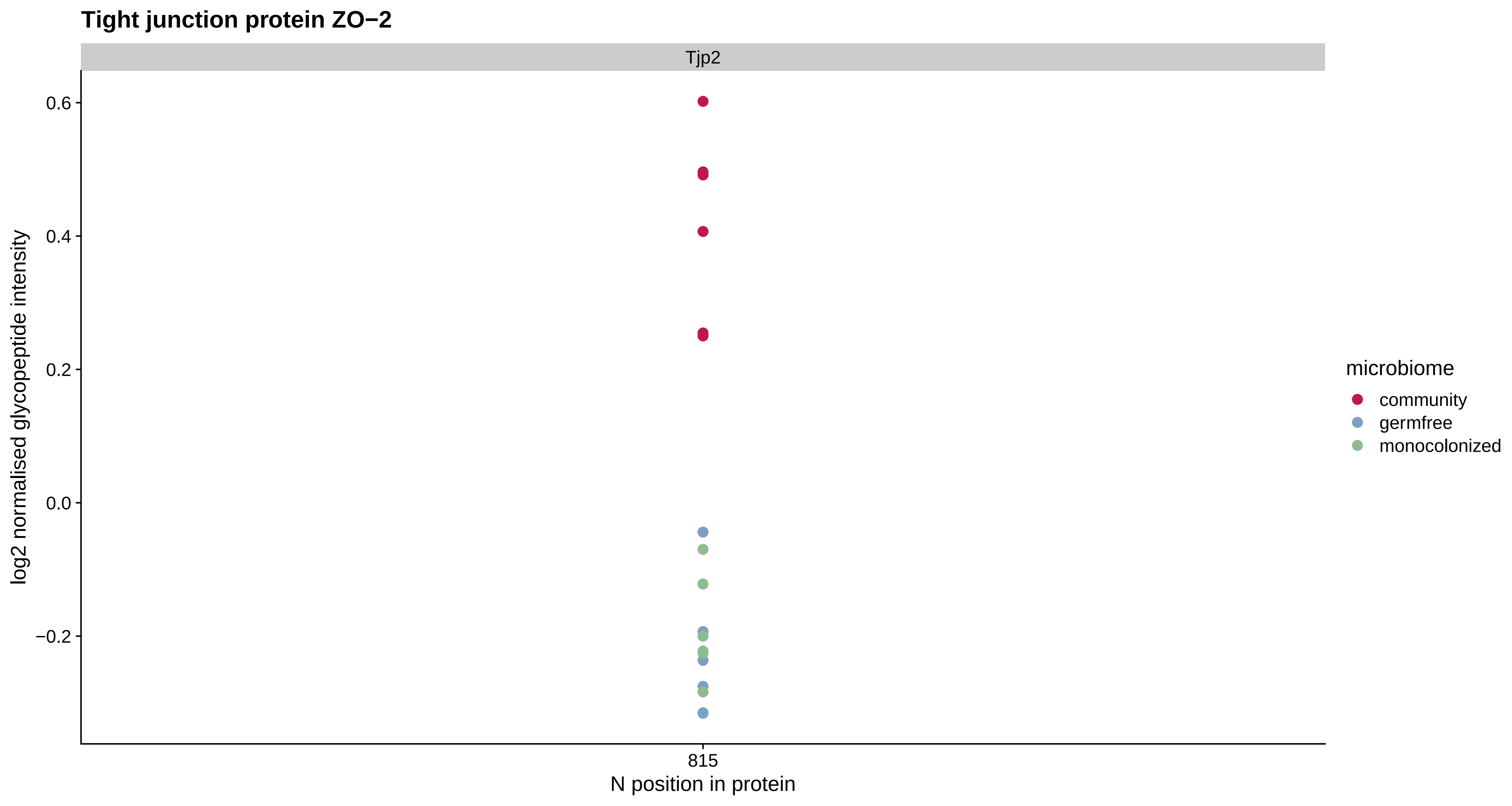

Tight junction protein ZO−2
Tjp2
0.6
0.4
0.2
0.0
−0.2
log2 normalised glycopeptide intensity
815
N position in protein
microbiome
community
germfree
monocolonized

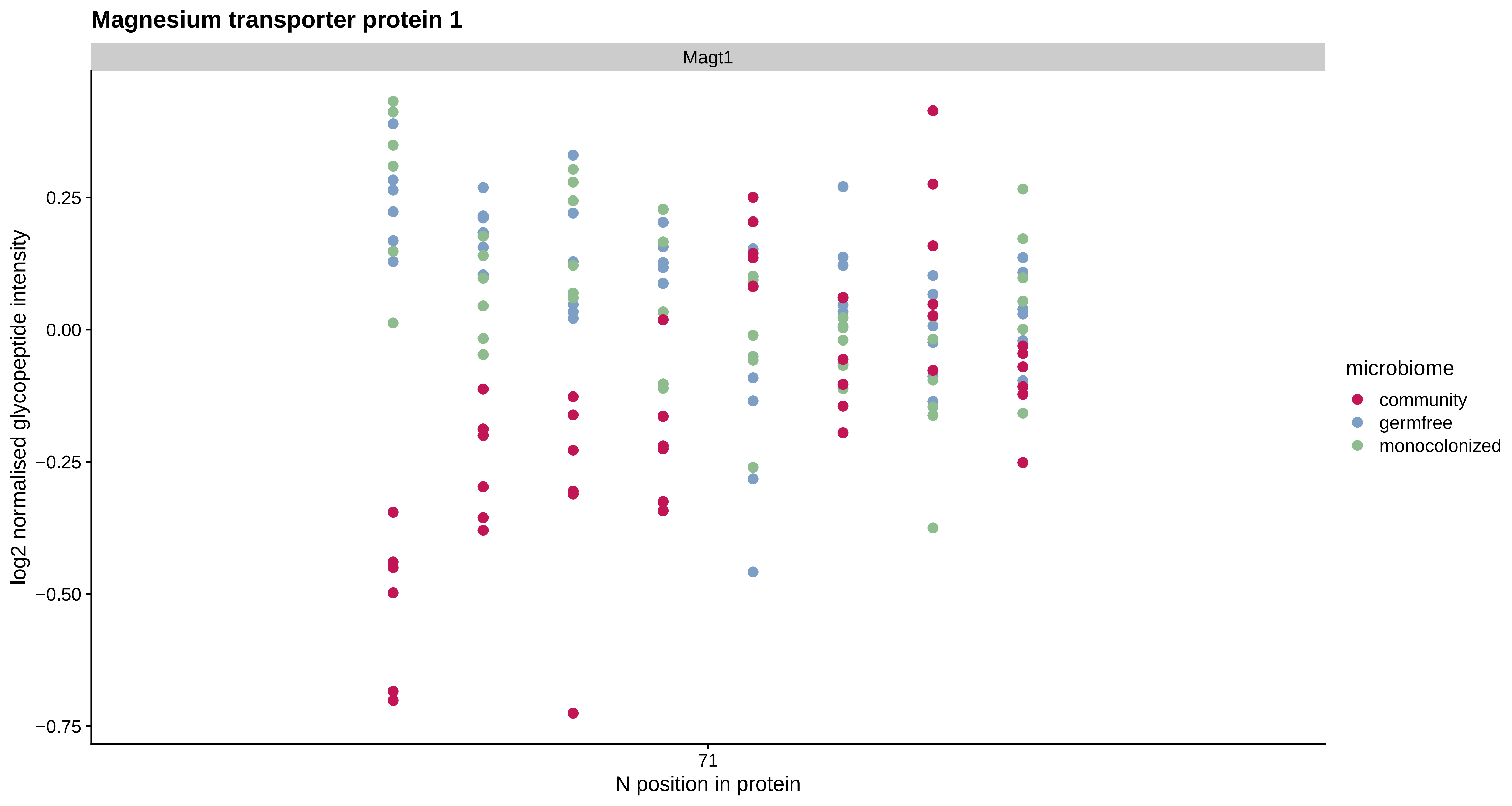

Magnesium transporter protein 1
Magt1
log2 normalised glycopeptide intensity
0.25
0.00
−0.25
−0.50
−0.75
71
N position in protein
microbiome
community
germfree
monocolonized

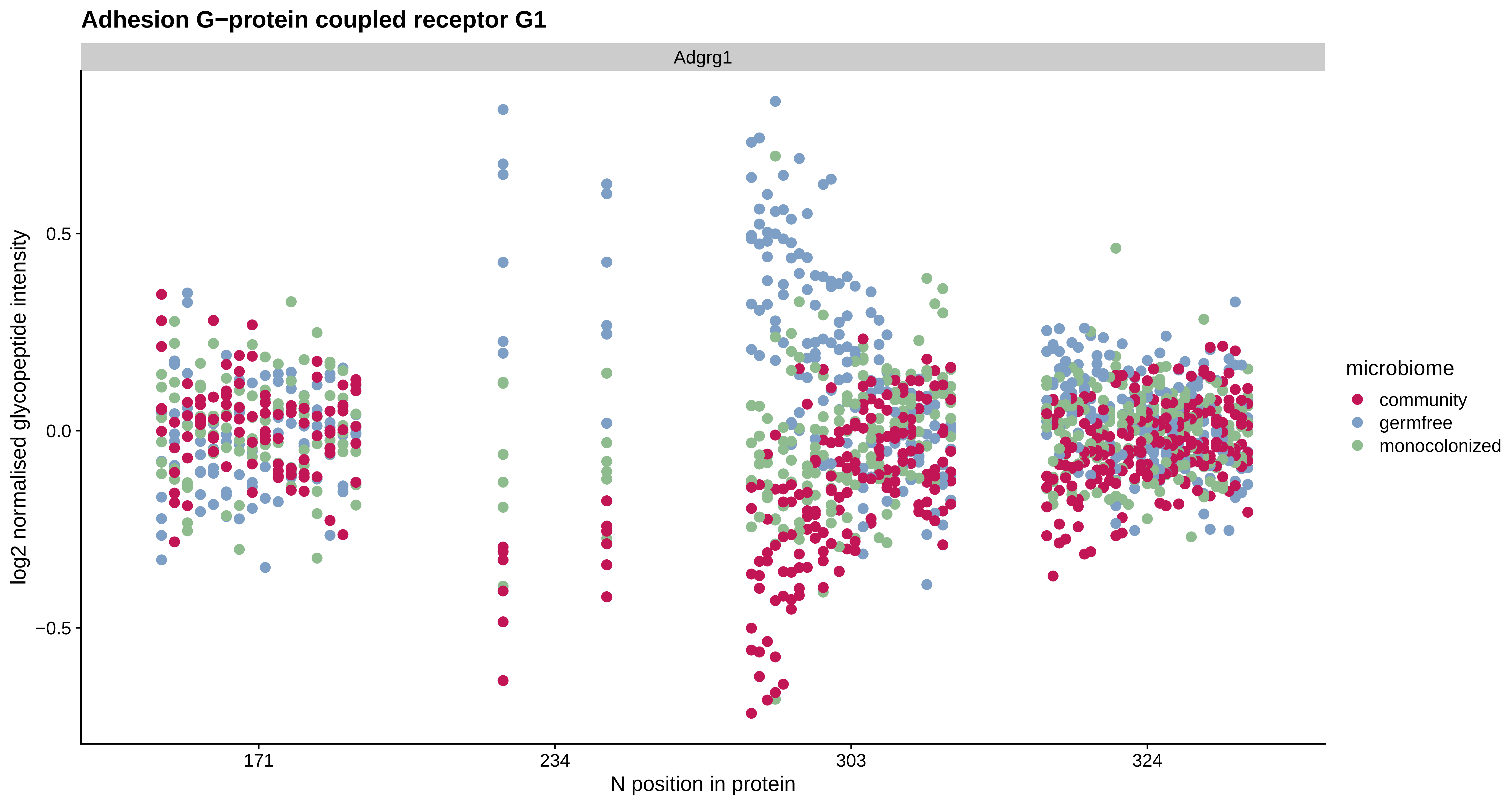
Adhesion G−protein coupled receptor G1
Adgrg1
log2 normalised glycopeptide intensity
0.5
0.0
−0.5
171
234
303
324
N position in protein
microbiome
community
germfree
monocolonized

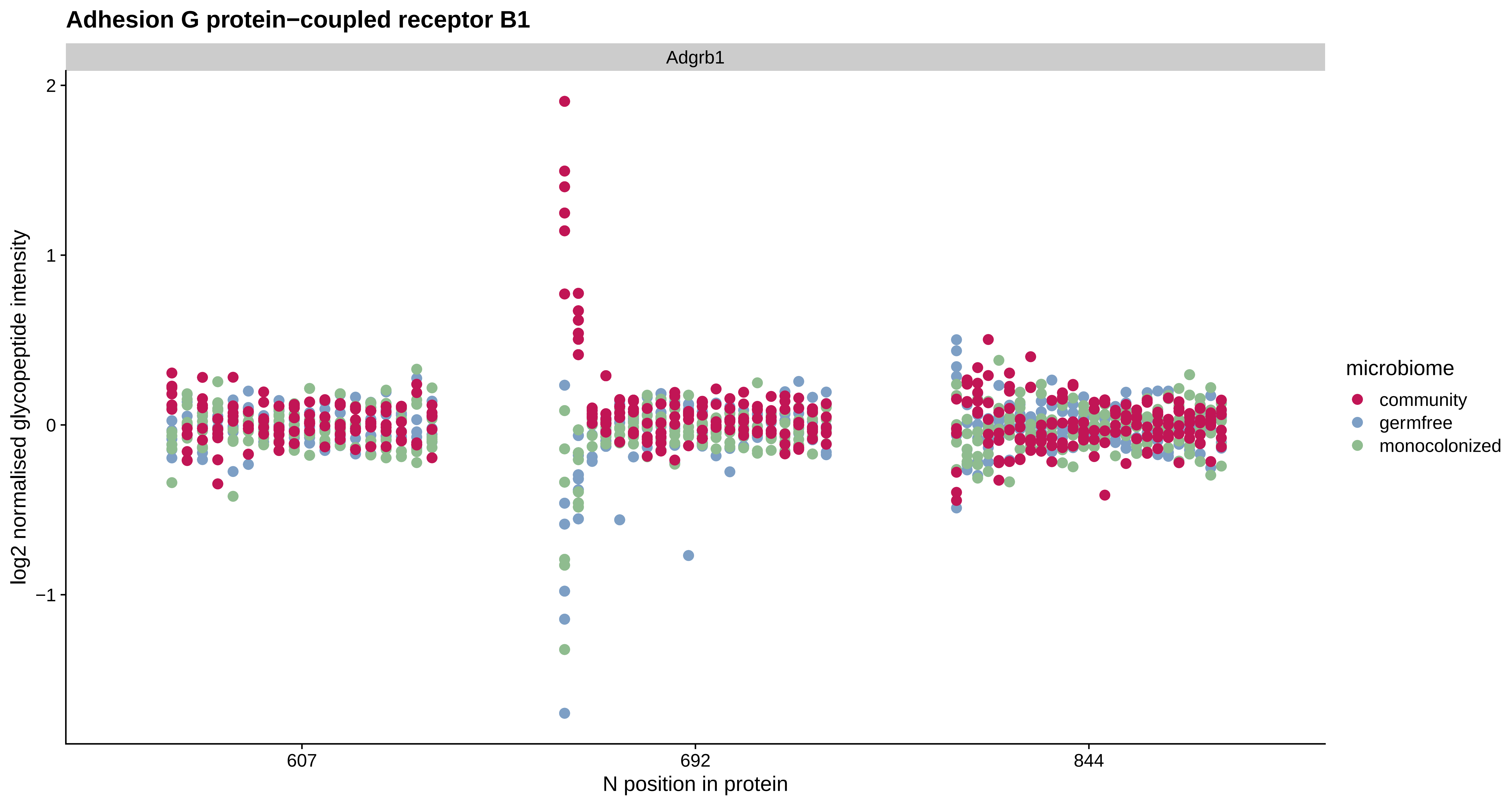

Adhesion G protein−coupled receptor B1
Adgrb1
log2 normalised glycopeptide intensity
2
1
0
−1
607
692
844
N position in protein
microbiome
community
germfree
monocolonized

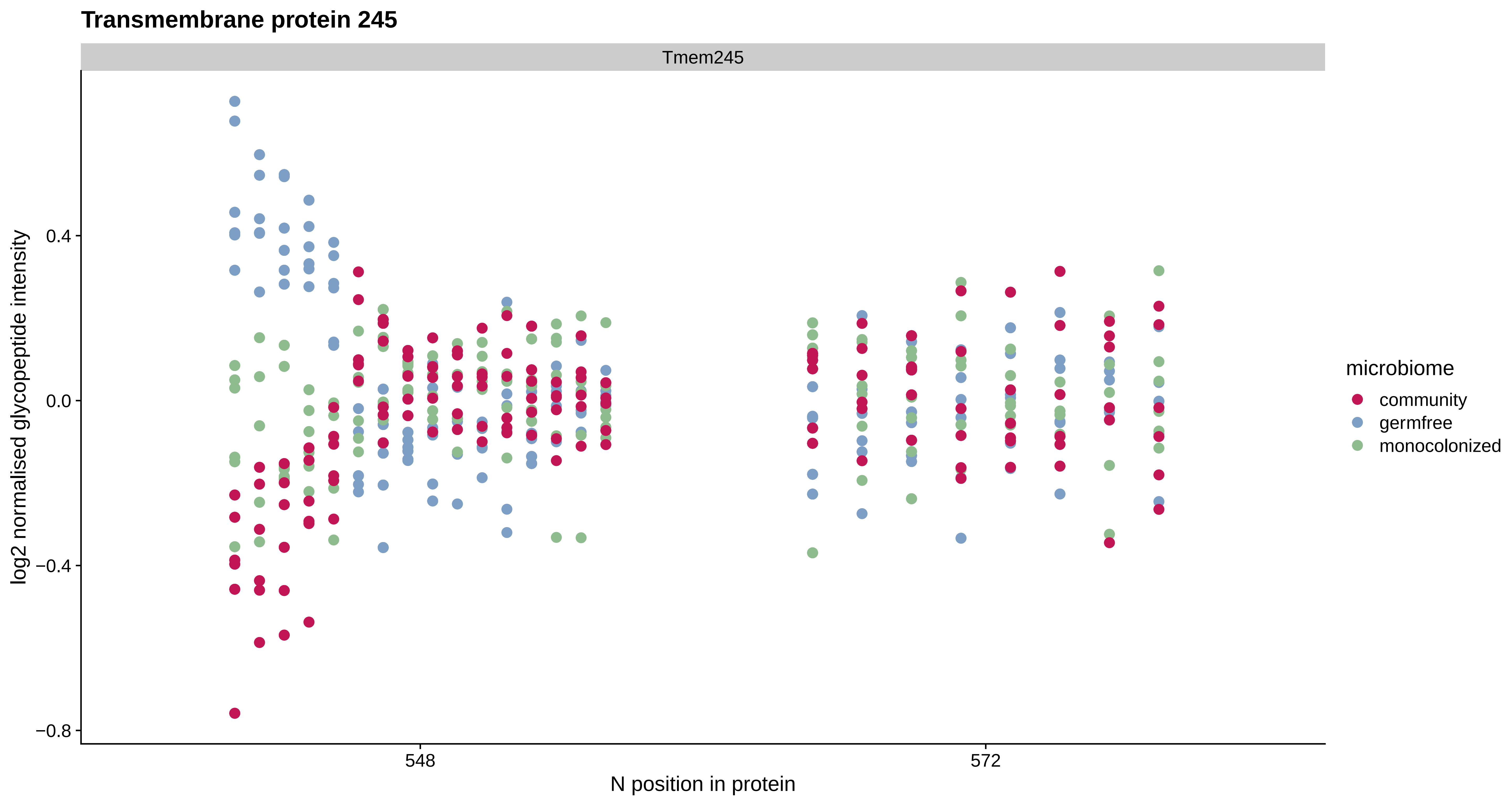

Transmembrane protein 245
Tmem245
log2 normalised glycopeptide intensity
0.4
0.0
−0.4
−0.8
548
572
N position in protein
microbiome
community
germfree
monocolonized

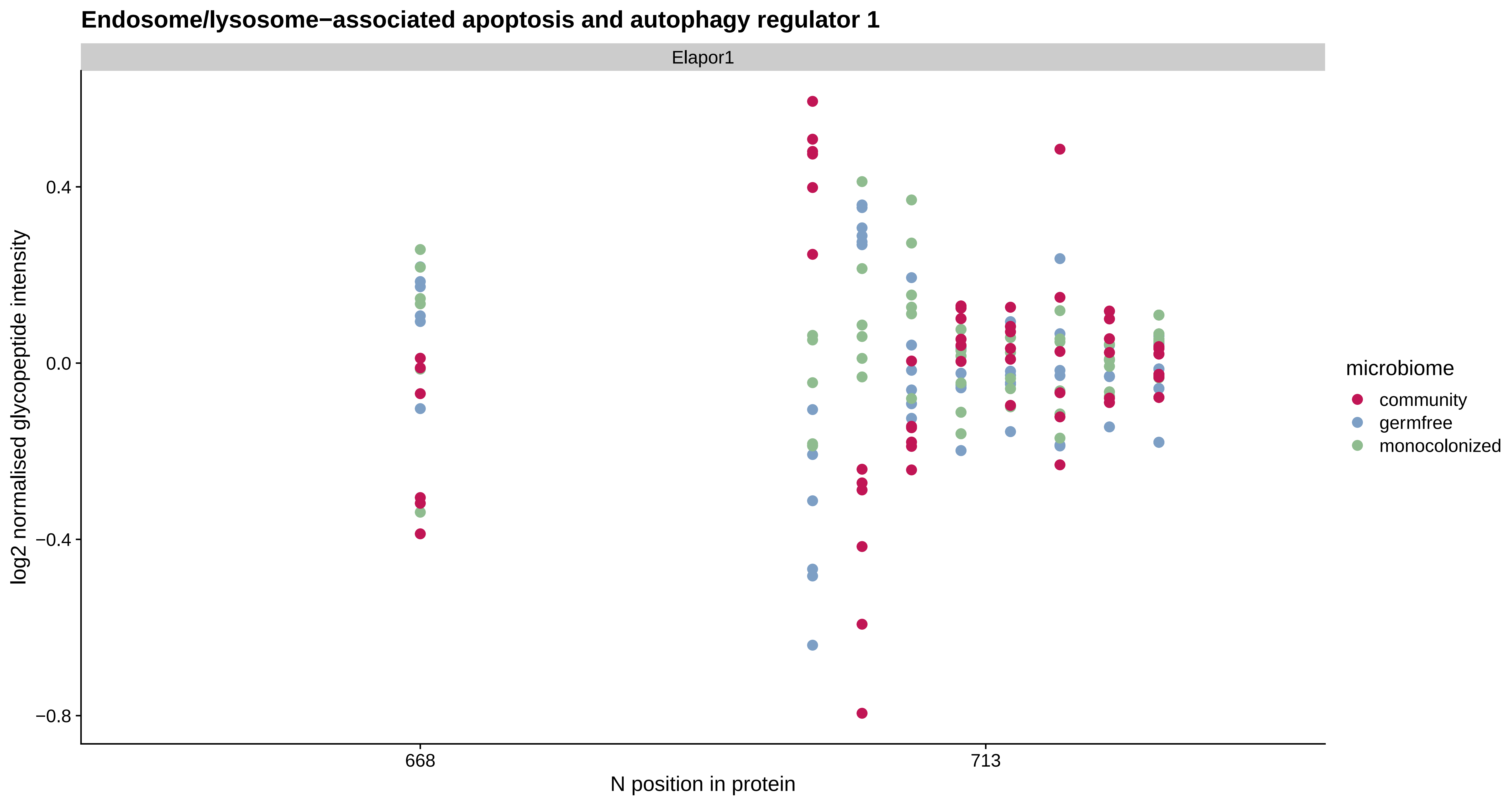

Endosome/lysosome−associated apoptosis and autophagy regulator 1
Elapor1
log2 normalised glycopeptide intensity
0.4
0.0
−0.4
−0.8
668
713
N position in protein
microbiome
community
germfree
monocolonized

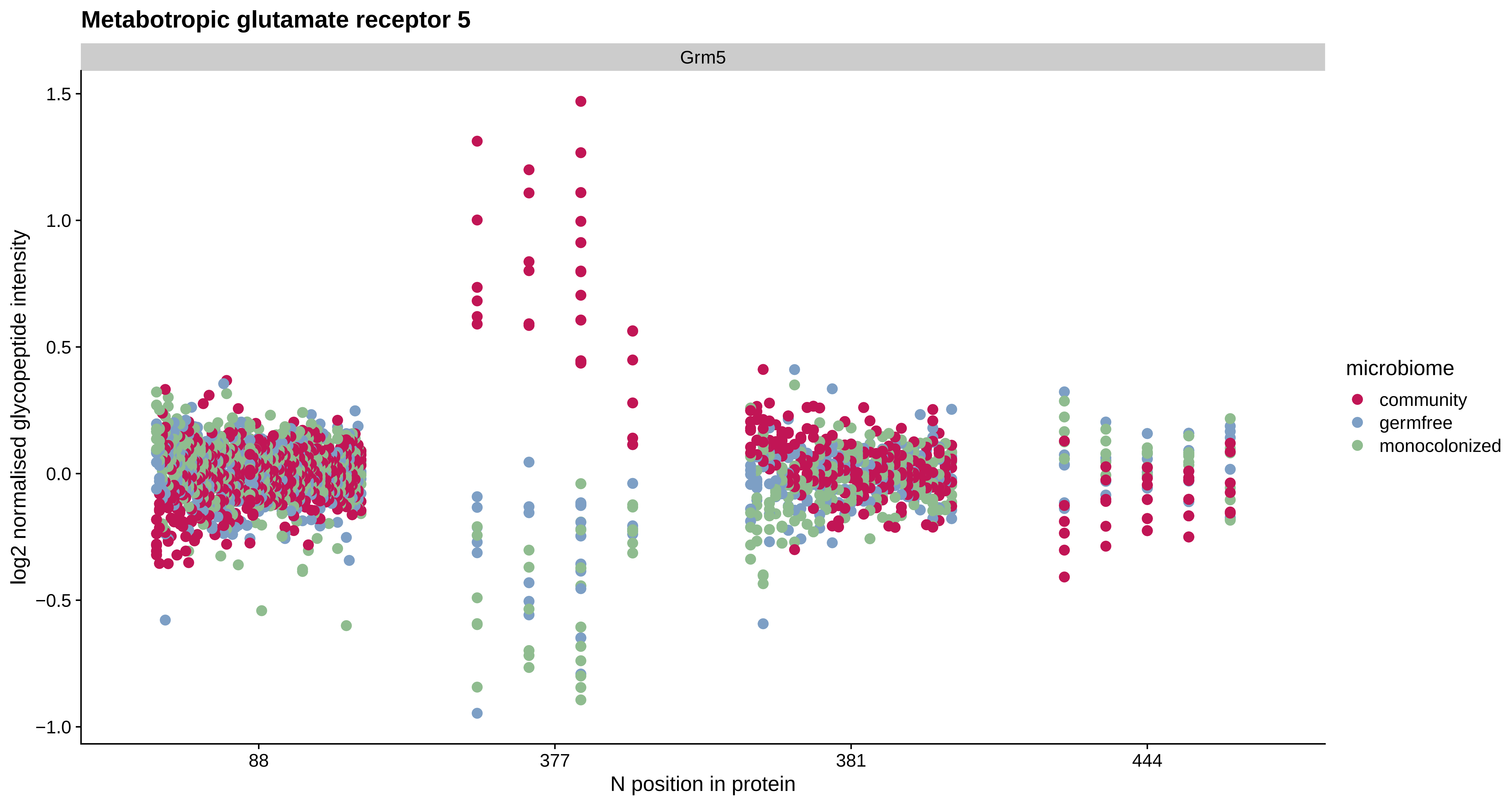
Metabotropic glutamate receptor 5
Grm5
1.5
1.0
0.5
0.0
−0.5
−1.0
log2 normalised glycopeptide intensity
88
377
381
444
N position in protein
microbiome
community
germfree
monocolonized

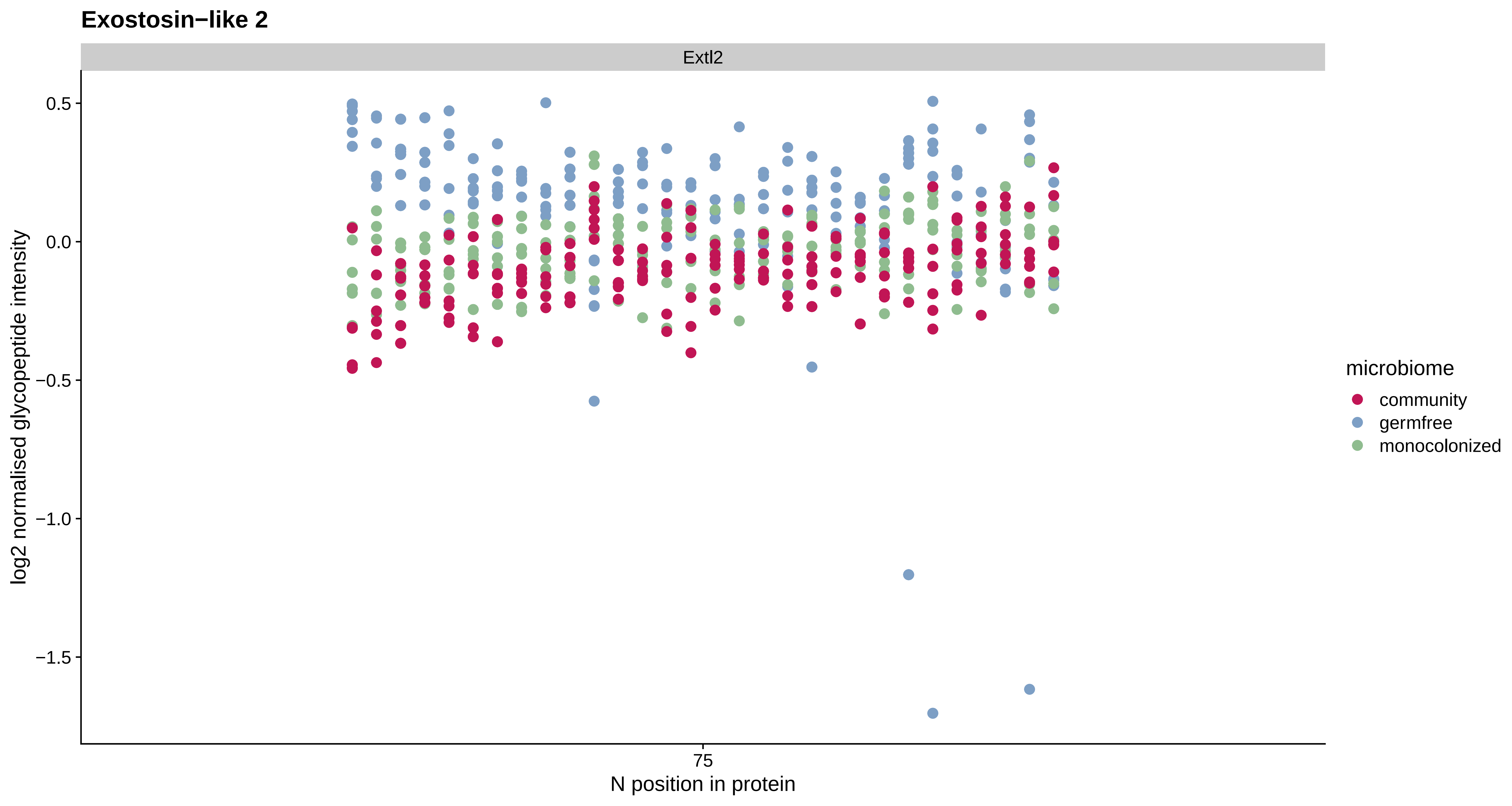
Exostosin−like 2
Extl2
0.5
0.0
−0.5
−1.0
−1.5
log2 normalised glycopeptide intensity
75
N position in protein
microbiome
community
germfree
monocolonized

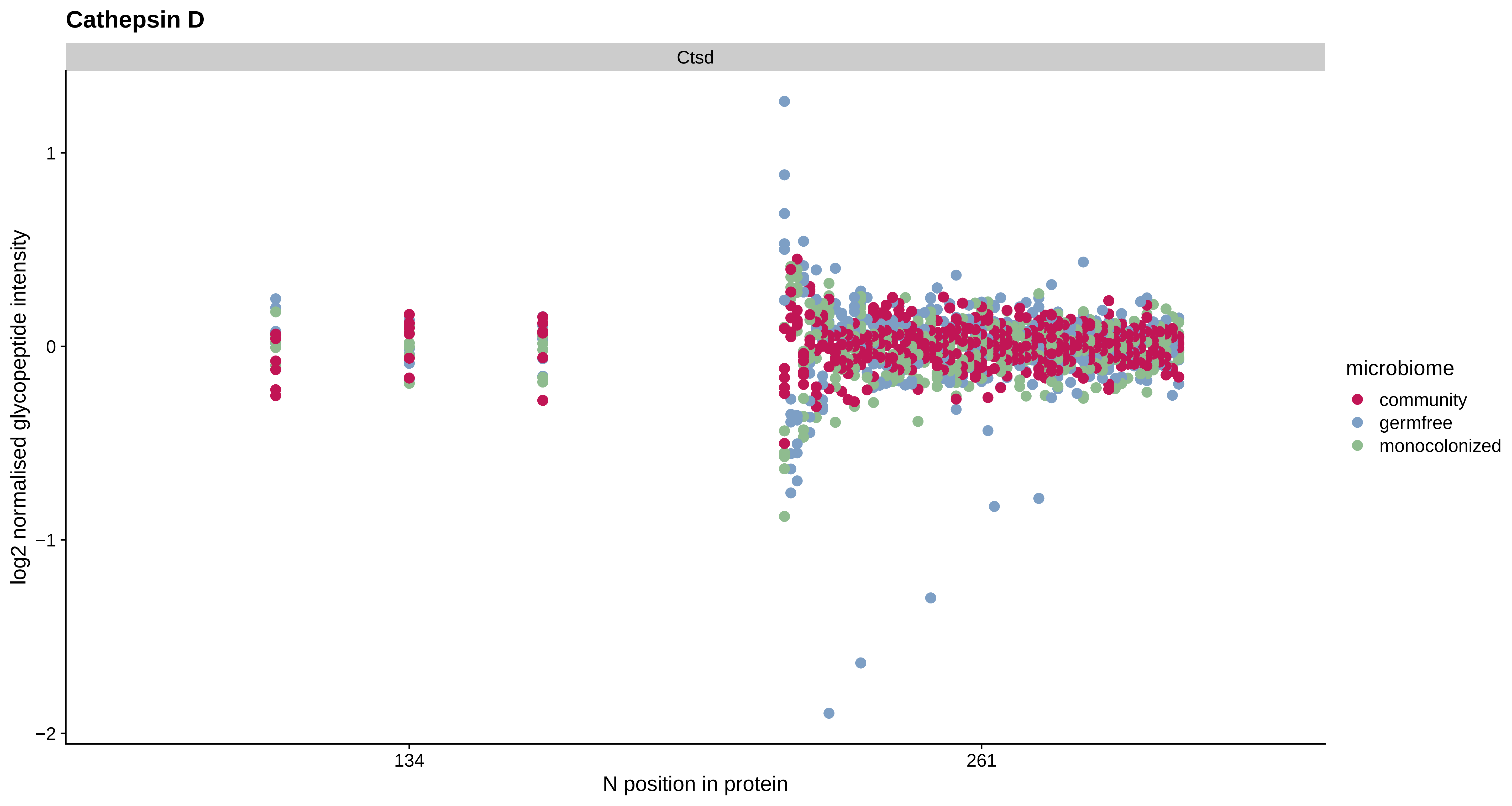

Cathepsin D
Ctsd
log2 normalised glycopeptide intensity
1
0
−1
−2
134
261
N position in protein
microbiome
community
germfree
monocolonized

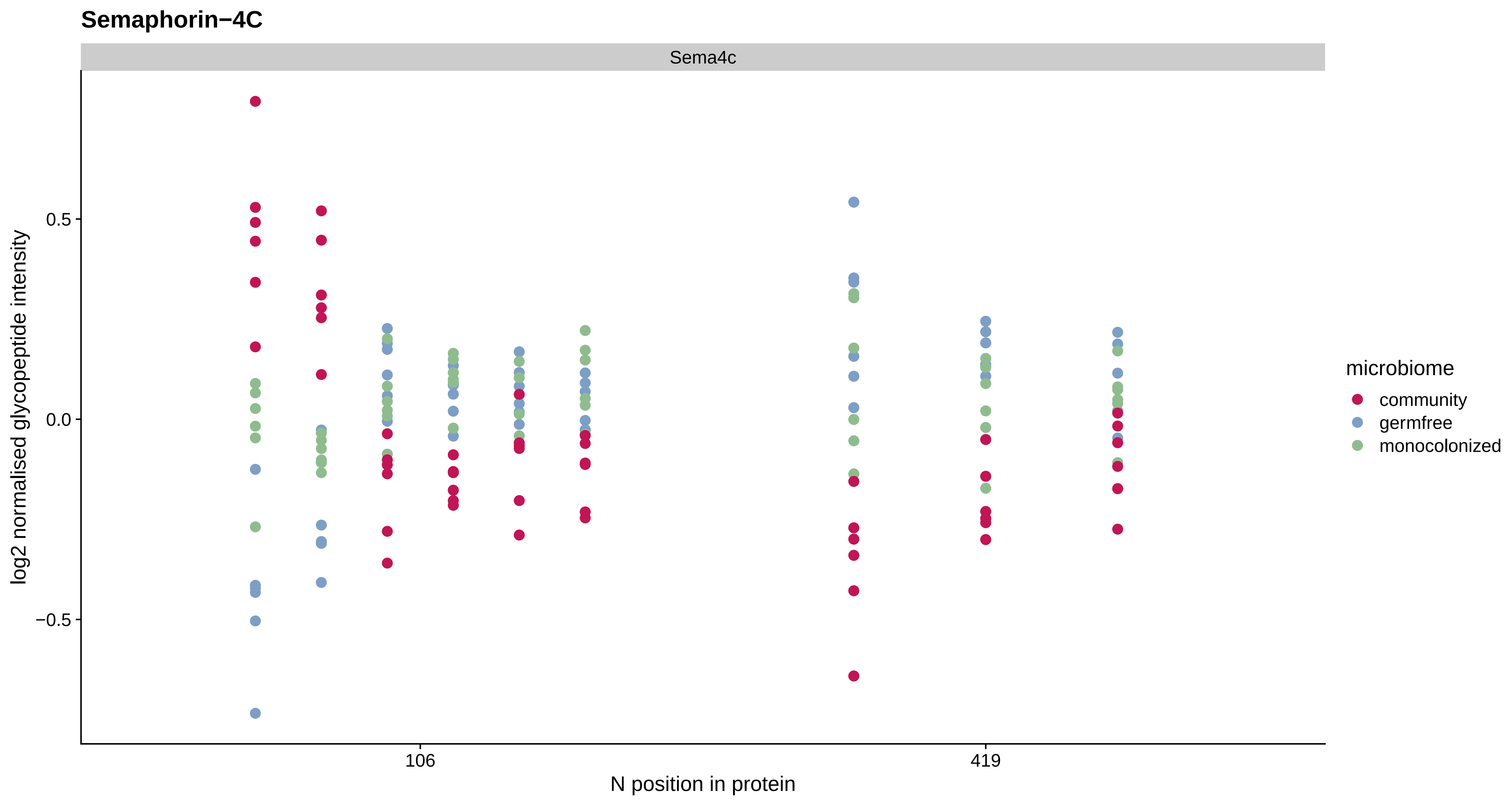

Semaphorin−4C
Sema4c
log2 normalised glycopeptide intensity
0.5
0.0
−0.5
106
419
N position in protein
microbiome
community
germfree
monocolonized

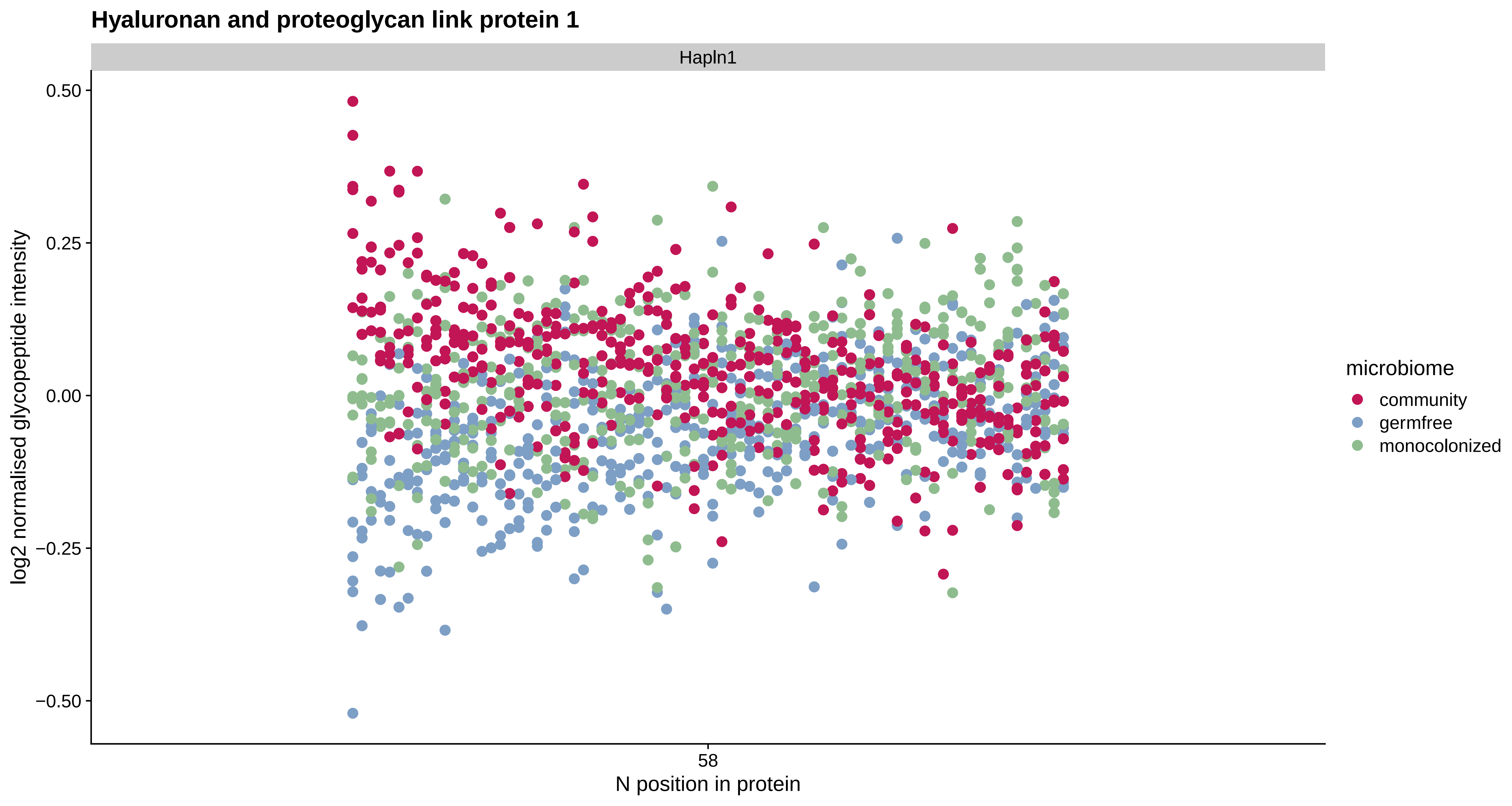
Hyaluronan and proteoglycan link protein 1
Hapln1
0.50
0.25
0.00
−0.25
−0.50
log2 normalised glycopeptide intensity
58
N position in protein
microbiome
community
germfree
monocolonized

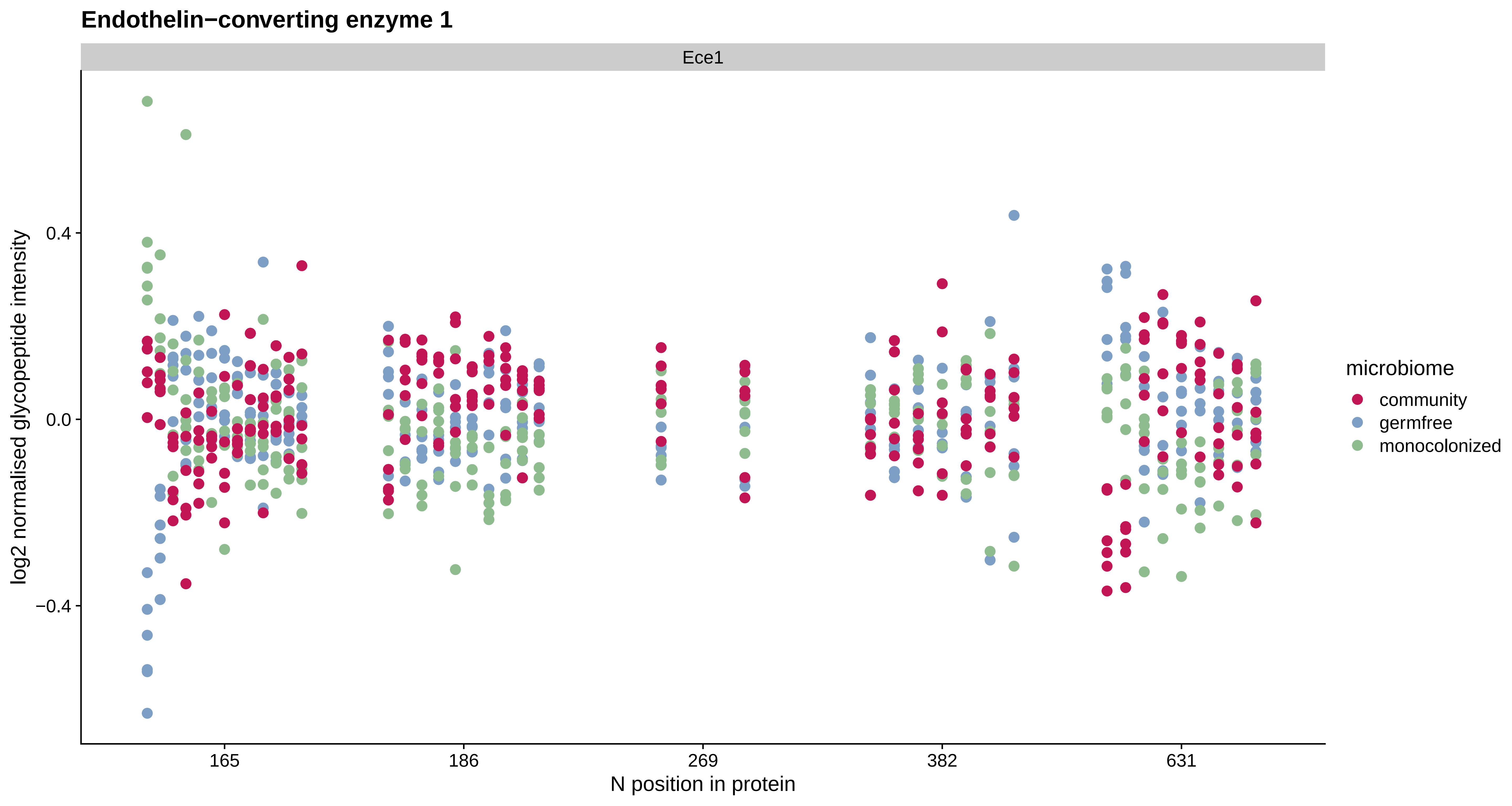

Endothelin−converting enzyme 1
Ece1
log2 normalised glycopeptide intensity
0.4
0.0
−0.4
165
186
269
382
631
N position in protein
microbiome
community
germfree
monocolonized

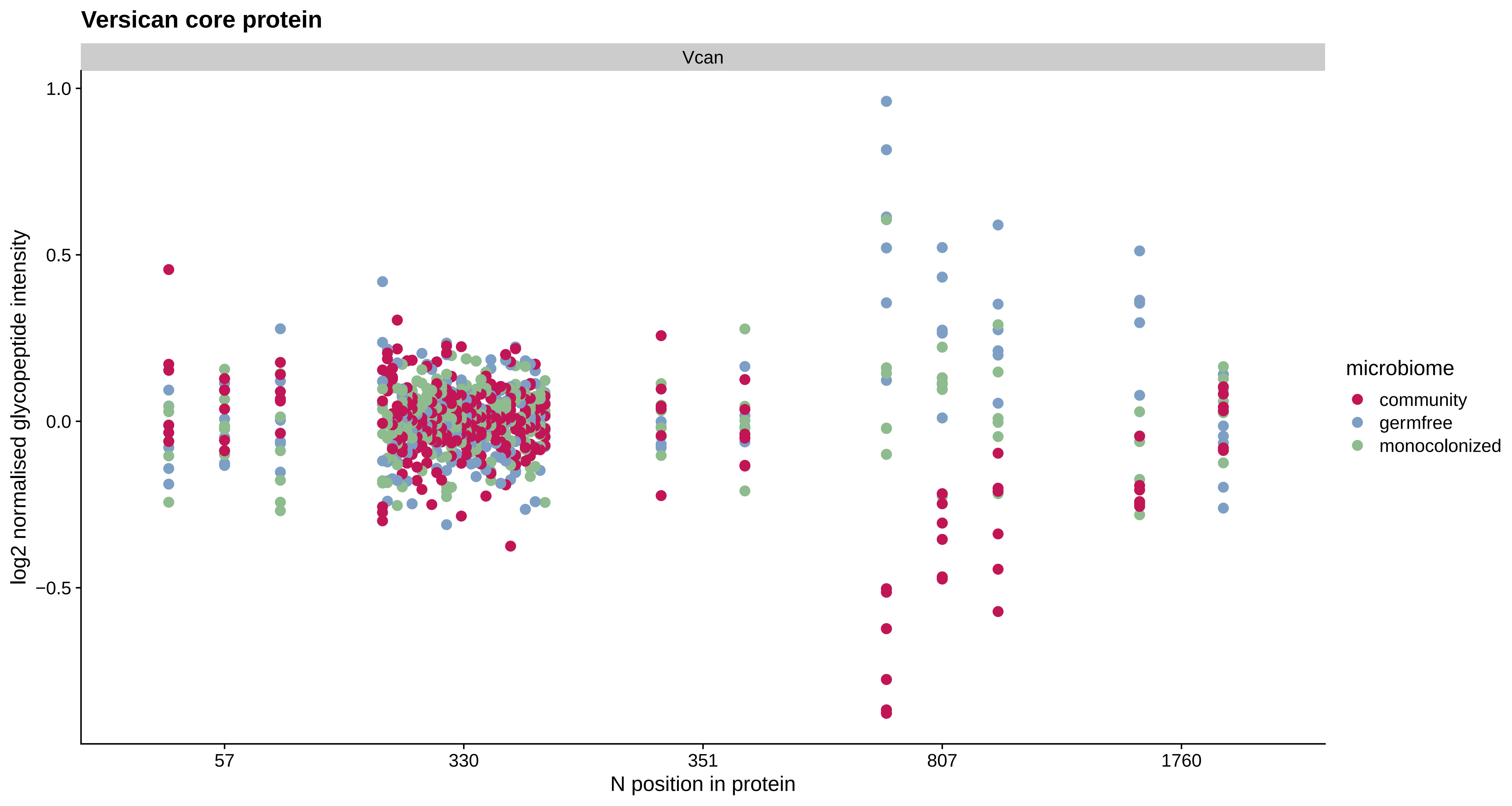
Versican core protein
Vcan
log2 normalised glycopeptide intensity
1.0
0.5
0.0
−0.5
57
330
351
807
1760
N position in protein
microbiome
community
germfree
monocolonized

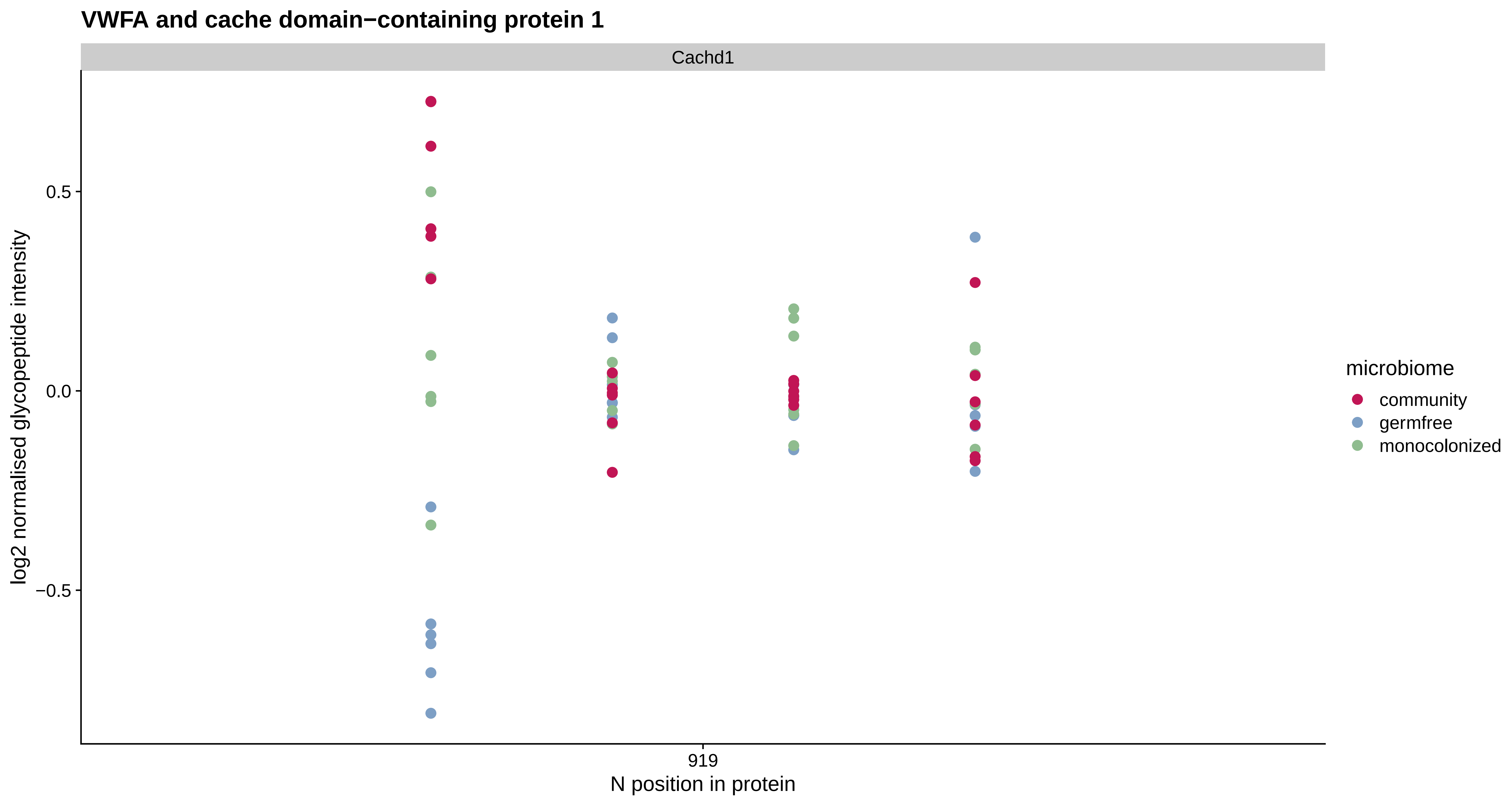

VWFA and cache domain−containing protein 1
Cachd1
0.5
0.0
−0.5
log2 normalised glycopeptide intensity
919
N position in protein
microbiome
community
germfree
monocolonized

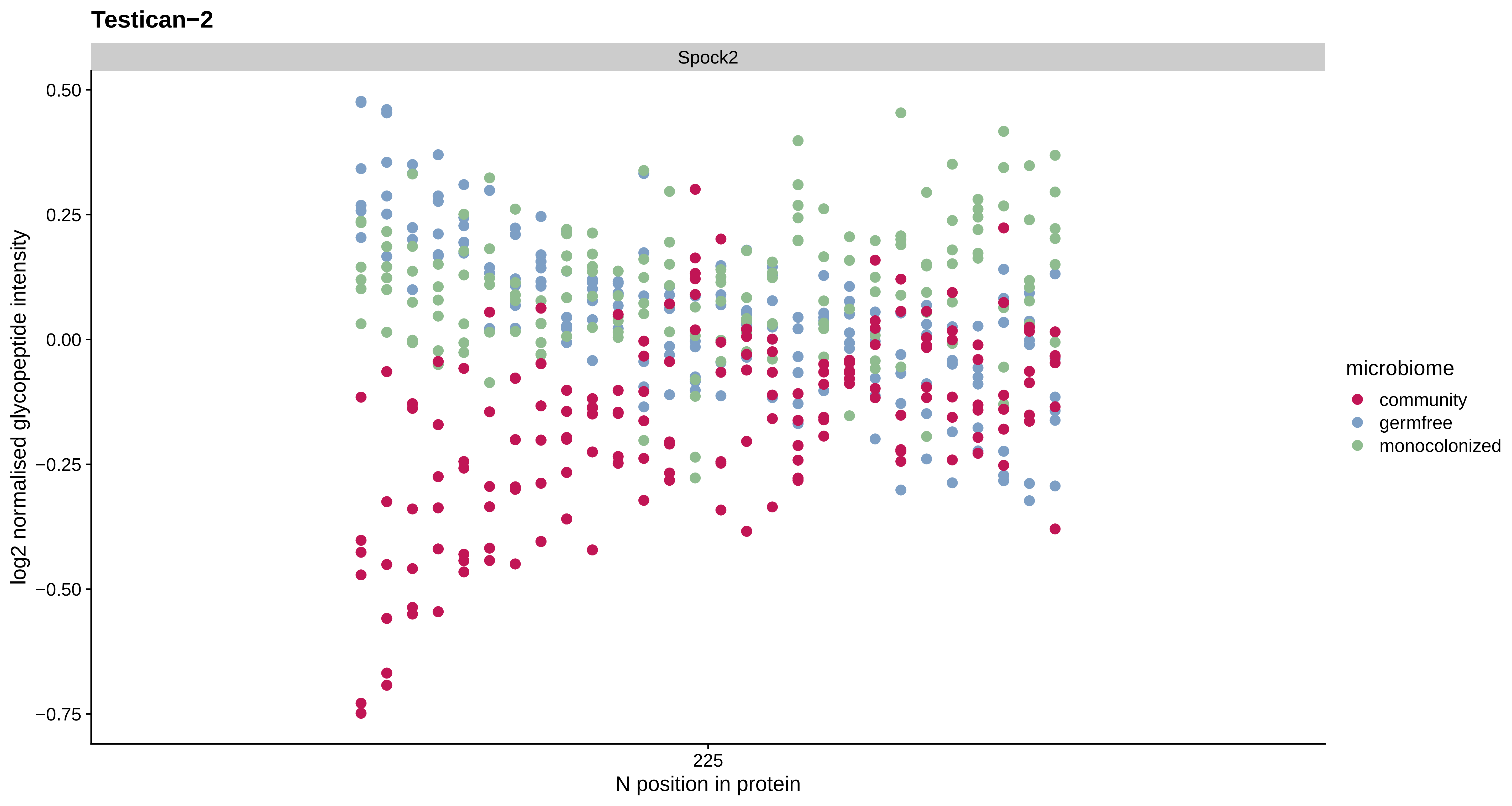

Testican−2
Spock2
0.50
0.25
0.00
−0.25
−0.50
−0.75
log2 normalised glycopeptide intensity
225
N position in protein
microbiome
community
germfree
monocolonized

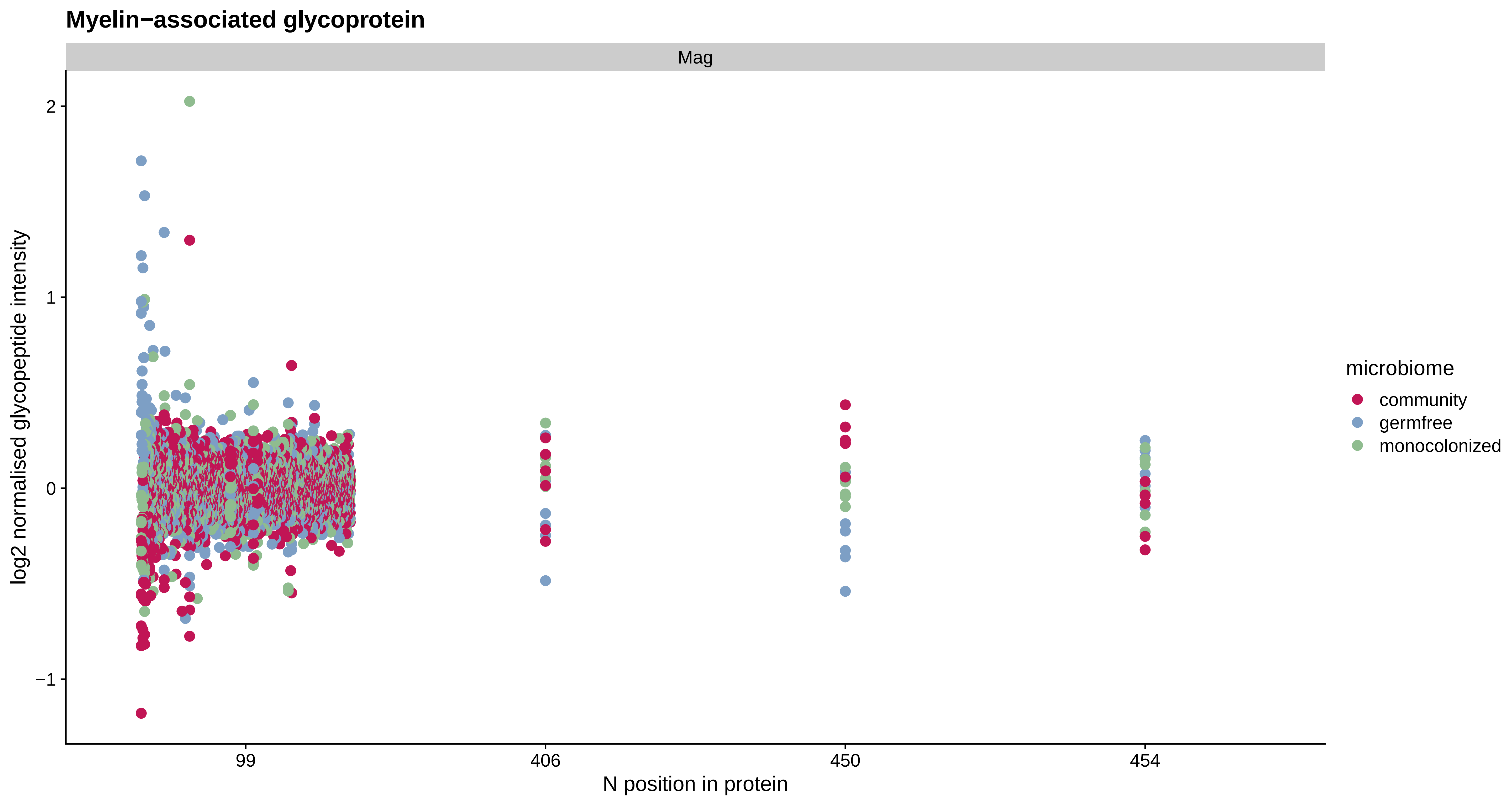
Myelin−associated glycoprotein
Mag
2
1
0
−1
log2 normalised glycopeptide intensity
99
406
450
454
N position in protein
microbiome
community
germfree
monocolonized

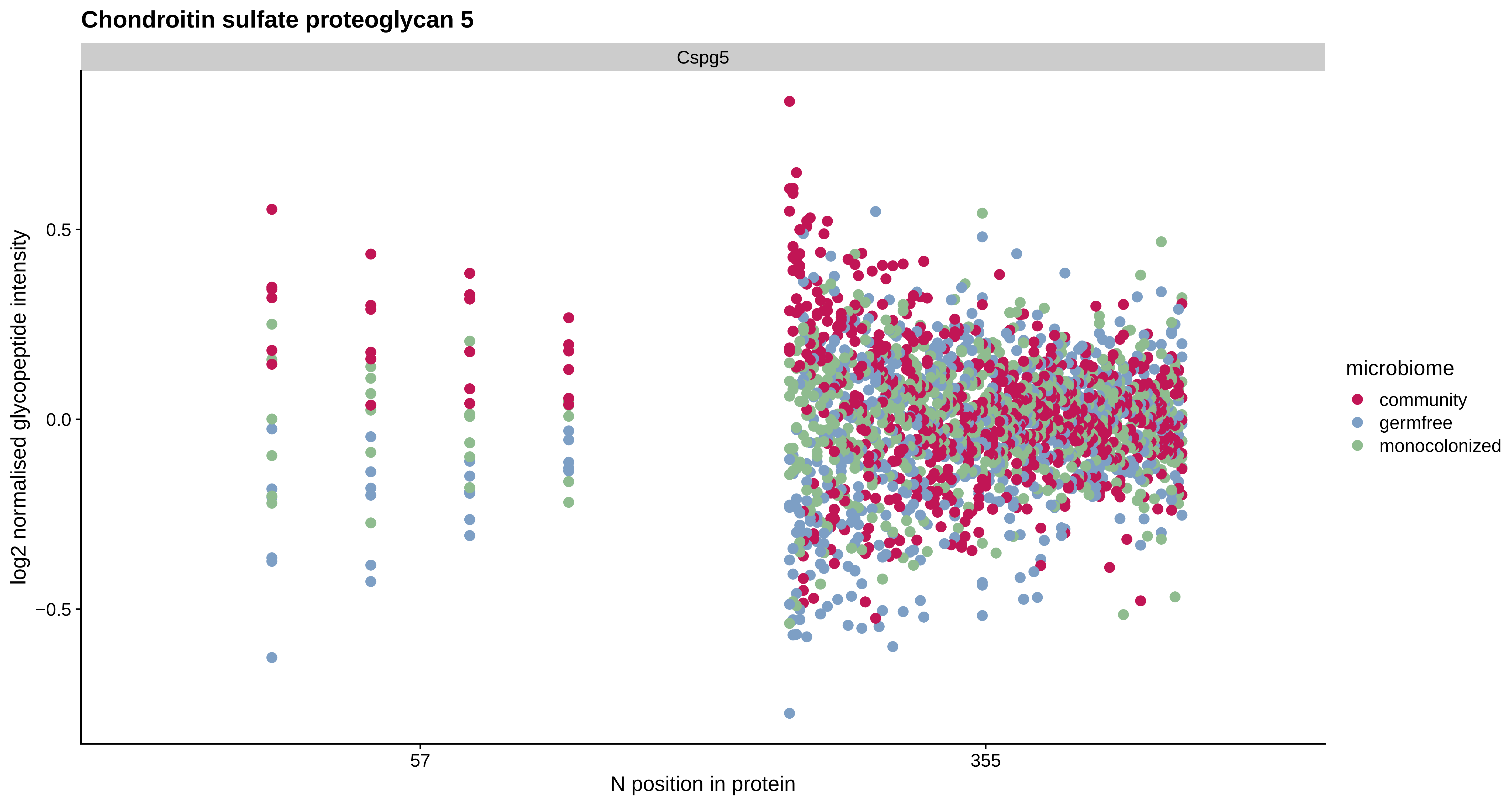
Chondroitin sulfate proteoglycan 5
Cspg5
log2 normalised glycopeptide intensity
0.5
0.0
−0.5
57
355
N position in protein
microbiome
community
germfree
monocolonized

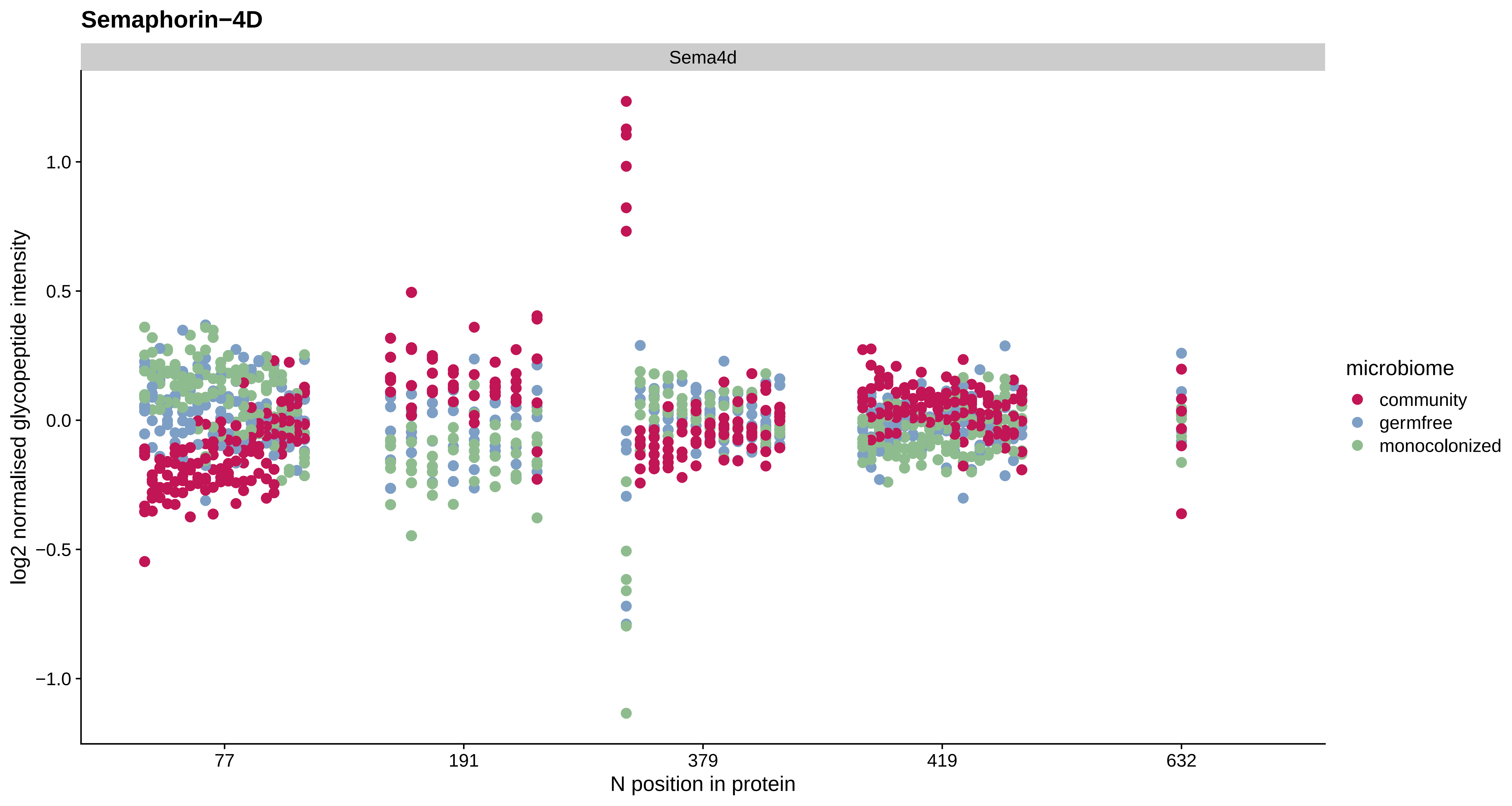
Semaphorin−4D
Sema4d
log2 normalised glycopeptide intensity
1.0
0.5
0.0
−0.5
−1.0
77
191
379
419
632
N position in protein
microbiome
community
germfree
monocolonized

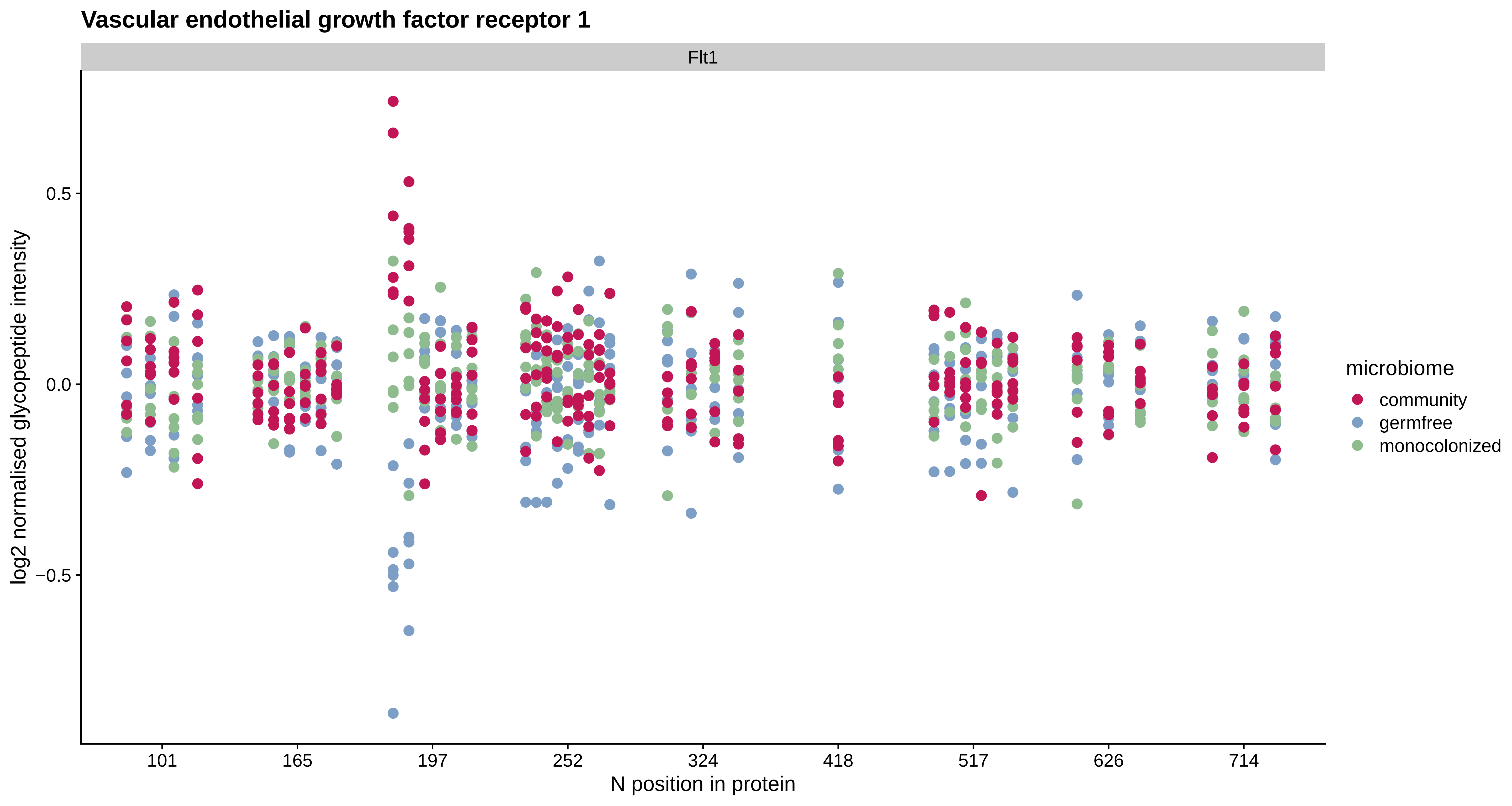

Vascular endothelial growth factor receptor 1
Flt1
log2 normalised glycopeptide intensity
0.5
0.0
−0.5
101
165
197
252
324
418
517
626
714
N position in protein
microbiome
community
germfree
monocolonized

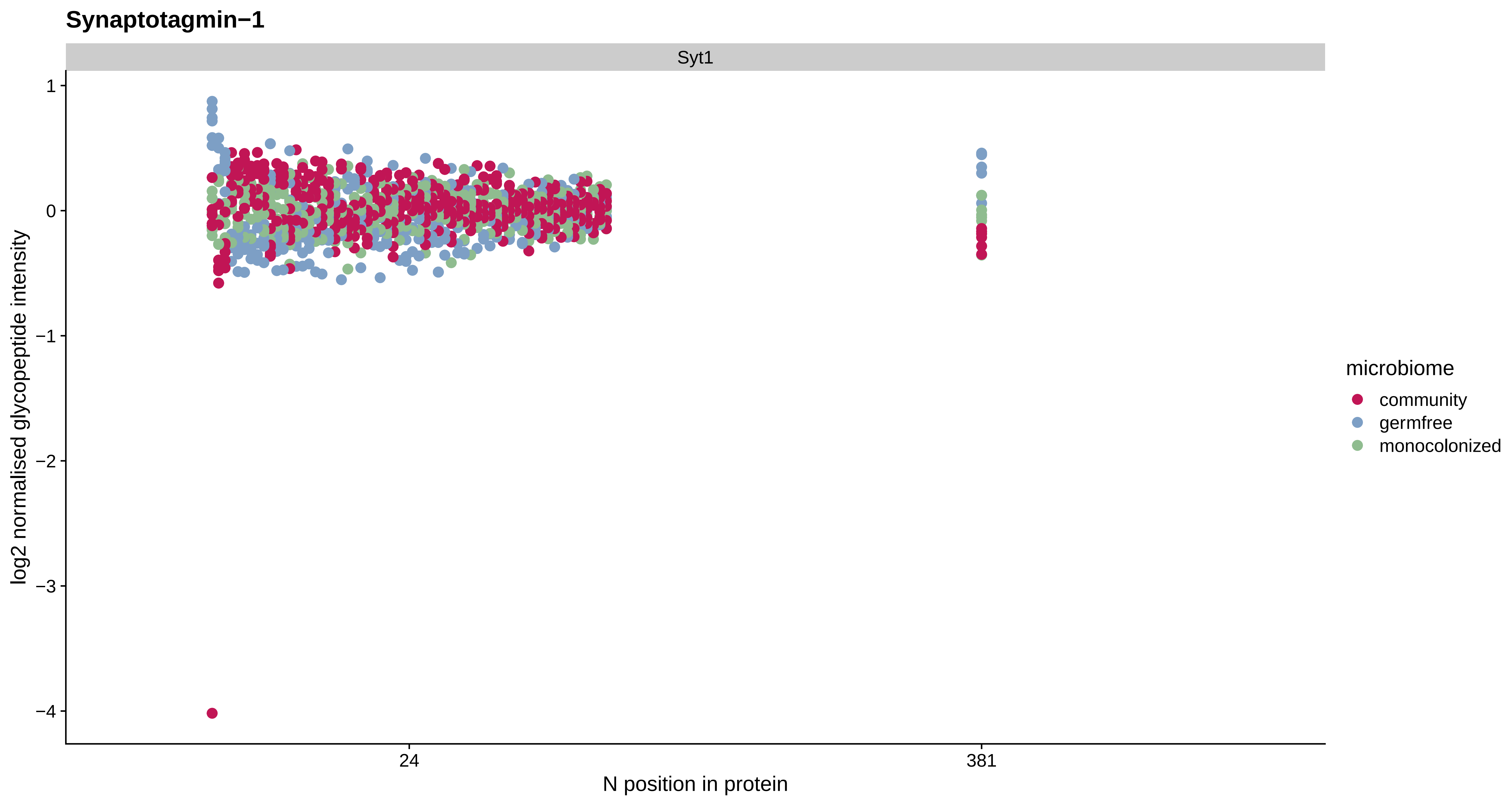
Synaptotagmin−1
Syt1
1
0
−1
−2
−3
−4
log2 normalised glycopeptide intensity
24
381
N position in protein
microbiome
community
germfree
monocolonized

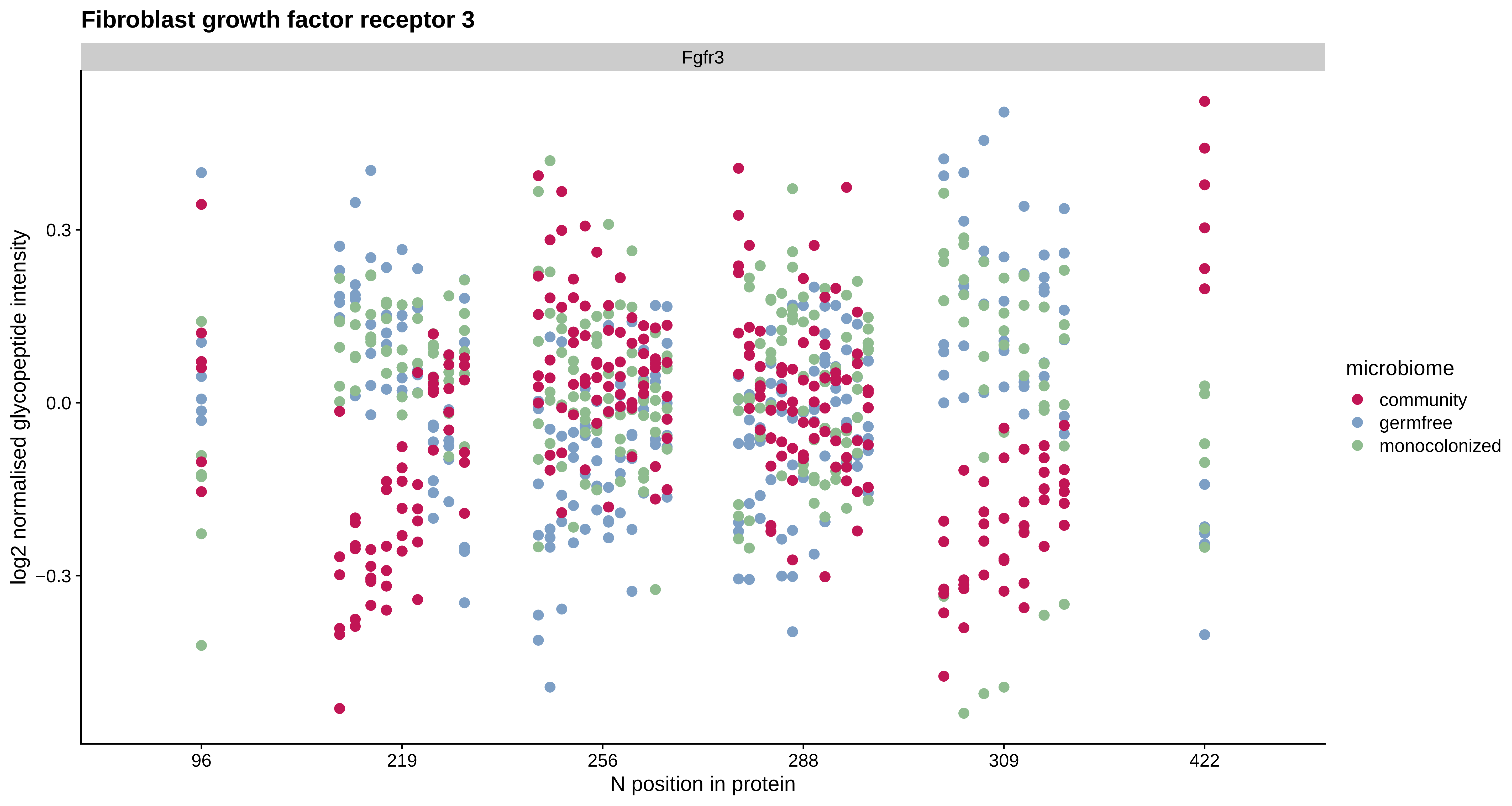

Fibroblast growth factor receptor 3
Fgfr3
log2 normalised glycopeptide intensity
0.3
0.0
−0.3
96
219
256
288
309
422
N position in protein
microbiome
community
germfree
monocolonized

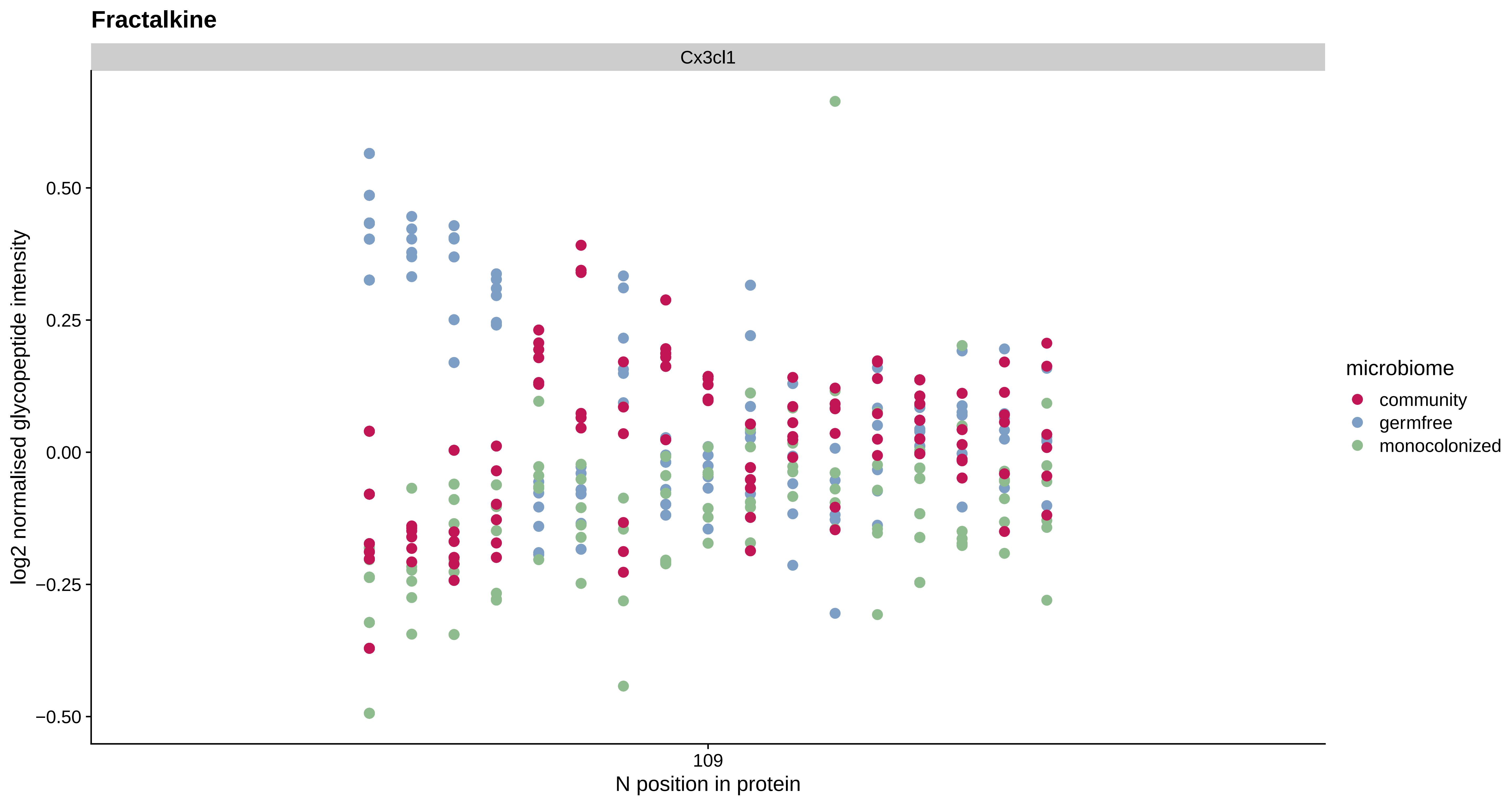
Fractalkine
Cx3cl1
0.50
0.25
0.00
−0.25
−0.50
log2 normalised glycopeptide intensity
109
N position in protein
microbiome
community
germfree
monocolonized

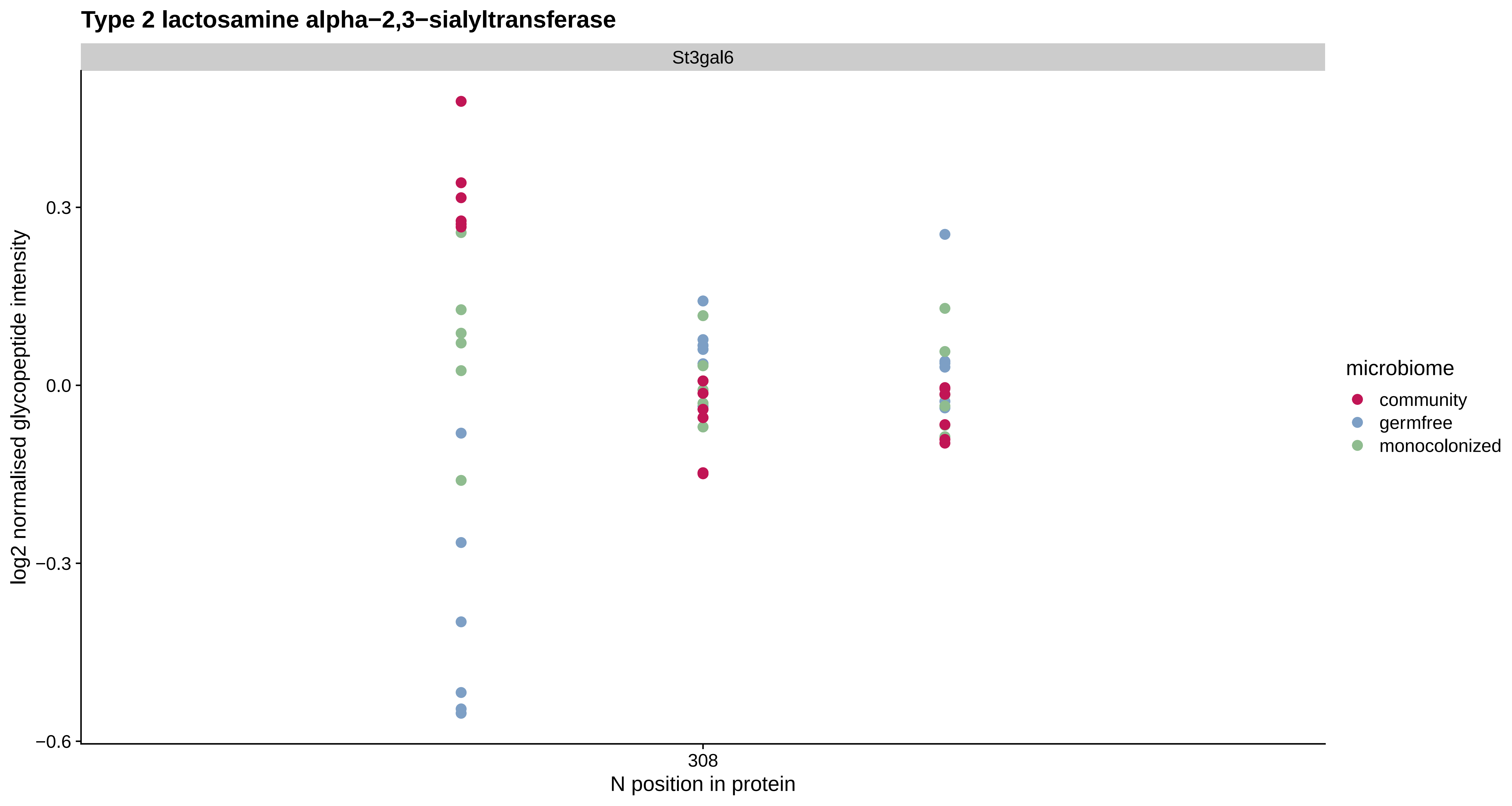
Type 2 lactosamine alpha−2,3−sialyltransferase
St3gal6
log2 normalised glycopeptide intensity
0.3
0.0
−0.3
−0.6
308
N position in protein
microbiome
community
germfree
monocolonized

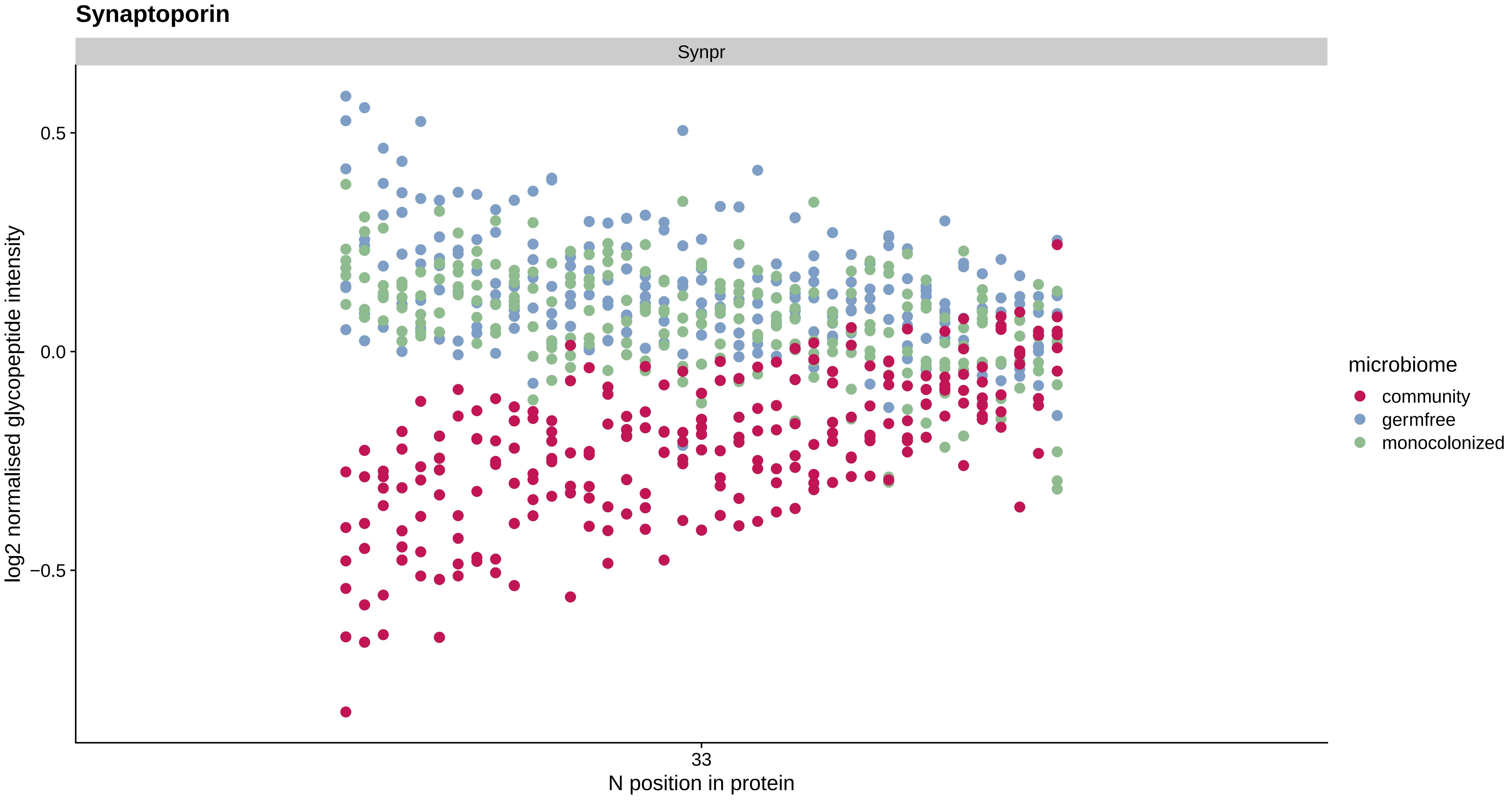
Synaptoporin
Synpr
log2 normalised glycopeptide intensity
0.5
0.0
−0.5
33
N position in protein
microbiome
community
germfree
monocolonized

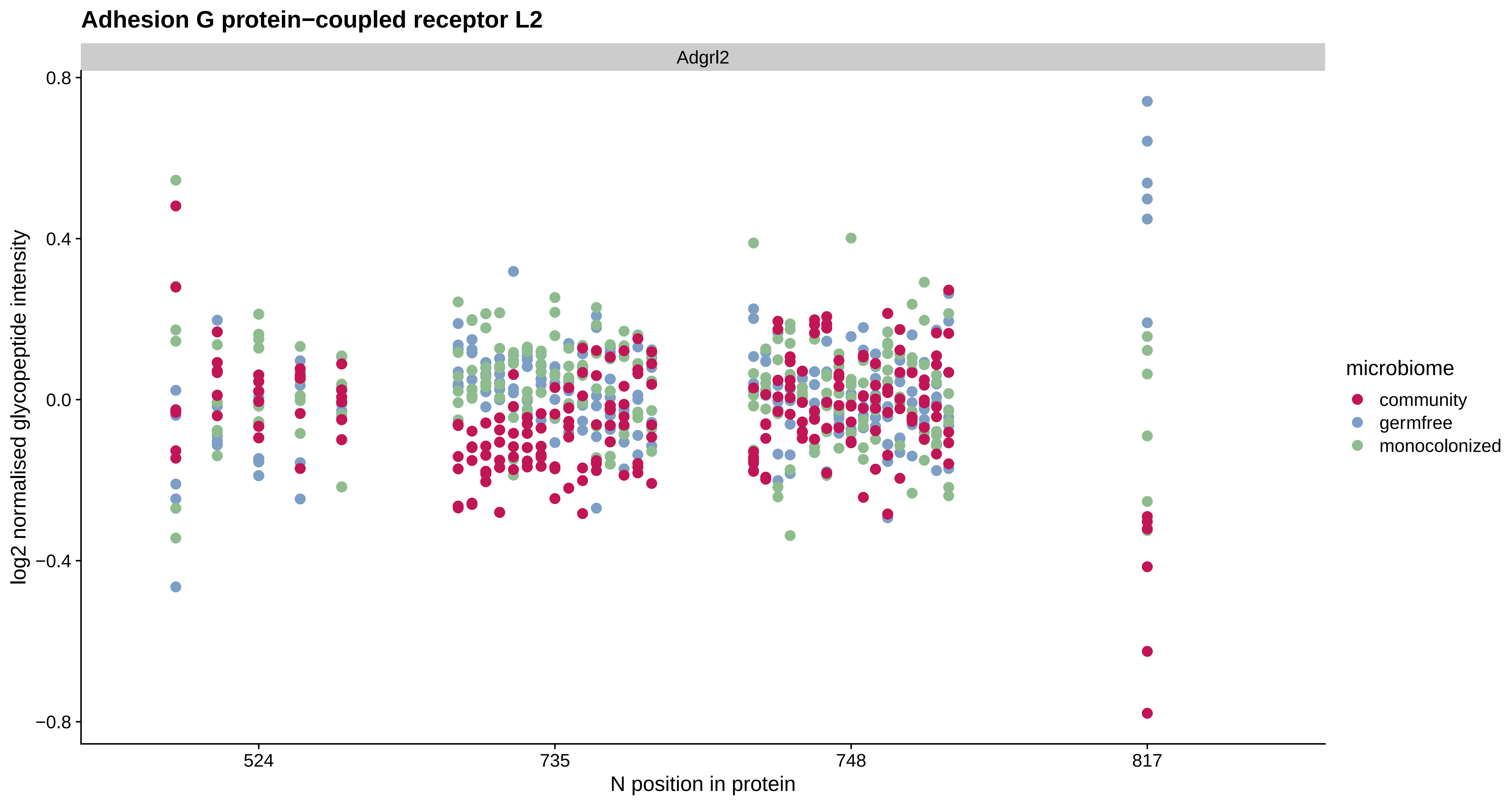

Adhesion G protein−coupled receptor L2
Adgrl2
0.8
0.4
0.0
−0.4
−0.8
log2 normalised glycopeptide intensity
524
735
748
817
N position in protein
microbiome
community
germfree
monocolonized

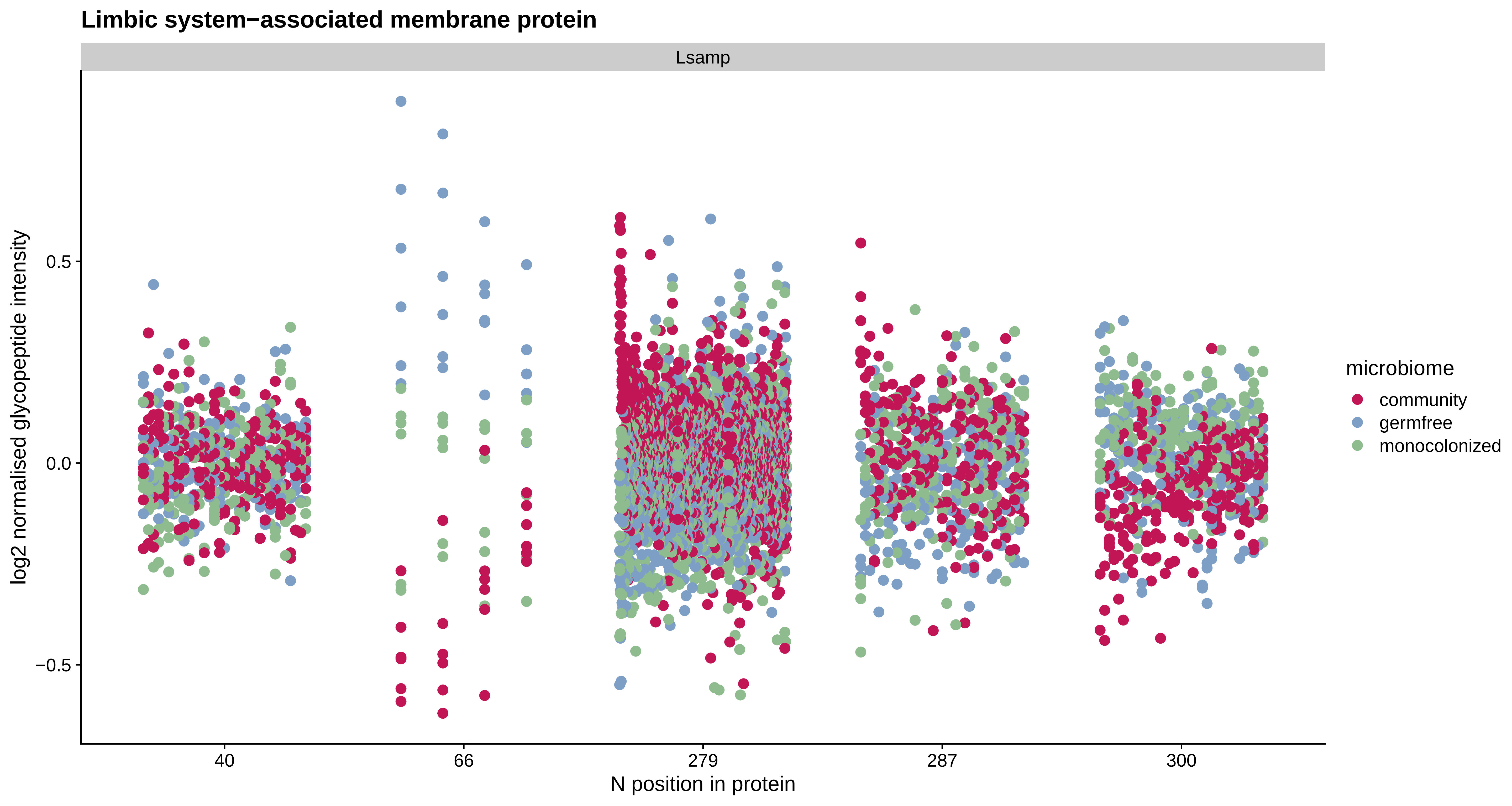

Limbic system−associated membrane protein
Lsamp
log2 normalised glycopeptide intensity
0.5
0.0
−0.5
40
66
279
287
300
N position in protein
microbiome
community
germfree
monocolonized

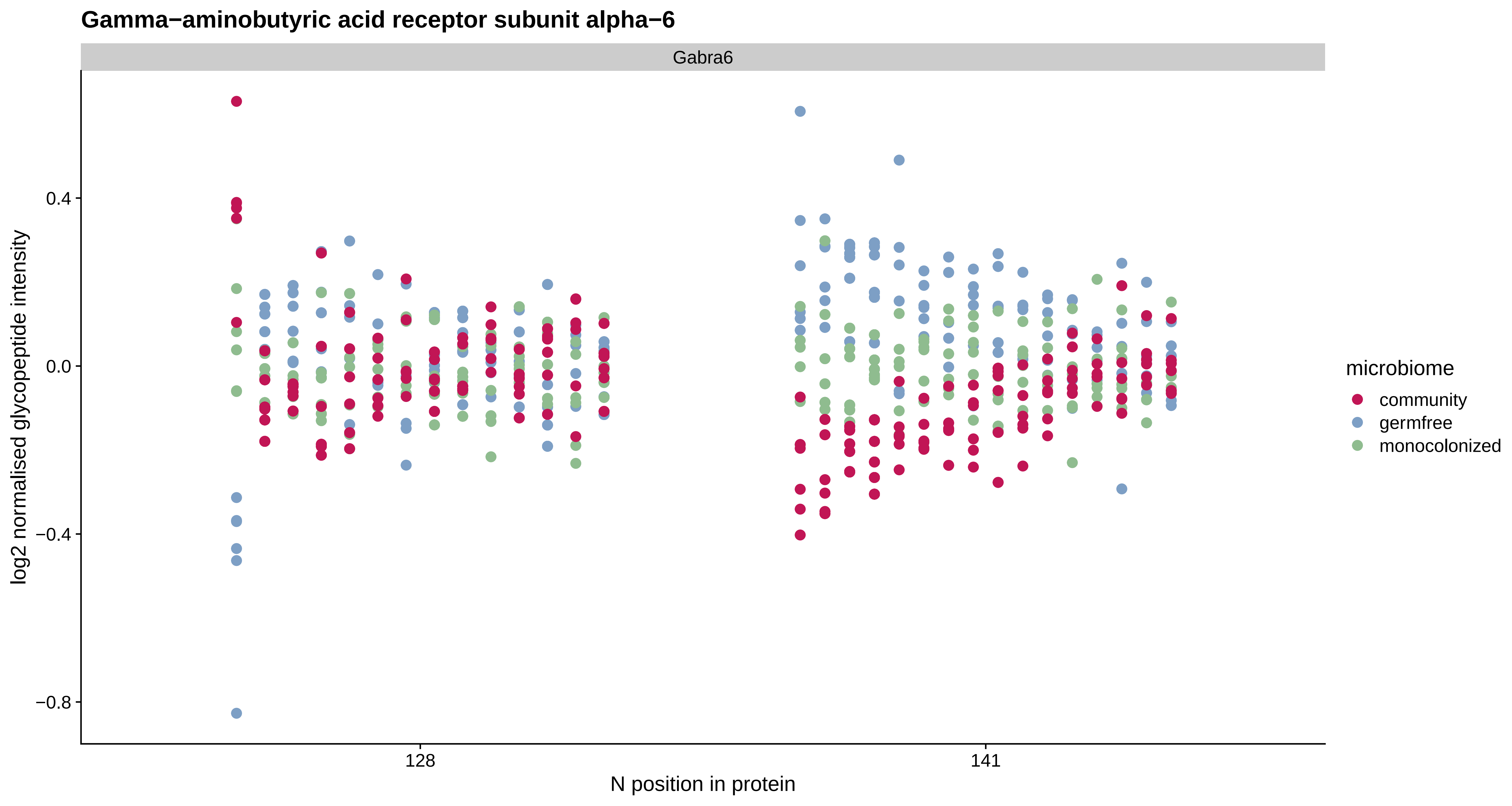
Gamma−aminobutyric acid receptor subunit alpha−6
Gabra6
log2 normalised glycopeptide intensity
0.4
0.0
−0.4
−0.8
128
141
N position in protein
microbiome
community
germfree
monocolonized

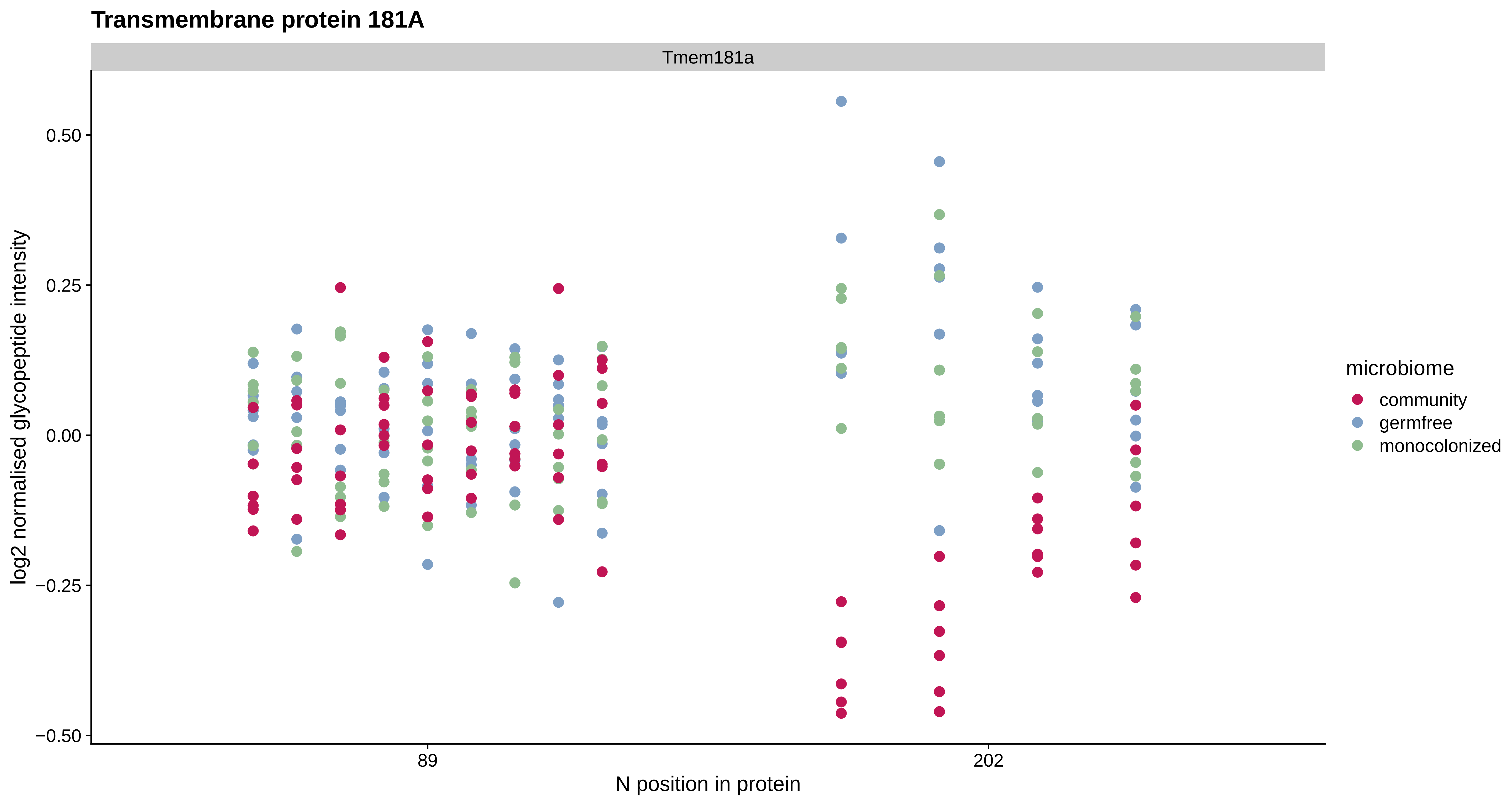
Transmembrane protein 181A
Tmem181a
log2 normalised glycopeptide intensity
0.50
0.25
0.00
−0.25
−0.50
89
202
N position in protein
microbiome
community
germfree
monocolonized

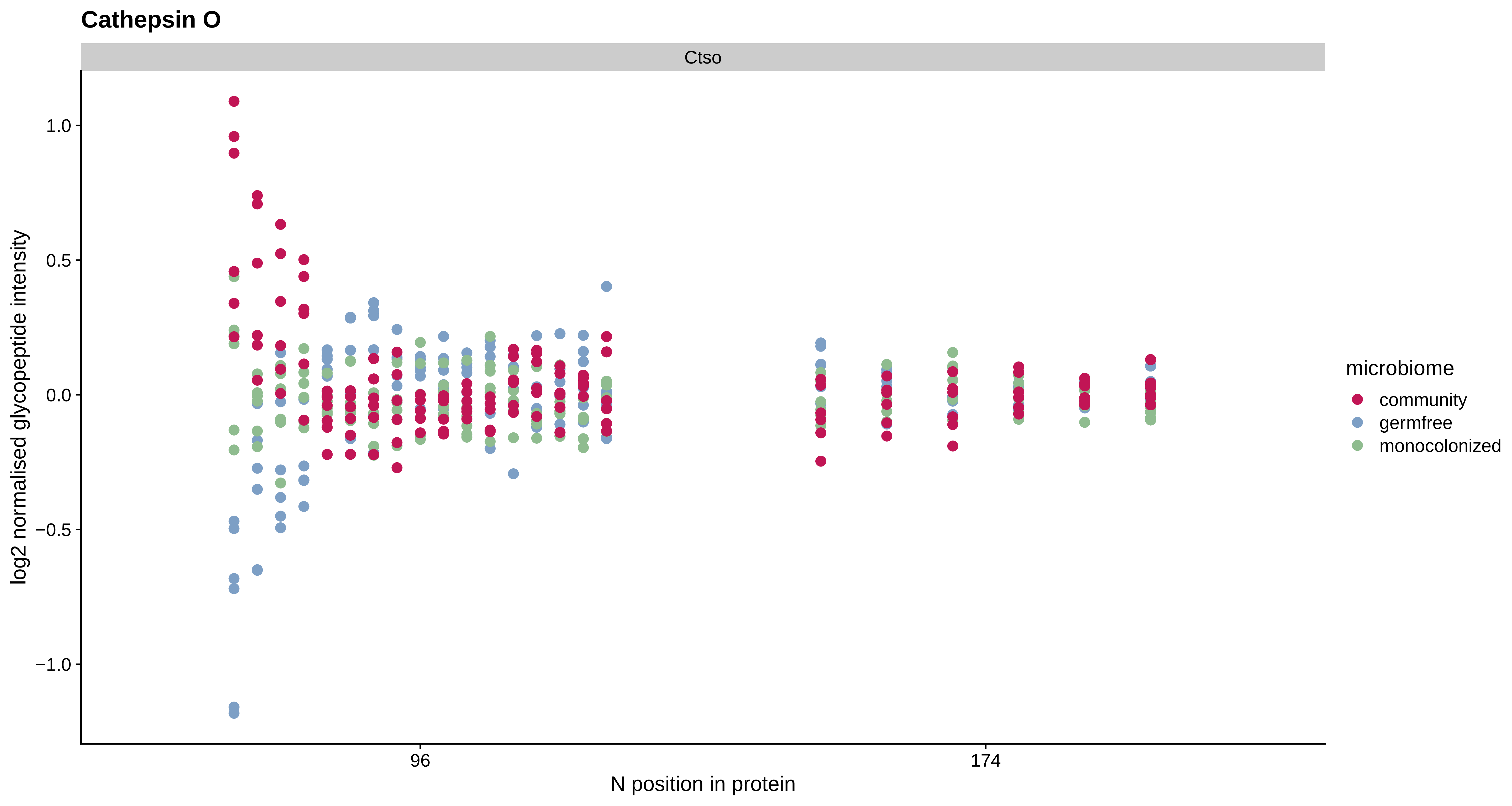

Cathepsin O
Ctso
log2 normalised glycopeptide intensity
1.0
0.5
0.0
−0.5
−1.0
96
174
N position in protein
microbiome
community
germfree
monocolonized

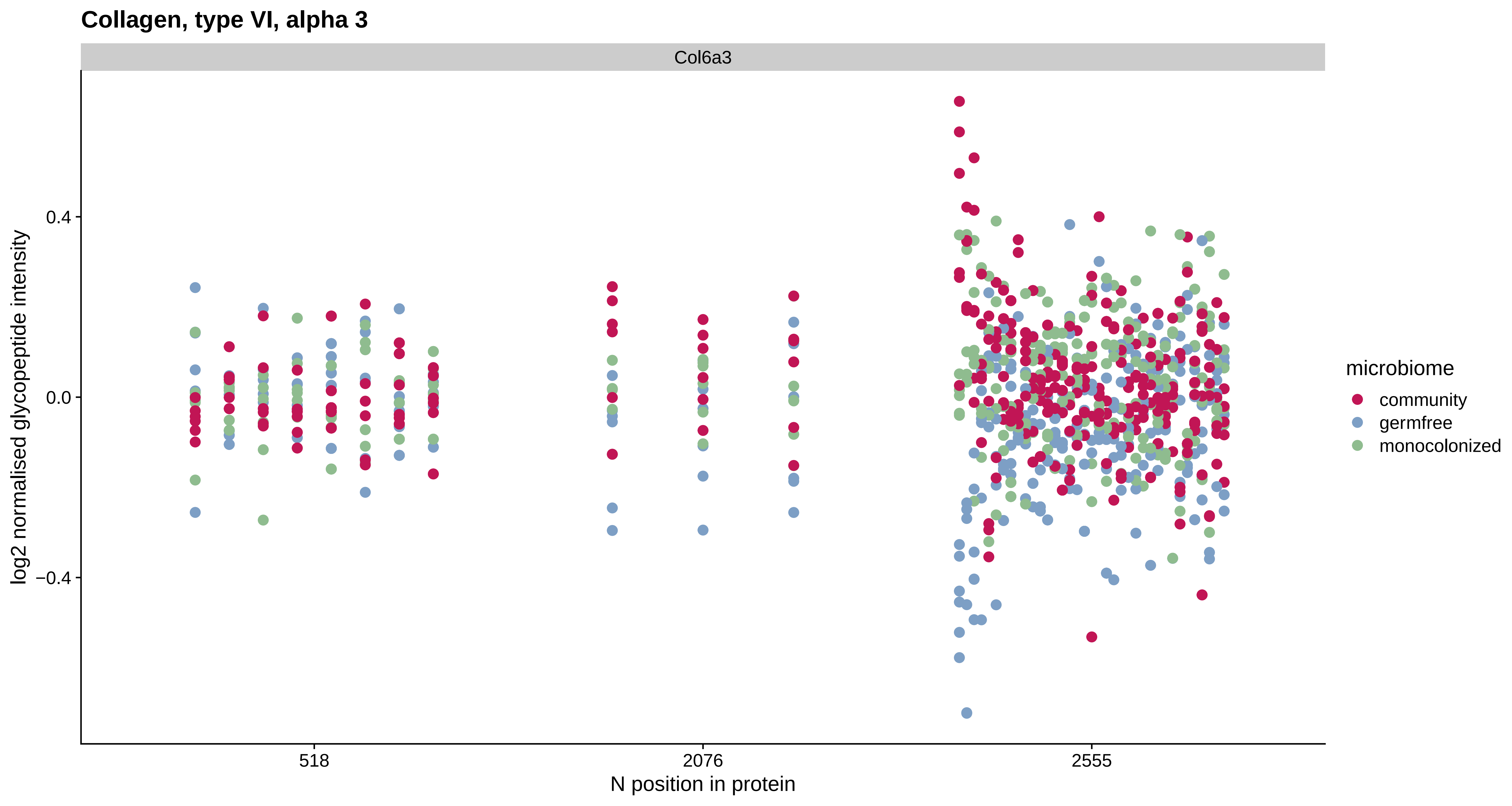

Collagen, type VI, alpha 3
Col6a3
log2 normalised glycopeptide intensity
0.4
0.0
−0.4
518
2076
2555
N position in protein
microbiome
community
germfree
monocolonized

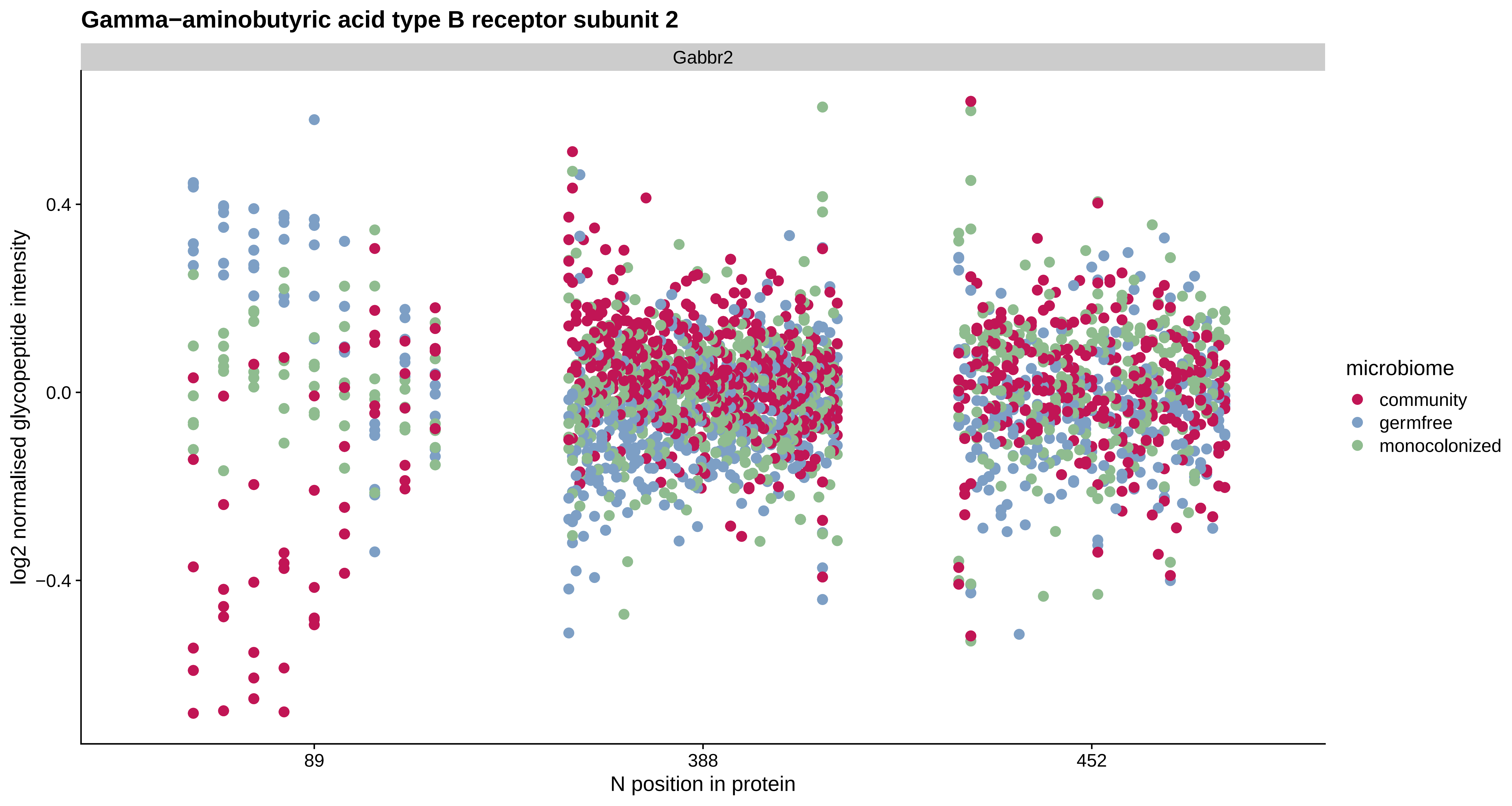

Gamma−aminobutyric acid type B receptor subunit 2
Gabbr2
log2 normalised glycopeptide intensity
0.4
0.0
−0.4
89
388
452
N position in protein
microbiome
community
germfree
monocolonized

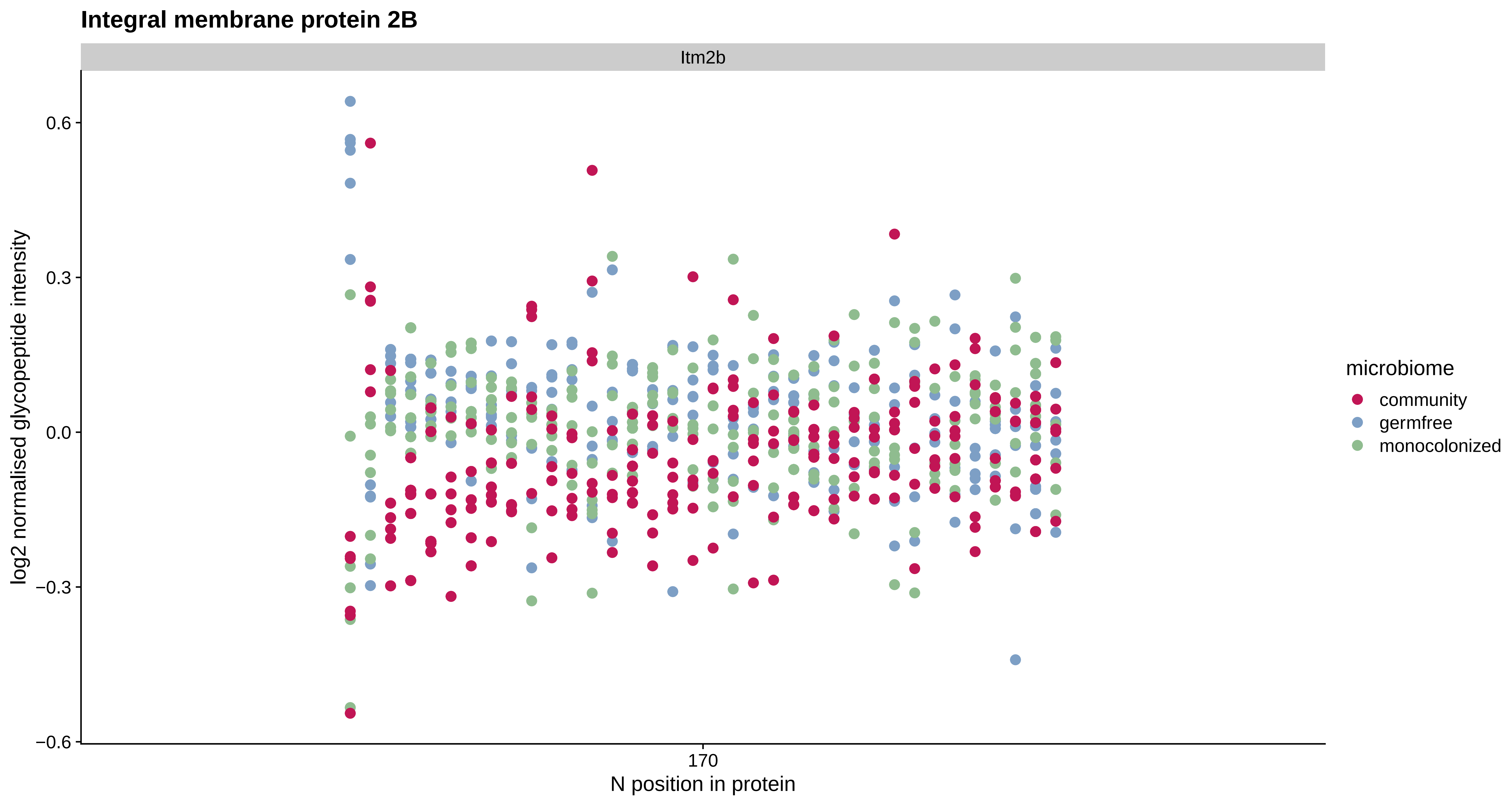

Integral membrane protein 2B
Itm2b
0.6
0.3
0.0
−0.3
−0.6
log2 normalised glycopeptide intensity
170
N position in protein
microbiome
community
germfree
monocolonized

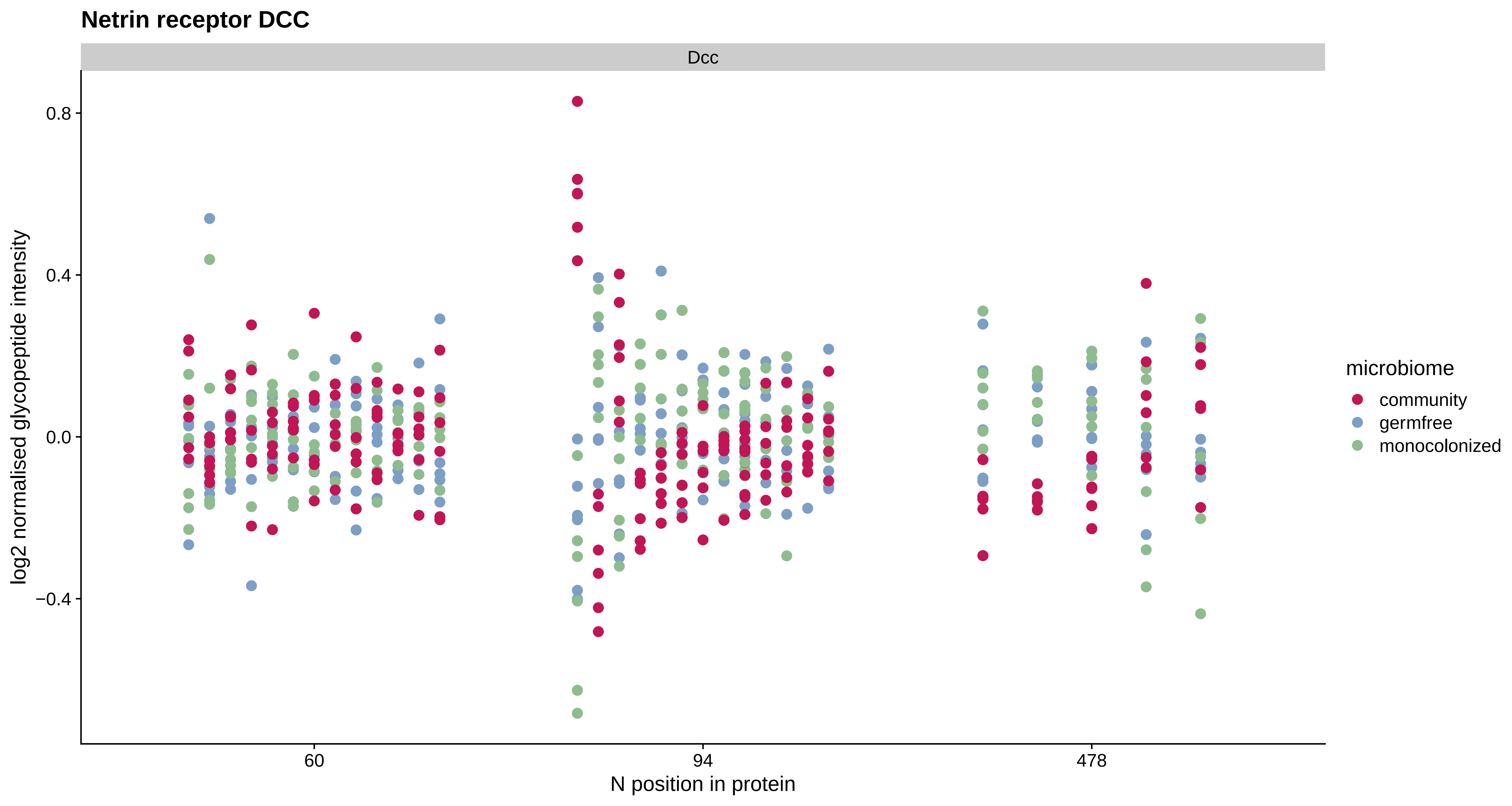

Netrin receptor DCC
Dcc
0.8
0.4
0.0
−0.4
log2 normalised glycopeptide intensity
60
94
478
N position in protein
microbiome
community
germfree
monocolonized

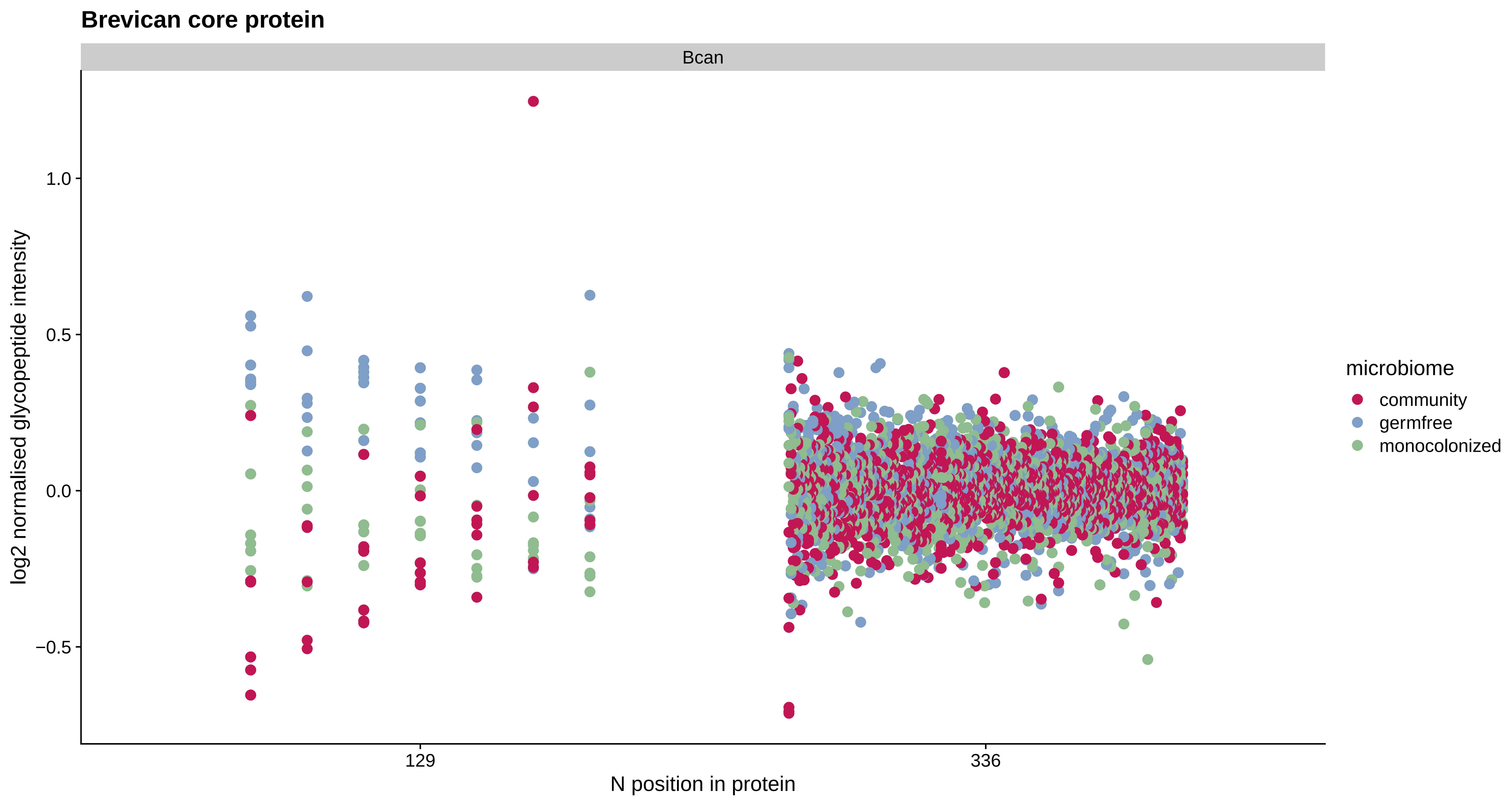
Brevican core protein
Bcan
log2 normalised glycopeptide intensity
1.0
0.5
0.0
−0.5
129
336
N position in protein
microbiome
community
germfree
monocolonized

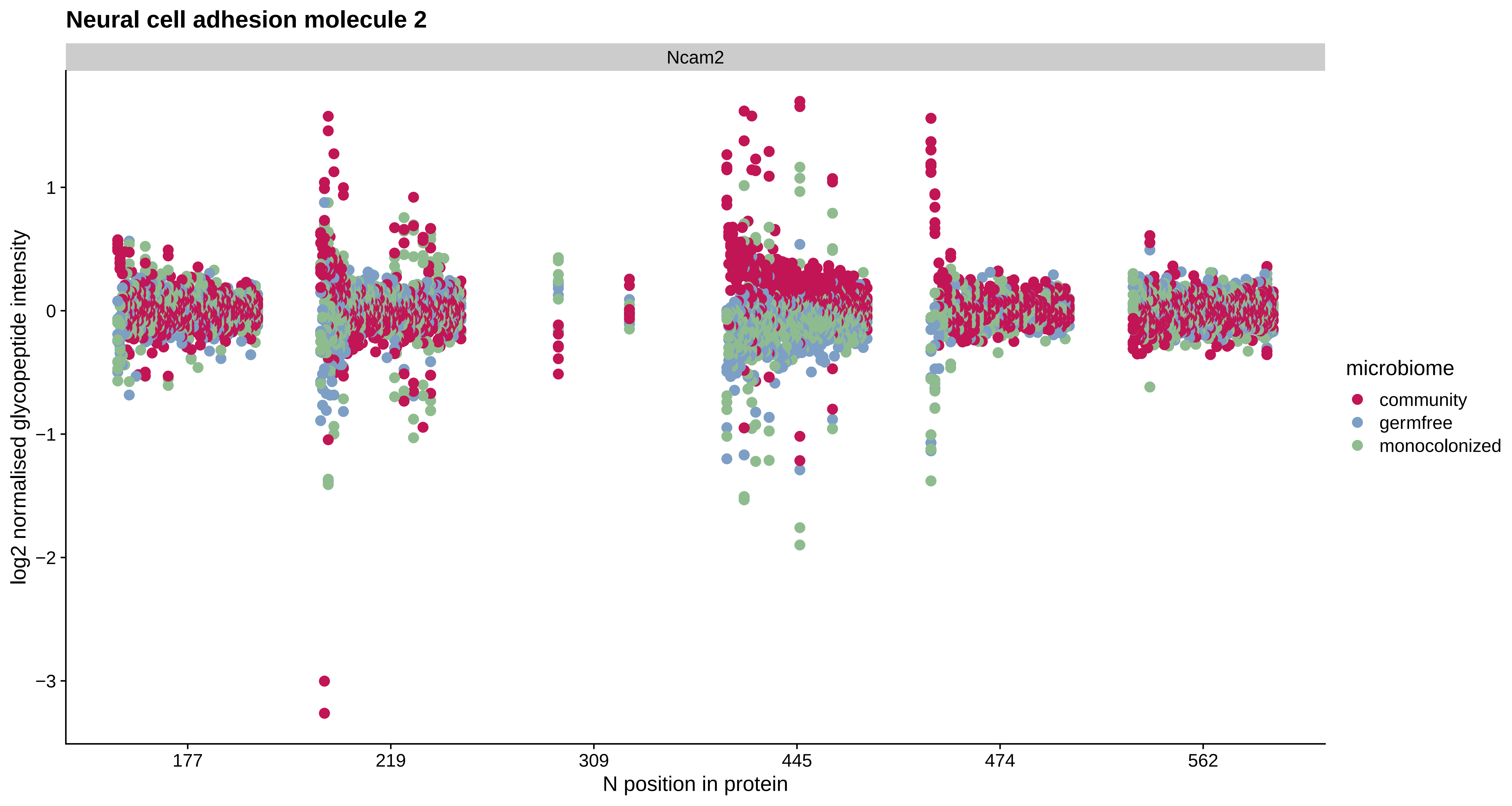
Neural cell adhesion molecule 2
Ncam2
log2 normalised glycopeptide intensity
1
0
−1
−2
−3
177
219
309
445
474
562
N position in protein
microbiome
community
germfree
monocolonized

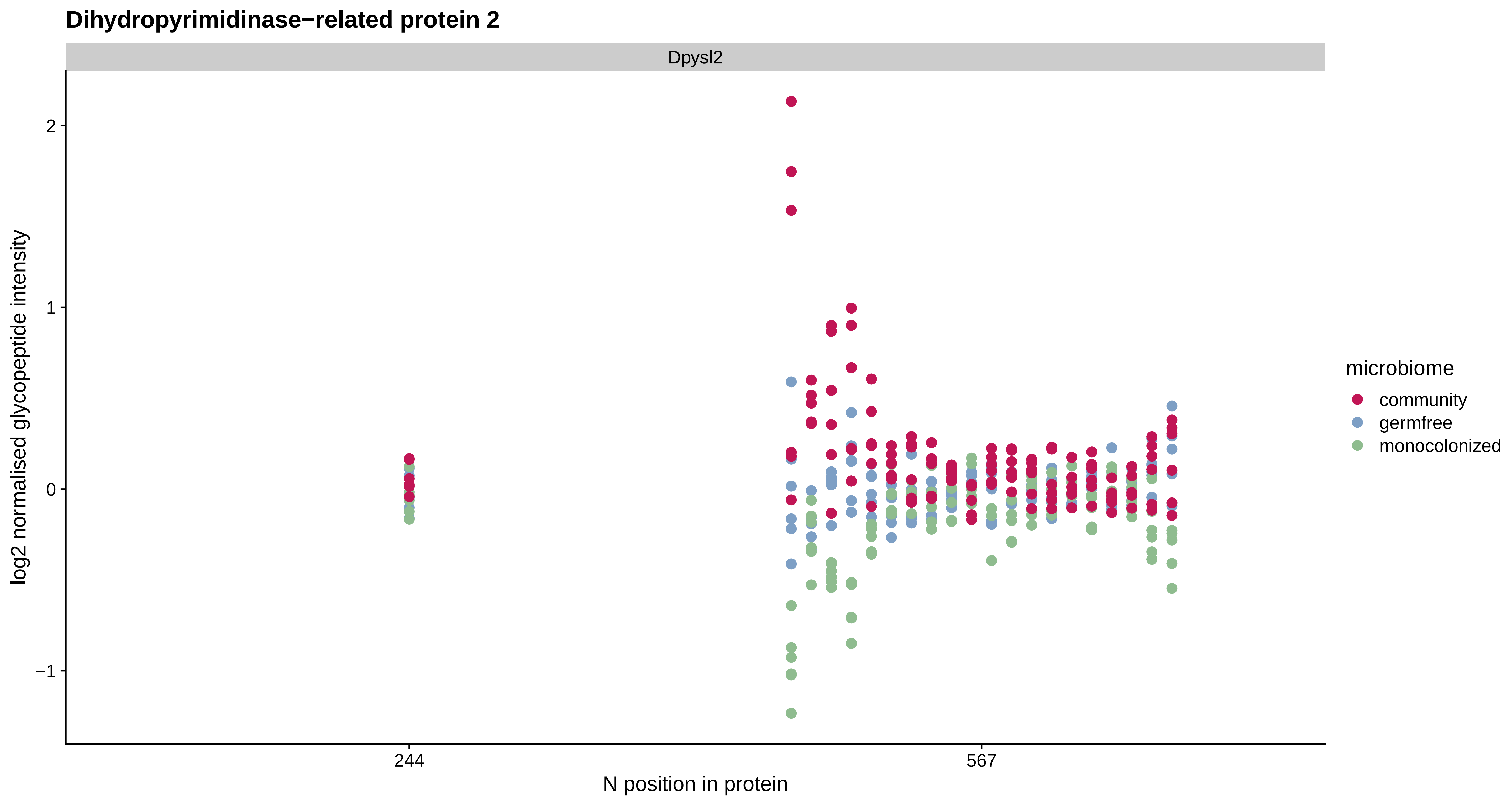
Dihydropyrimidinase−related protein 2
Dpysl2
log2 normalised glycopeptide intensity
2
1
0
−1
244
567
N position in protein
microbiome
community
germfree
monocolonized

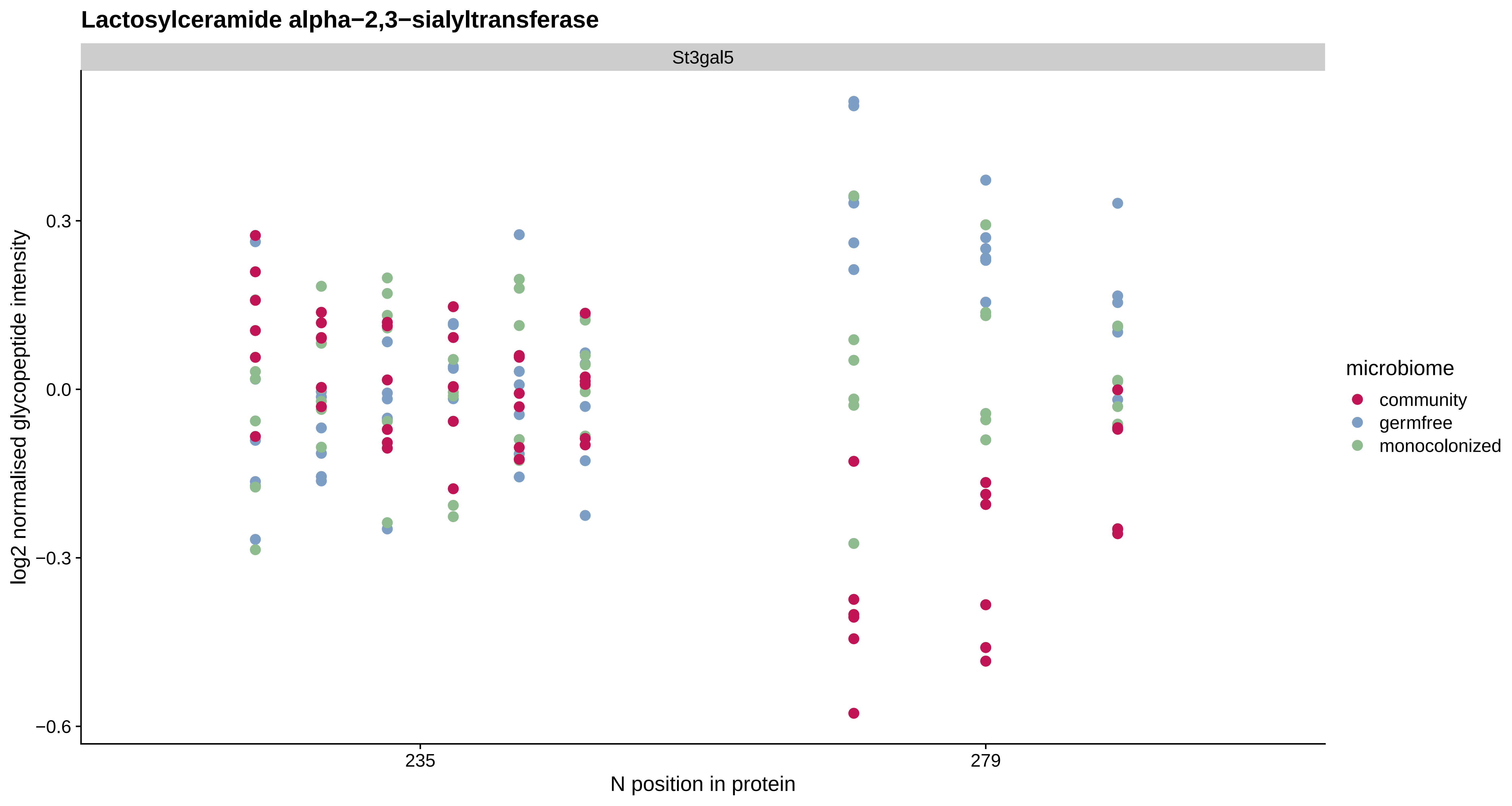

Lactosylceramide alpha−2,3−sialyltransferase
St3gal5
0.3
0.0
−0.3
−0.6
log2 normalised glycopeptide intensity
235
279
N position in protein
microbiome
community
germfree
monocolonized

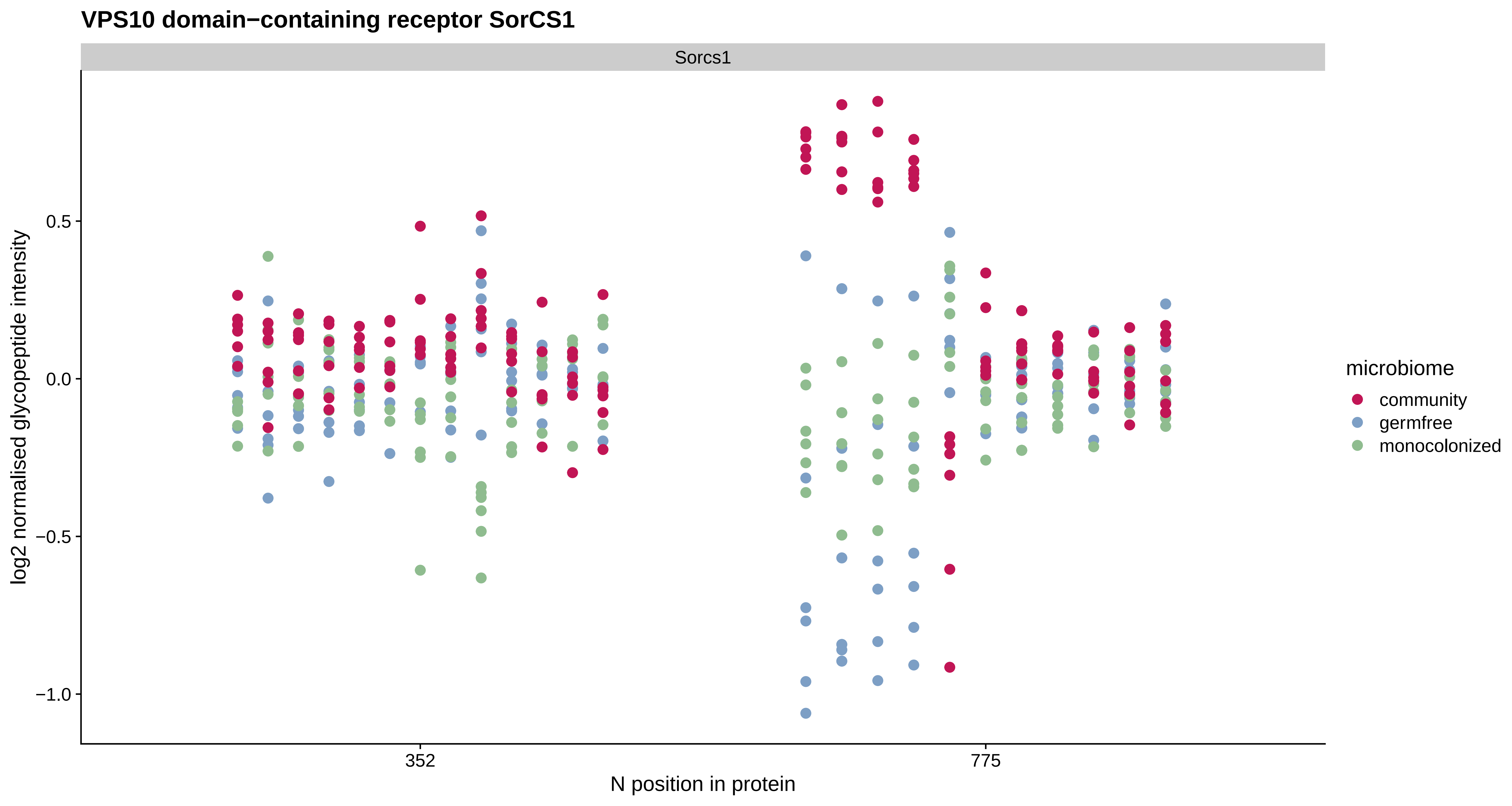
VPS10 domain−containing receptor SorCS1
Sorcs1
log2 normalised glycopeptide intensity
0.5
0.0
−0.5
−1.0
352
775
N position in protein
microbiome
community
germfree
monocolonized

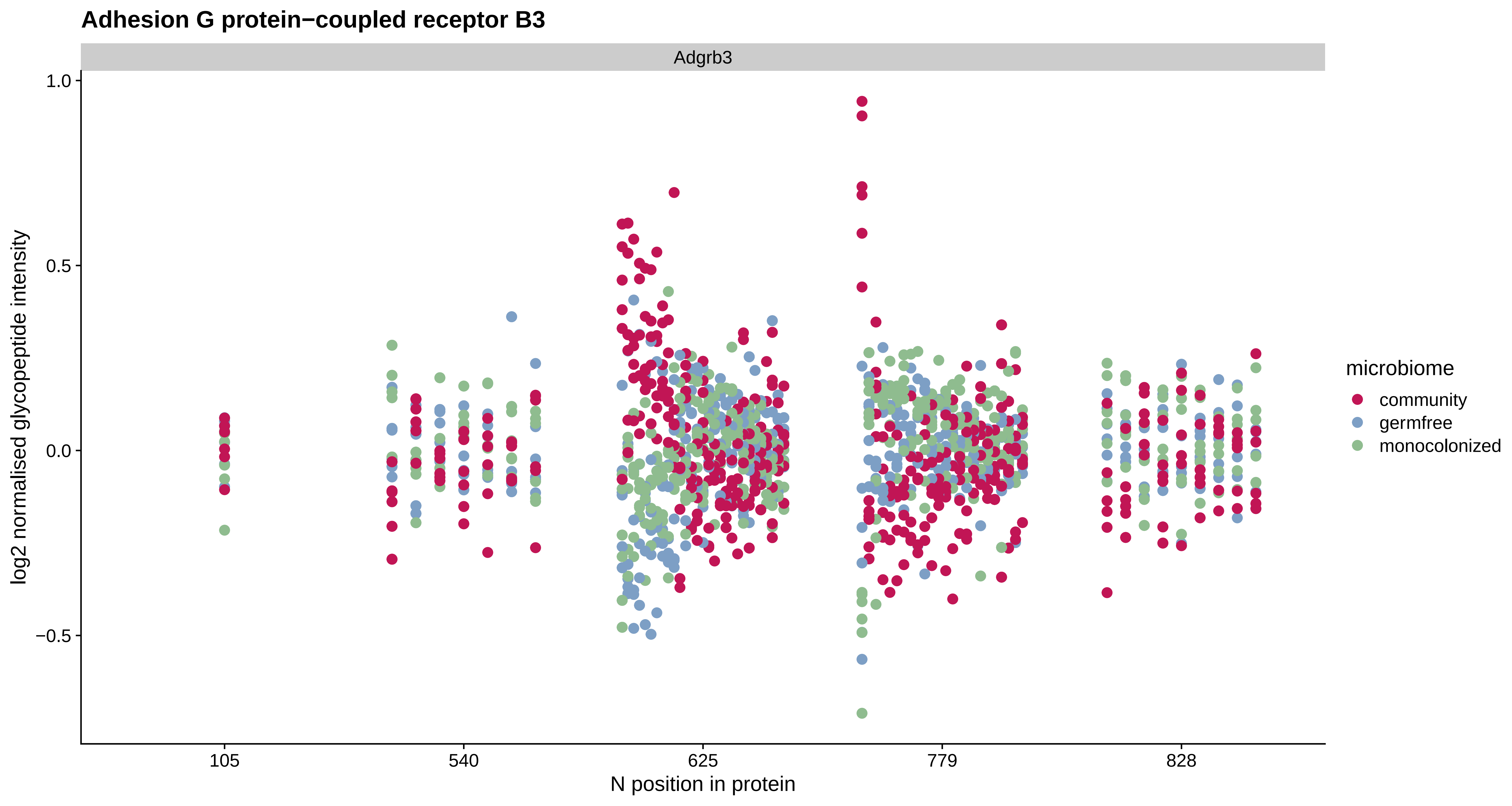

Adhesion G protein−coupled receptor B3
Adgrb3
log2 normalised glycopeptide intensity
1.0
0.5
0.0
−0.5
105
540
625
779
828
N position in protein
microbiome
community
germfree
monocolonized

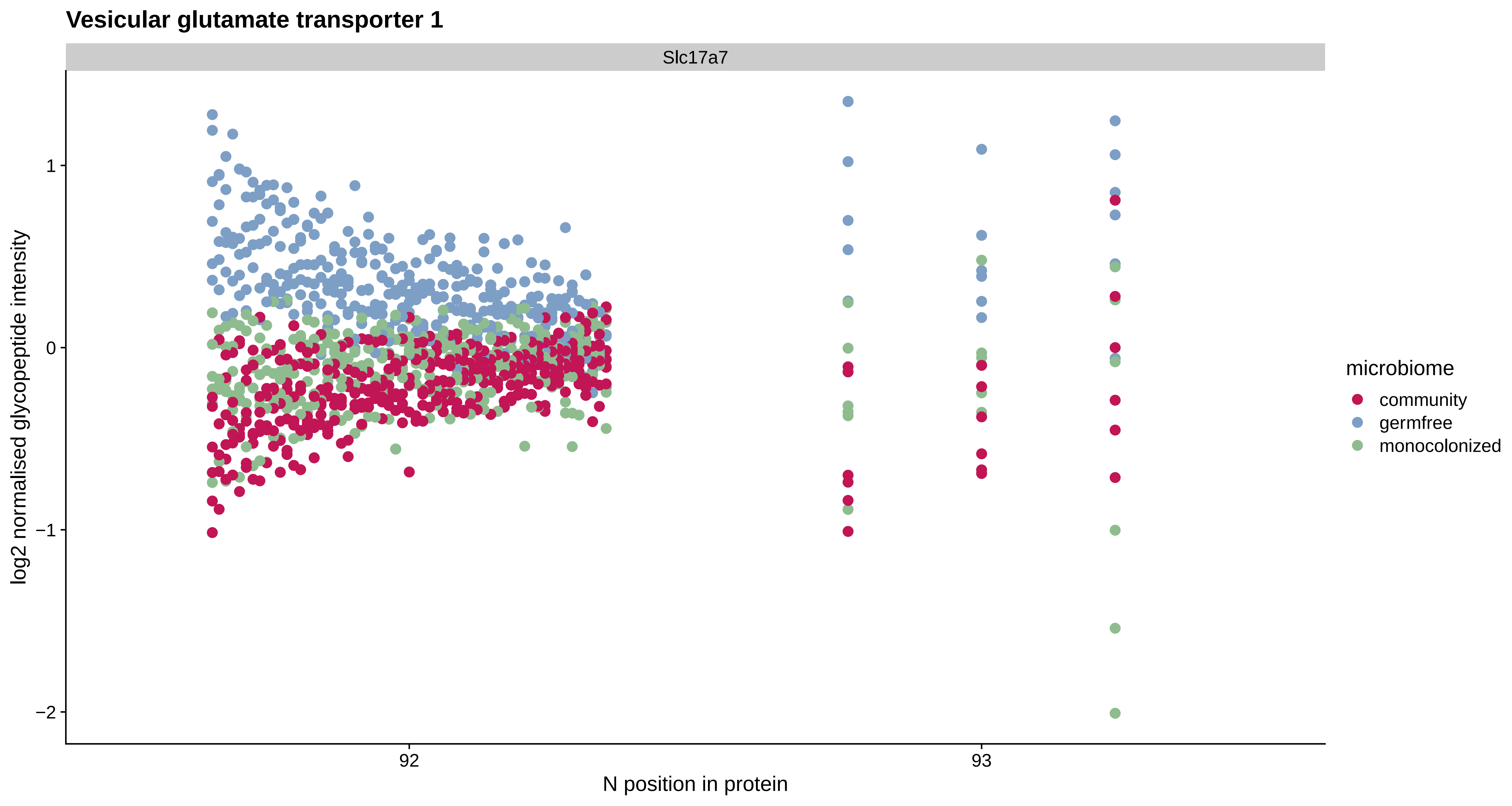

Vesicular glutamate transporter 1
Slc17a7
log2 normalised glycopeptide intensity
1
0
−1
−2
92
93
N position in protein
microbiome
community
germfree
monocolonized

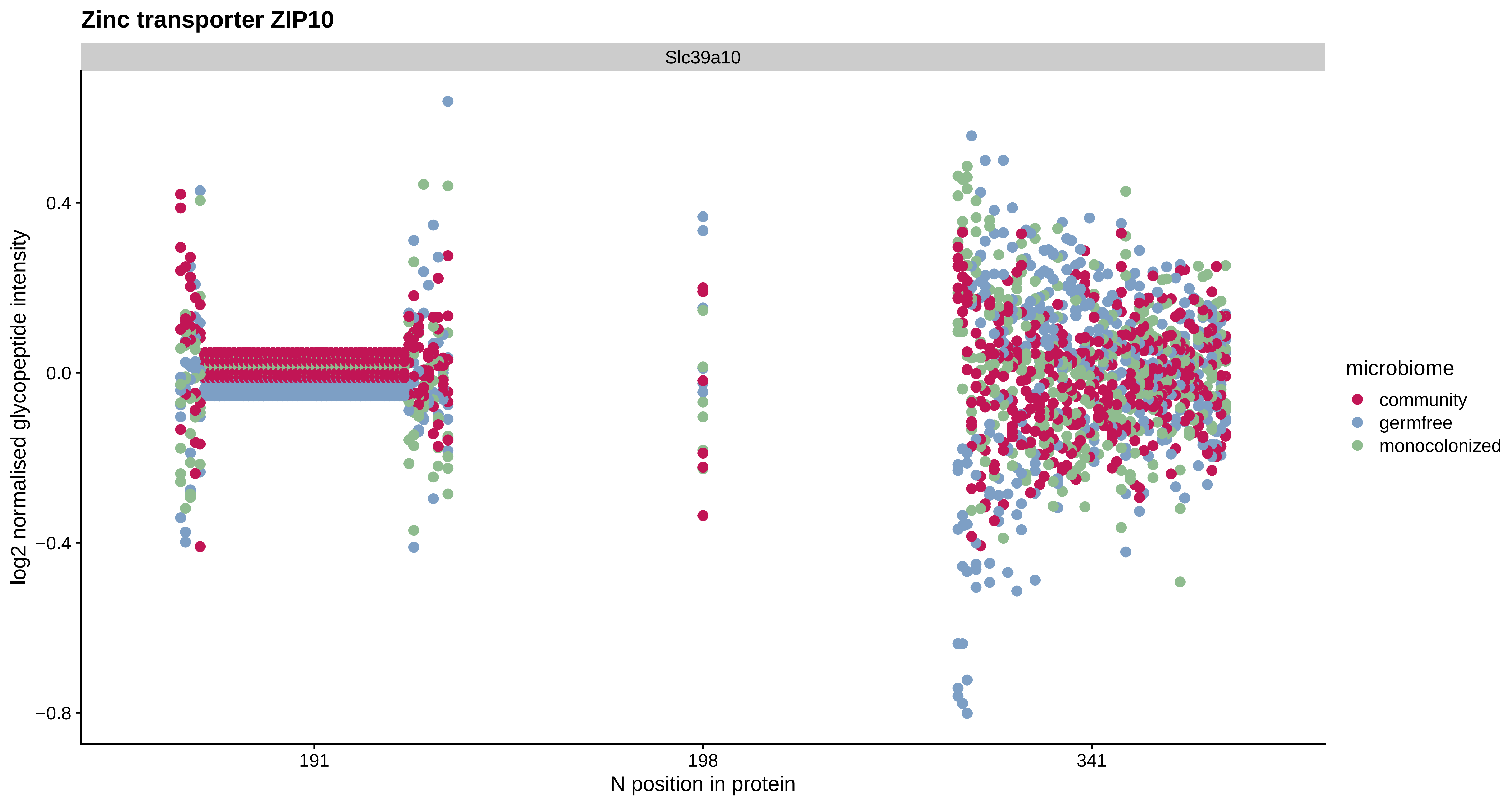

Zinc transporter ZIP10
Slc39a10
log2 normalised glycopeptide intensity
0.4
0.0
−0.4
−0.8
191
198
341
N position in protein
microbiome
community
germfree
monocolonized

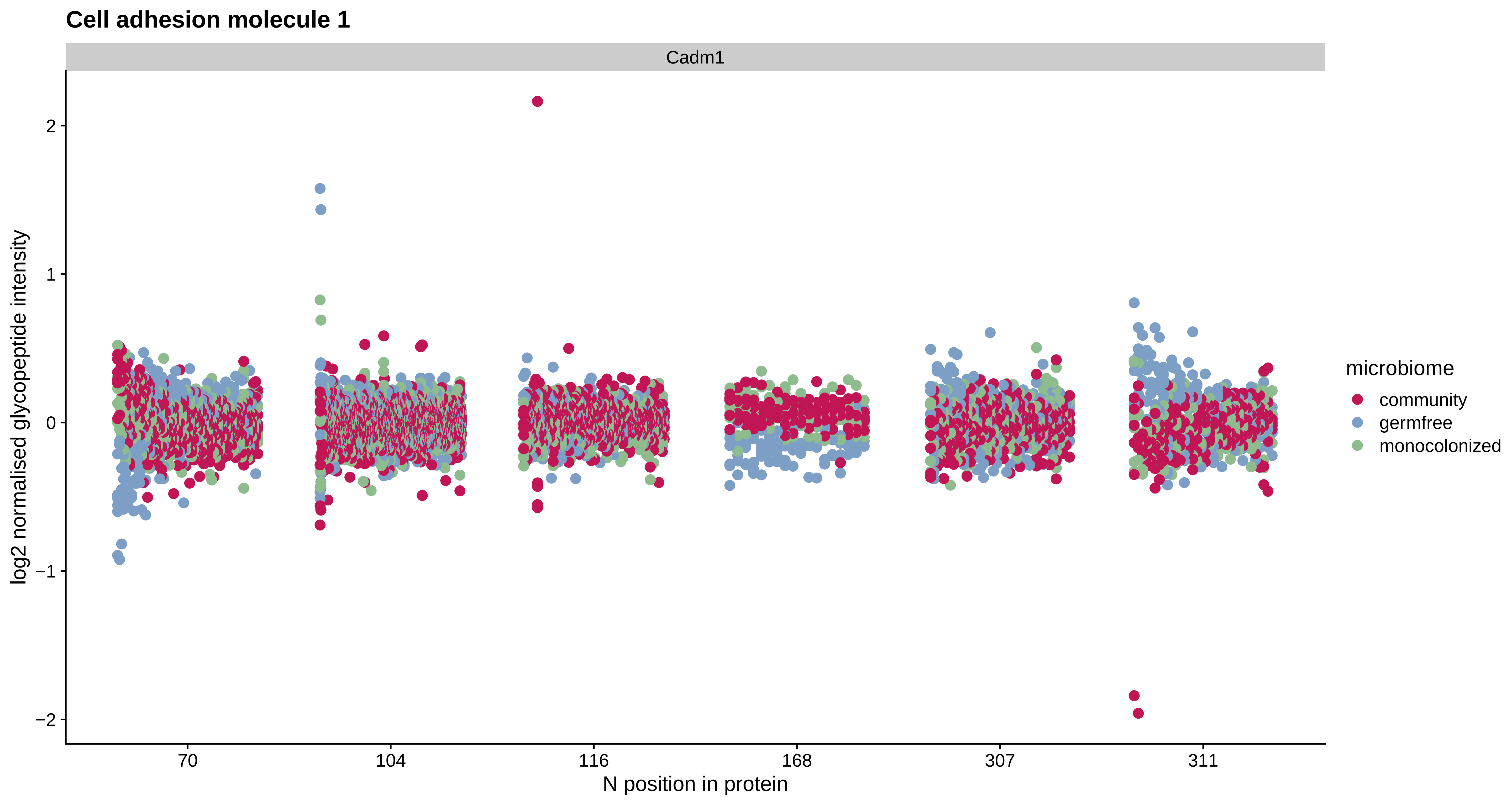

Cell adhesion molecule 1
Cadm1
log2 normalised glycopeptide intensity
2
1
0
−1
−2
70
104
116
168
307
311
N position in protein
microbiome
community
germfree
monocolonized

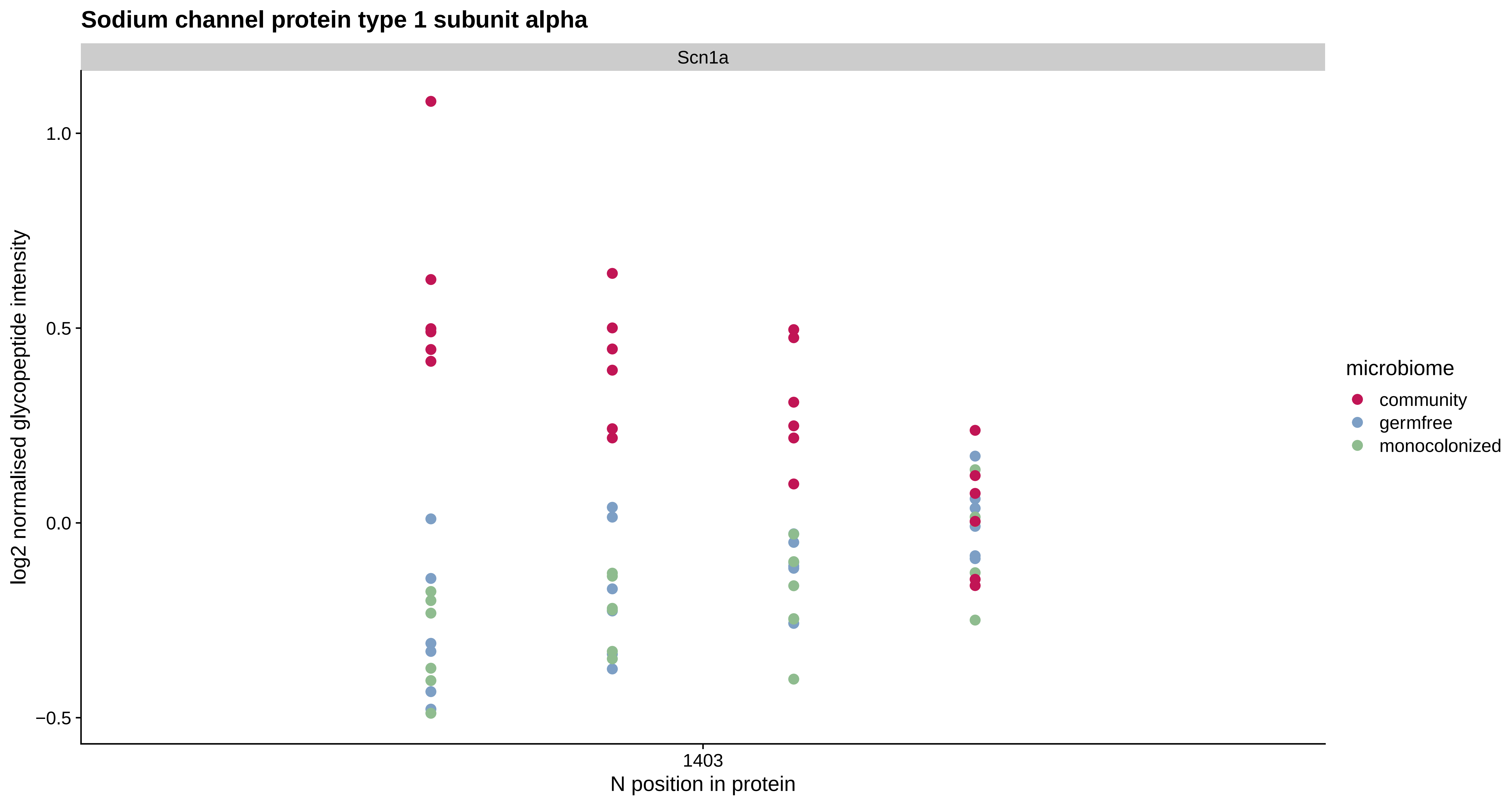

Sodium channel protein type 1 subunit alpha
Scn1a
1.0
0.5
0.0
−0.5
log2 normalised glycopeptide intensity
1403
N position in protein
microbiome
community
germfree
monocolonized

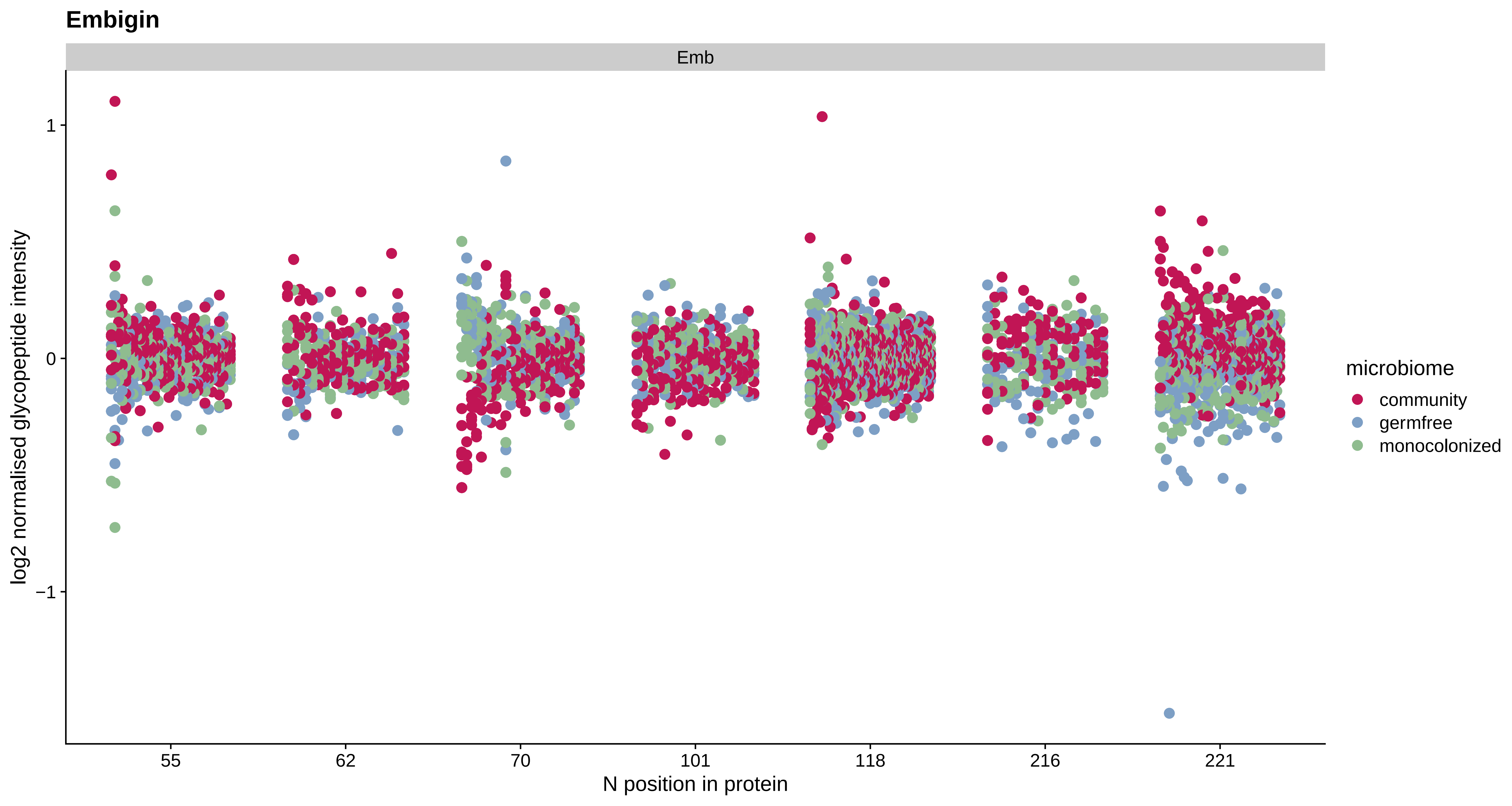

Embigin
Emb
log2 normalised glycopeptide intensity
1
0
−1
55
62
70
101
118
216
221
N position in protein
microbiome
community
germfree
monocolonized

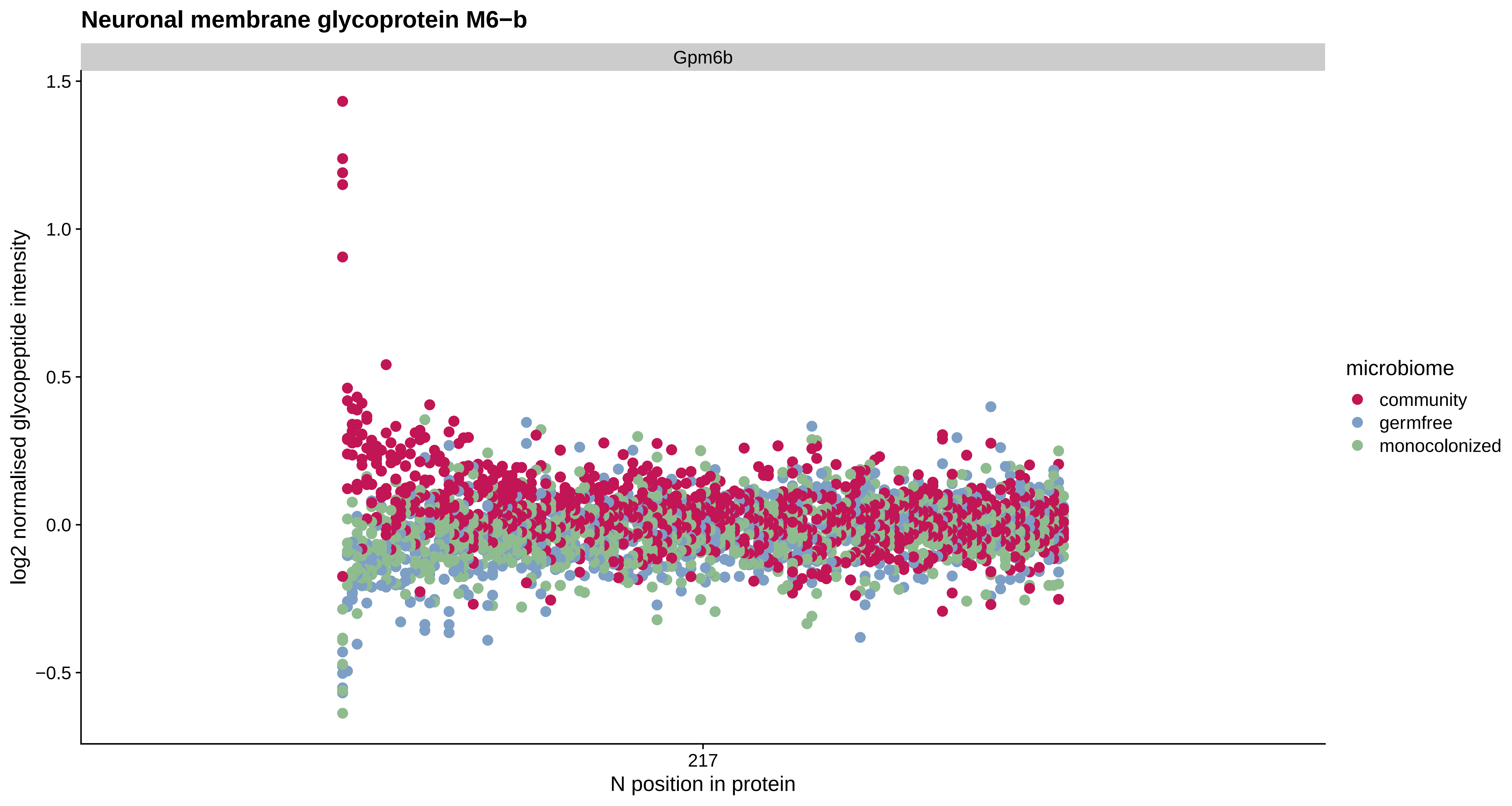

Neuronal membrane glycoprotein M6−b
Gpm6b
1.5
1.0
0.5
0.0
−0.5
log2 normalised glycopeptide intensity
217
N position in protein
microbiome
community
germfree
monocolonized

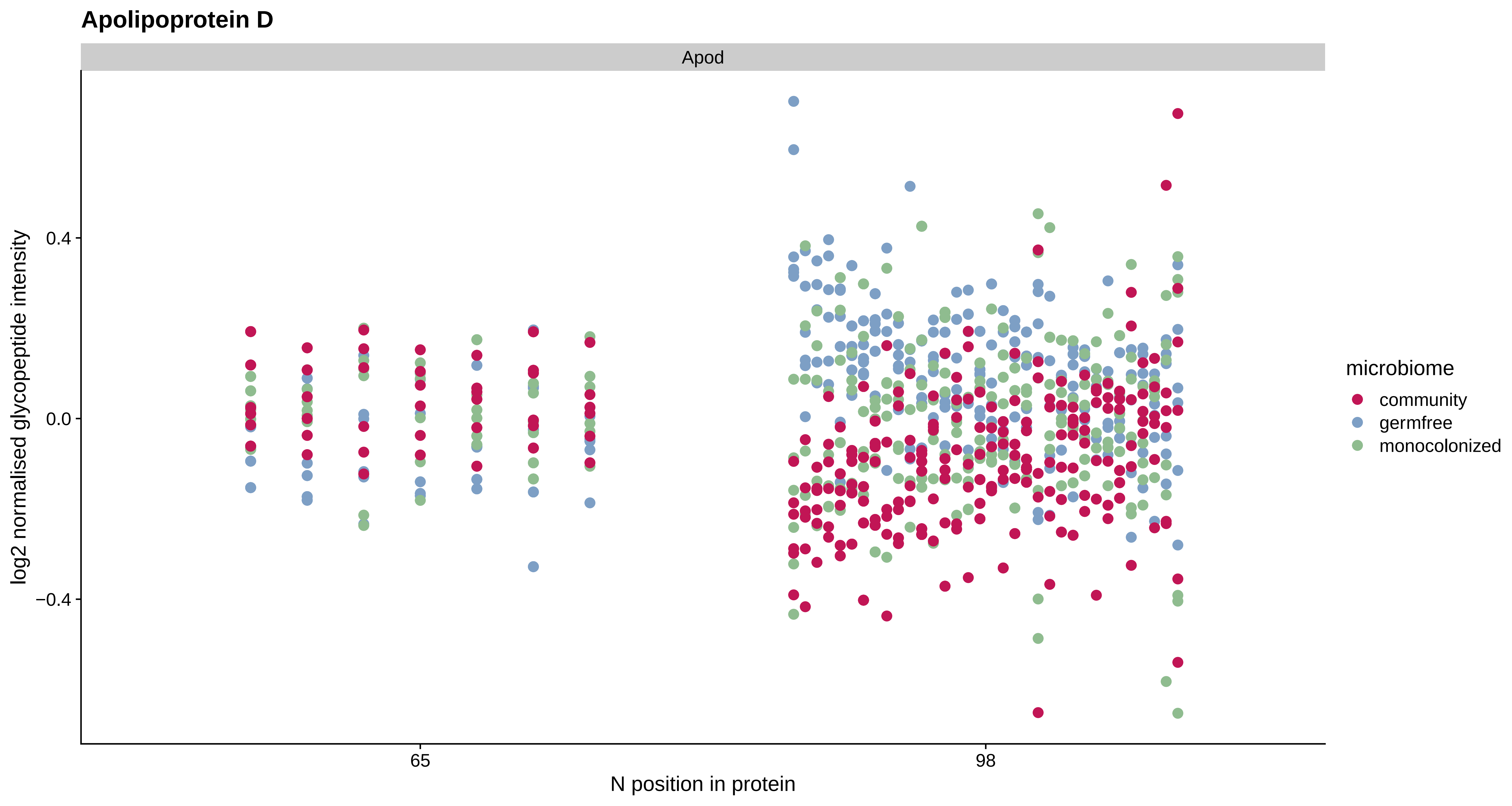
Apolipoprotein D
Apod
log2 normalised glycopeptide intensity
0.4
0.0
−0.4
65
98
N position in protein
microbiome
community
germfree
monocolonized

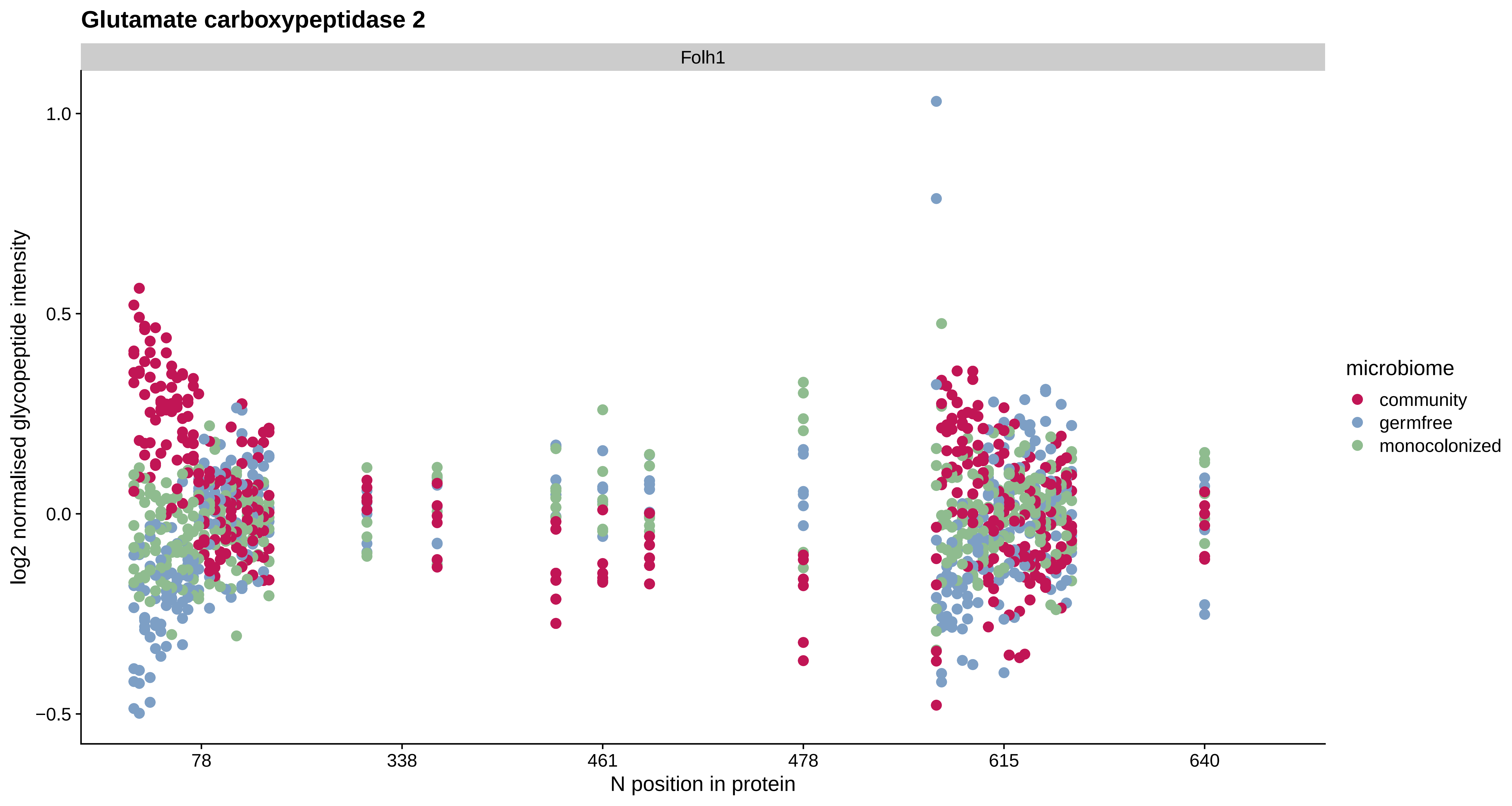

Glutamate carboxypeptidase 2
Folh1
1.0
0.5
0.0
−0.5
log2 normalised glycopeptide intensity
78
338
461
478
615
640
N position in protein
microbiome
community
germfree
monocolonized

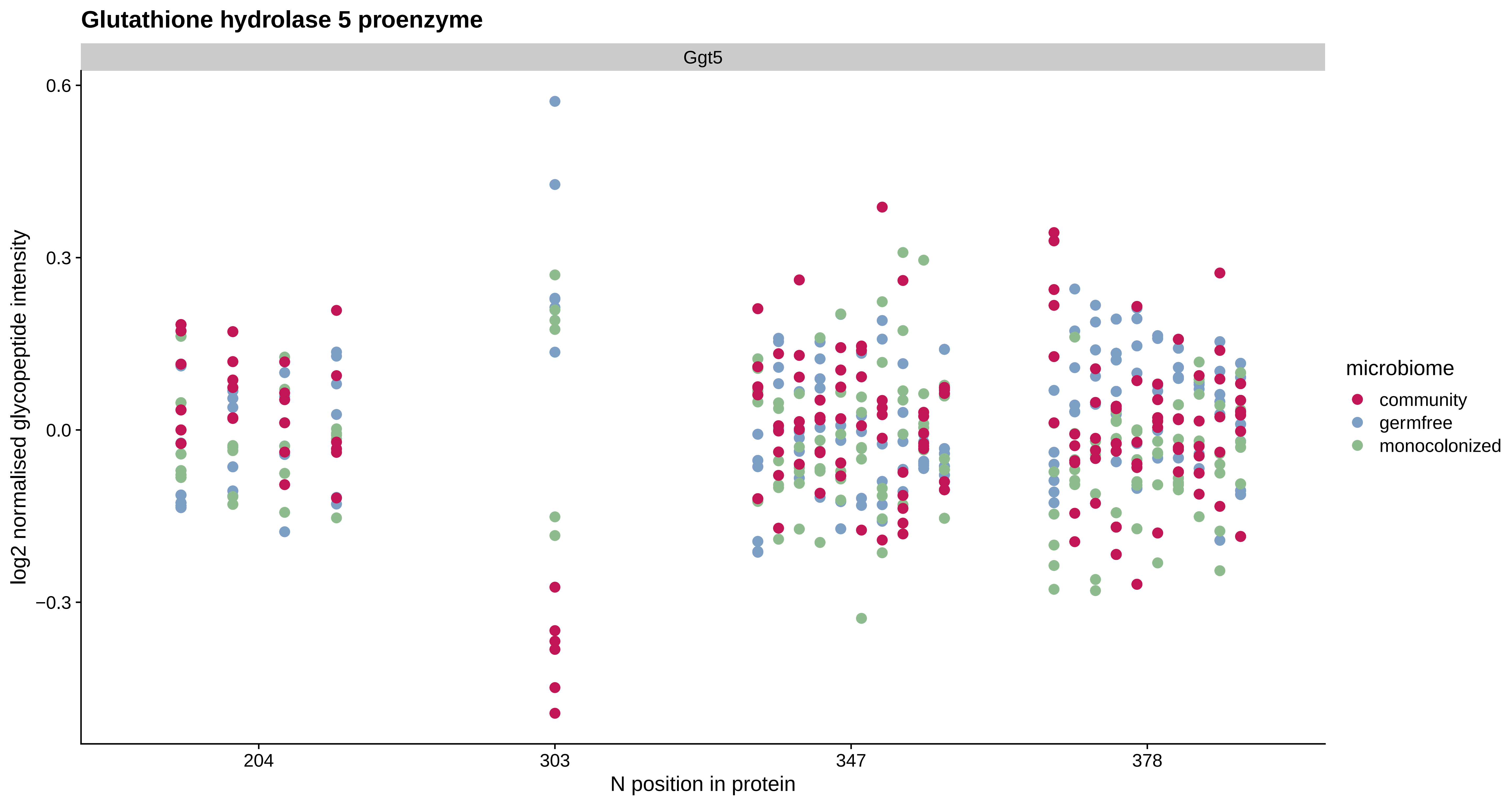
Glutathione hydrolase 5 proenzyme
Ggt5
0.6
0.3
0.0
−0.3
log2 normalised glycopeptide intensity
204
303
347
378
N position in protein
microbiome
community
germfree
monocolonized

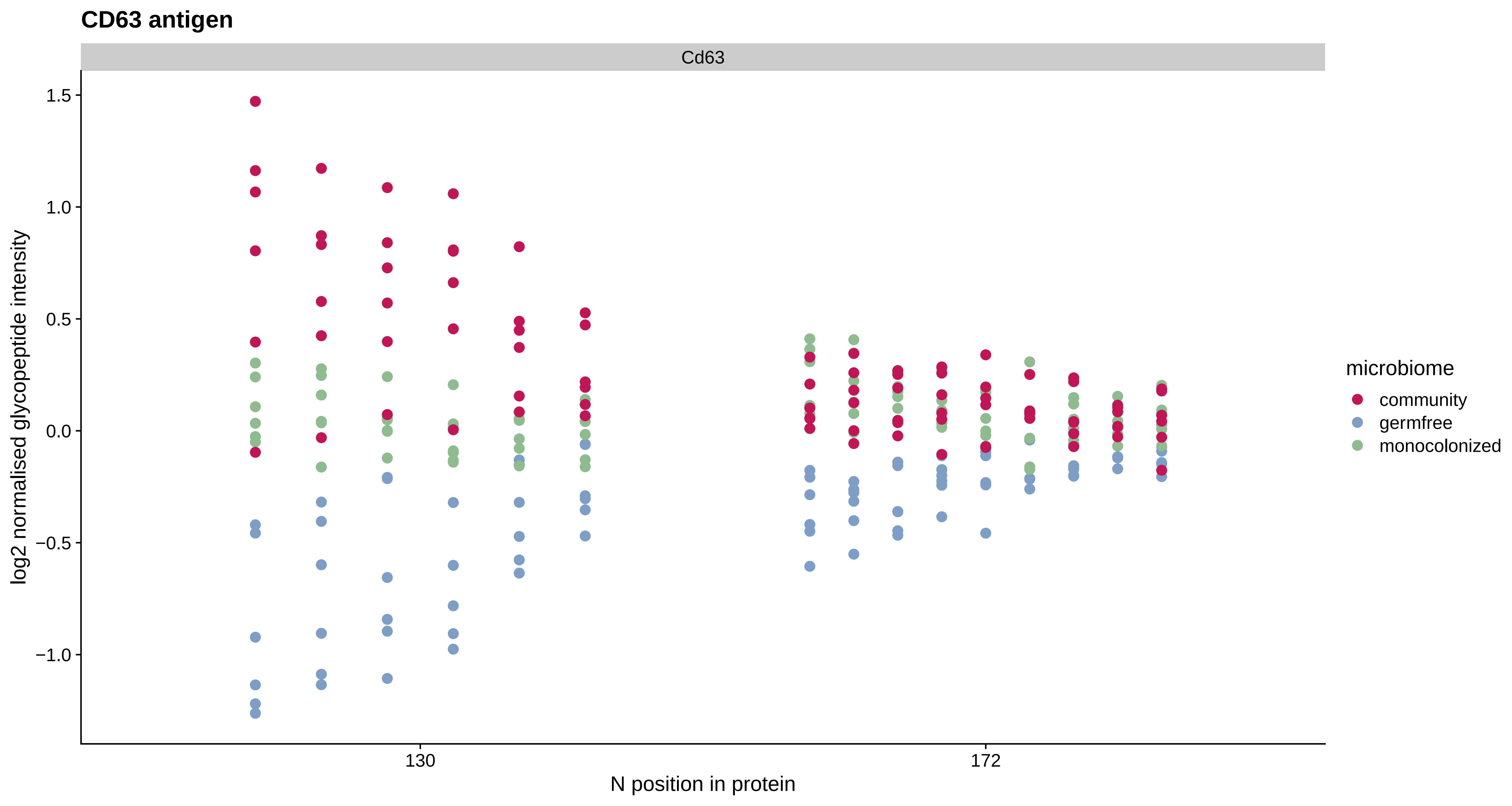

CD63 antigen
Cd63
1.5
1.0
0.5
0.0
−0.5
−1.0
log2 normalised glycopeptide intensity
130
172
N position in protein
microbiome
community
germfree
monocolonized

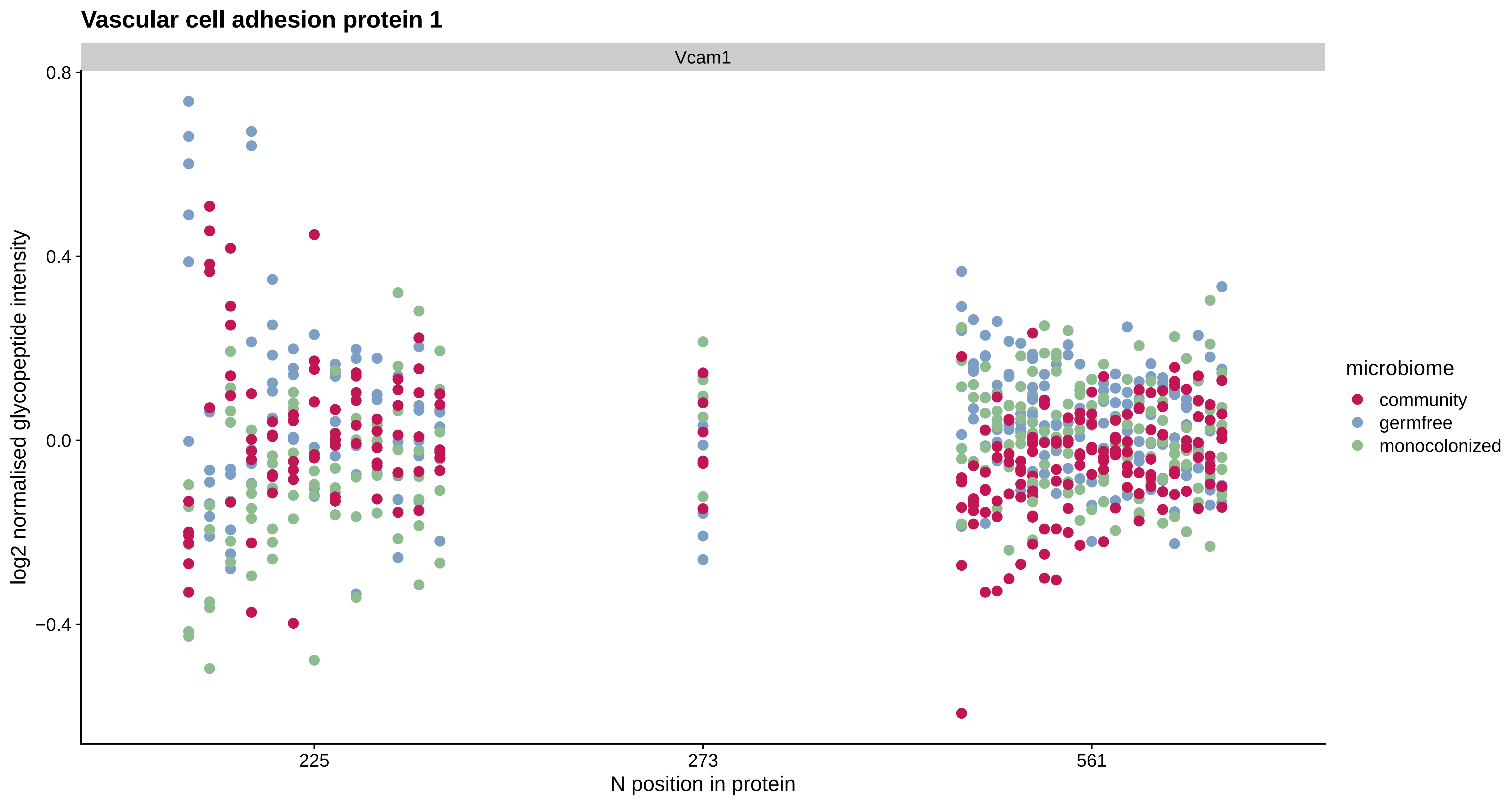
Vascular cell adhesion protein 1
Vcam1
0.8
0.4
0.0
−0.4
log2 normalised glycopeptide intensity
225
273
561
N position in protein
microbiome
community
germfree
monocolonized

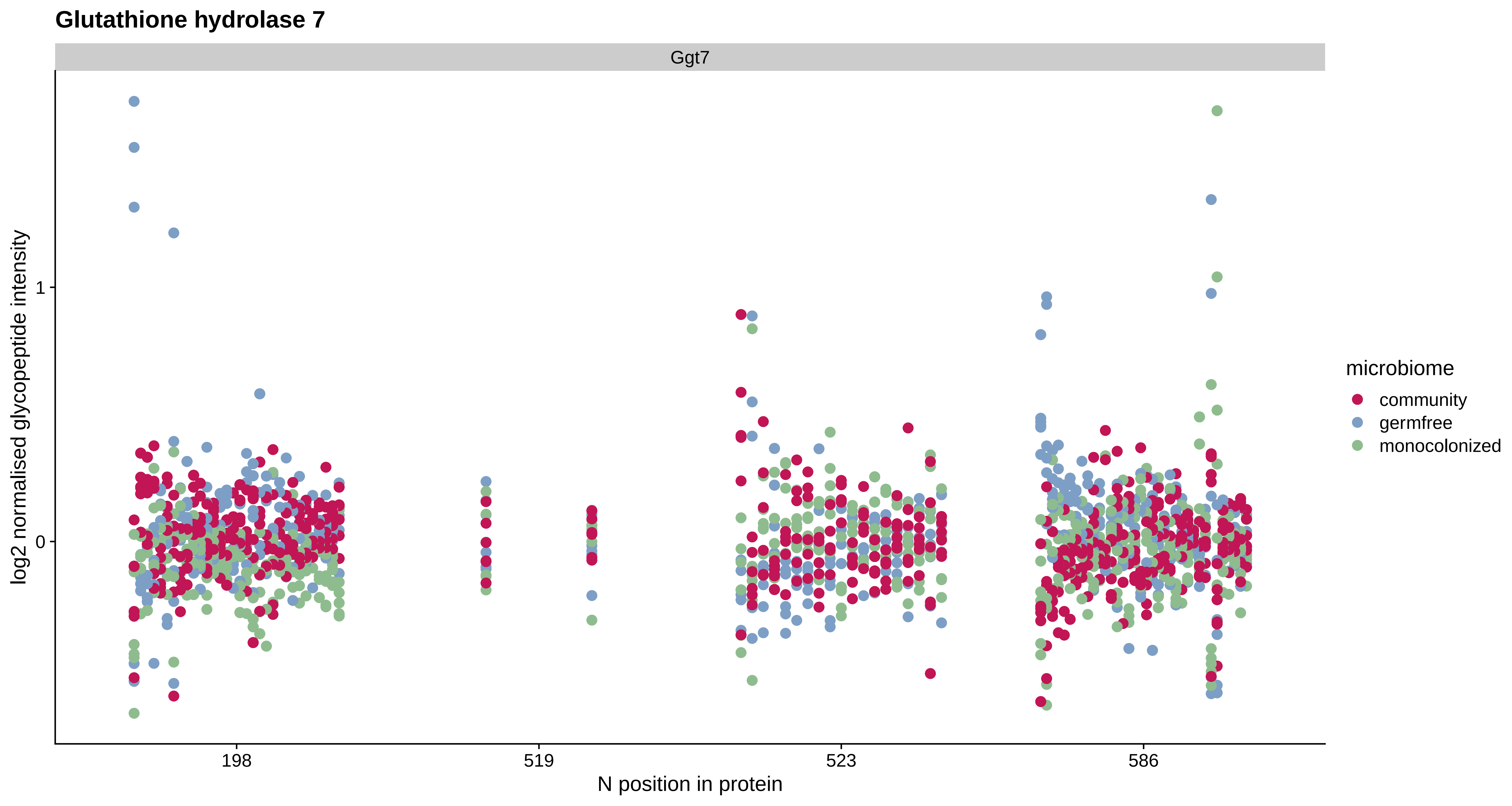
Glutathione hydrolase 7
Ggt7
log2 normalised glycopeptide intensity
1
0
198
519
523
586
N position in protein
microbiome
community
germfree
monocolonized

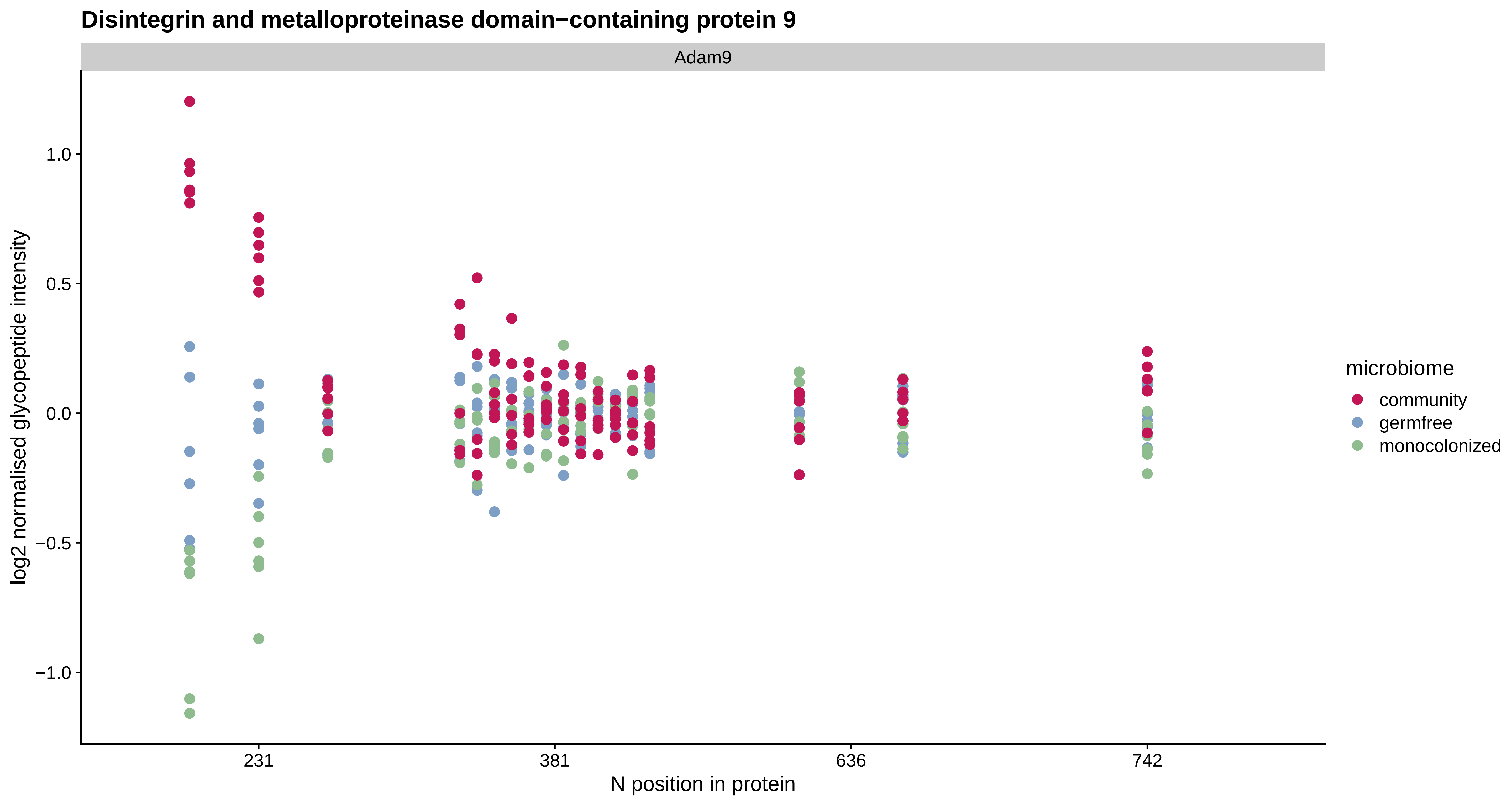

Disintegrin and metalloproteinase domain−containing protein 9
Adam9
log2 normalised glycopeptide intensity
1.0
0.5
0.0
−0.5
−1.0
231
381
636
742
N position in protein
microbiome
community
germfree
monocolonized

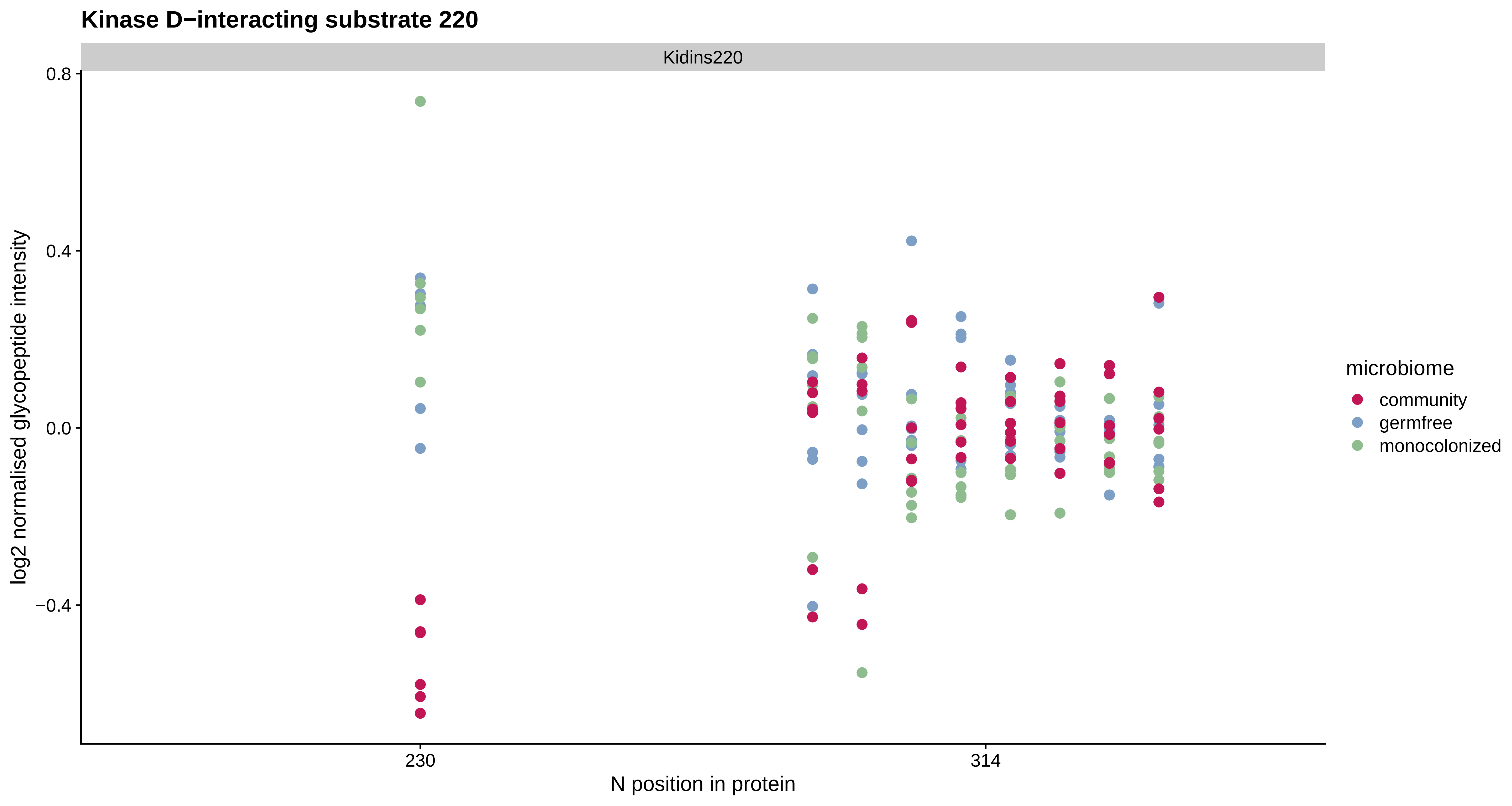
Kinase D−interacting substrate 220
Kidins220
0.8
0.4
0.0
−0.4
log2 normalised glycopeptide intensity
230
314
N position in protein
microbiome
community
germfree
monocolonized

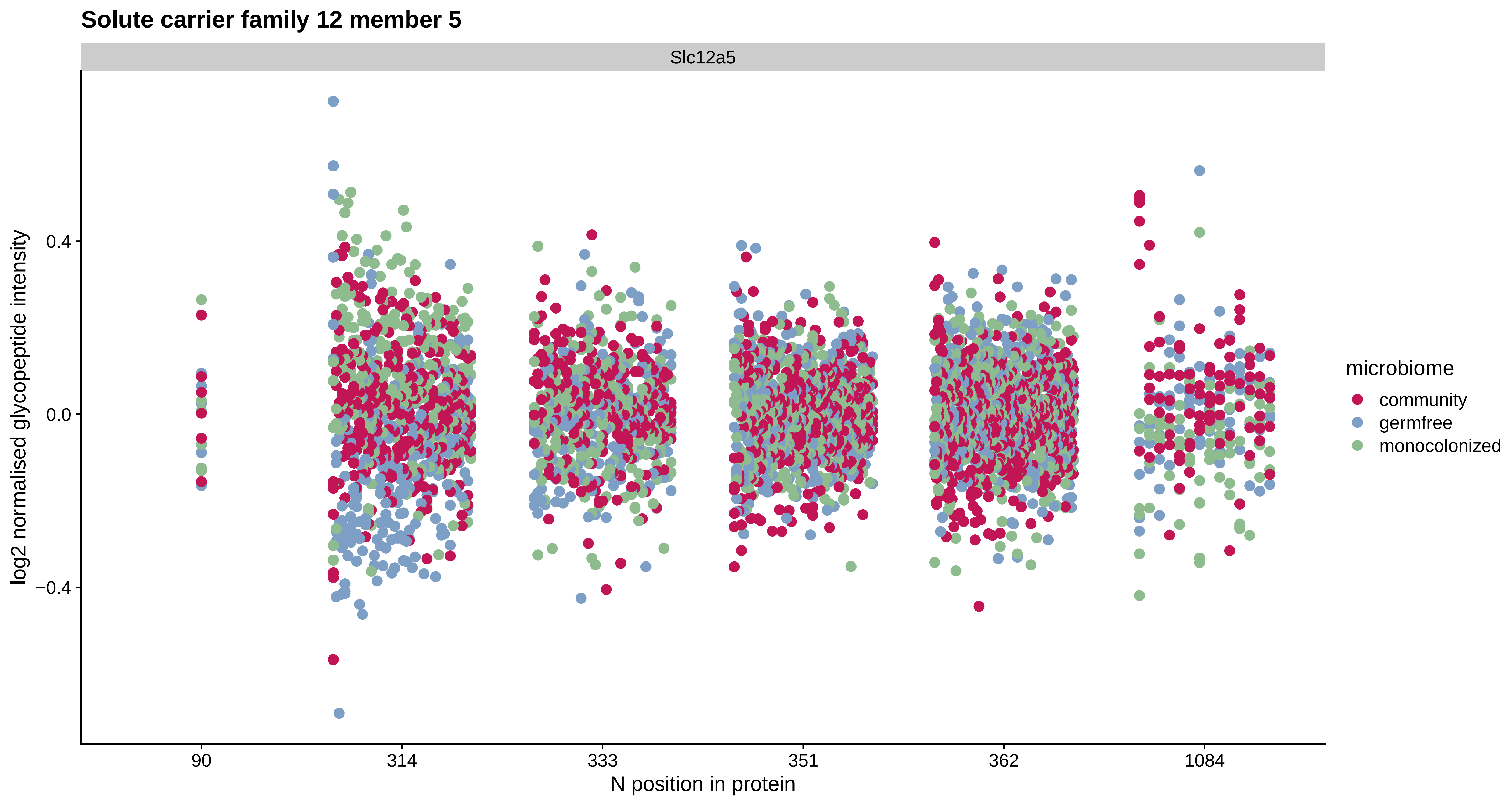
Solute carrier family 12 member 5
Slc12a5
log2 normalised glycopeptide intensity
0.4
0.0
−0.4
90
314
333
351
362
1084
N position in protein
microbiome
community
germfree
monocolonized

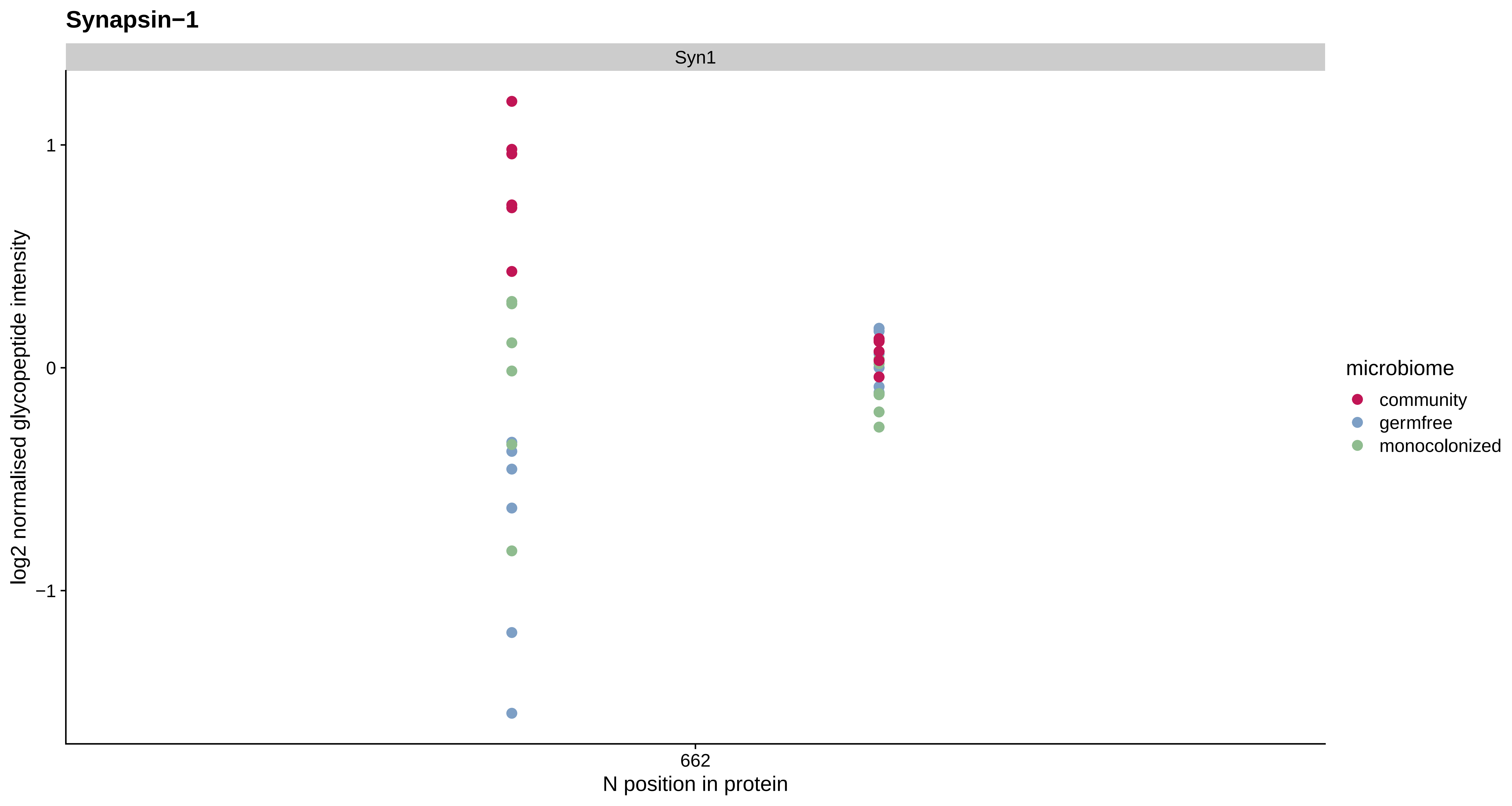

Synapsin−1
Syn1
log2 normalised glycopeptide intensity
1
0
−1
662
N position in protein
microbiome
community
germfree
monocolonized

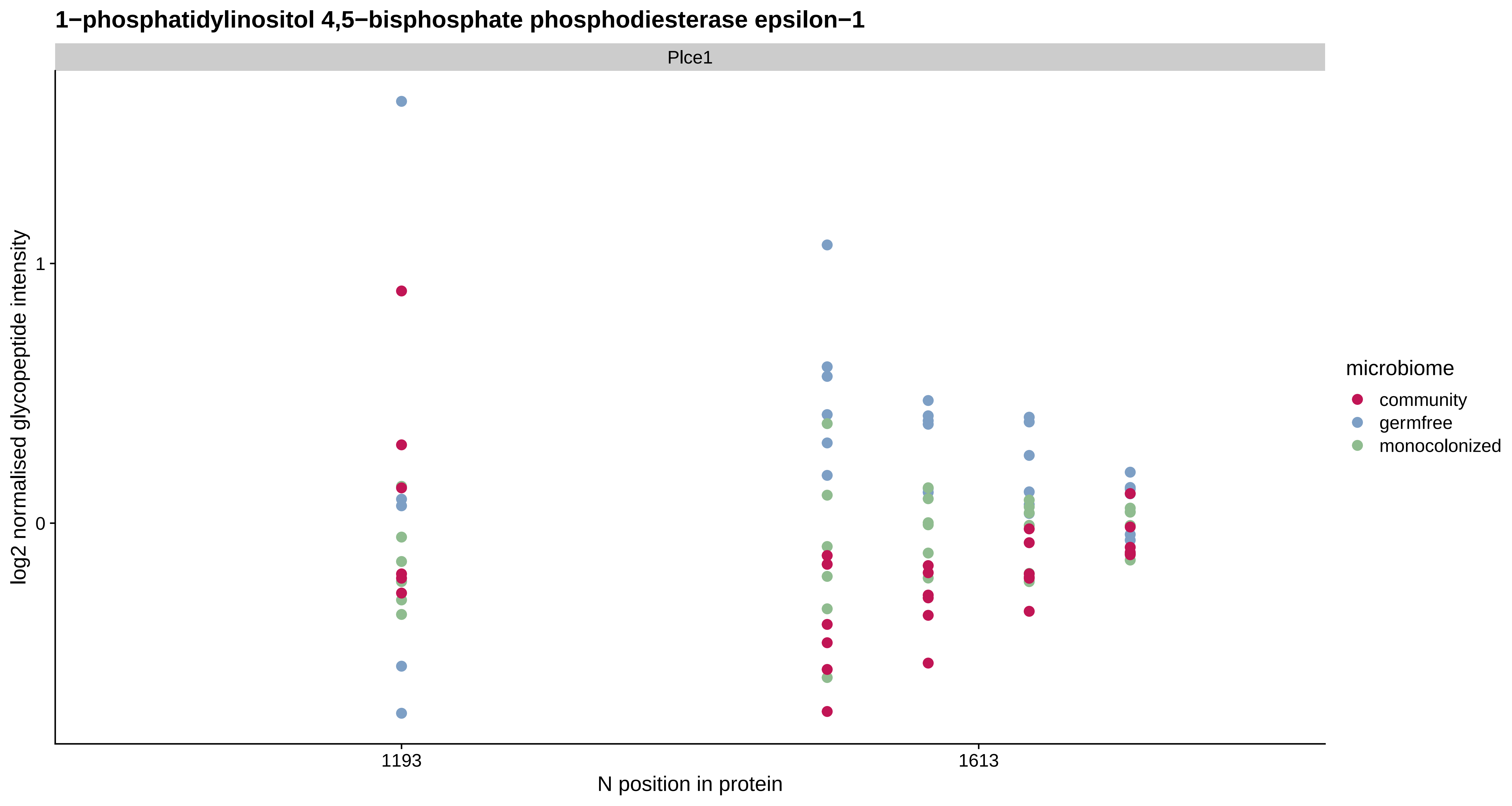

1−phosphatidylinositol 4,5−bisphosphate phosphodiesterase epsilon−1
Plce1
log2 normalised glycopeptide intensity
1
0
1193
1613
N position in protein
microbiome
community
germfree
monocolonized

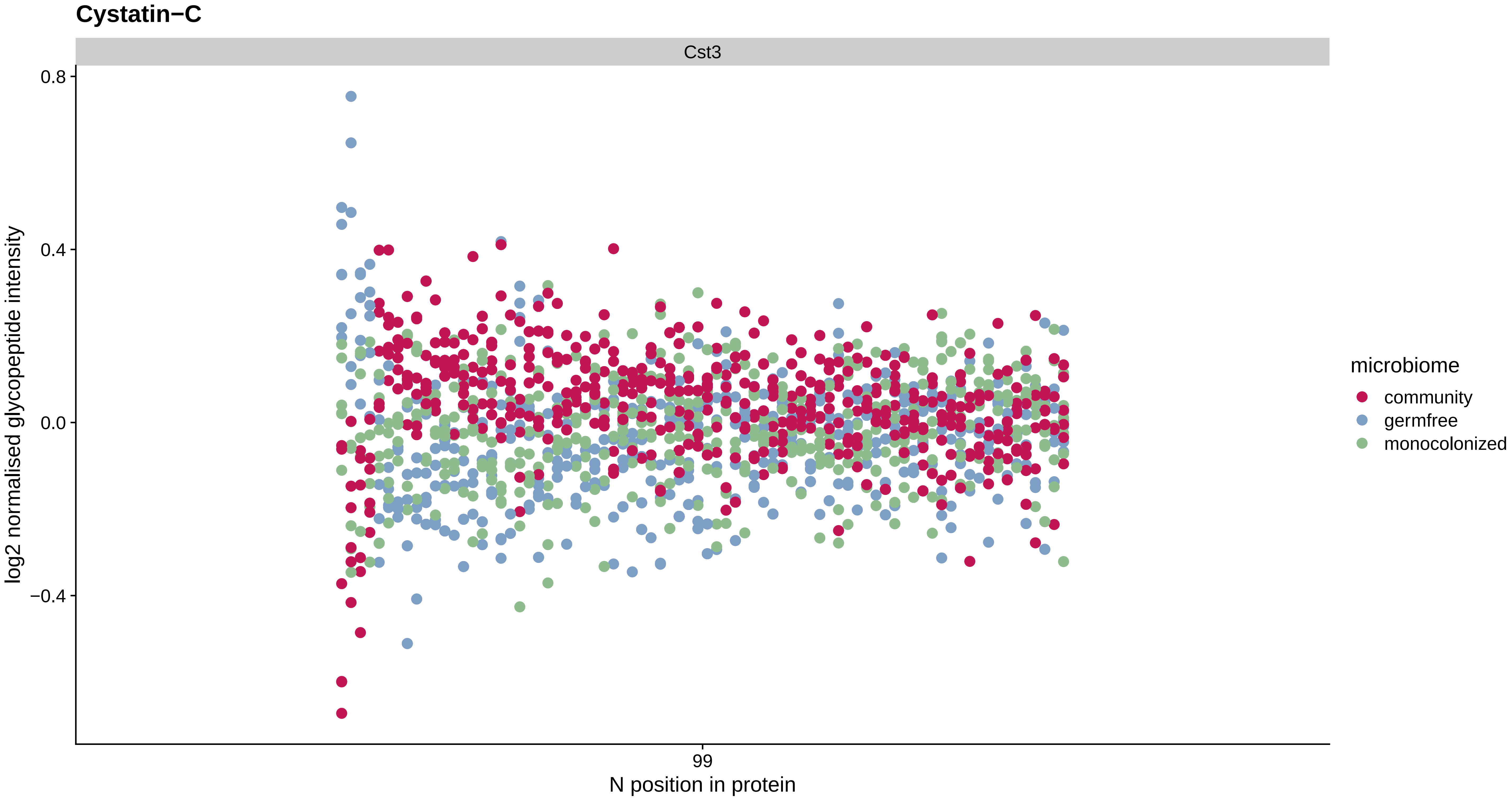
Cystatin−C
Cst3
0.8
0.4
0.0
−0.4
log2 normalised glycopeptide intensity
99
N position in protein
microbiome
community
germfree
monocolonized

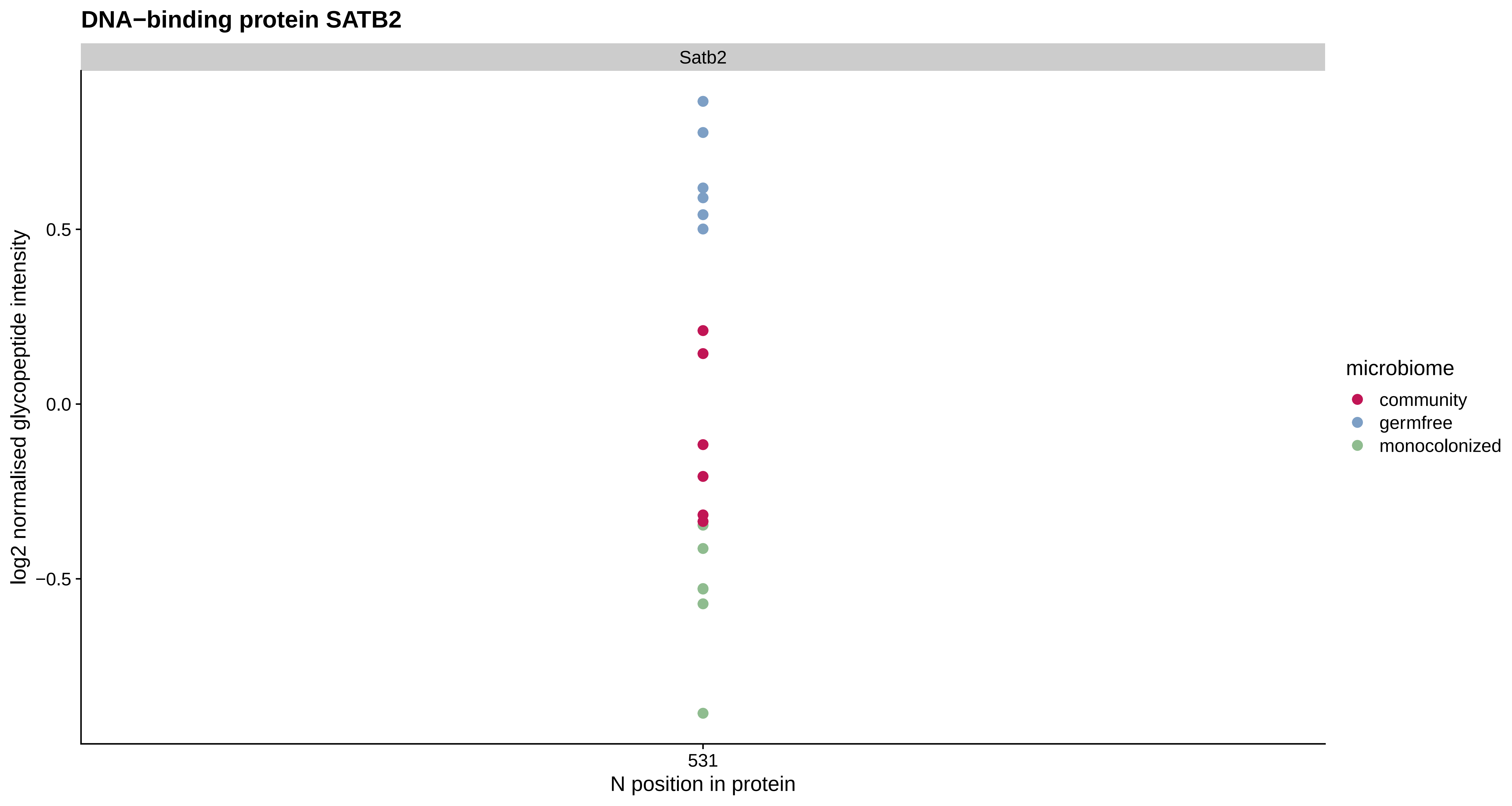

DNA−binding protein SATB2
Satb2
log2 normalised glycopeptide intensity
0.5
0.0
−0.5
531
N position in protein
microbiome
community
germfree
monocolonized

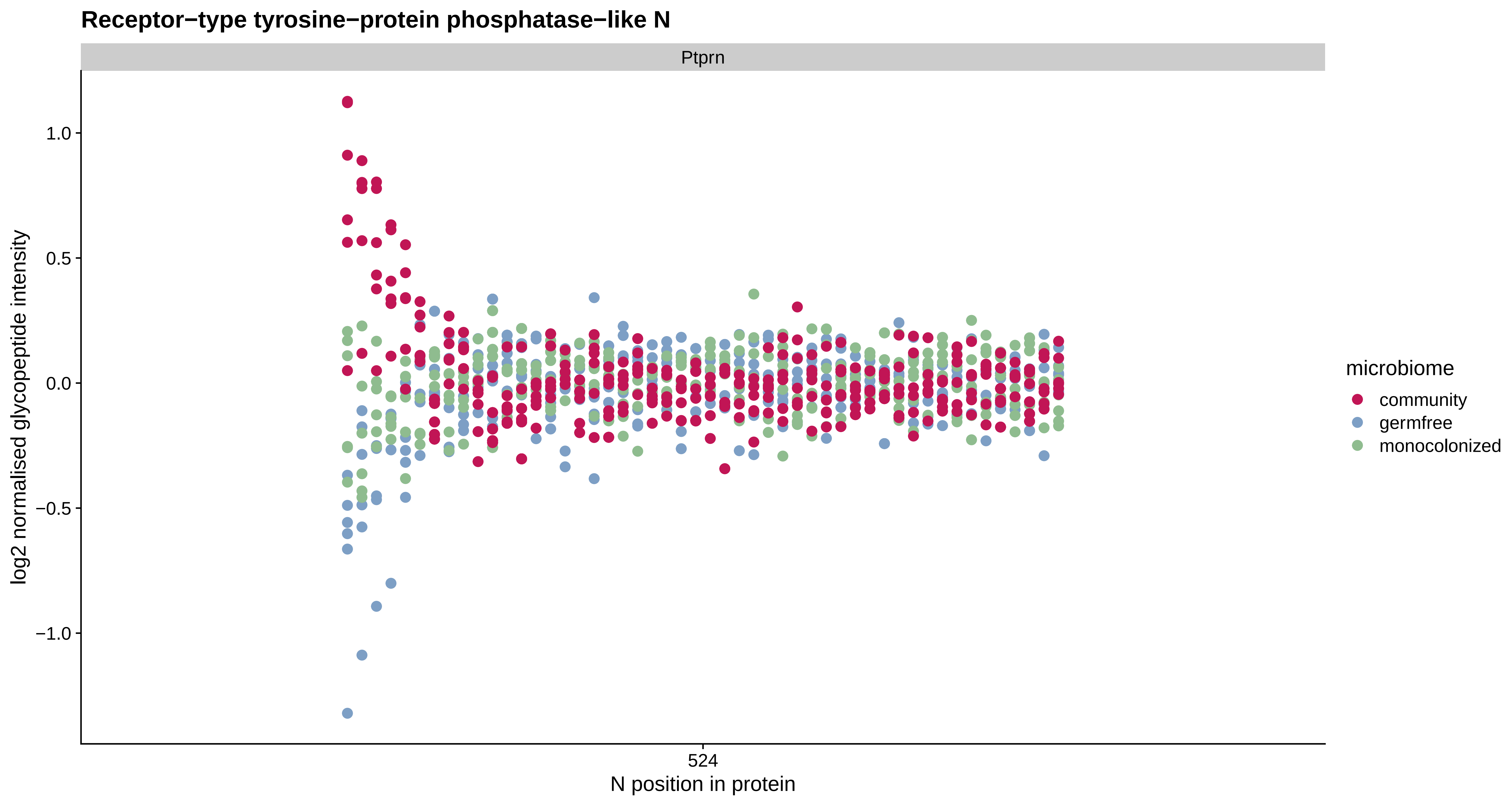

Receptor−type tyrosine−protein phosphatase−like N
Ptprn
1.0
0.5
0.0
−0.5
−1.0
log2 normalised glycopeptide intensity
524
N position in protein
microbiome
community
germfree
monocolonized

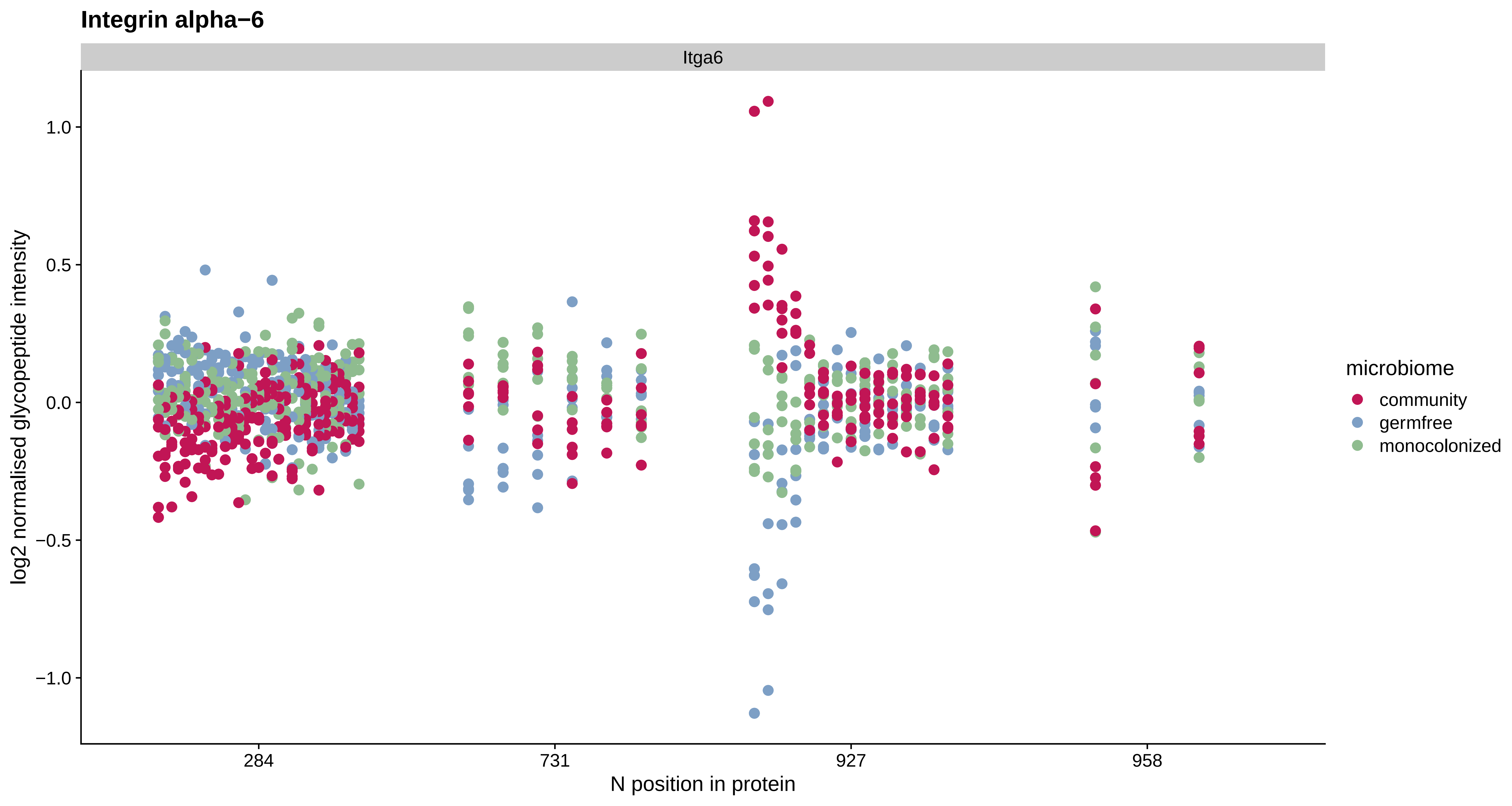

Integrin alpha−6
Itga6
log2 normalised glycopeptide intensity
1.0
0.5
0.0
−0.5
−1.0
284
731
927
958
N position in protein
microbiome
community
germfree
monocolonized

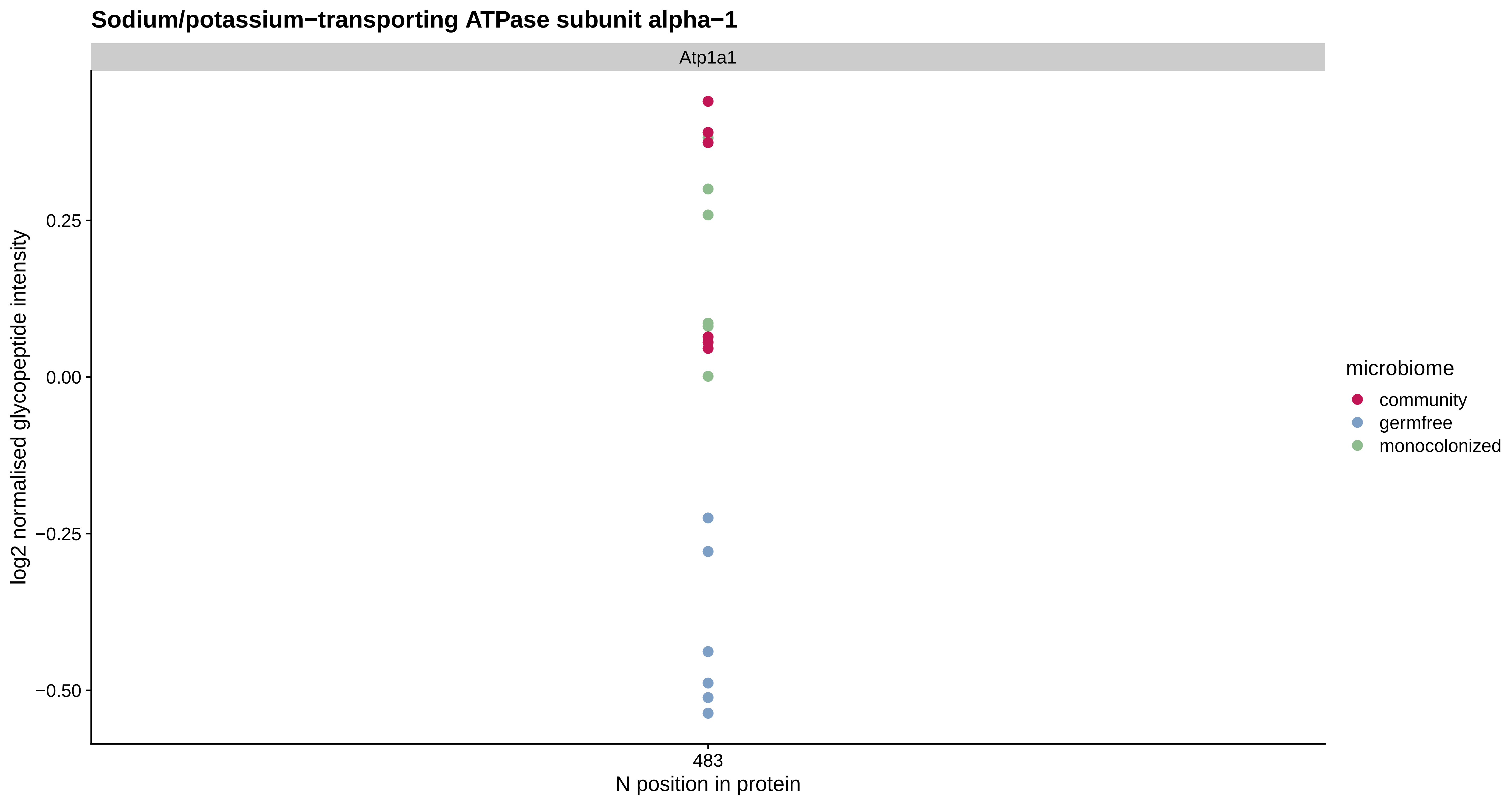
Sodium/potassium−transporting ATPase subunit alpha−1
Atp1a1
log2 normalised glycopeptide intensity
0.25
0.00
−0.25
−0.50
483
N position in protein
microbiome
community
germfree
monocolonized

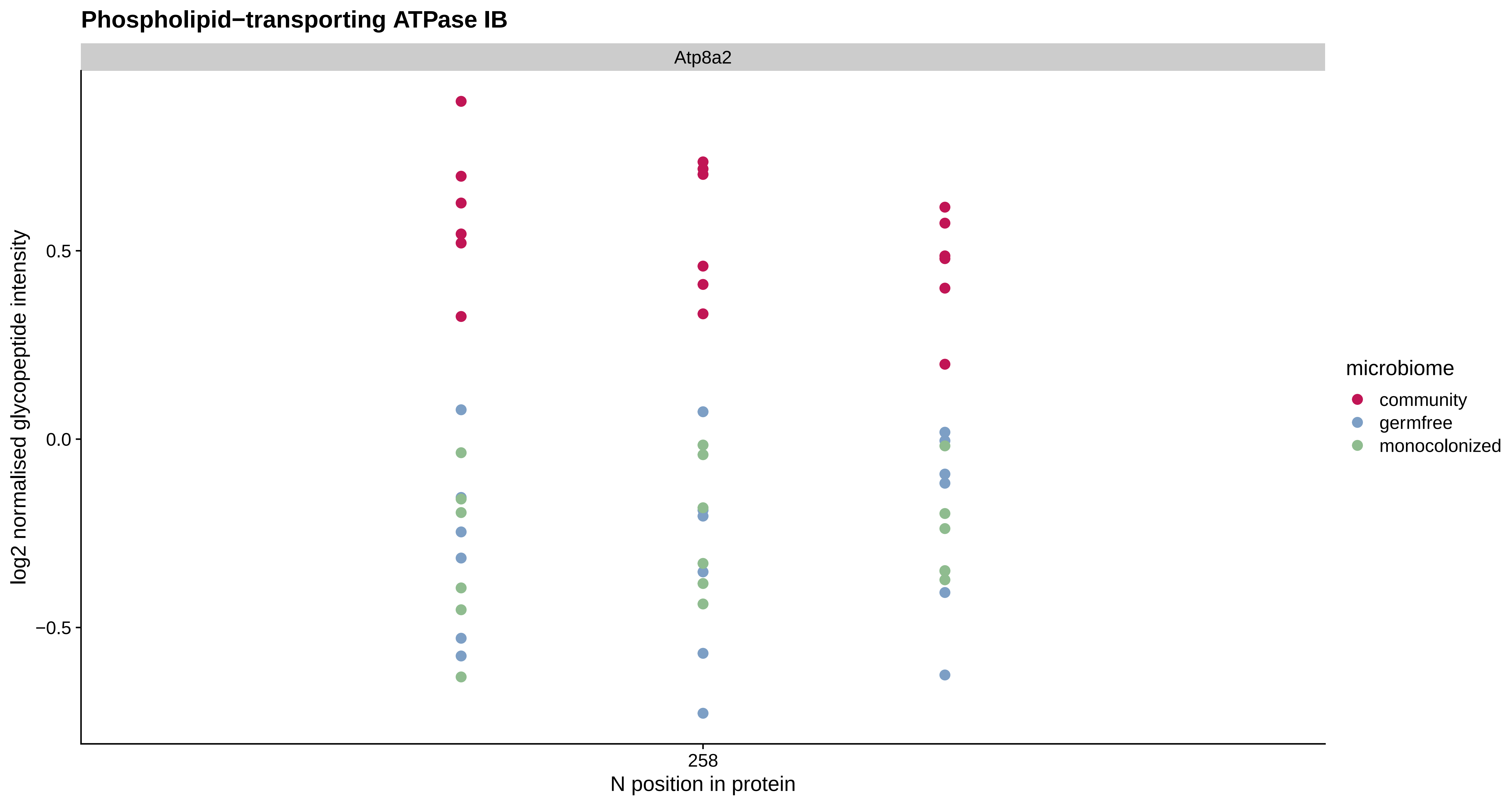

Phospholipid−transporting ATPase IB
Atp8a2
log2 normalised glycopeptide intensity
0.5
0.0
−0.5
258
N position in protein
microbiome
community
germfree
monocolonized

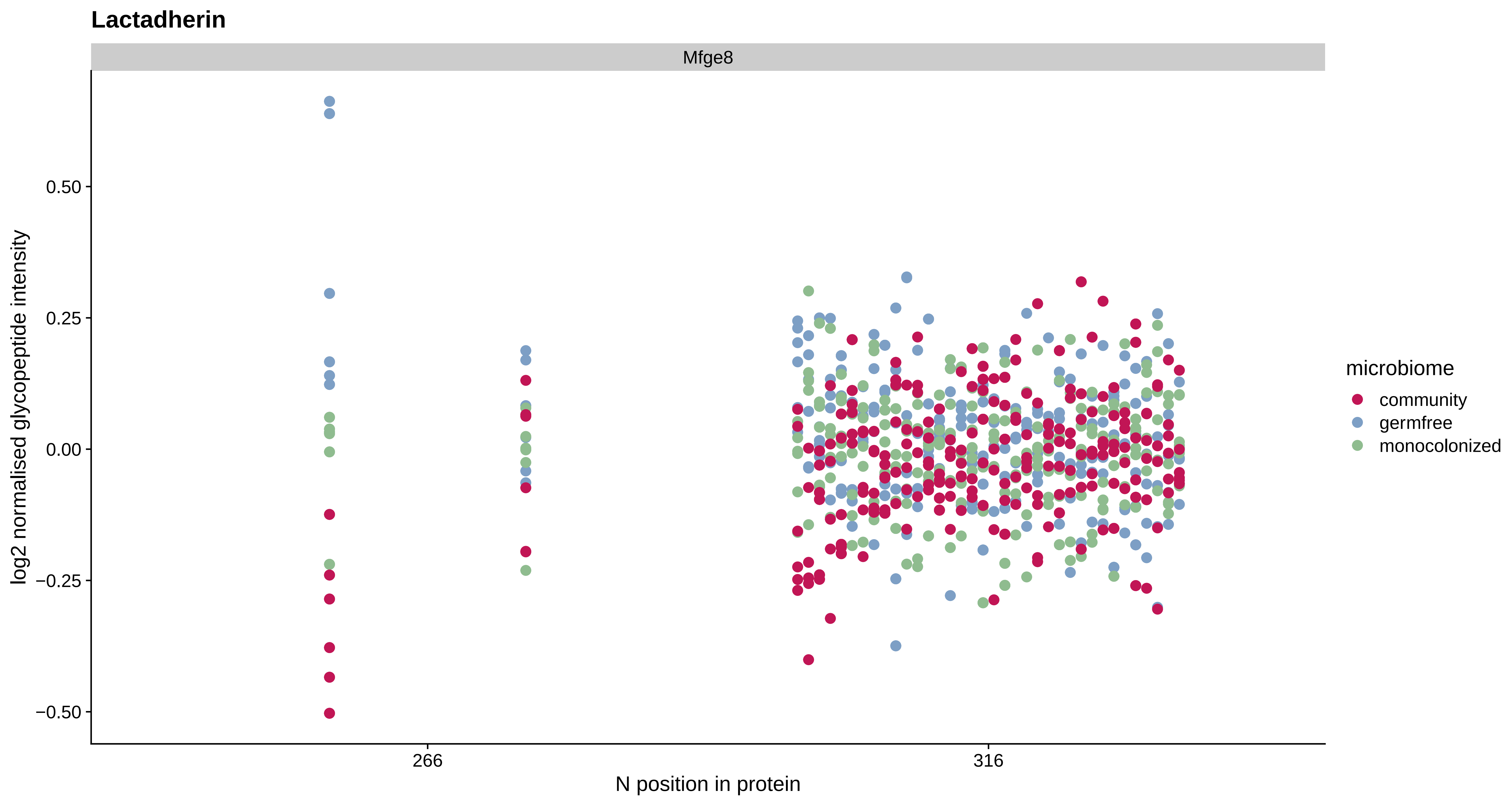
Lactadherin
Mfge8
log2 normalised glycopeptide intensity
0.50
0.25
0.00
−0.25
−0.50
266
316
N position in protein
microbiome
community
germfree
monocolonized

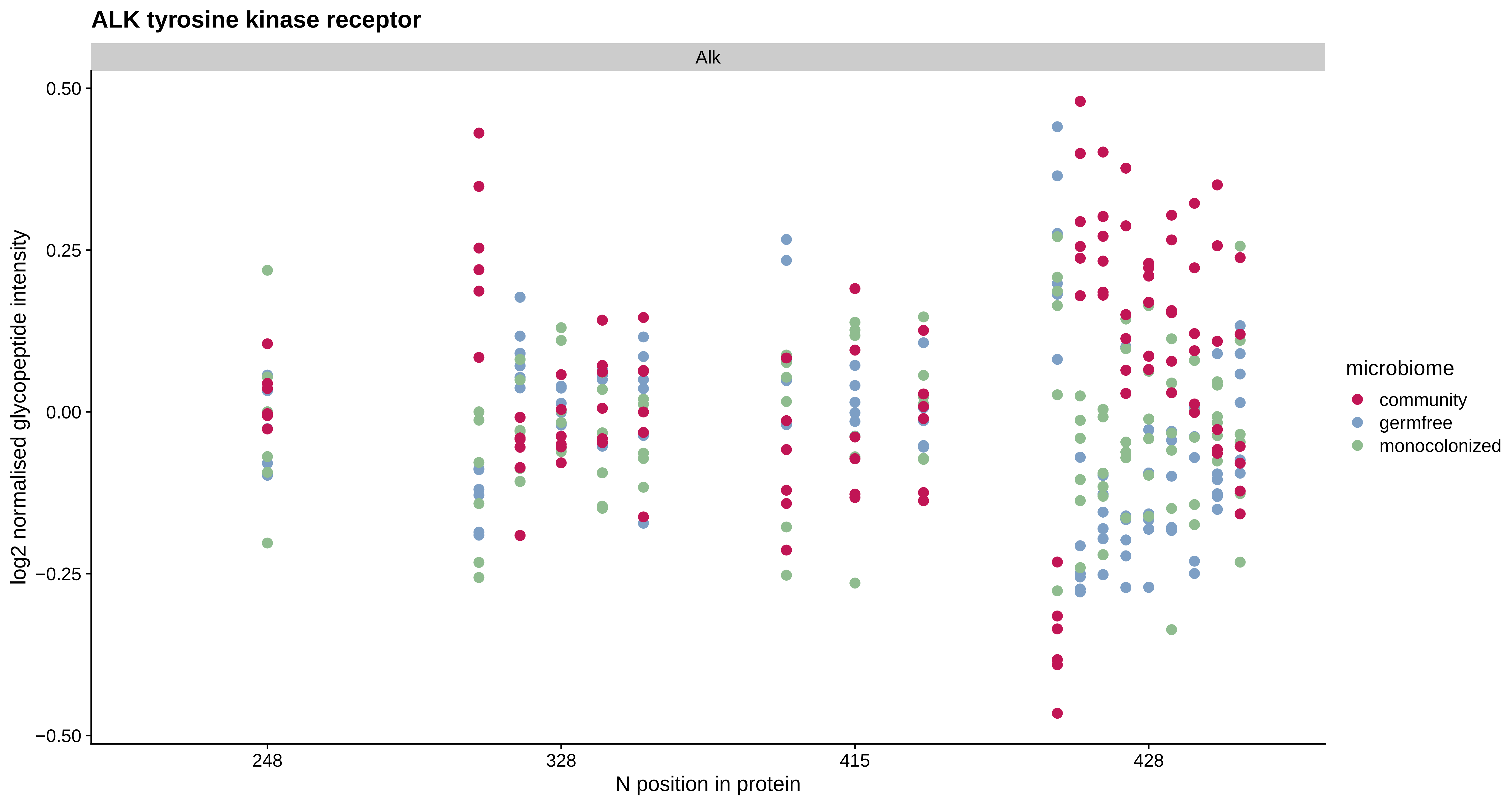

ALK tyrosine kinase receptor
Alk
0.50
0.25
0.00
−0.25
−0.50
log2 normalised glycopeptide intensity
248
328
415
428
N position in protein
microbiome
community
germfree
monocolonized

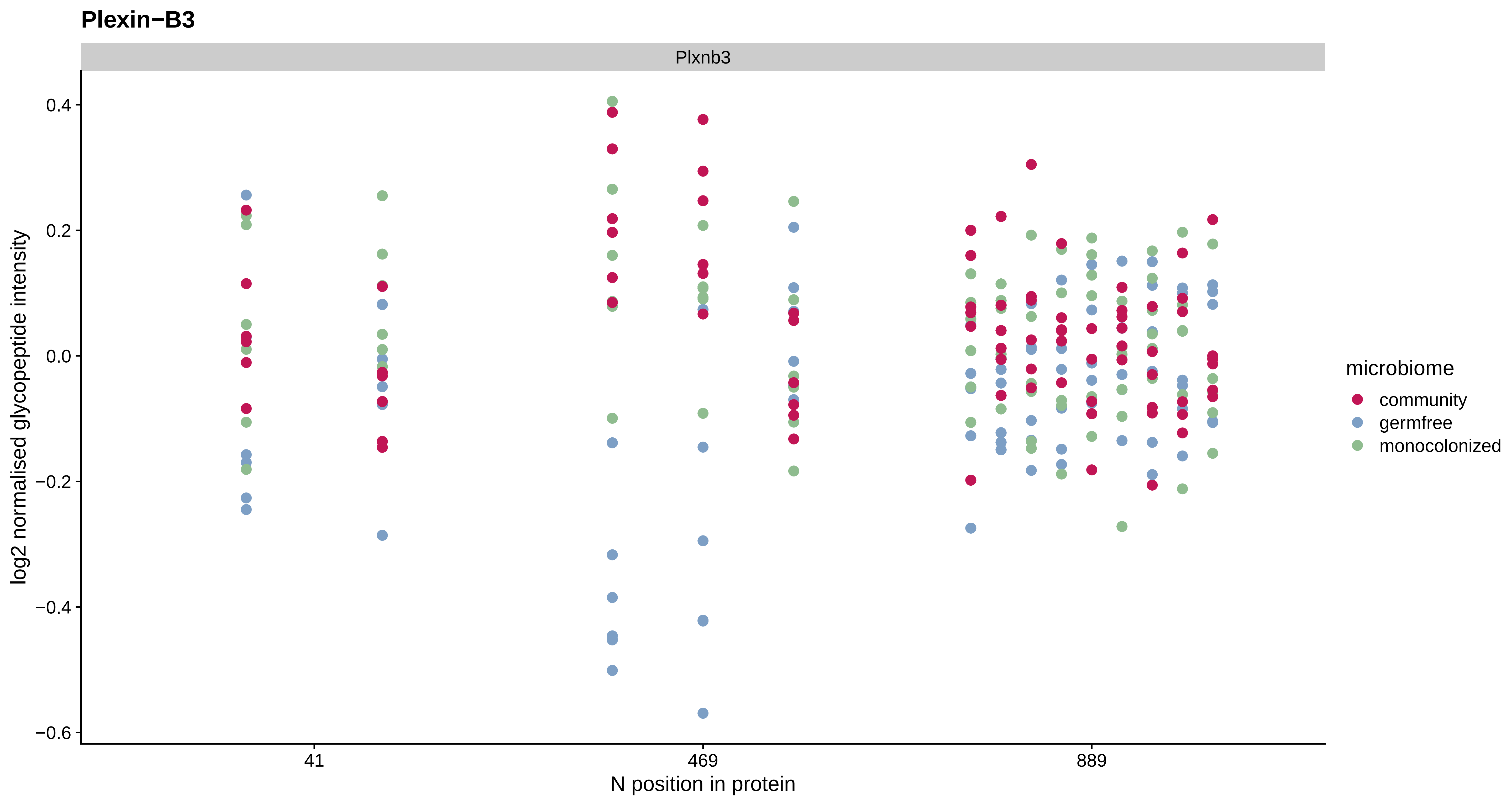

Plexin−B3
Plxnb3
0.4
0.2
0.0
−0.2
−0.4
−0.6
log2 normalised glycopeptide intensity
41
469
889
N position in protein
microbiome
community
germfree
monocolonized

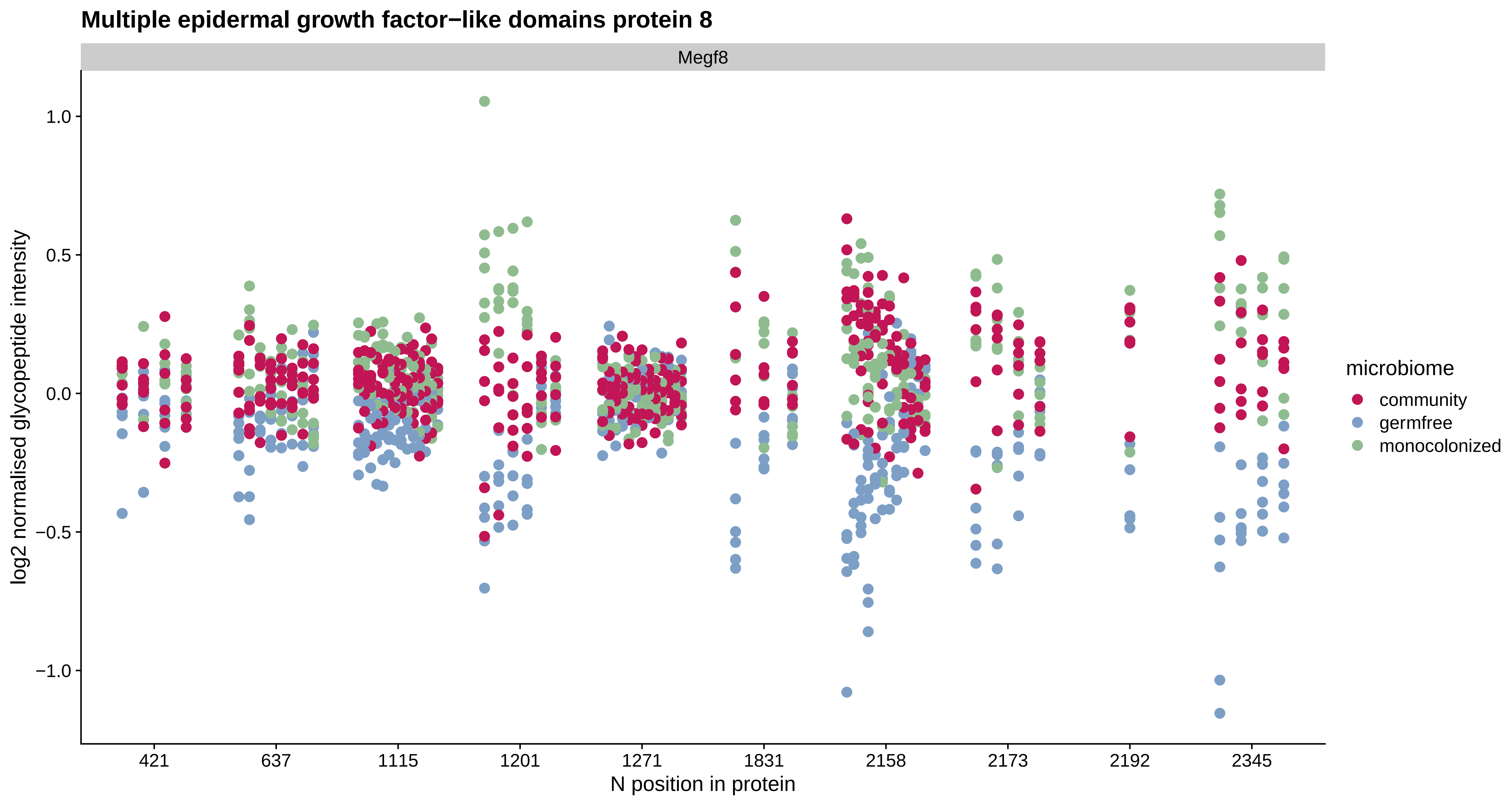

Multiple epidermal growth factor−like domains protein 8
Megf8
log2 normalised glycopeptide intensity
1.0
0.5
0.0
−0.5
−1.0
421
637
1115
1201
1271
1831
2158
2173
2192
2345
N position in protein
microbiome
community
germfree
monocolonized

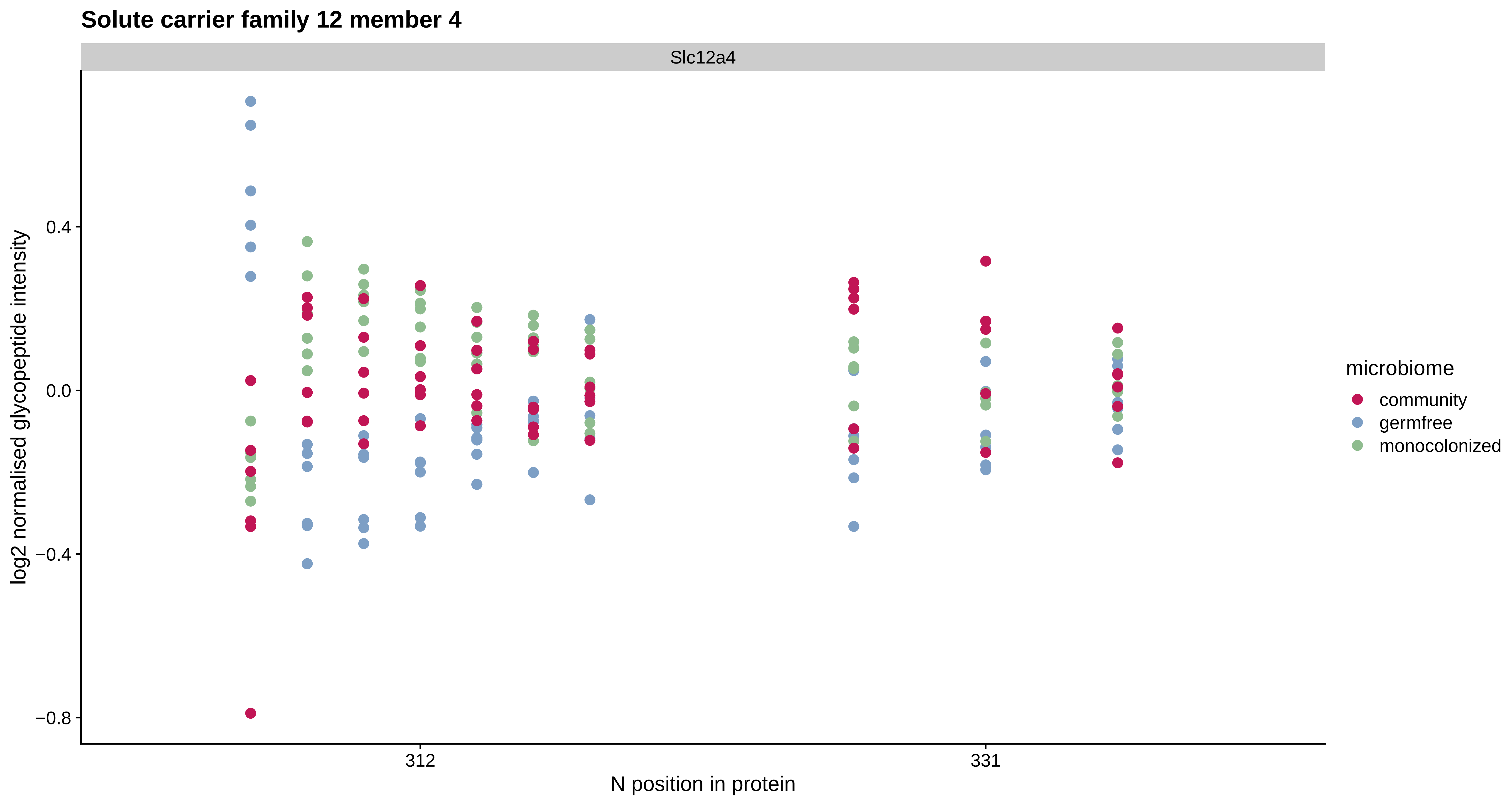
Solute carrier family 12 member 4
Slc12a4
log2 normalised glycopeptide intensity
0.4
0.0
−0.4
−0.8
312
331
N position in protein
microbiome
community
germfree
monocolonized

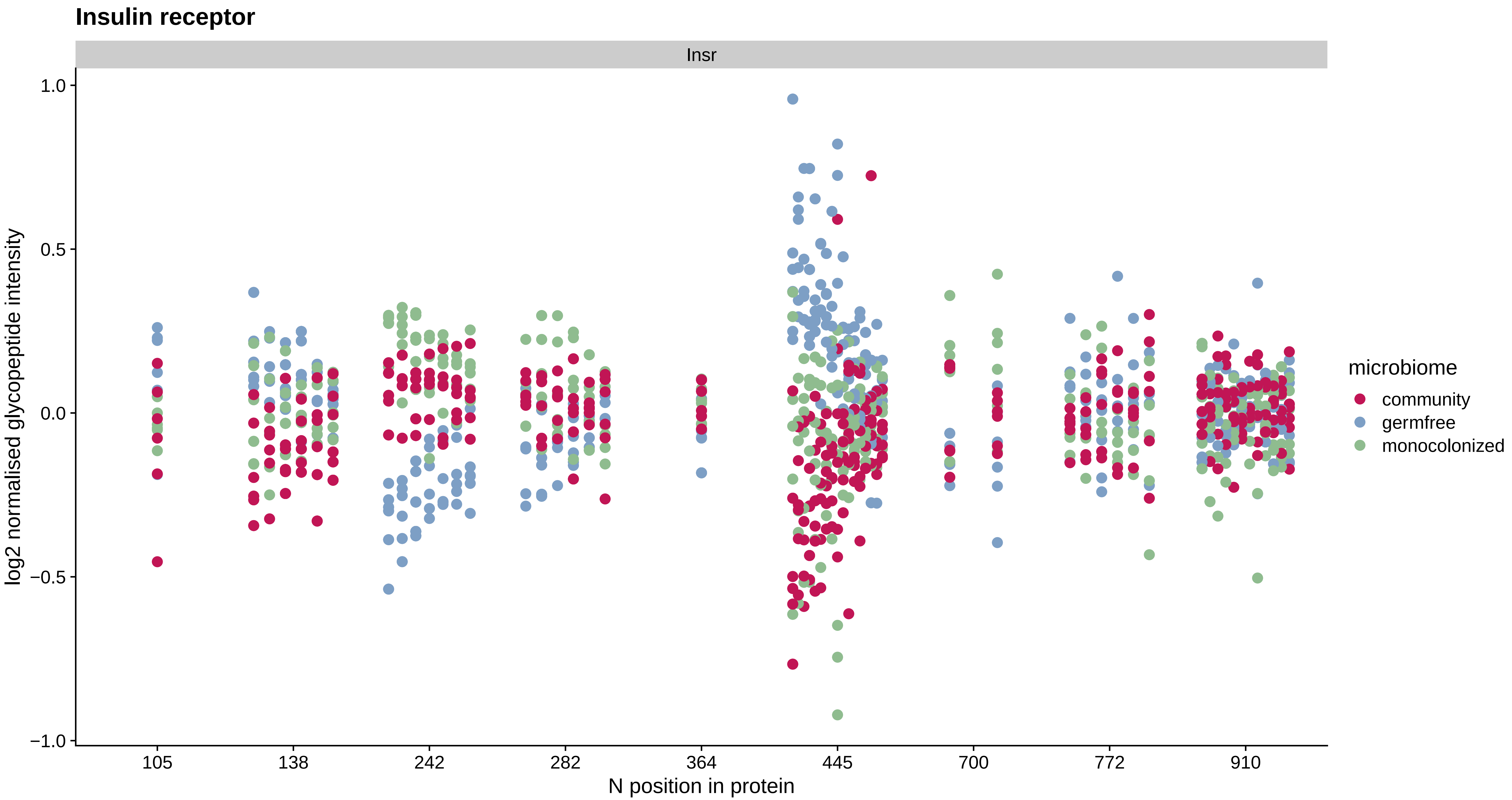
Insulin receptor
Insr
log2 normalised glycopeptide intensity
1.0
0.5
0.0
−0.5
−1.0
105
138
242
282
364
445
700
772
910
N position in protein
microbiome
community
germfree
monocolonized

### Pregnancy zone protein

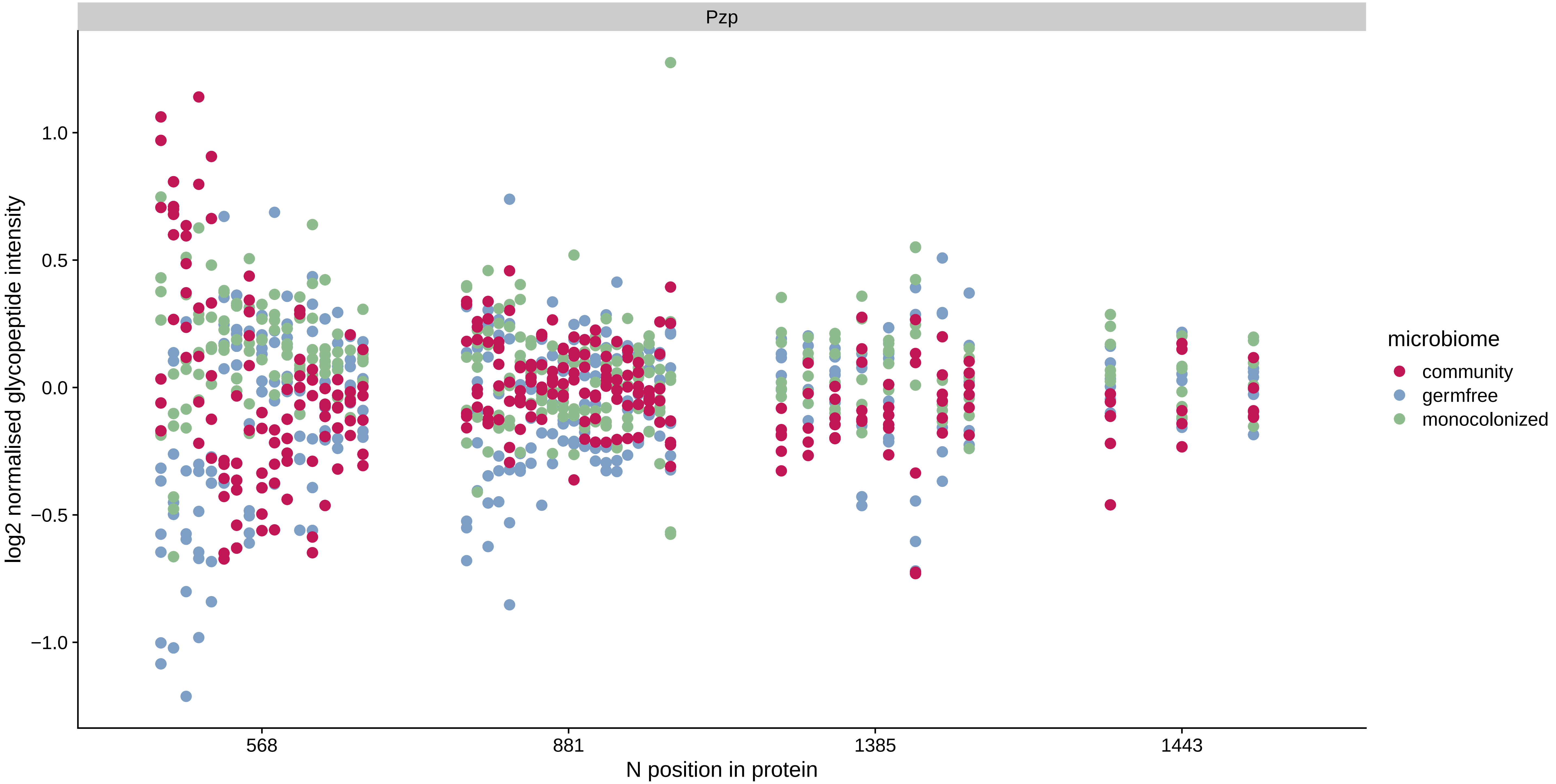

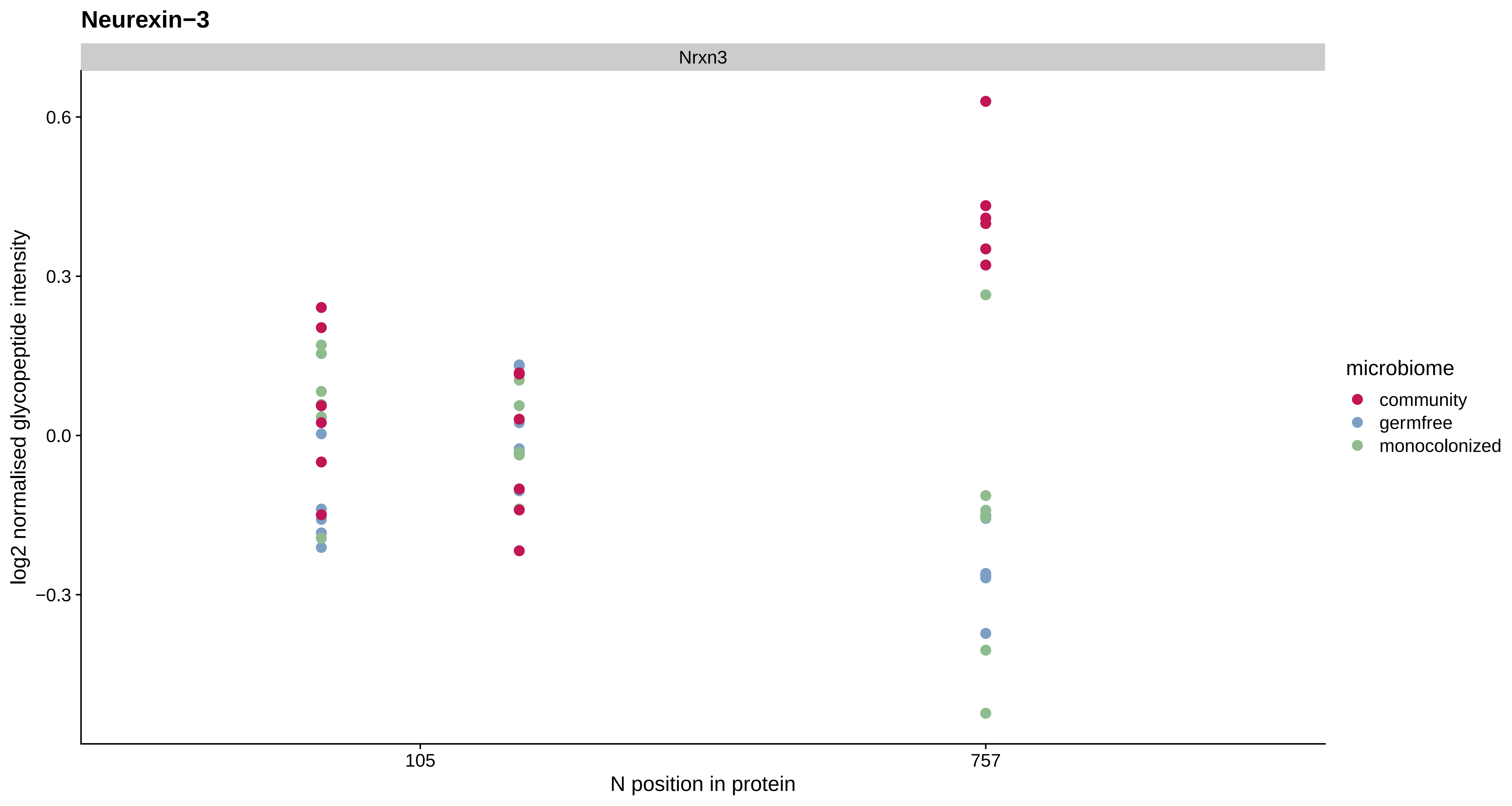
Neurexin−3
Nrxn3
log2 normalised glycopeptide intensity
0.6
0.3
0.0
−0.3
105
757
N position in protein
microbiome
community
germfree
monocolonized

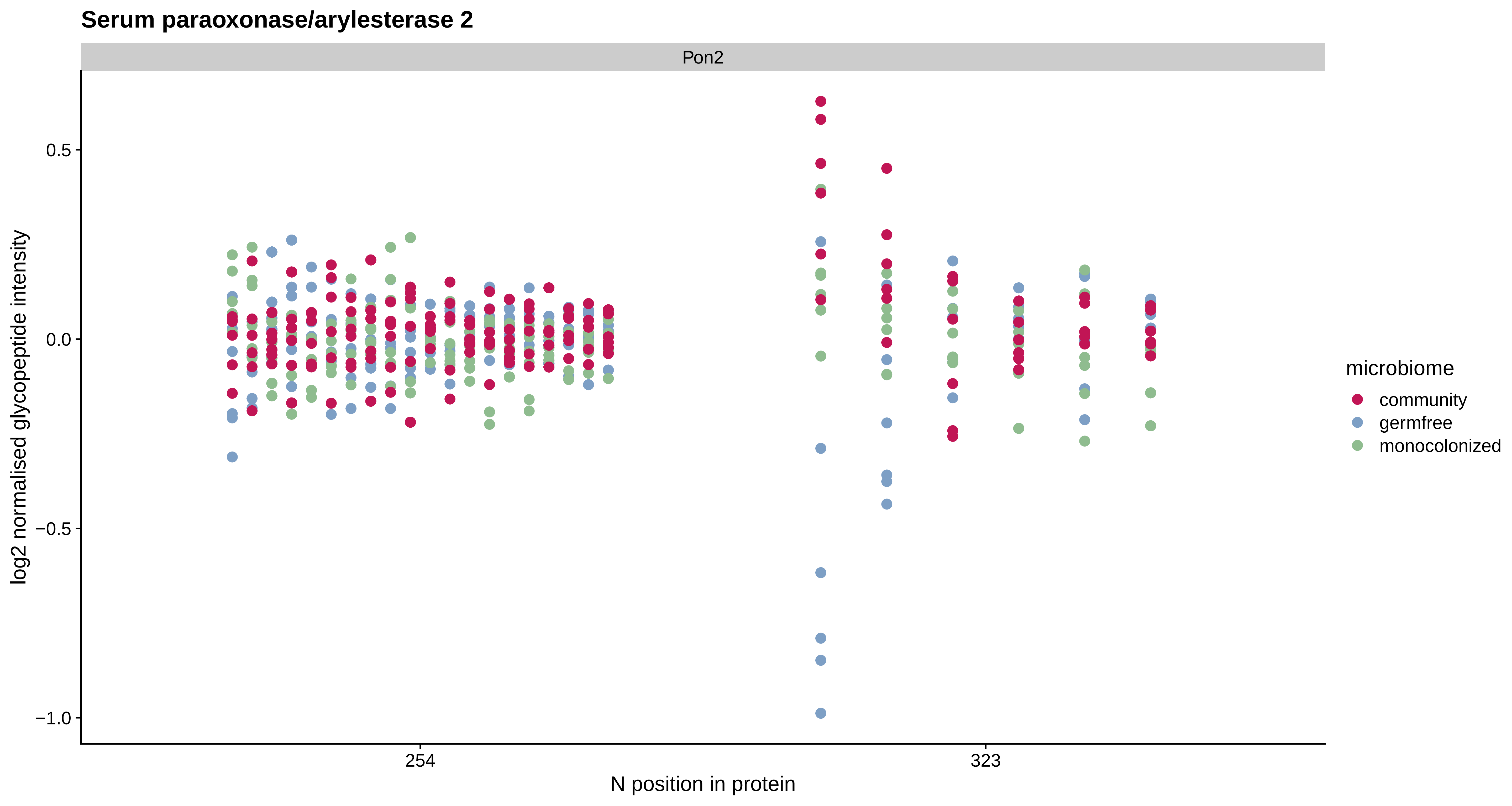
Serum paraoxonase/arylesterase 2
Pon2
log2 normalised glycopeptide intensity
0.5
0.0
−0.5
−1.0
254
323
N position in protein
microbiome
community
germfree
monocolonized

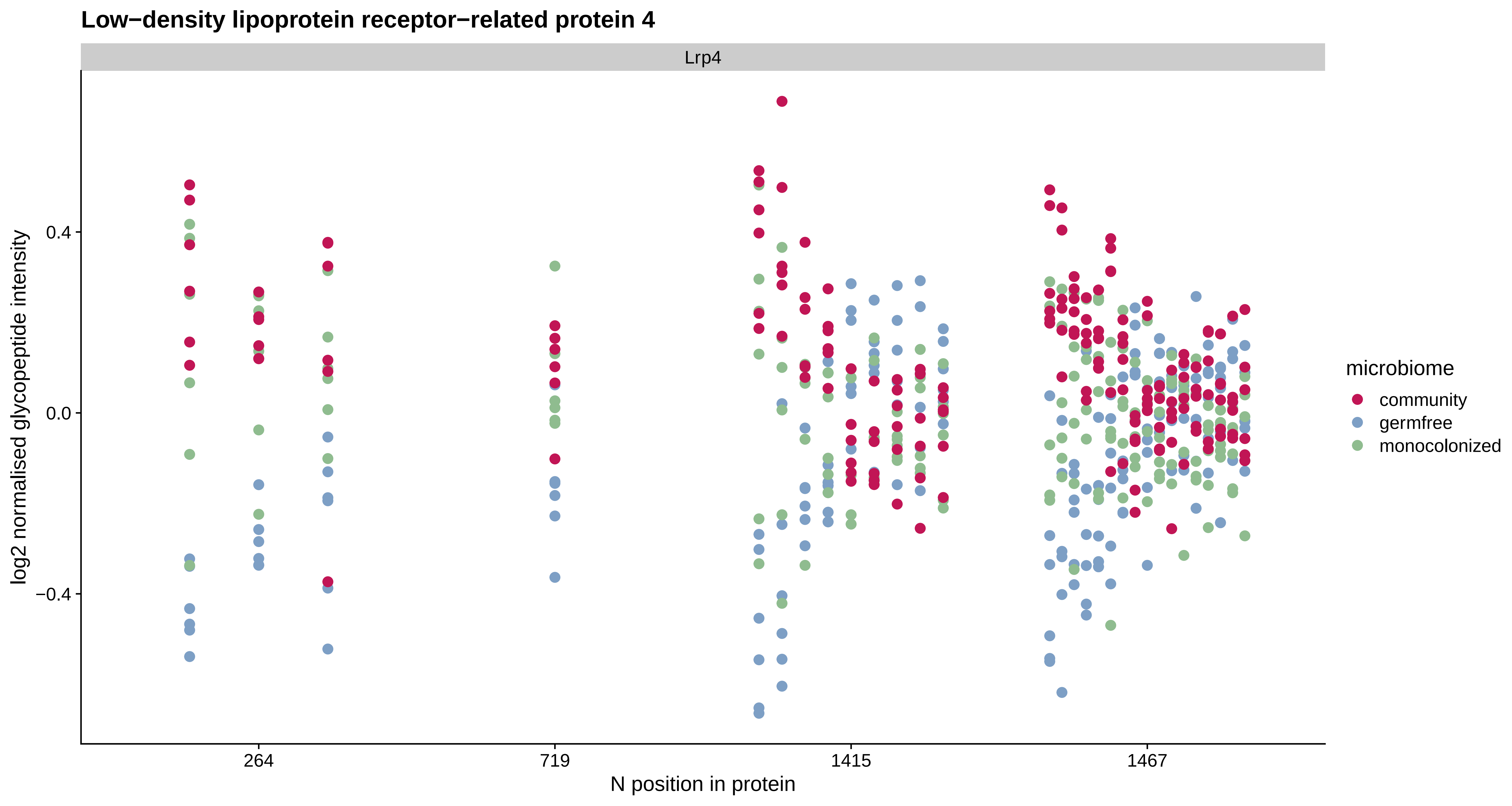

Low−density lipoprotein receptor−related protein 4
Lrp4
log2 normalised glycopeptide intensity
0.4
0.0
−0.4
264
719
1415
1467
N position in protein
microbiome
community
germfree
monocolonized

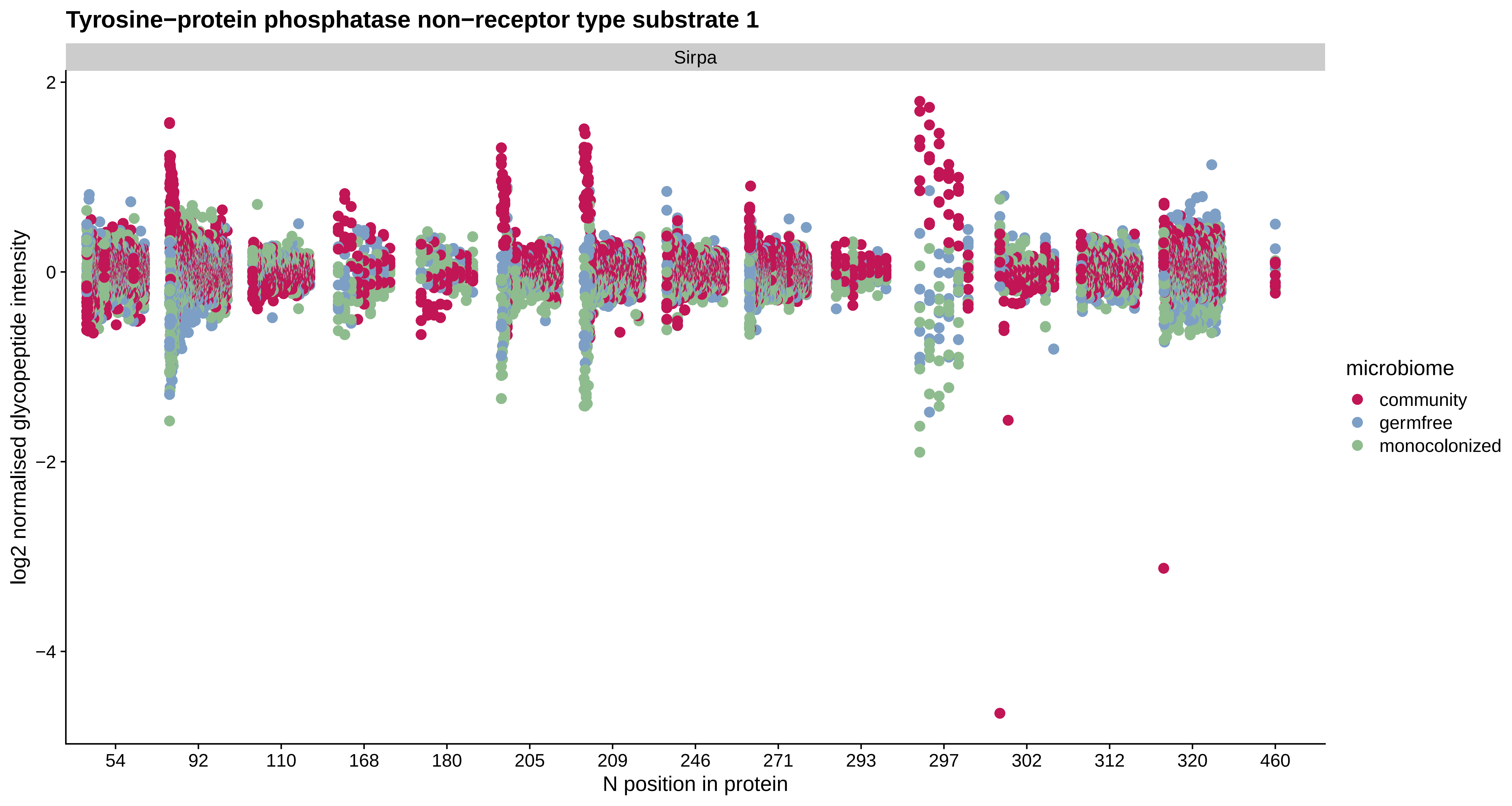
Tyrosine−protein phosphatase non−receptor type substrate 1
Sirpa
log2 normalised glycopeptide intensity
2
0
−2
−4
54
92
110
168
180
205
209
246
271
293
297
302
312
320
460
N position in protein
microbiome
community
germfree
monocolonized

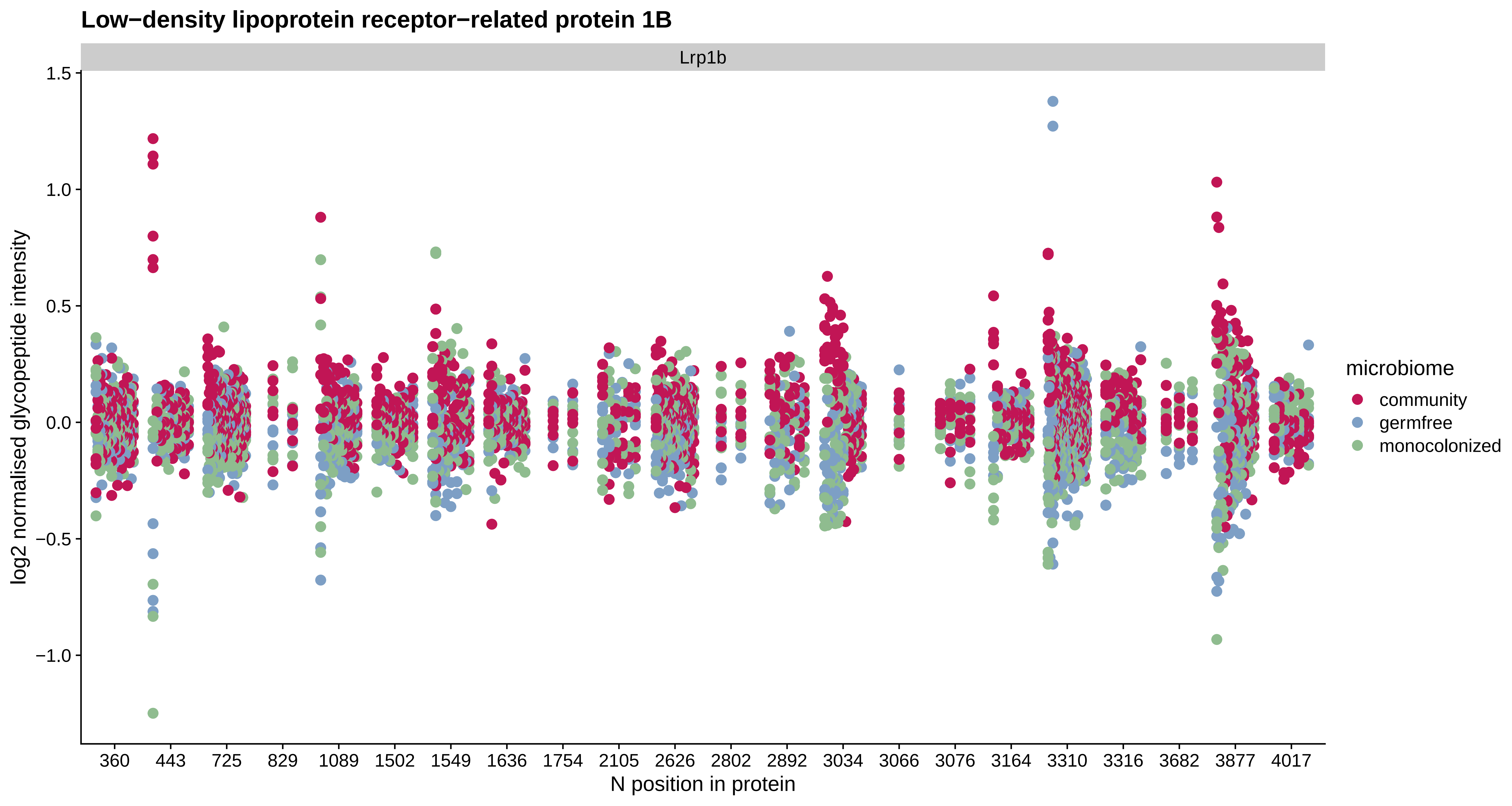

Low−density lipoprotein receptor−related protein 1B
Lrp1b
log2 normalised glycopeptide intensity
1.5
1.0
0.5
0.0
−0.5
−1.0
360
443
725
829
1089
1502
1549
1636
1754
2105
2626
2802
2892
3034
3066
3076
3164
3310
3316
3682
3877
4017
N position in protein
microbiome
community
germfree
monocolonized

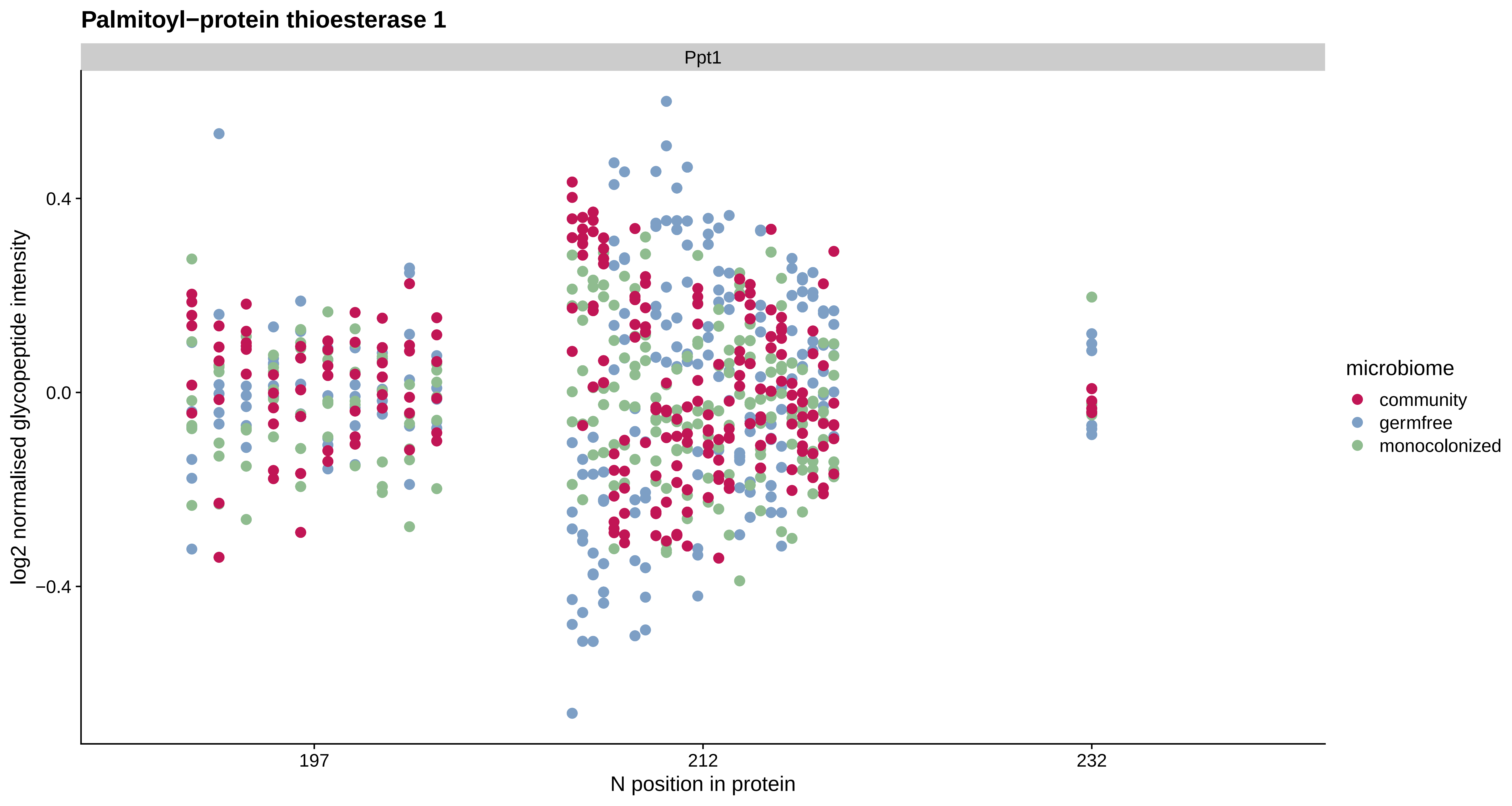

Palmitoyl−protein thioesterase 1
Ppt1
log2 normalised glycopeptide intensity
0.4
0.0
−0.4
197
212
232
N position in protein
microbiome
community
germfree
monocolonized

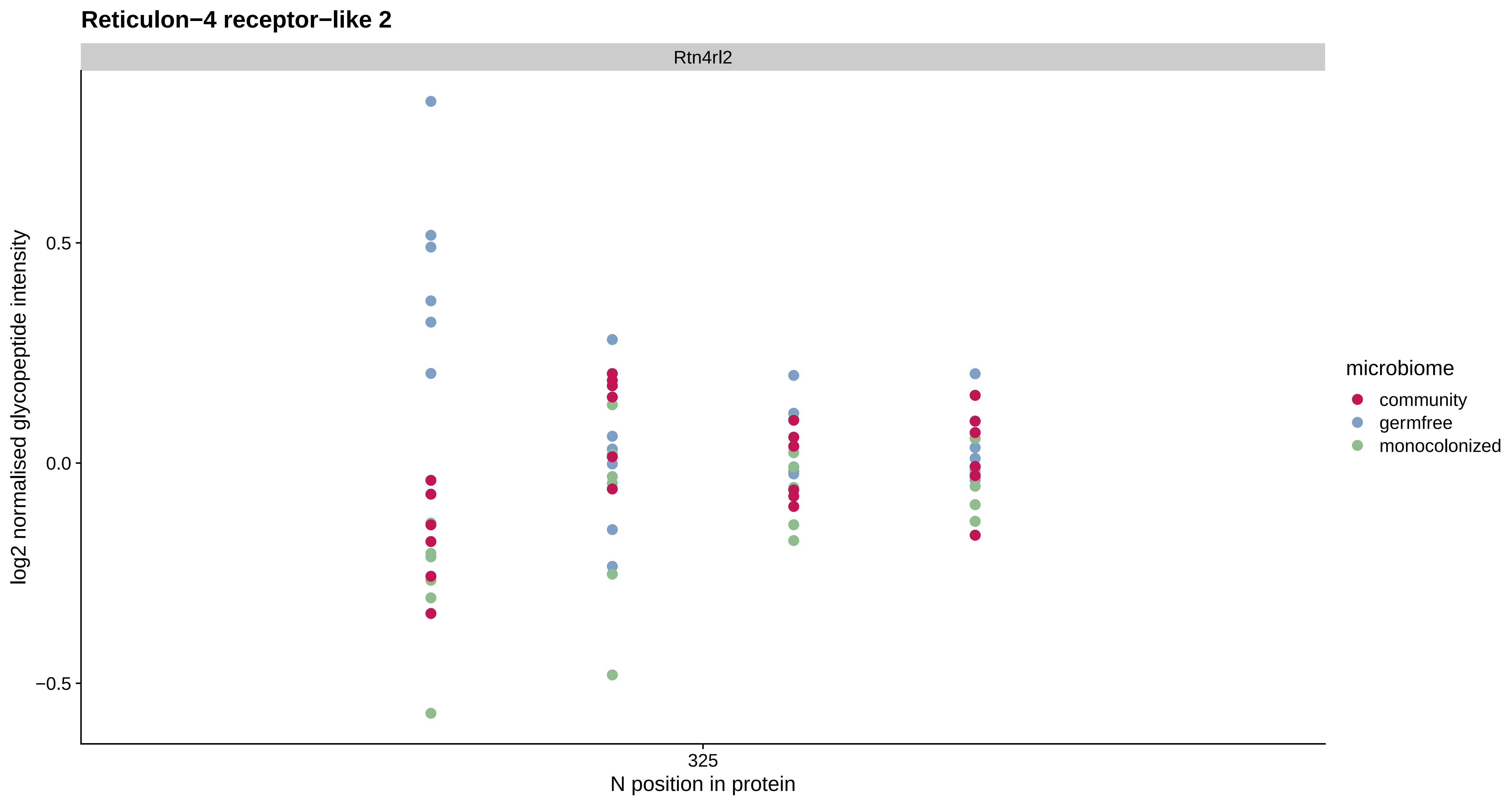
Reticulon−4 receptor−like 2
Rtn4rl2
log2 normalised glycopeptide intensity
0.5
0.0
−0.5
325
N position in protein
microbiome
community
germfree
monocolonized

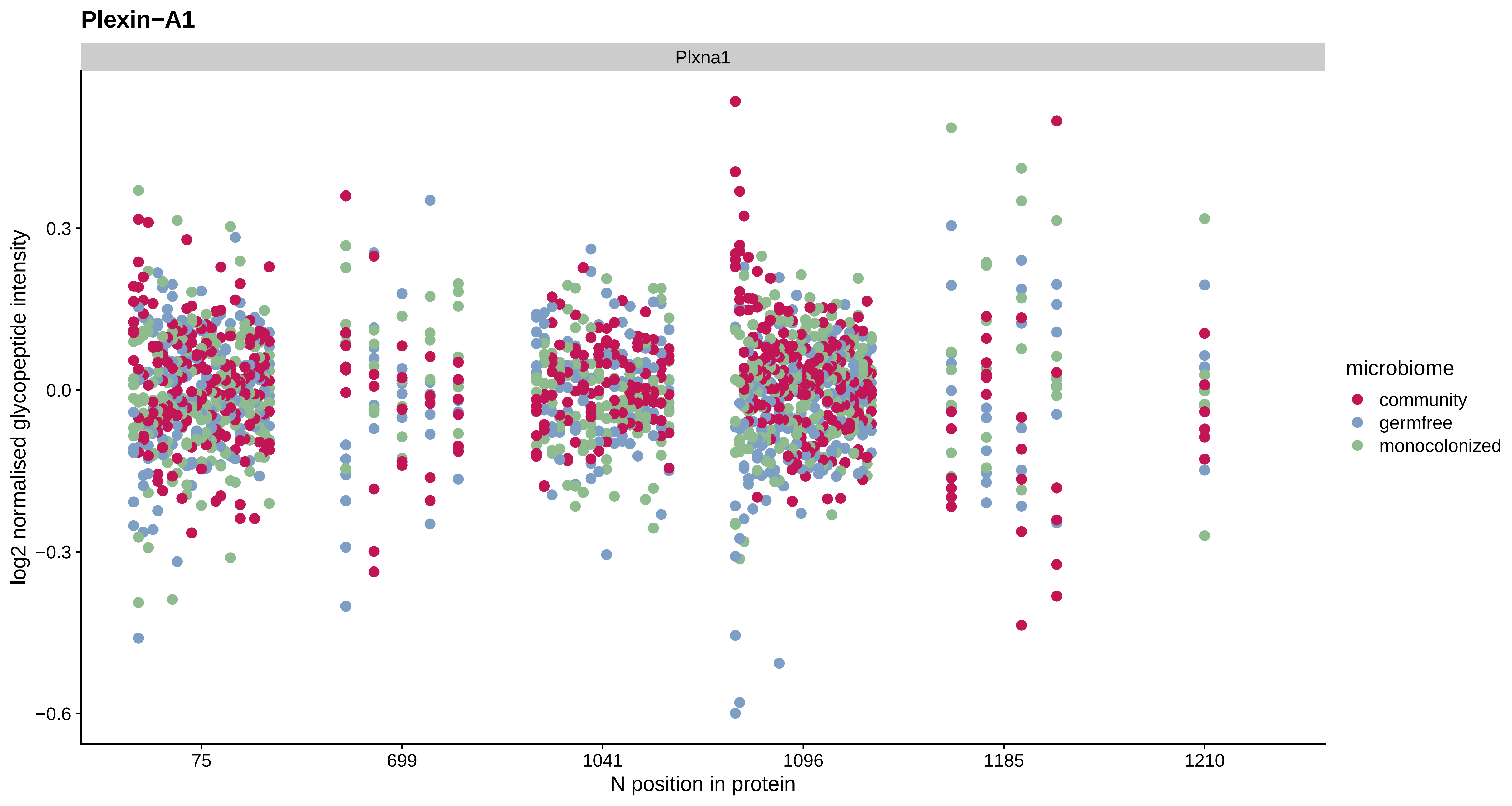
Plexin−A1
Plxna1
log2 normalised glycopeptide intensity
0.3
0.0
−0.3
−0.6
75
699
1041
1096
1185
1210
N position in protein
microbiome
community
germfree
monocolonized

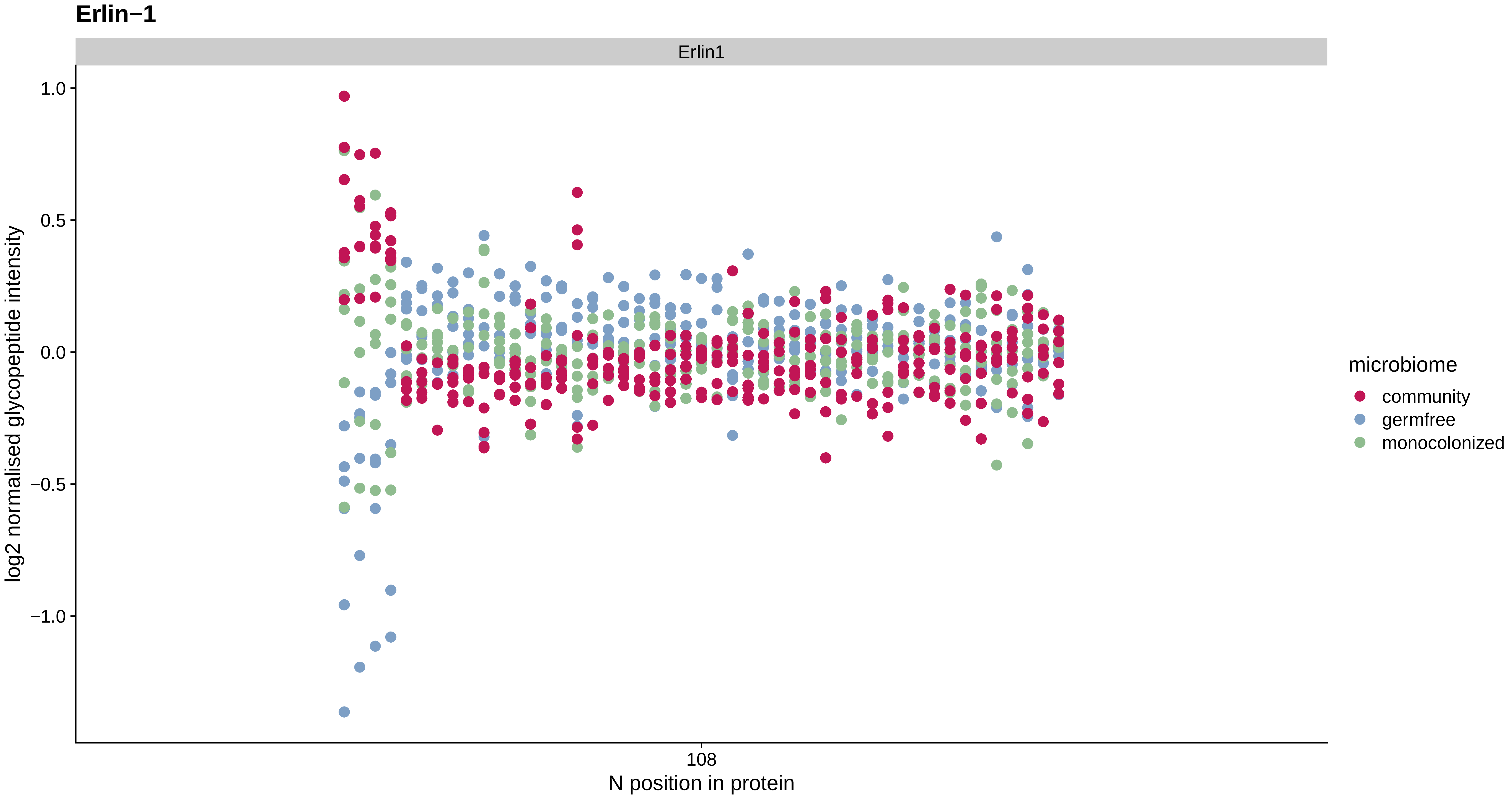
Erlin−1
Erlin1
1.0
0.5
0.0
−0.5
−1.0
log2 normalised glycopeptide intensity
108
N position in protein
microbiome
community
germfree
monocolonized

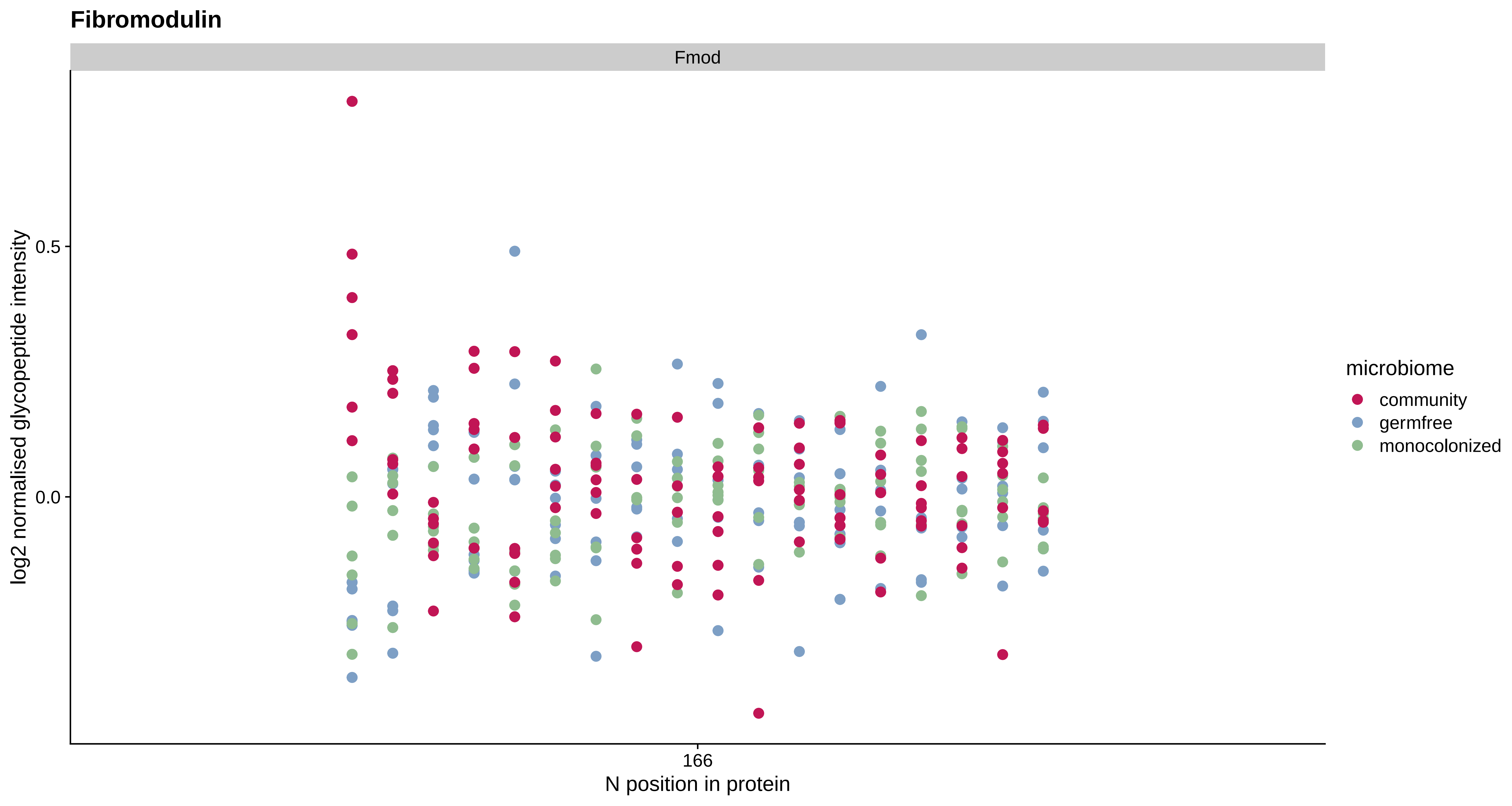

Fibromodulin
Fmod
log2 normalised glycopeptide intensity
0.5
0.0
166
N position in protein
microbiome
community
germfree
monocolonized

#### Neural cell adhesion molecule L1

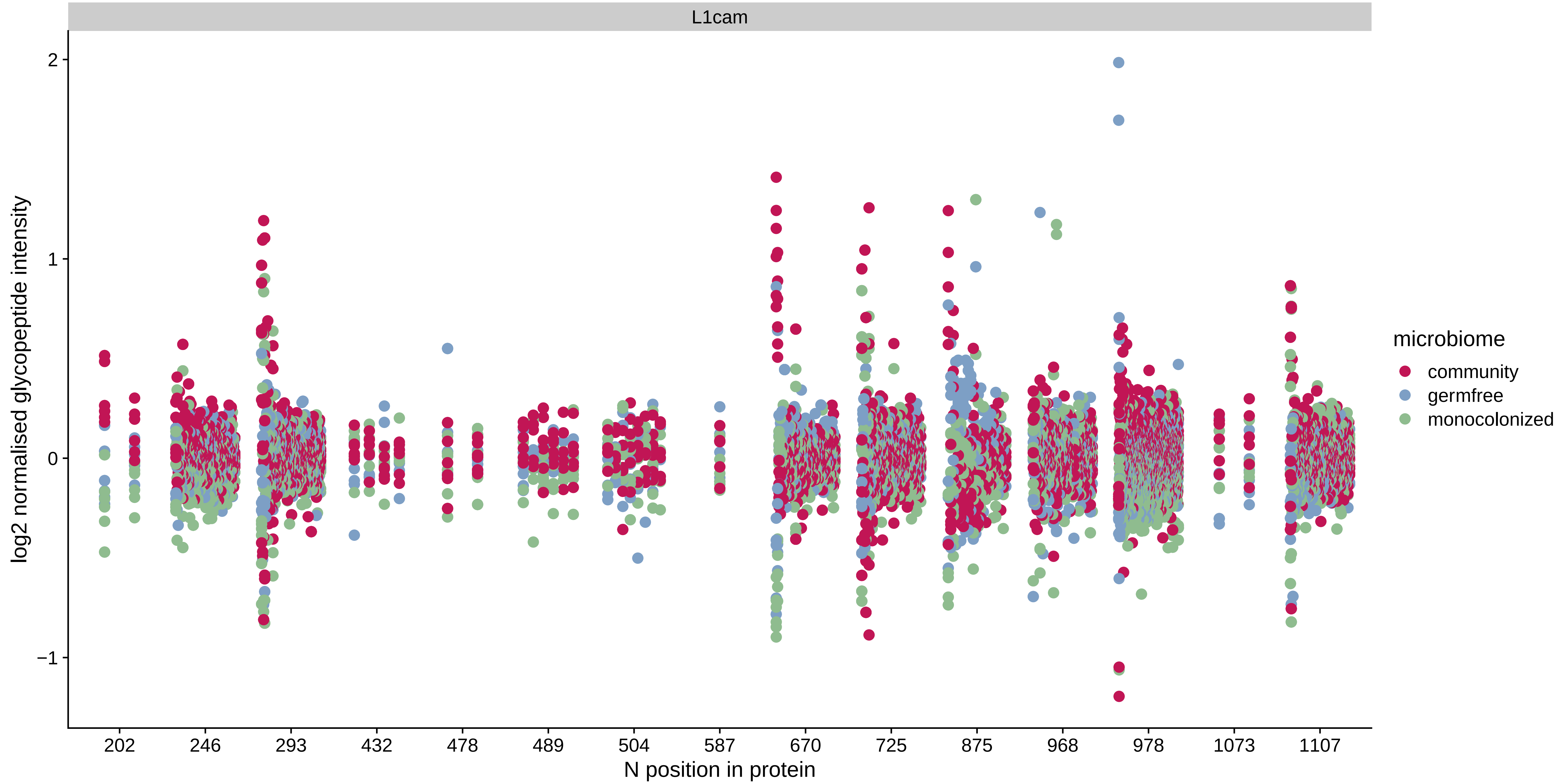

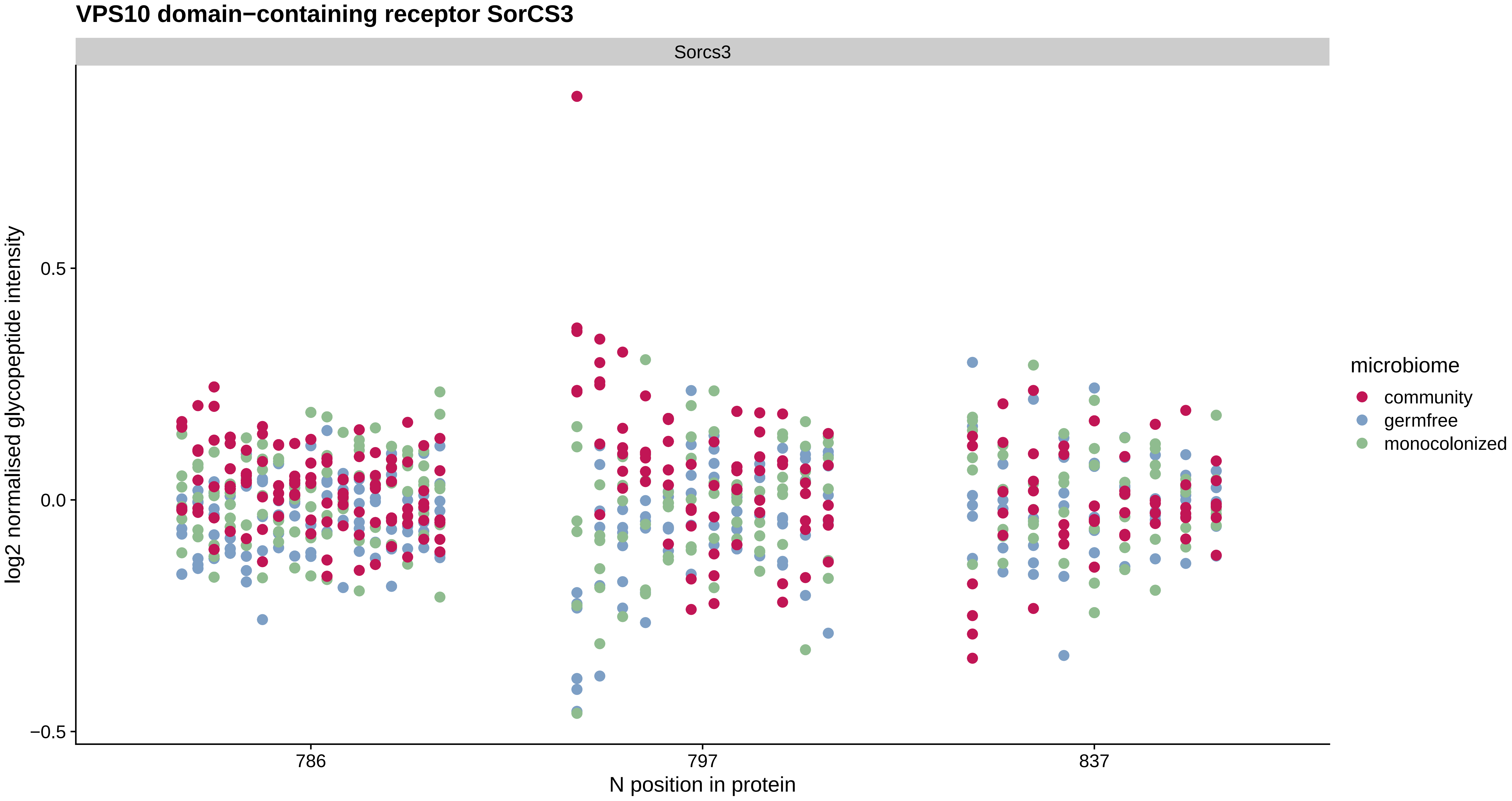

VPS10 domain−containing receptor SorCS3
Sorcs3
log2 normalised glycopeptide intensity
0.5
0.0
−0.5
786
797
837
N position in protein
microbiome
community
germfree
monocolonized

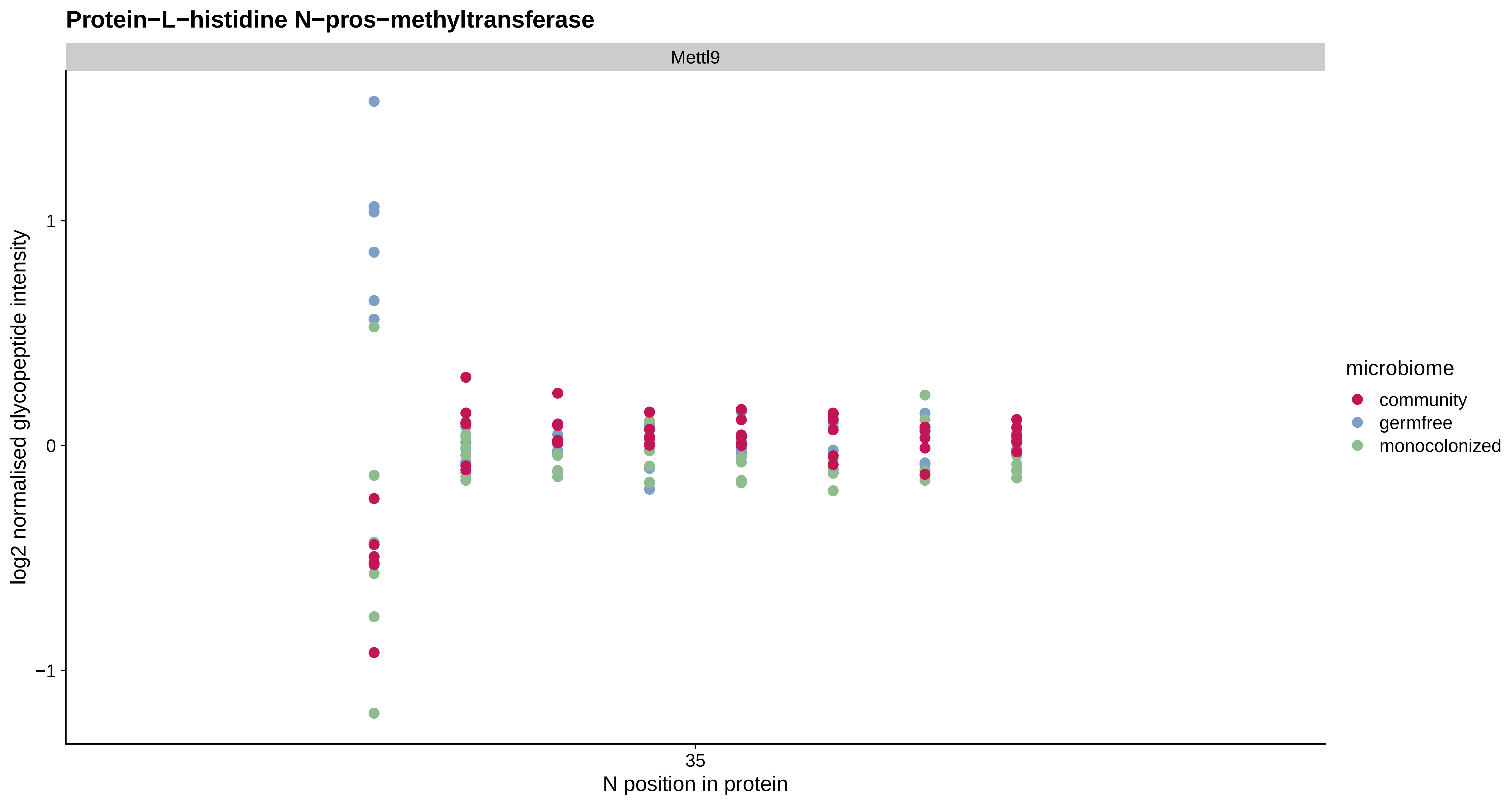
Protein−L−histidine N−pros−methyltransferase
Mettl9
log2 normalised glycopeptide intensity
1
0
−1
35
N position in protein
microbiome
community
germfree
monocolonized

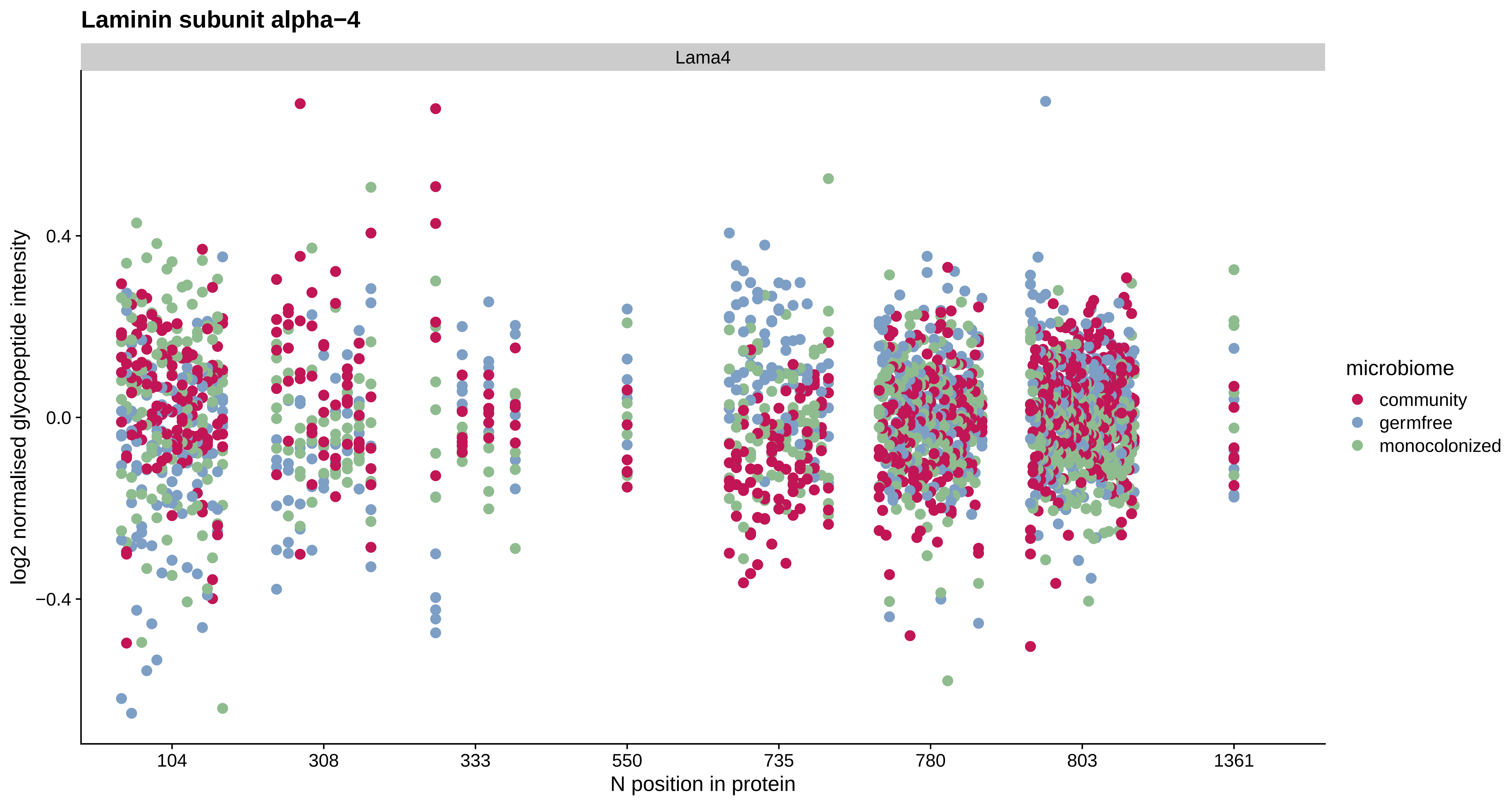

Laminin subunit alpha−4
Lama4
log2 normalised glycopeptide intensity
0.4
0.0
−0.4
104
308
333
550
735
780
803
1361
N position in protein
microbiome
community
germfree
monocolonized

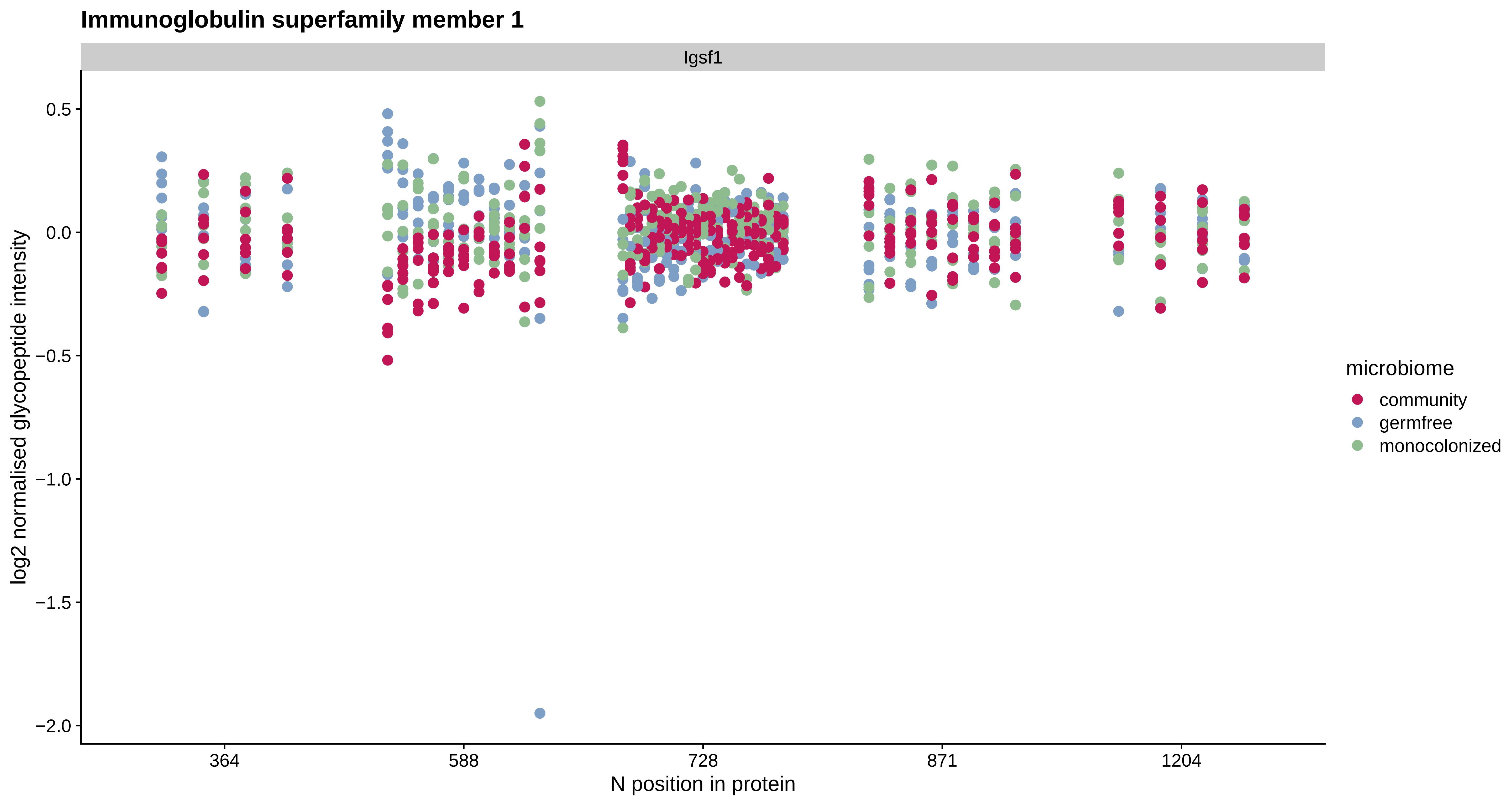
Immunoglobulin superfamily member 1
Igsf1
log2 normalised glycopeptide intensity
0.5
0.0
−0.5
−1.0
−1.5
−2.0
364
588
728
871
1204
N position in protein
microbiome
community
germfree
monocolonized

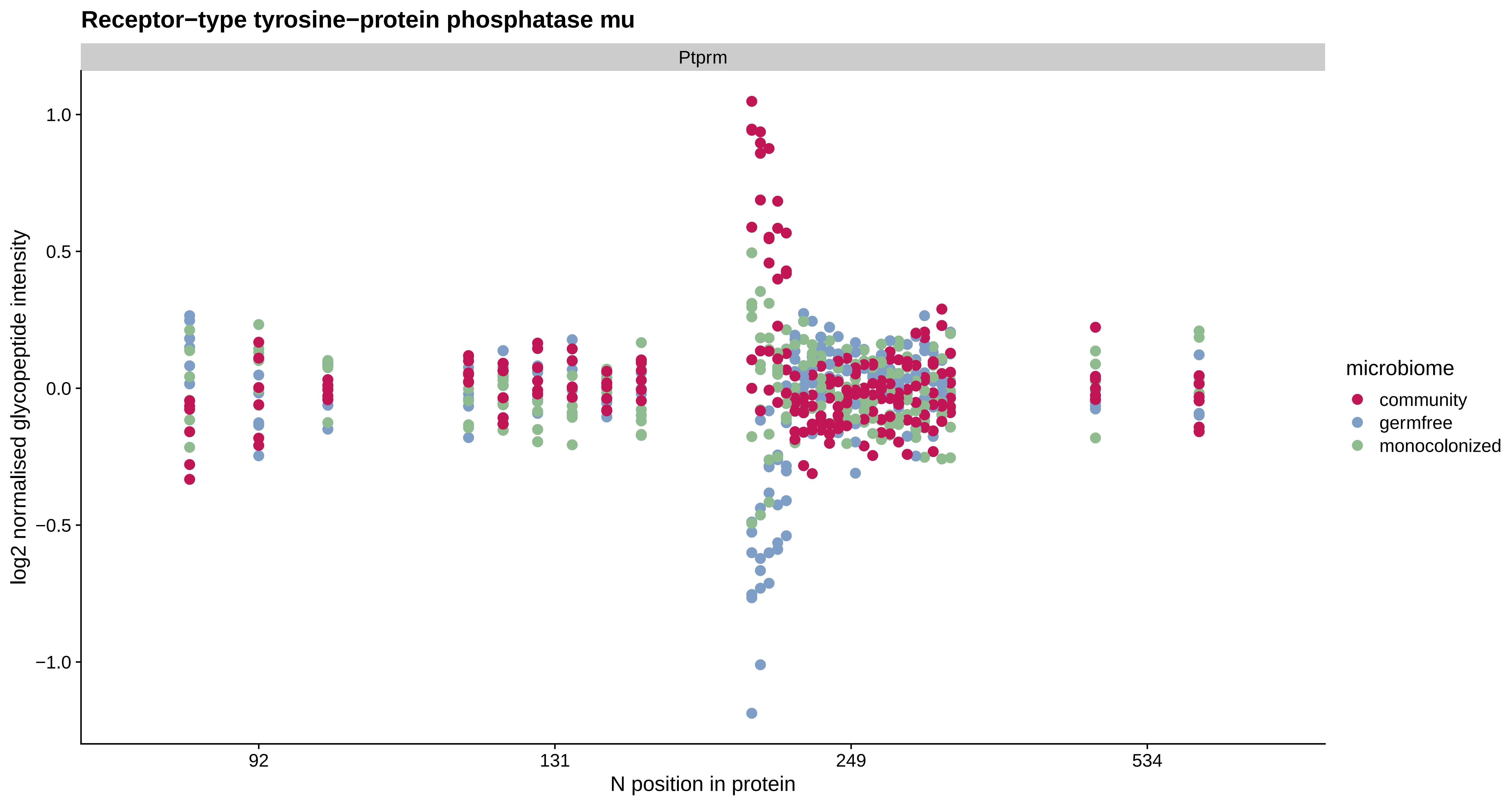

Receptor−type tyrosine−protein phosphatase mu
Ptprm
log2 normalised glycopeptide intensity
1.0
0.5
0.0
−0.5
−1.0
92
131
249
534
N position in protein
microbiome
community
germfree
monocolonized

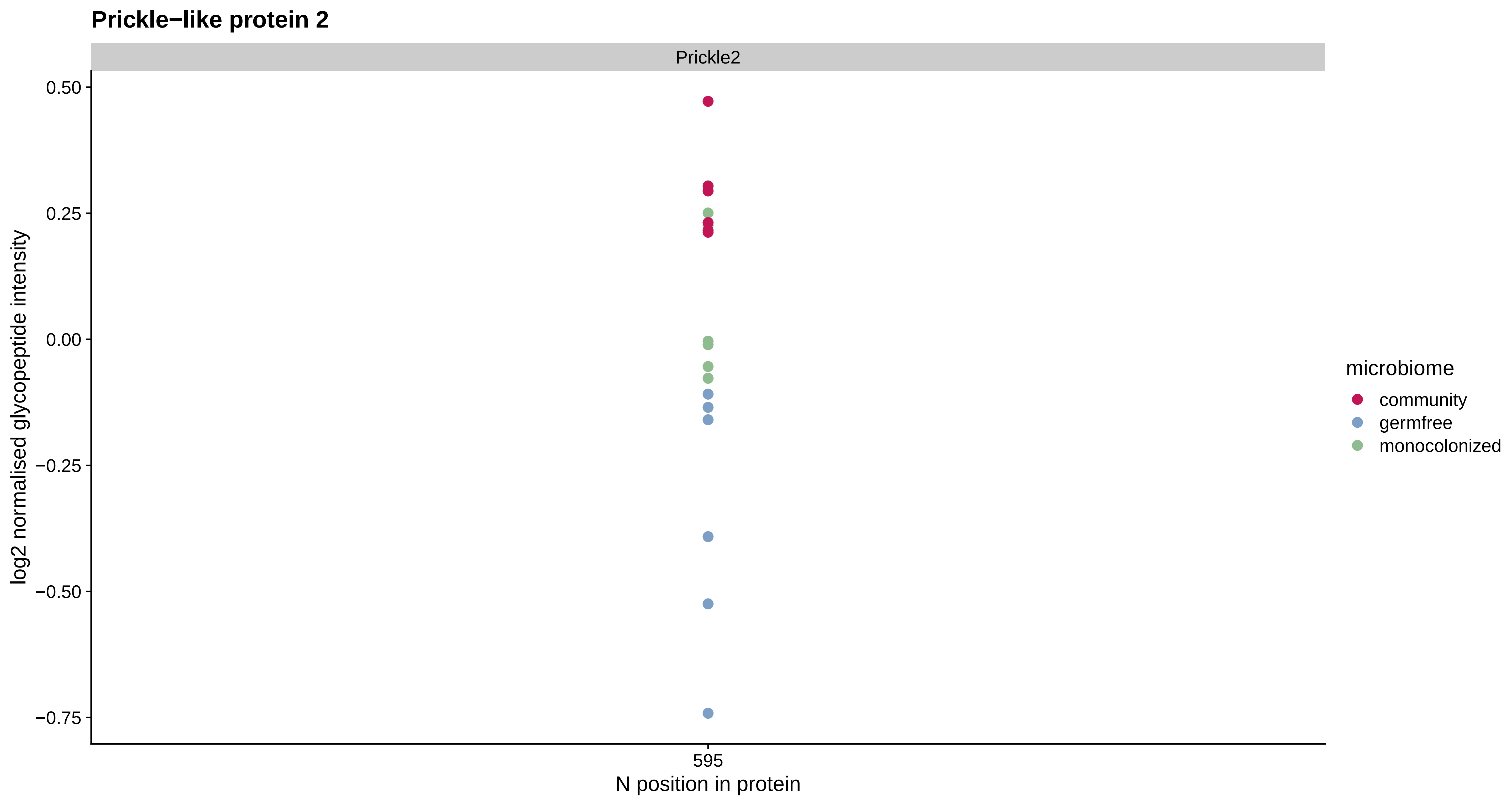
Prickle−like protein 2
Prickle2
0.50
0.25
0.00
−0.25
−0.50
−0.75
log2 normalised glycopeptide intensity
595
N position in protein
microbiome
community
germfree
monocolonized

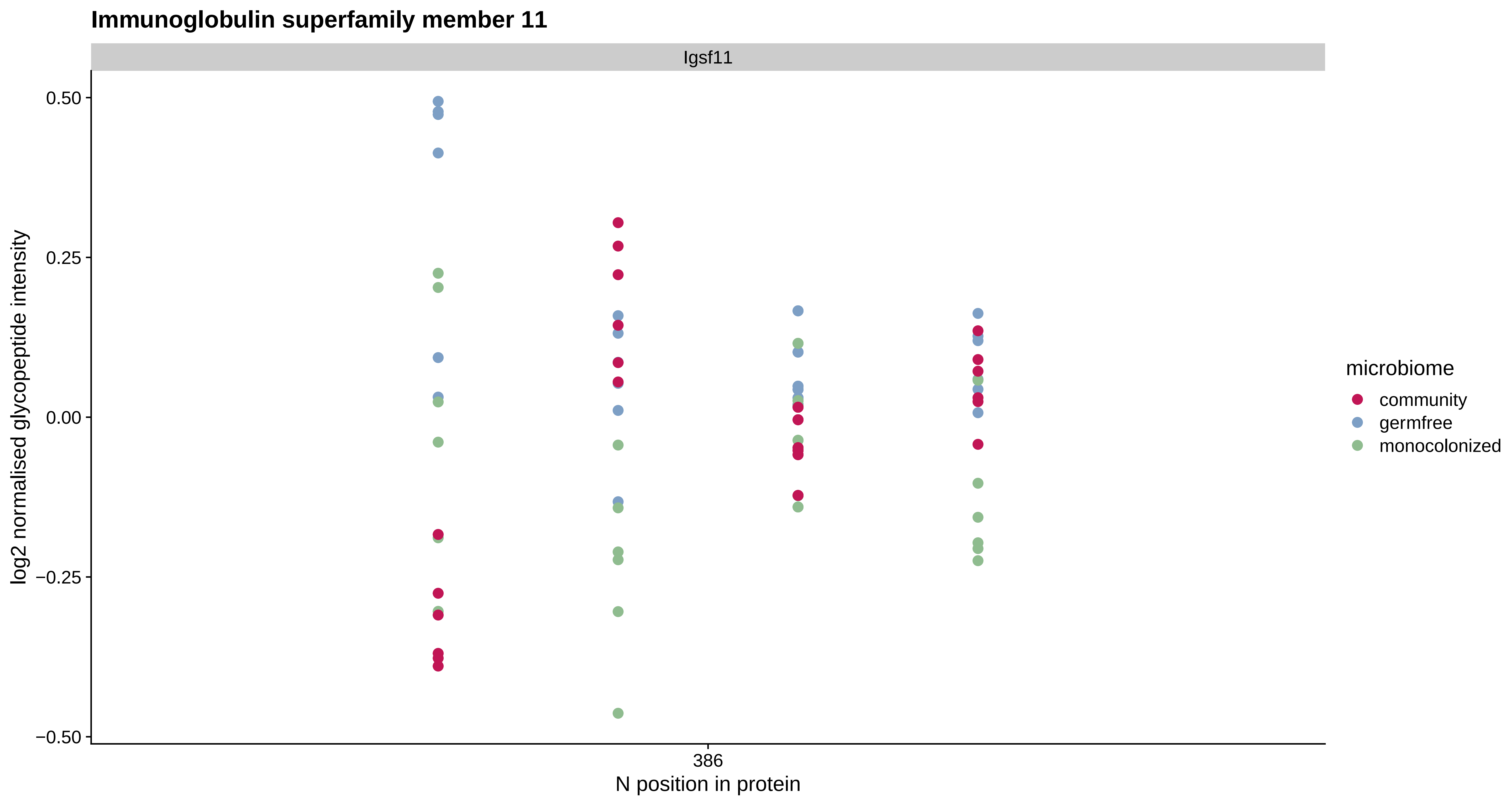
Immunoglobulin superfamily member 11
Igsf11
0.50
0.25
0.00
−0.25
−0.50
log2 normalised glycopeptide intensity
386
N position in protein
microbiome
community
germfree
monocolonized

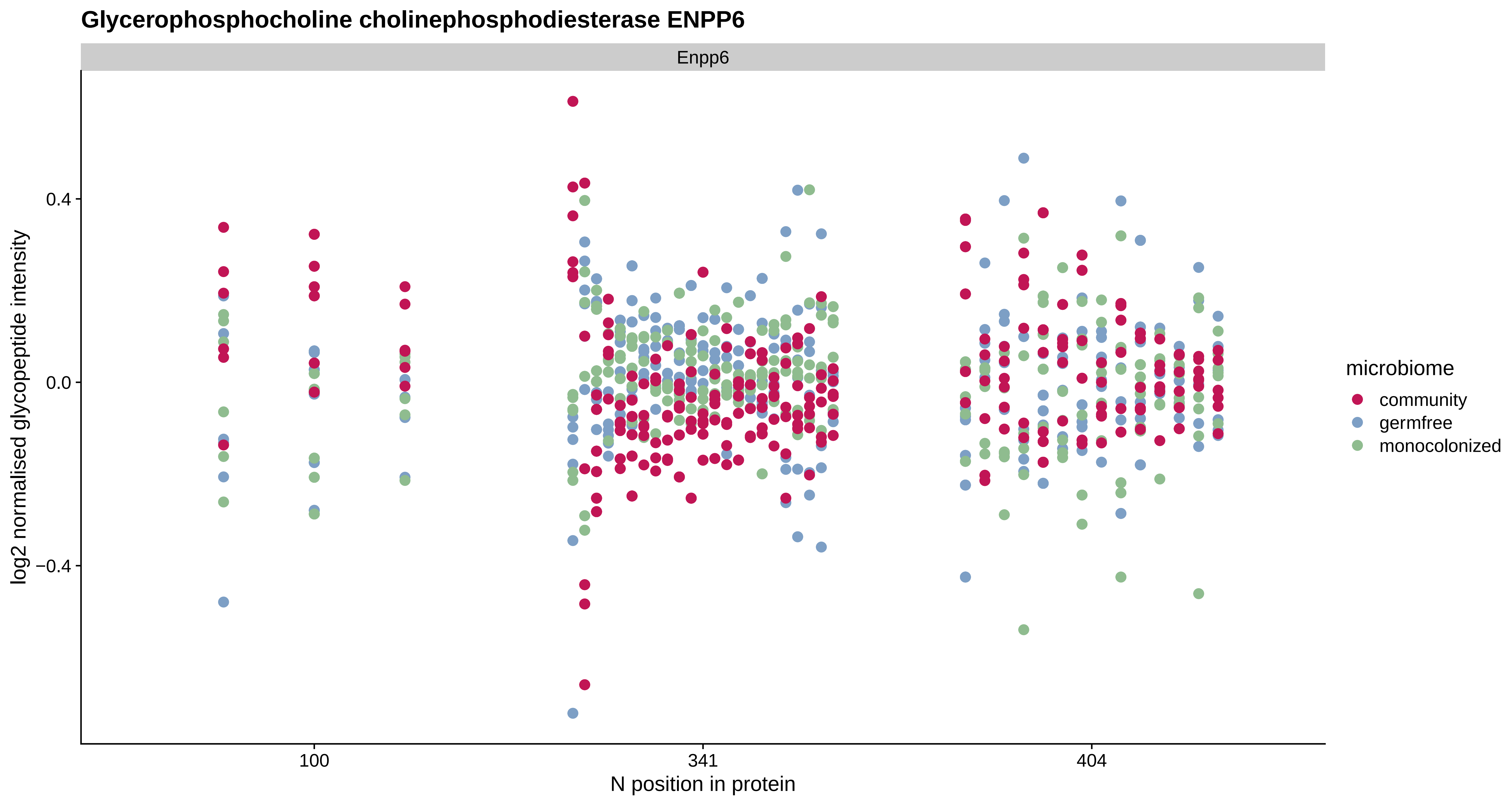
Glycerophosphocholine cholinephosphodiesterase ENPP6
Enpp6
0.4
0.0
−0.4
log2 normalised glycopeptide intensity
100
341
404
N position in protein
microbiome
community
germfree
monocolonized

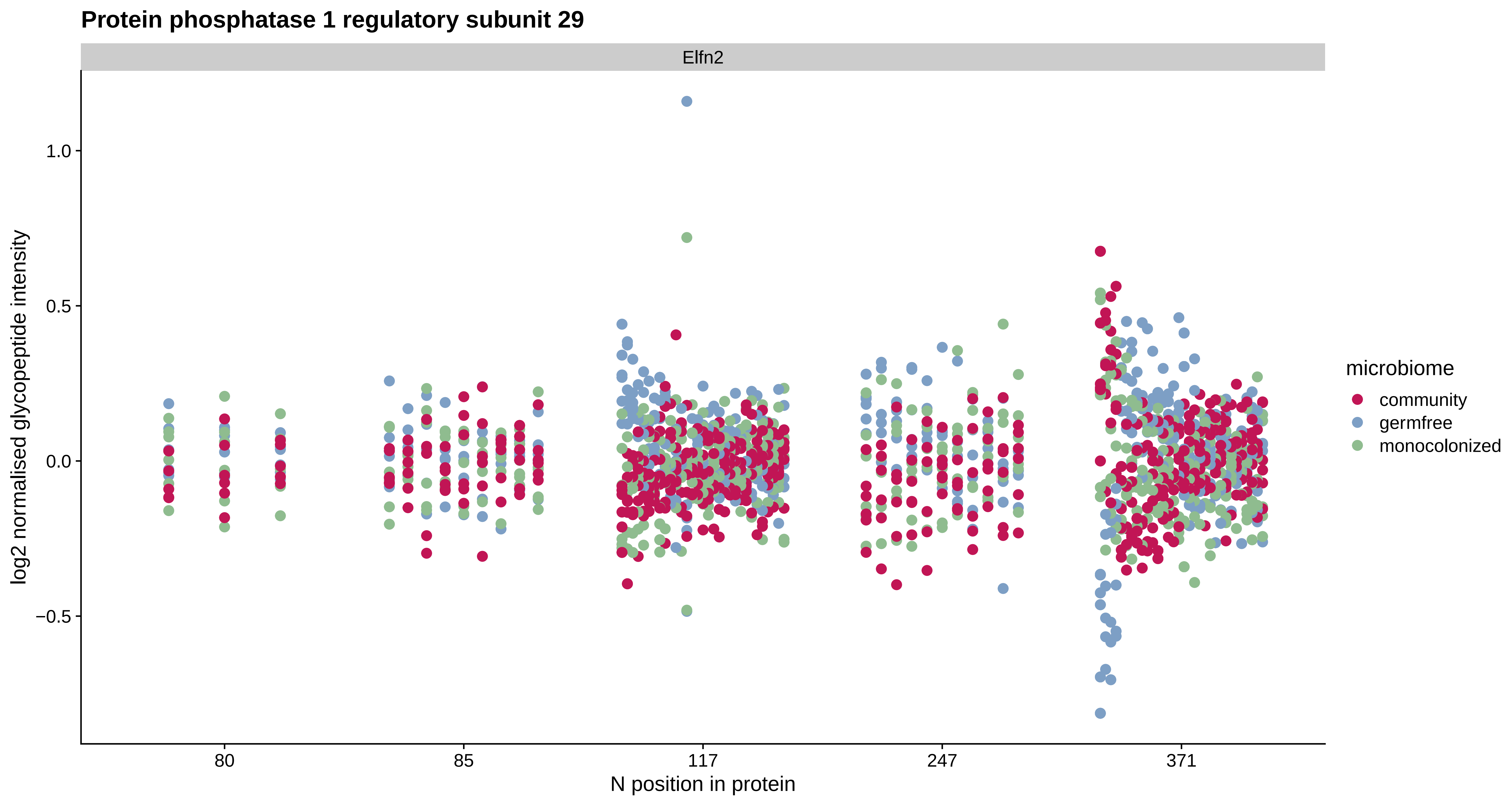

Protein phosphatase 1 regulatory subunit 29
Elfn2
log2 normalised glycopeptide intensity
1.0
0.5
0.0
−0.5
80
85
117
247
371
N position in protein
microbiome
community
germfree
monocolonized

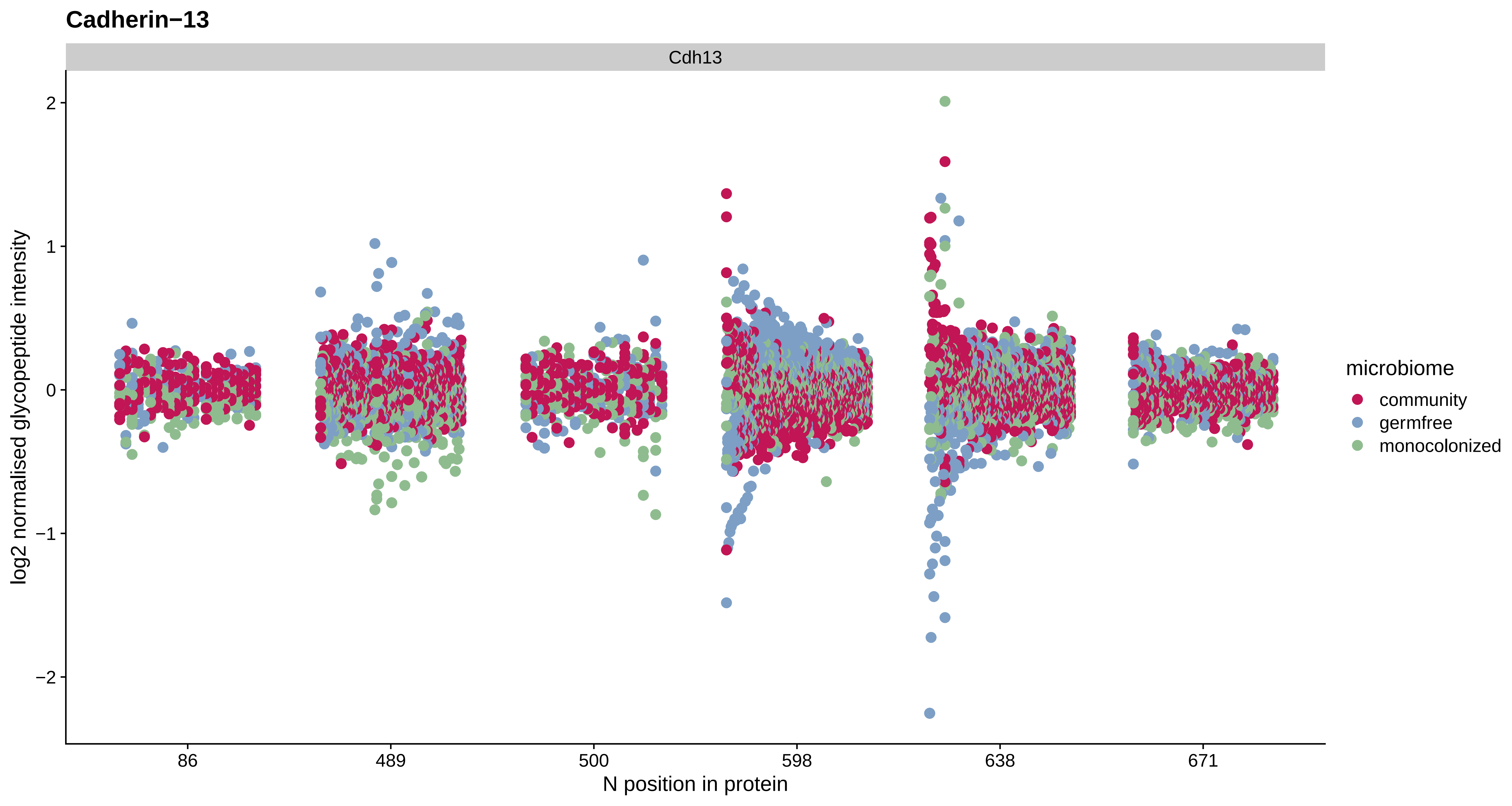

Cadherin−13
Cdh13
log2 normalised glycopeptide intensity
2
1
0
−1
−2
86
489
500
598
638
671
N position in protein
microbiome
community
germfree
monocolonized

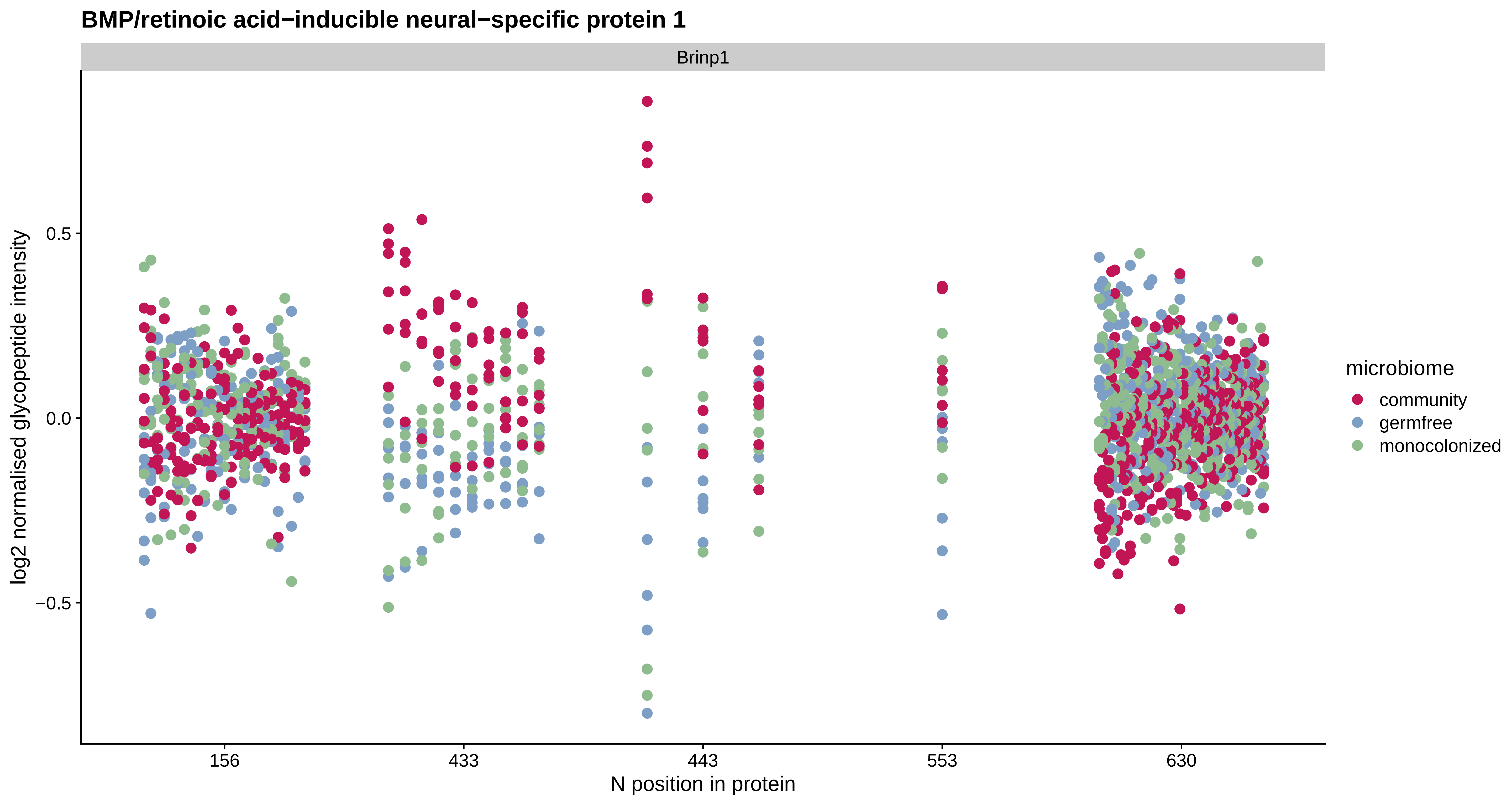
BMP/retinoic acid−inducible neural−specific protein 1
Brinp1
log2 normalised glycopeptide intensity
0.5
0.0
−0.5
156
433
443
553
630
N position in protein
microbiome
community
germfree
monocolonized

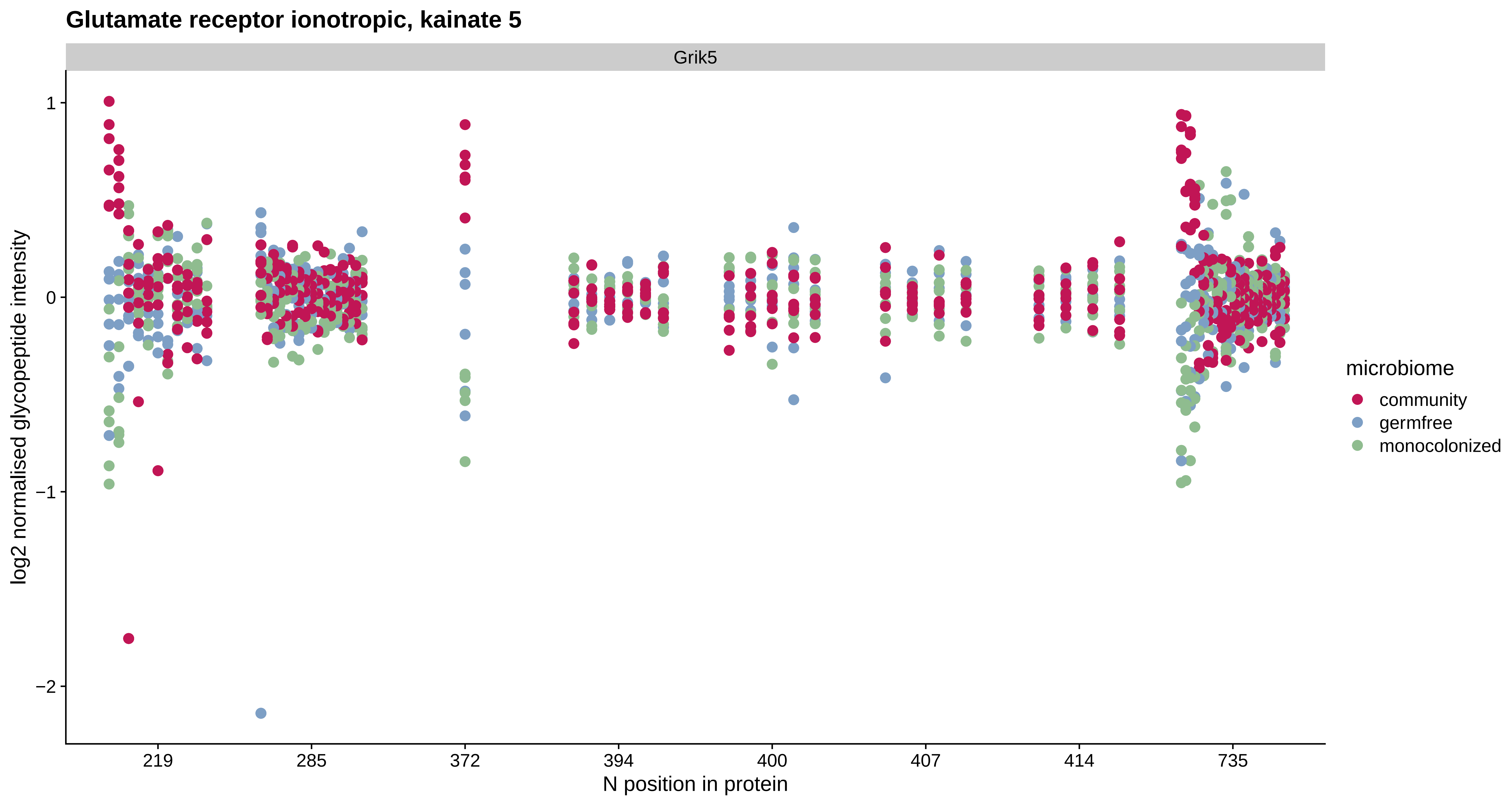
Glutamate receptor ionotropic, kainate 5
Grik5
log2 normalised glycopeptide intensity
1
0
−1
−2
219
285
372
394
400
407
414
735
N position in protein
microbiome
community
germfree
monocolonized

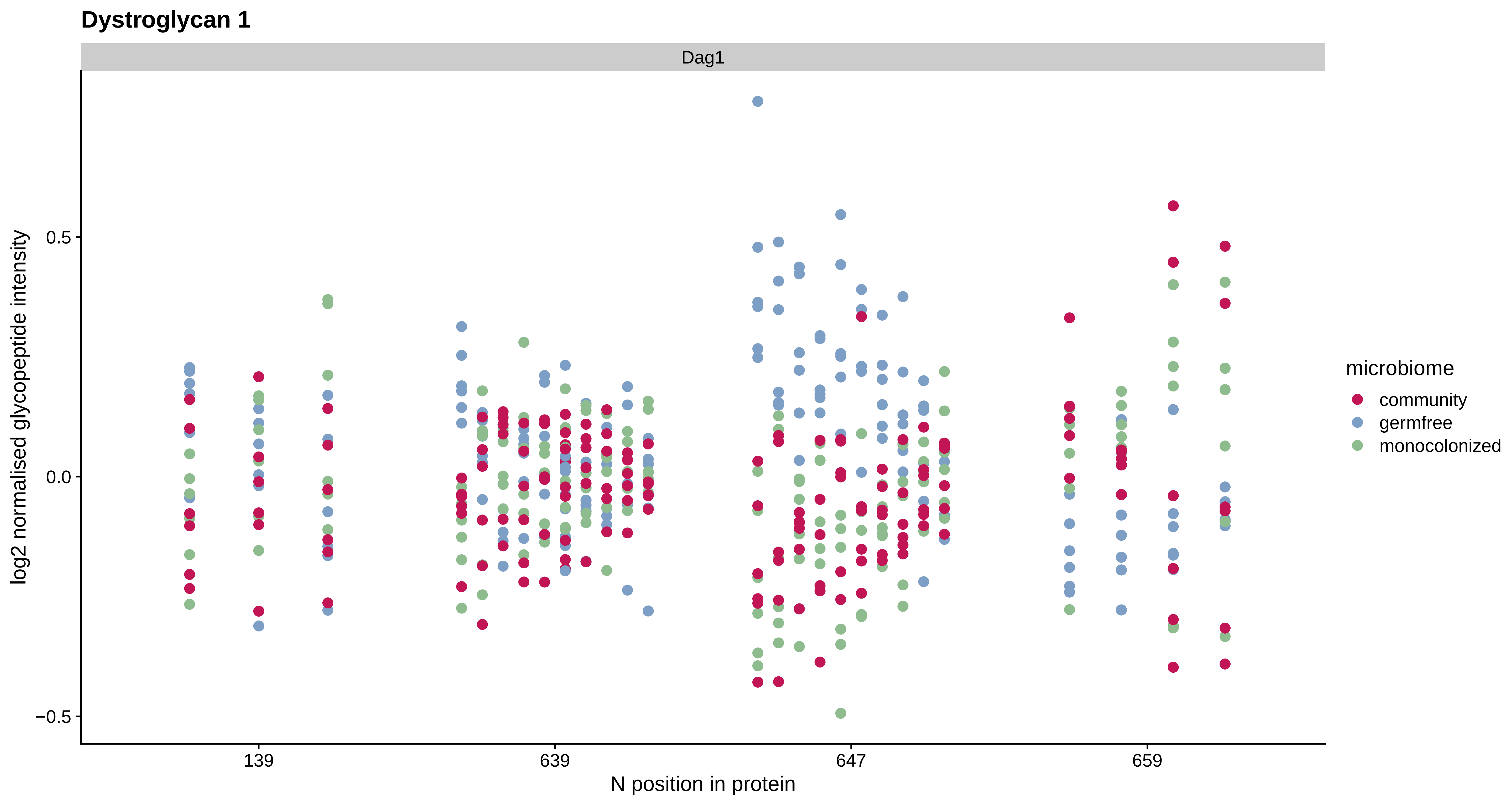
Dystroglycan 1
Dag1
log2 normalised glycopeptide intensity
0.5
0.0
−0.5
139
639
647
659
N position in protein
microbiome
community
germfree
monocolonized

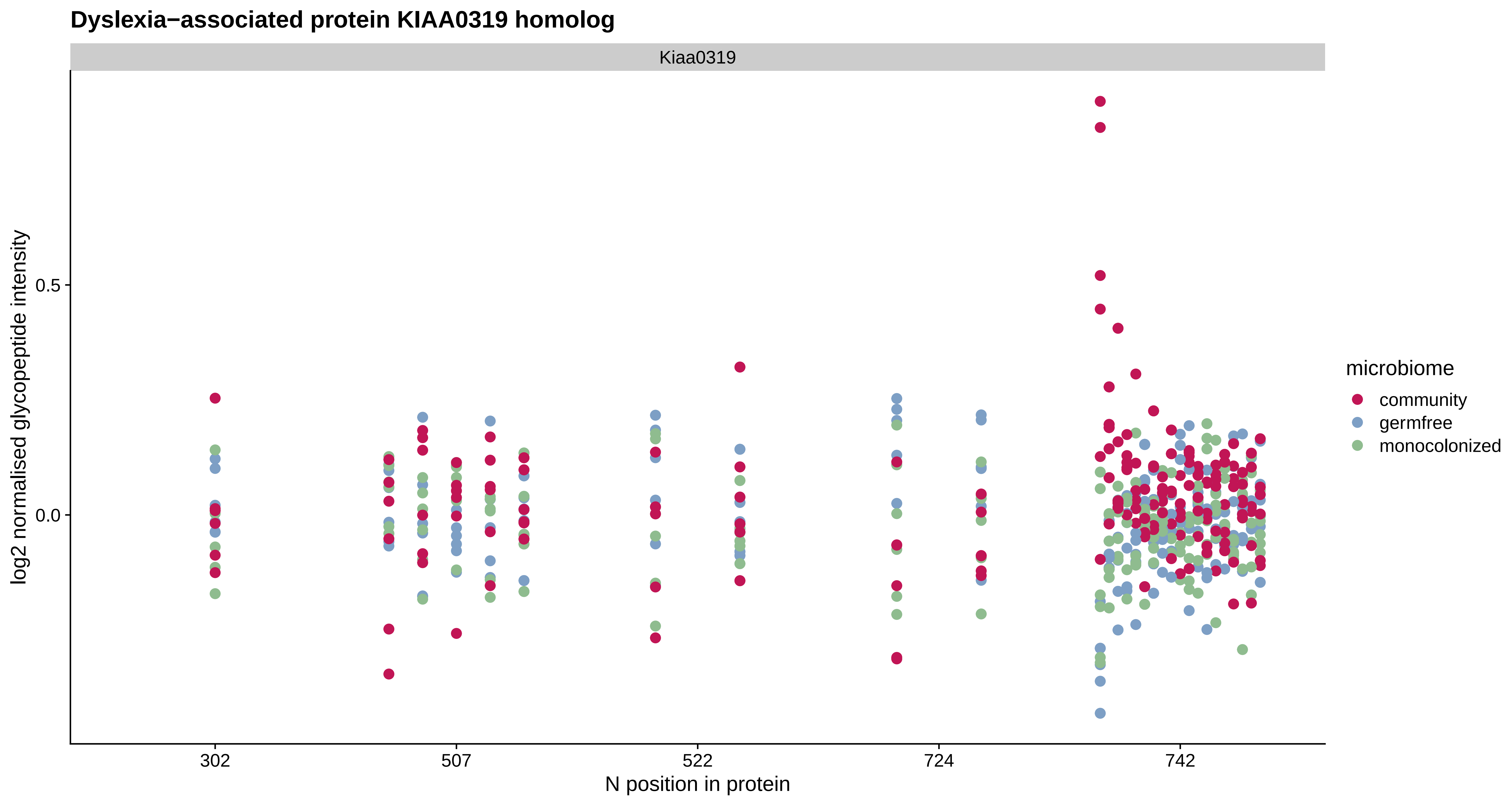

Dyslexia−associated protein KIAA0319 homolog
Kiaa0319
log2 normalised glycopeptide intensity
0.5
0.0
302
507
522
724
742
N position in protein
microbiome
community
germfree
monocolonized

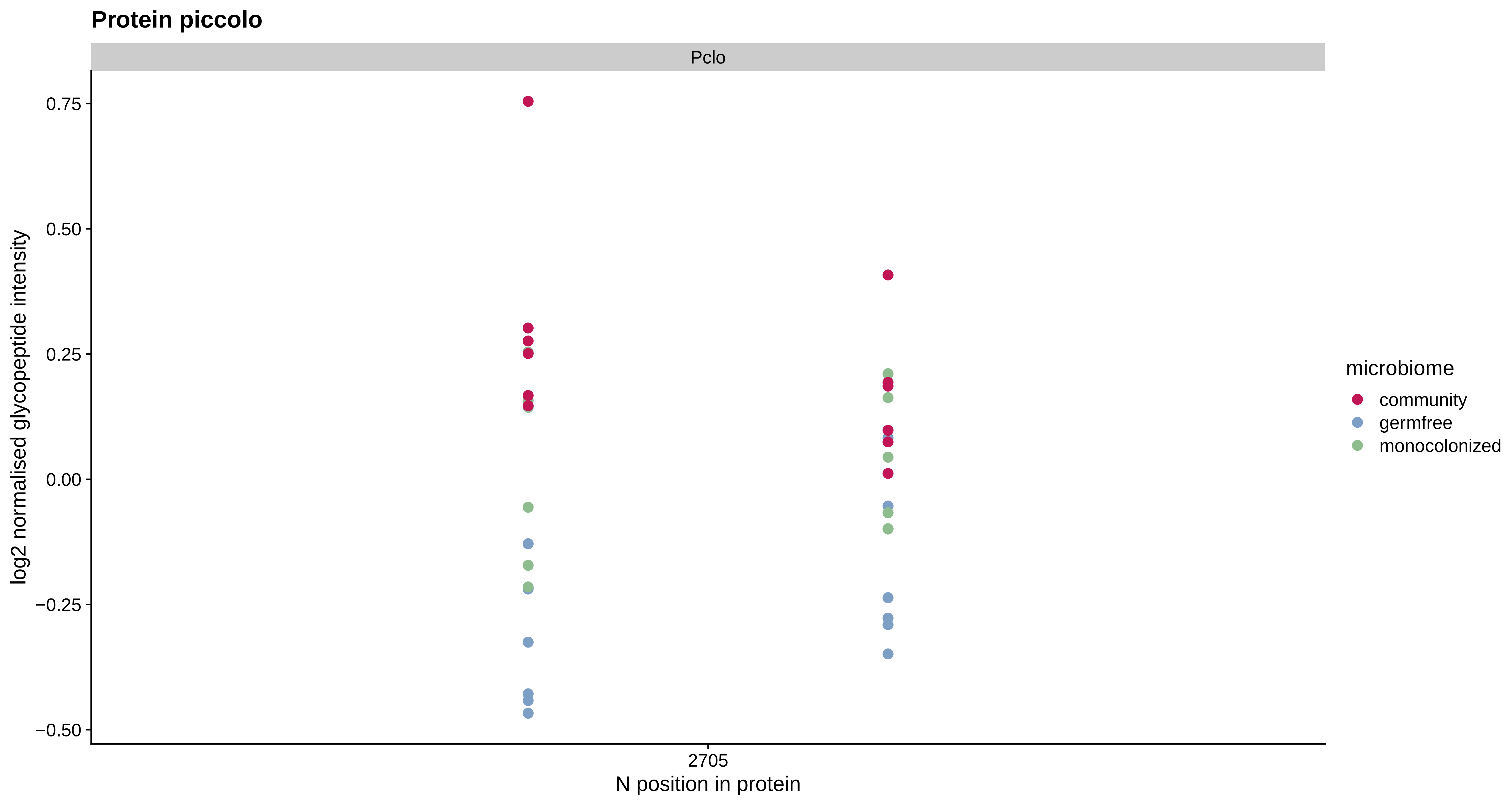
Protein piccolo
Pclo
0.75
0.50
0.25
0.00
−0.25
−0.50
log2 normalised glycopeptide intensity
2705
N position in protein
microbiome
community
germfree
monocolonized

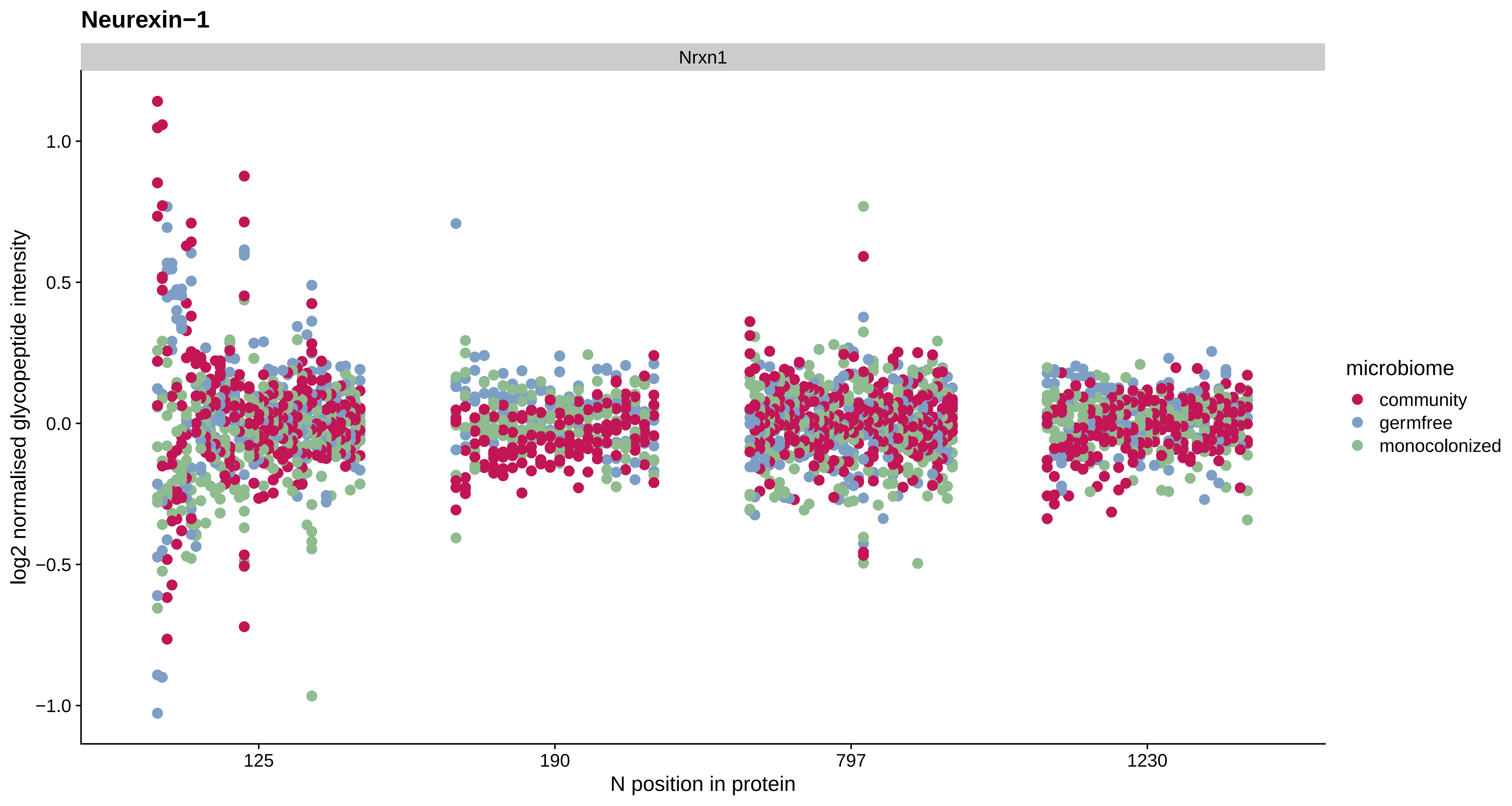

Neurexin−1
Nrxn1
log2 normalised glycopeptide intensity
1.0
0.5
0.0
−0.5
−1.0
125
190
797
1230
N position in protein
microbiome
community
germfree
monocolonized

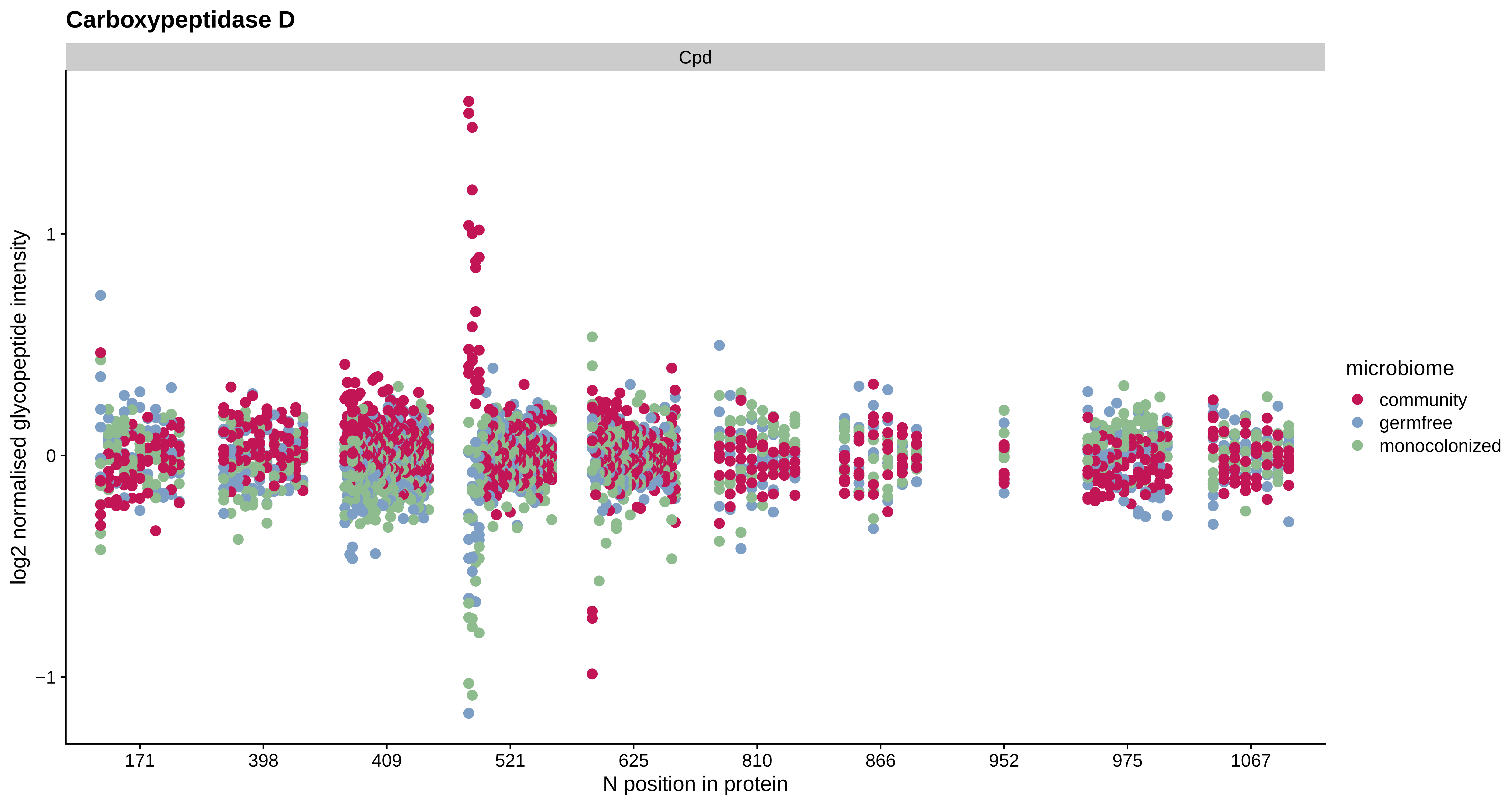
Carboxypeptidase D
Cpd
log2 normalised glycopeptide intensity
1
0
−1
171
398
409
521
625
810
866
952
975
1067
N position in protein
microbiome
community
germfree
monocolonized

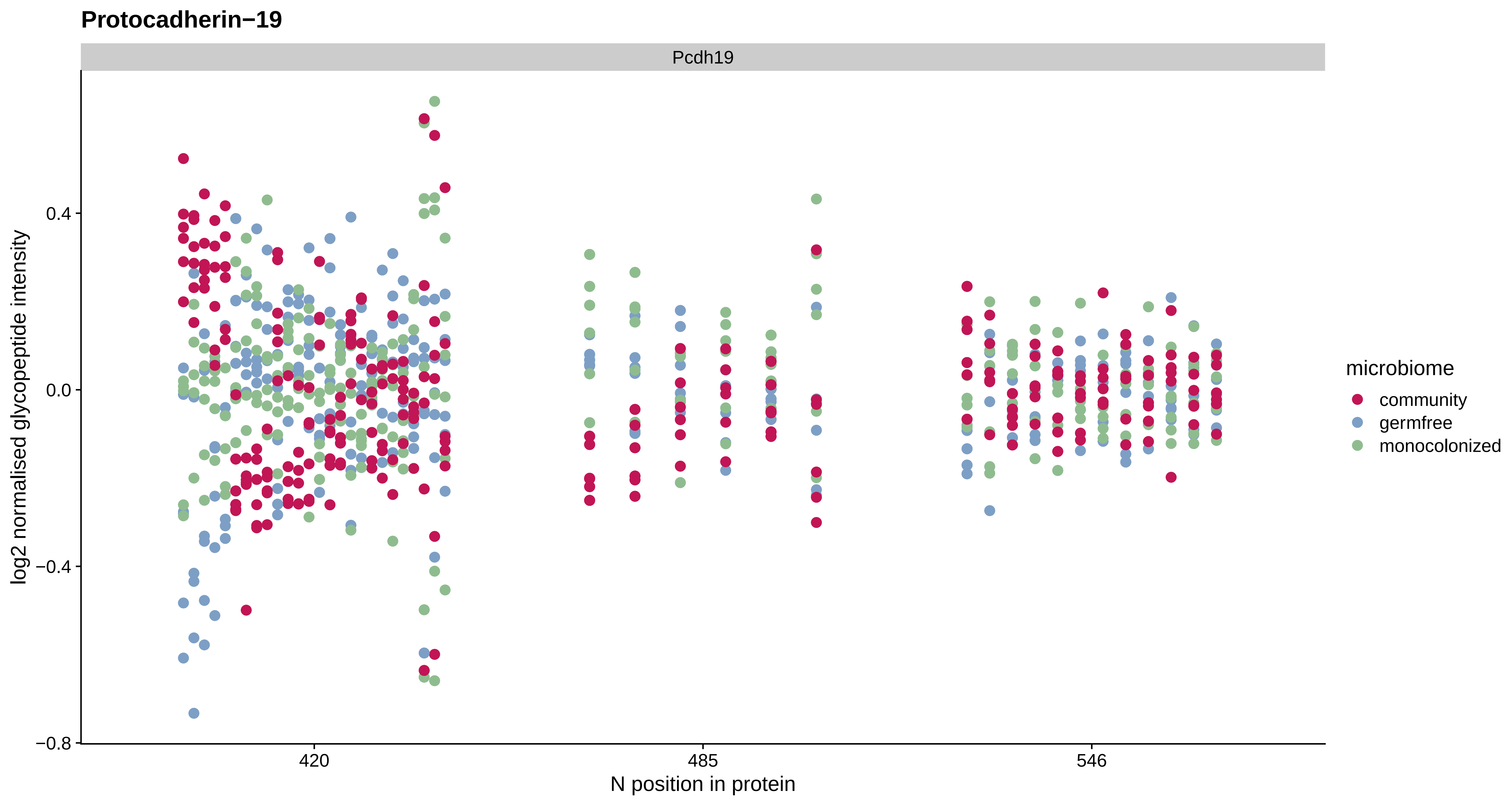
Protocadherin−19
Pcdh19
log2 normalised glycopeptide intensity
0.4
0.0
−0.4
−0.8
420
485
546
N position in protein
microbiome
community
germfree
monocolonized

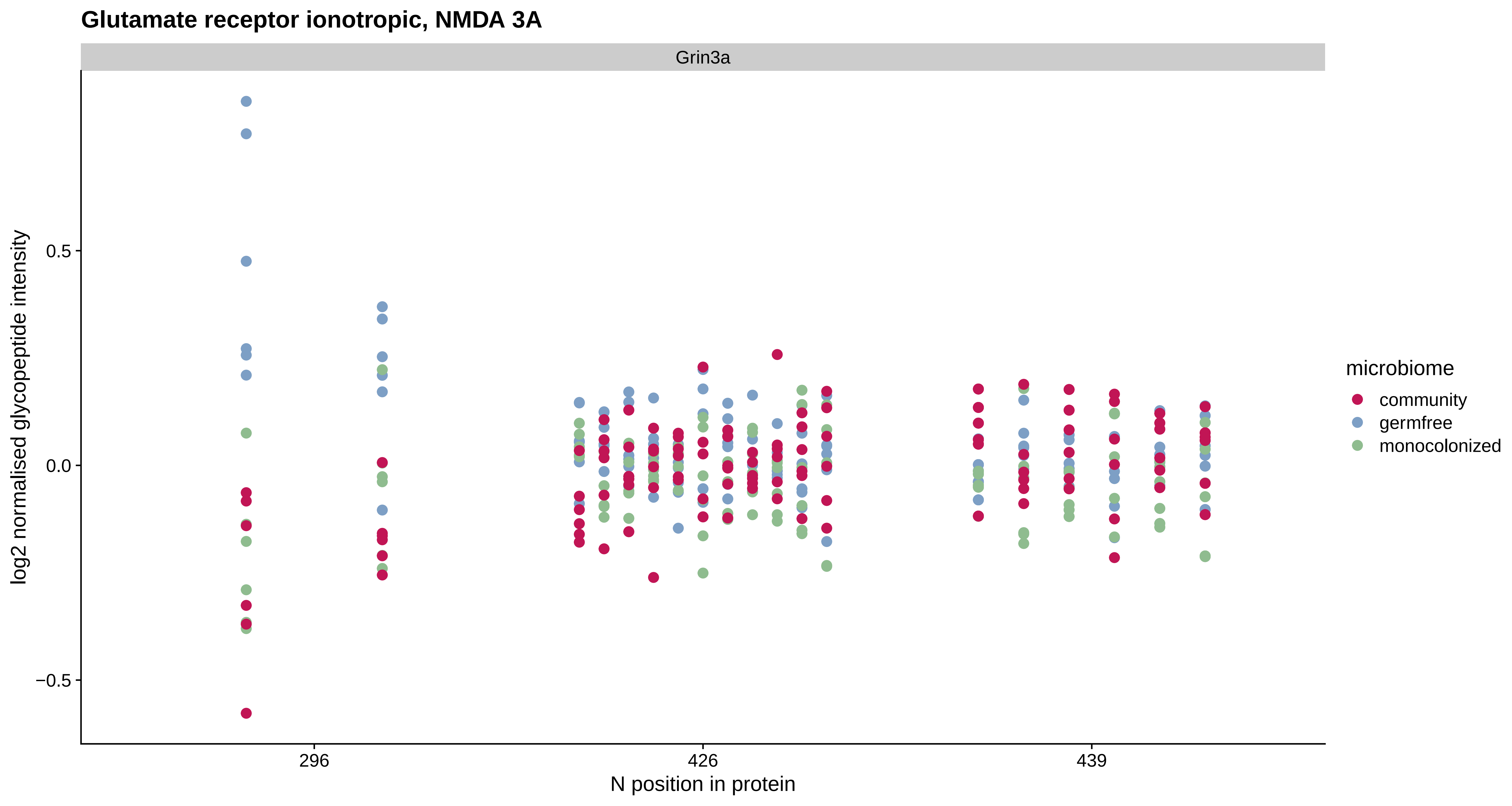
Glutamate receptor ionotropic, NMDA 3A
Grin3a
log2 normalised glycopeptide intensity
0.5
0.0
−0.5
296
426
439
N position in protein
microbiome
community
germfree
monocolonized

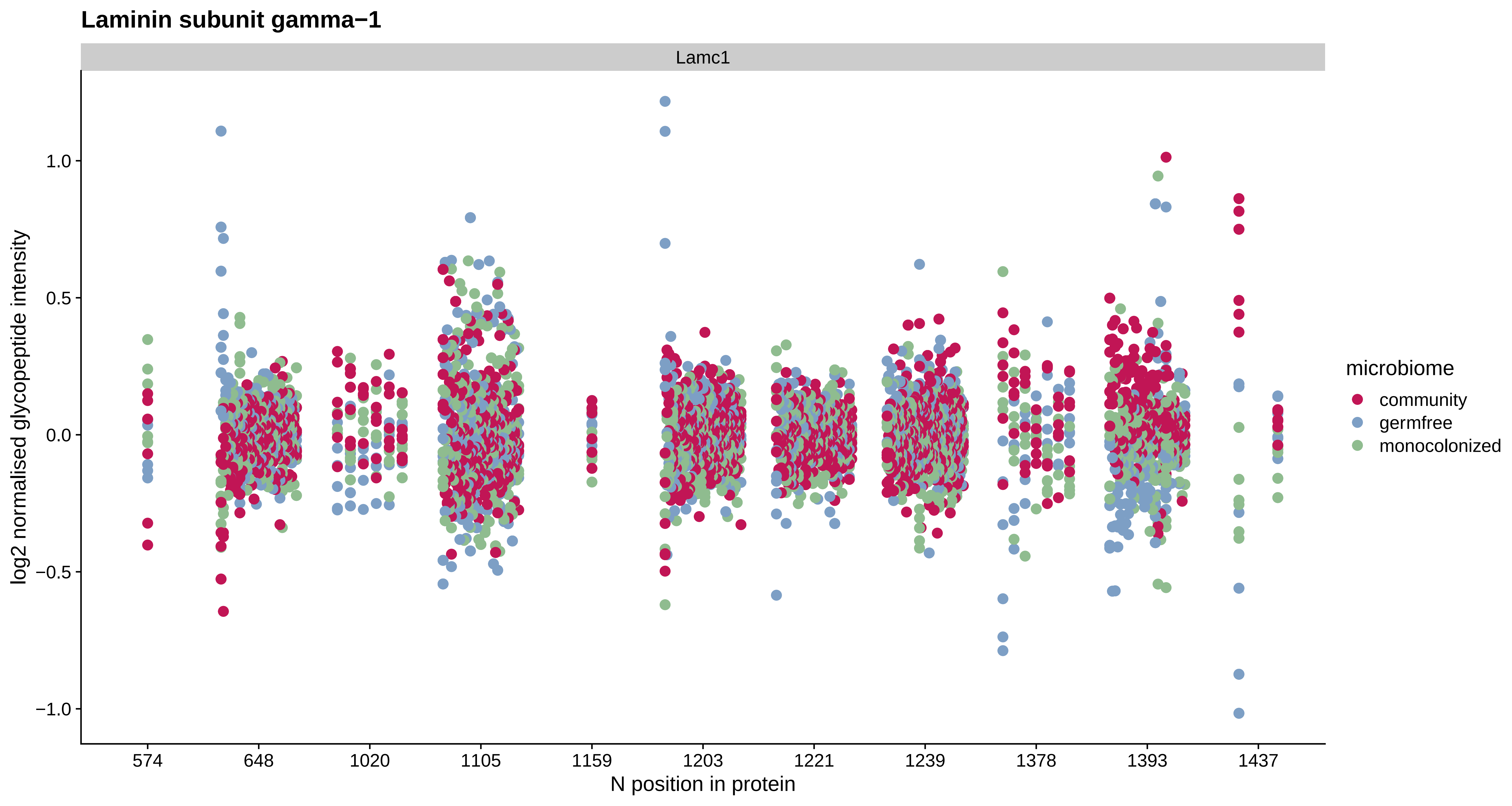
Laminin subunit gamma−1
Lamc1
log2 normalised glycopeptide intensity
1.0
0.5
0.0
−0.5
−1.0
574
648
1020
1105
1159
1203
1221
1239
1378
1393
1437
N position in protein
microbiome
community
germfree
monocolonized

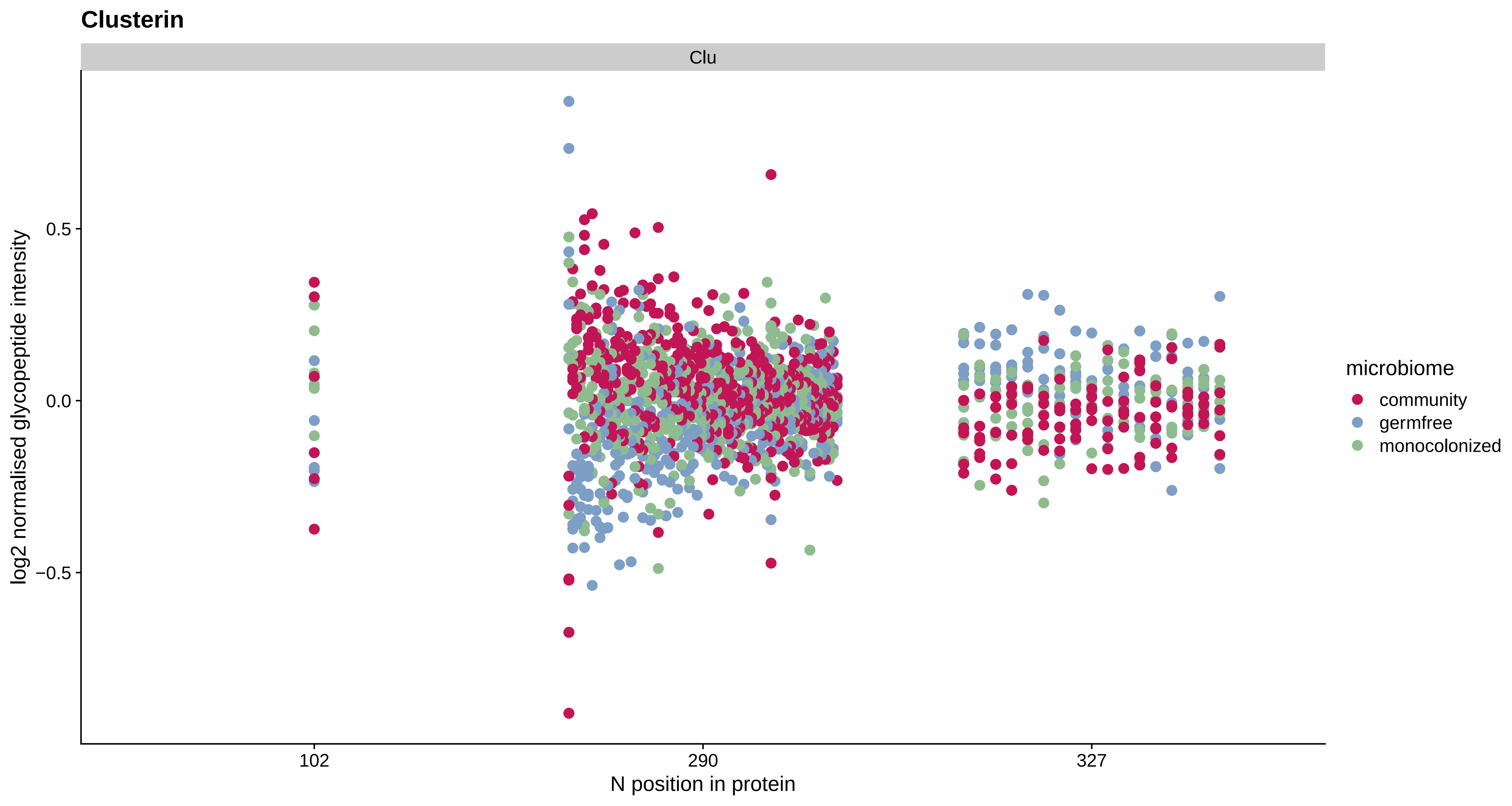
Clusterin
Clu
log2 normalised glycopeptide intensity
0.5
0.0
−0.5
102
290
327
N position in protein
microbiome
community
germfree
monocolonized

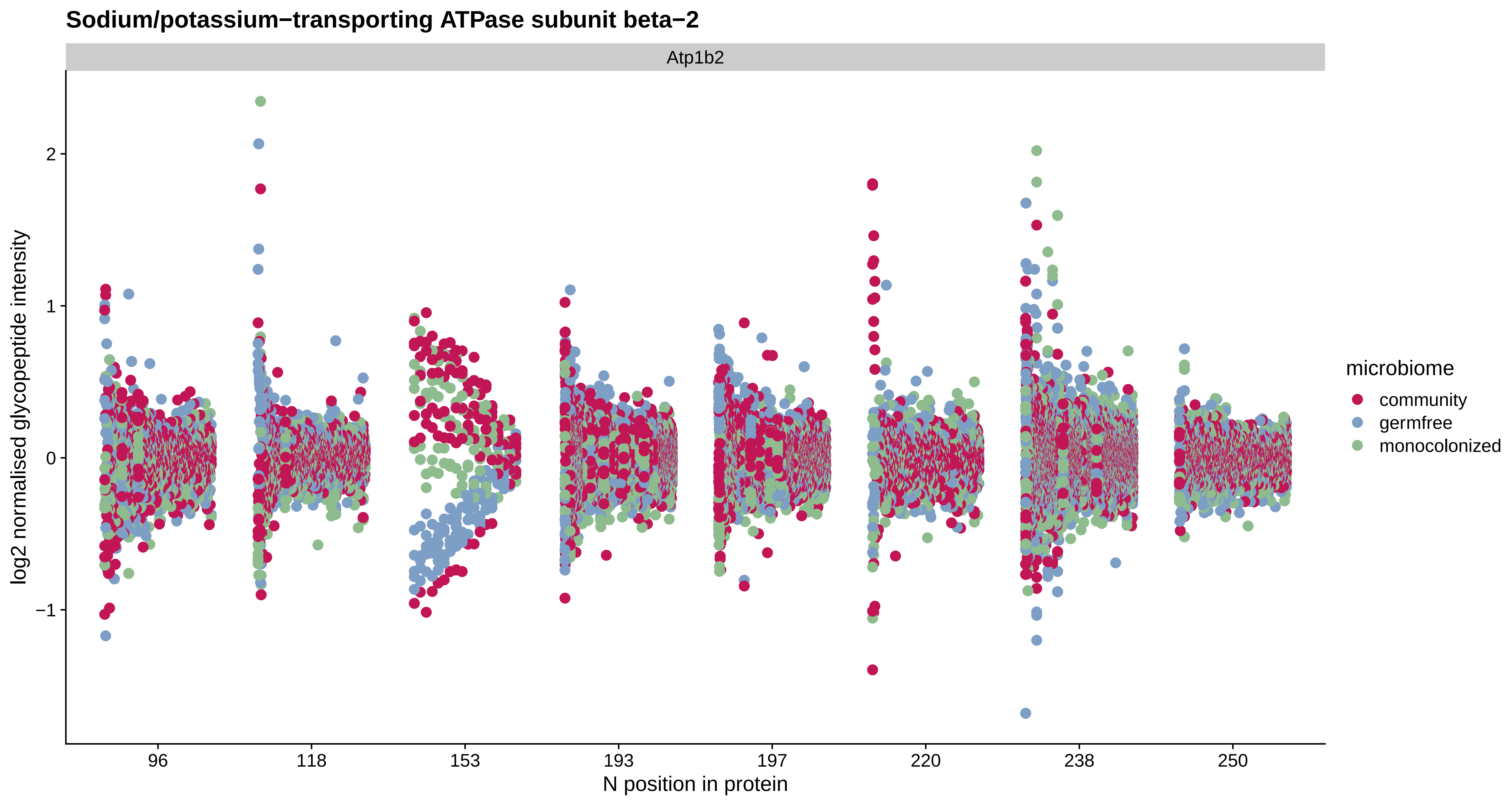

Sodium/potassium−transporting ATPase subunit beta−2
Atp1b2
log2 normalised glycopeptide intensity
2
1
0
−1
96
118
153
193
197
220
238
250
N position in protein
microbiome
community
germfree
monocolonized

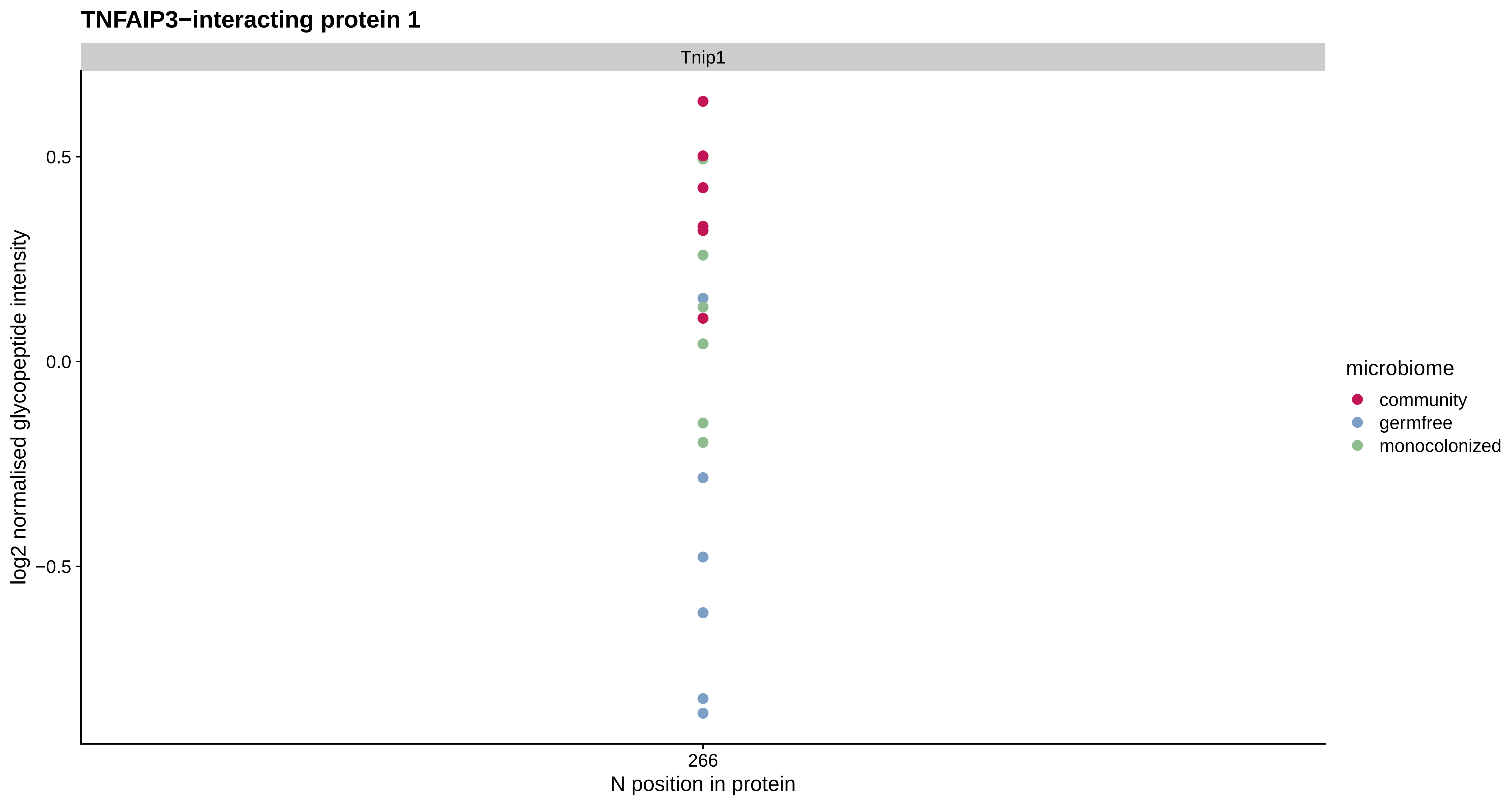
TNFAIP3−interacting protein 1
Tnip1
0.5
0.0
−0.5
log2 normalised glycopeptide intensity
266
N position in protein
microbiome
community
germfree
monocolonized

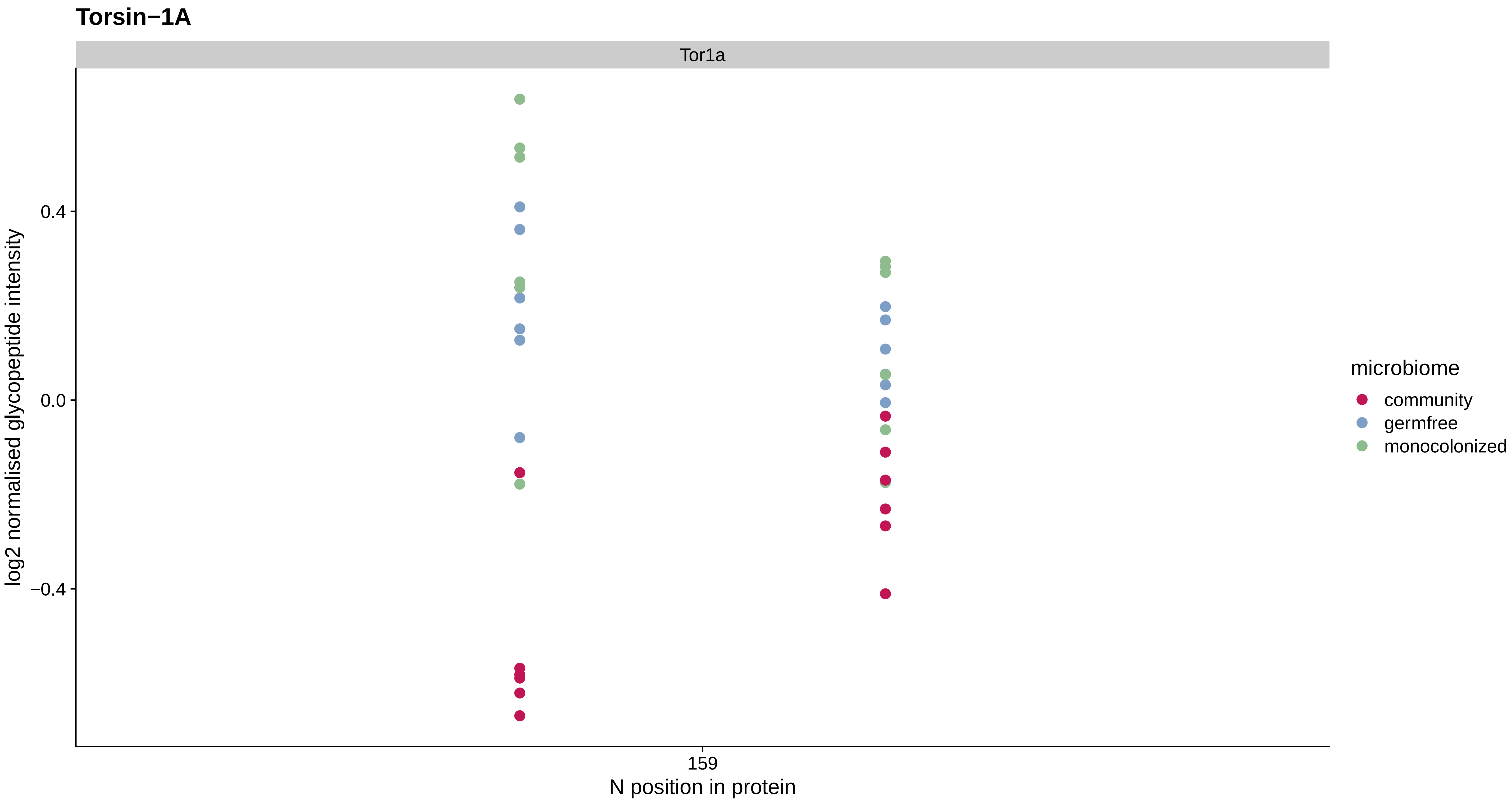
Torsin−1A
Tor1a
log2 normalised glycopeptide intensity
0.4
0.0
−0.4
159
N position in protein
microbiome
community
germfree
monocolonized

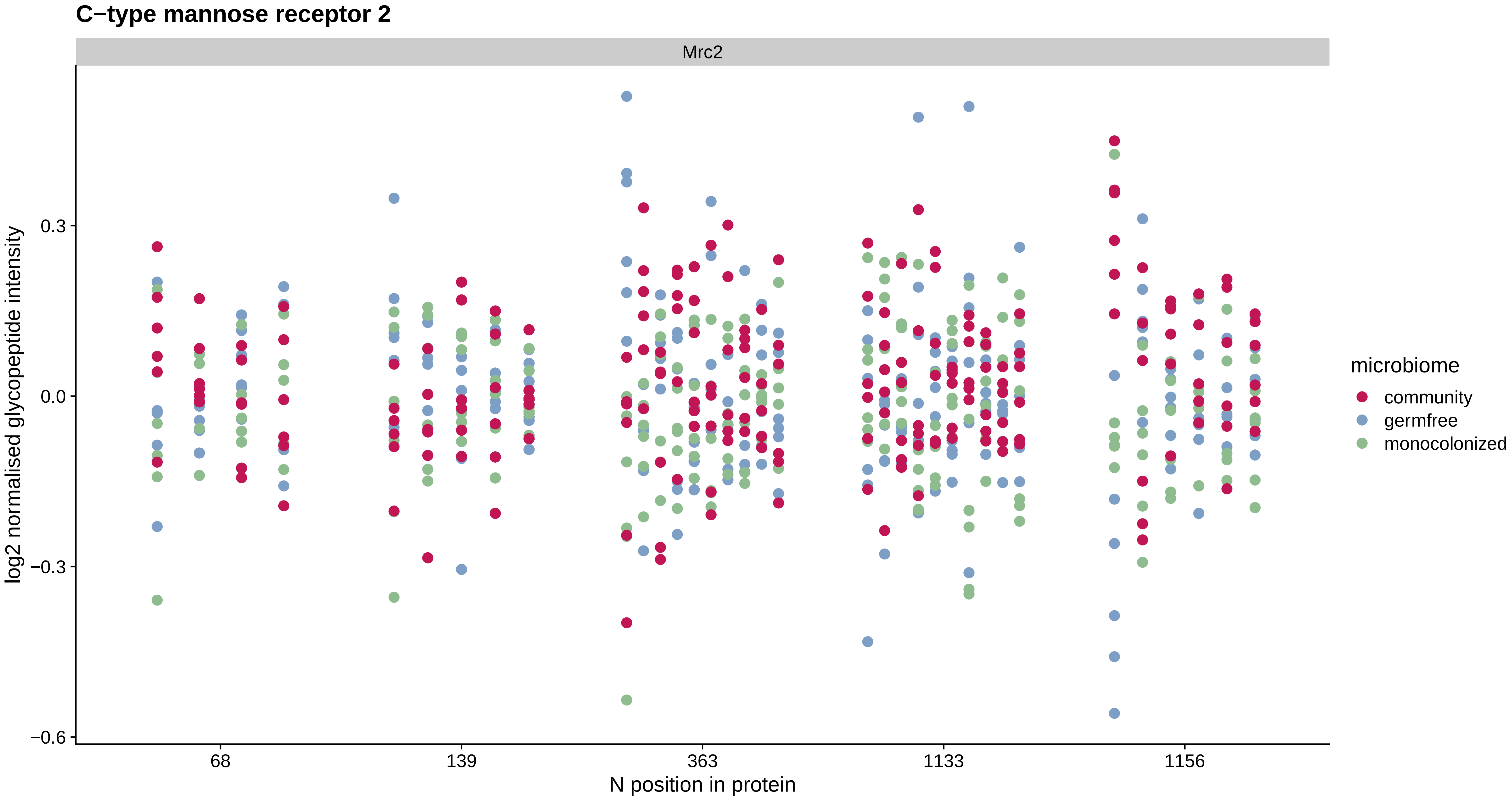

C−type mannose receptor 2
Mrc2
log2 normalised glycopeptide intensity
0.3
0.0
−0.3
−0.6
68
139
363
1133
1156
N position in protein
microbiome
community
germfree
monocolonized

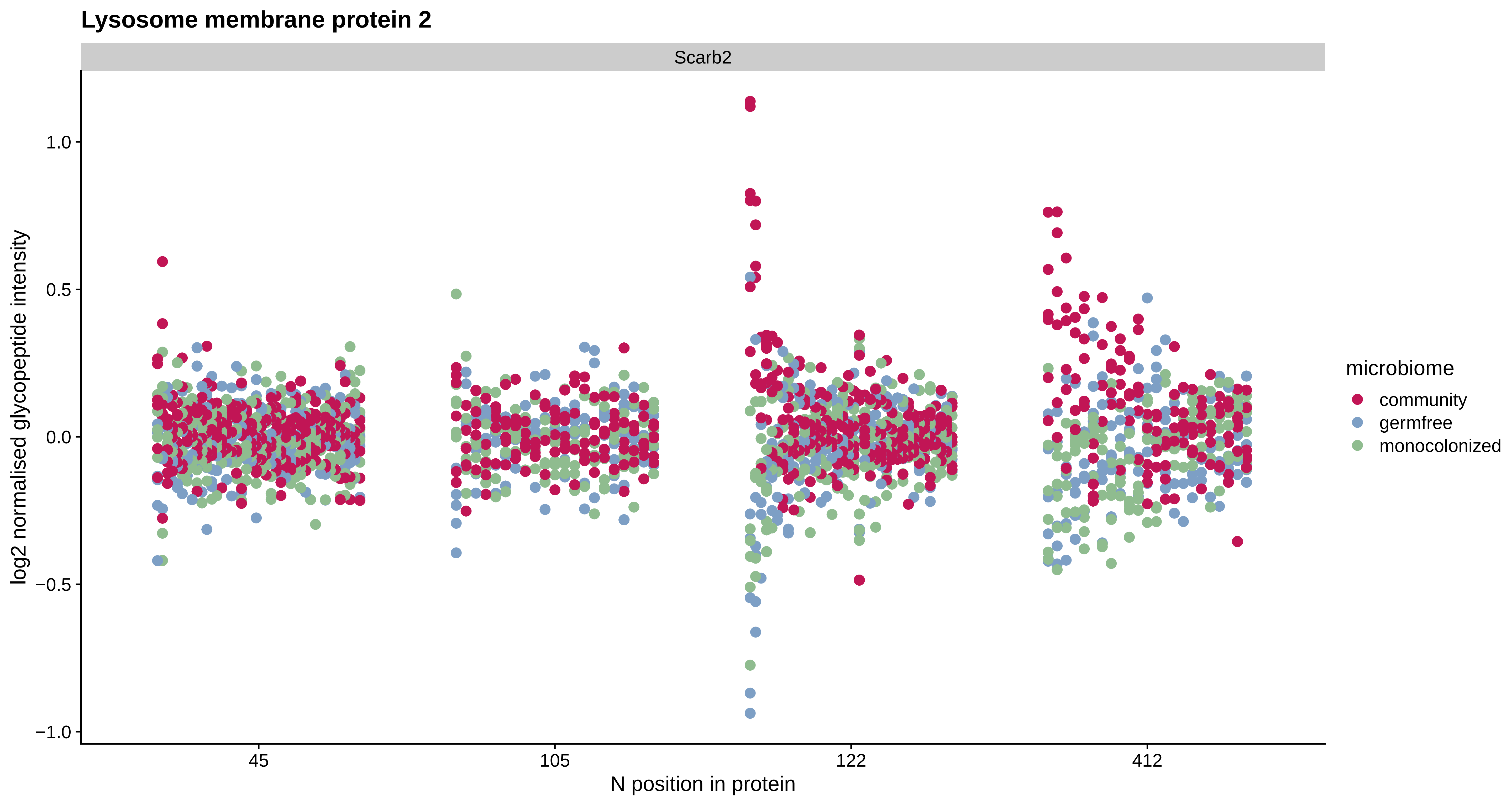
Lysosome membrane protein 2
Scarb2
log2 normalised glycopeptide intensity
1.0
0.5
0.0
−0.5
−1.0
45
105
122
412
N position in protein
microbiome
community
germfree
monocolonized

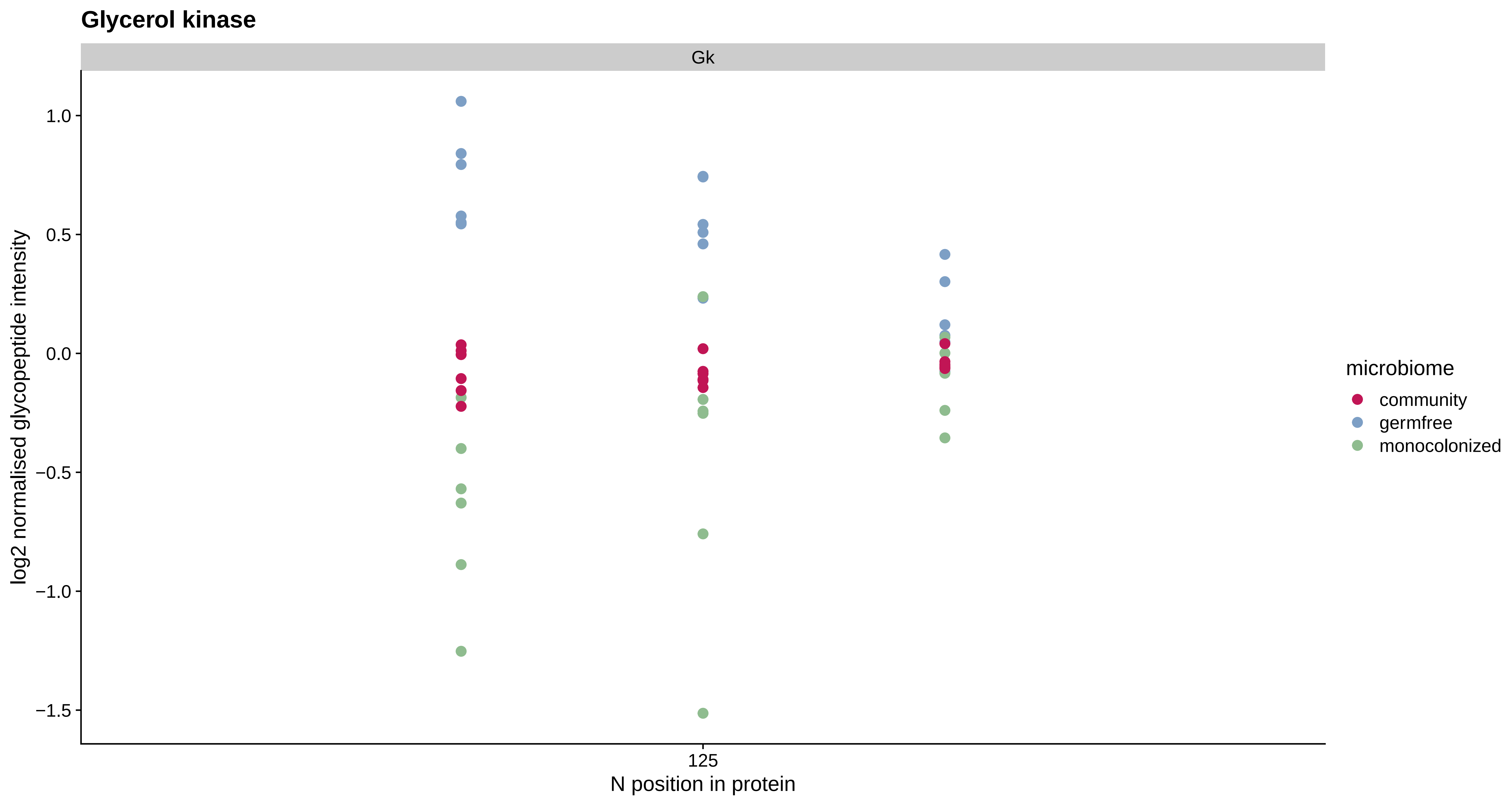

Glycerol kinase
Gk
1.0
0.5
0.0
−0.5
−1.0
−1.5
log2 normalised glycopeptide intensity
125
N position in protein
microbiome
community
germfree
monocolonized

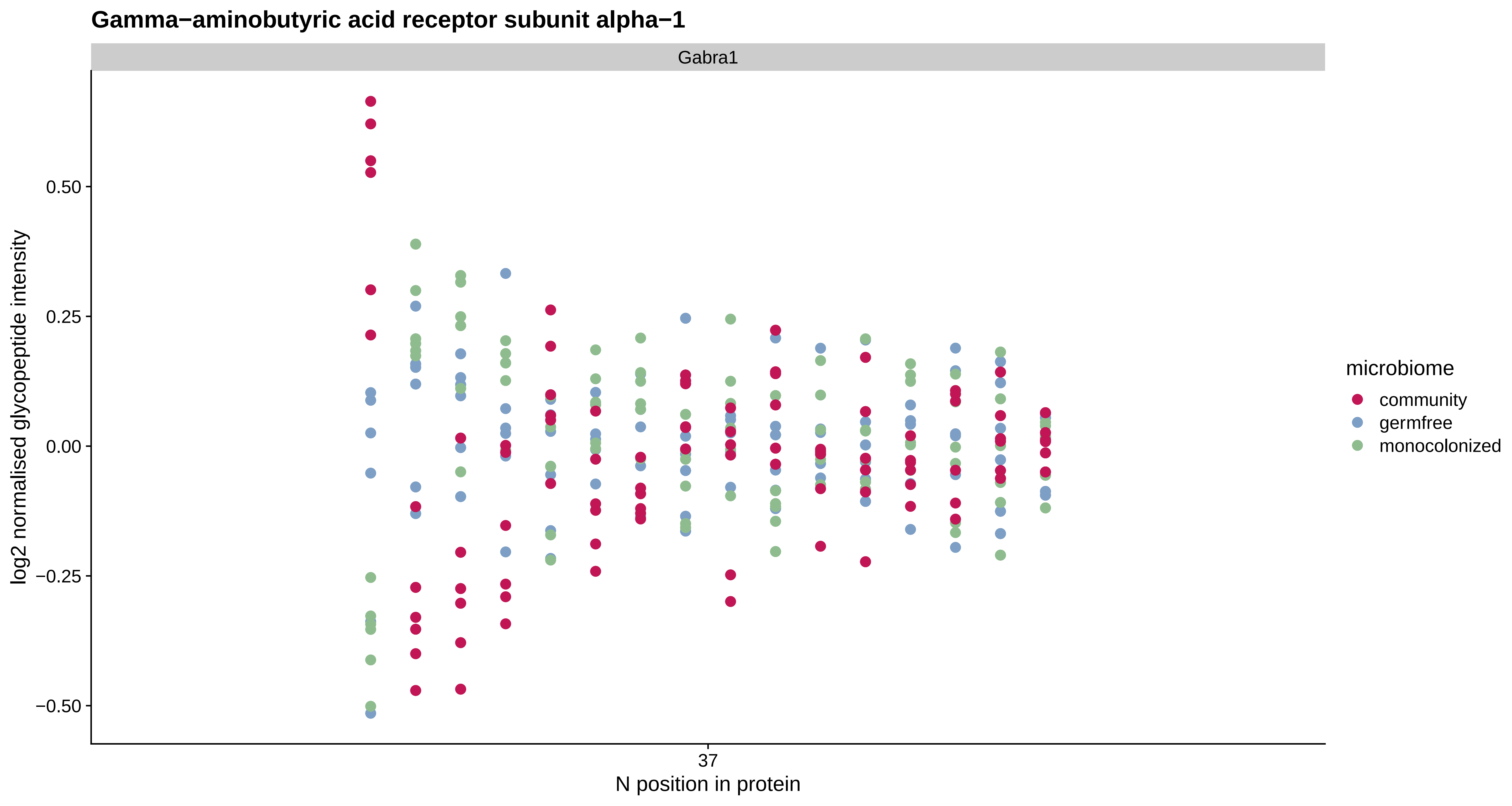

Gamma−aminobutyric acid receptor subunit alpha−1
Gabra1
0.50
0.25
0.00
−0.25
−0.50
log2 normalised glycopeptide intensity
37
N position in protein
microbiome
community
germfree
monocolonized

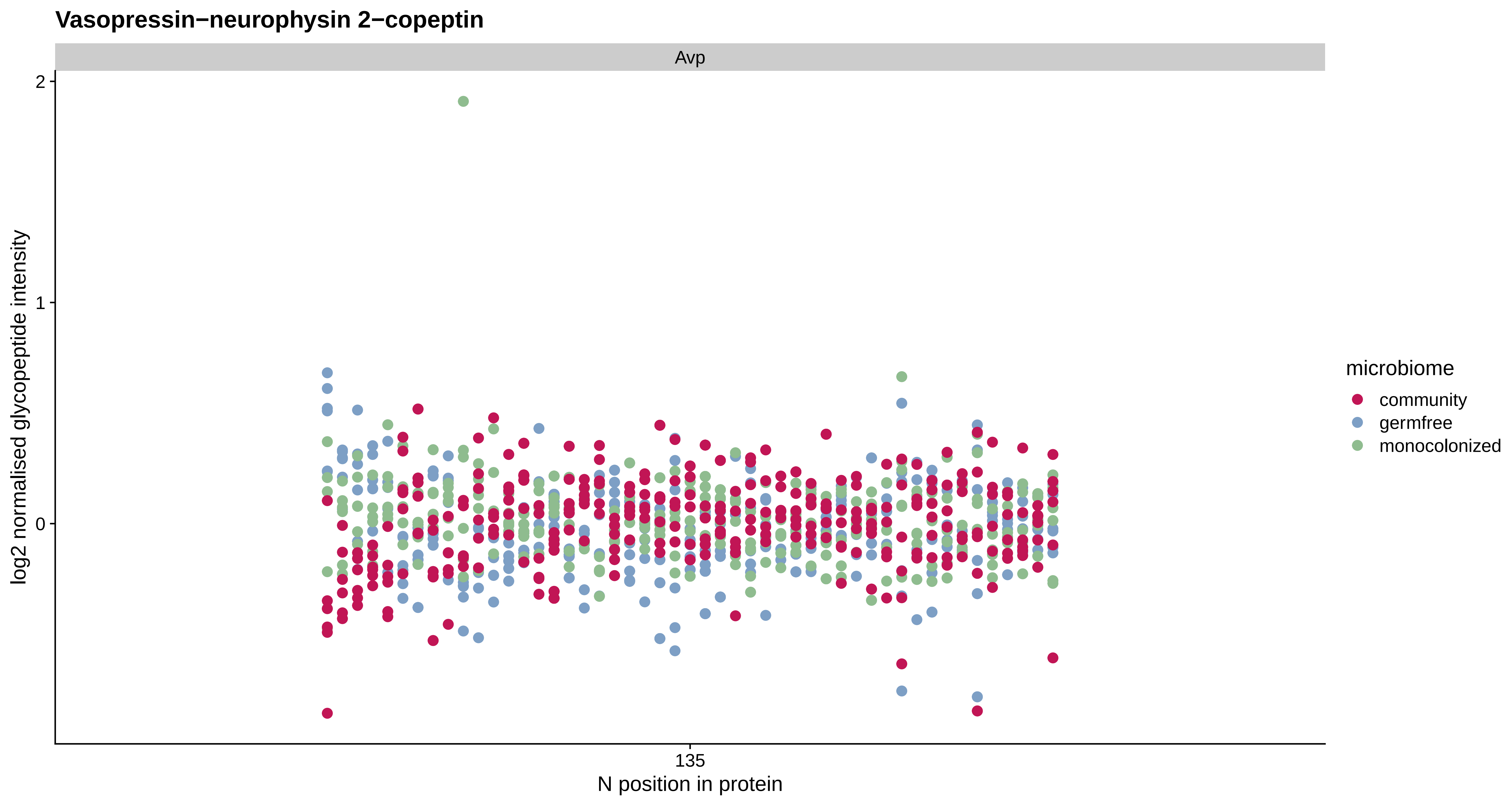

Vasopressin−neurophysin 2−copeptin
Avp
2
1
0
log2 normalised glycopeptide intensity
135
N position in protein
microbiome
community
germfree
monocolonized

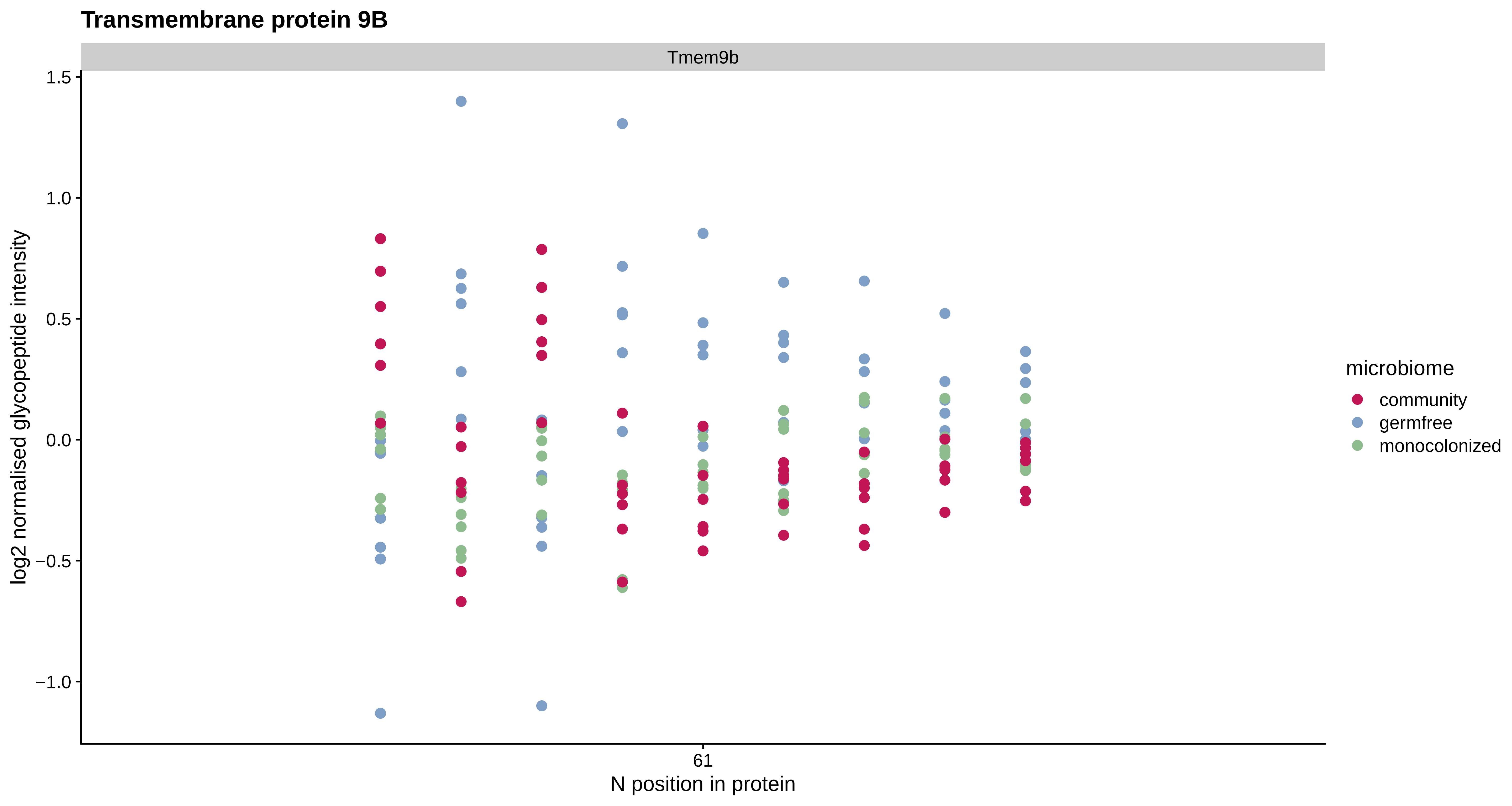
Transmembrane protein 9B
Tmem9b
1.5
1.0
0.5
0.0
−0.5
−1.0
log2 normalised glycopeptide intensity
61
N position in protein
microbiome
community
germfree
monocolonized

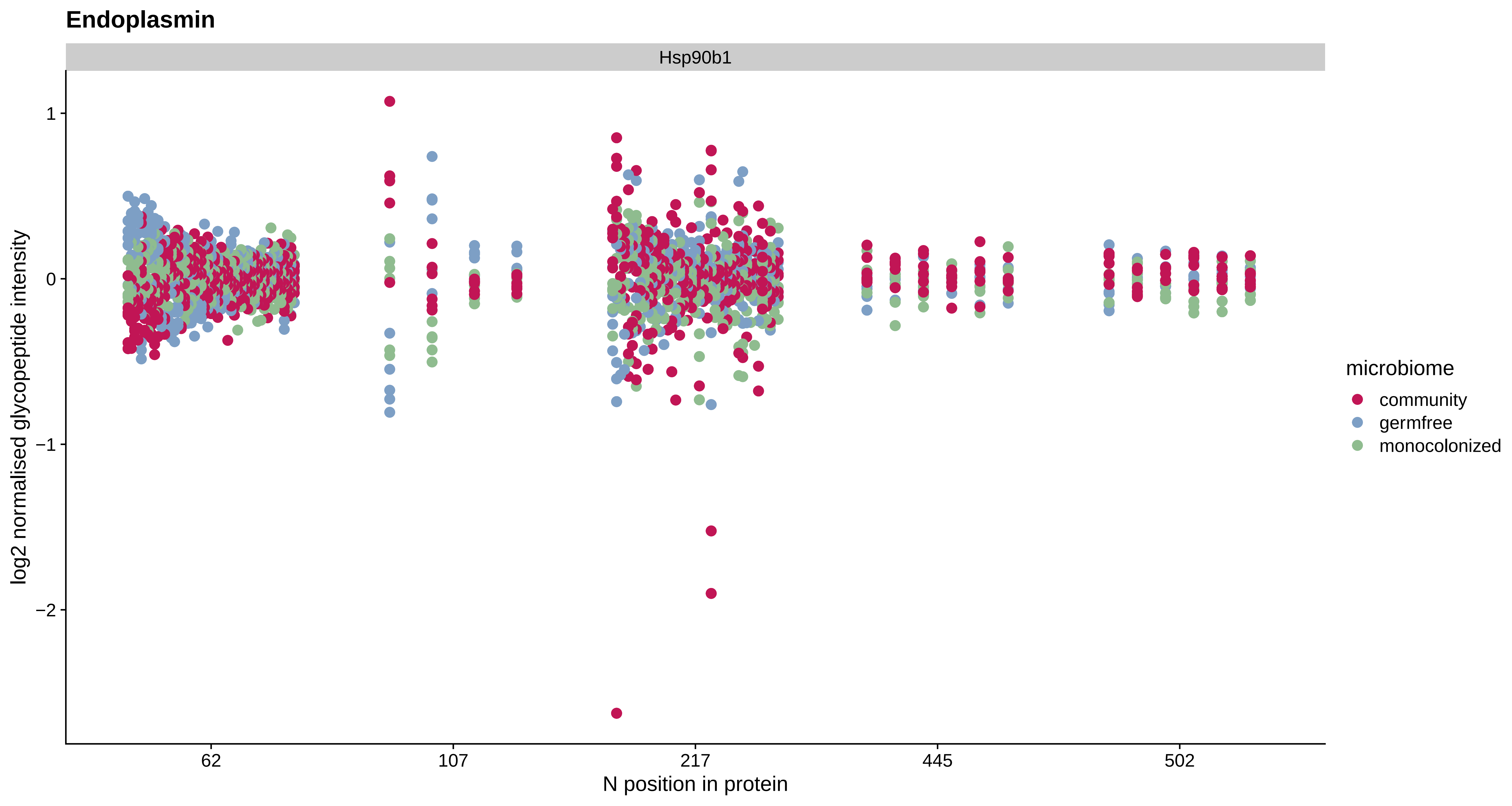
Endoplasmin
Hsp90b1
log2 normalised glycopeptide intensity
1
0
−1
−2
62
107
217
445
502
N position in protein
microbiome
community
germfree
monocolonized

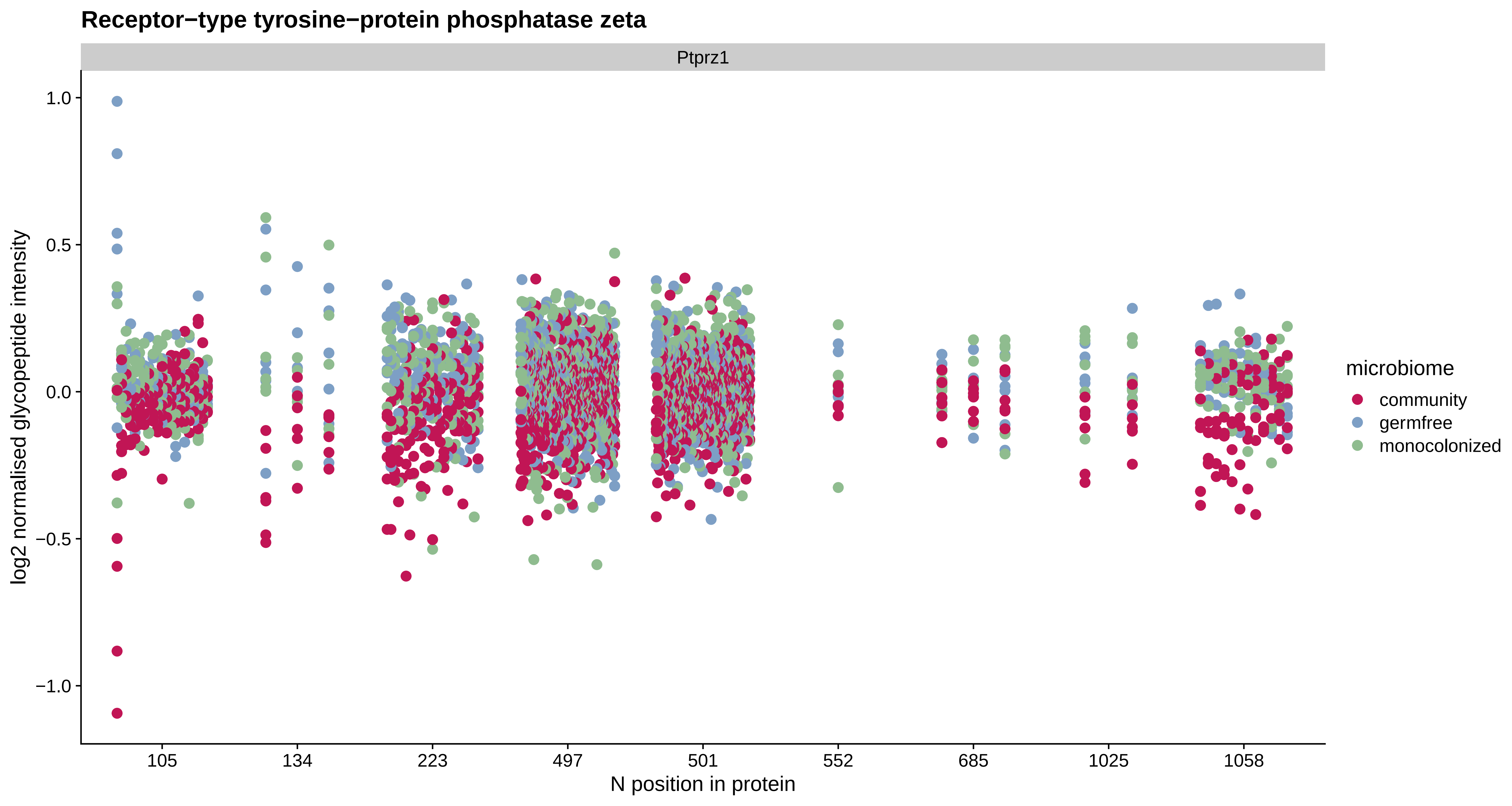
Receptor−type tyrosine−protein phosphatase zeta
Ptprz1
log2 normalised glycopeptide intensity
1.0
0.5
0.0
−0.5
−1.0
105
134
223
497
501
552
685
1025
1058
N position in protein
microbiome
community
germfree
monocolonized

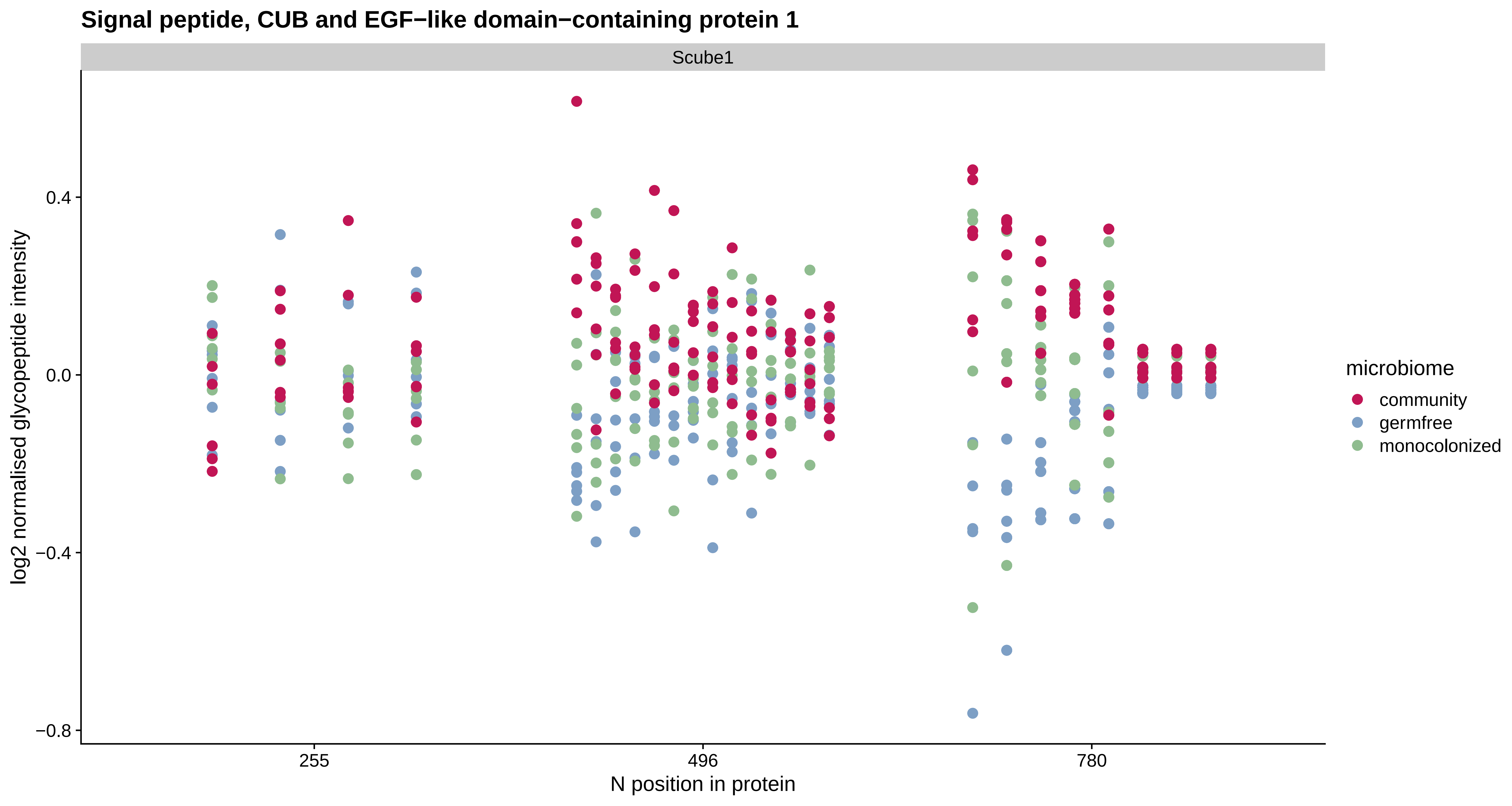

Signal peptide, CUB and EGF−like domain−containing protein 1
Scube1
log2 normalised glycopeptide intensity
0.4
0.0
−0.4
−0.8
255
496
780
N position in protein
microbiome
community
germfree
monocolonized

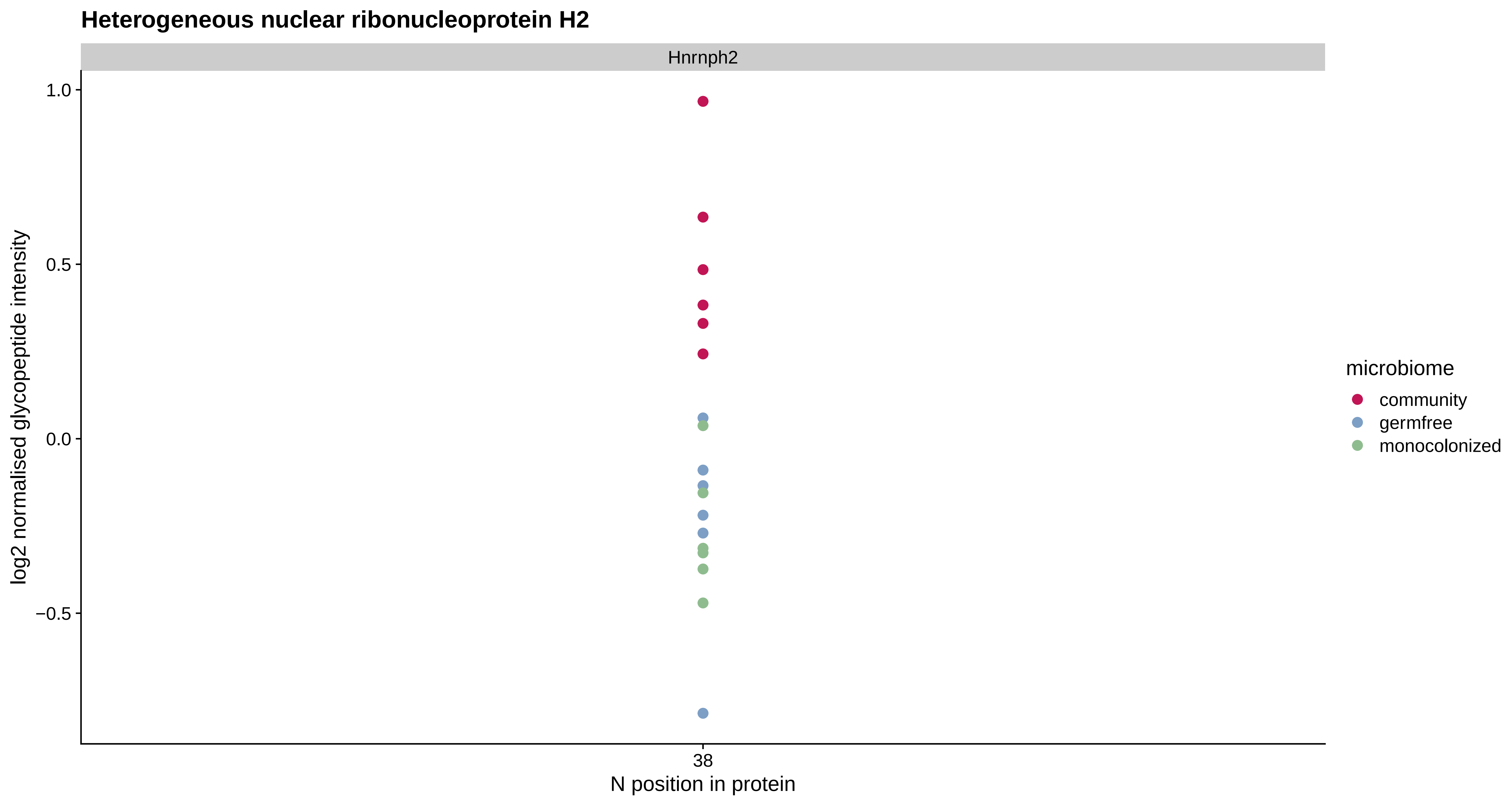
Heterogeneous nuclear ribonucleoprotein H2
Hnrnph2
1.0
0.5
0.0
−0.5
log2 normalised glycopeptide intensity
38
N position in protein
microbiome
community
germfree
monocolonized

### Major prion protein

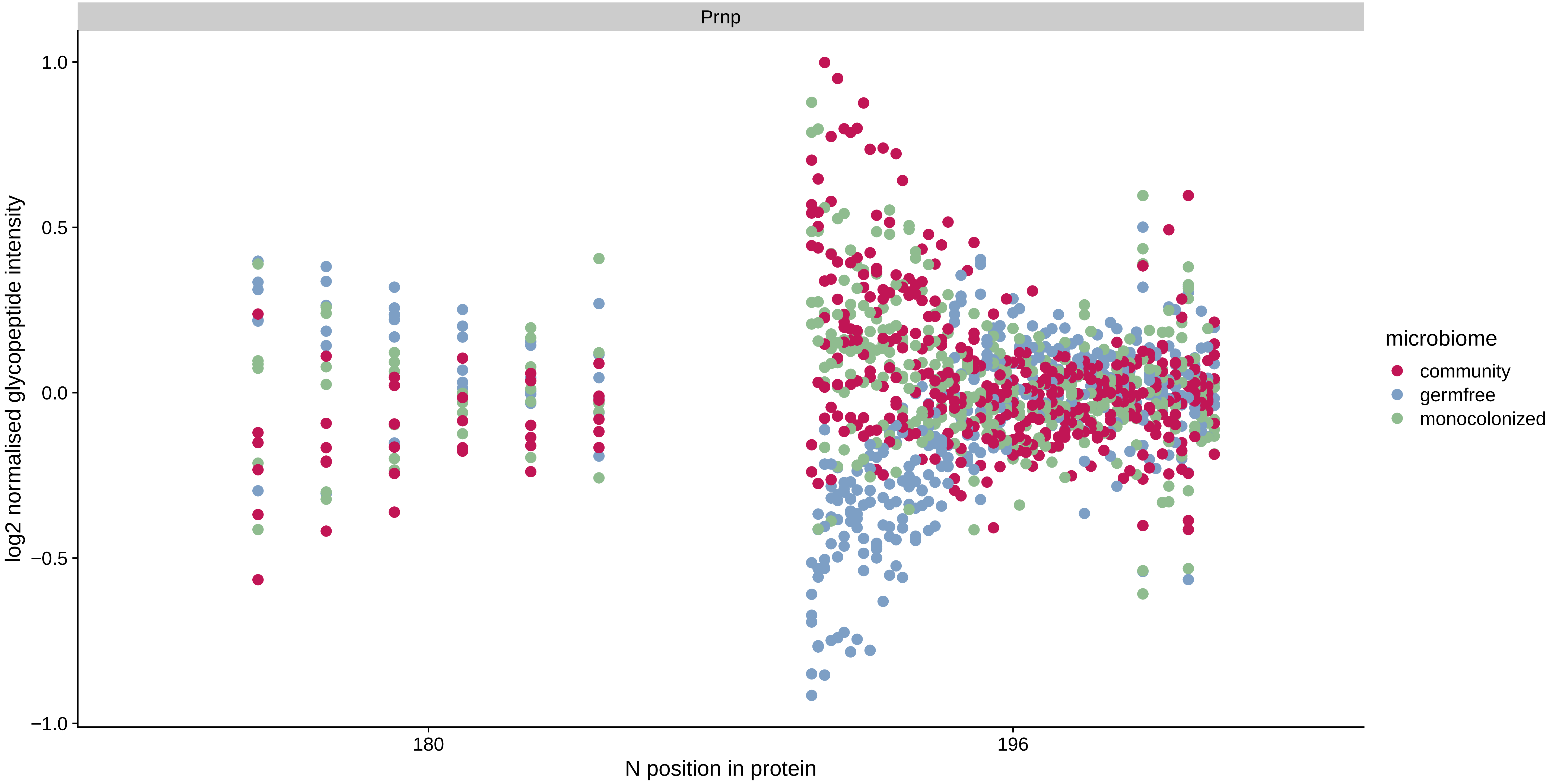

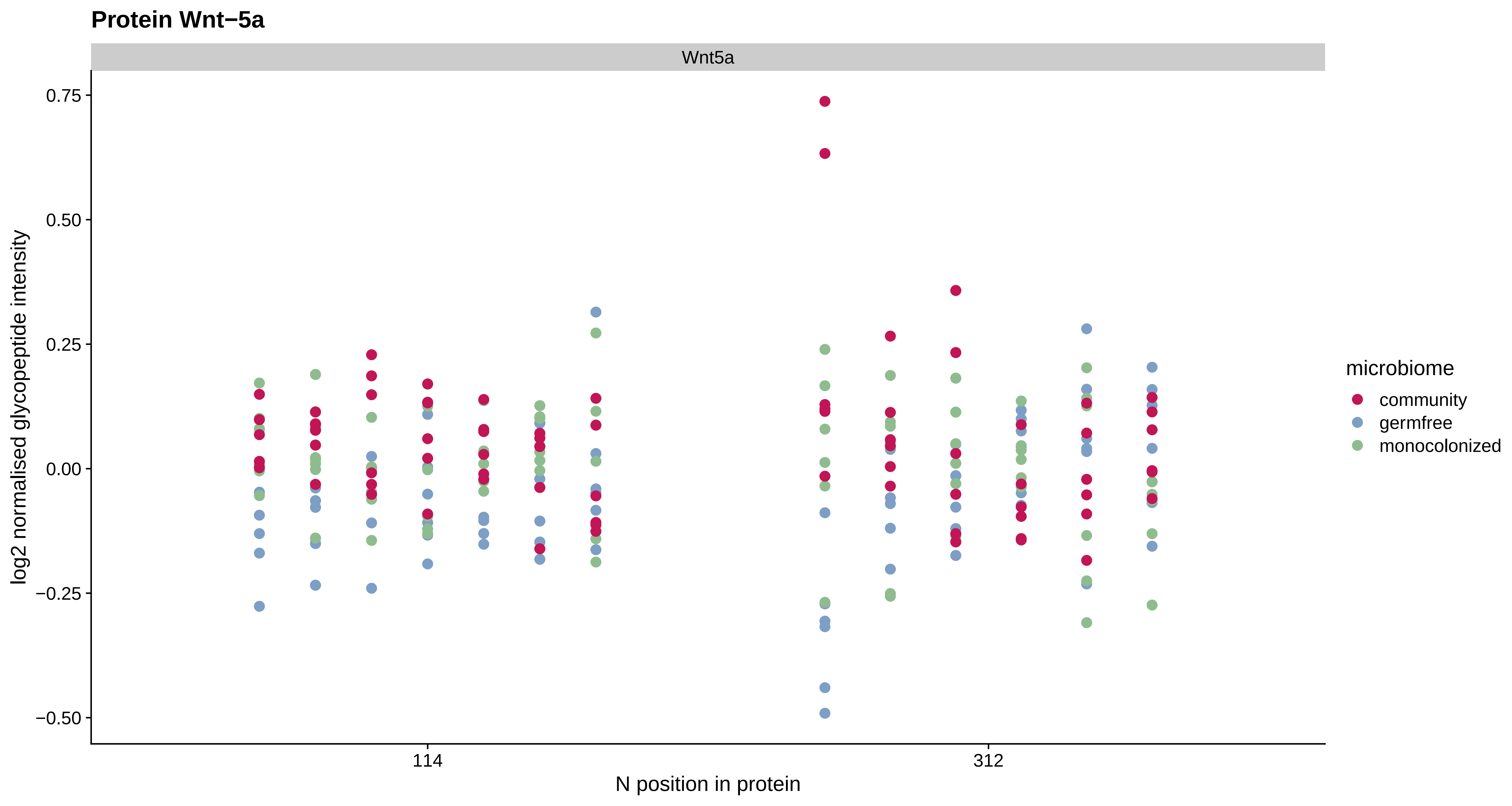
Protein Wnt−5a
Wnt5a
0.75
0.50
0.25
0.00
−0.25
−0.50
log2 normalised glycopeptide intensity
114
312
N position in protein
microbiome
community
germfree
monocolonized

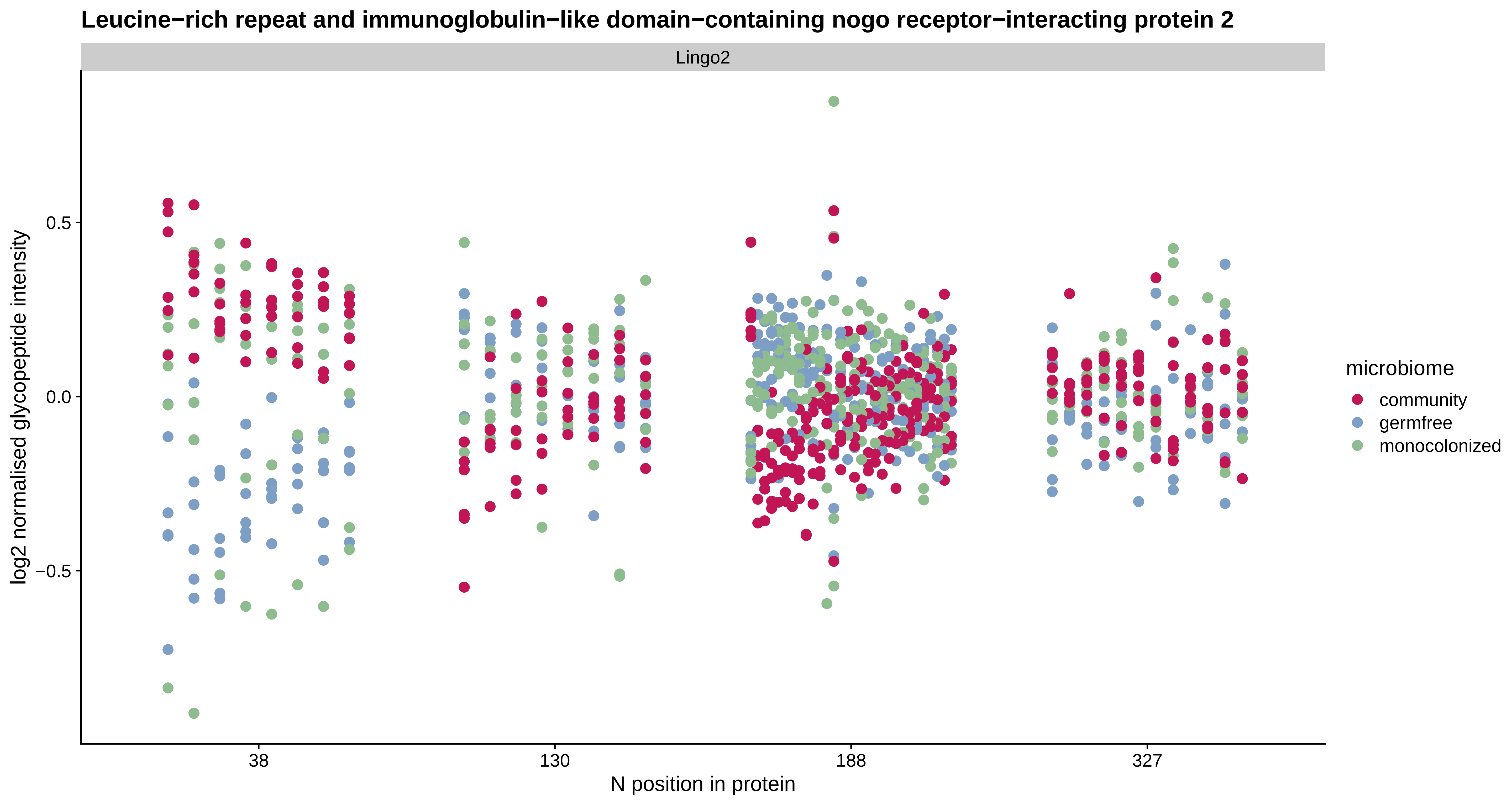

Leucine−rich repeat and immunoglobulin−like domain−containing nogo receptor−interacting protein 2
Lingo2
log2 normalised glycopeptide intensity
0.5
0.0
−0.5
38
130
188
327
N position in protein
microbiome
community
germfree
monocolonized

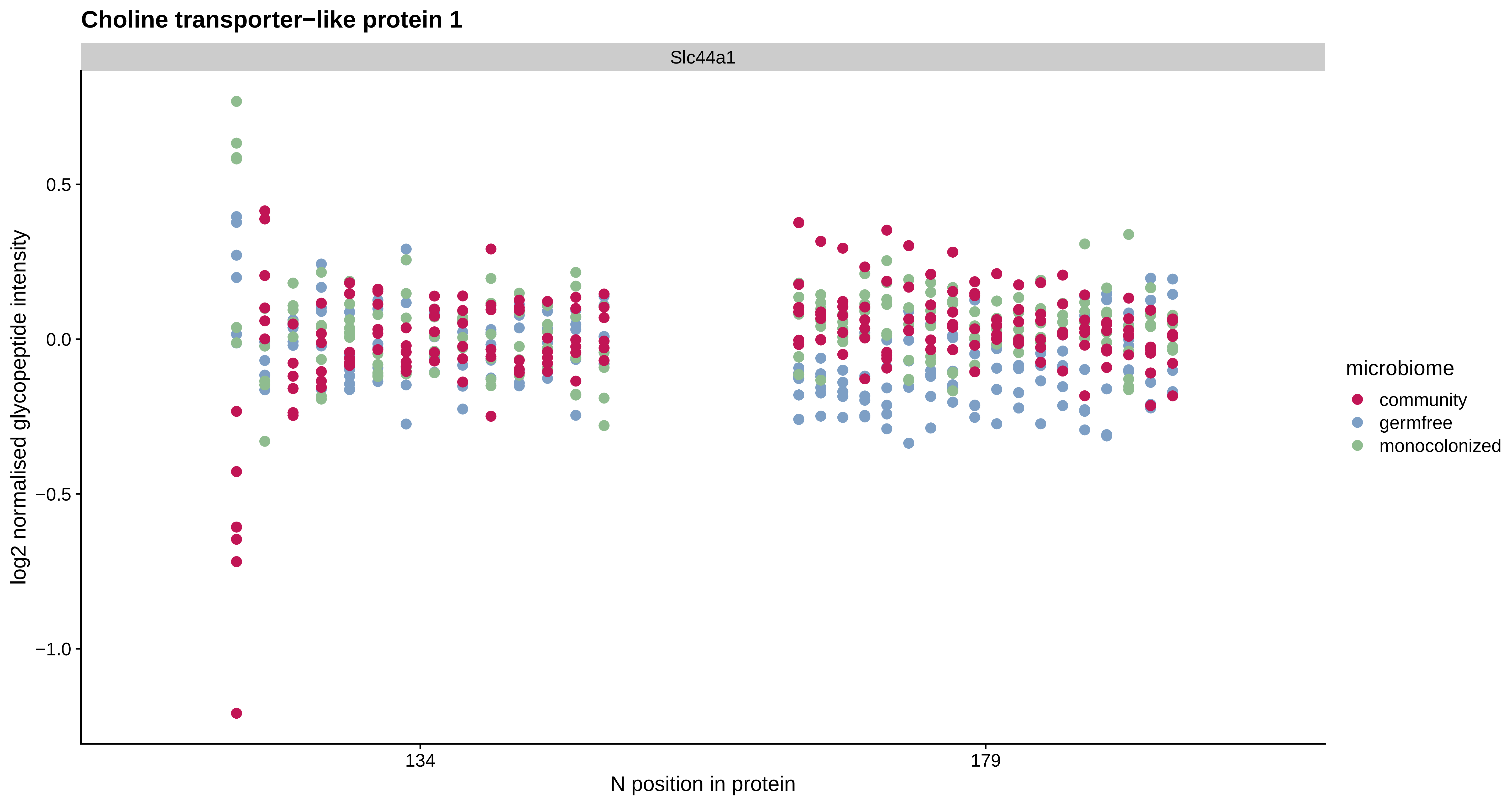
Choline transporter−like protein 1
Slc44a1
log2 normalised glycopeptide intensity
0.5
0.0
−0.5
−1.0
134
179
N position in protein
microbiome
community
germfree
monocolonized

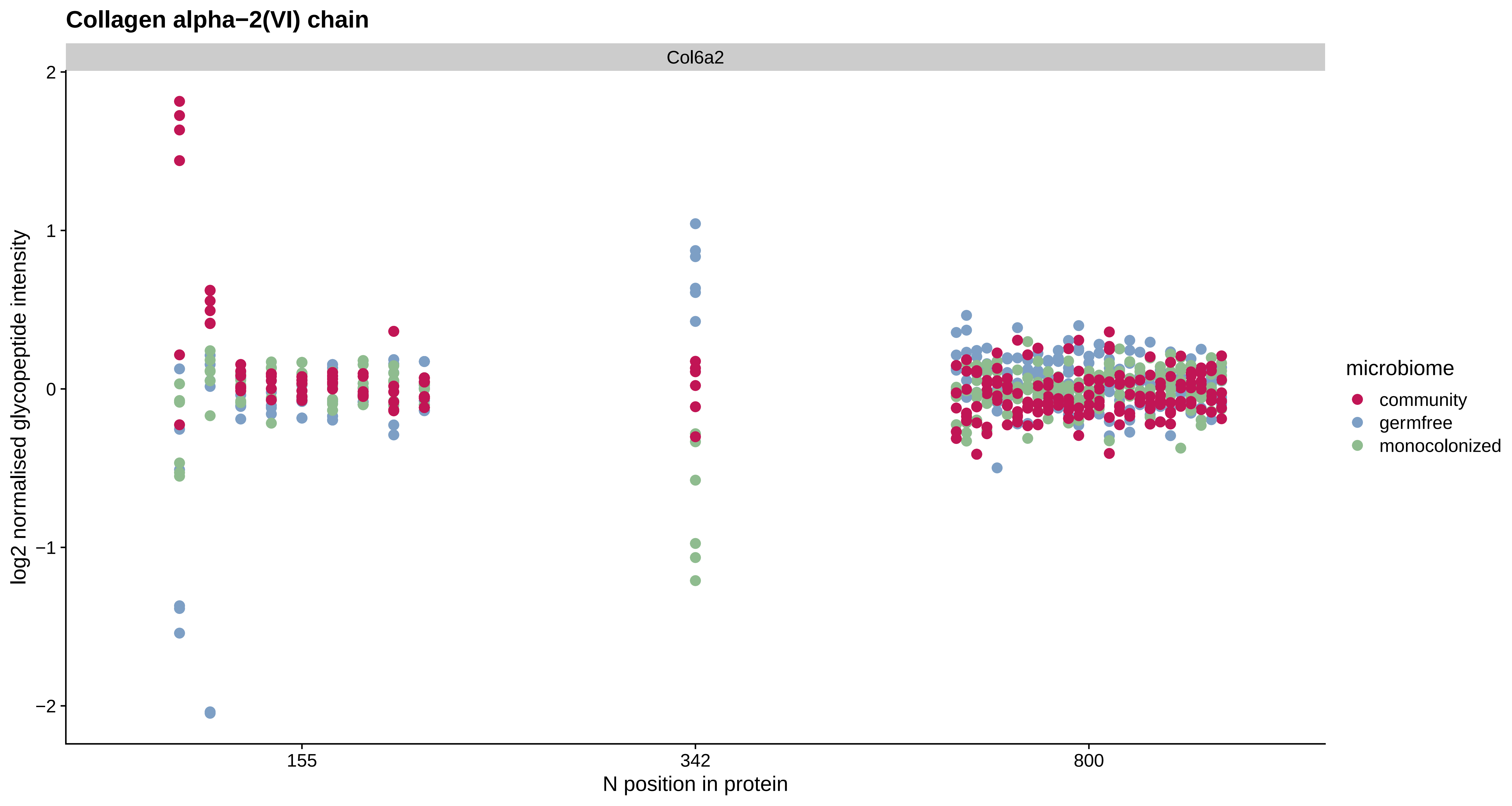

Collagen alpha−2(VI) chain
Col6a2
log2 normalised glycopeptide intensity
2
1
0
−1
−2
155
342
800
N position in protein
microbiome
community
germfree
monocolonized

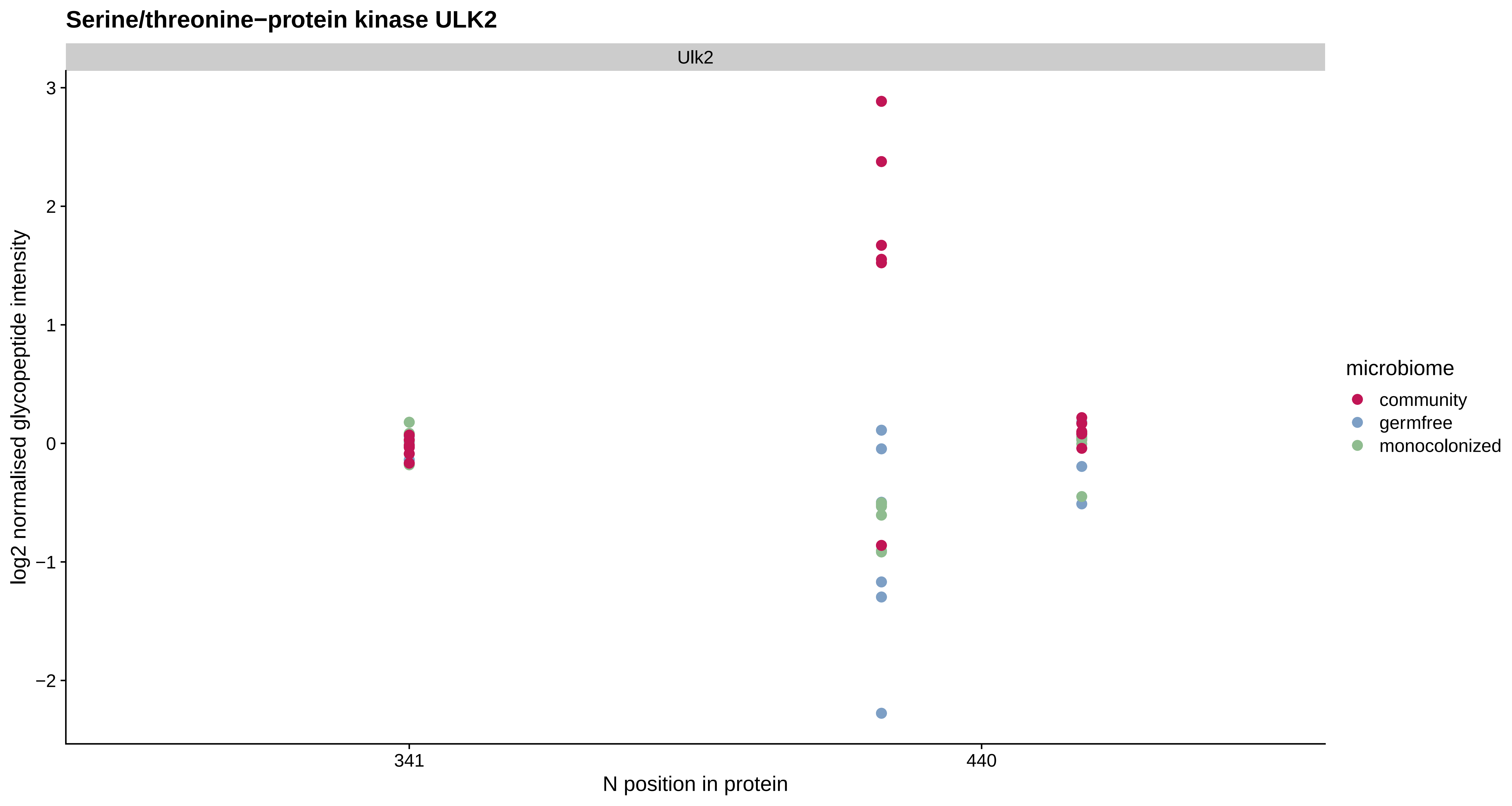
Serine/threonine−protein kinase ULK2
Ulk2
log2 normalised glycopeptide intensity
3
2
1
0
−1
−2
341
440
N position in protein
microbiome
community
germfree
monocolonized

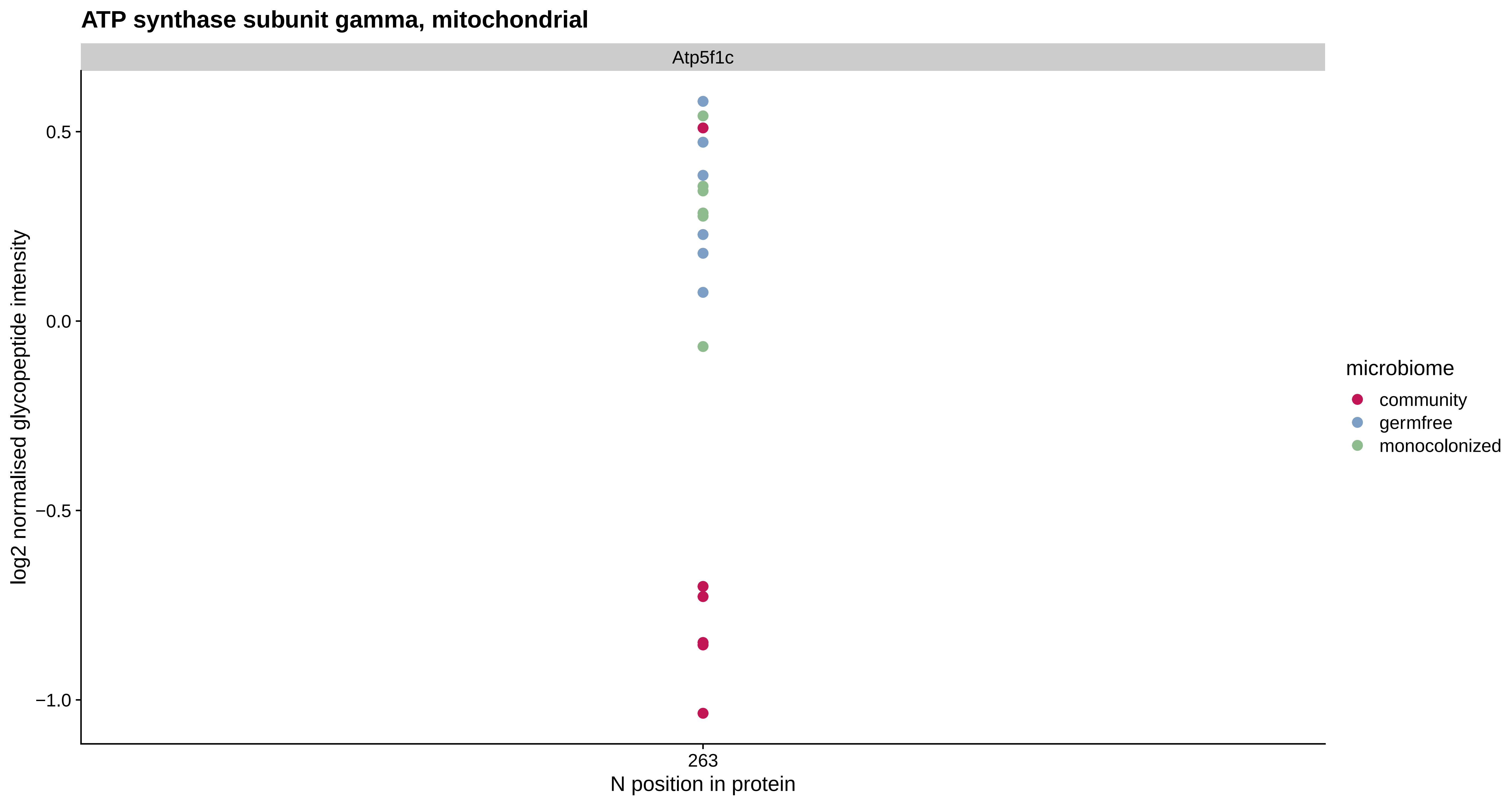

ATP synthase subunit gamma, mitochondrial
Atp5f1c
0.5
0.0
−0.5
−1.0
log2 normalised glycopeptide intensity
263
N position in protein
microbiome
community
germfree
monocolonized

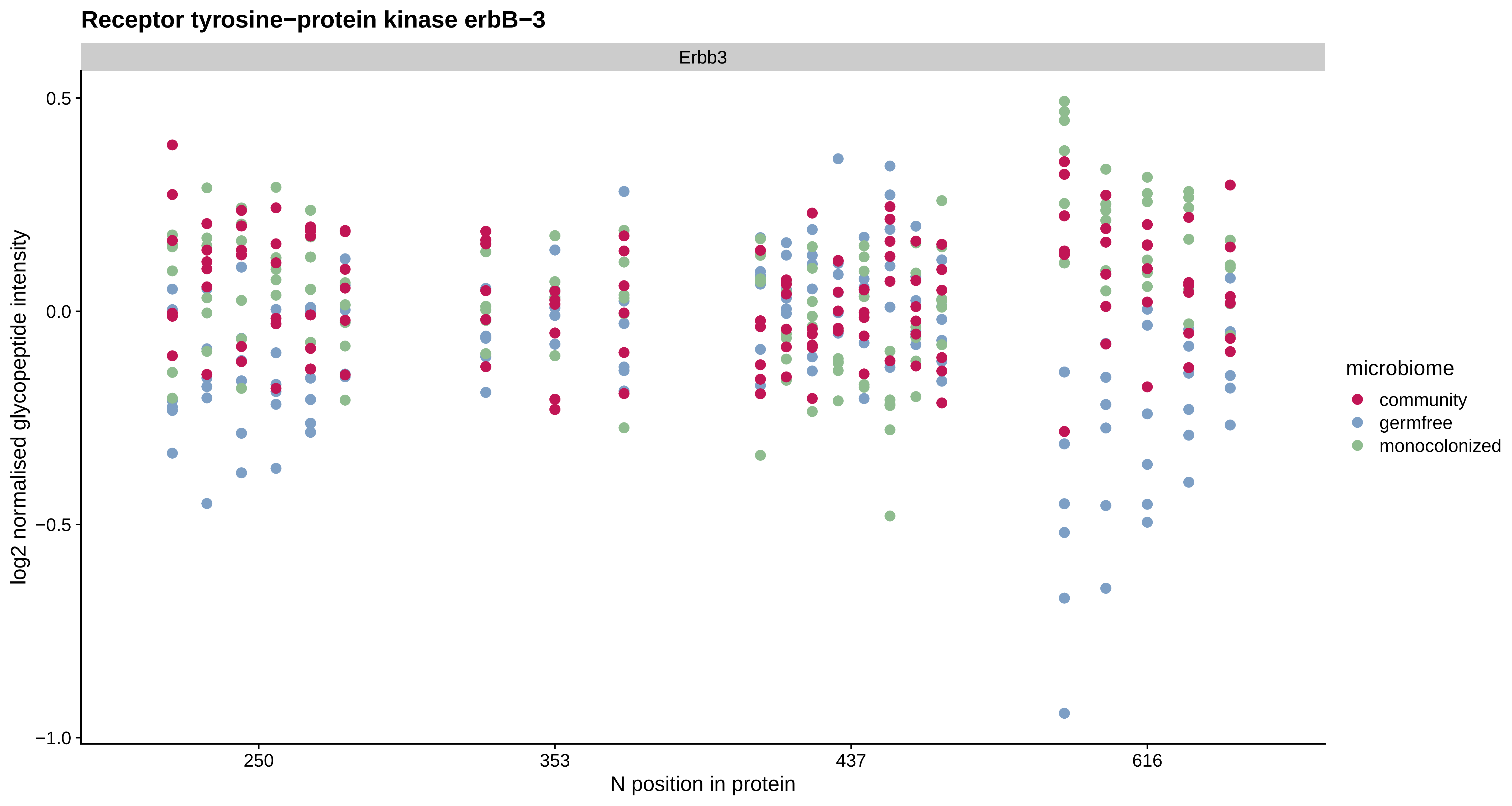

Receptor tyrosine−protein kinase erbB−3
Erbb3
0.5
0.0
−0.5
−1.0
log2 normalised glycopeptide intensity
250
353
437
616
N position in protein
microbiome
community
germfree
monocolonized

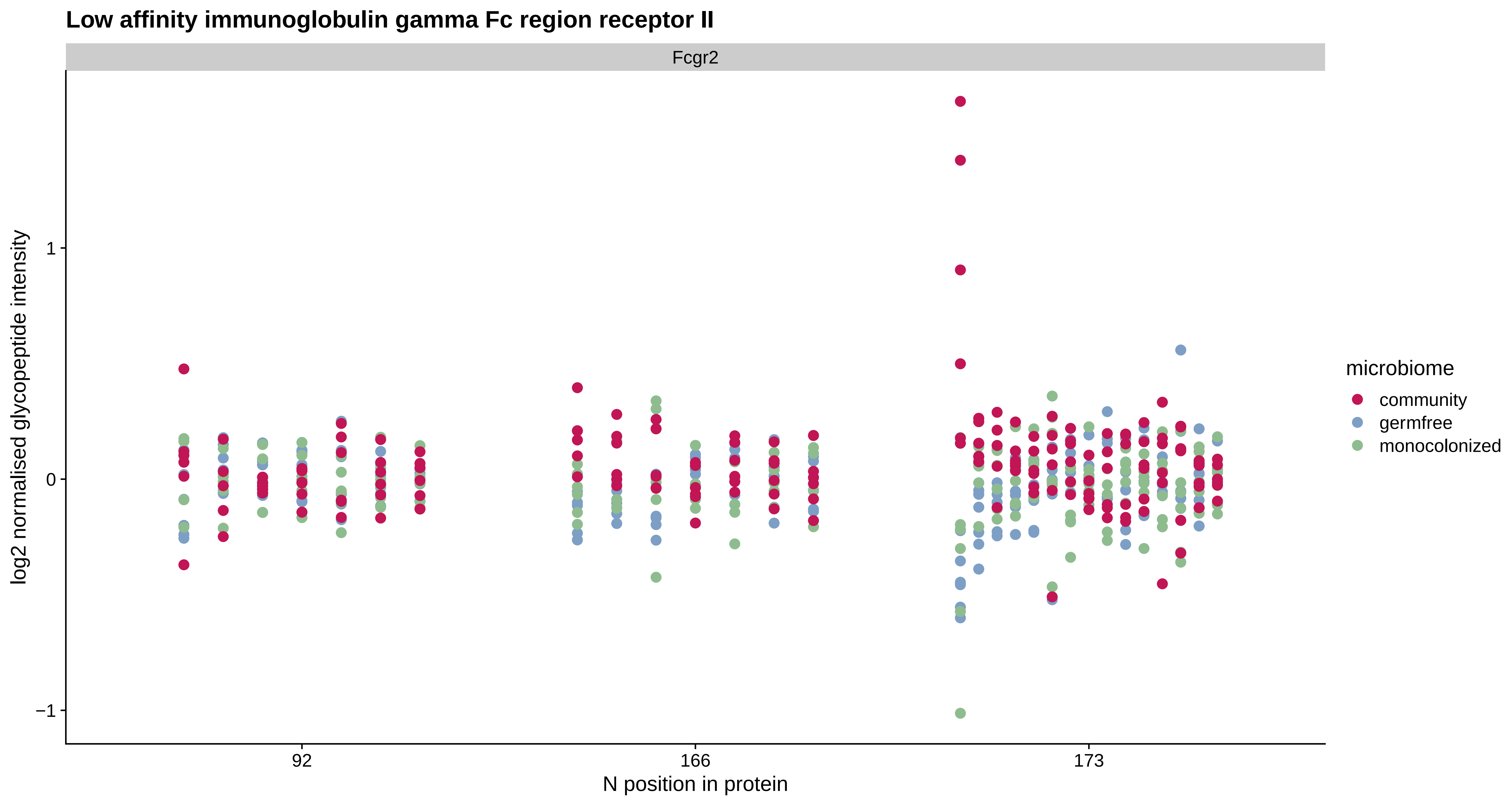
Low affinity immunoglobulin gamma Fc region receptor II
Fcgr2
log2 normalised glycopeptide intensity
1
0
−1
92
166
173
N position in protein
microbiome
community
germfree
monocolonized

#### ATP−binding cassette sub−family A member 2

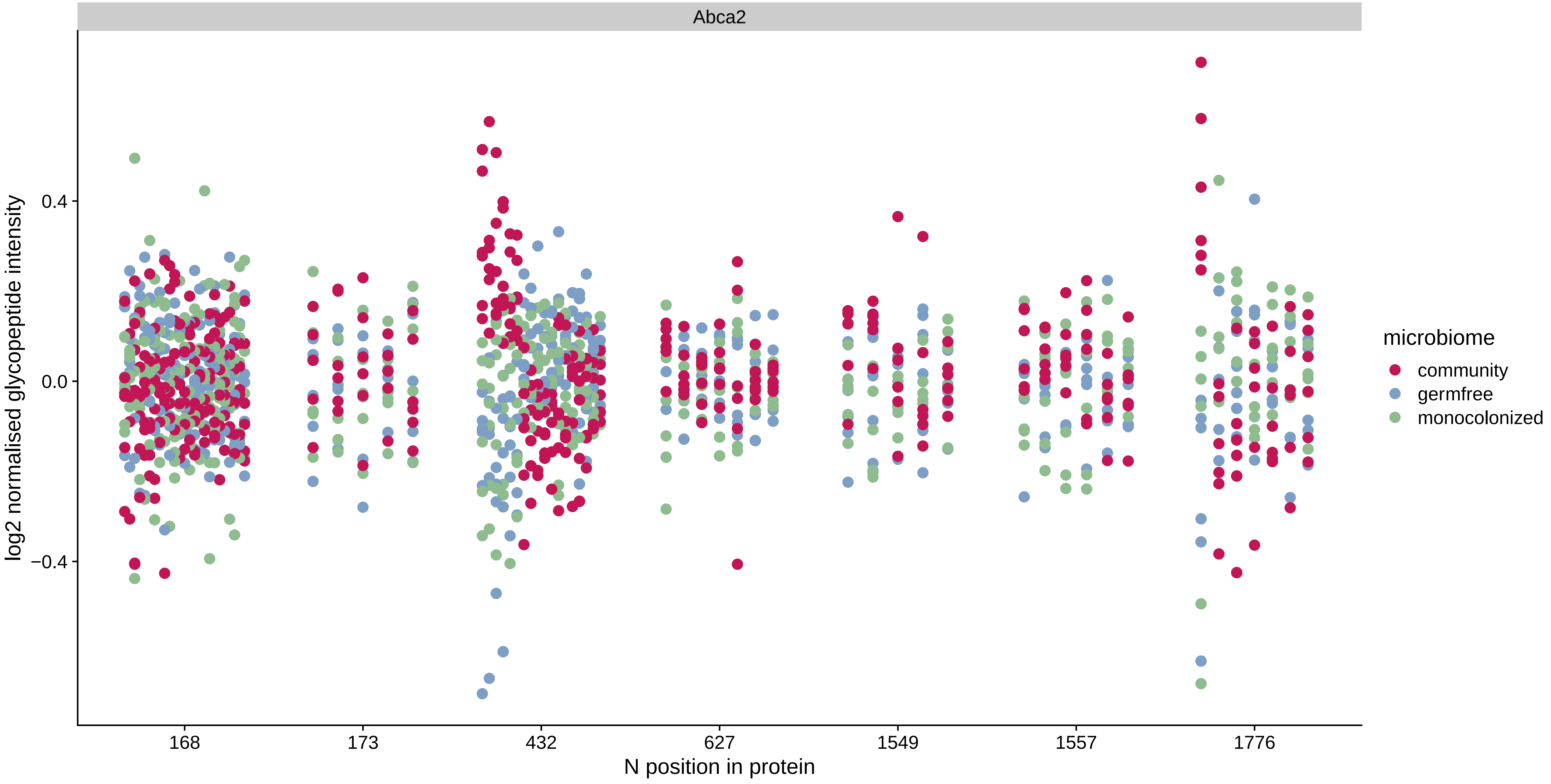

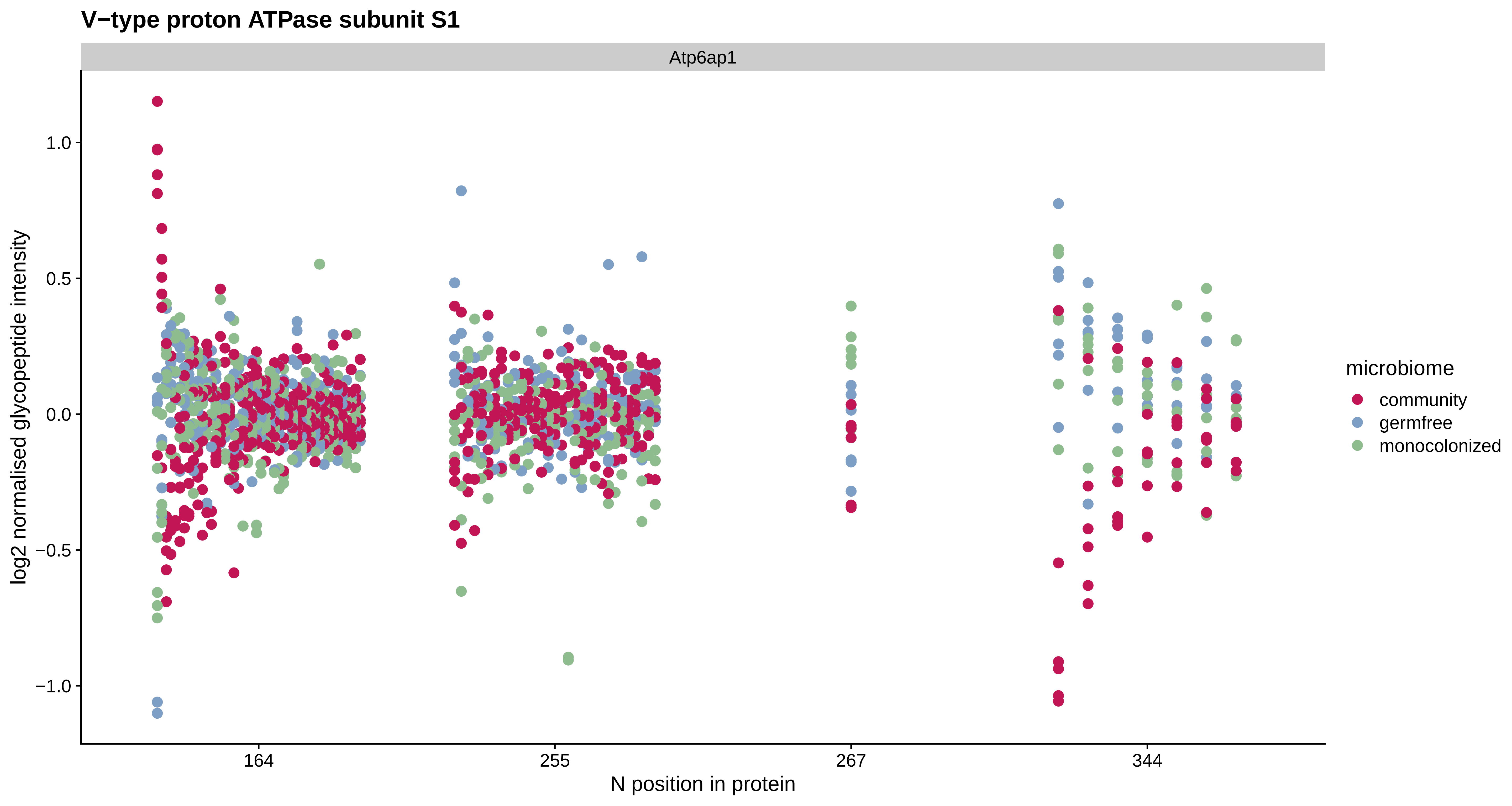

V−type proton ATPase subunit S1
Atp6ap1
log2 normalised glycopeptide intensity
1.0
0.5
0.0
−0.5
−1.0
164
255
267
344
N position in protein
microbiome
community
germfree
monocolonized

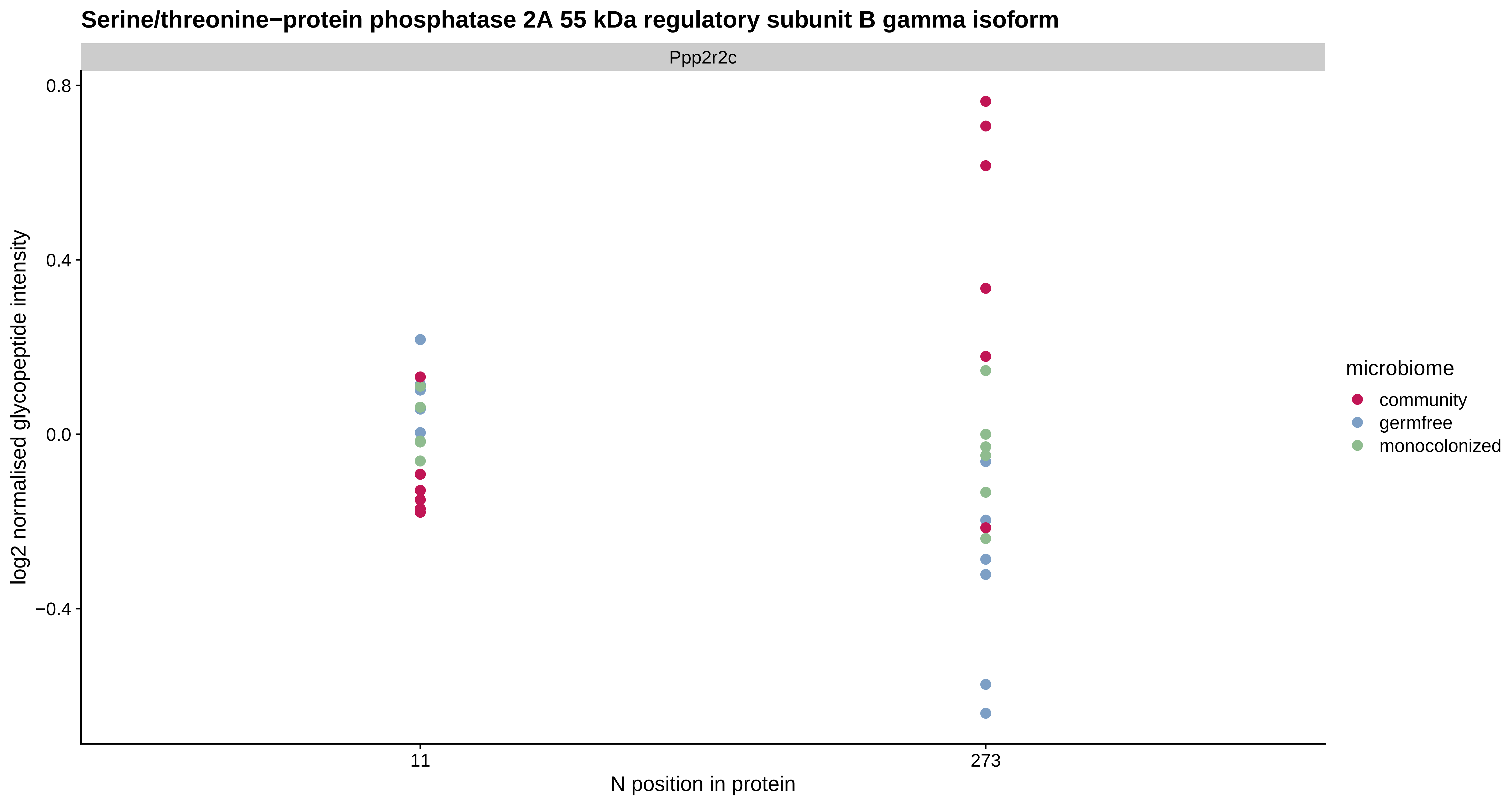

Serine/threonine−protein phosphatase 2A 55 kDa regulatory subunit B gamma isoform
Ppp2r2c
0.8
0.4
0.0
−0.4
log2 normalised glycopeptide intensity
11
273
N position in protein
microbiome
community
germfree
monocolonized

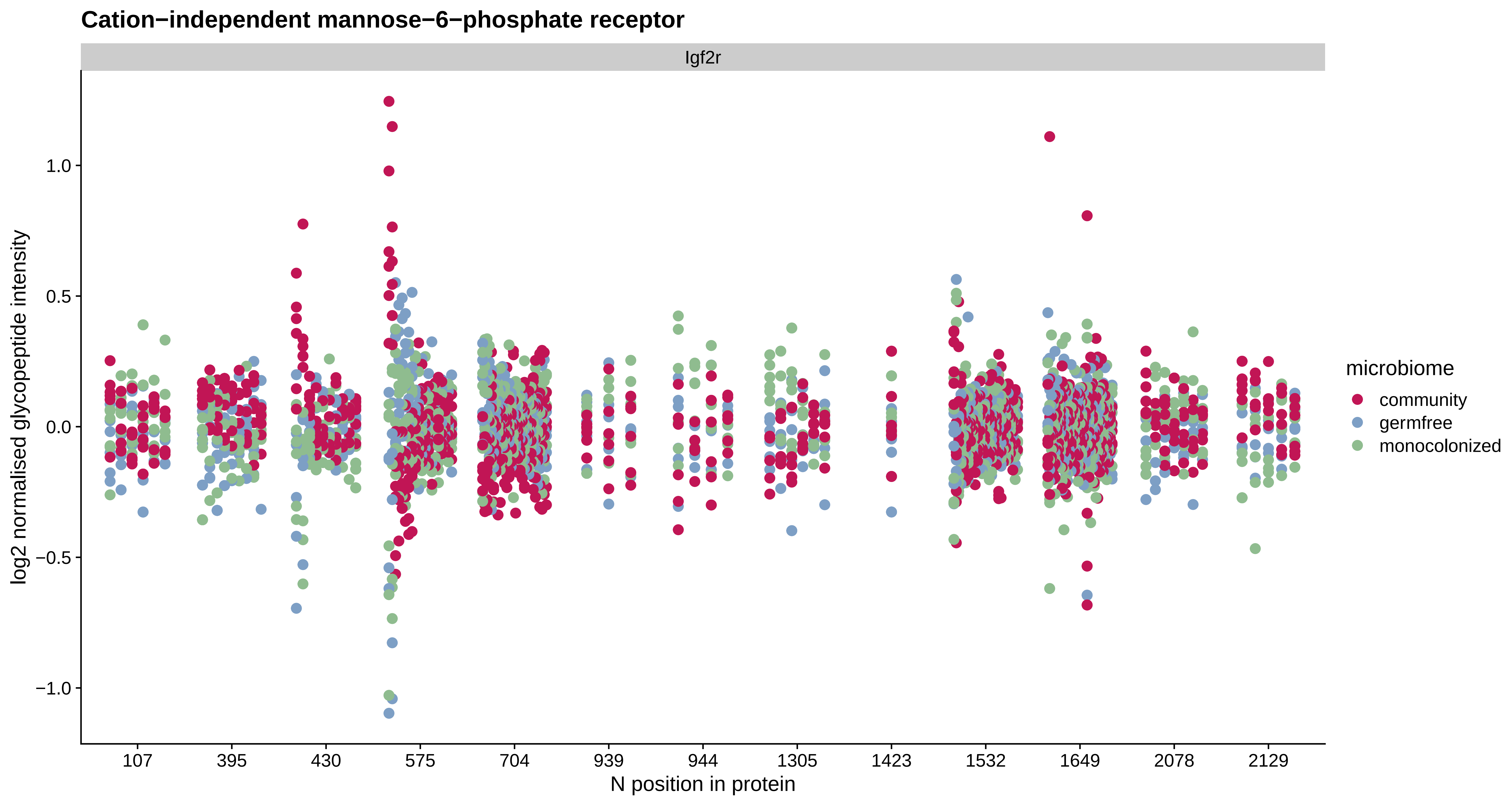

Cation−independent mannose−6−phosphate receptor
Igf2r
log2 normalised glycopeptide intensity
1.0
0.5
0.0
−0.5
−1.0
107
395
430
575
704
939
944
1305
1423
1532
1649
2078
2129
N position in protein
microbiome
community
germfree
monocolonized

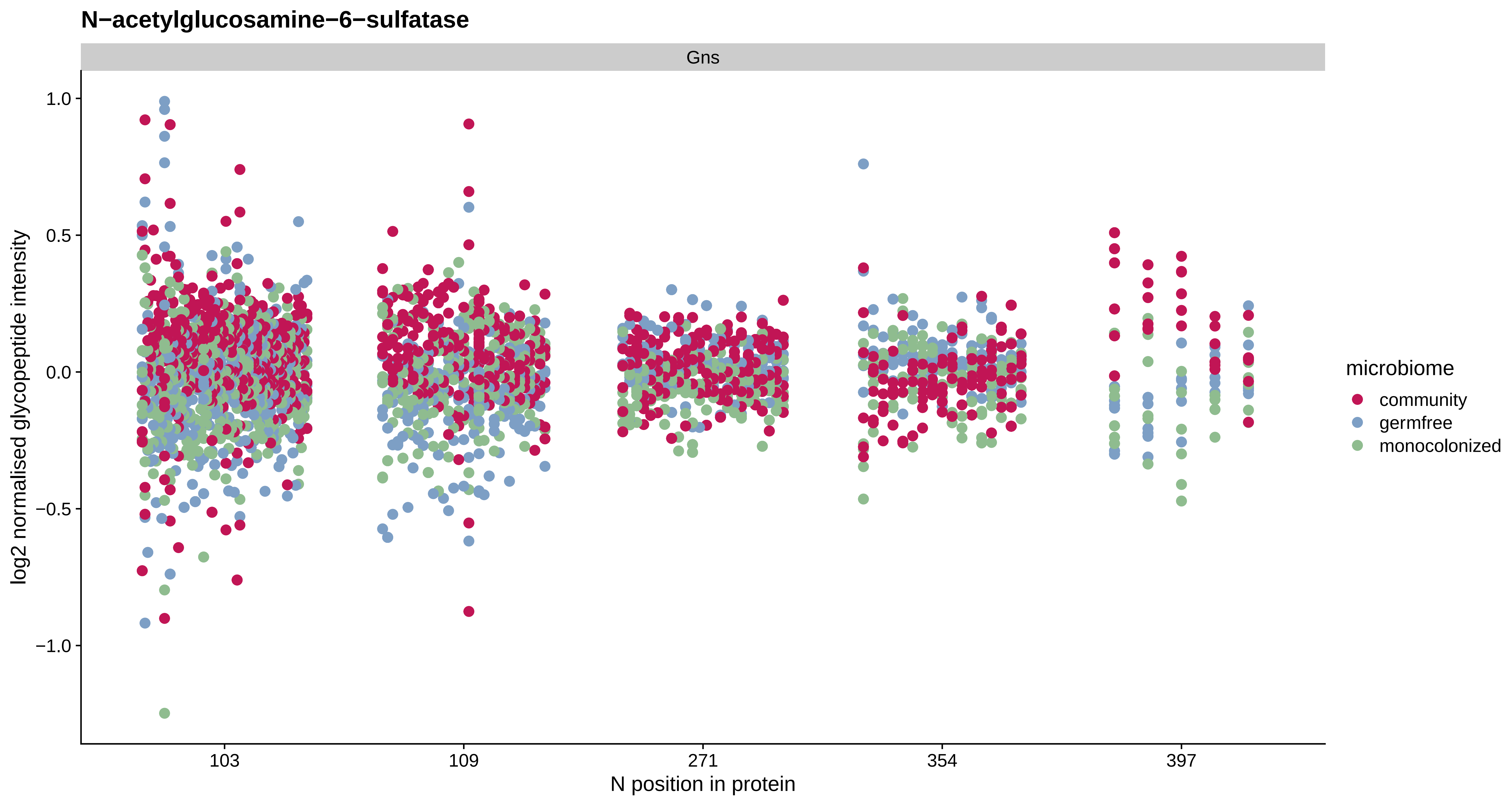
N−acetylglucosamine−6−sulfatase
Gns
1.0
0.5
0.0
−0.5
−1.0
log2 normalised glycopeptide intensity
103
109
271
354
397
N position in protein
microbiome
community
germfree
monocolonized

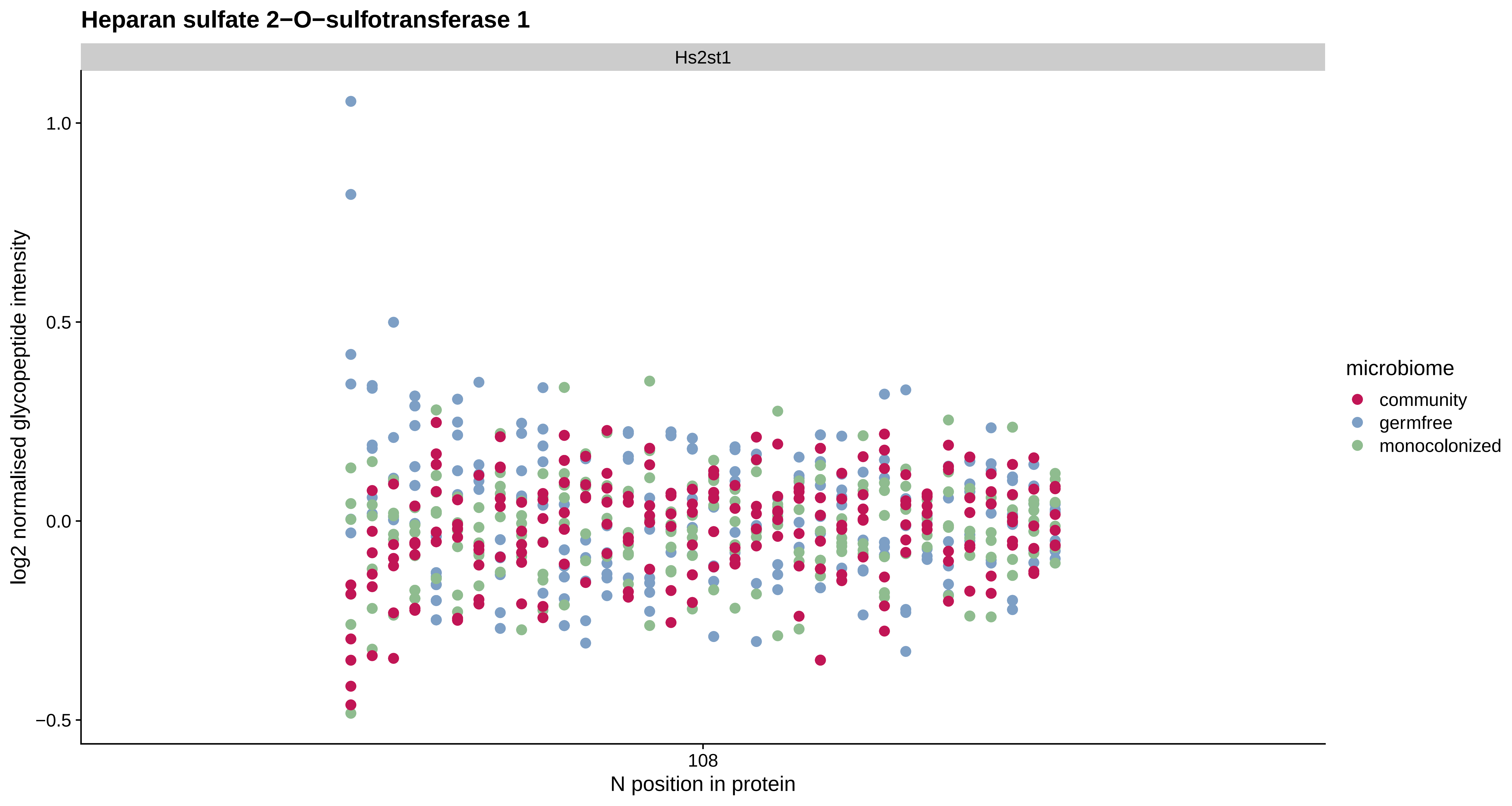

Heparan sulfate 2−O−sulfotransferase 1
Hs2st1
log2 normalised glycopeptide intensity
1.0
0.5
0.0
−0.5
108
N position in protein
microbiome
community
germfree
monocolonized

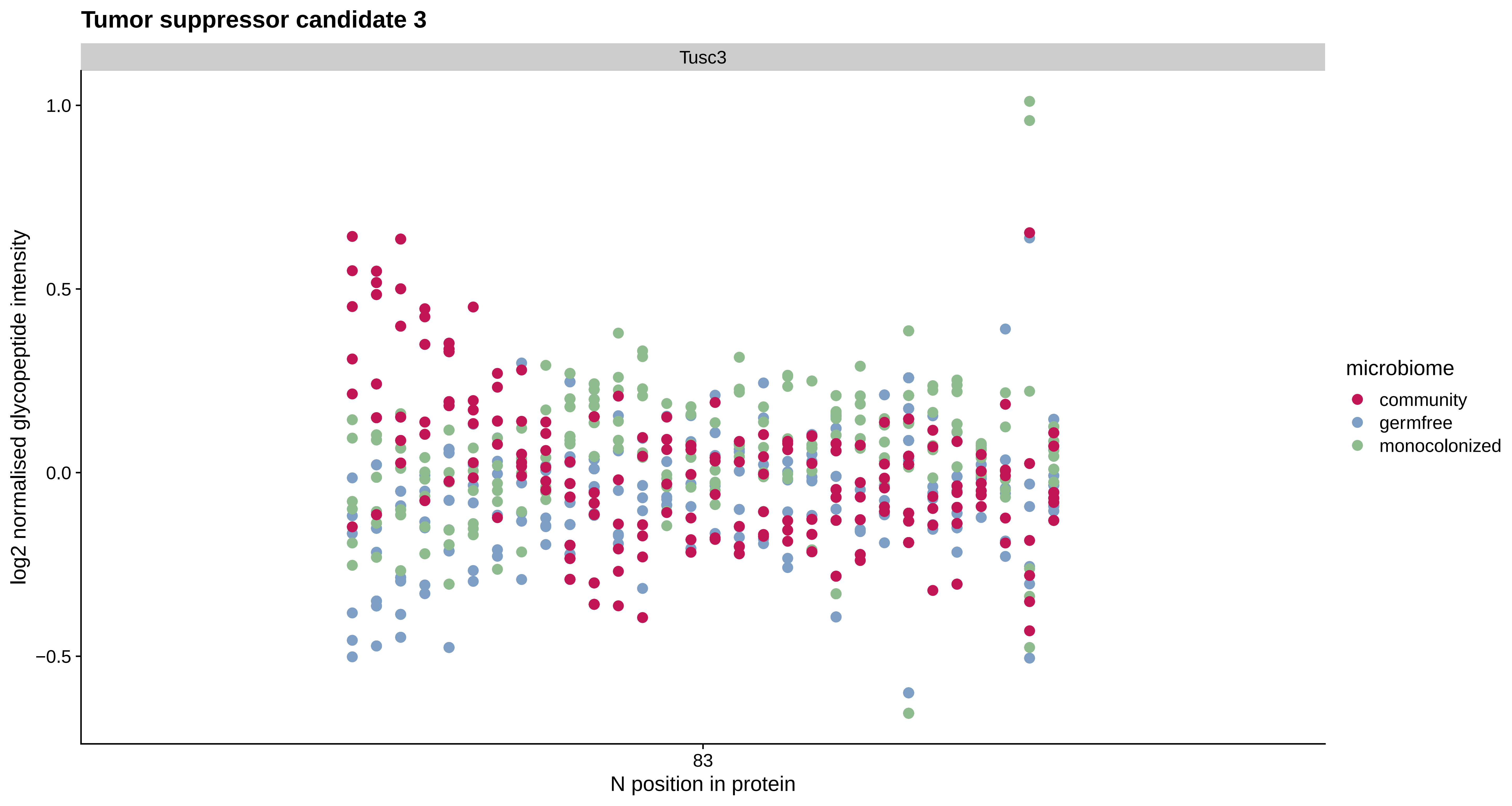

Tumor suppressor candidate 3
Tusc3
1.0
0.5
0.0
−0.5
log2 normalised glycopeptide intensity
83
N position in protein
microbiome
community
germfree
monocolonized

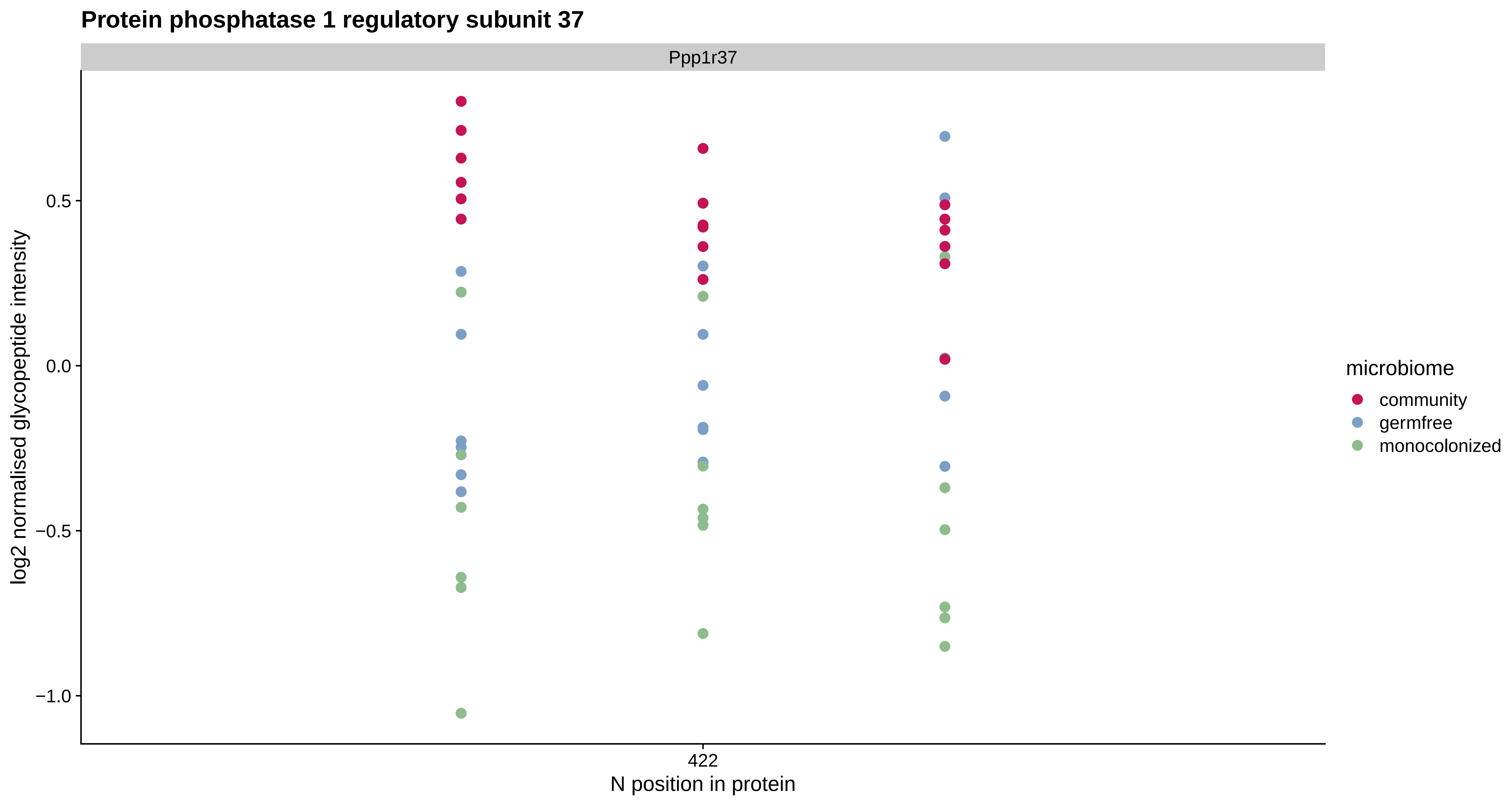
Protein phosphatase 1 regulatory subunit 37
Ppp1r37
log2 normalised glycopeptide intensity
0.5
0.0
−0.5
−1.0
422
N position in protein
microbiome
community
germfree
monocolonized

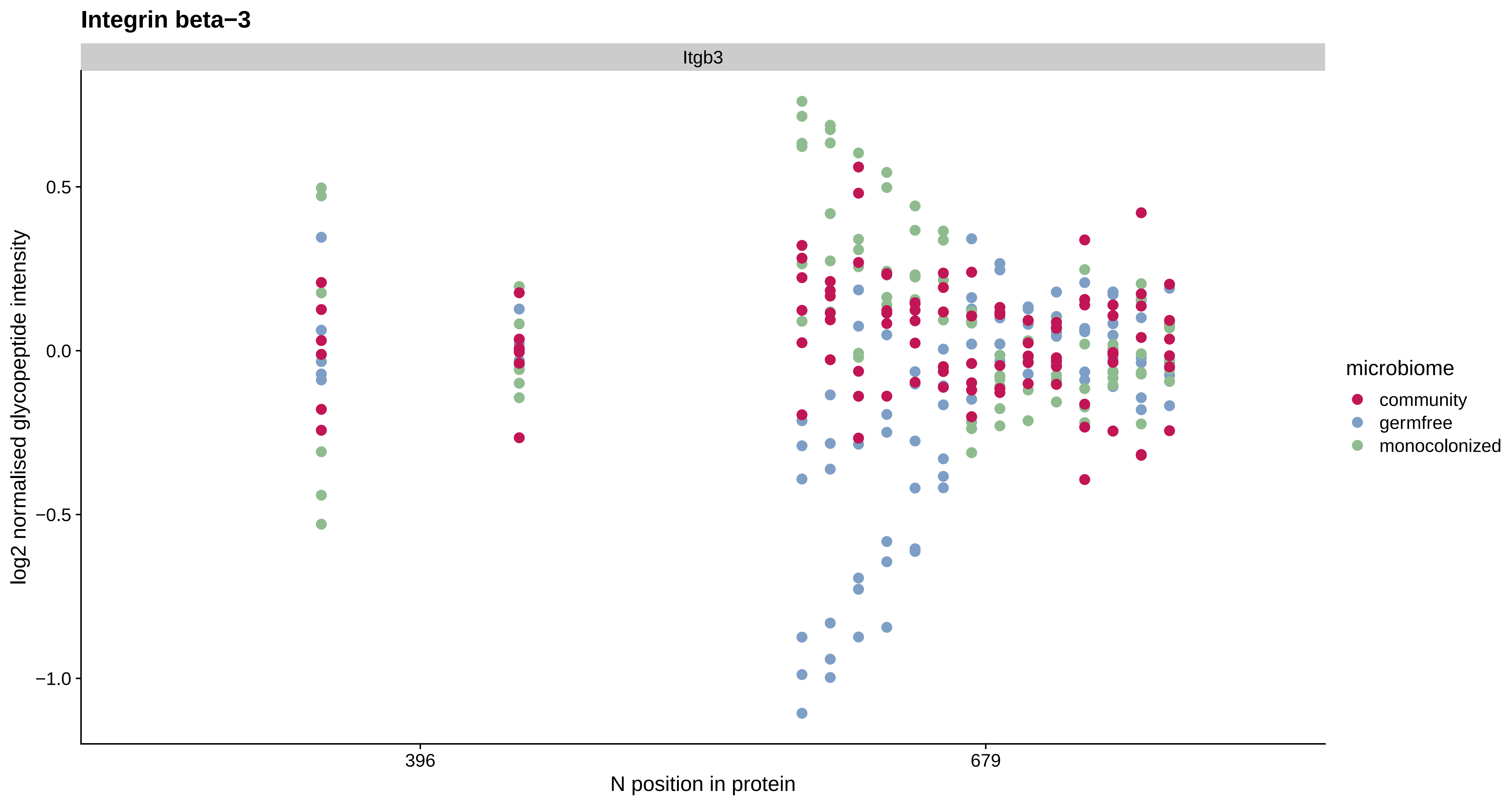
Integrin beta−3
Itgb3
0.5
0.0
−0.5
−1.0
log2 normalised glycopeptide intensity
396
679
N position in protein
microbiome
community
germfree
monocolonized

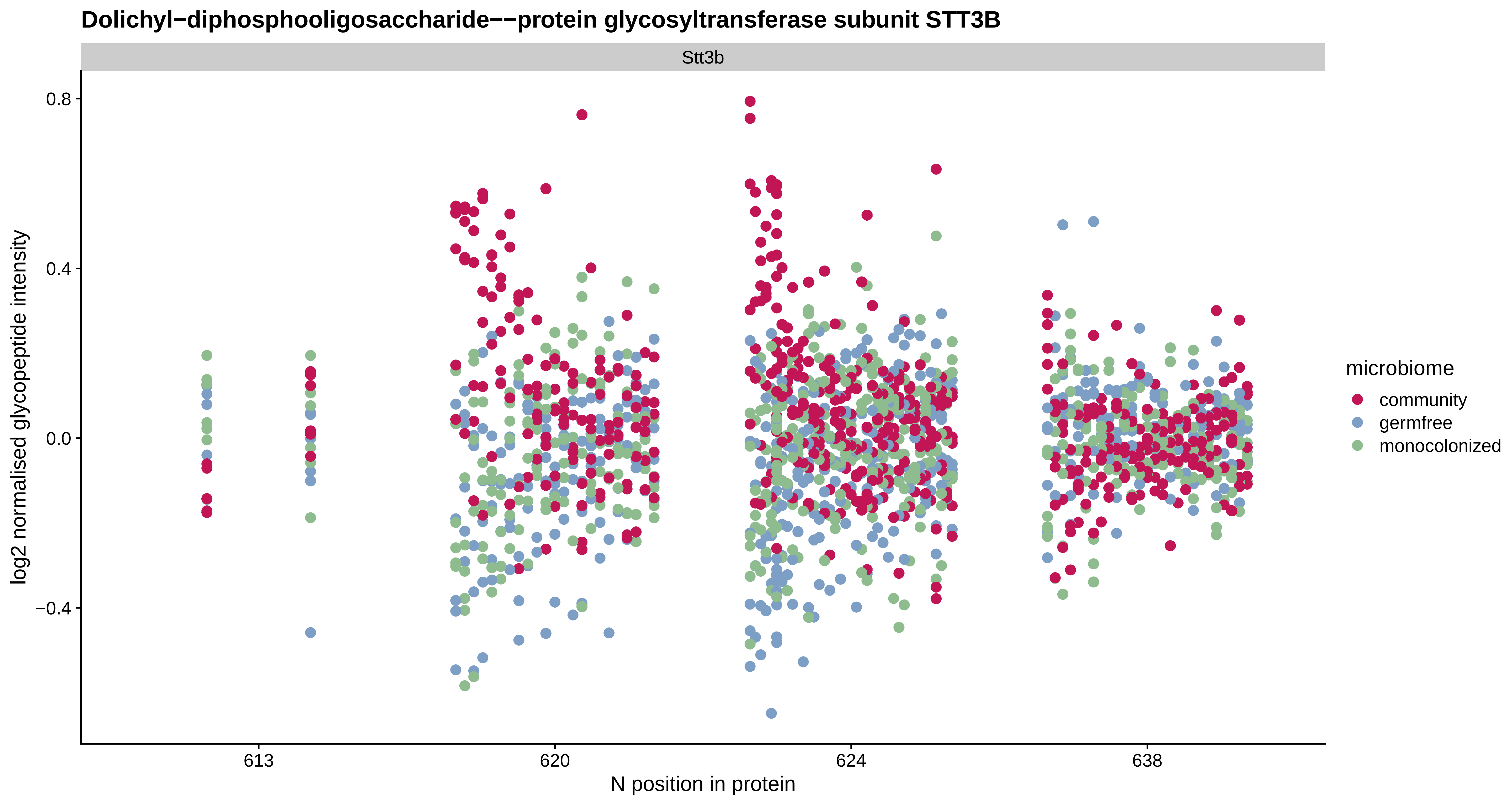
Dolichyl−diphosphooligosaccharide−−protein glycosyltransferase subunit STT3B
Stt3b
0.8
0.4
0.0
−0.4
log2 normalised glycopeptide intensity
613
620
624
638
N position in protein
microbiome
community
germfree
monocolonized

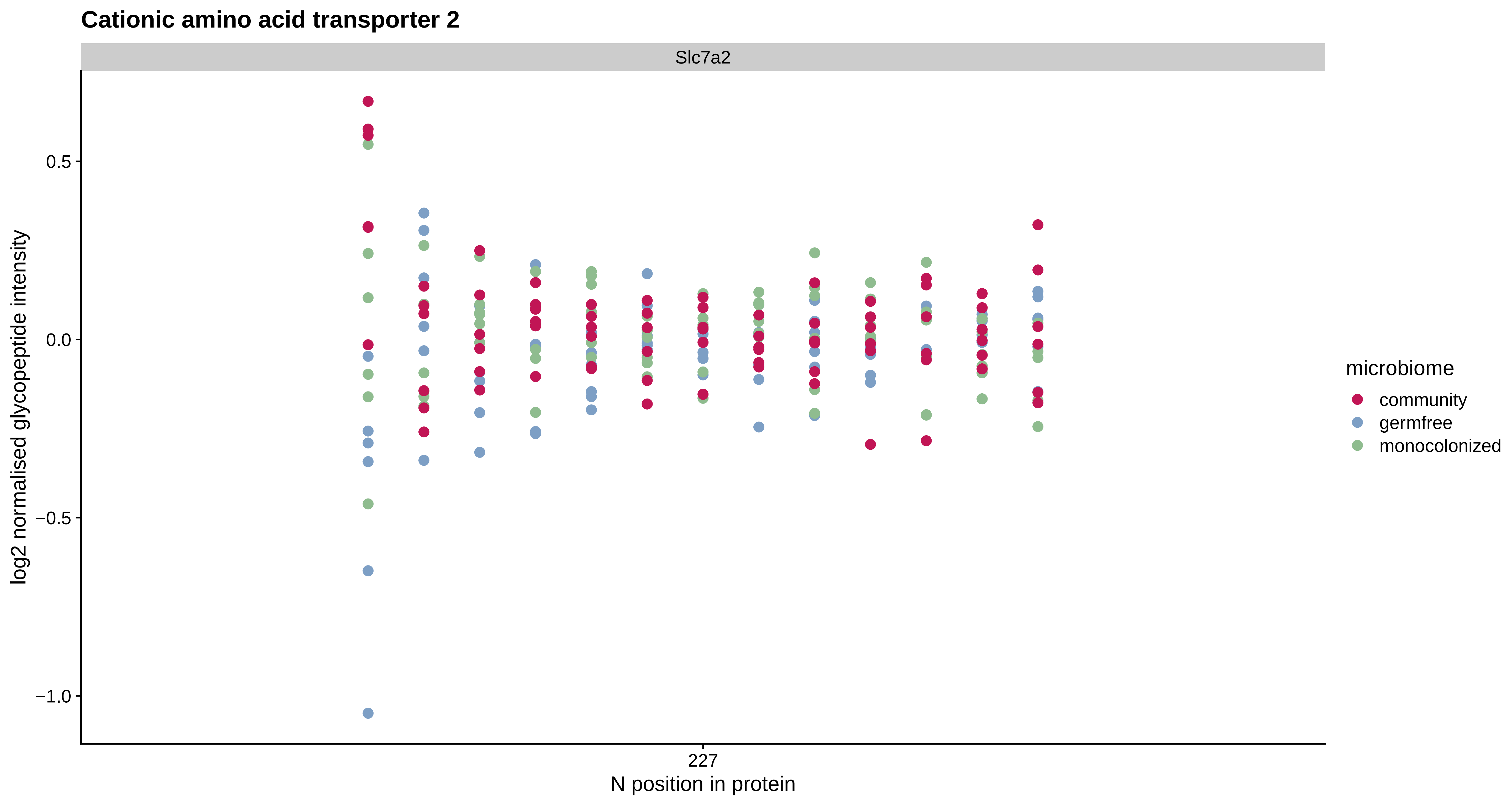
Cationic amino acid transporter 2
Slc7a2
log2 normalised glycopeptide intensity
0.5
0.0
−0.5
−1.0
227
N position in protein
microbiome
community
germfree
monocolonized

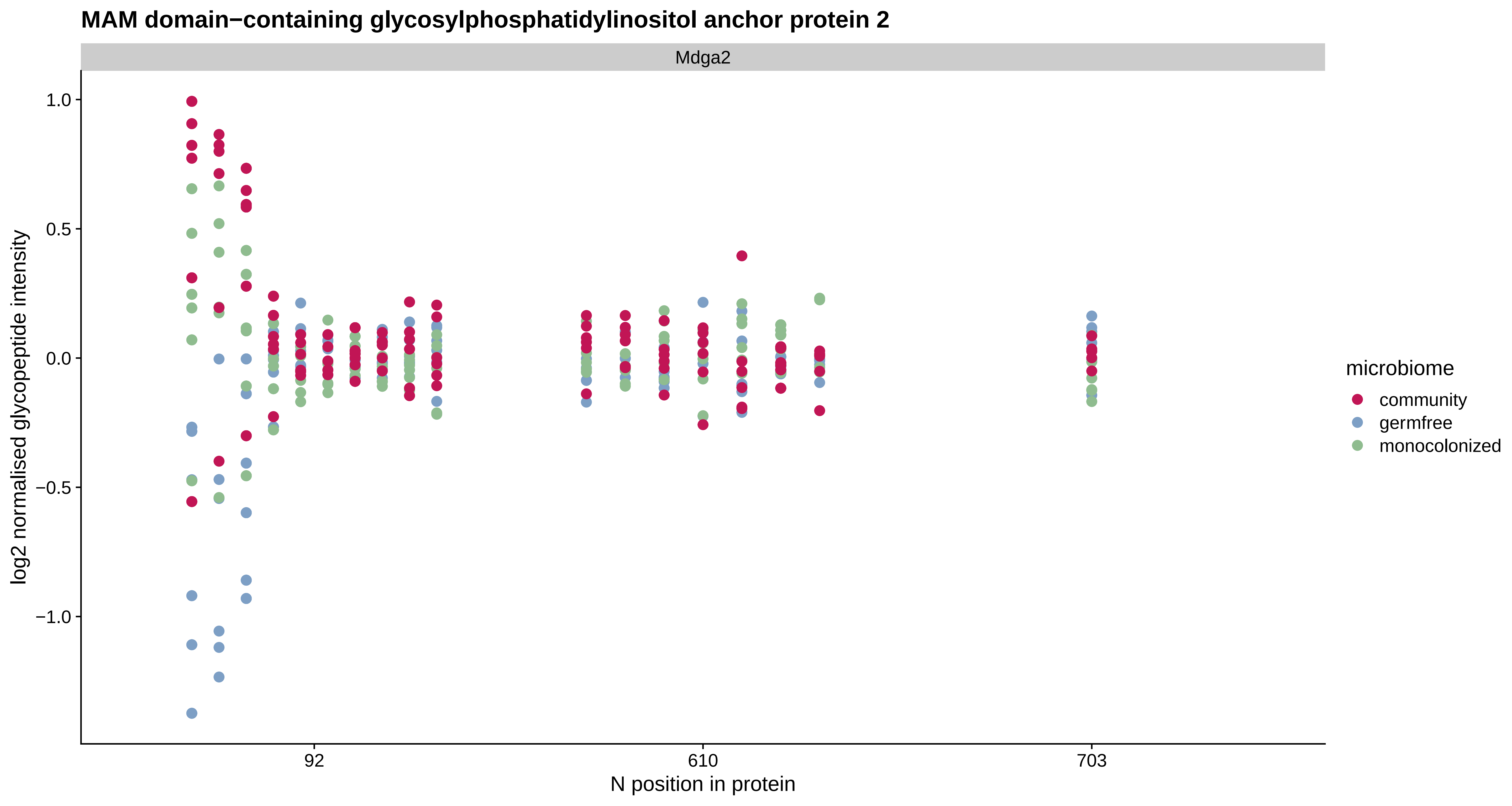
MAM domain−containing glycosylphosphatidylinositol anchor protein 2
Mdga2
1.0
0.5
0.0
−0.5
−1.0
log2 normalised glycopeptide intensity
92
610
703
N position in protein
microbiome
community
germfree
monocolonized

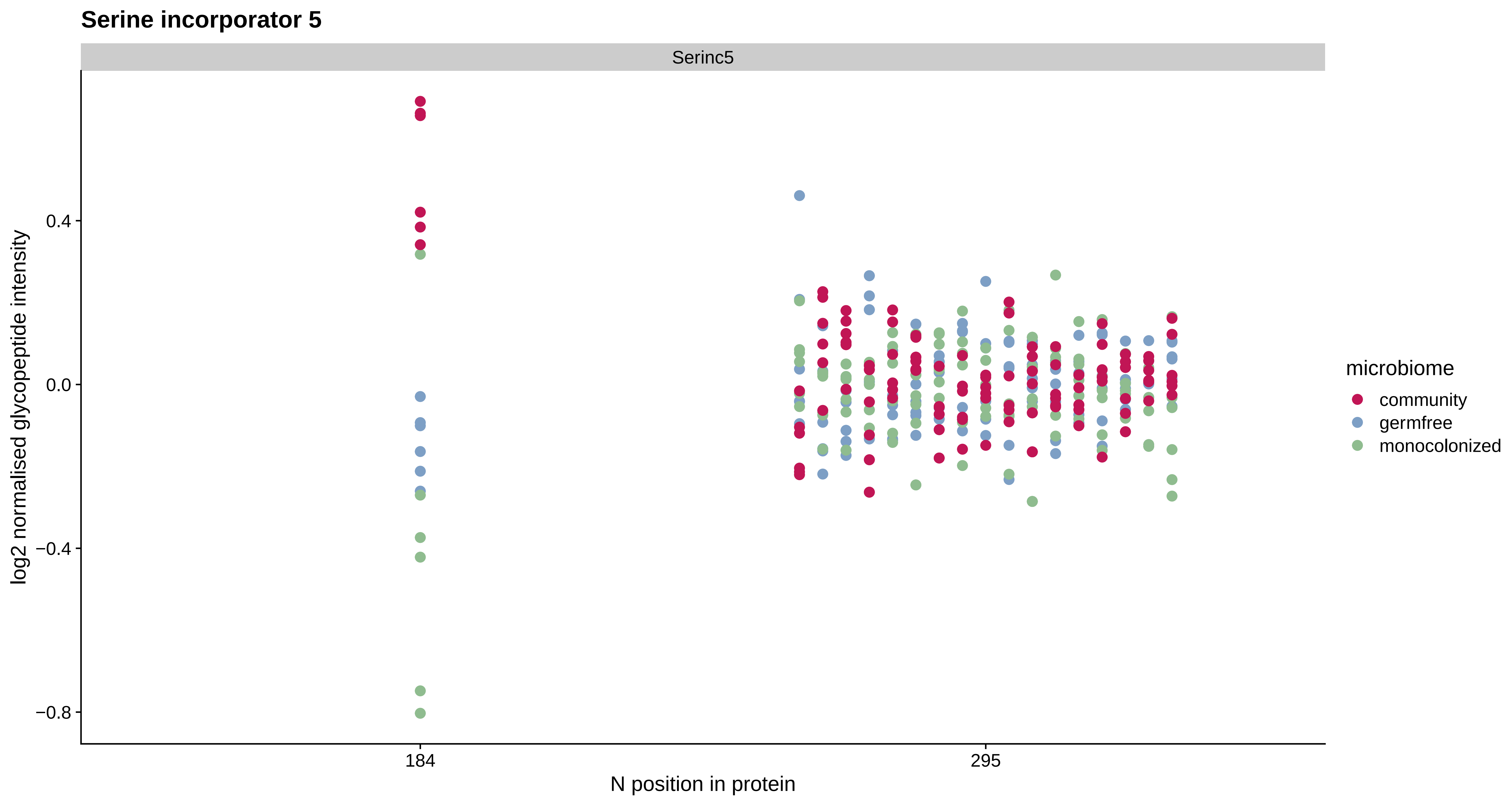

Serine incorporator 5
Serinc5
log2 normalised glycopeptide intensity
0.4
0.0
−0.4
−0.8
184
295
N position in protein
microbiome
community
germfree
monocolonized

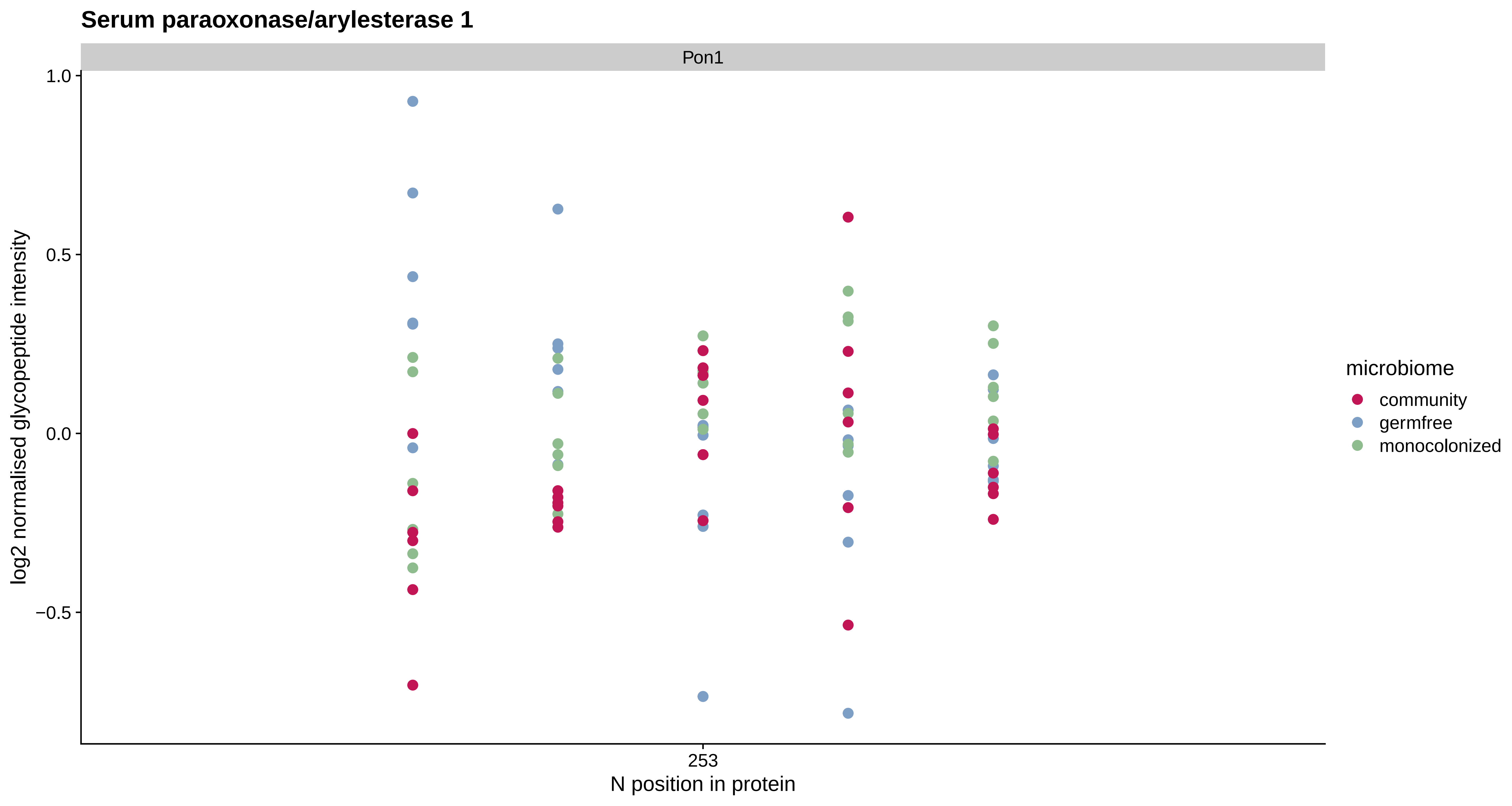
Serum paraoxonase/arylesterase 1
Pon1
1.0
0.5
0.0
−0.5
log2 normalised glycopeptide intensity
253
N position in protein
microbiome
community
germfree
monocolonized

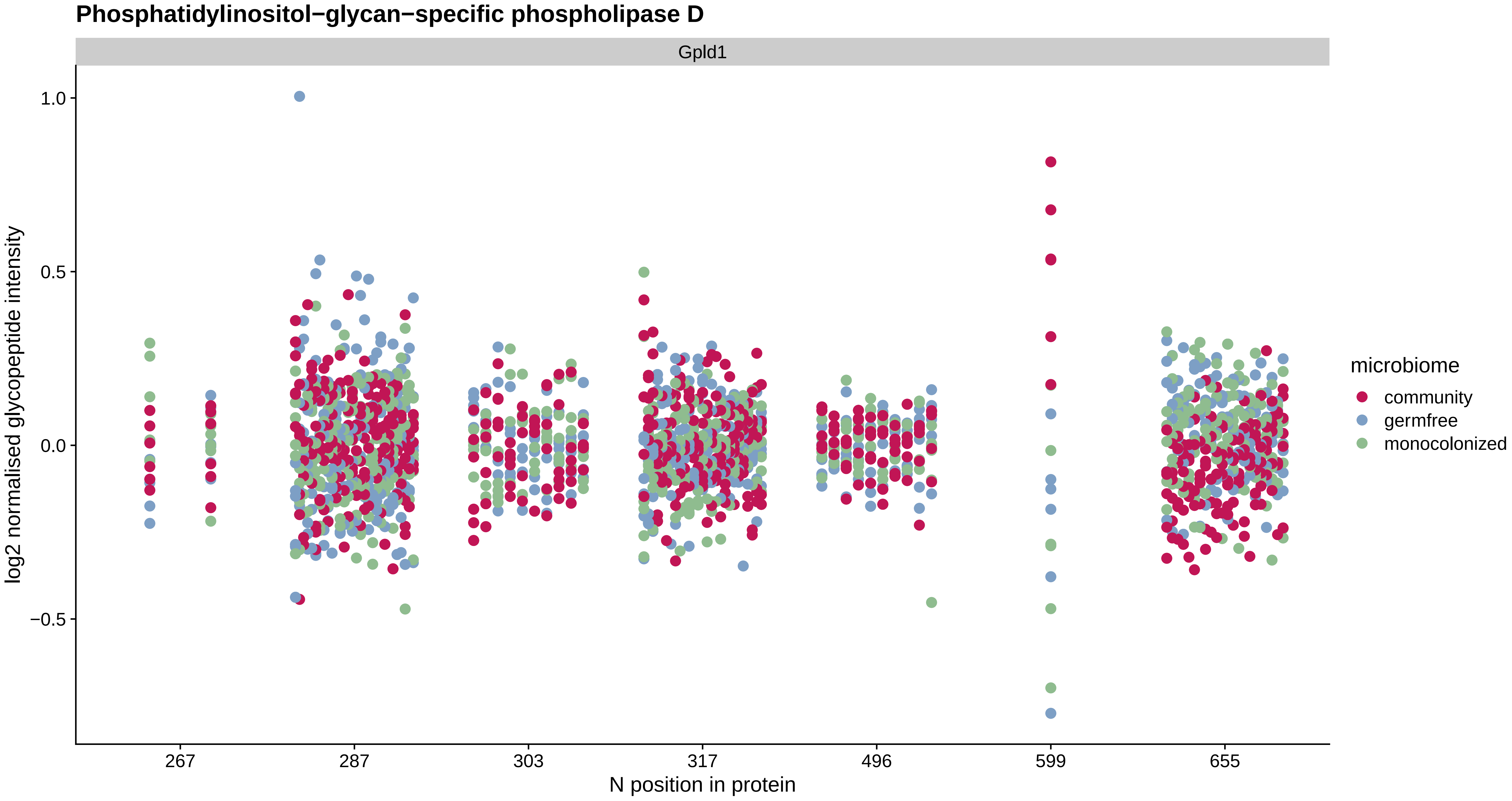

Phosphatidylinositol−glycan−specific phospholipase D
Gpld1
1.0
0.5
0.0
−0.5
log2 normalised glycopeptide intensity
267
287
303
317
496
599
655
N position in protein
microbiome
community
germfree
monocolonized

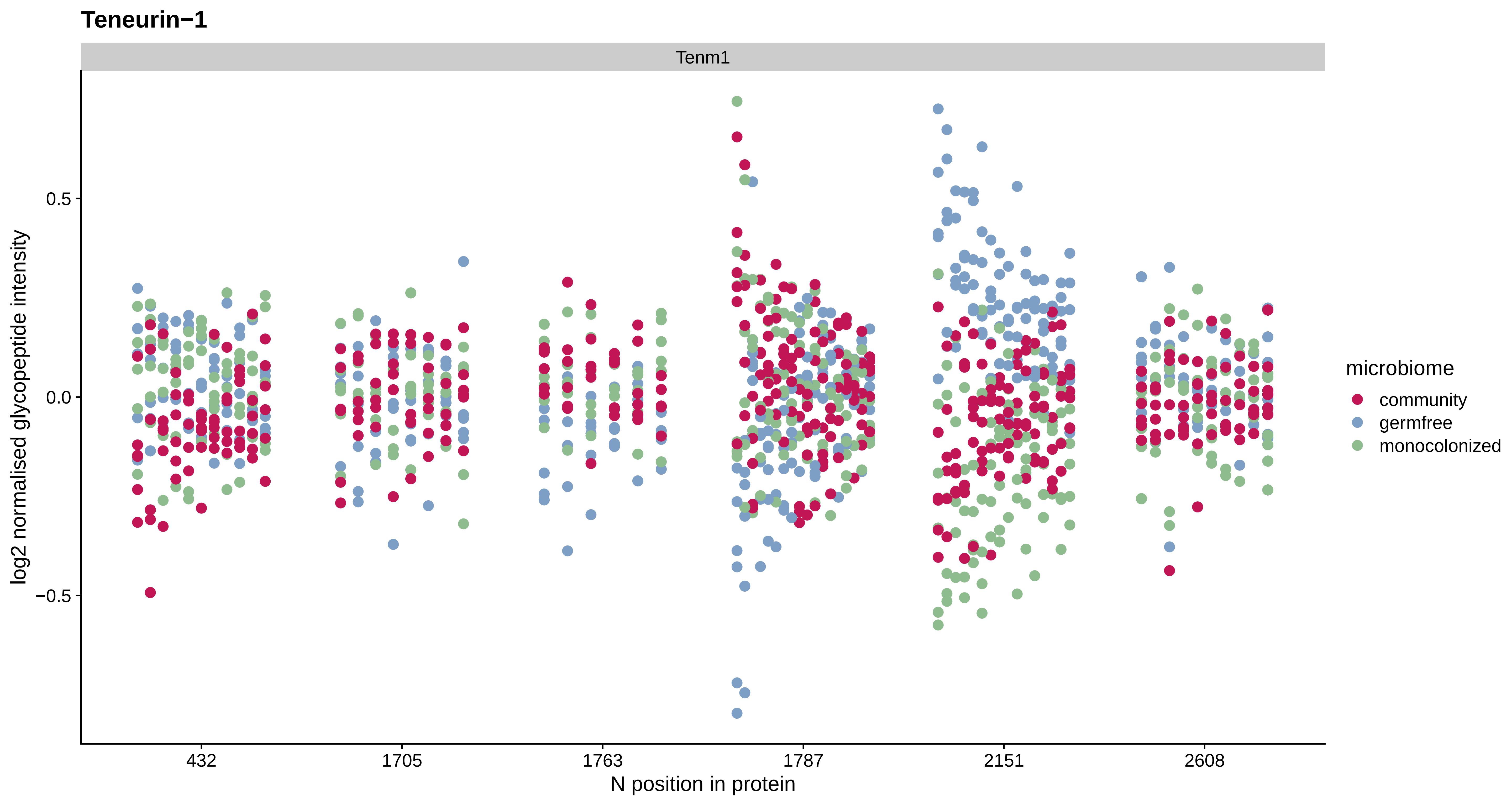
Teneurin−1
Tenm1
0.5
0.0
−0.5
log2 normalised glycopeptide intensity
432
1705
1763
1787
2151
2608
N position in protein
microbiome
community
germfree
monocolonized

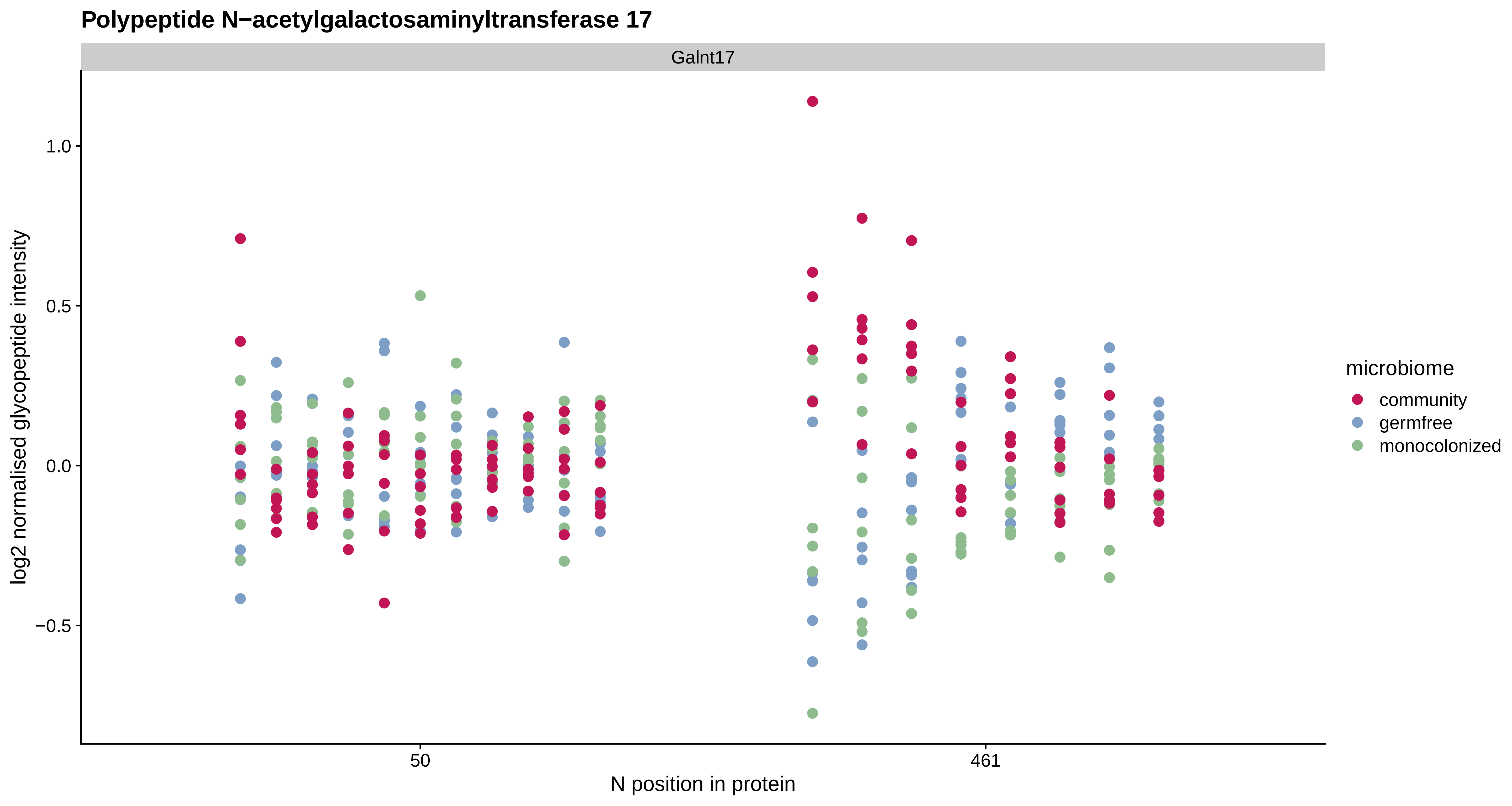
Polypeptide N−acetylgalactosaminyltransferase 17
Galnt17
log2 normalised glycopeptide intensity
1.0
0.5
0.0
−0.5
50
461
N position in protein
microbiome
community
germfree
monocolonized

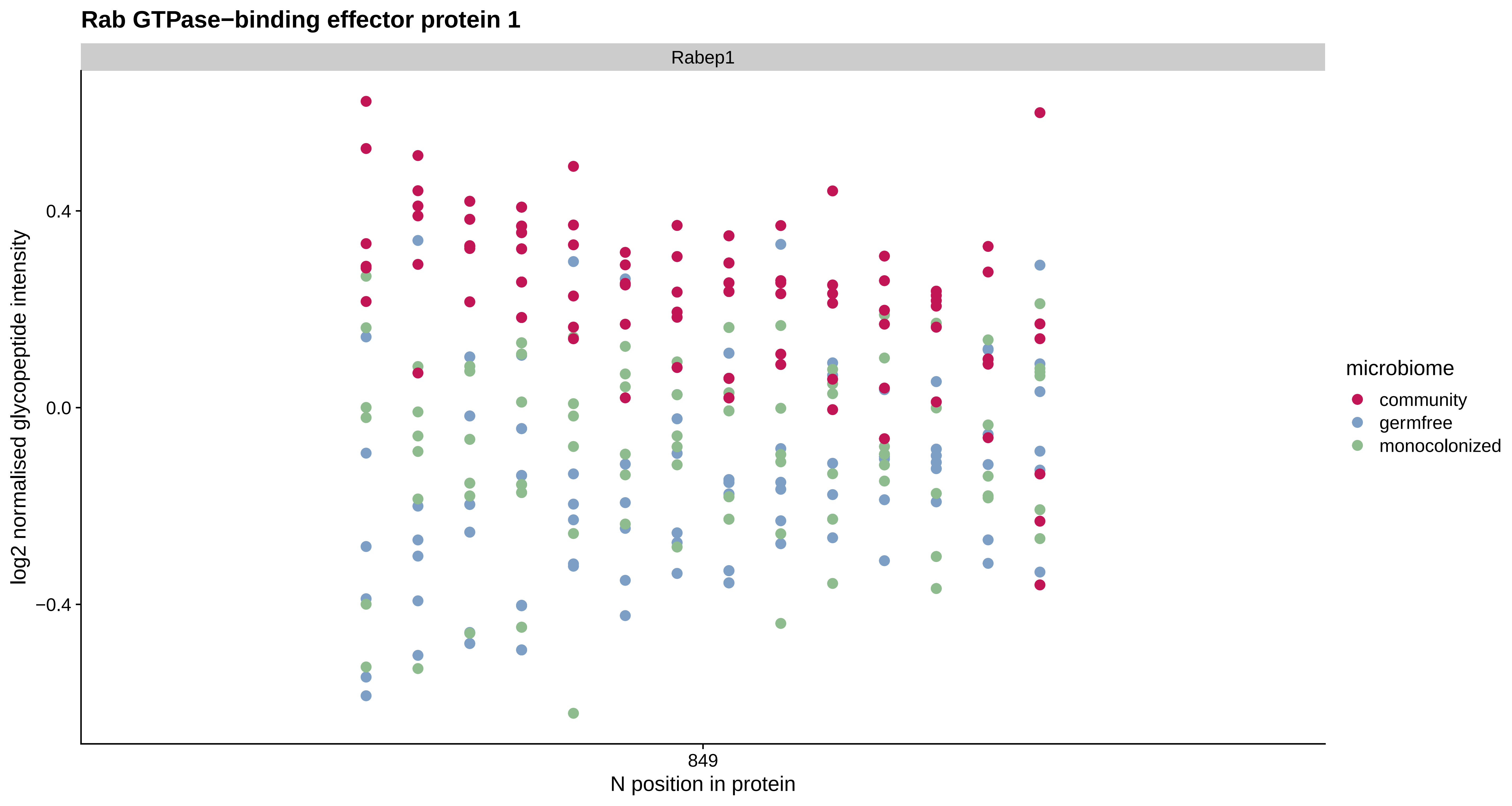

Rab GTPase−binding effector protein 1
Rabep1
log2 normalised glycopeptide intensity
0.4
0.0
−0.4
849
N position in protein
microbiome
community
germfree
monocolonized

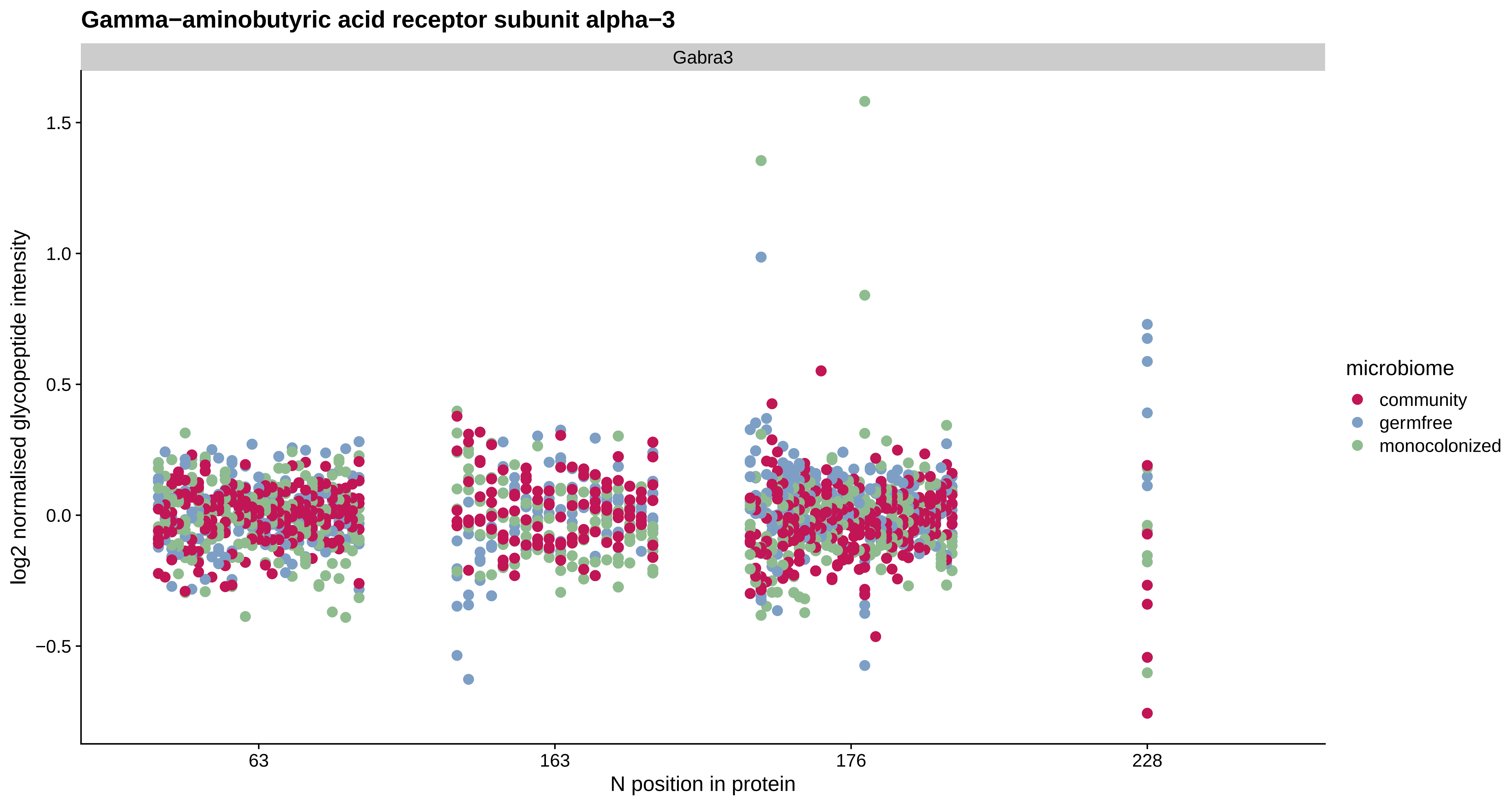

Gamma−aminobutyric acid receptor subunit alpha−3
Gabra3
log2 normalised glycopeptide intensity
1.5
1.0
0.5
0.0
−0.5
63
163
176
228
N position in protein
microbiome
community
germfree
monocolonized

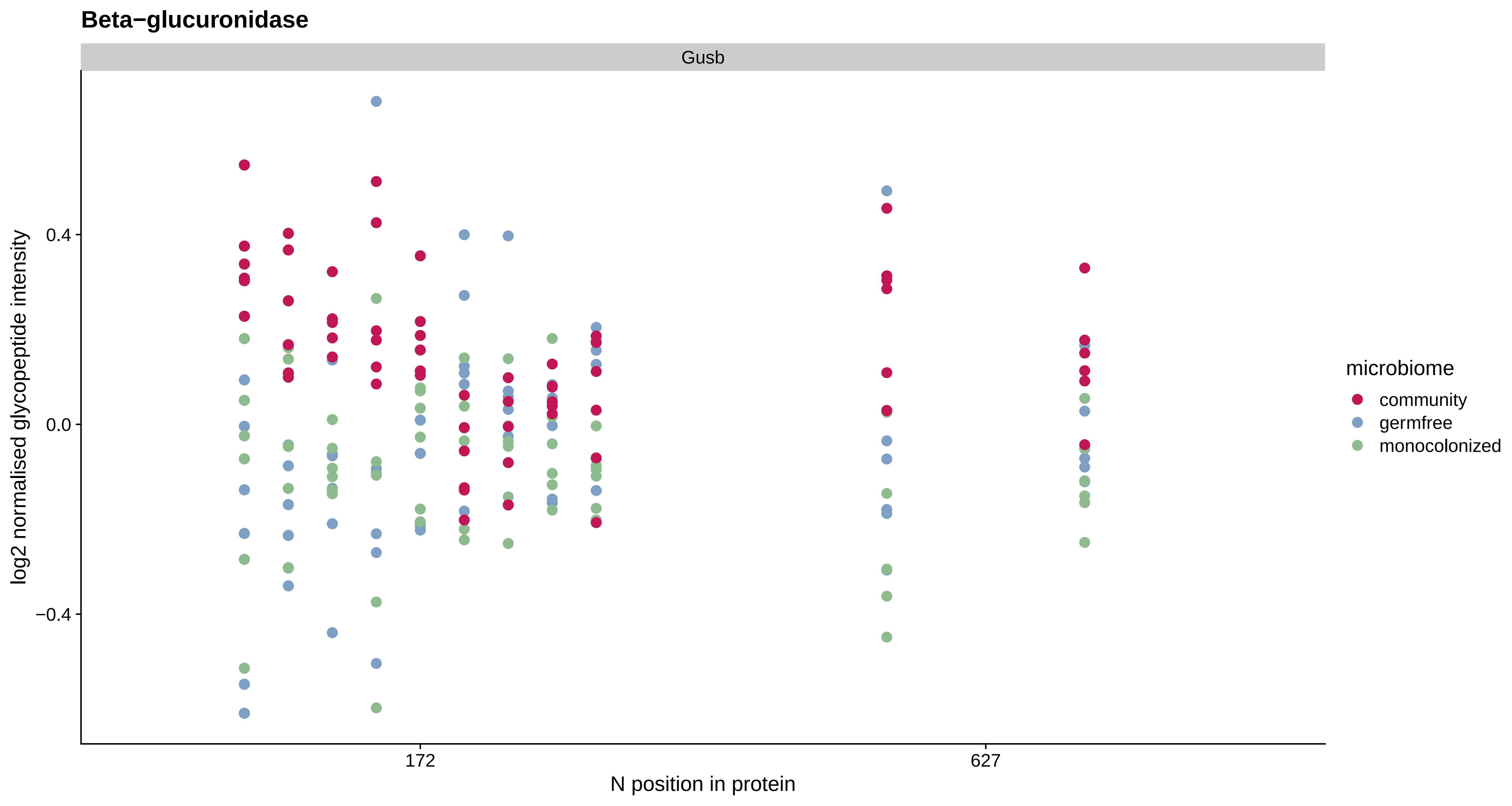
Beta−glucuronidase
Gusb
log2 normalised glycopeptide intensity
0.4
0.0
−0.4
172
627
N position in protein
microbiome
community
germfree
monocolonized

#### Nucleotide exchange factor SIL1

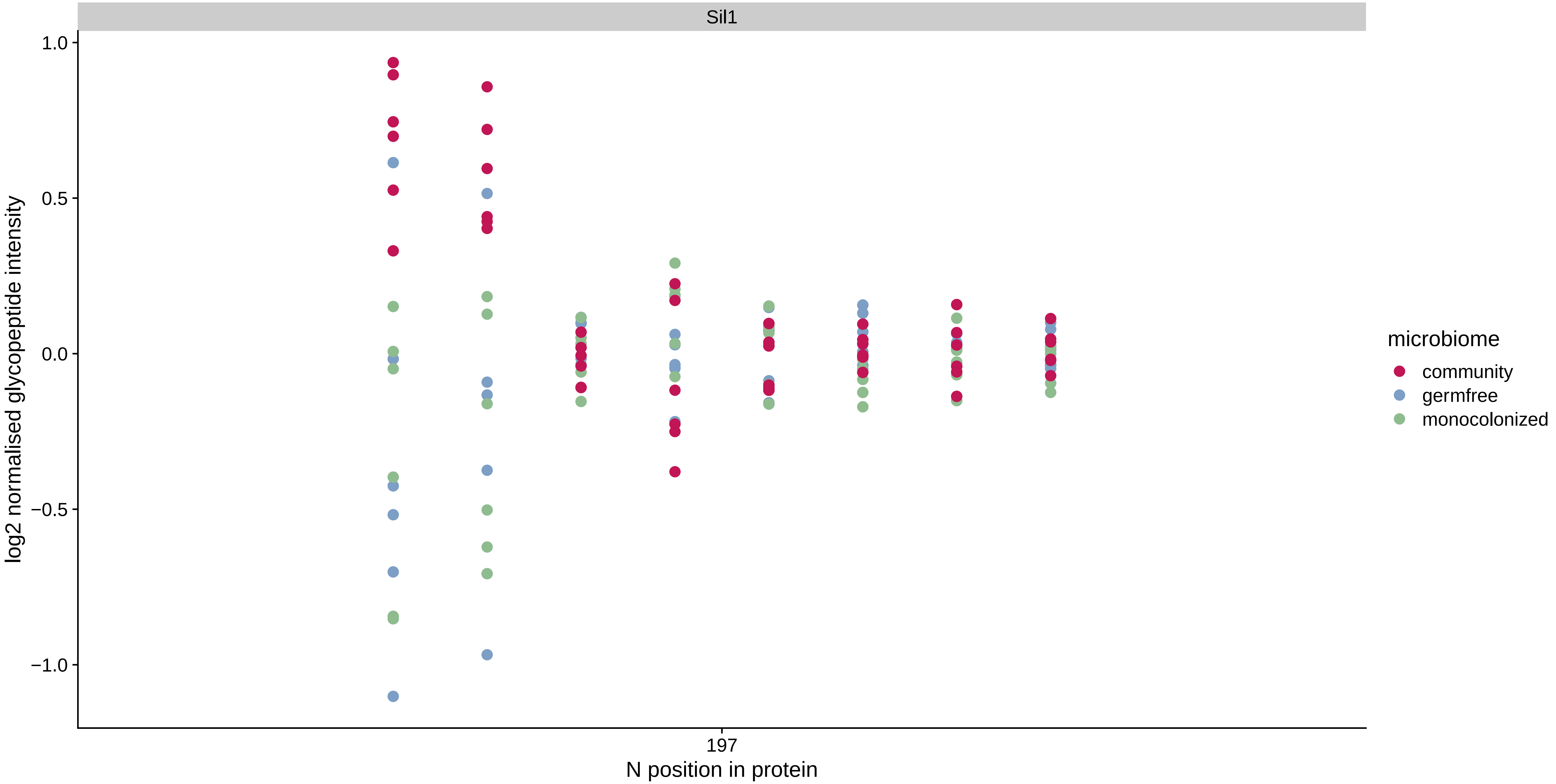

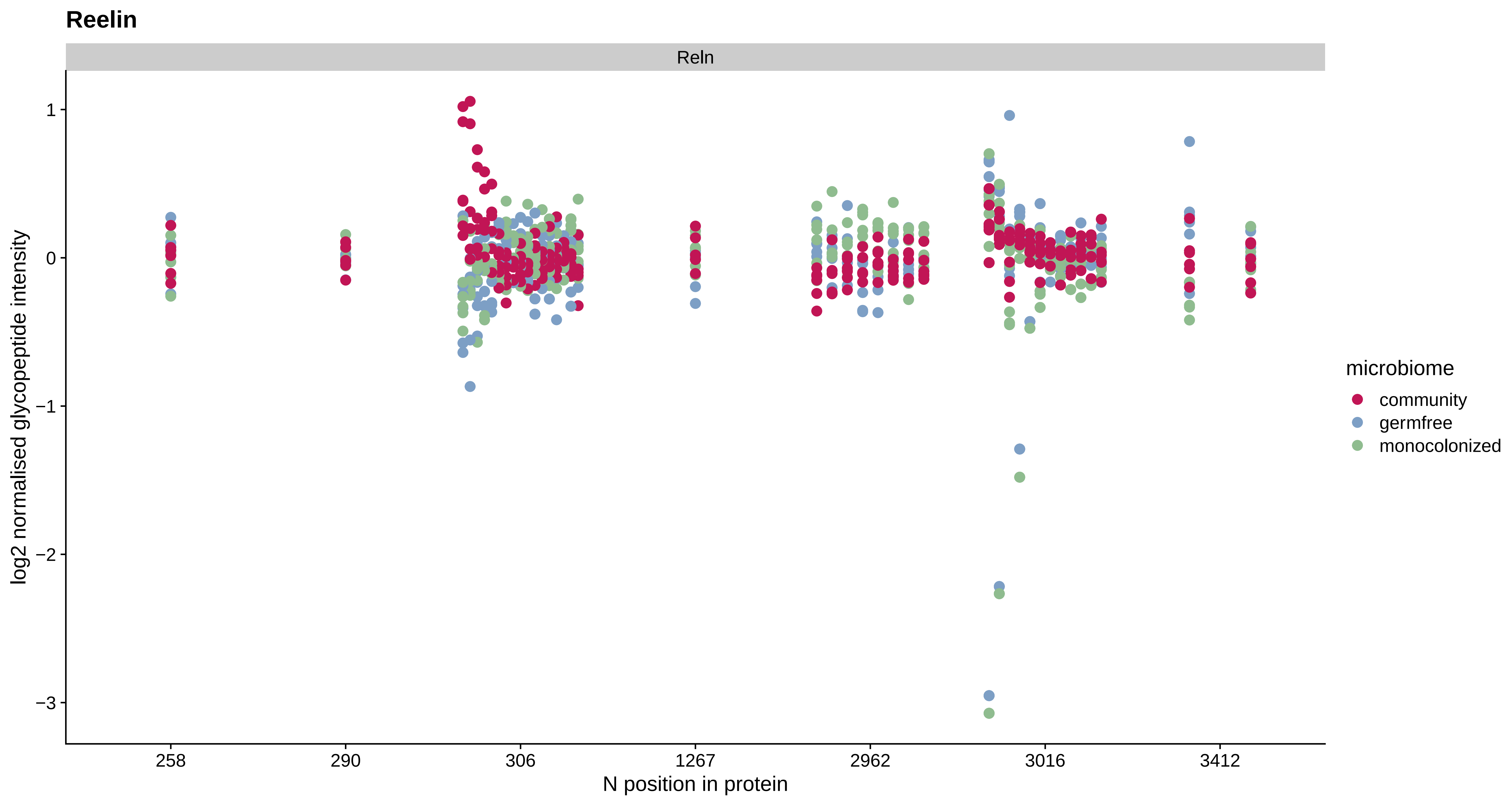
Reelin
Reln
1
0
−1
−2
−3
log2 normalised glycopeptide intensity
258
290
306
1267
2962
3016
3412
N position in protein
microbiome
community
germfree
monocolonized

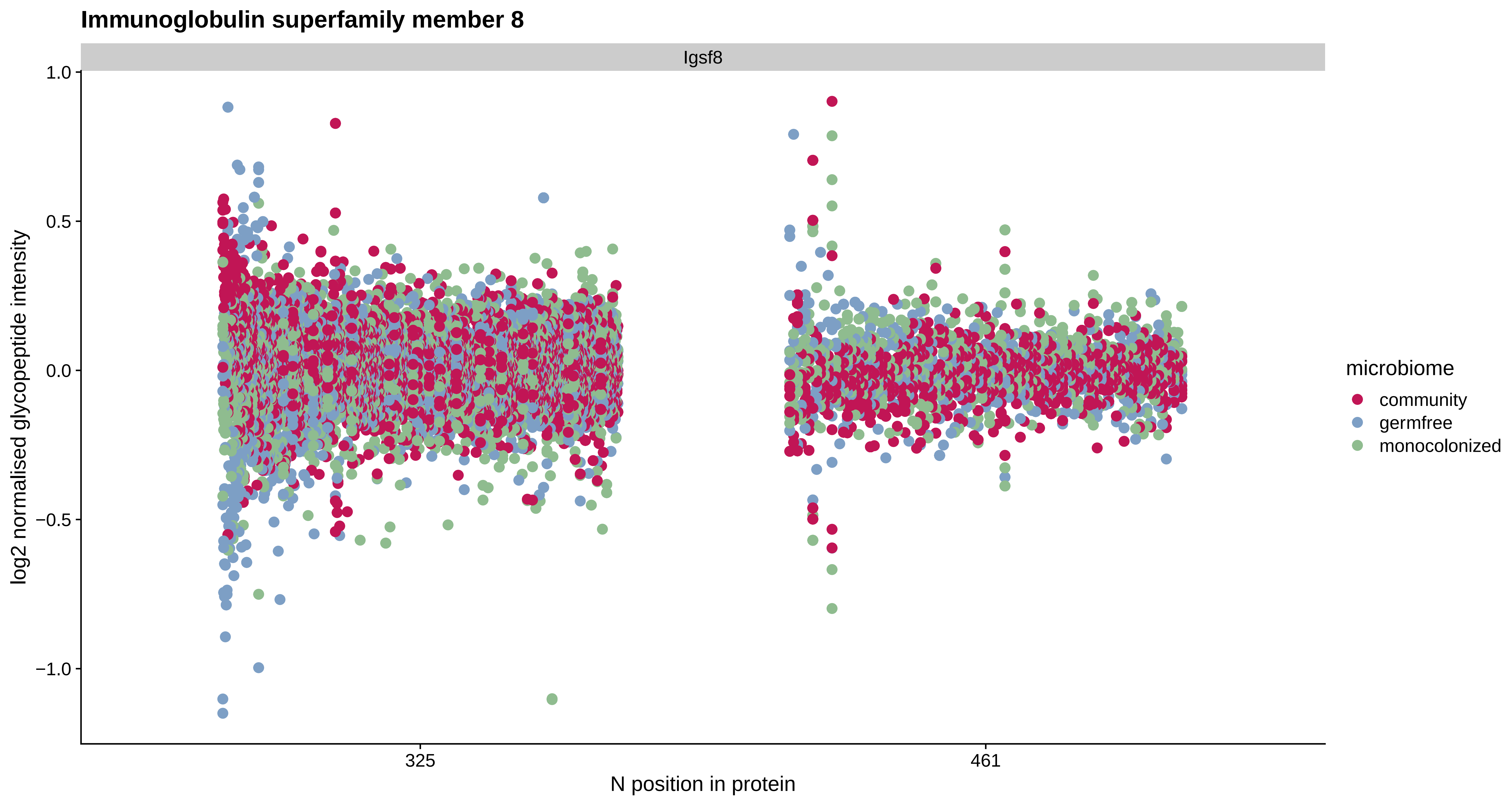

Immunoglobulin superfamily member 8
Igsf8
log2 normalised glycopeptide intensity
1.0
0.5
0.0
−0.5
−1.0
325
461
N position in protein
microbiome
community
germfree
monocolonized

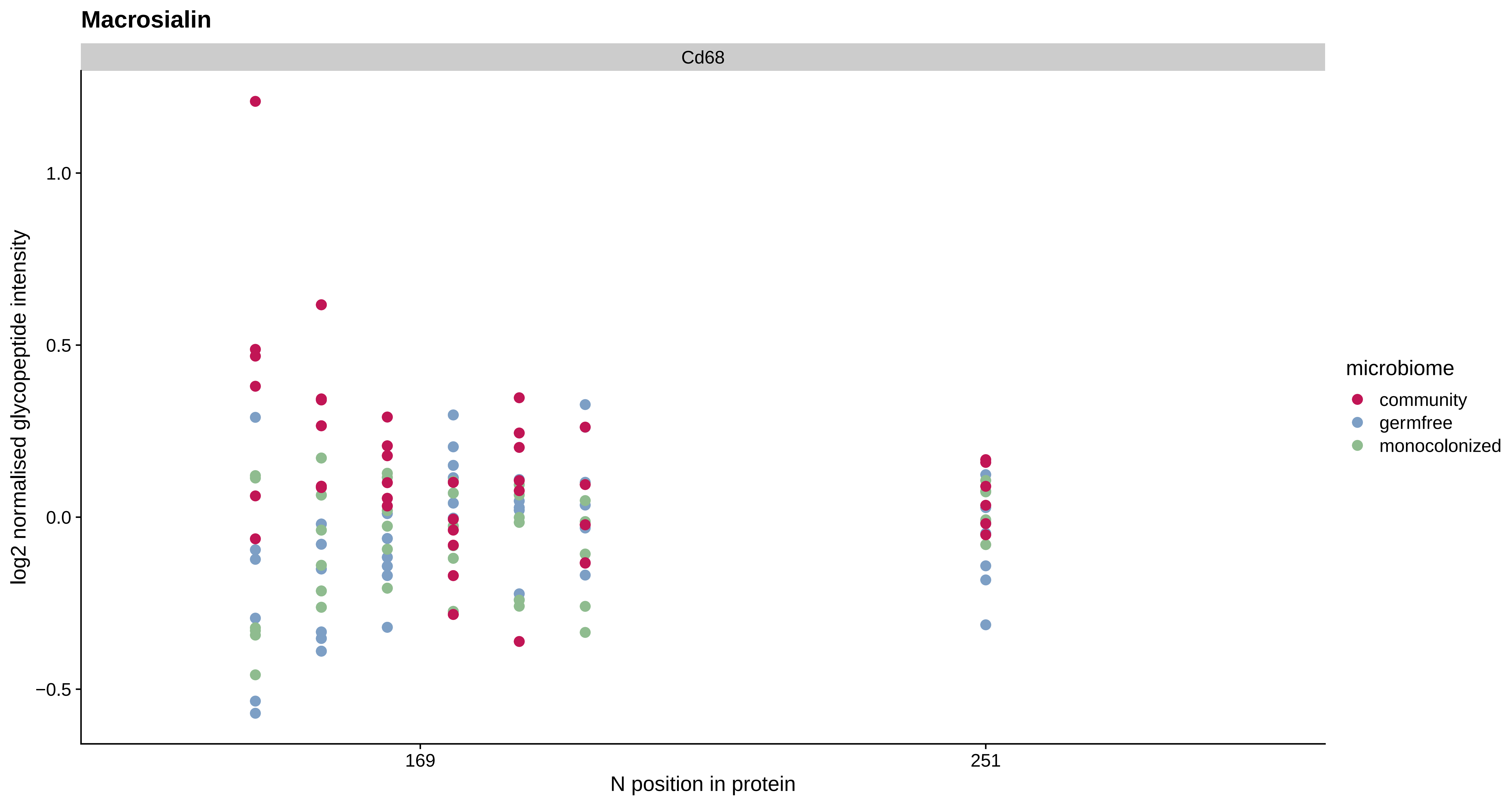
Macrosialin
Cd68
log2 normalised glycopeptide intensity
1.0
0.5
0.0
−0.5
169
251
N position in protein
microbiome
community
germfree
monocolonized

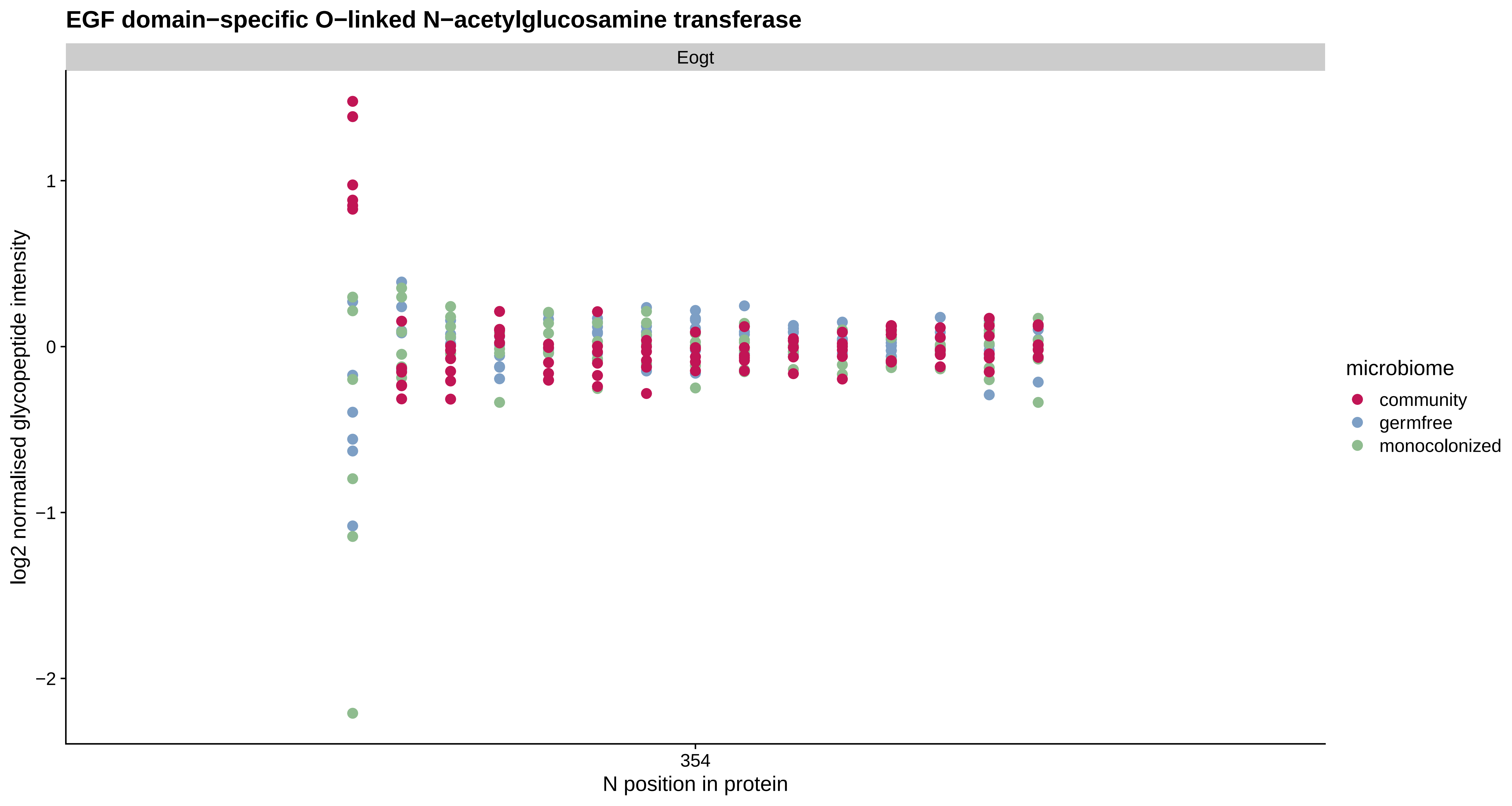

EGF domain−specific O−linked N−acetylglucosamine transferase
Eogt
log2 normalised glycopeptide intensity
1
0
−1
−2
354
N position in protein
microbiome
community
germfree
monocolonized

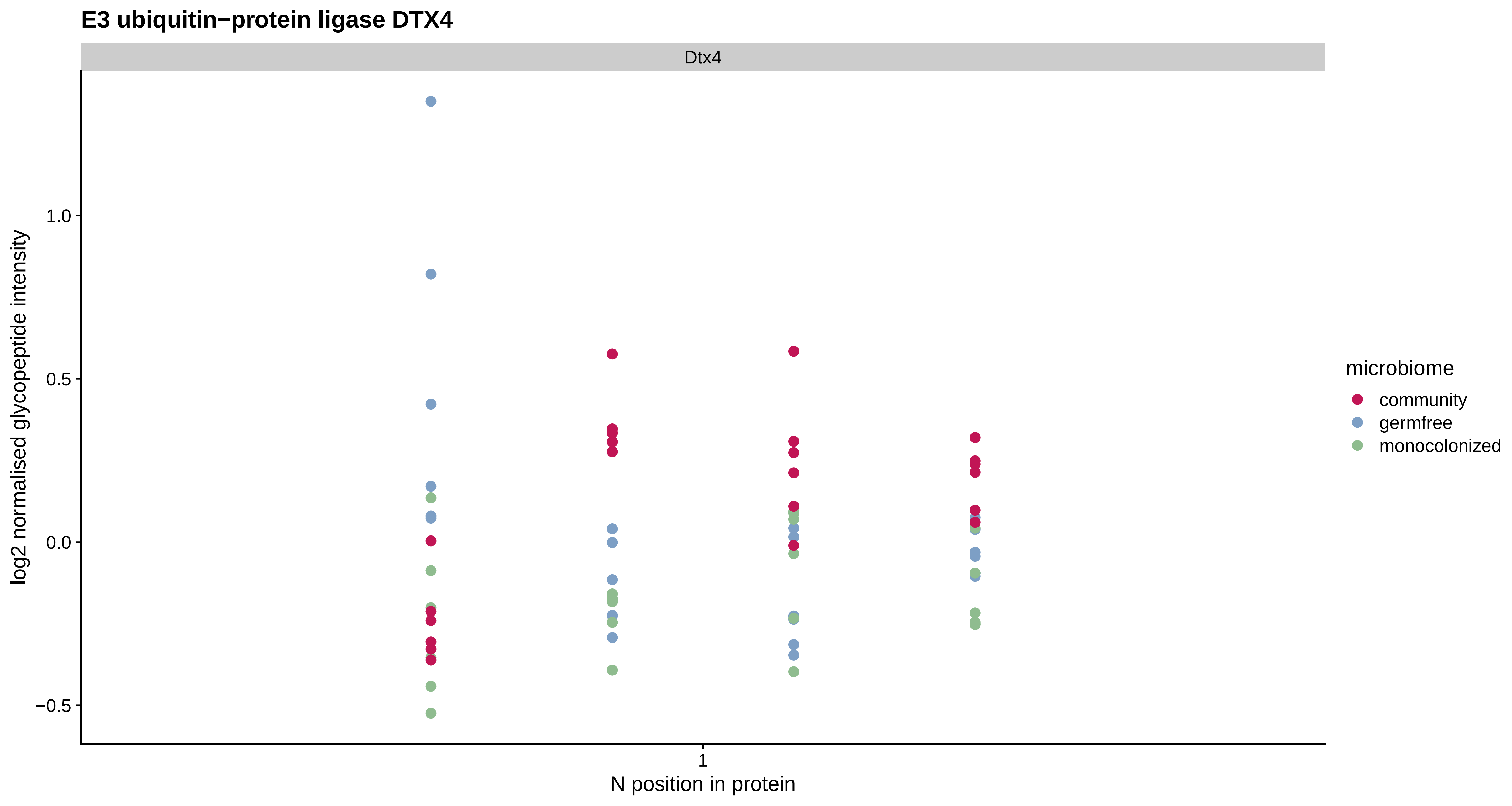
E3 ubiquitin−protein ligase DTX4
Dtx4
log2 normalised glycopeptide intensity
1.0
0.5
0.0
−0.5
1
N position in protein
microbiome
community
germfree
monocolonized

**Glutamate receptor ionotropic, delta−2**

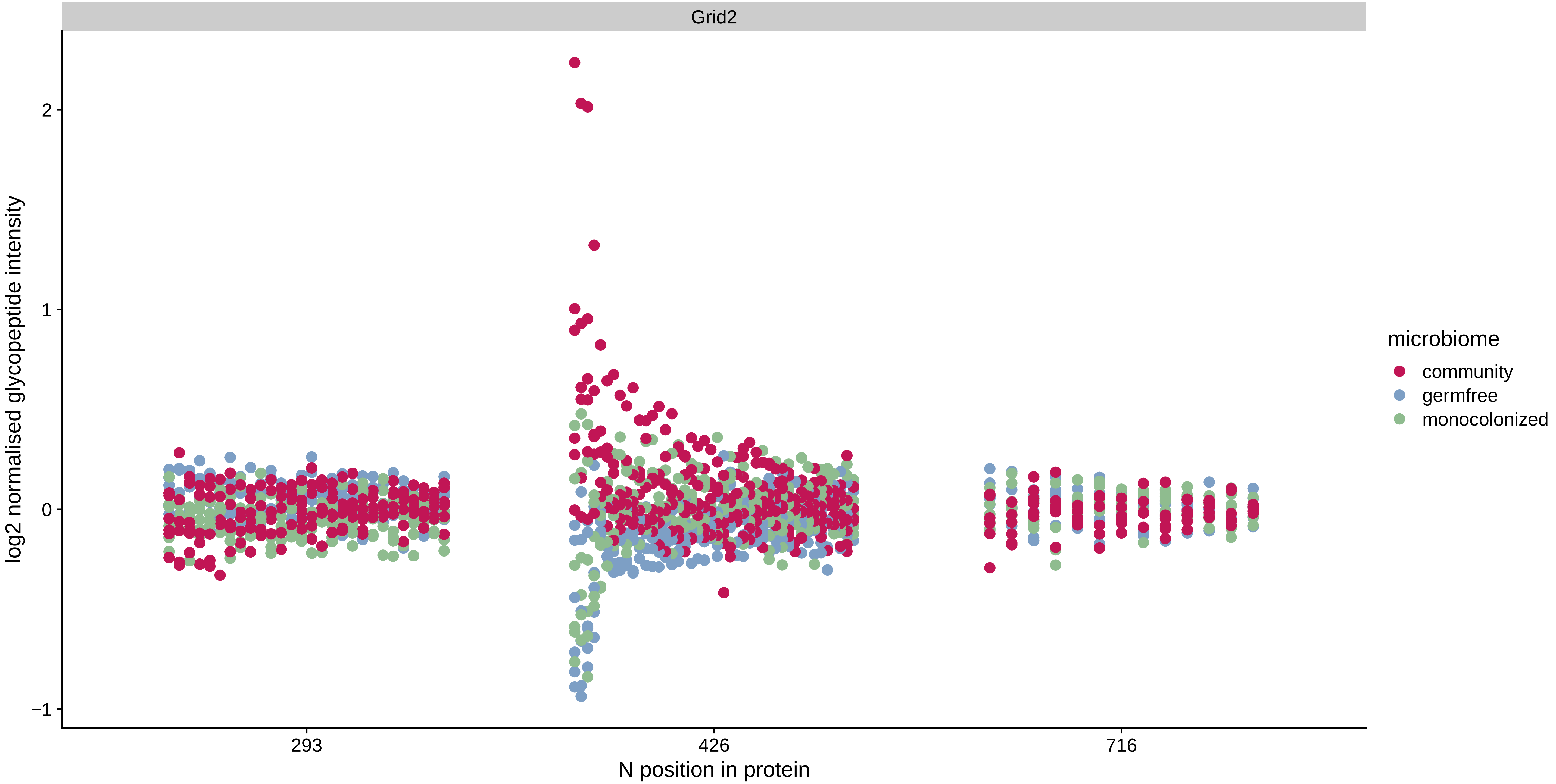

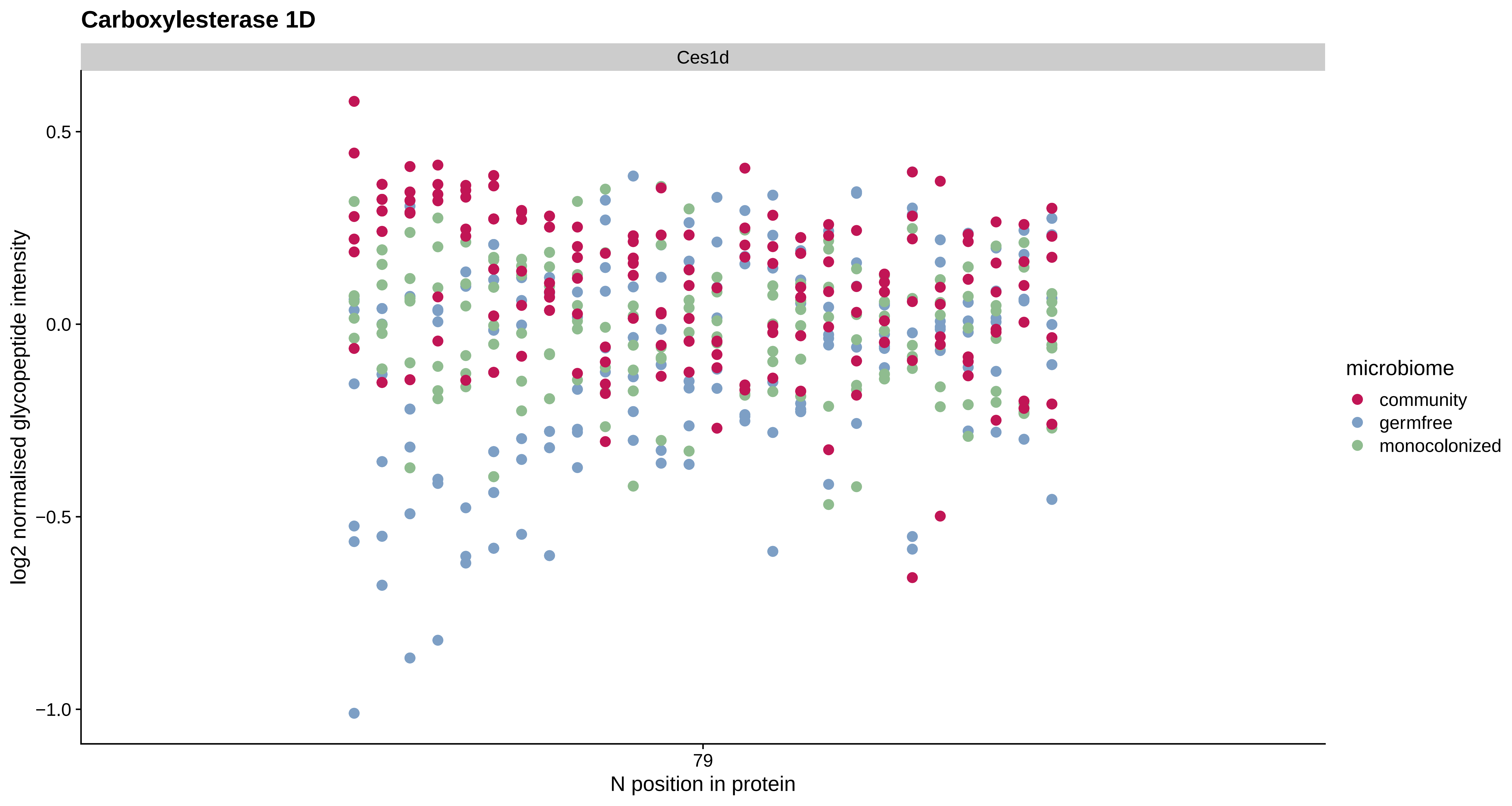
Carboxylesterase 1D
Ces1d
0.5
0.0
−0.5
−1.0
log2 normalised glycopeptide intensity
79
N position in protein
microbiome
community
germfree
monocolonized

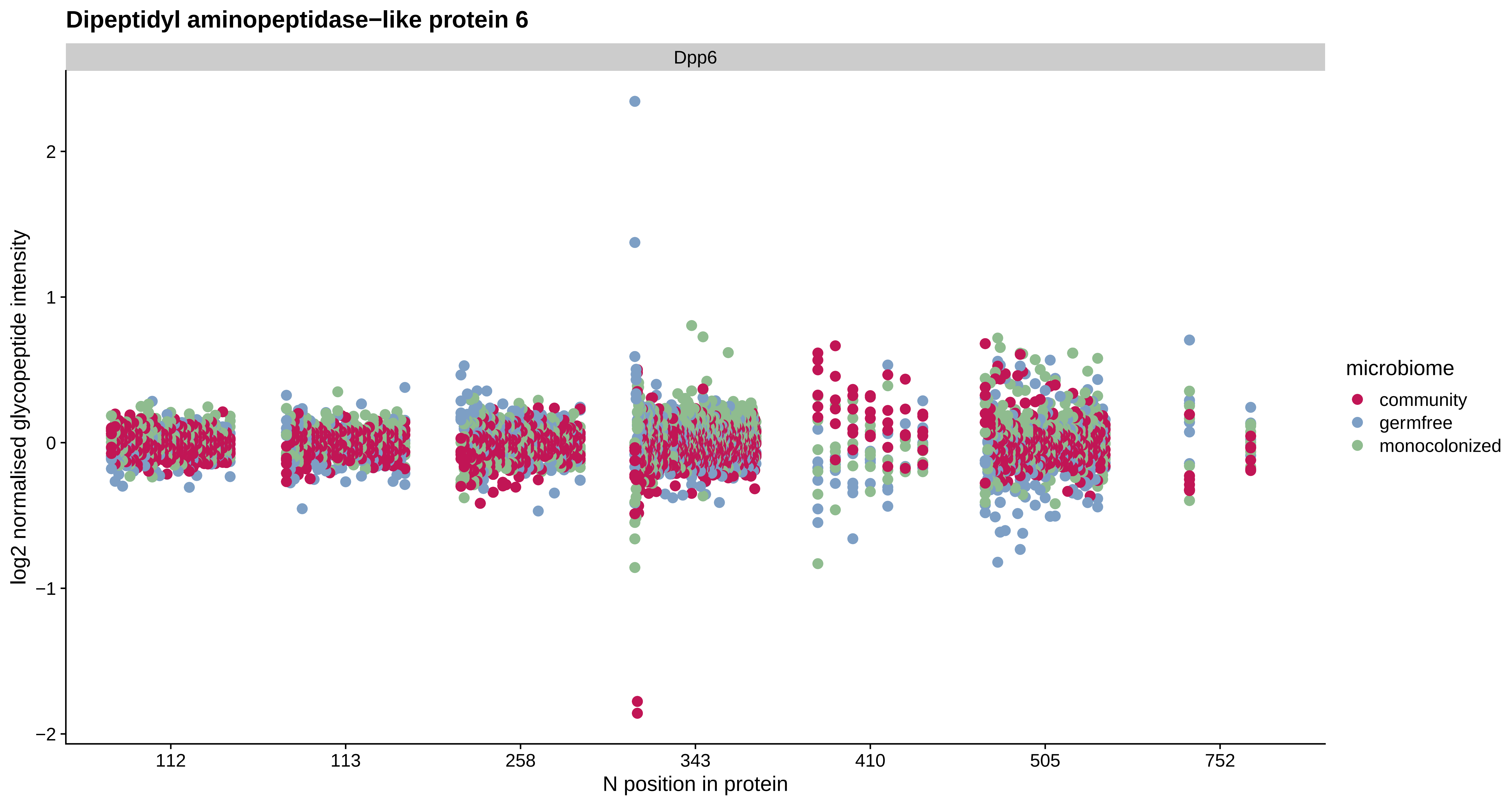

Dipeptidyl aminopeptidase−like protein 6
Dpp6
log2 normalised glycopeptide intensity
2
1
0
−1
−2
112
113
258
343
410
505
752
N position in protein
microbiome
community
germfree
monocolonized

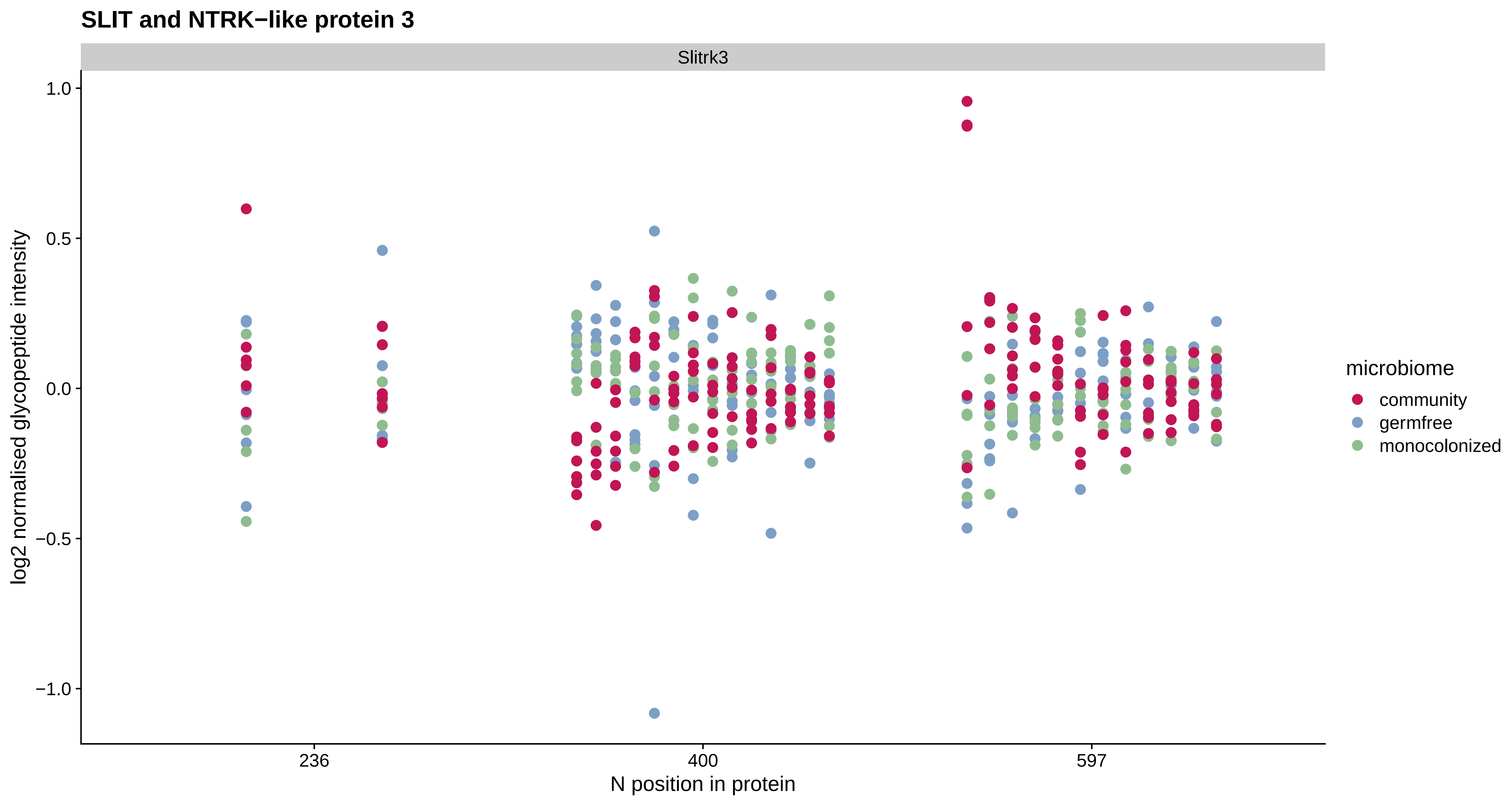
SLIT and NTRK−like protein 3
Slitrk3
1.0
0.5
0.0
−0.5
−1.0
log2 normalised glycopeptide intensity
236
400
597
N position in protein
microbiome
community
germfree
monocolonized

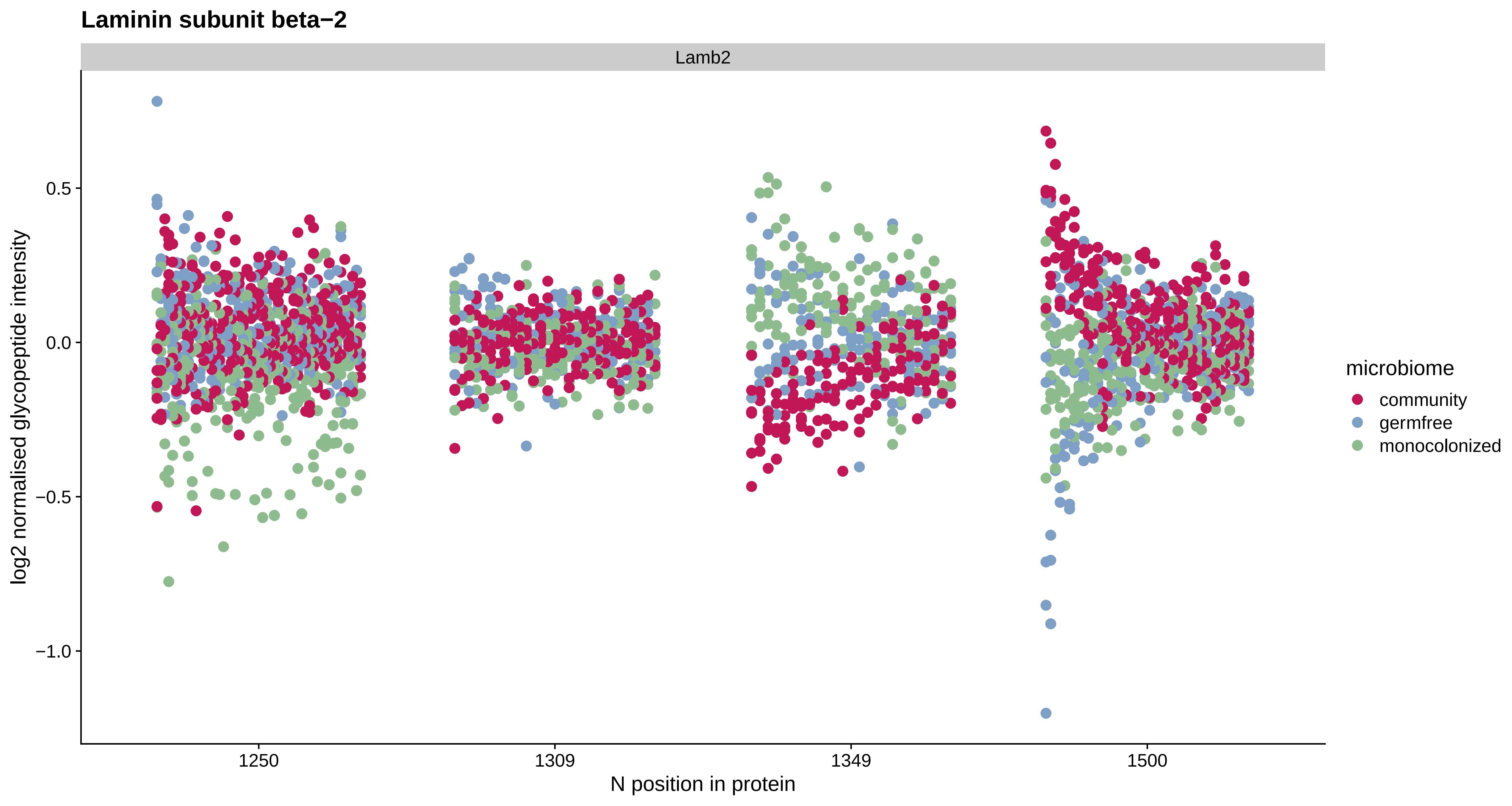

Laminin subunit beta−2
Lamb2
0.5
0.0
−0.5
−1.0
log2 normalised glycopeptide intensity
1250
1309
1349
1500
N position in protein
microbiome
community
germfree
monocolonized

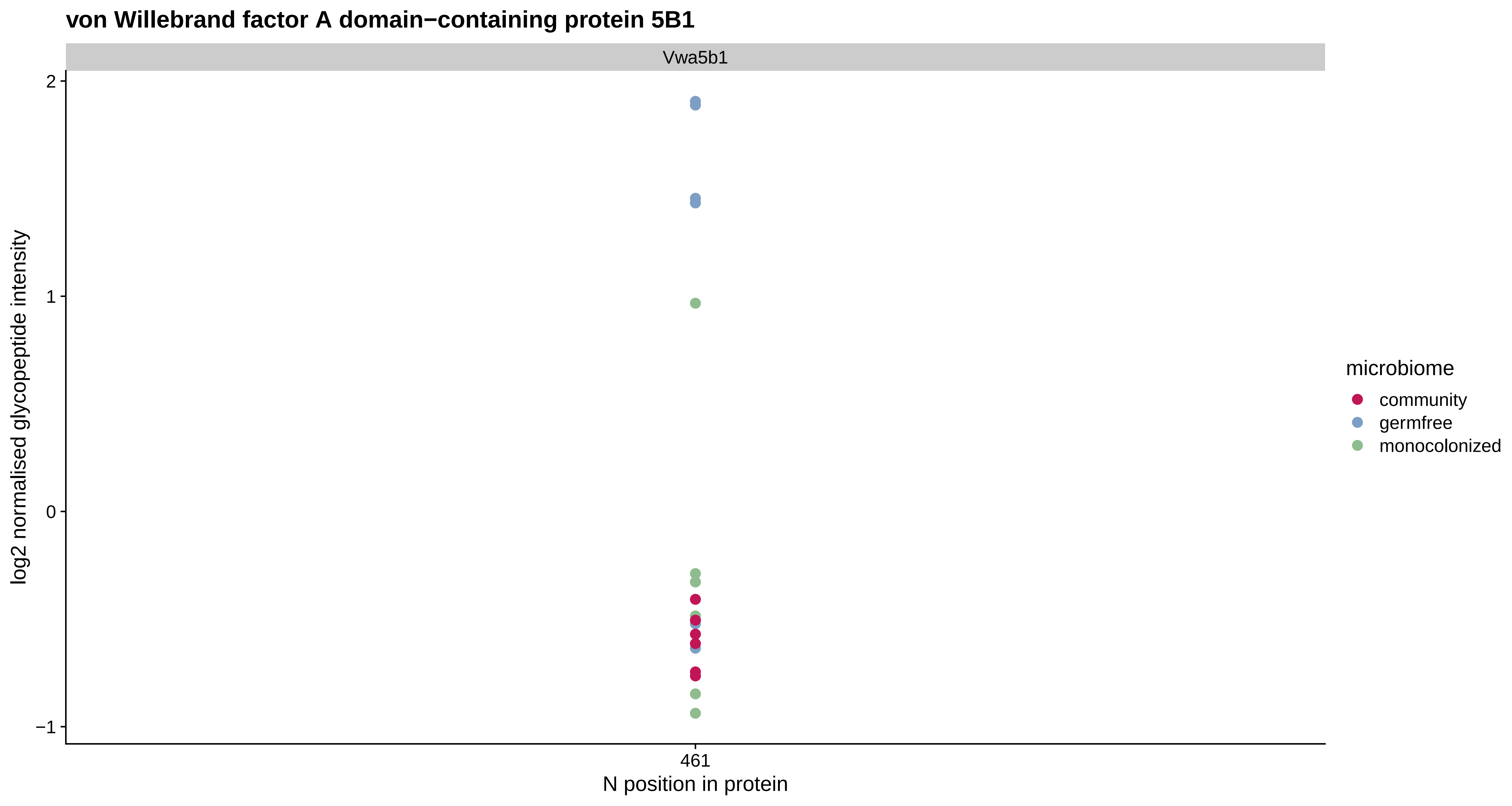

von Willebrand factor A domain−containing protein 5B1
Vwa5b1
log2 normalised glycopeptide intensity
2
1
0
−1
461
N position in protein
microbiome
community
germfree
monocolonized

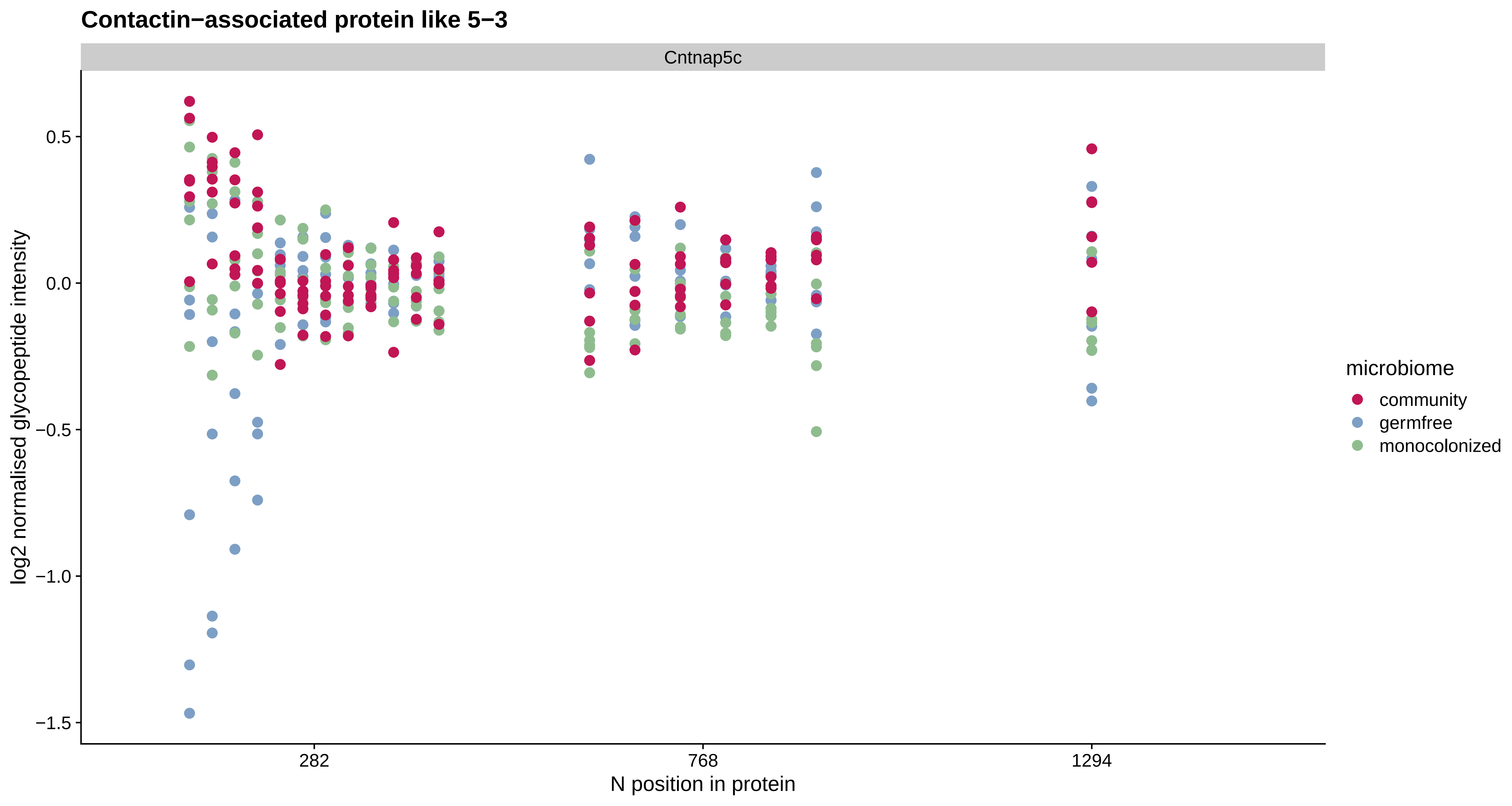
Contactin−associated protein like 5−3
Cntnap5c
log2 normalised glycopeptide intensity
0.5
0.0
−0.5
−1.0
−1.5
282
768
1294
N position in protein
microbiome
community
germfree
monocolonized

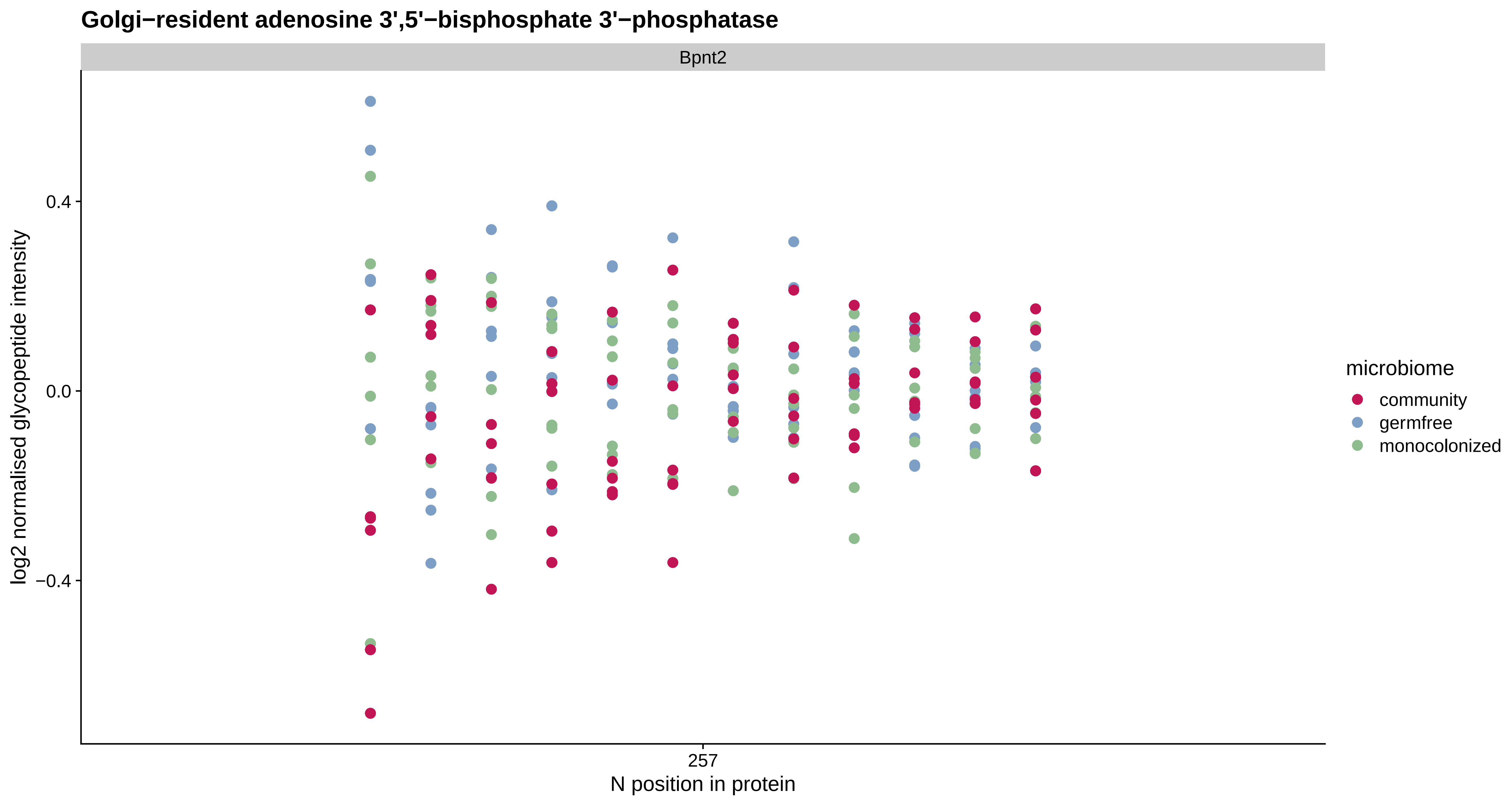
Golgi−resident adenosine 3',5'−bisphosphate 3'−phosphatase
Bpnt2
0.4
0.0
−0.4
log2 normalised glycopeptide intensity
257
N position in protein
microbiome
community
germfree
monocolonized

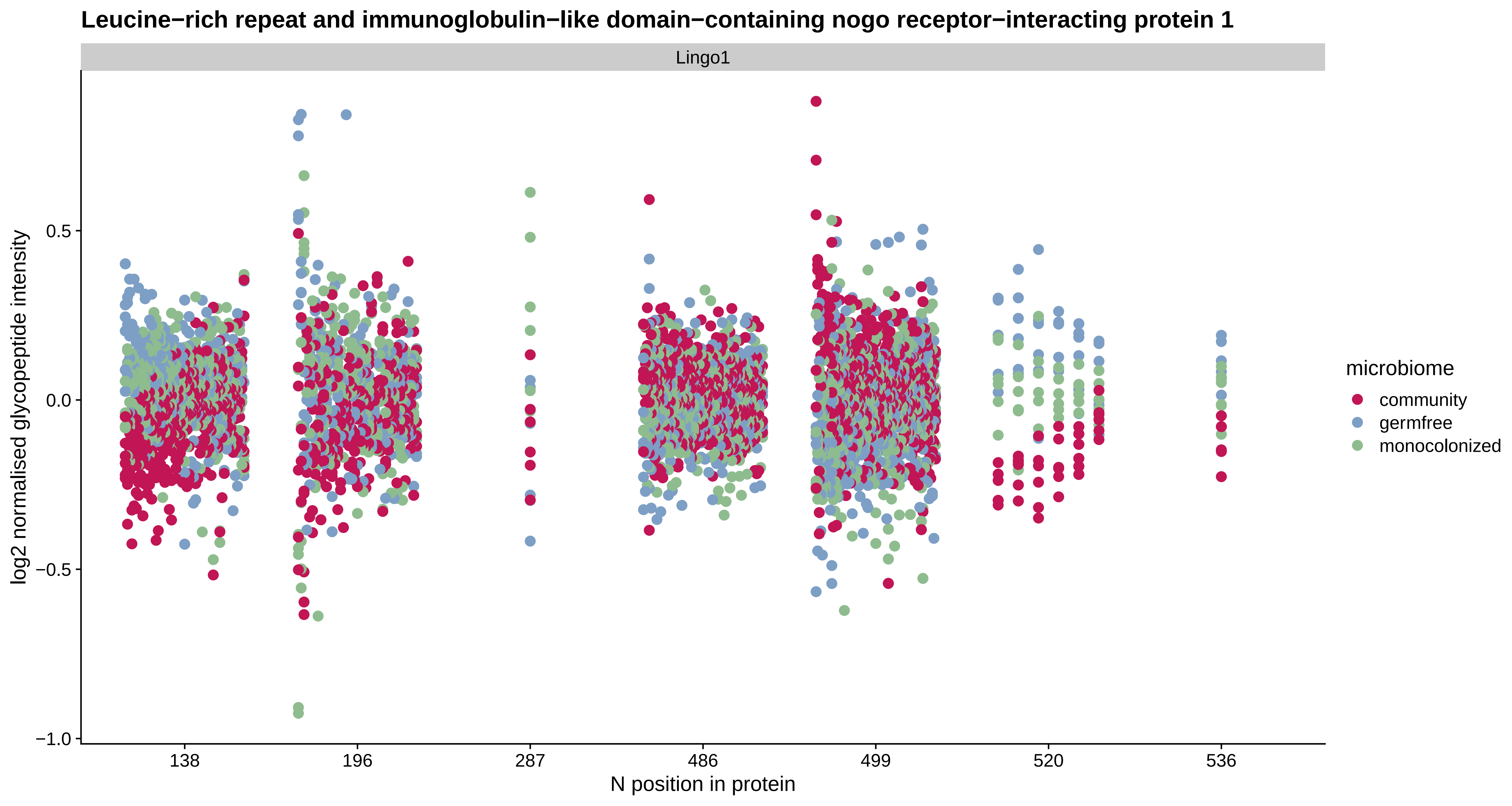
Leucine−rich repeat and immunoglobulin−like domain−containing nogo receptor−interacting protein 1
Lingo1
log2 normalised glycopeptide intensity
0.5
0.0
−0.5
−1.0
138
196
287
486
499
520
536
N position in protein
microbiome
community
germfree
monocolonized

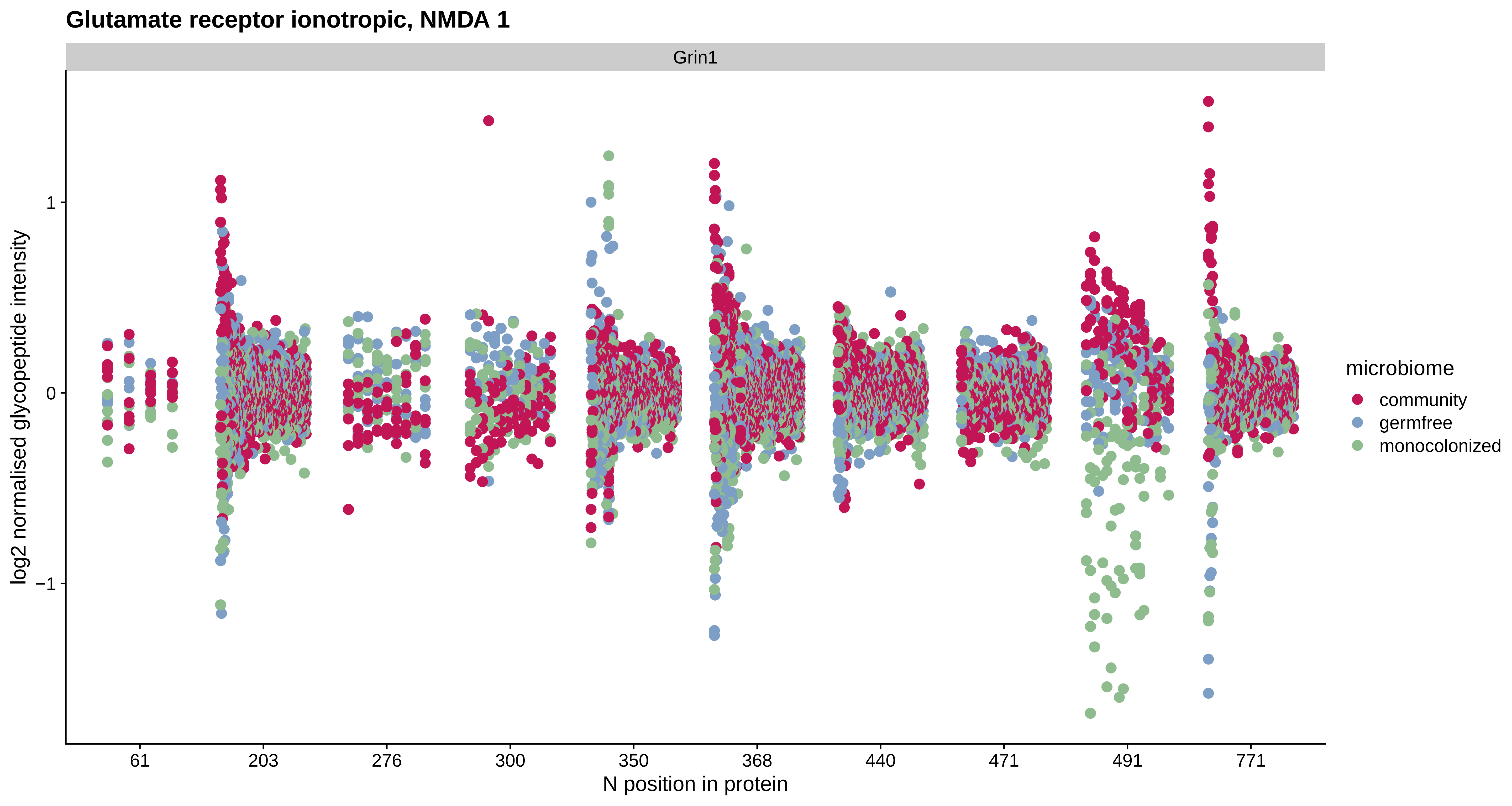
Glutamate receptor ionotropic, NMDA 1
Grin1
log2 normalised glycopeptide intensity
1
0
−1
61
203
276
300
350
368
440
471
491
771
N position in protein
microbiome
community
germfree
monocolonized

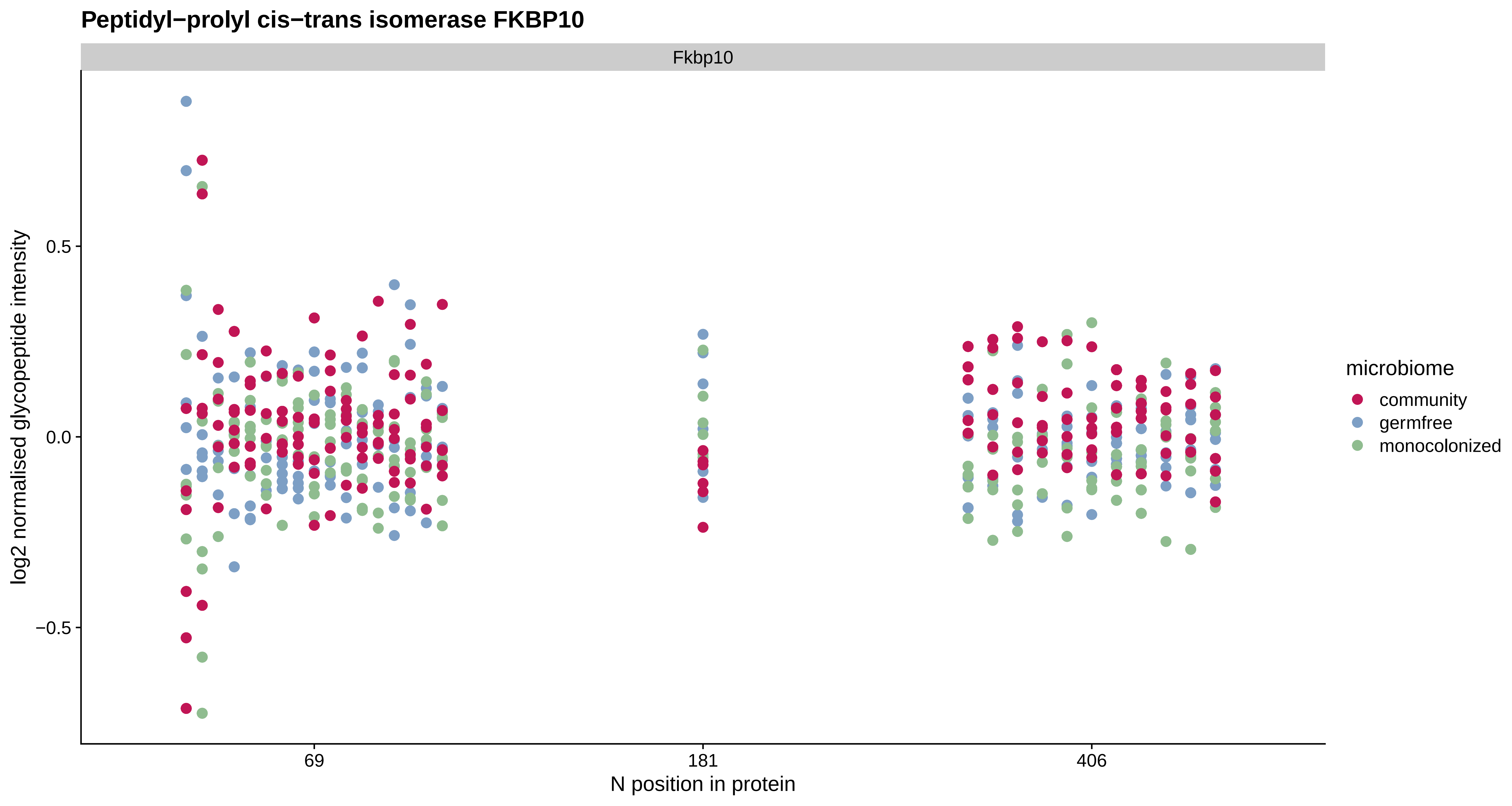
Peptidyl−prolyl cis−trans isomerase FKBP10
Fkbp10
log2 normalised glycopeptide intensity
0.5
0.0
−0.5
69
181
406
N position in protein
microbiome
community
germfree
monocolonized

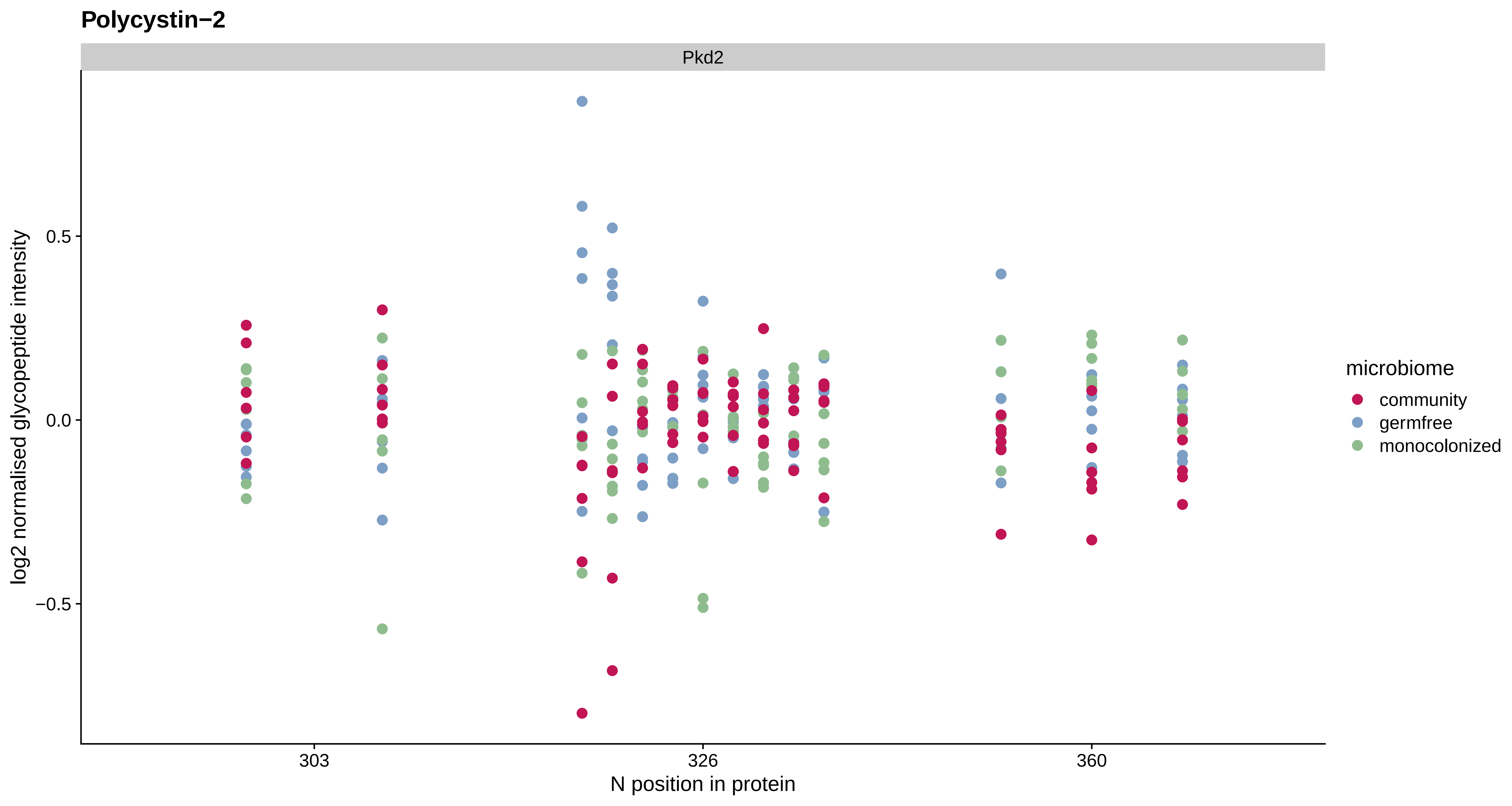
Polycystin−2
Pkd2
0.5
0.0
−0.5
log2 normalised glycopeptide intensity
303
326
360
N position in protein
microbiome
community
germfree
monocolonized

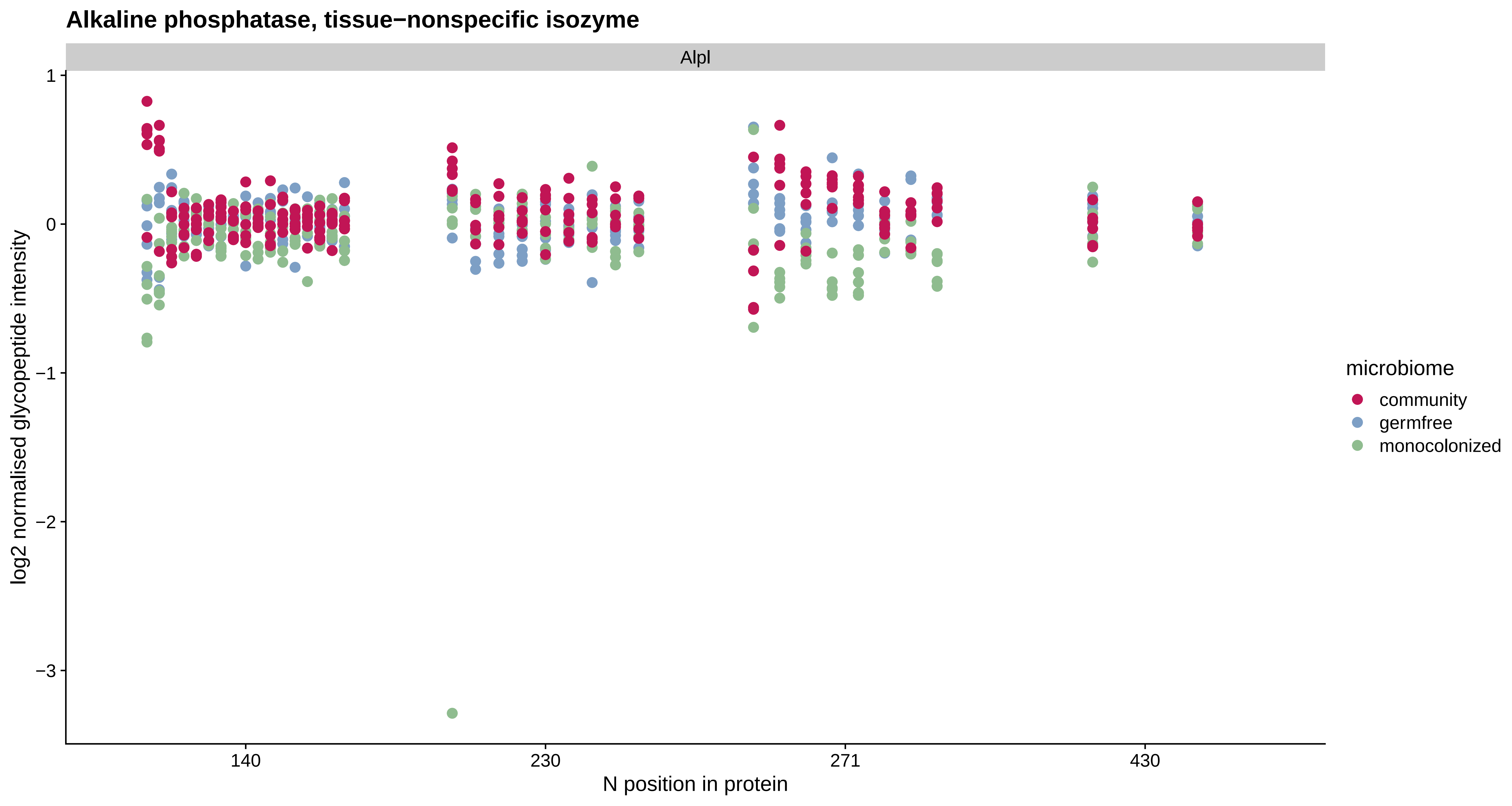
Alkaline phosphatase, tissue−nonspecific isozyme
Alpl
1
0
−1
−2
−3
log2 normalised glycopeptide intensity
140
230
271
430
N position in protein
microbiome
community
germfree
monocolonized

#### Amyloid beta precursor like protein 1

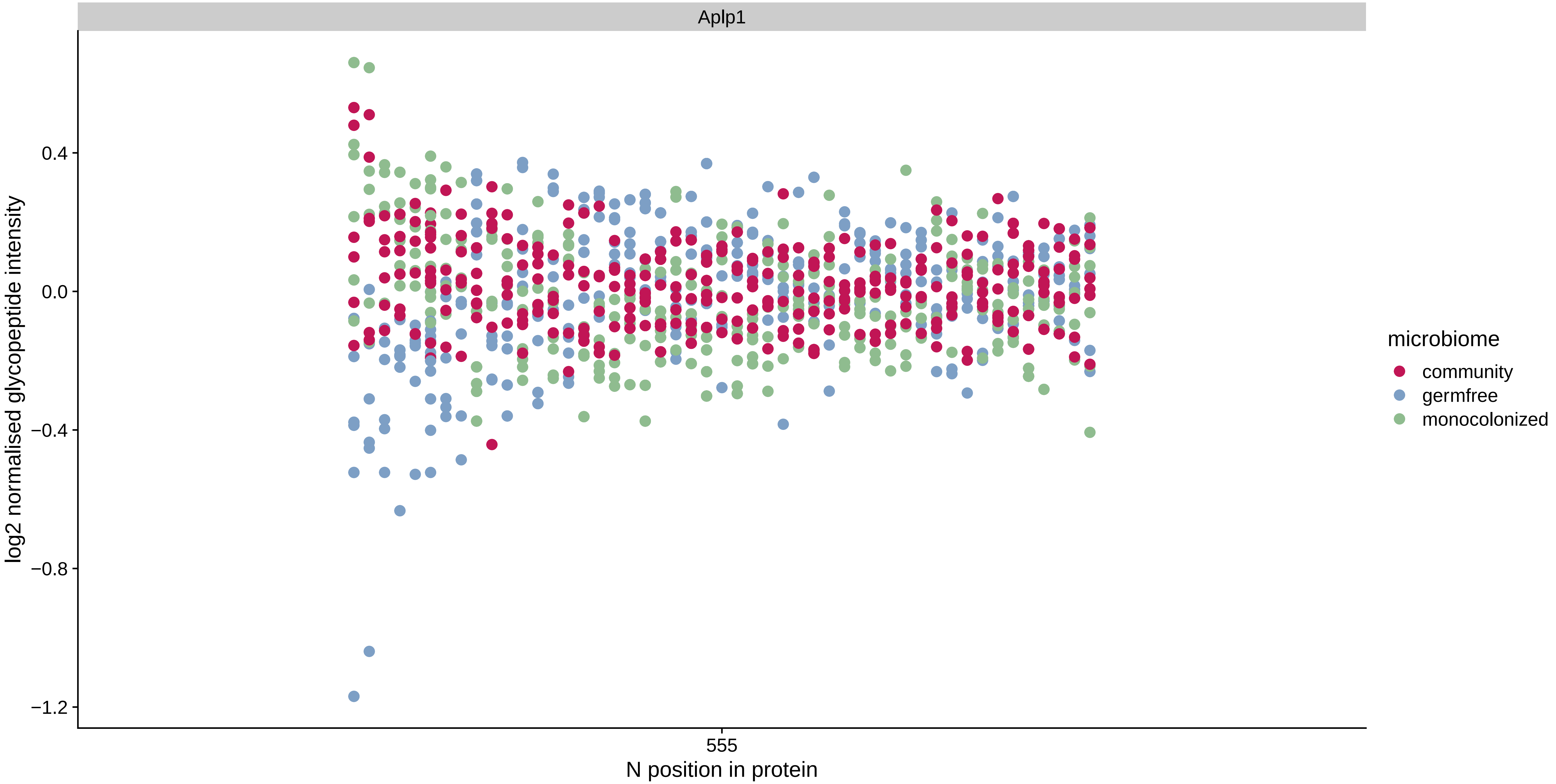

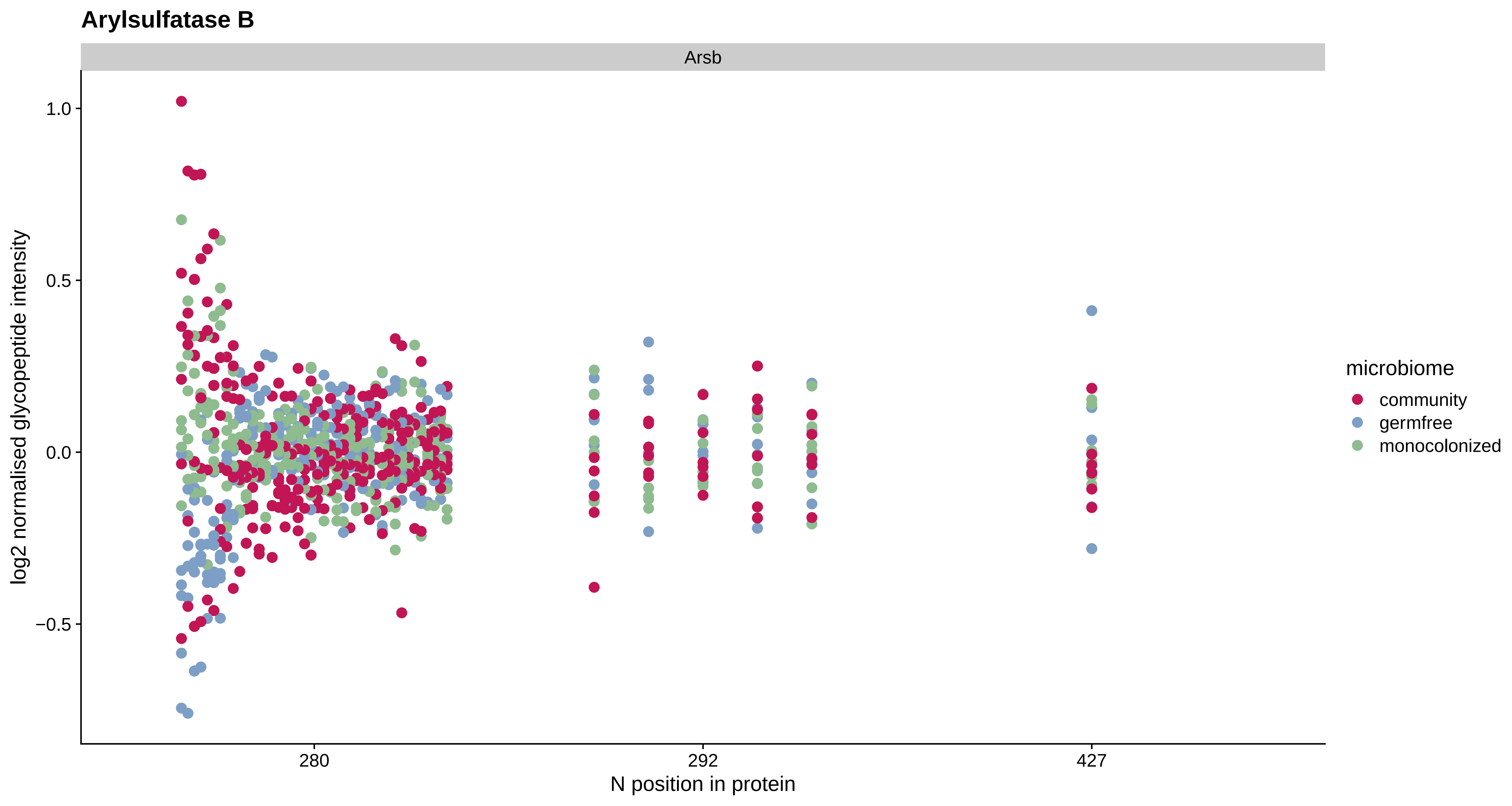
Arylsulfatase B
Arsb
1.0
0.5
0.0
−0.5
log2 normalised glycopeptide intensity
280
292
427
N position in protein
microbiome
community
germfree
monocolonized

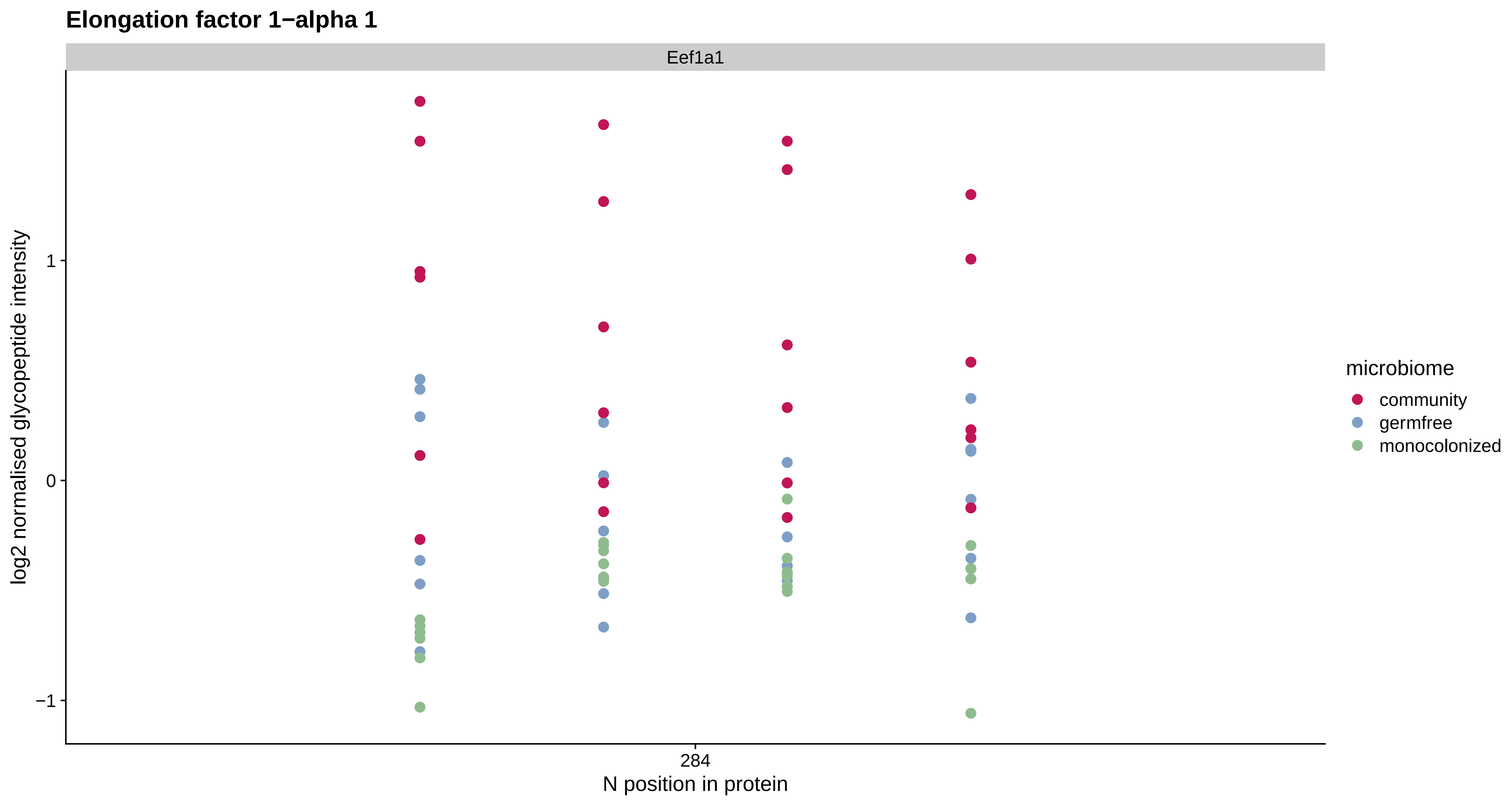
Elongation factor 1−alpha 1
Eef1a1
log2 normalised glycopeptide intensity
1
0
−1
284
N position in protein
microbiome
community
germfree
monocolonized

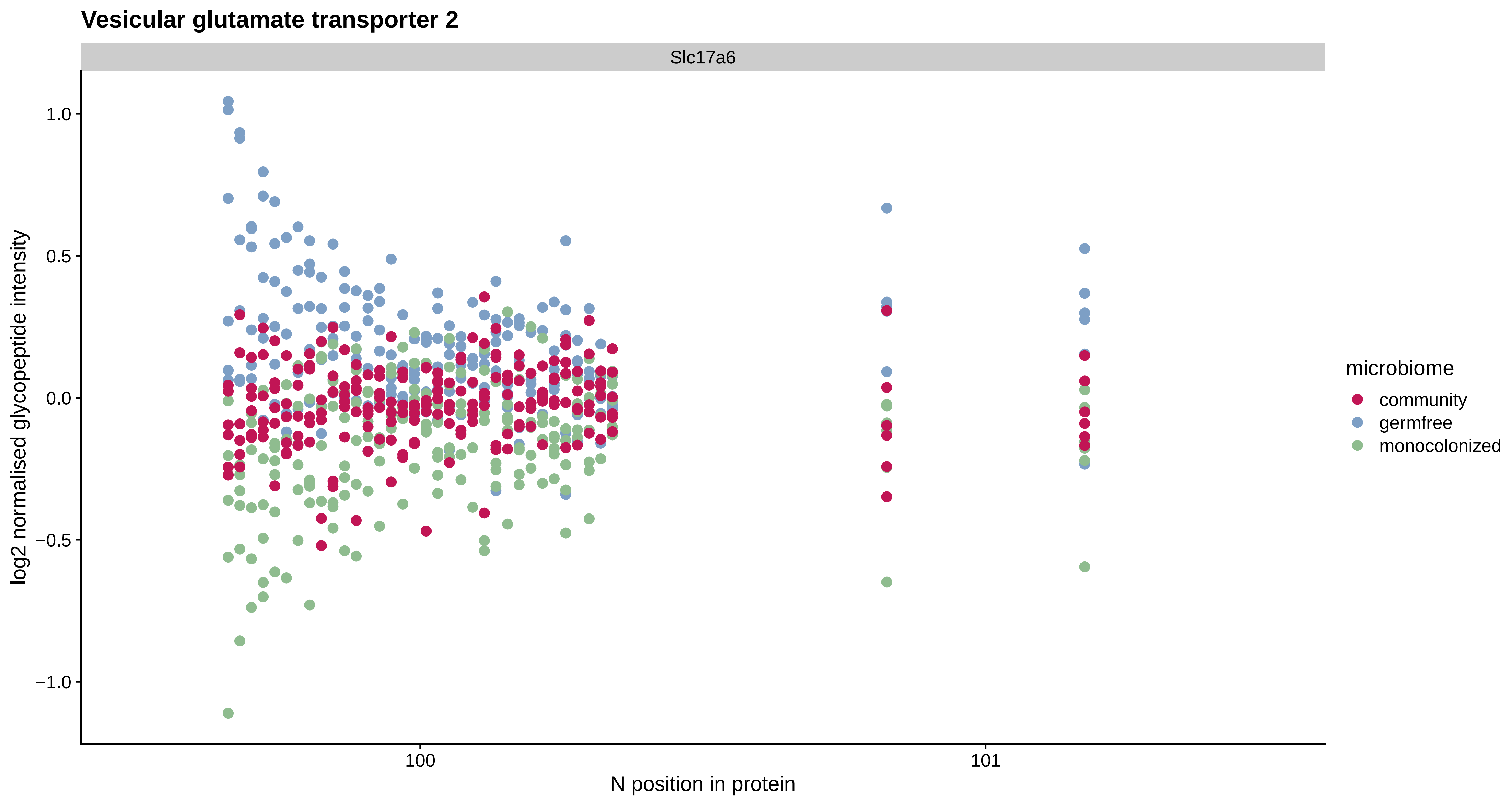
Vesicular glutamate transporter 2
Slc17a6
log2 normalised glycopeptide intensity
1.0
0.5
0.0
−0.5
−1.0
100
101
N position in protein
microbiome
community
germfree
monocolonized

#### Leucine−rich repeat serine/threonine−protein kinase 2

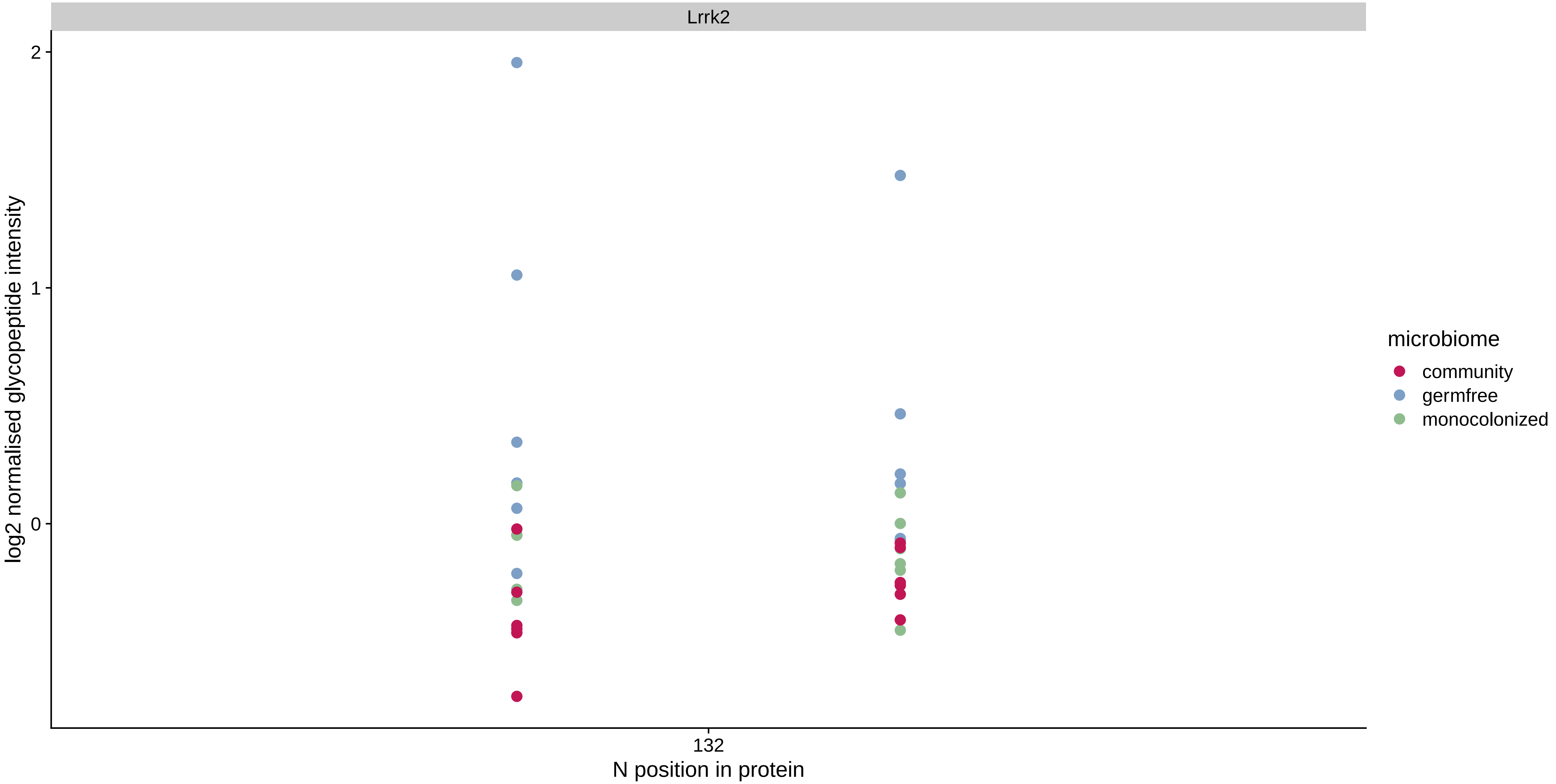

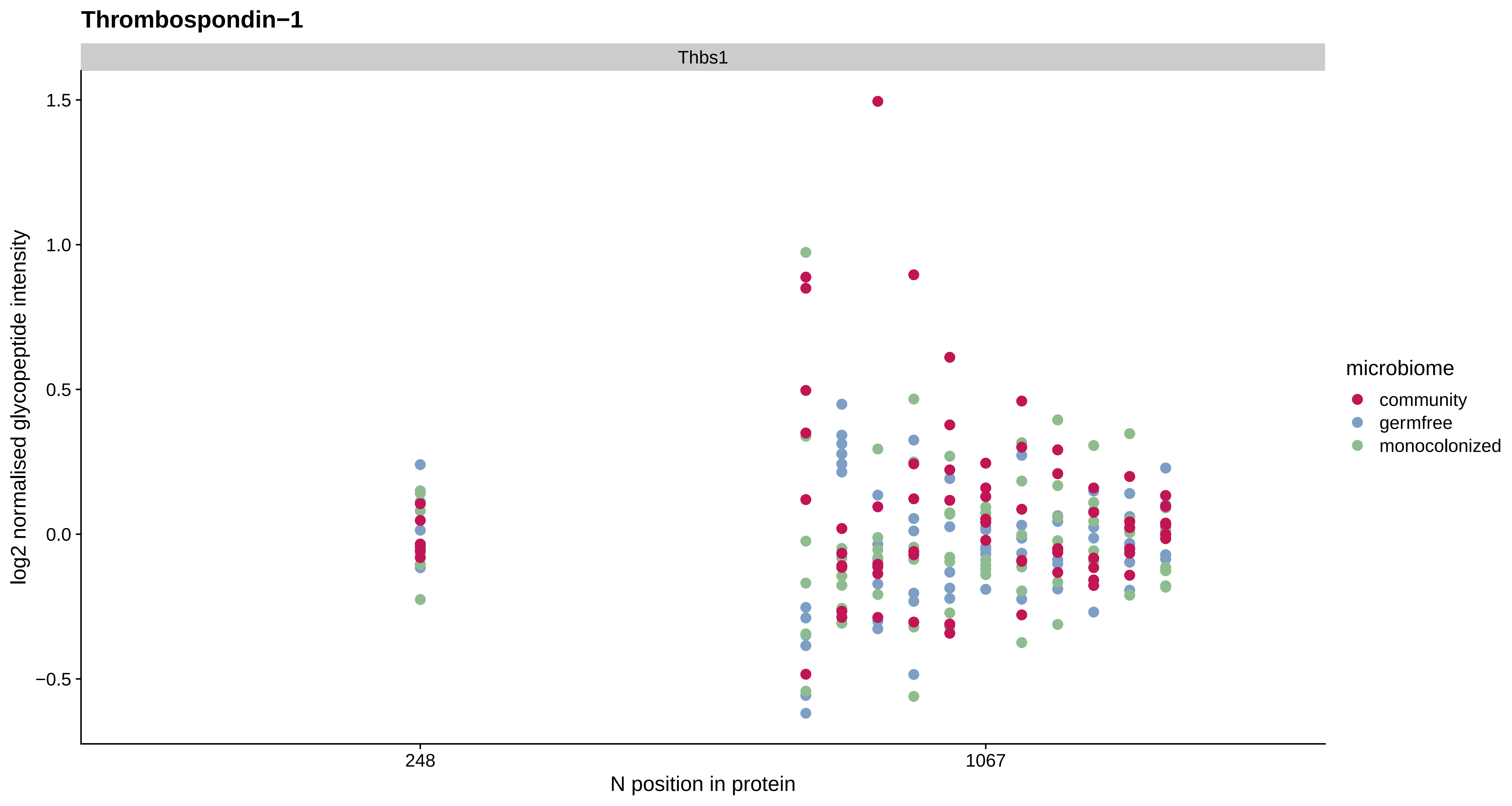
Thrombospondin−1
Thbs1
1.5
1.0
0.5
0.0
−0.5
log2 normalised glycopeptide intensity
248
1067
N position in protein
microbiome
community
germfree
monocolonized

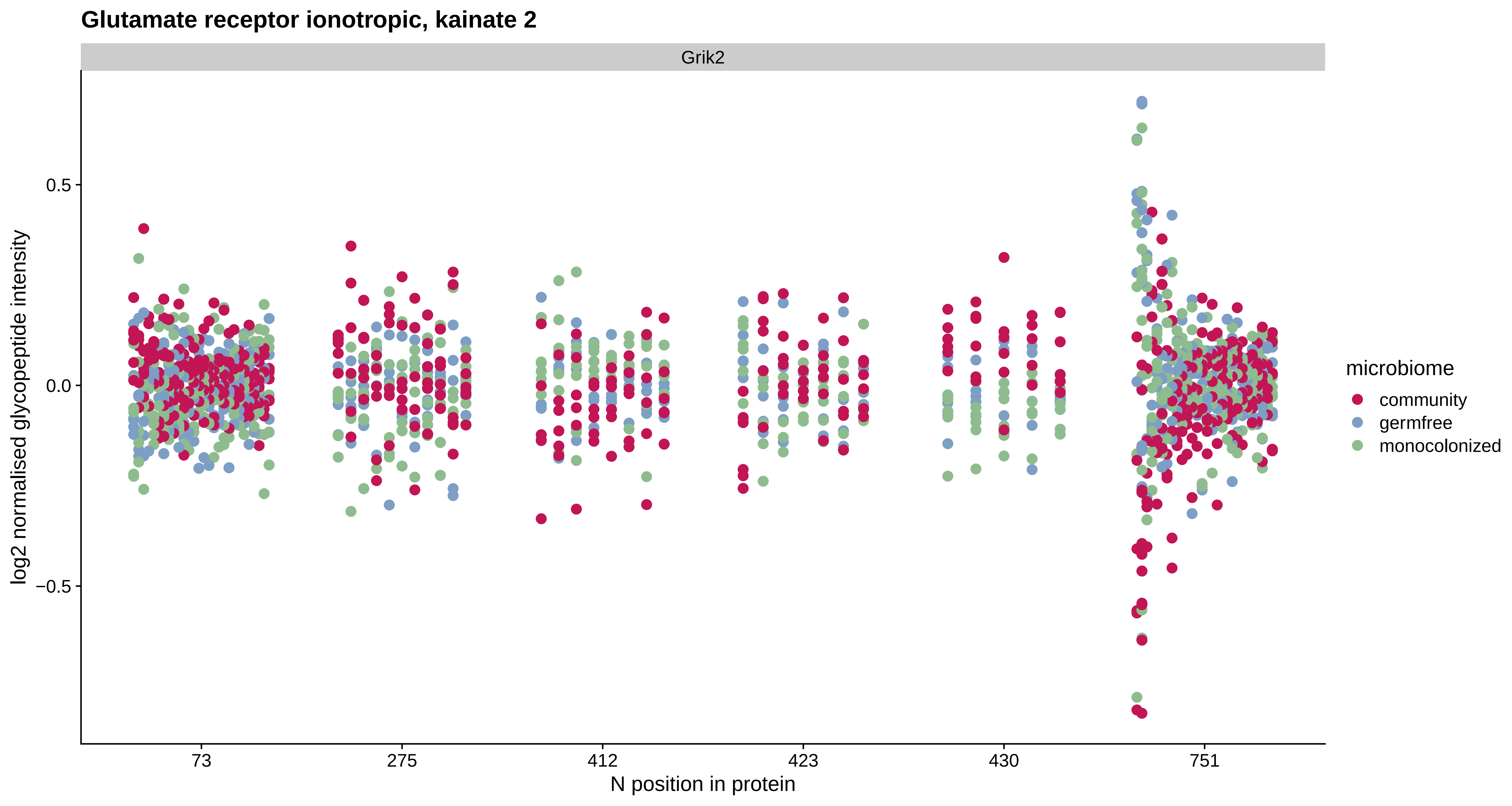
Glutamate receptor ionotropic, kainate 2
Grik2
log2 normalised glycopeptide intensity
0.5
0.0
−0.5
73
275
412
423
430
751
N position in protein
microbiome
community
germfree
monocolonized

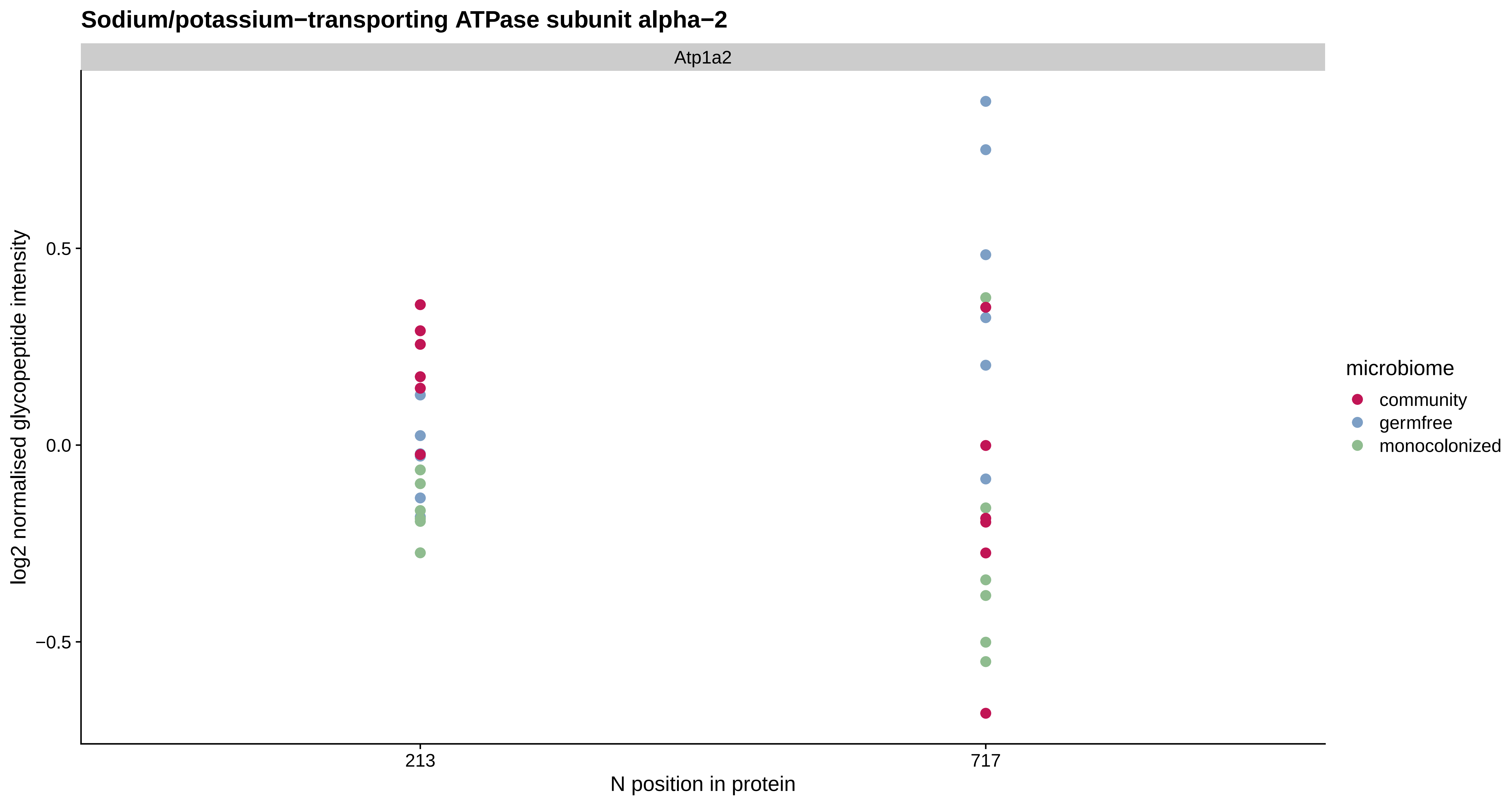
Sodium/potassium−transporting ATPase subunit alpha−2
Atp1a2
log2 normalised glycopeptide intensity
0.5
0.0
−0.5
213
717
N position in protein
microbiome
community
germfree
monocolonized

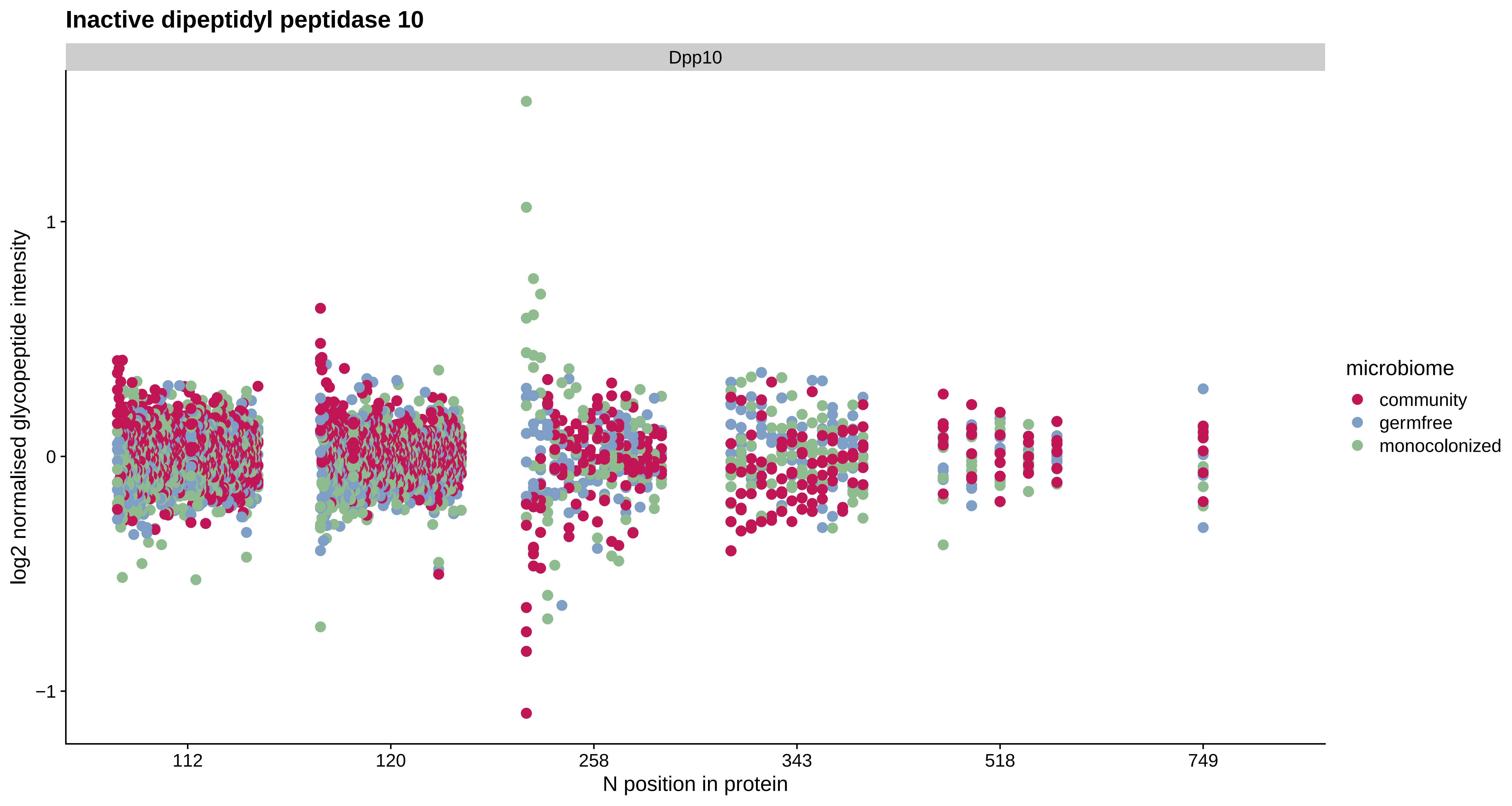
Inactive dipeptidyl peptidase 10
Dpp10
log2 normalised glycopeptide intensity
1
0
−1
112
120
258
343
518
749
N position in protein
microbiome
community
germfree
monocolonized

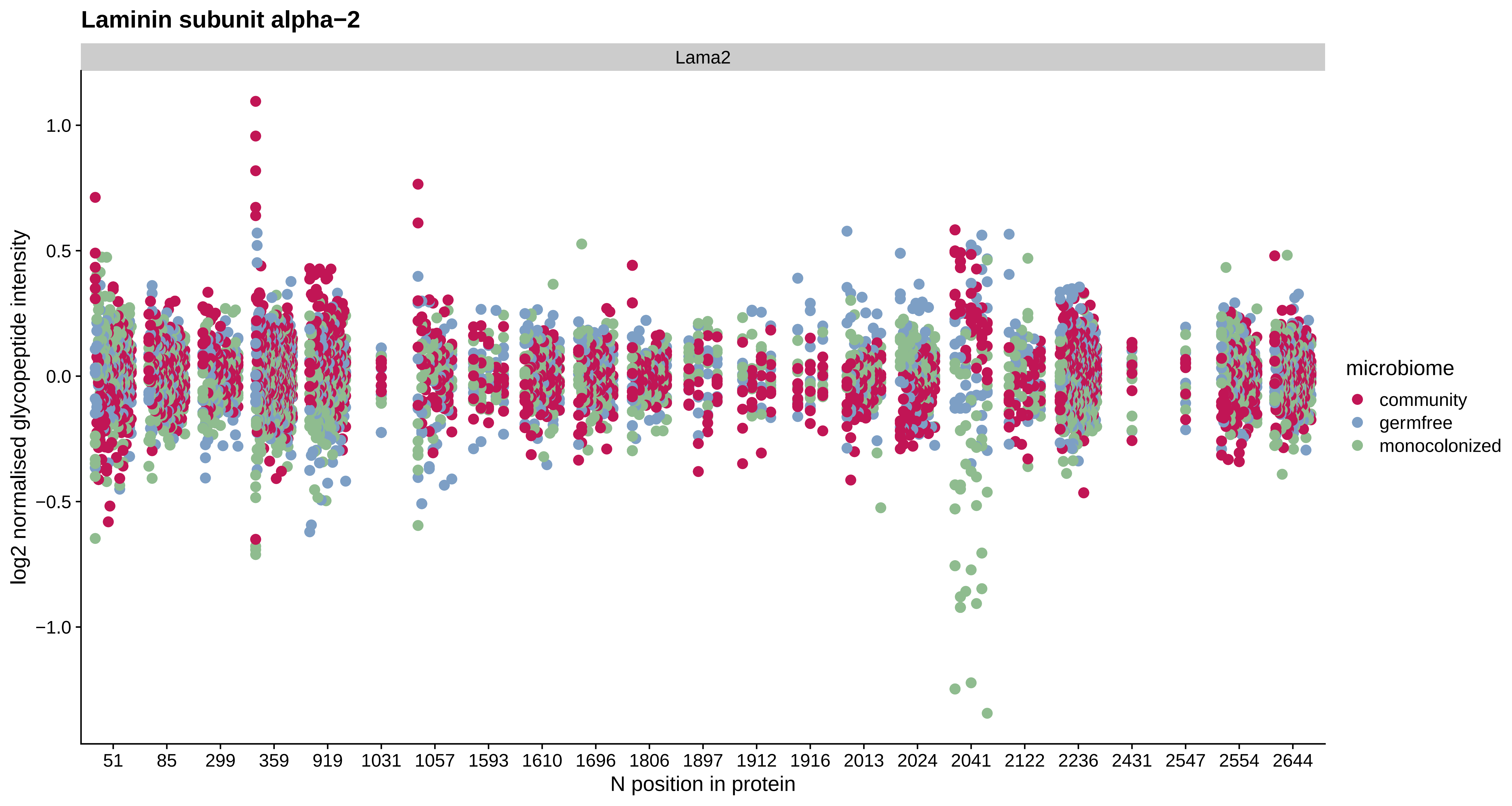

Laminin subunit alpha−2
Lama2
log2 normalised glycopeptide intensity
1.0
0.5
0.0
−0.5
−1.0
51
85
299
359
919
1031
1057
1593
1610
1696
1806
1897
1912
1916
2013
2024
2041
2122
2236
2431
2547
2554
2644
N position in protein
microbiome
community
germfree
monocolonized

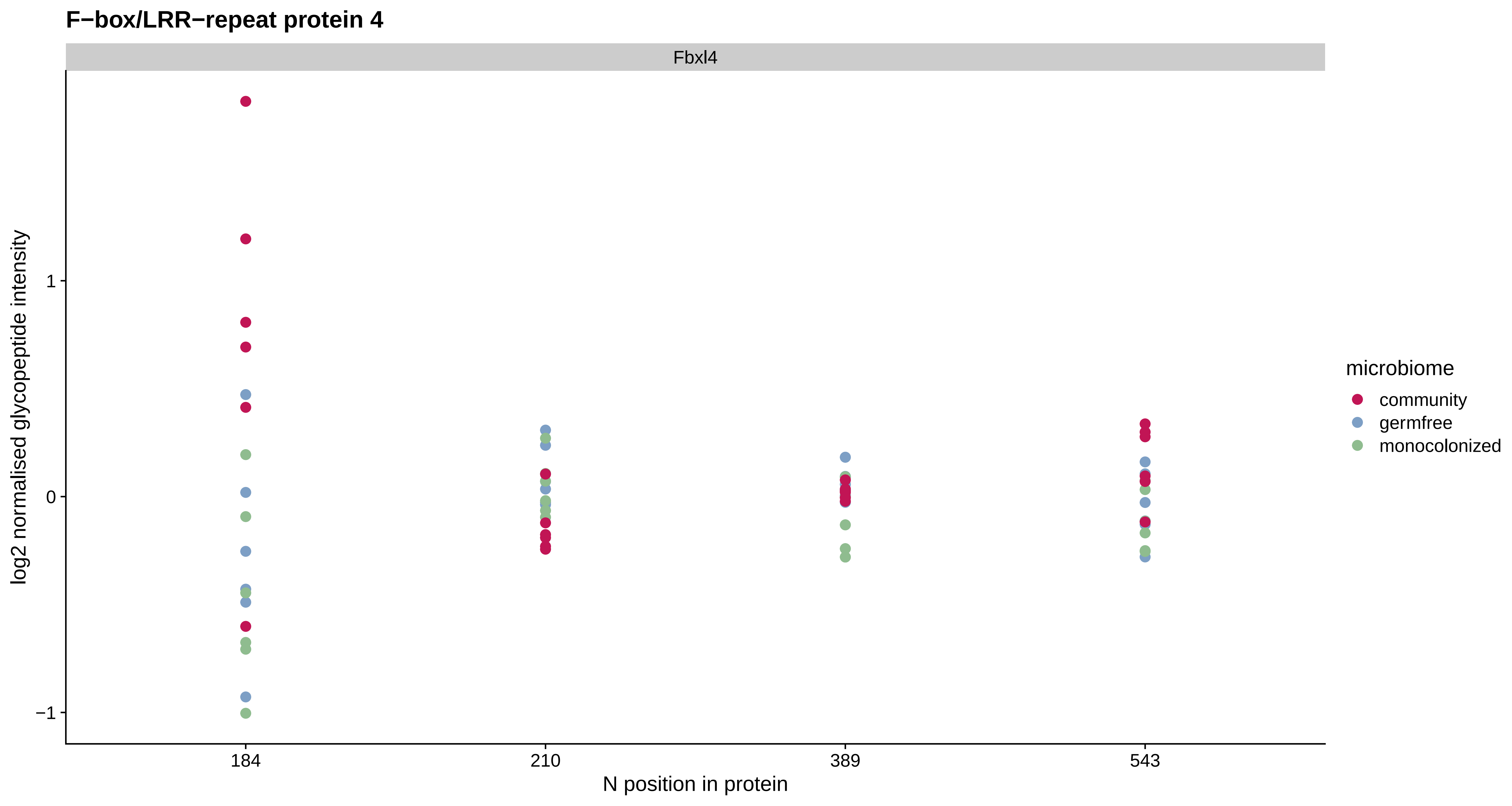
F−box/LRR−repeat protein 4
Fbxl4
log2 normalised glycopeptide intensity
1
0
−1
184
210
389
543
N position in protein
microbiome
community
germfree
monocolonized

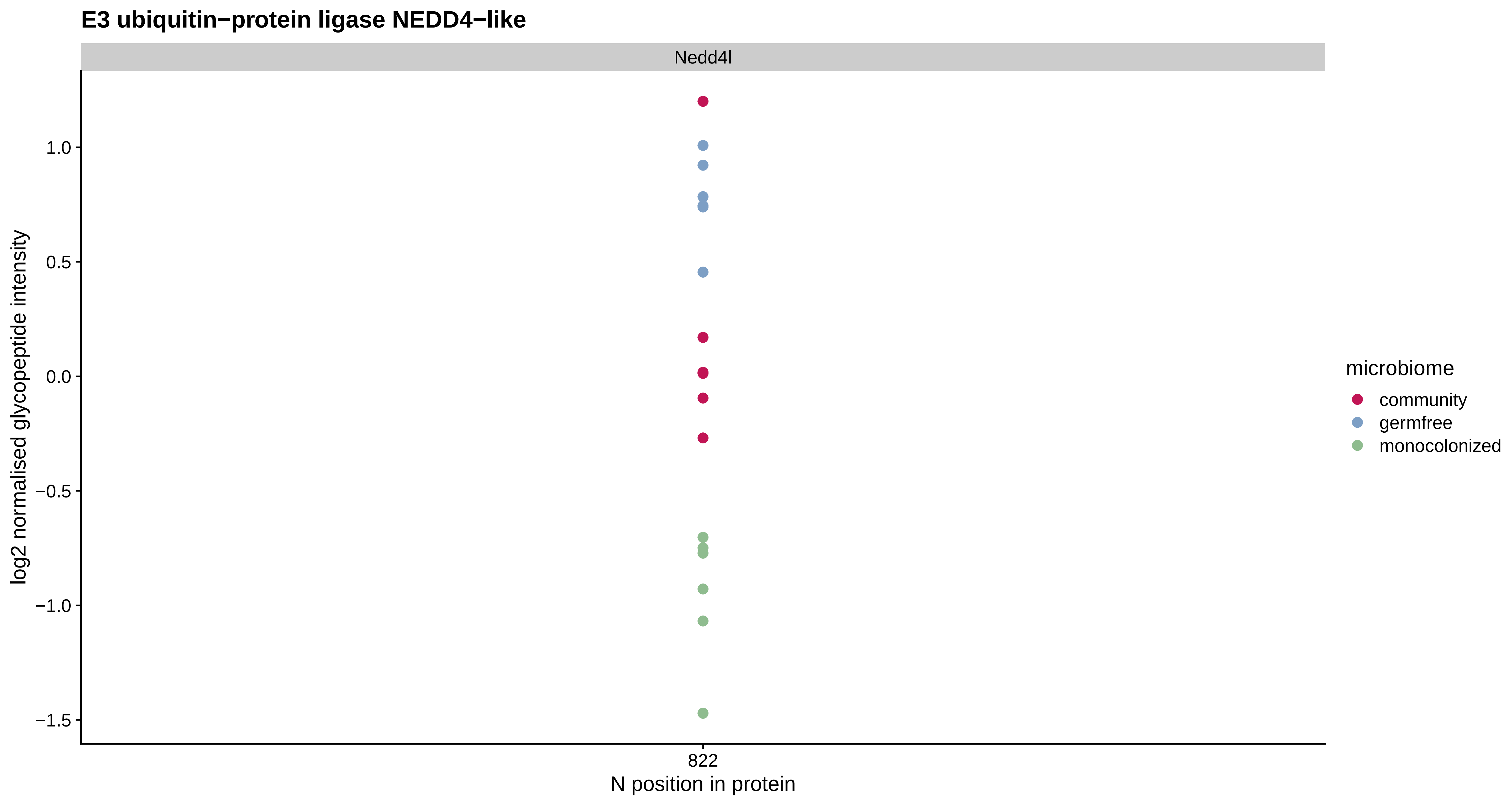

E3 ubiquitin−protein ligase NEDD4−like
Nedd4l
log2 normalised glycopeptide intensity
1.0
0.5
0.0
−0.5
−1.0
−1.5
822
N position in protein
microbiome
community
germfree
monocolonized

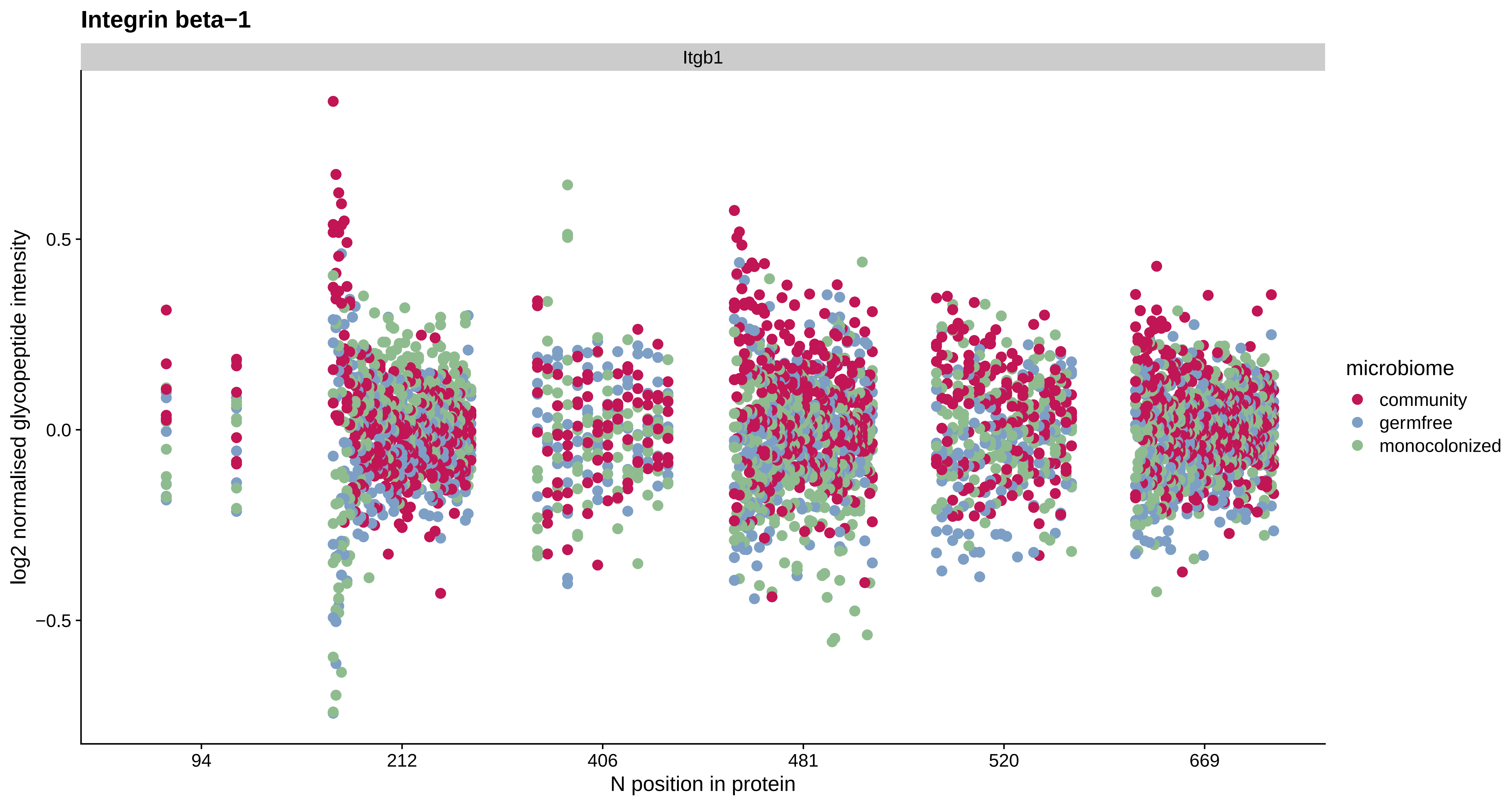
Integrin beta−1
Itgb1
log2 normalised glycopeptide intensity
0.5
0.0
−0.5
94
212
406
481
520
669
N position in protein
microbiome
community
germfree
monocolonized

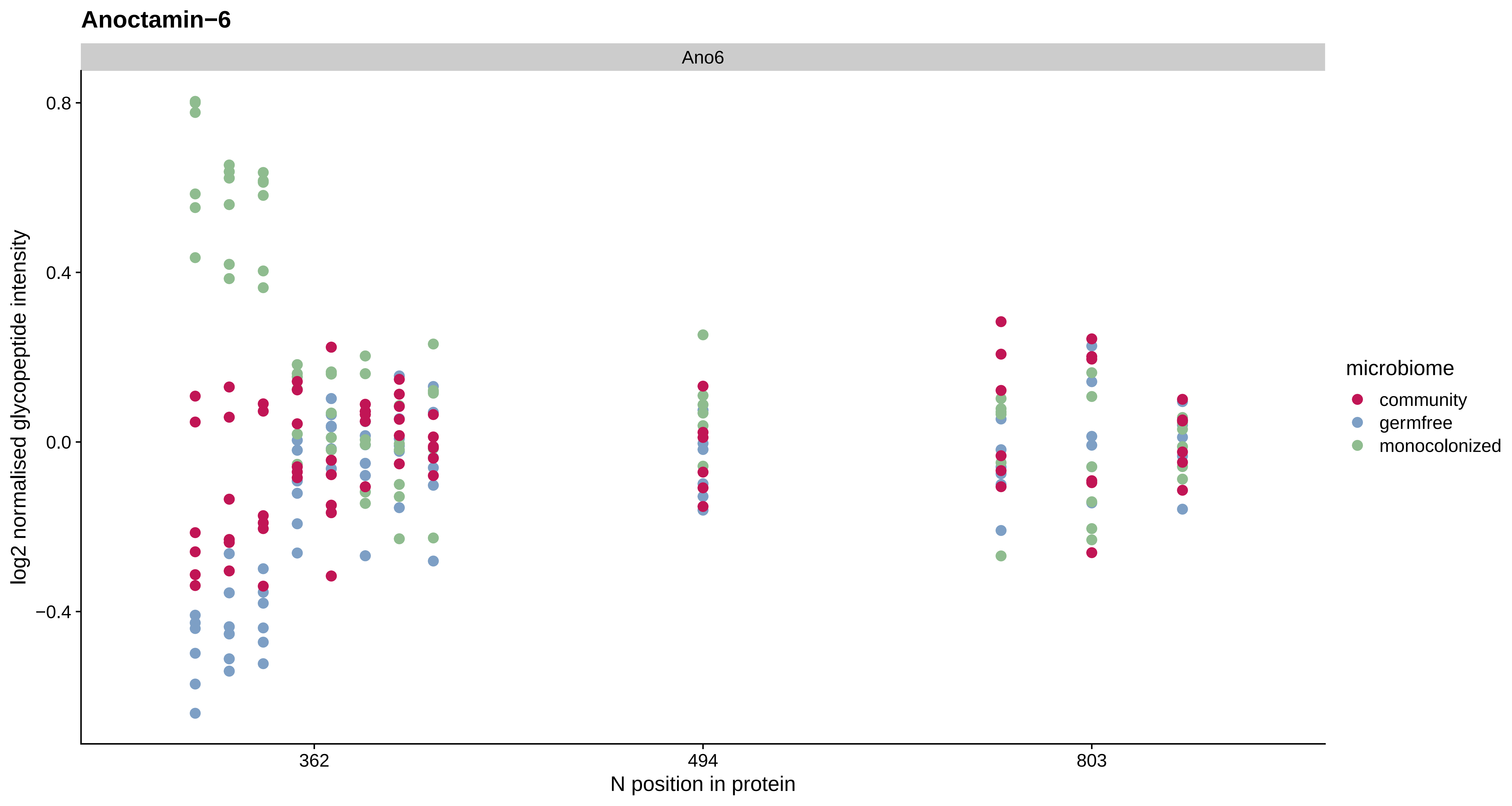

Anoctamin−6
Ano6
0.8
0.4
0.0
−0.4
log2 normalised glycopeptide intensity
362
494
803
N position in protein
microbiome
community
germfree
monocolonized

#### Mitochondrial glutamate carrier 1

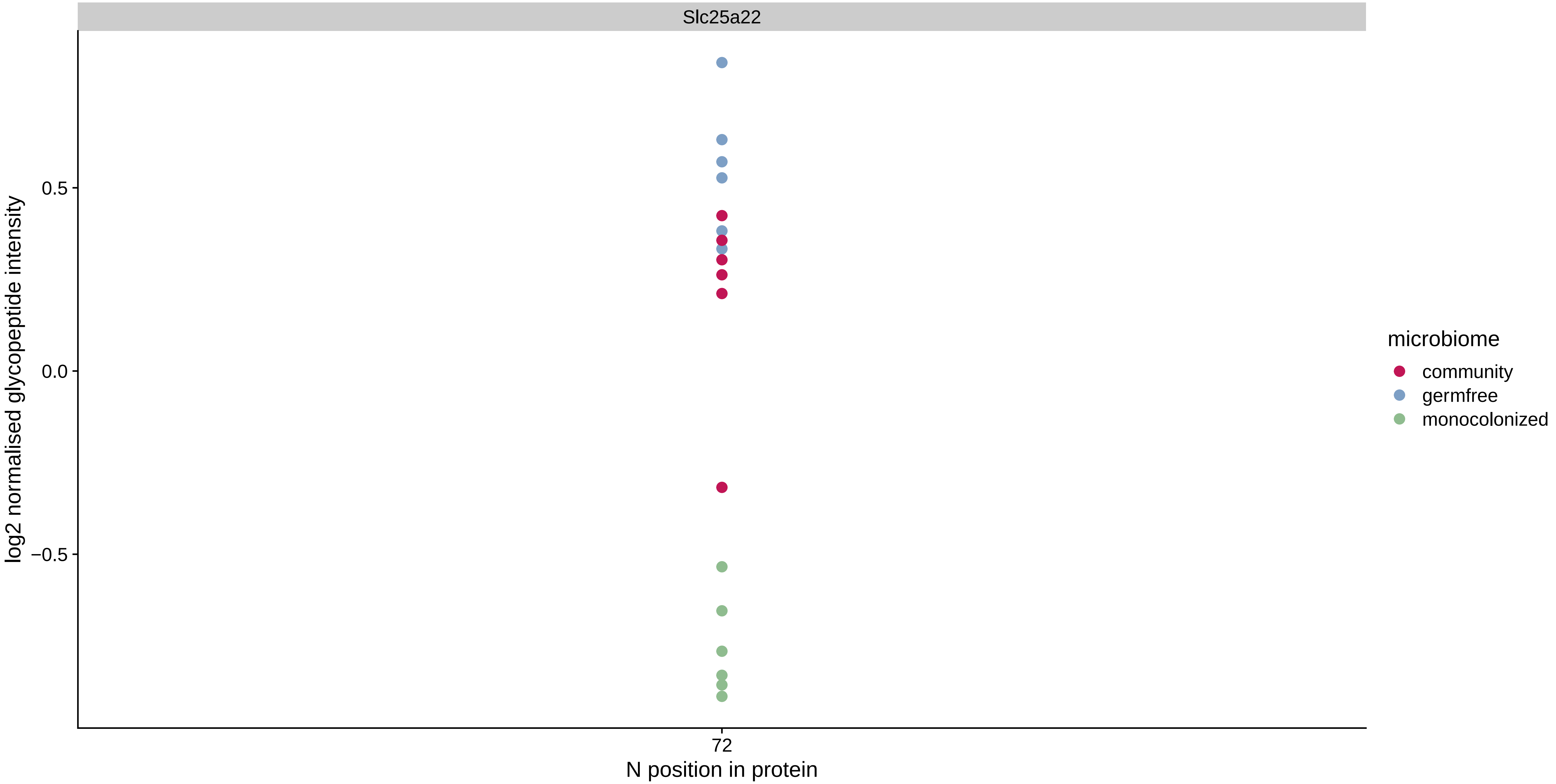

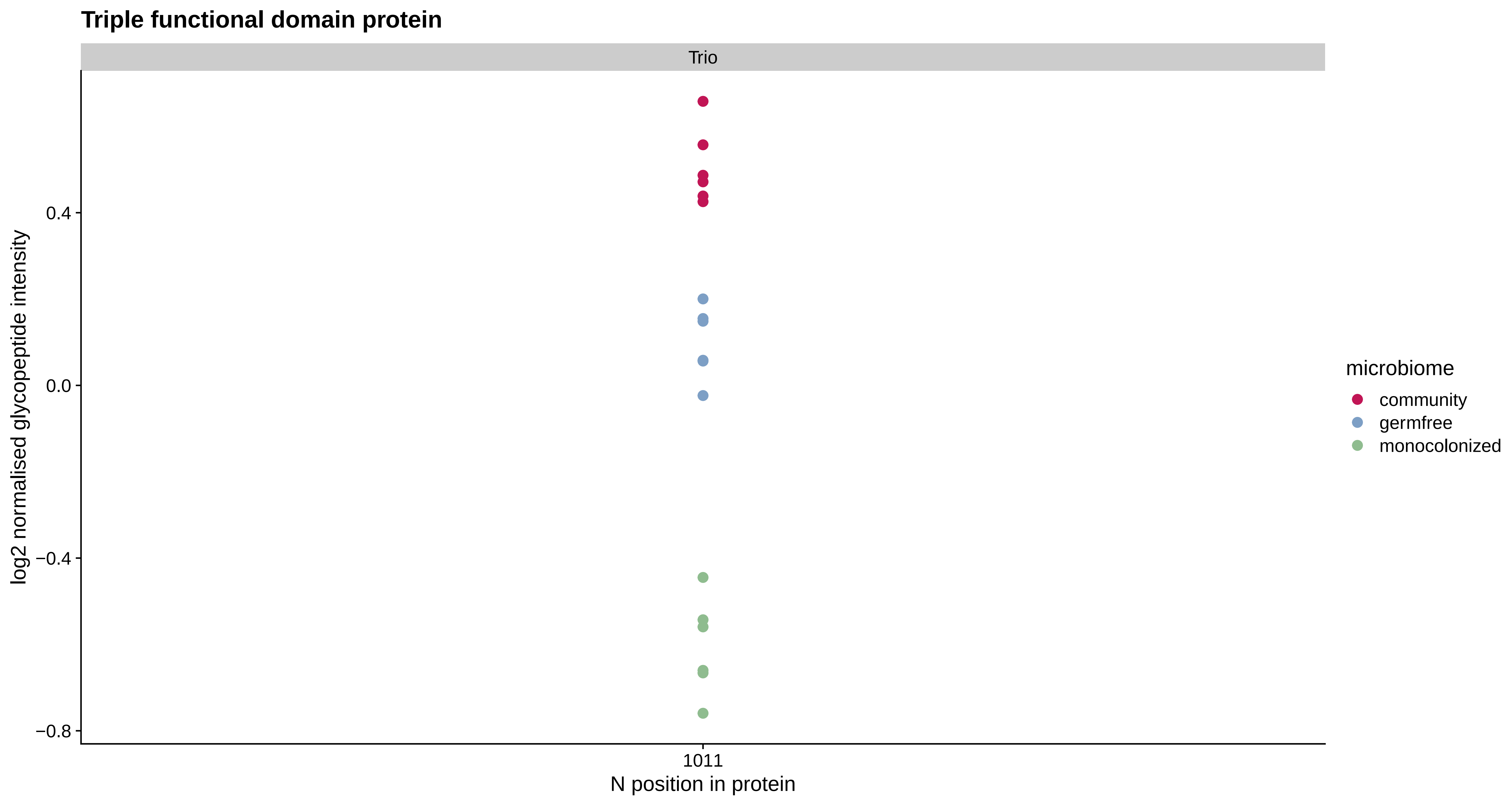

Triple functional domain protein
Trio
log2 normalised glycopeptide intensity
0.4
0.0
−0.4
−0.8
1011
N position in protein
microbiome
community
germfree
monocolonized

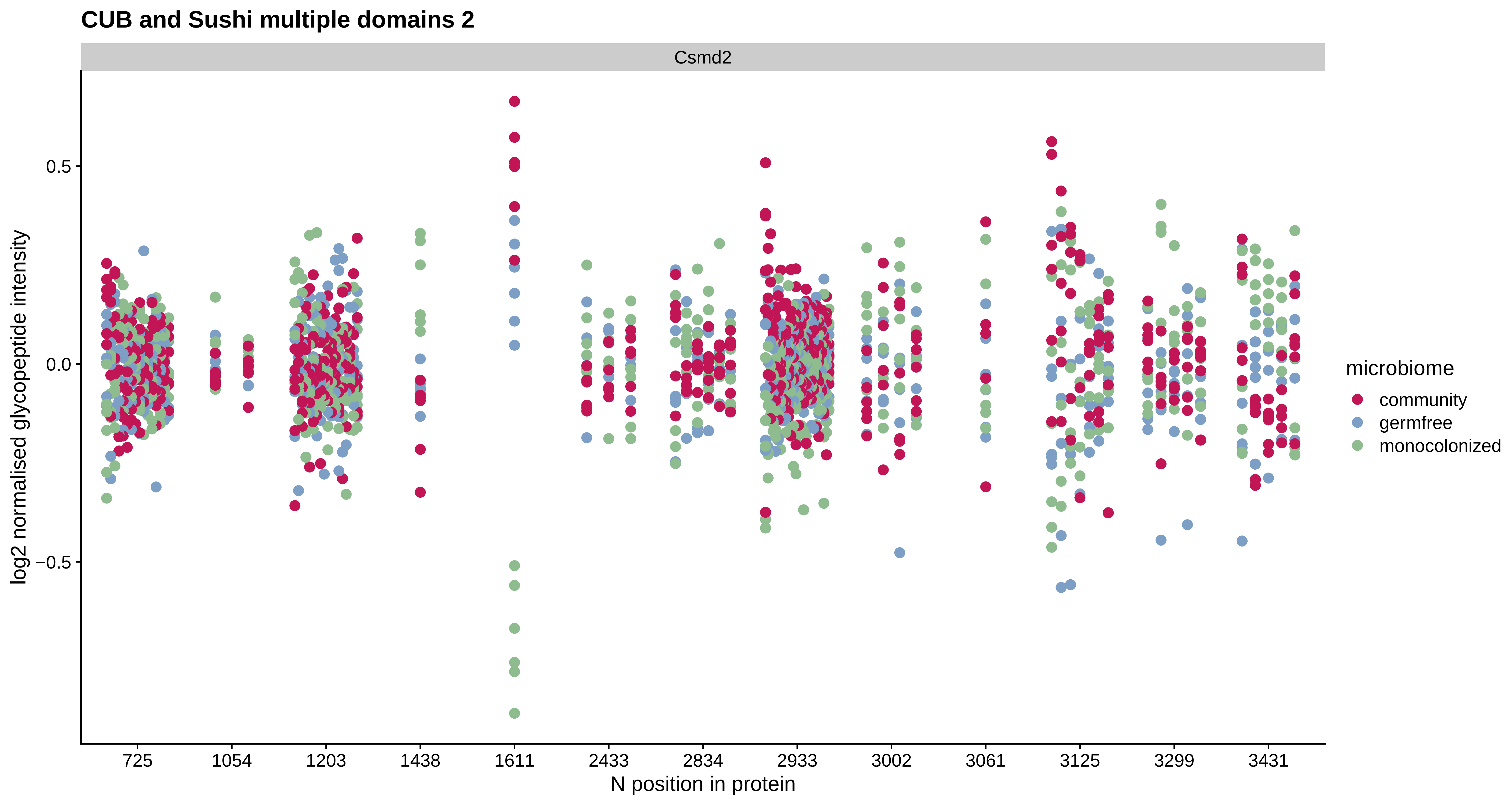

CUB and Sushi multiple domains 2
Csmd2
log2 normalised glycopeptide intensity
0.5
0.0
−0.5
725
1054
1203
1438
1611
2433
2834
2933
3002
3061
3125
3299
3431
N position in protein
microbiome
community
germfree
monocolonized

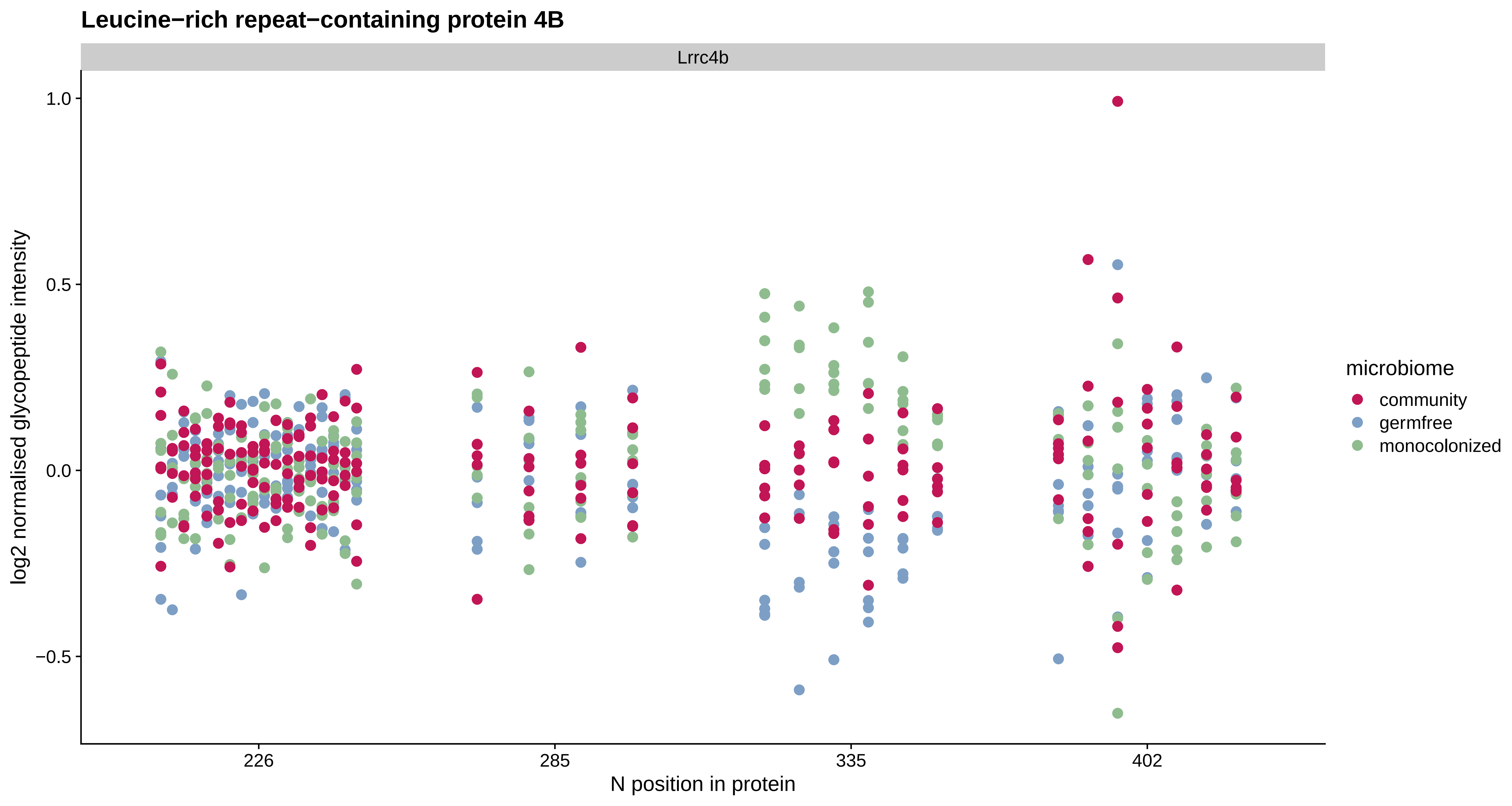

Leucine−rich repeat−containing protein 4B
Lrrc4b
log2 normalised glycopeptide intensity
1.0
0.5
0.0
−0.5
226
285
335
402
N position in protein
microbiome
community
germfree
monocolonized

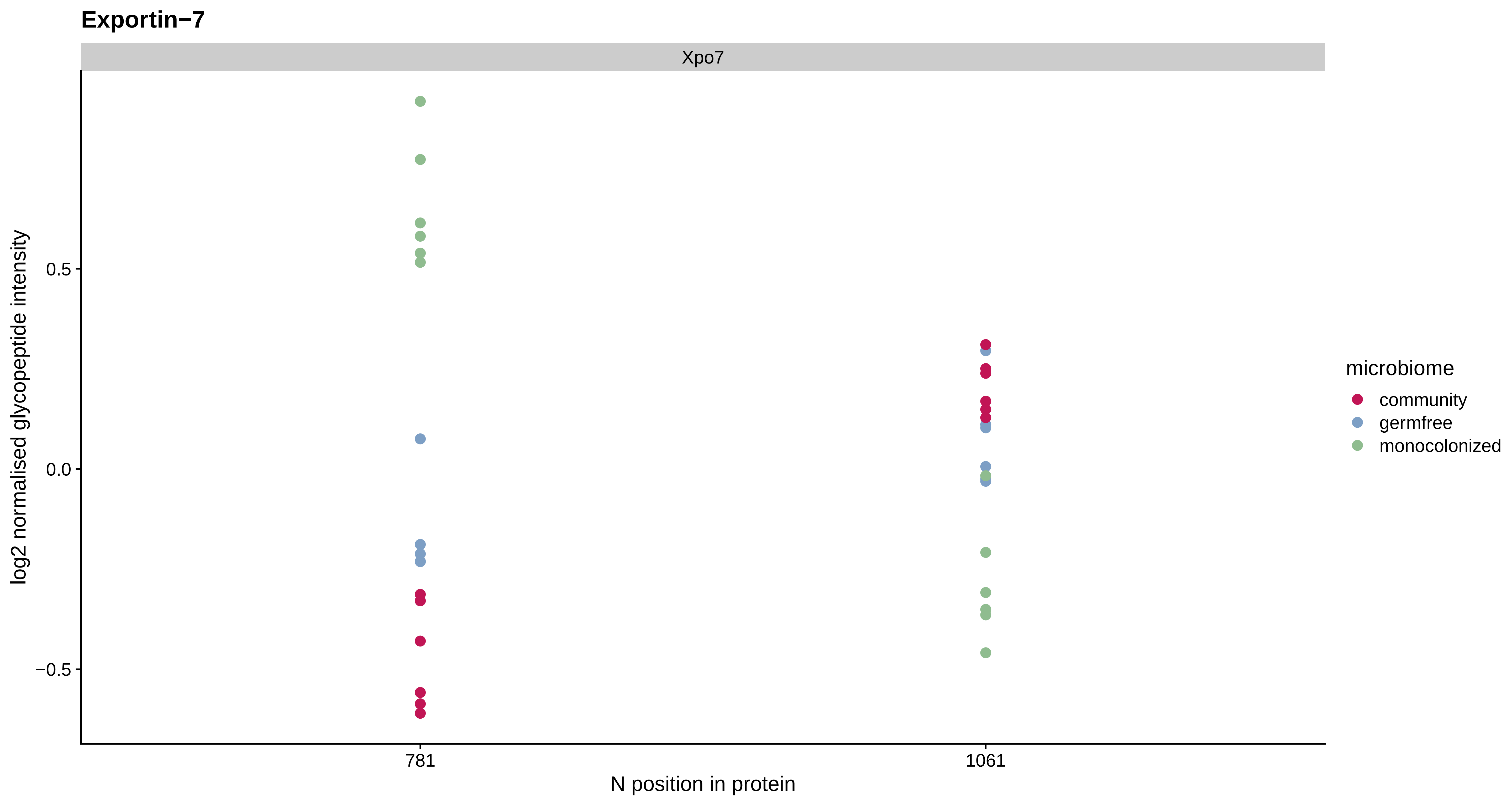
Exportin−7
Xpo7
log2 normalised glycopeptide intensity
0.5
0.0
−0.5
781
1061
N position in protein
microbiome
community
germfree
monocolonized

Cell surface glycoprotein MUC18

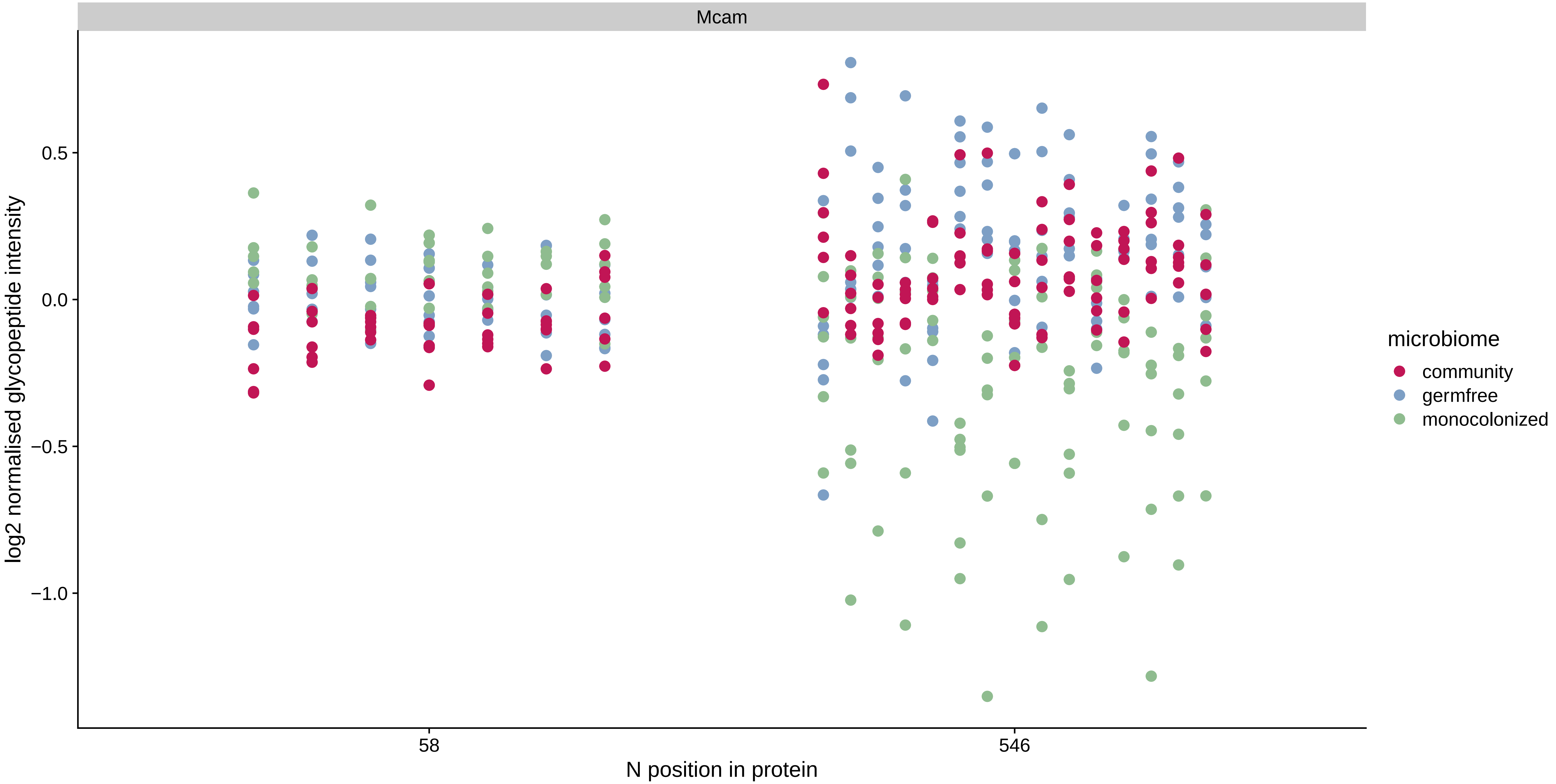

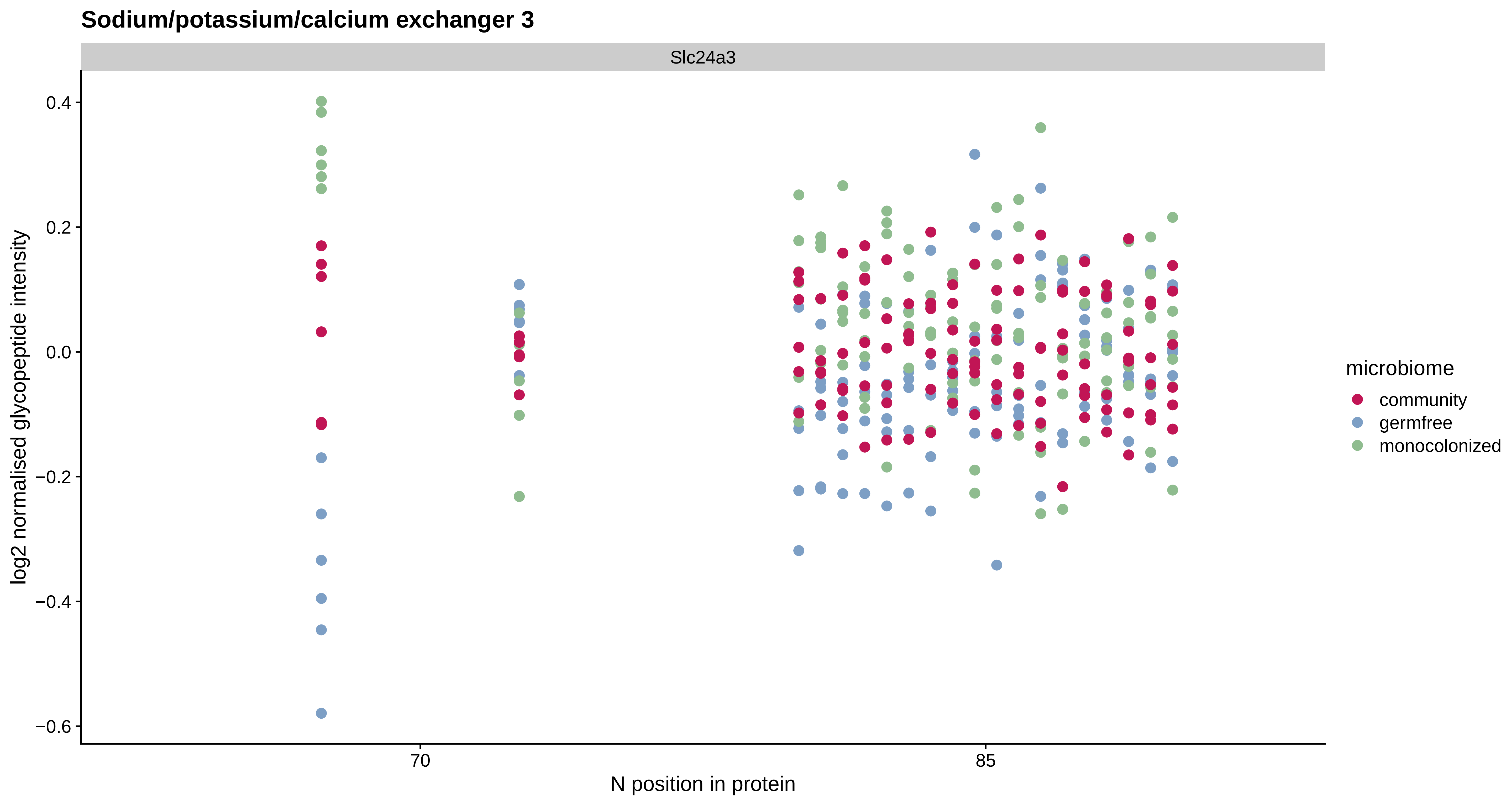
Sodium/potassium/calcium exchanger 3
Slc24a3
log2 normalised glycopeptide intensity
0.4
0.2
0.0
−0.2
−0.4
−0.6
70
85
N position in protein
microbiome
community
germfree
monocolonized

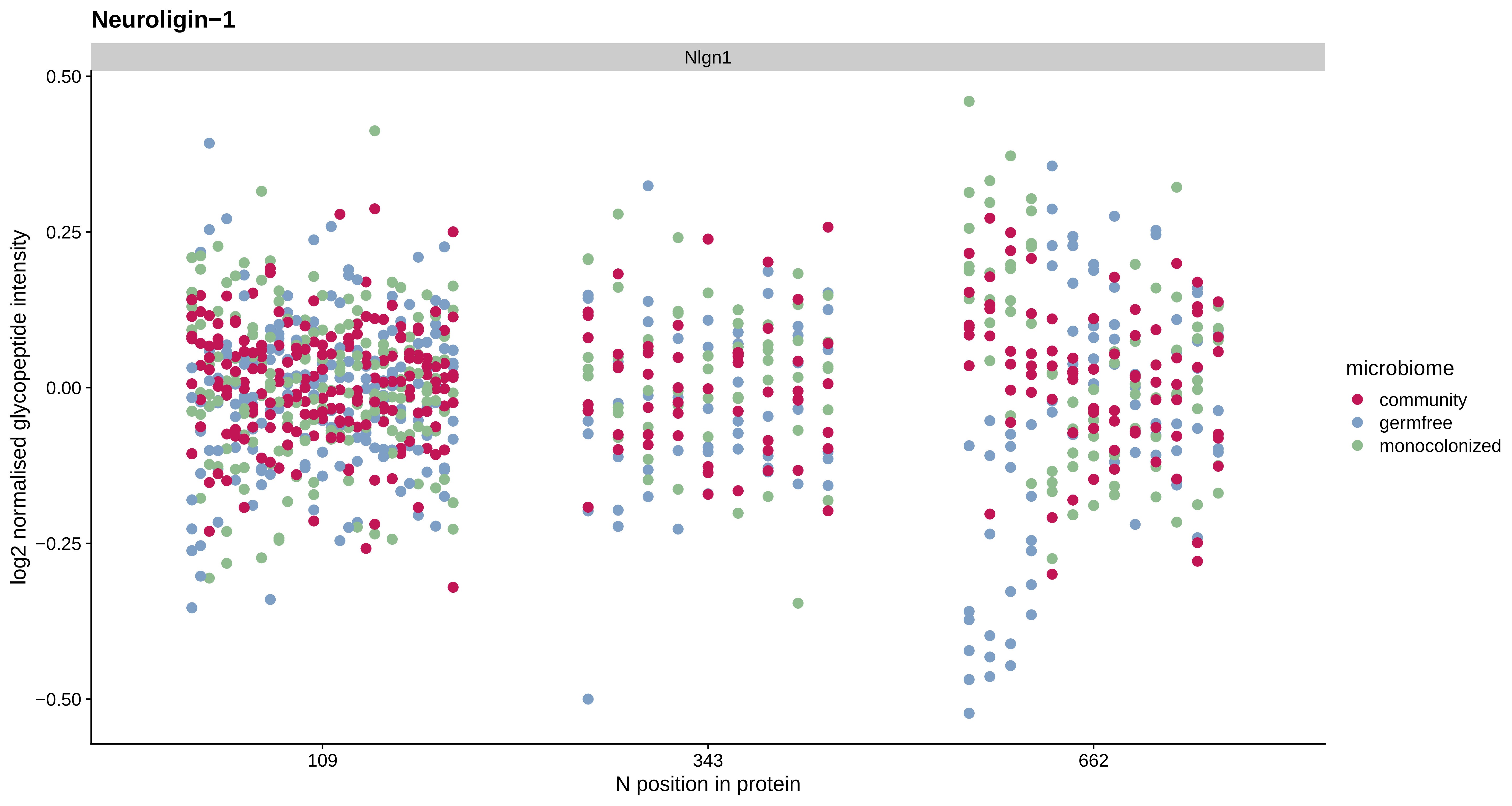
Neuroligin−1
Nlgn1
0.50
0.25
0.00
−0.25
−0.50
log2 normalised glycopeptide intensity
109
343
662
N position in protein
microbiome
community
germfree
monocolonized

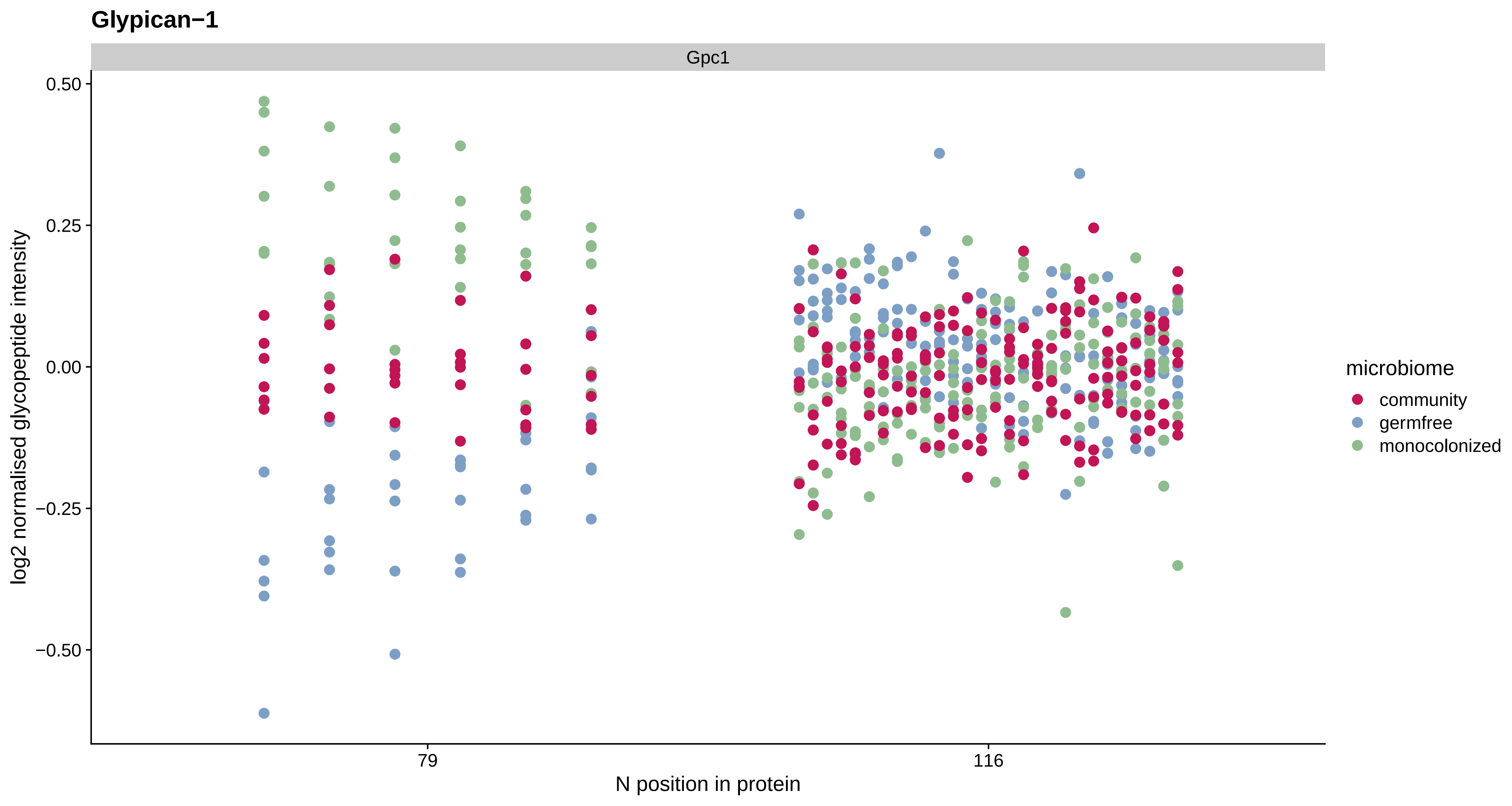
Glypican−1
Gpc1
0.50
0.25
0.00
−0.25
−0.50
log2 normalised glycopeptide intensity
79
116
N position in protein
microbiome
community
germfree
monocolonized

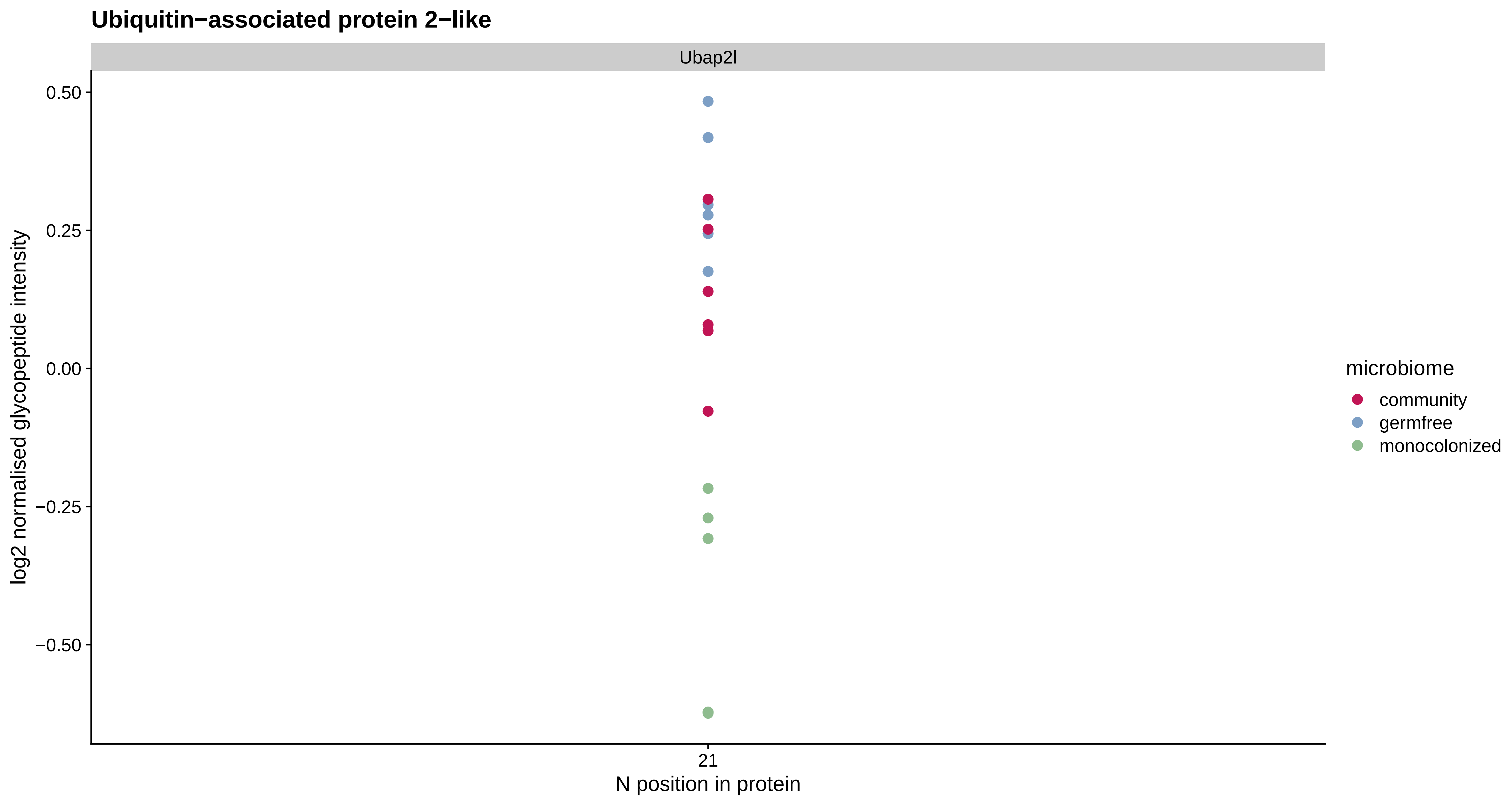
Ubiquitin−associated protein 2−like
Ubap2l
0.50
0.25
0.00
−0.25
−0.50
log2 normalised glycopeptide intensity
21
N position in protein
microbiome
community
germfree
monocolonized

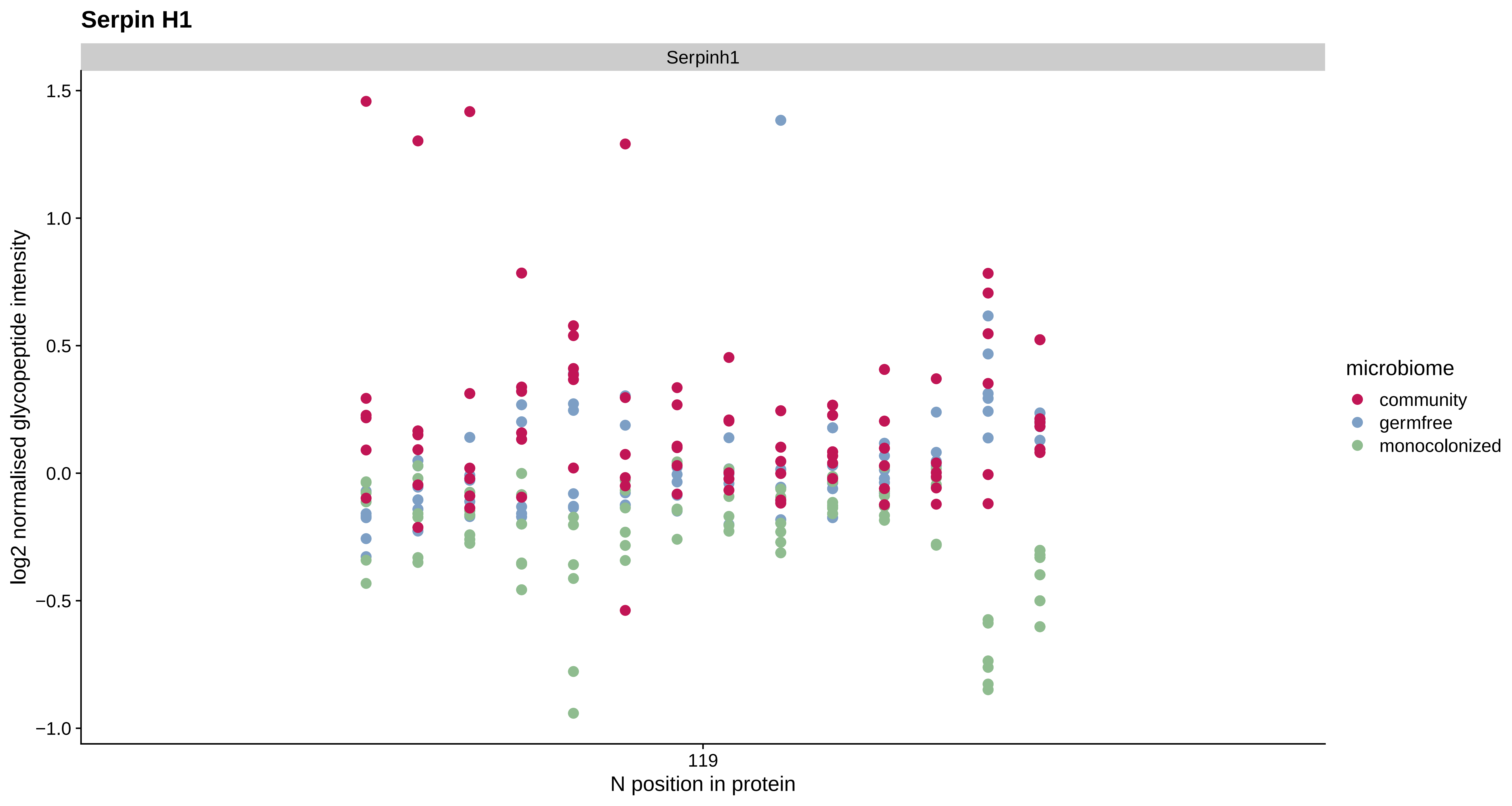
Serpin H1
Serpinh1
1.5
1.0
0.5
0.0
−0.5
−1.0
log2 normalised glycopeptide intensity
119
N position in protein
microbiome
community
germfree
monocolonized

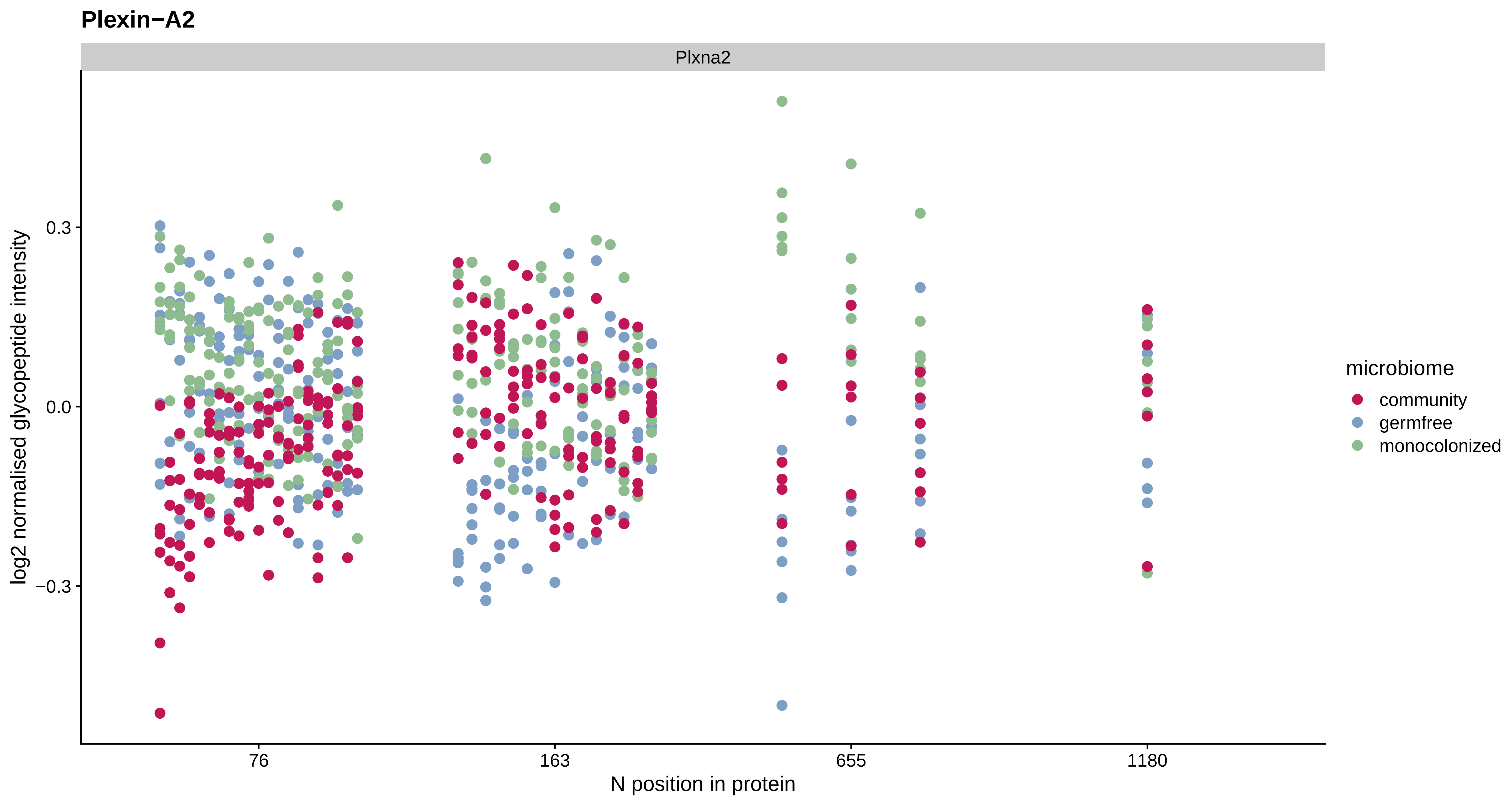

Plexin−A2
Plxna2
log2 normalised glycopeptide intensity
0.3
0.0
−0.3
76
163
655
1180
N position in protein
microbiome
community
germfree
monocolonized

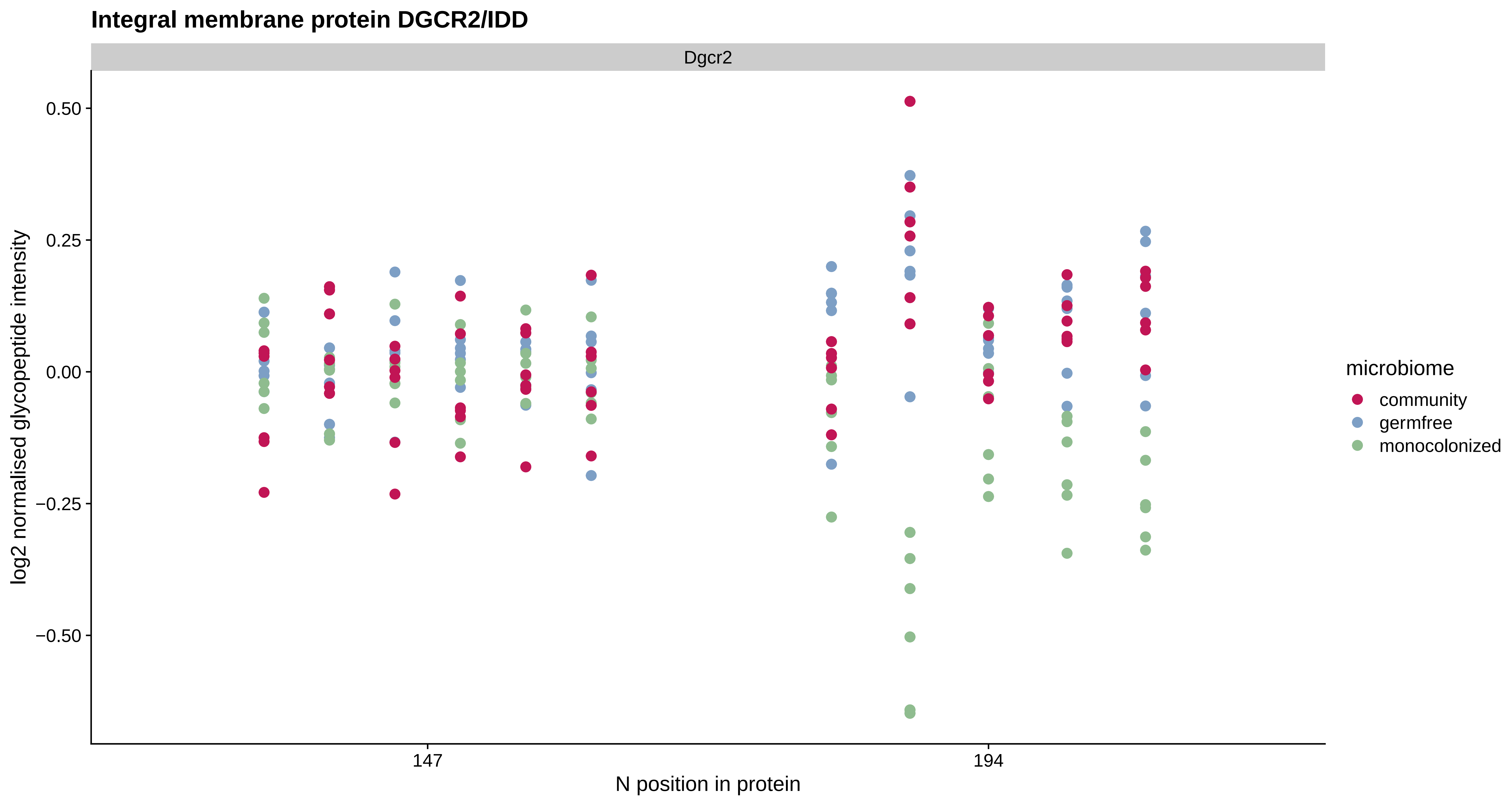

Integral membrane protein DGCR2/IDD
Dgcr2
0.50
0.25
0.00
−0.25
−0.50
log2 normalised glycopeptide intensity
147
194
N position in protein
microbiome
community
germfree
monocolonized

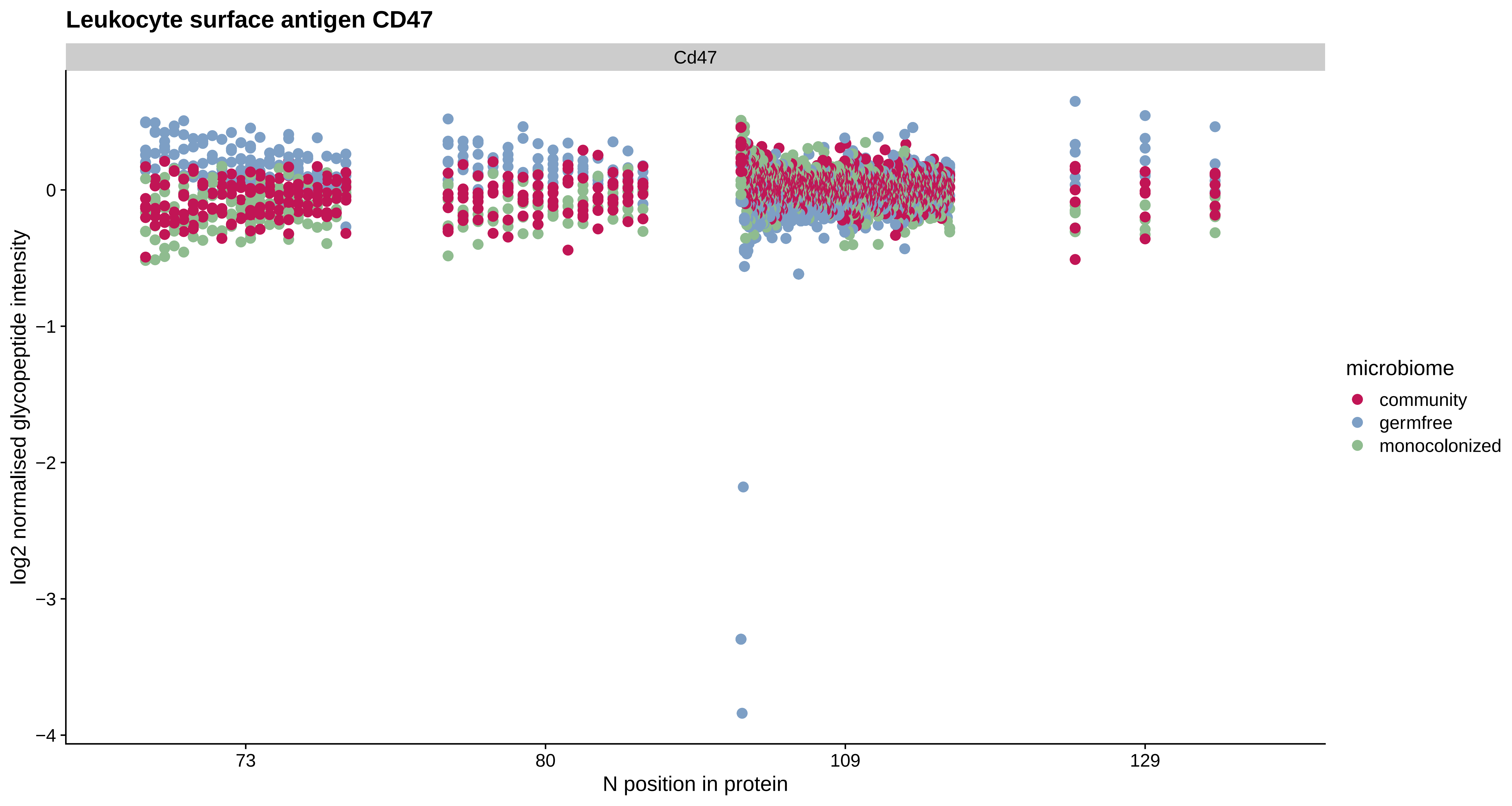

Leukocyte surface antigen CD47
Cd47
log2 normalised glycopeptide intensity
0
−1
−2
−3
−4
73
80
109
129
N position in protein
microbiome
community
germfree
monocolonized

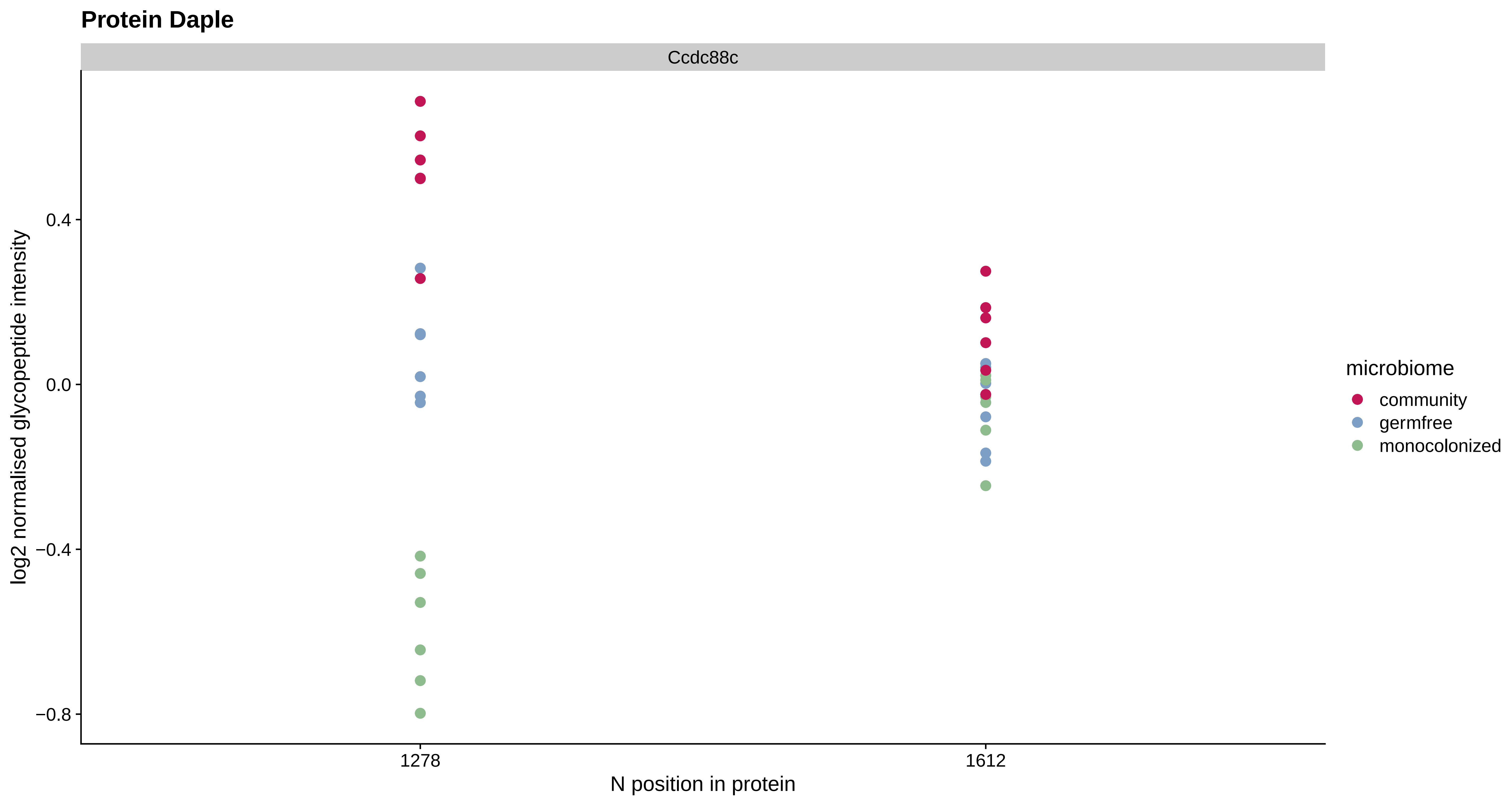
Protein Daple
Ccdc88c
log2 normalised glycopeptide intensity
0.4
0.0
−0.4
−0.8
1278
1612
N position in protein
microbiome
community
germfree
monocolonized

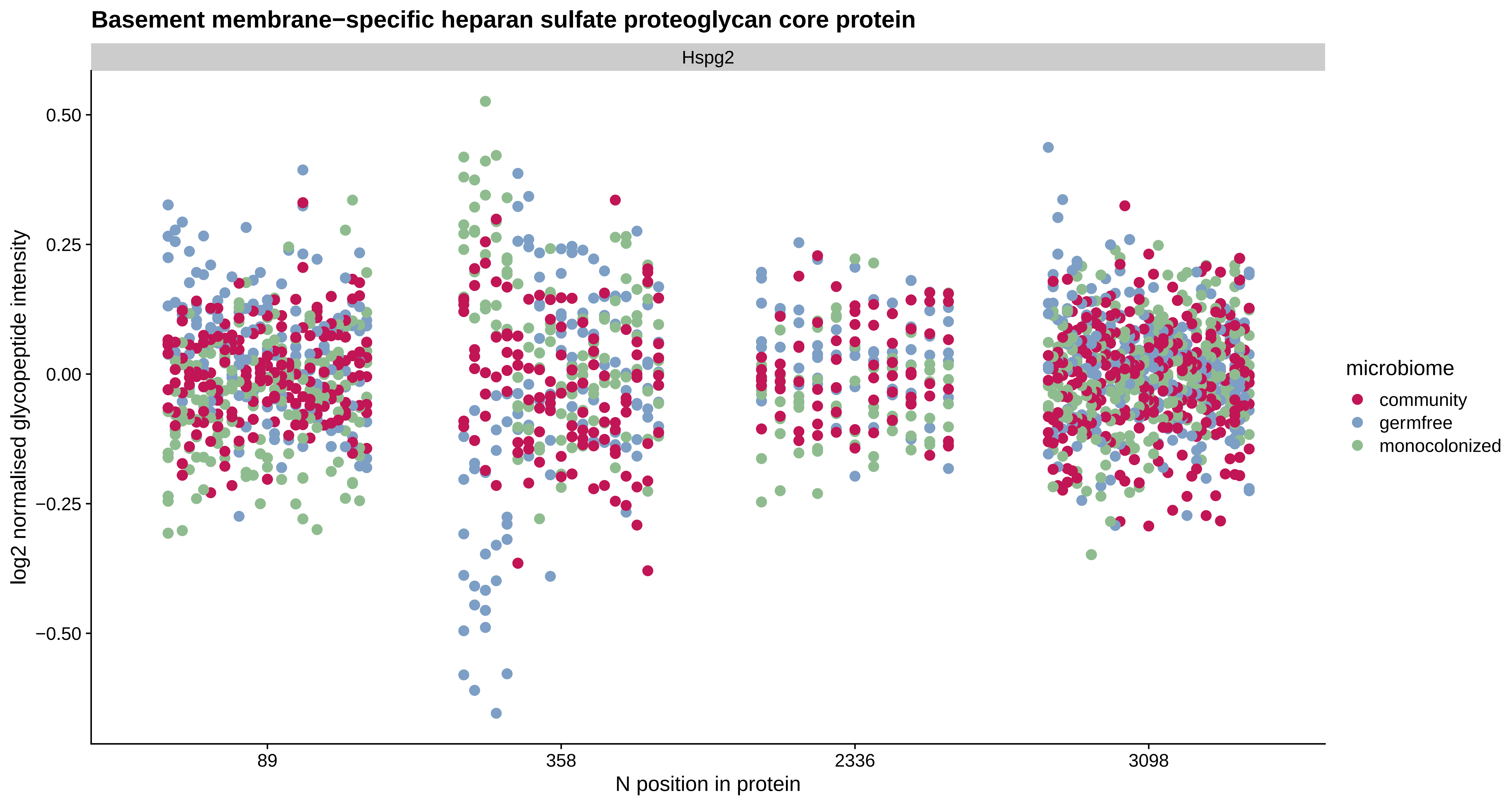

Basement membrane−specific heparan sulfate proteoglycan core protein
Hspg2
0.50
0.25
0.00
−0.25
−0.50
log2 normalised glycopeptide intensity
89
358
2336
3098
N position in protein
microbiome
community
germfree
monocolonized

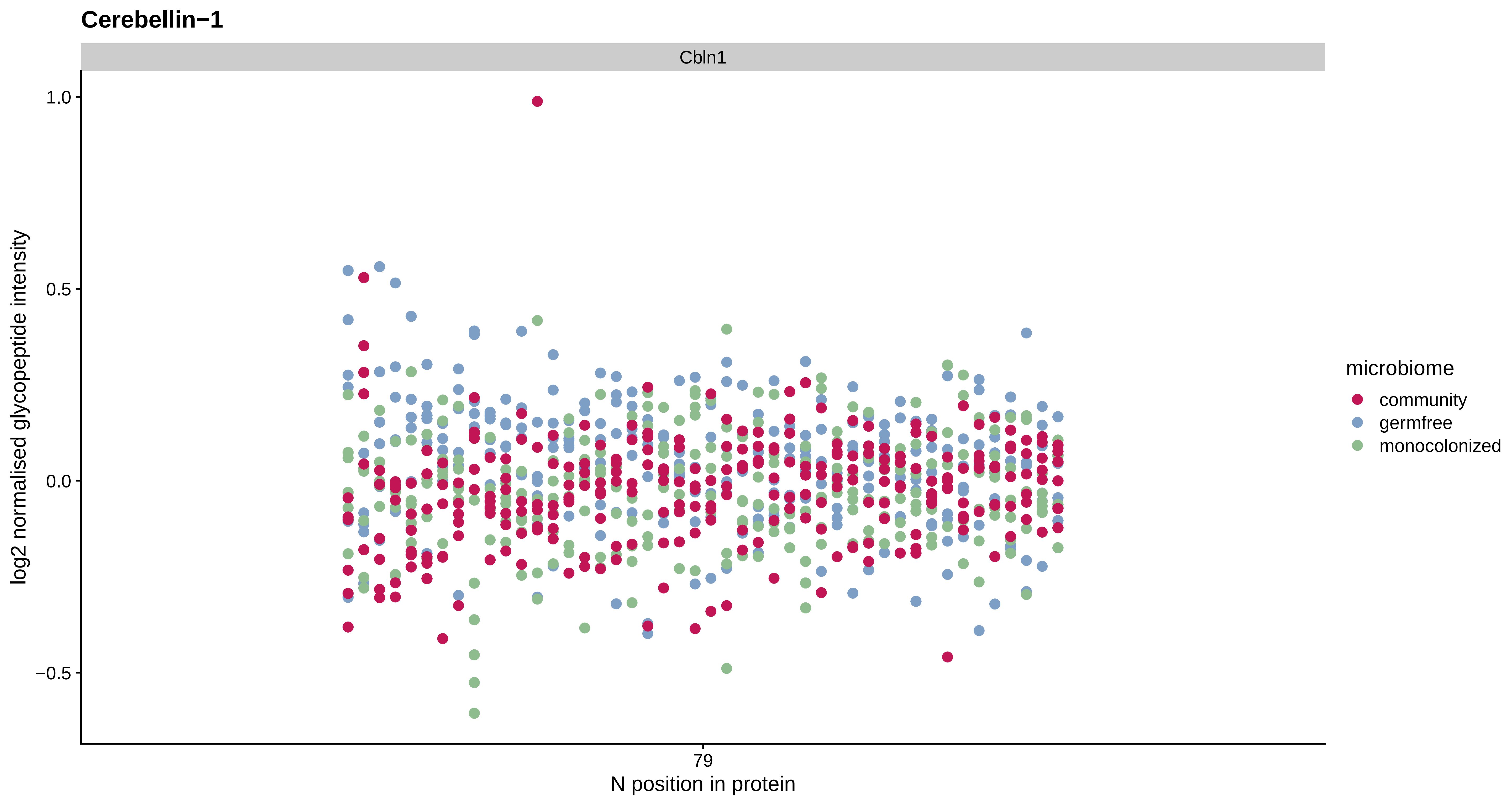

Cerebellin−1
Cbln1
1.0
0.5
0.0
−0.5
log2 normalised glycopeptide intensity
79
N position in protein
microbiome
community
germfree
monocolonized

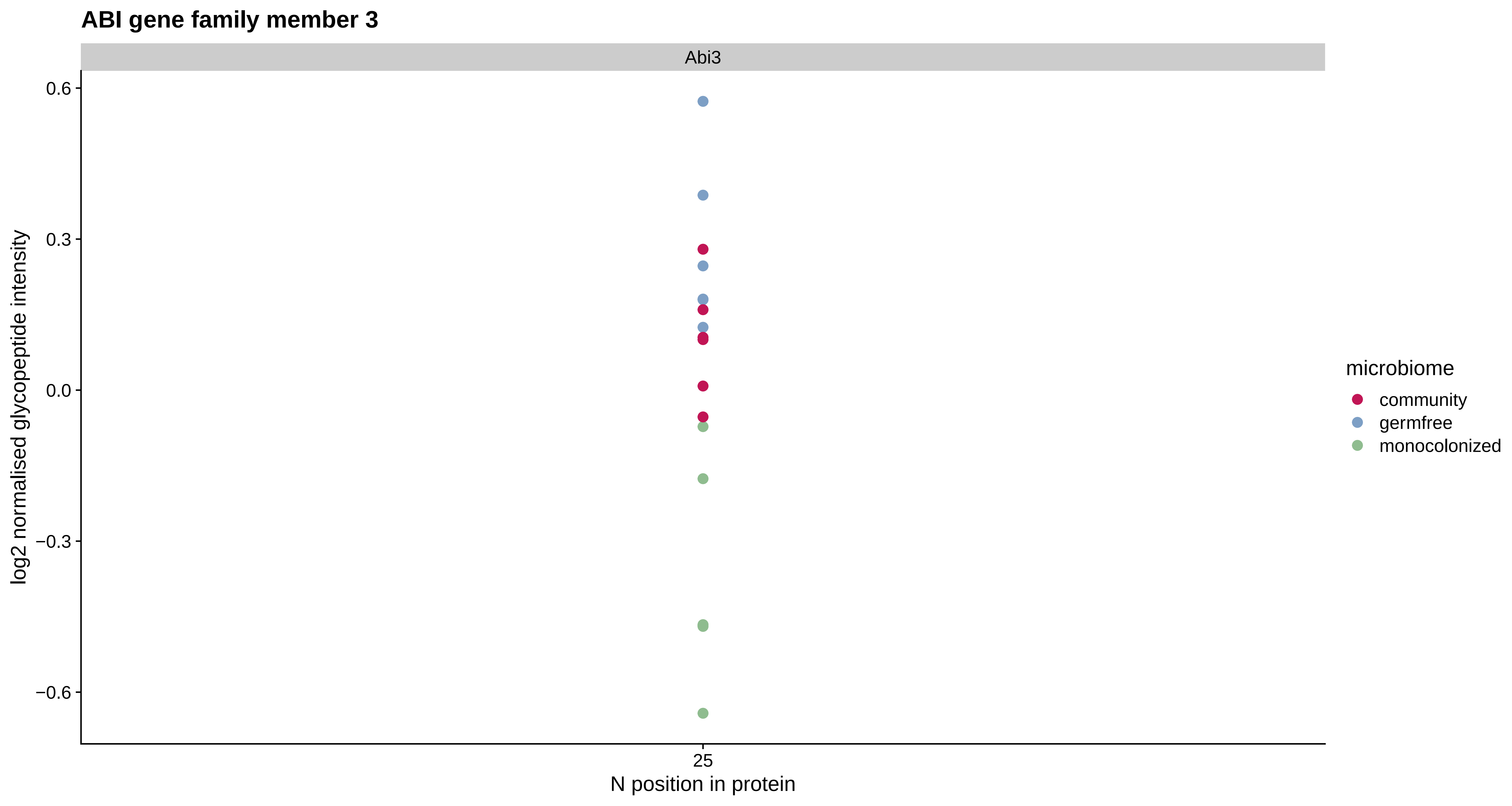
ABI gene family member 3
Abi3
0.6
0.3
0.0
−0.3
−0.6
log2 normalised glycopeptide intensity
25
N position in protein
microbiome
community
germfree
monocolonized

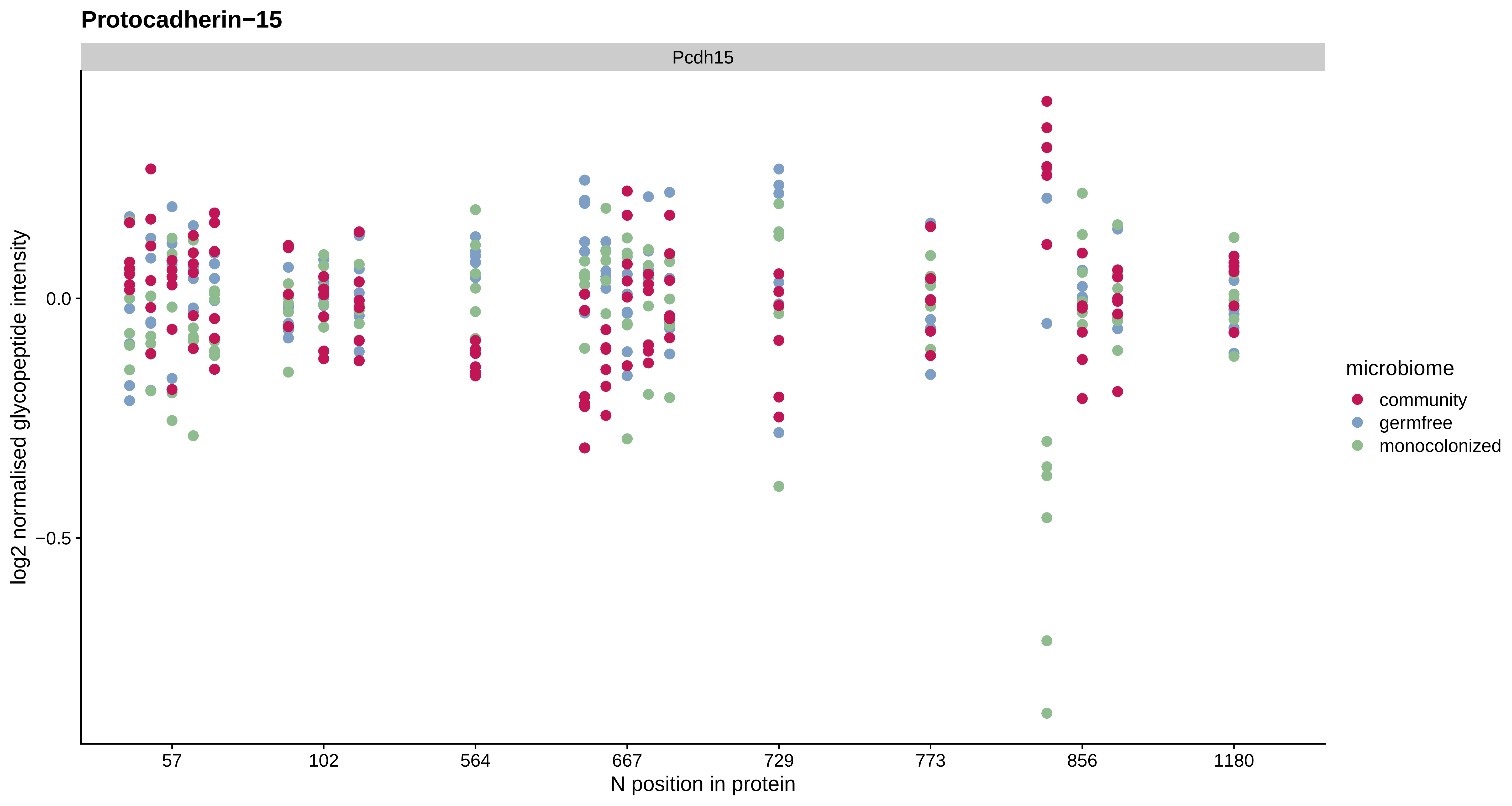
Protocadherin−15
Pcdh15
log2 normalised glycopeptide intensity
0.0
−0.5
57
102
564
667
729
773
856
1180
N position in protein
microbiome
community
germfree
monocolonized

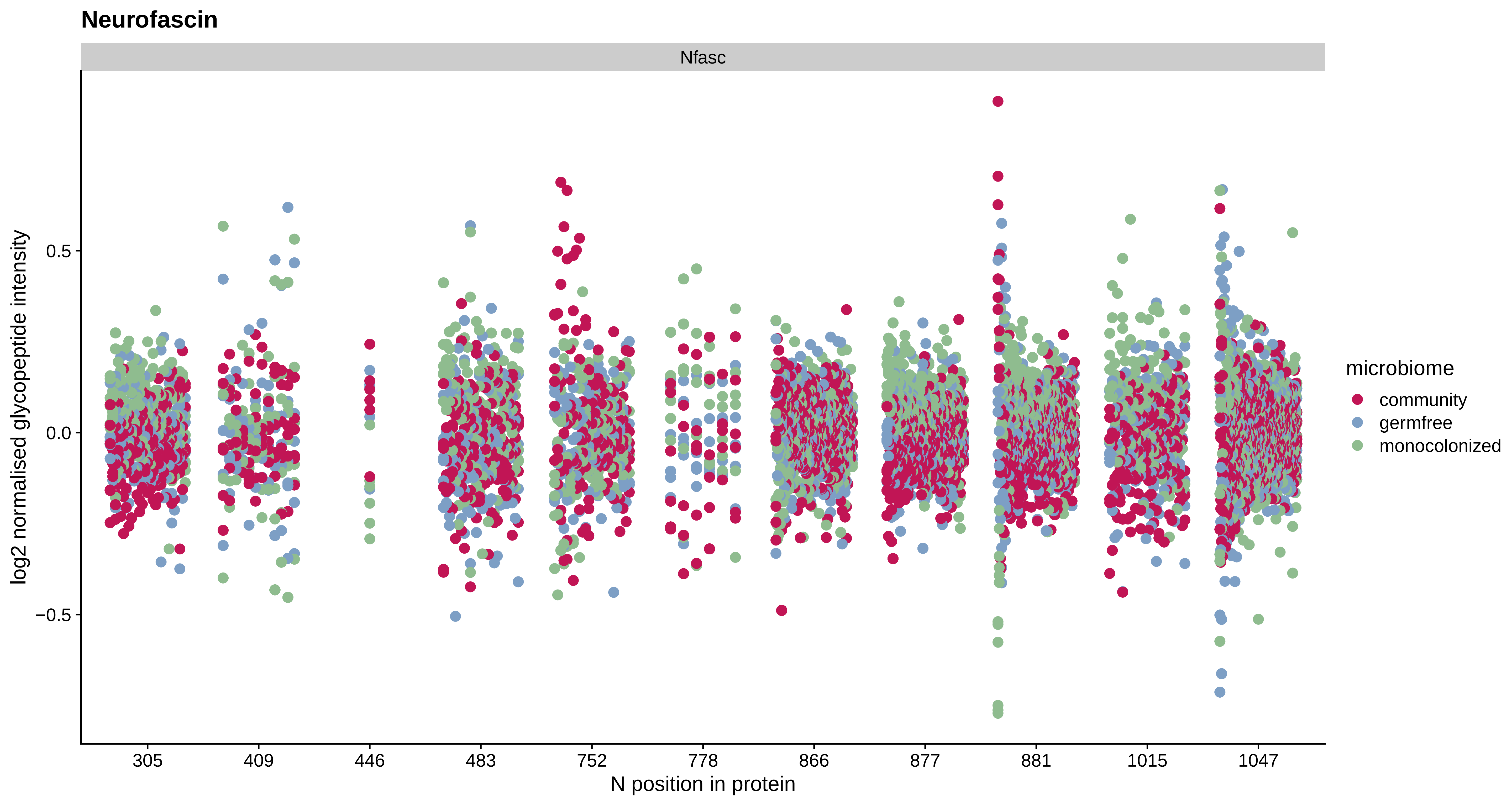
Neurofascin
Nfasc
log2 normalised glycopeptide intensity
0.5
0.0
−0.5
305
409
446
483
752
778
866
877
881
1015
1047
N position in protein
microbiome
community
germfree
monocolonized

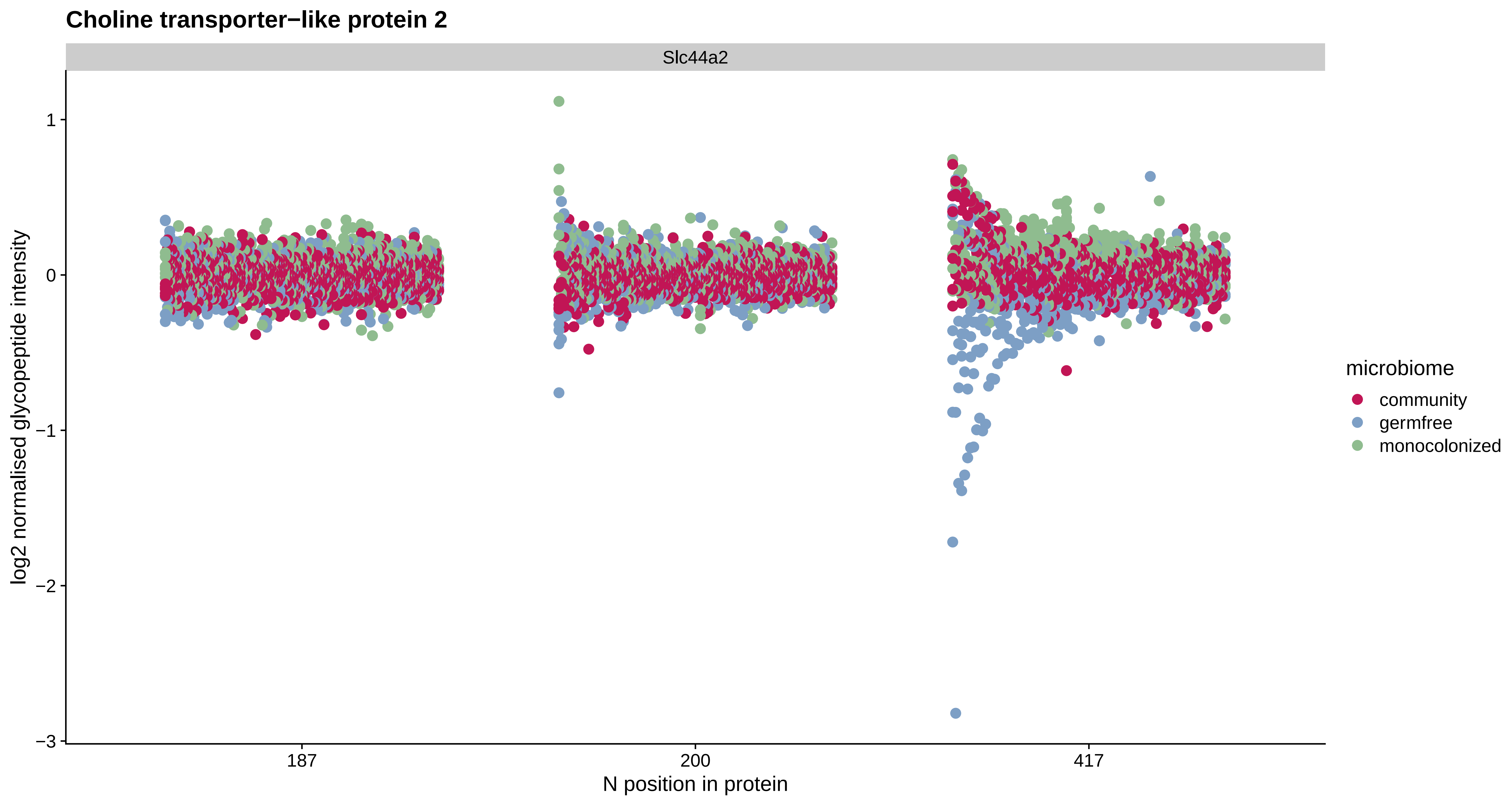
Choline transporter−like protein 2
Slc44a2
log2 normalised glycopeptide intensity
1
0
−1
−2
−3
187
200
417
N position in protein
microbiome
community
germfree
monocolonized

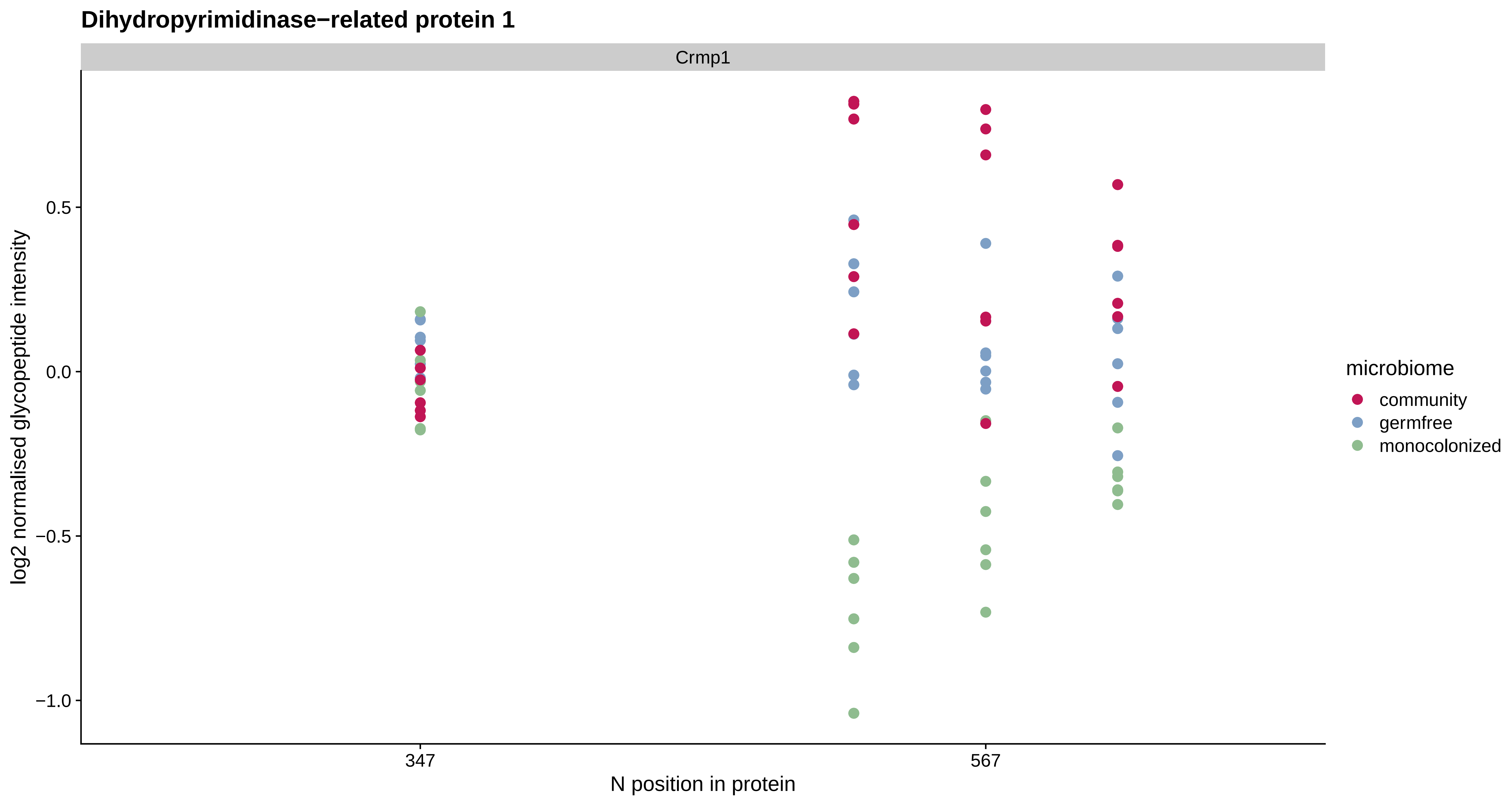
Dihydropyrimidinase−related protein 1
Crmp1
log2 normalised glycopeptide intensity
0.5
0.0
−0.5
−1.0
347
567
N position in protein
microbiome
community
germfree
monocolonized

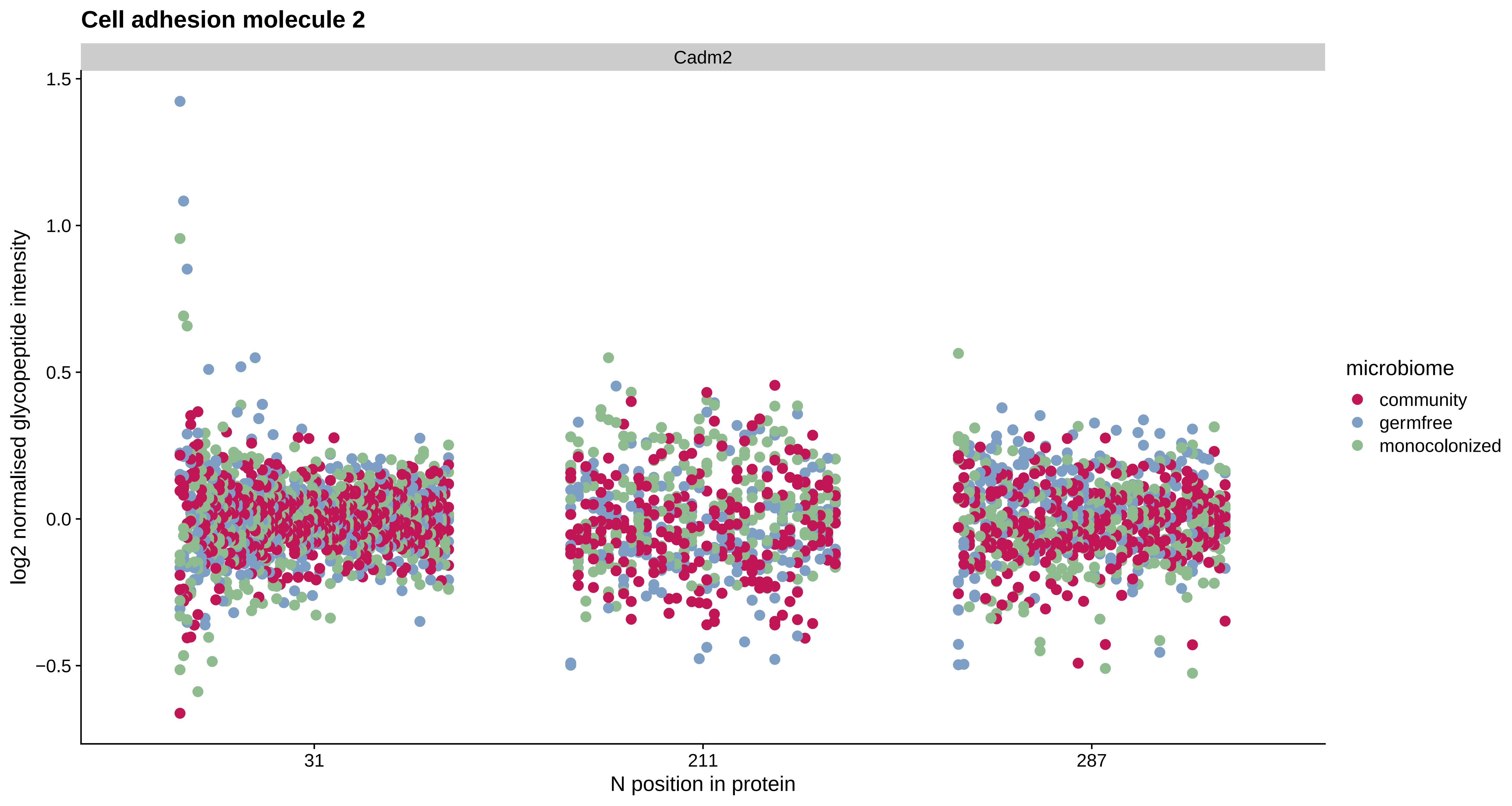

Cell adhesion molecule 2
Cadm2
1.5
1.0
0.5
0.0
−0.5
log2 normalised glycopeptide intensity
31
211
287
N position in protein
microbiome
community
germfree
monocolonized

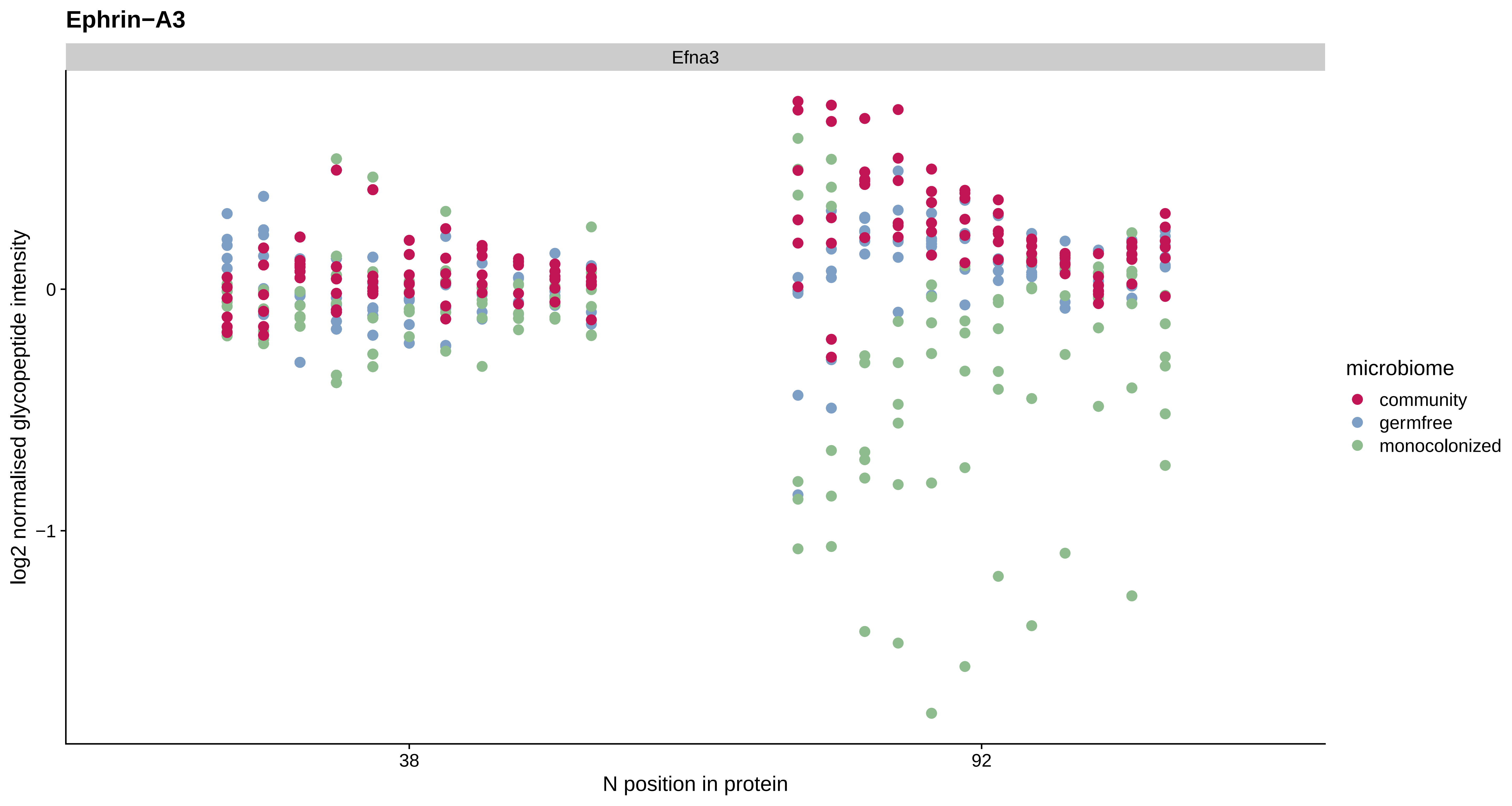
Ephrin−A3
Efna3
log2 normalised glycopeptide intensity
0
−1
38
92
N position in protein
microbiome
community
germfree
monocolonized

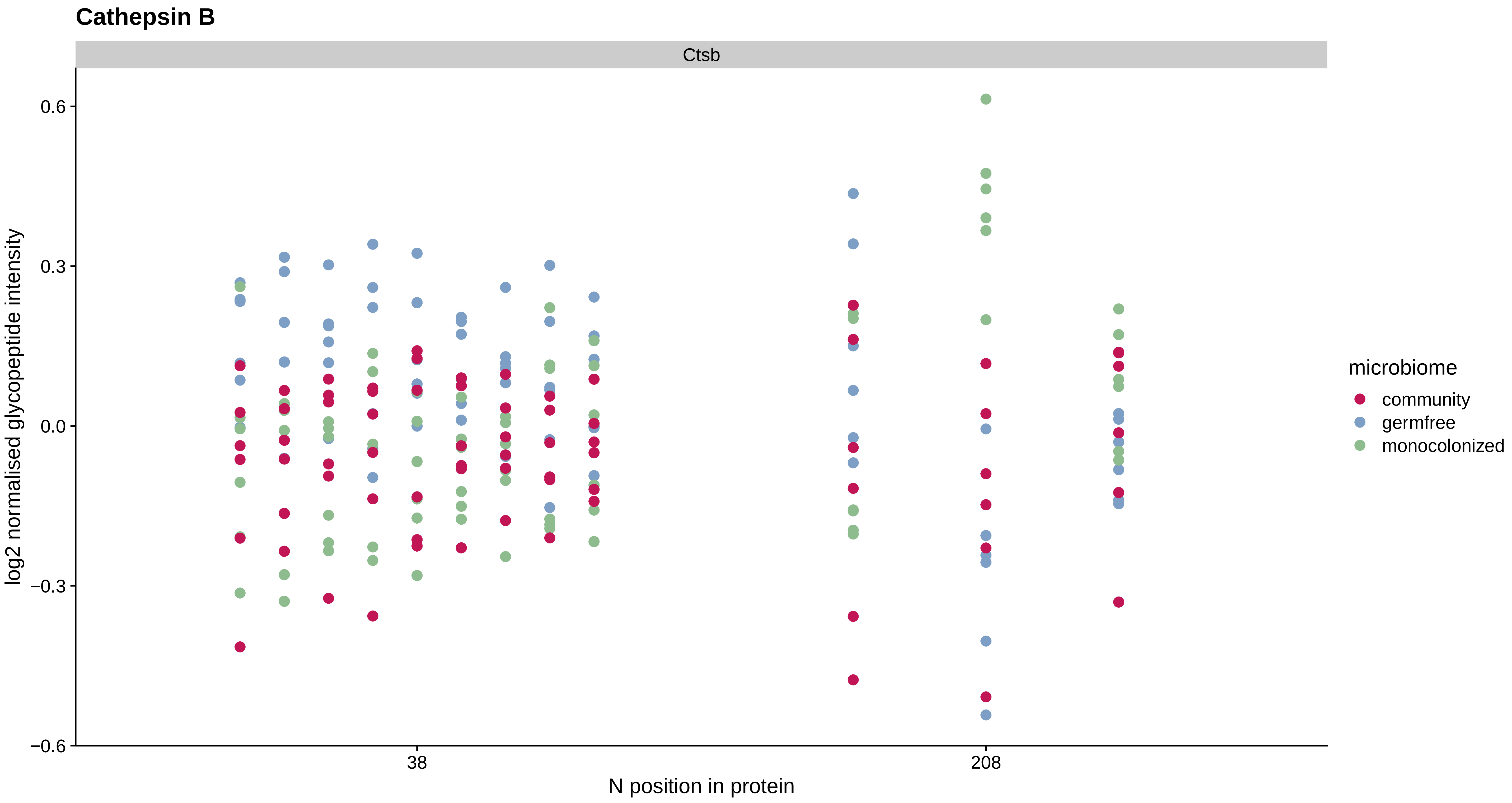

Cathepsin B
Ctsb
0.6
0.3
0.0
−0.3
−0.6
log2 normalised glycopeptide intensity
38
208
N position in protein
microbiome
community
germfree
monocolonized

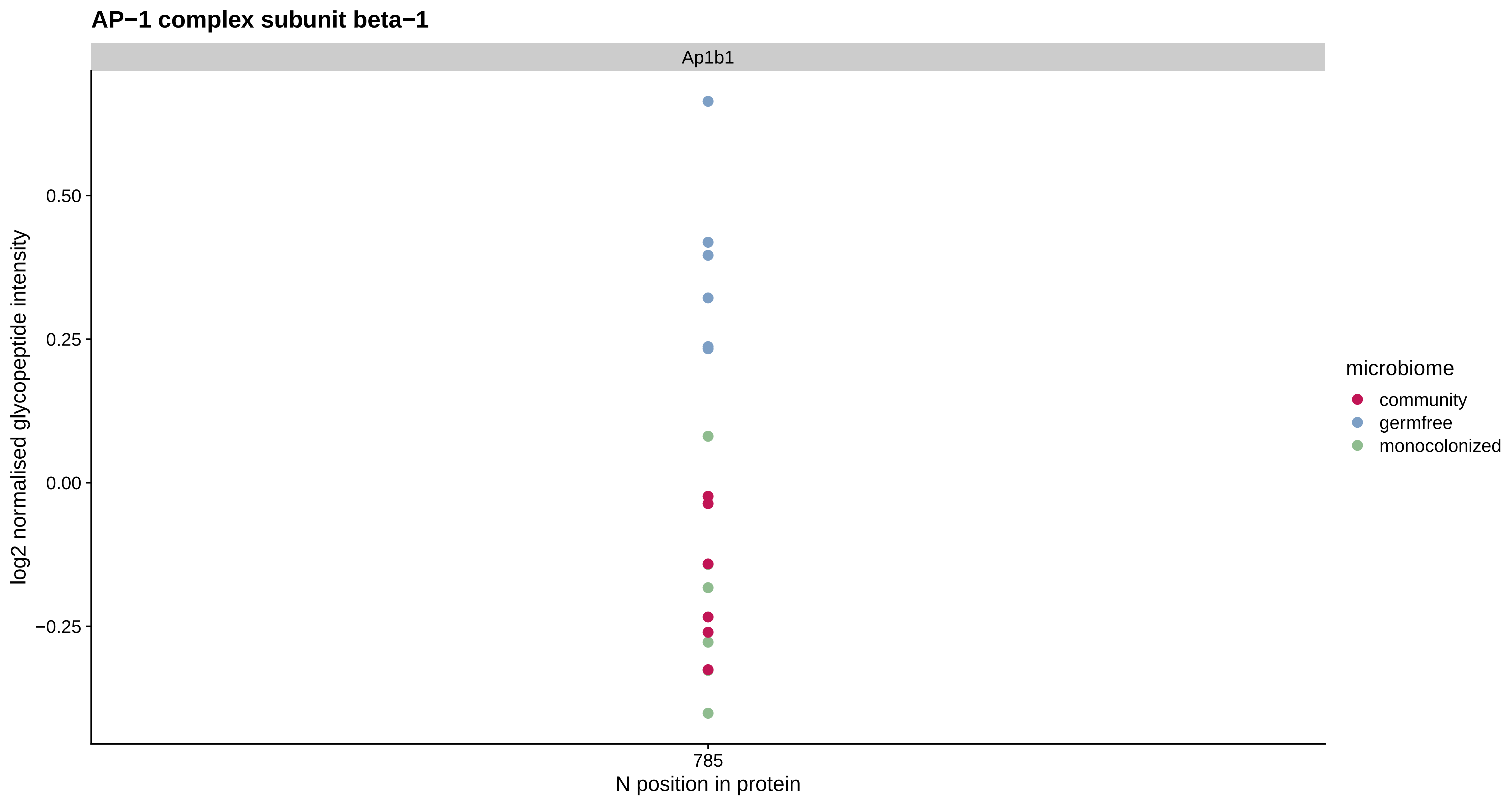
AP−1 complex subunit beta−1
Ap1b1
log2 normalised glycopeptide intensity
0.50
0.25
0.00
−0.25
785
N position in protein
microbiome
community
germfree
monocolonized

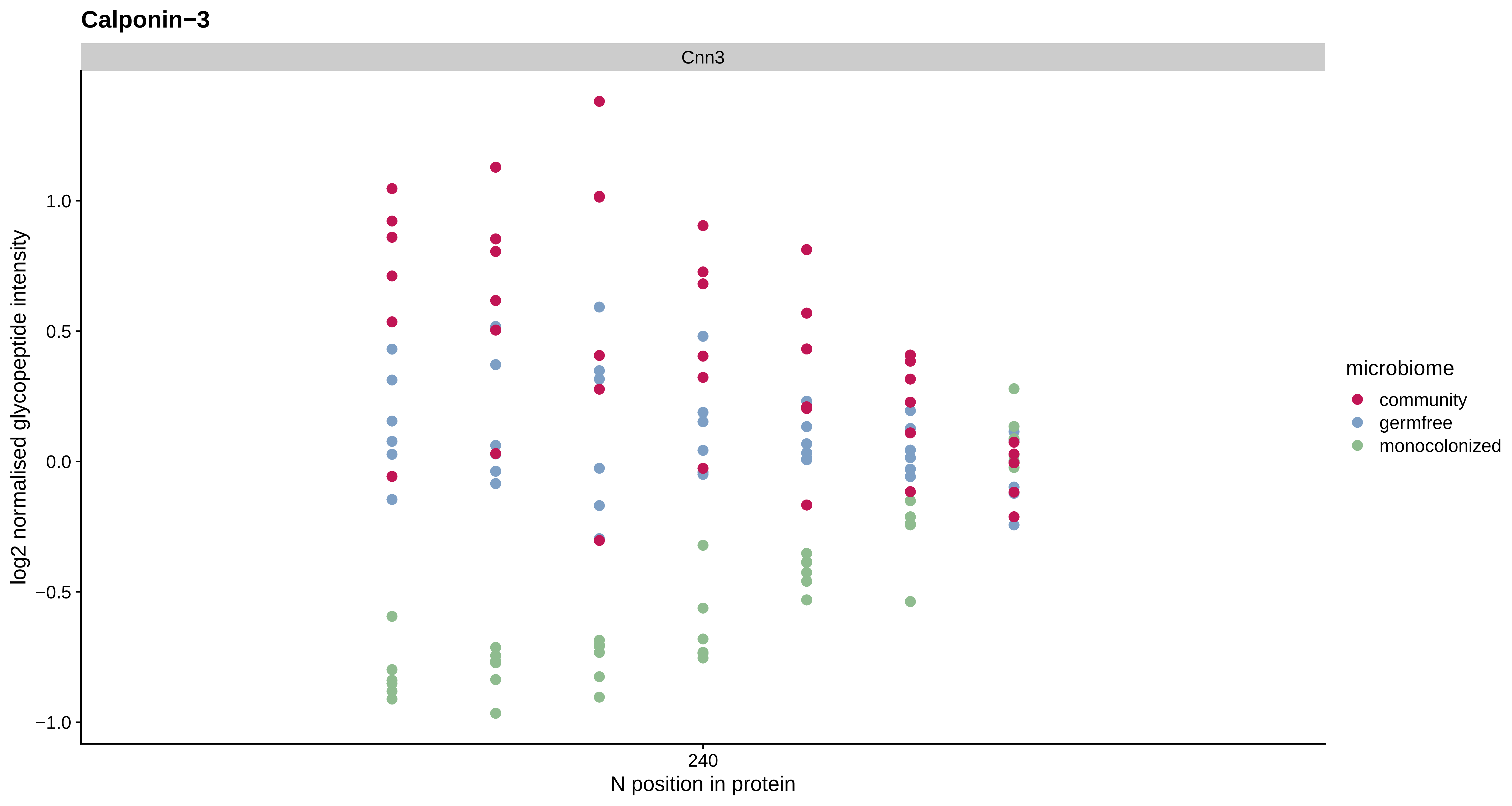

Calponin−3
Cnn3
1.0
0.5
0.0
−0.5
−1.0
log2 normalised glycopeptide intensity
240
N position in protein
microbiome
community
germfree
monocolonized

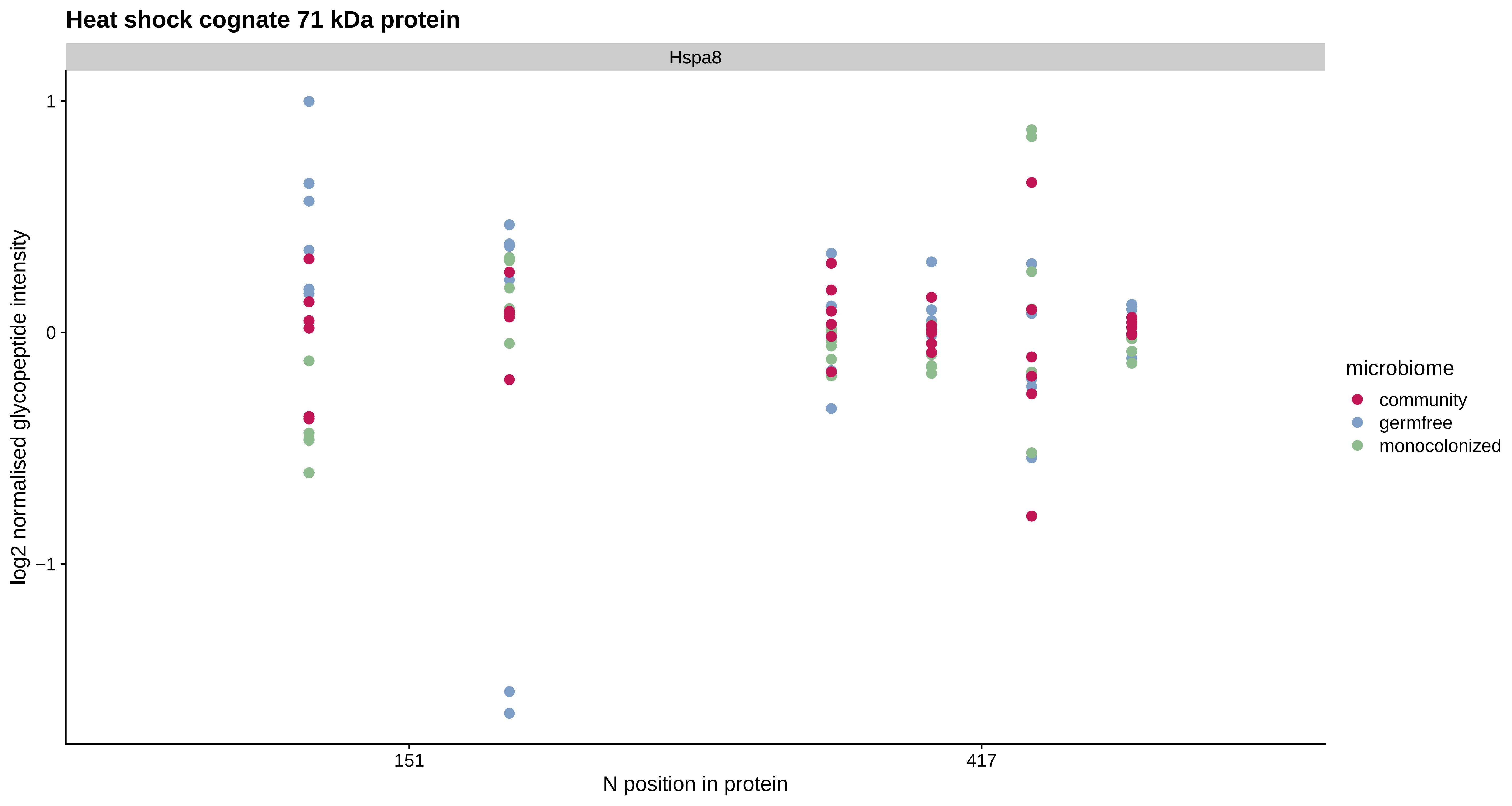

Heat shock cognate 71 kDa protein
Hspa8
1
0
−1
log2 normalised glycopeptide intensity
151
417
N position in protein
microbiome
community
germfree
monocolonized

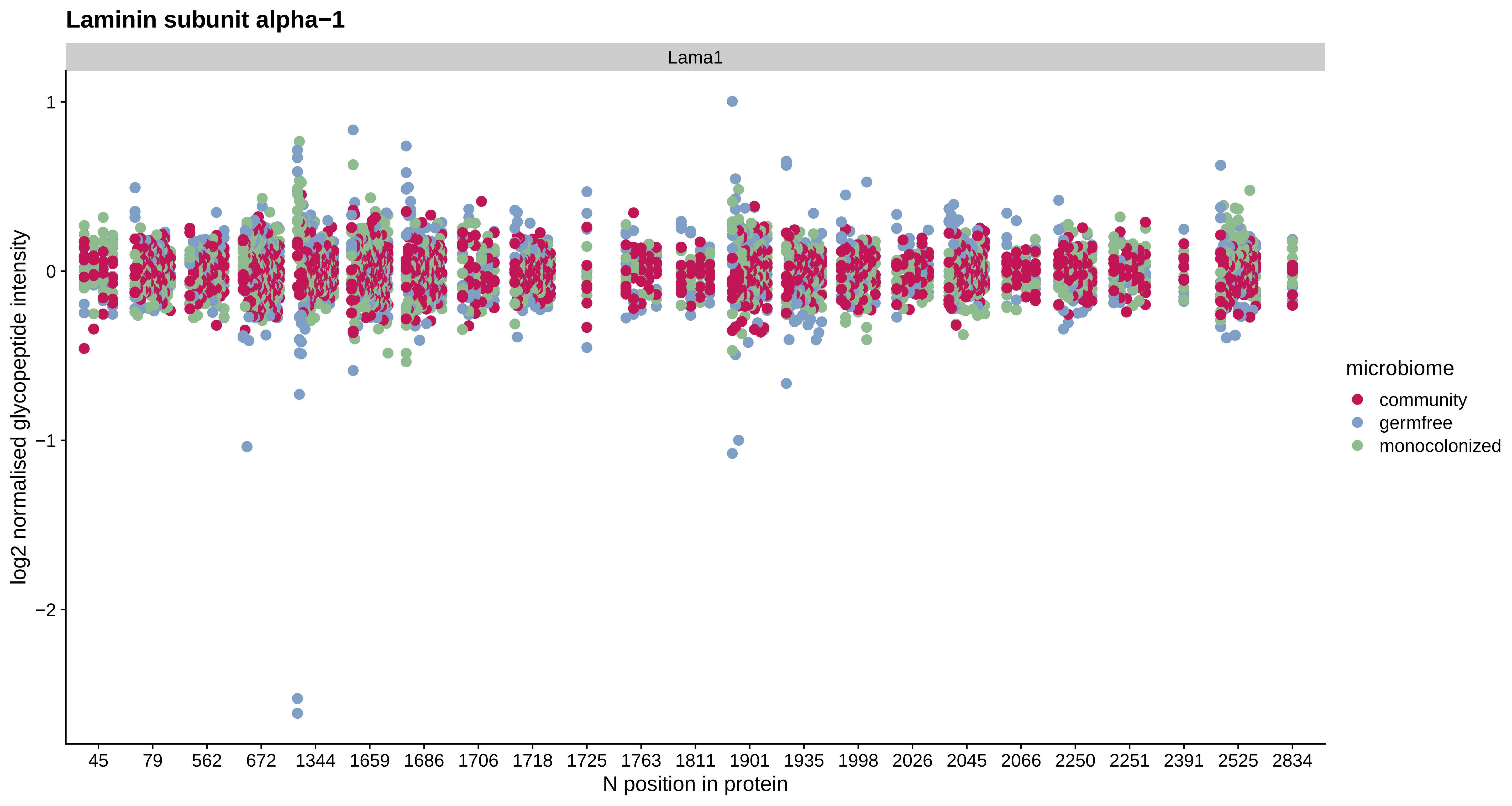
Laminin subunit alpha−1
Lama1
log2 normalised glycopeptide intensity
1
0
−1
−2
45
79
562
672
1344
1659
1686
1706
1718
1725
1763
1811
1901
1935
1998
2026
2045
2066
2250
2251
2391
2525
2834
N position in protein
microbiome
community
germfree
monocolonized

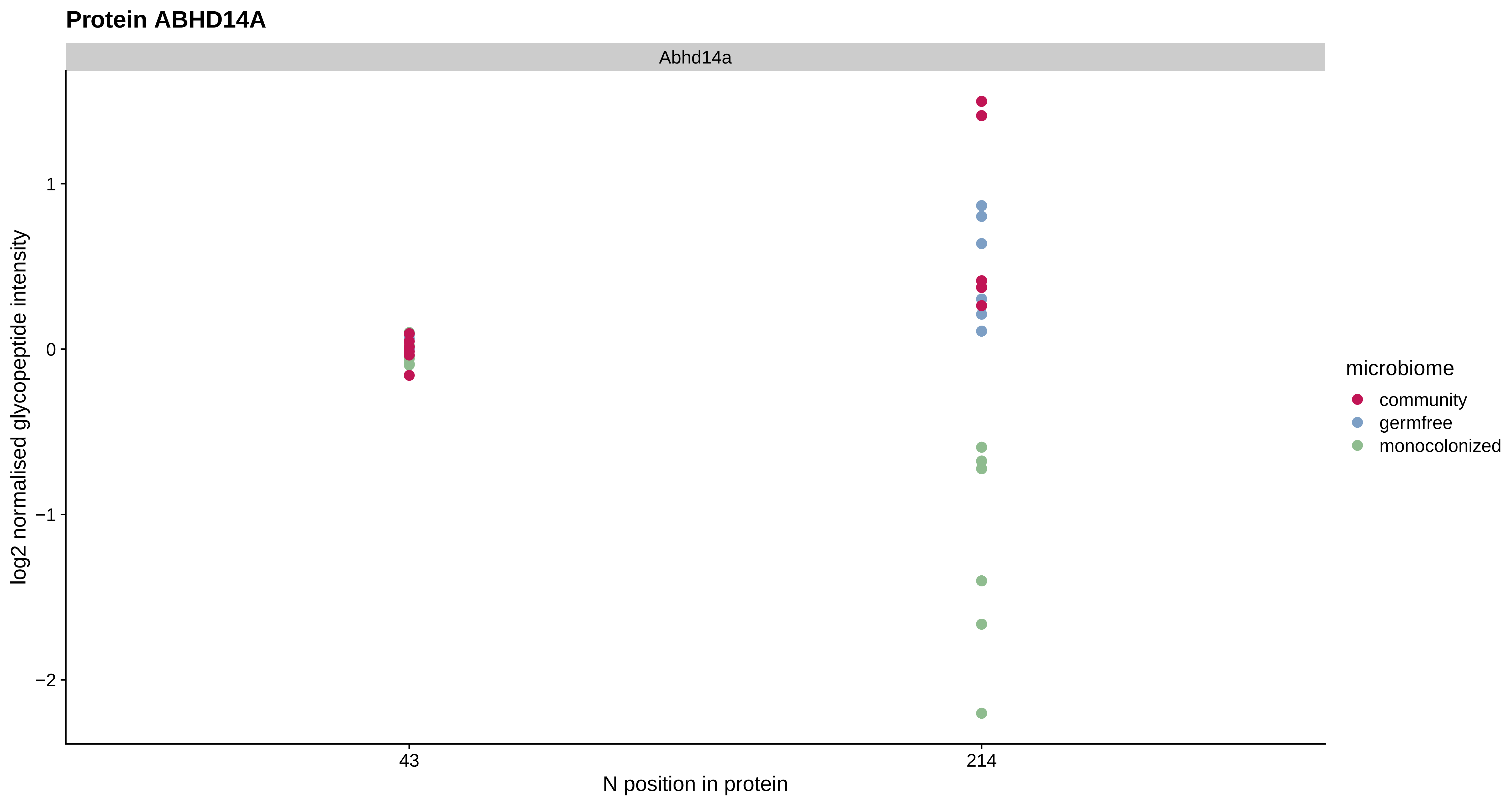
Protein ABHD14A
Abhd14a
log2 normalised glycopeptide intensity
1
0
−1
−2
43
214
N position in protein
microbiome
community
germfree
monocolonized

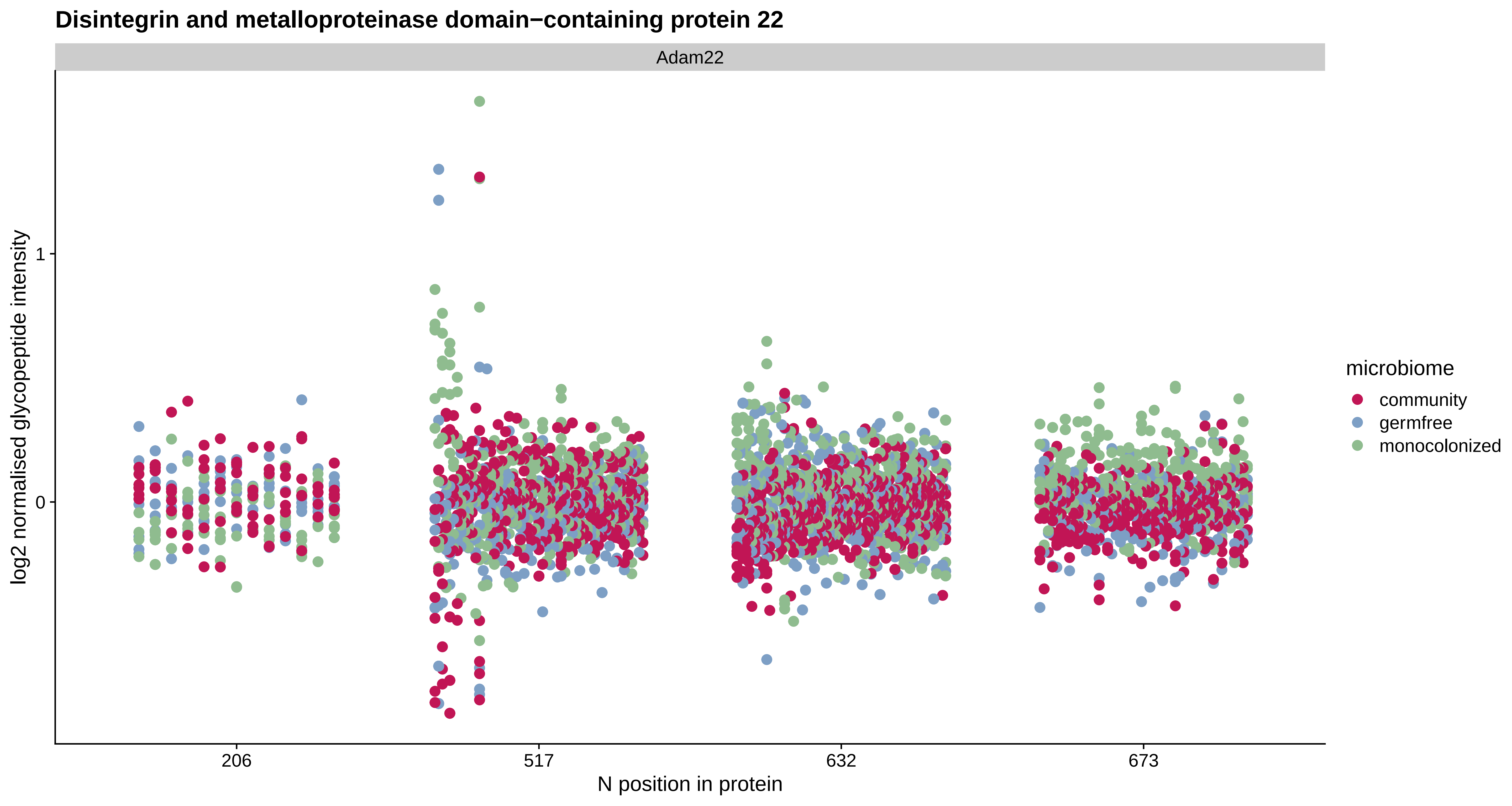
Disintegrin and metalloproteinase domain−containing protein 22
Adam22
log2 normalised glycopeptide intensity
1
0
206
517
632
673
N position in protein
microbiome
community
germfree
monocolonized

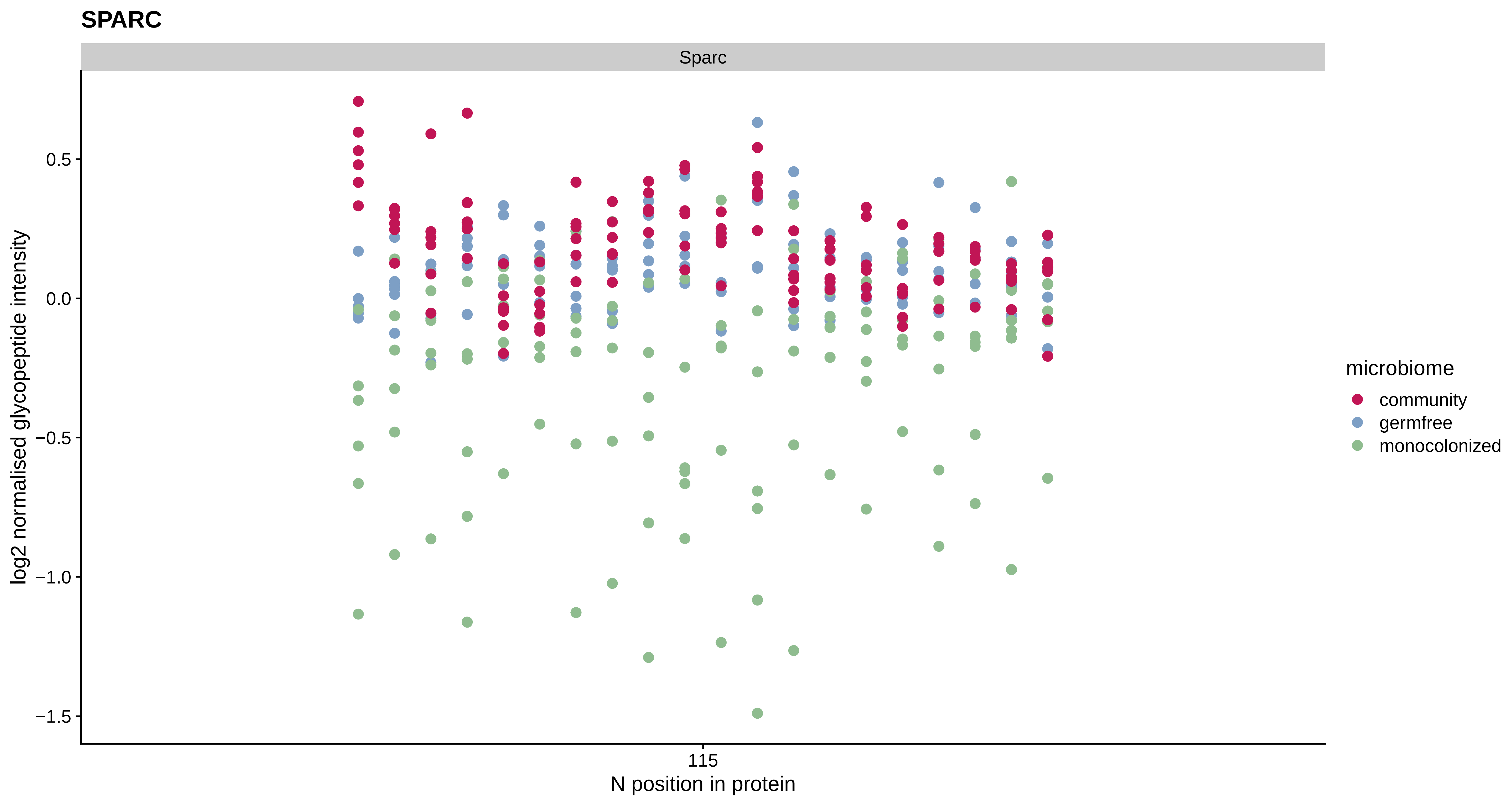

SPARC
Sparc
0.5
0.0
−0.5
−1.0
−1.5
log2 normalised glycopeptide intensity
115
N position in protein
microbiome
community
germfree
monocolonized

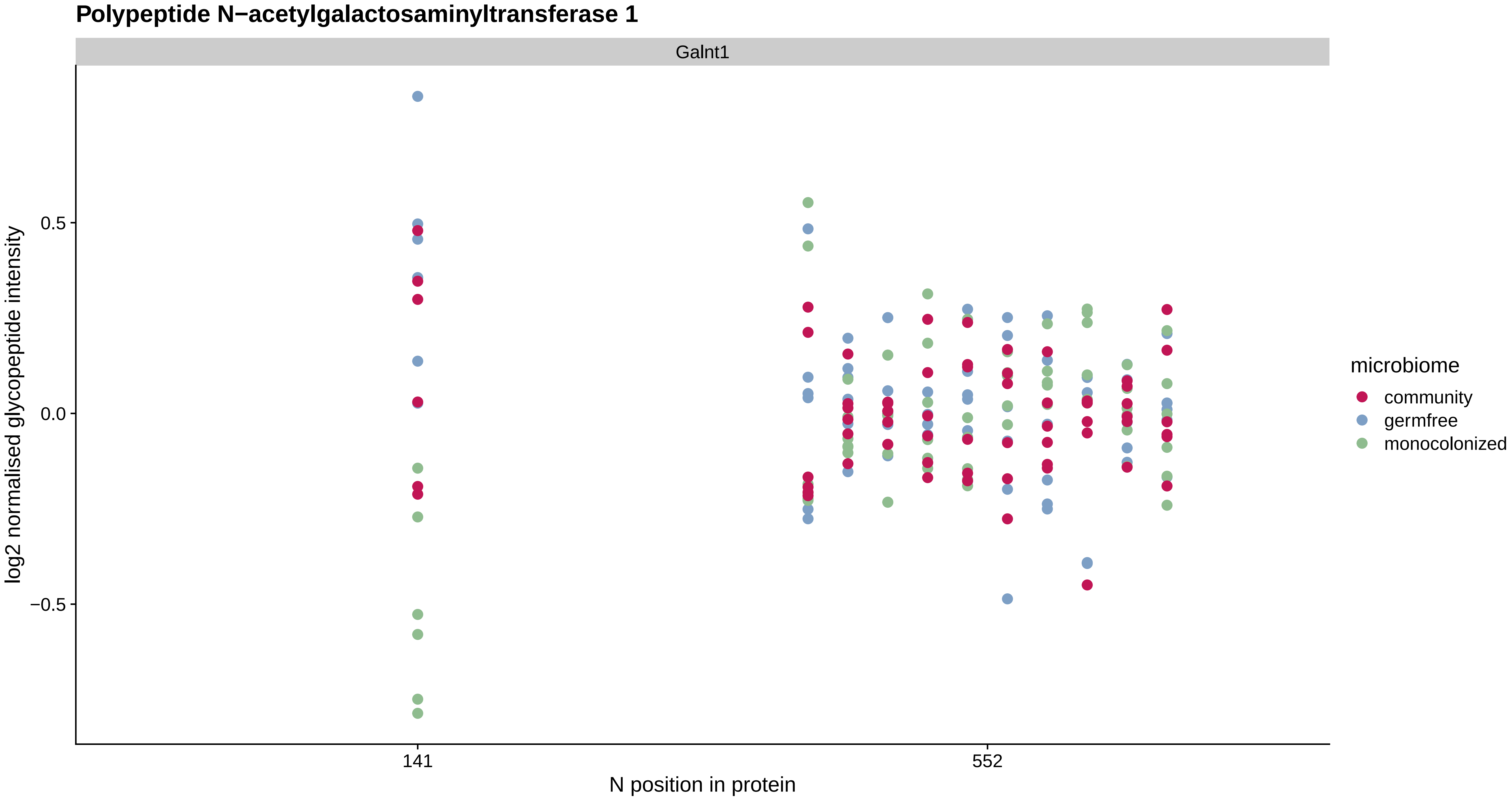
Polypeptide N−acetylgalactosaminyltransferase 1
Galnt1
0.5
0.0
−0.5
log2 normalised glycopeptide intensity
141
552
N position in protein
microbiome
community
germfree
monocolonized

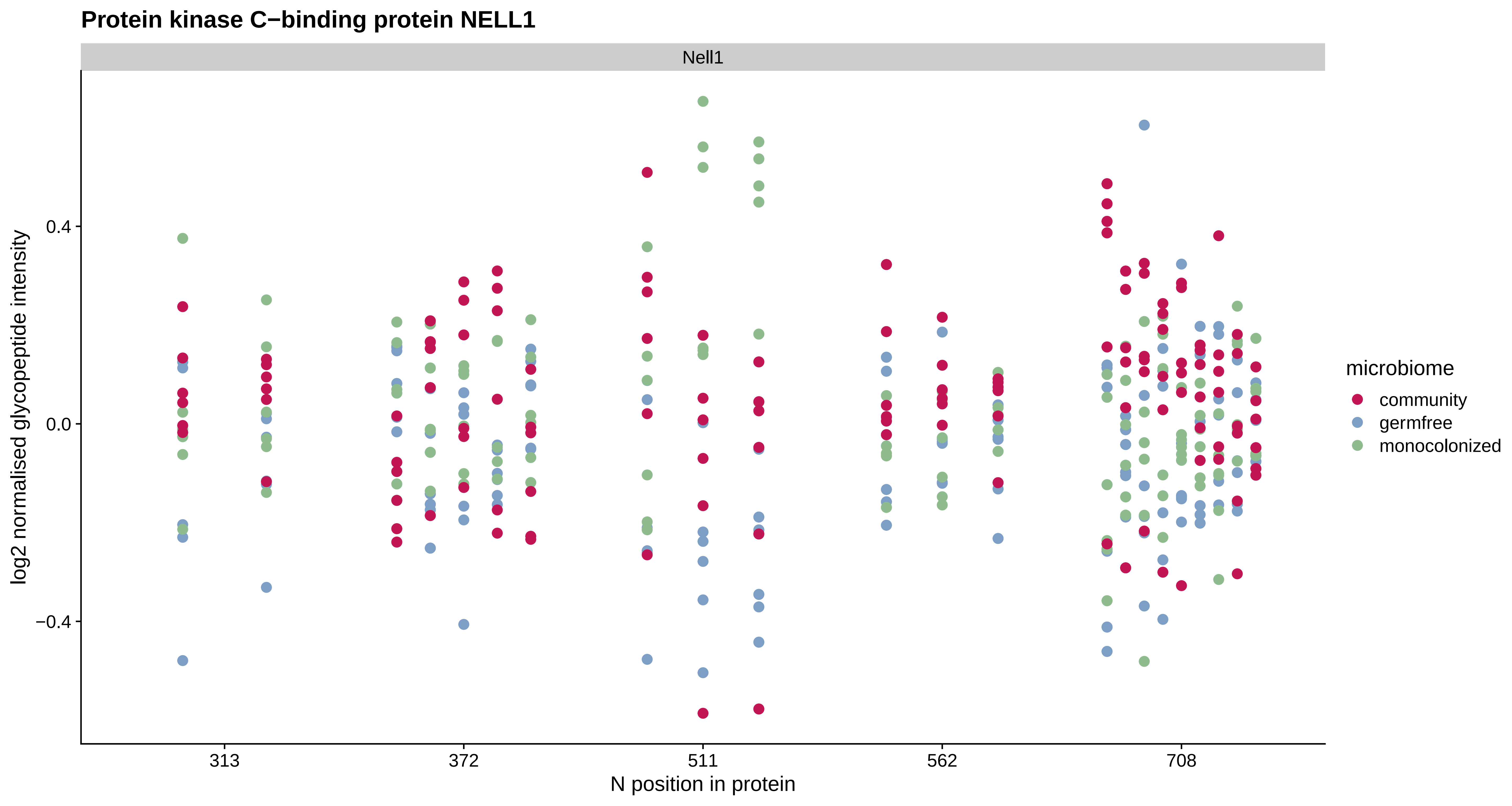
Protein kinase C−binding protein NELL1
Nell1
log2 normalised glycopeptide intensity
0.4
0.0
−0.4
313
372
511
562
708
N position in protein
microbiome
community
germfree
monocolonized

#### Low−density lipoprotein receptor−related protein 8

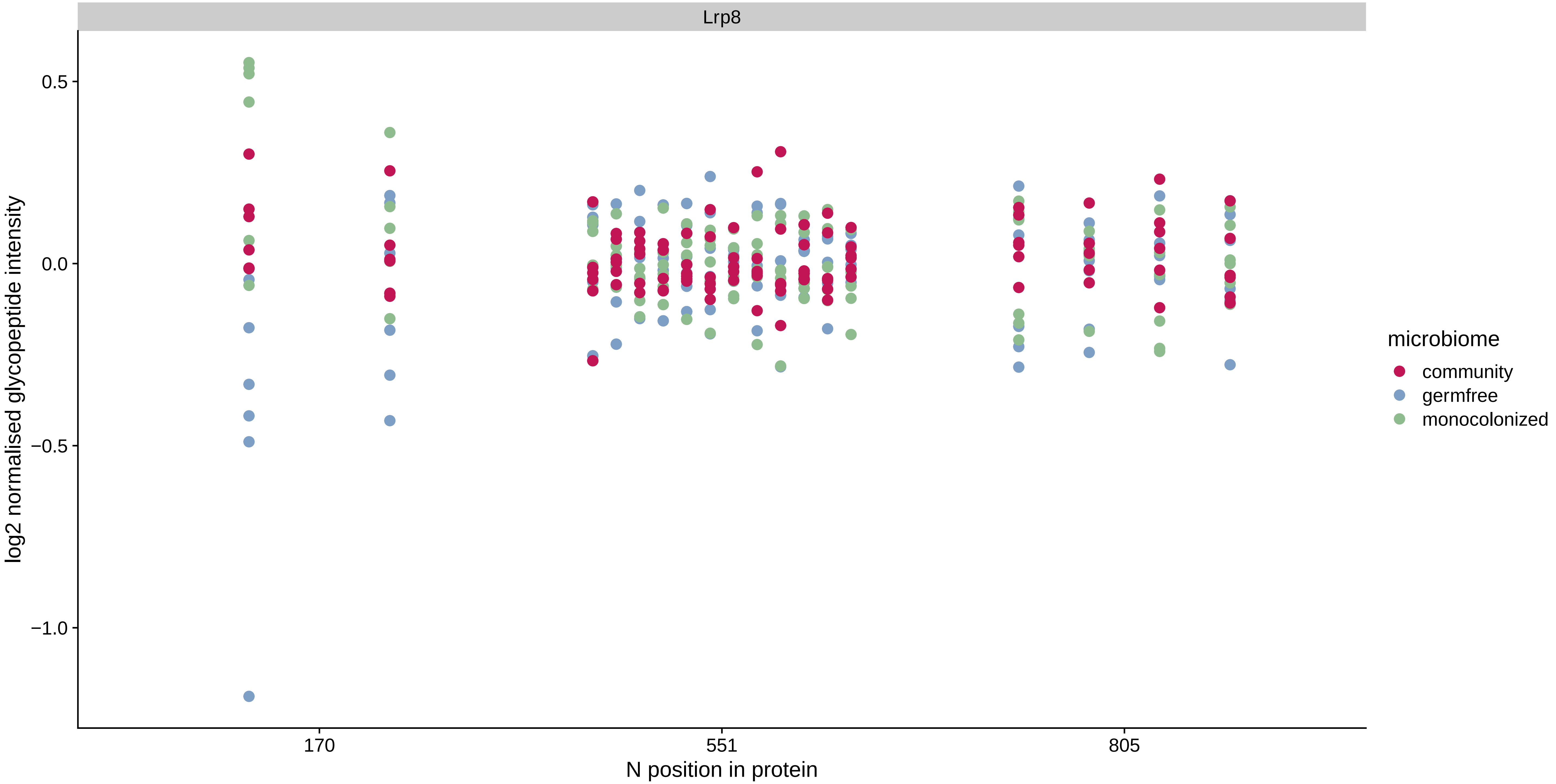

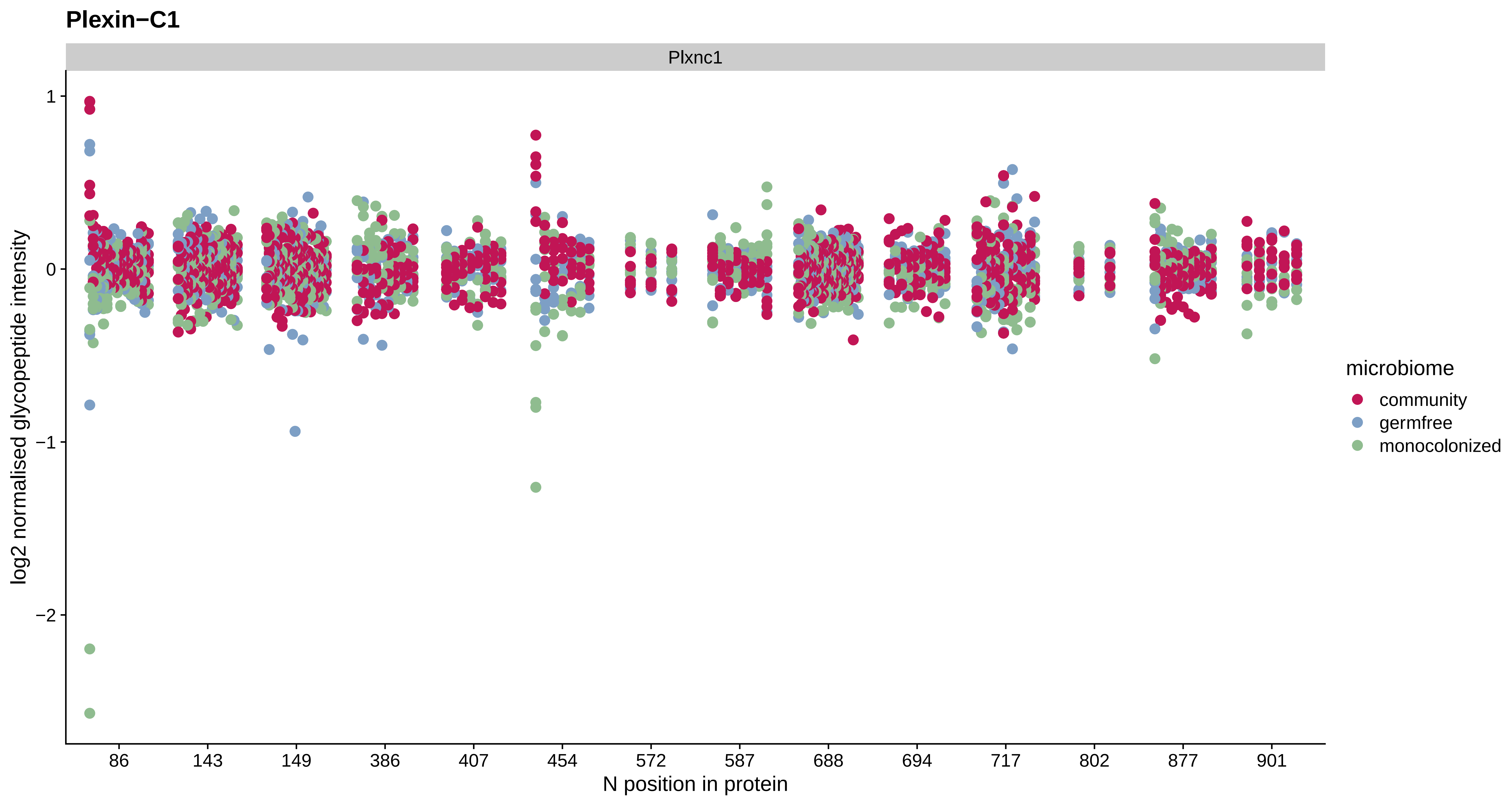
Plexin−C1
Plxnc1
log2 normalised glycopeptide intensity
1
0
−1
−2
86
143
149
386
407
454
572
587
688
694
717
802
877
901
N position in protein
microbiome
community
germfree
monocolonized

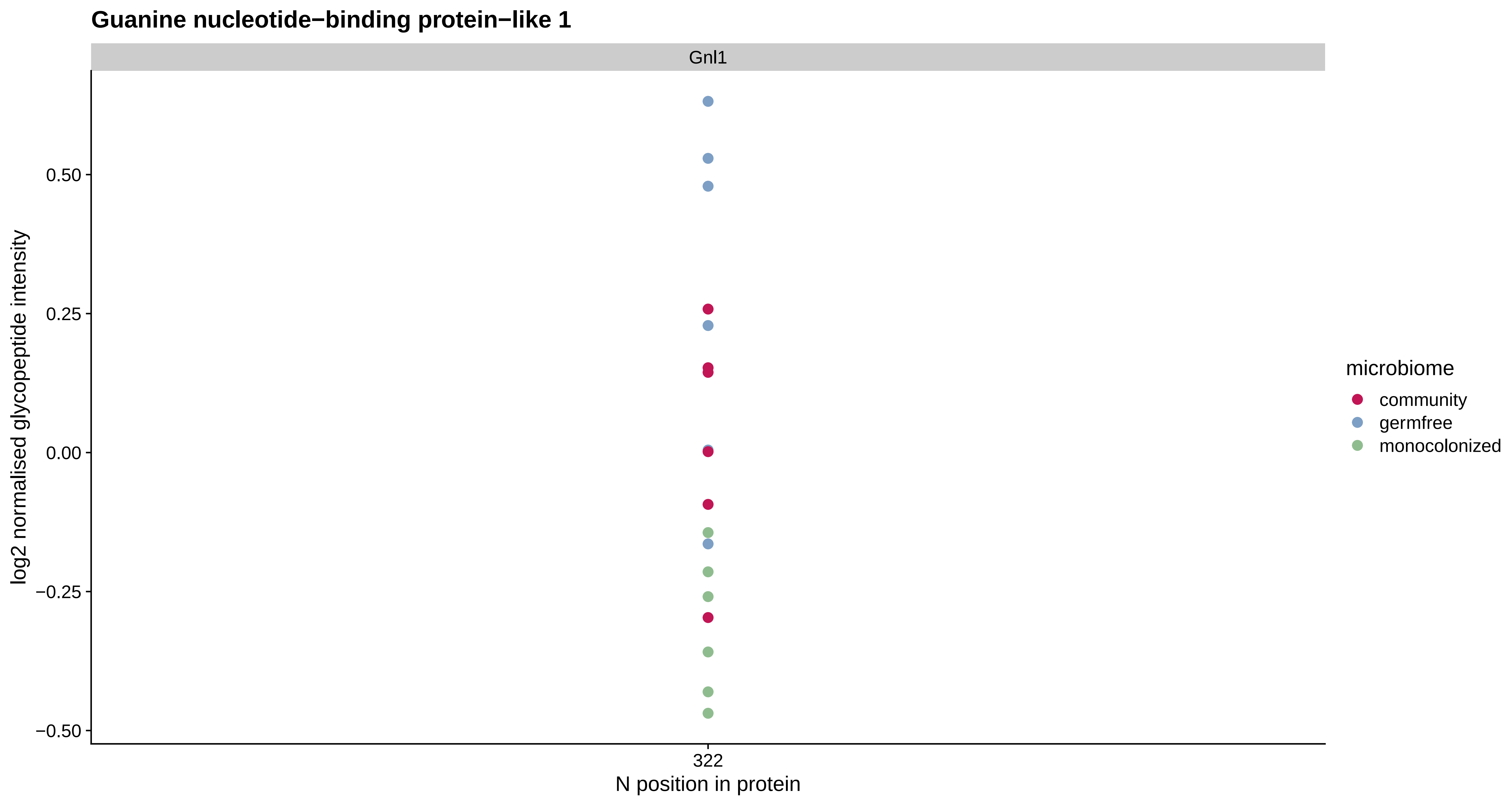
Guanine nucleotide−binding protein−like 1
Gnl1
log2 normalised glycopeptide intensity
0.50
0.25
0.00
−0.25
−0.50
322
N position in protein
microbiome
community
germfree
monocolonized

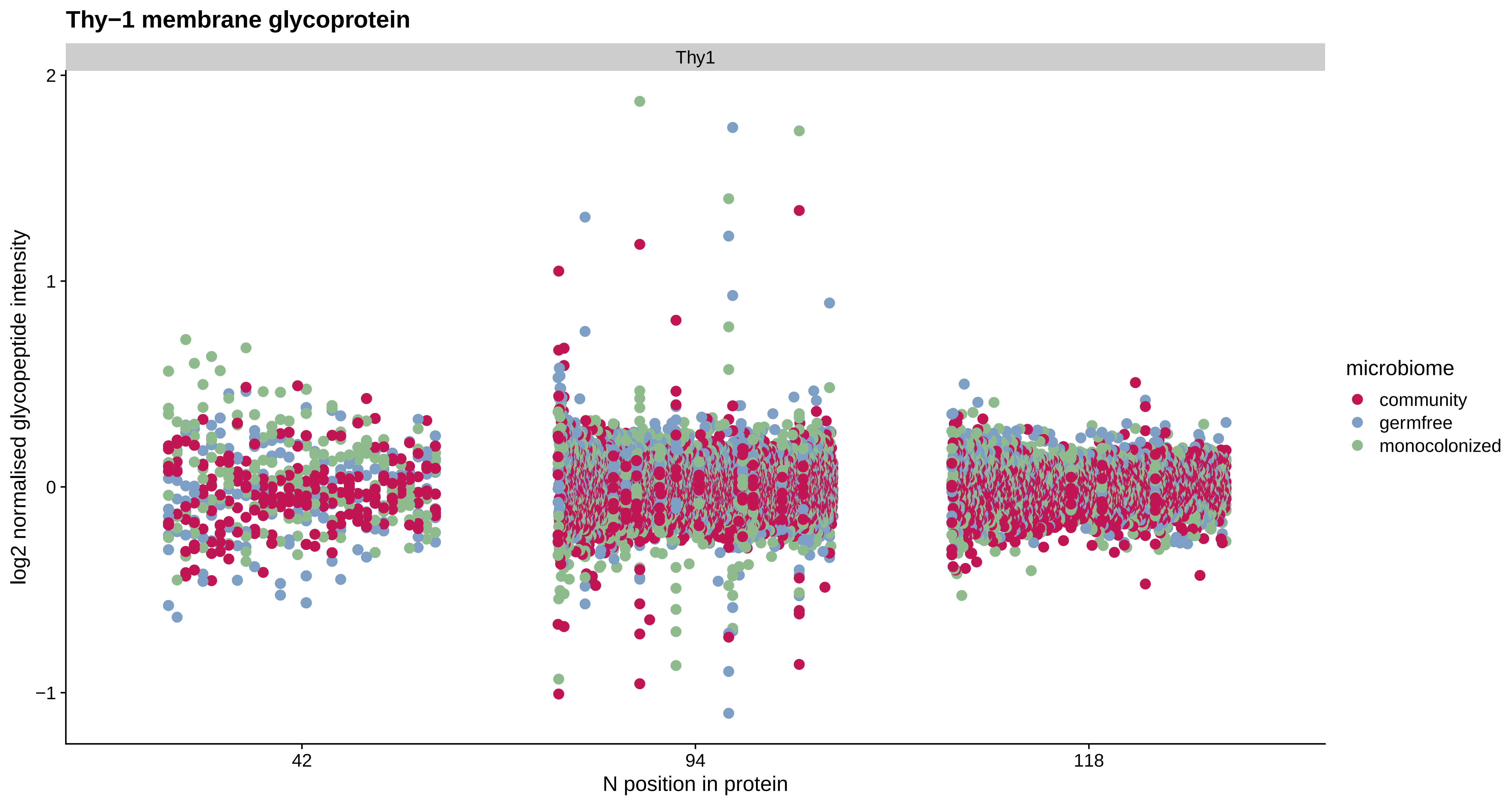
Thy−1 membrane glycoprotein
Thy1
2
1
0
−1
log2 normalised glycopeptide intensity
42
94
118
N position in protein
microbiome
community
germfree
monocolonized

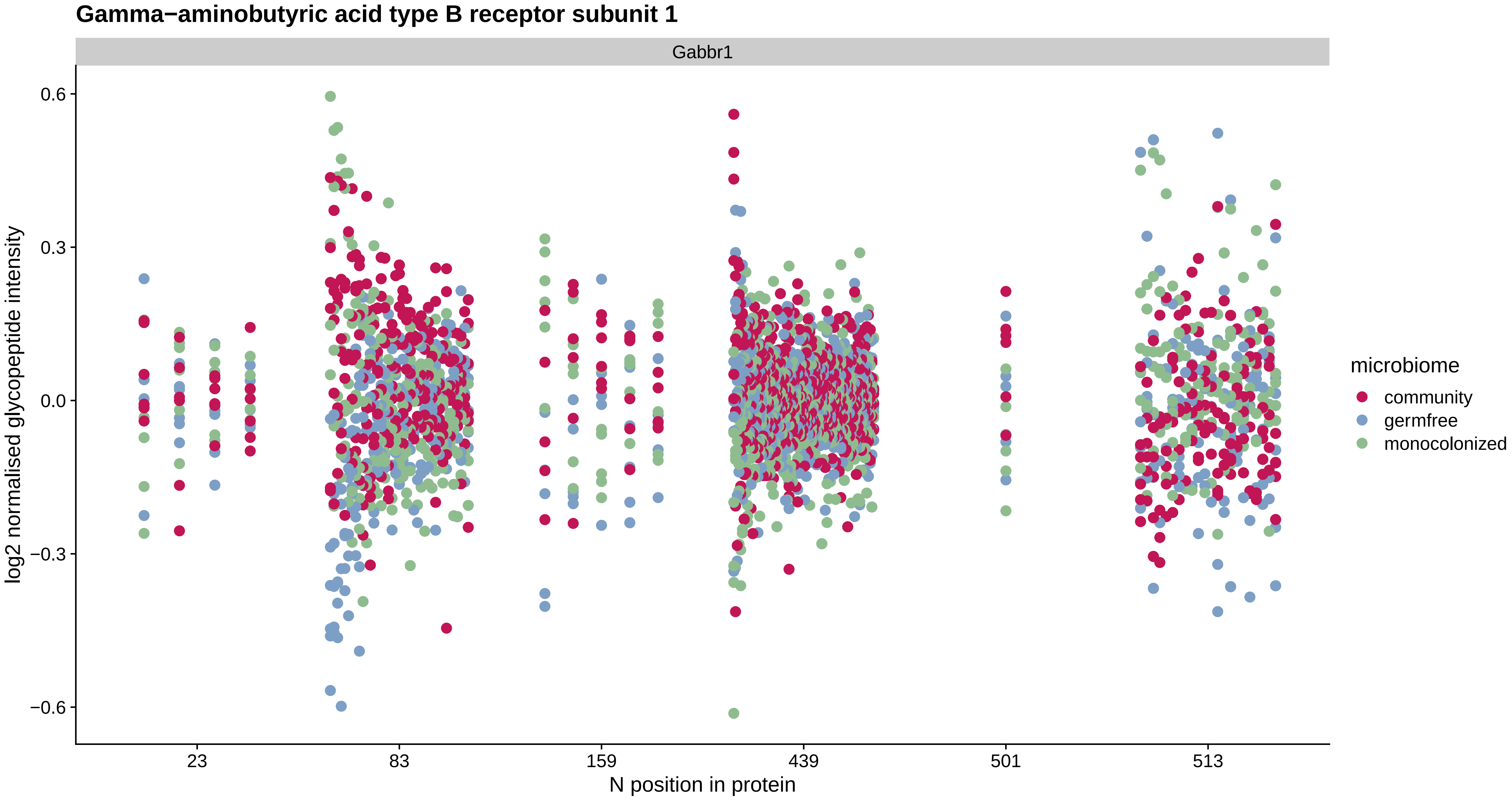
Gamma−aminobutyric acid type B receptor subunit 1
Gabbr1
log2 normalised glycopeptide intensity
0.6
0.3
0.0
−0.3
−0.6
23
83
159
439
501
513
N position in protein
microbiome
community
germfree
monocolonized

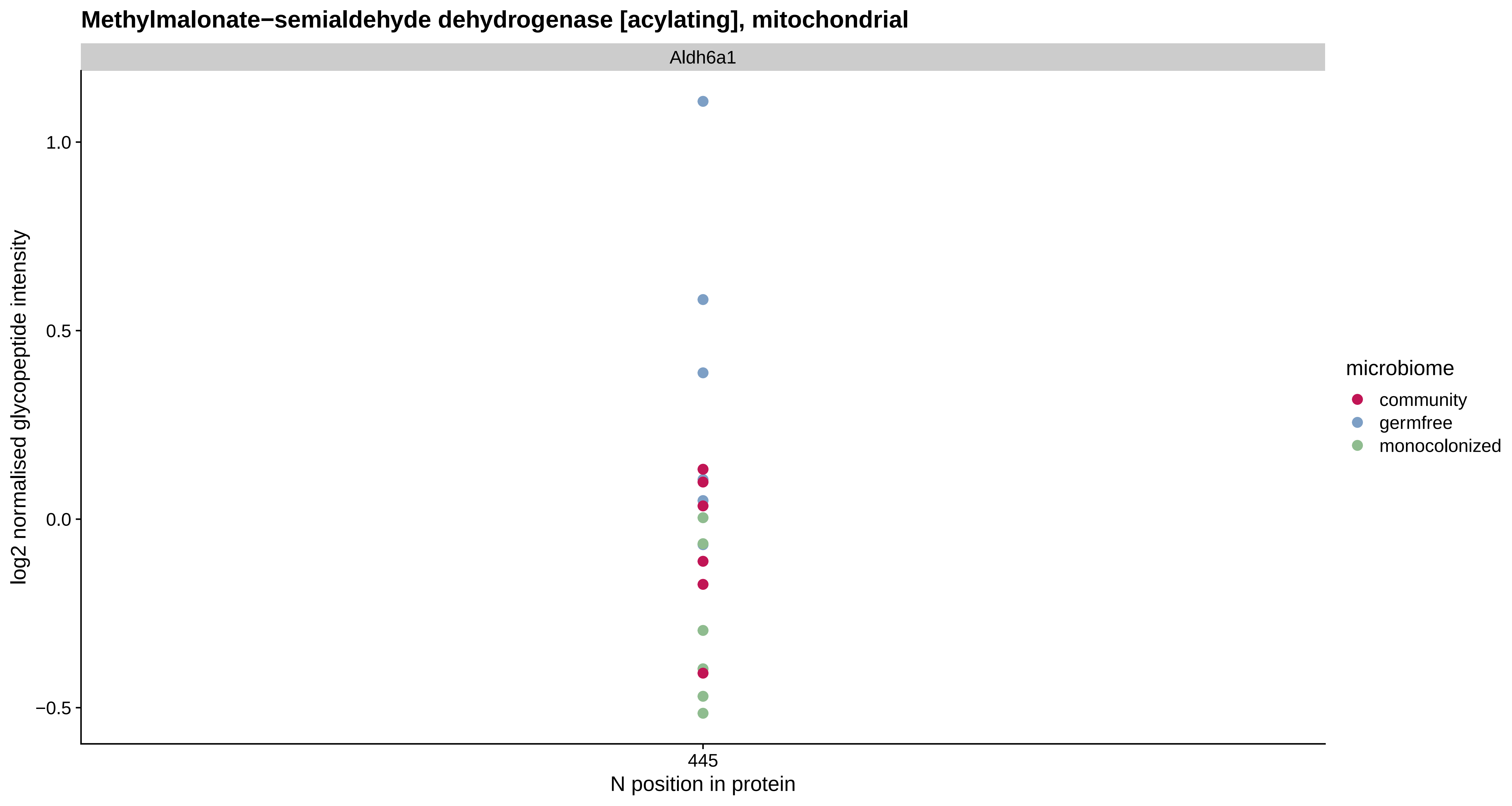

Methylmalonate−semialdehyde dehydrogenase [acylating], mitochondrial
Aldh6a1
1.0
0.5
0.0
−0.5
log2 normalised glycopeptide intensity
445
N position in protein
microbiome
community
germfree
monocolonized

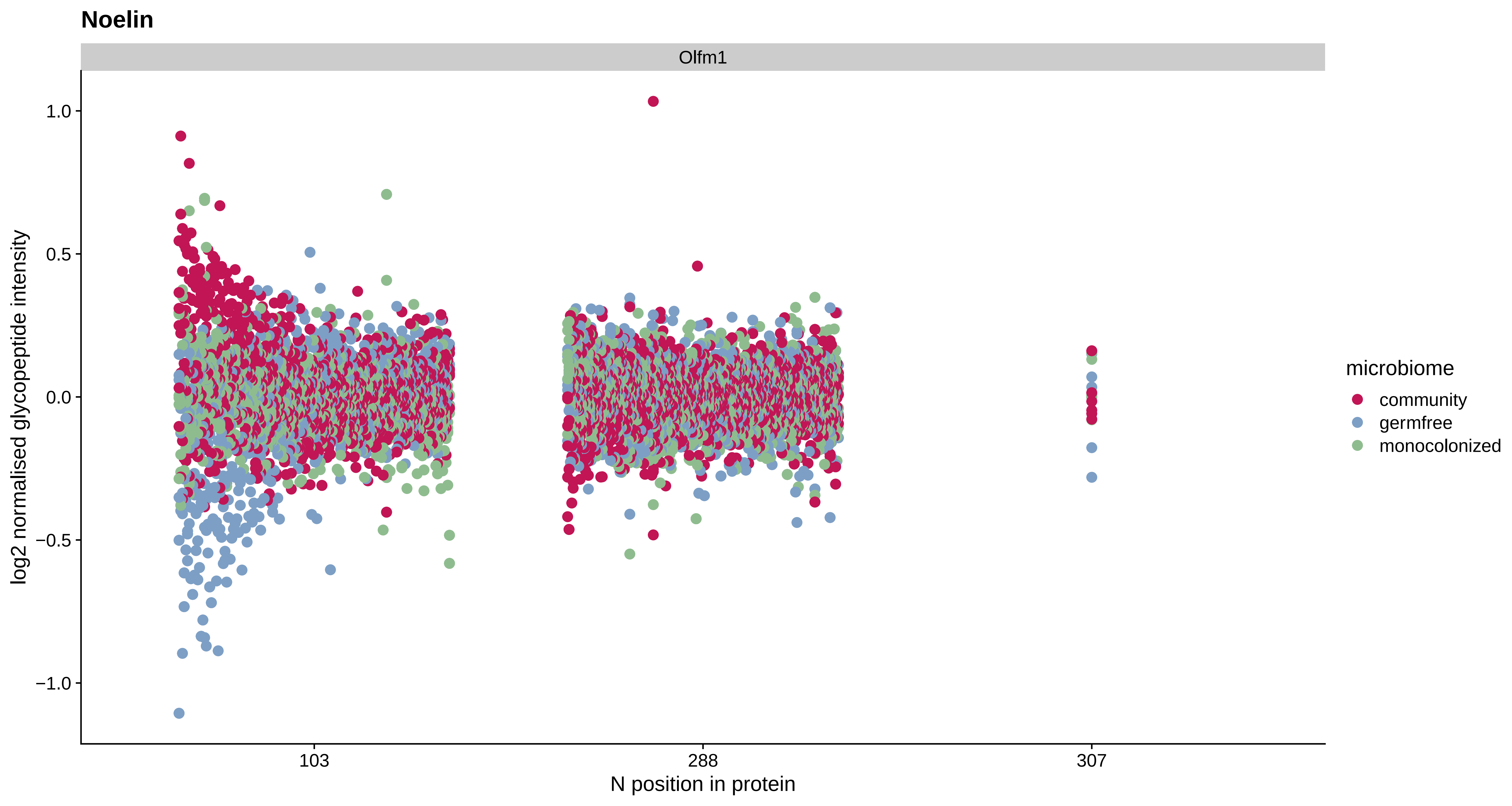

Noelin
Olfm1
log2 normalised glycopeptide intensity
1.0
0.5
0.0
−0.5
−1.0
103
288
307
N position in protein
microbiome
community
germfree
monocolonized

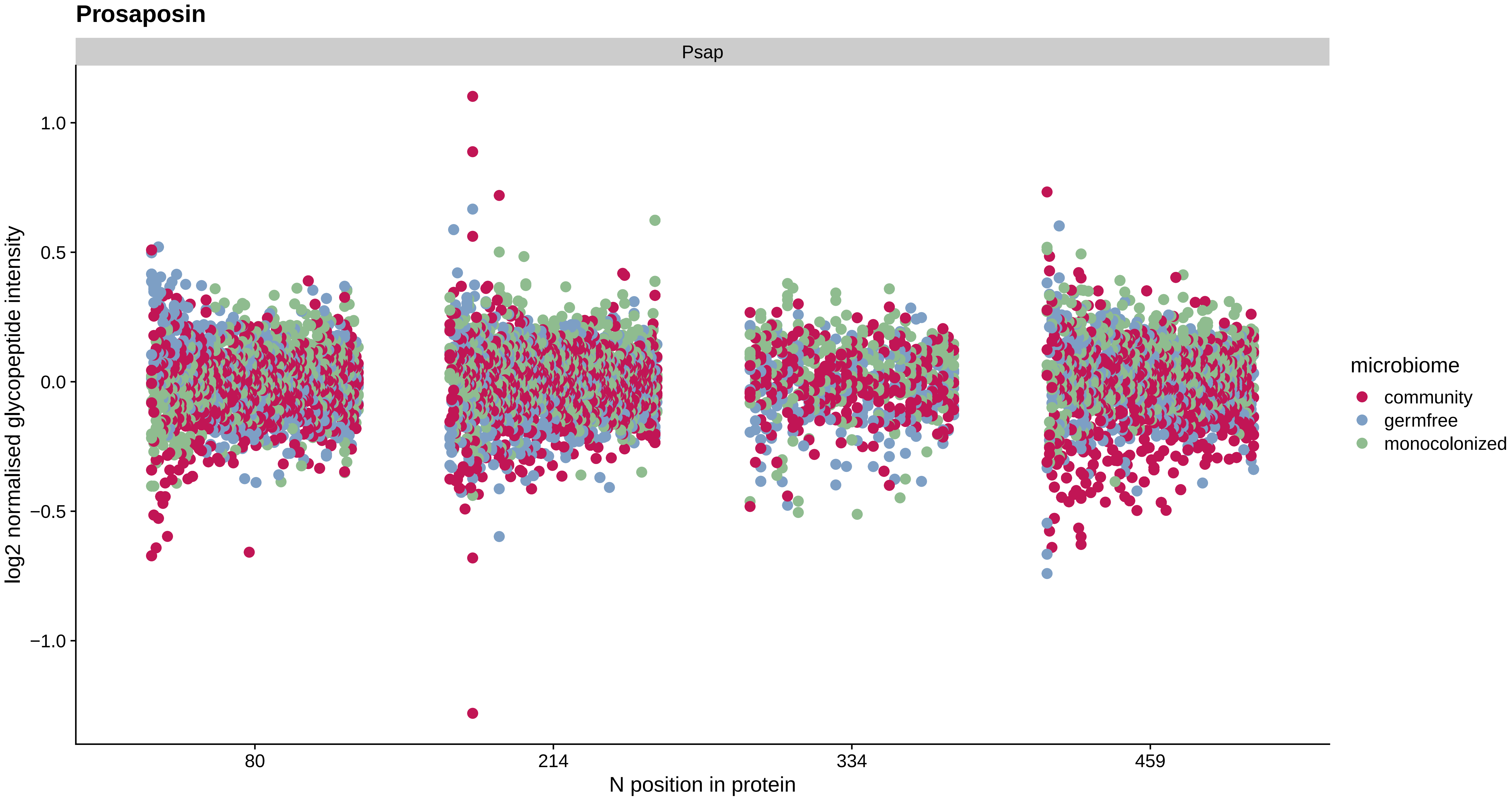
Prosaposin
Psap
1.0
0.5
0.0
−0.5
−1.0
log2 normalised glycopeptide intensity
80
214
334
459
N position in protein
microbiome
community
germfree
monocolonized

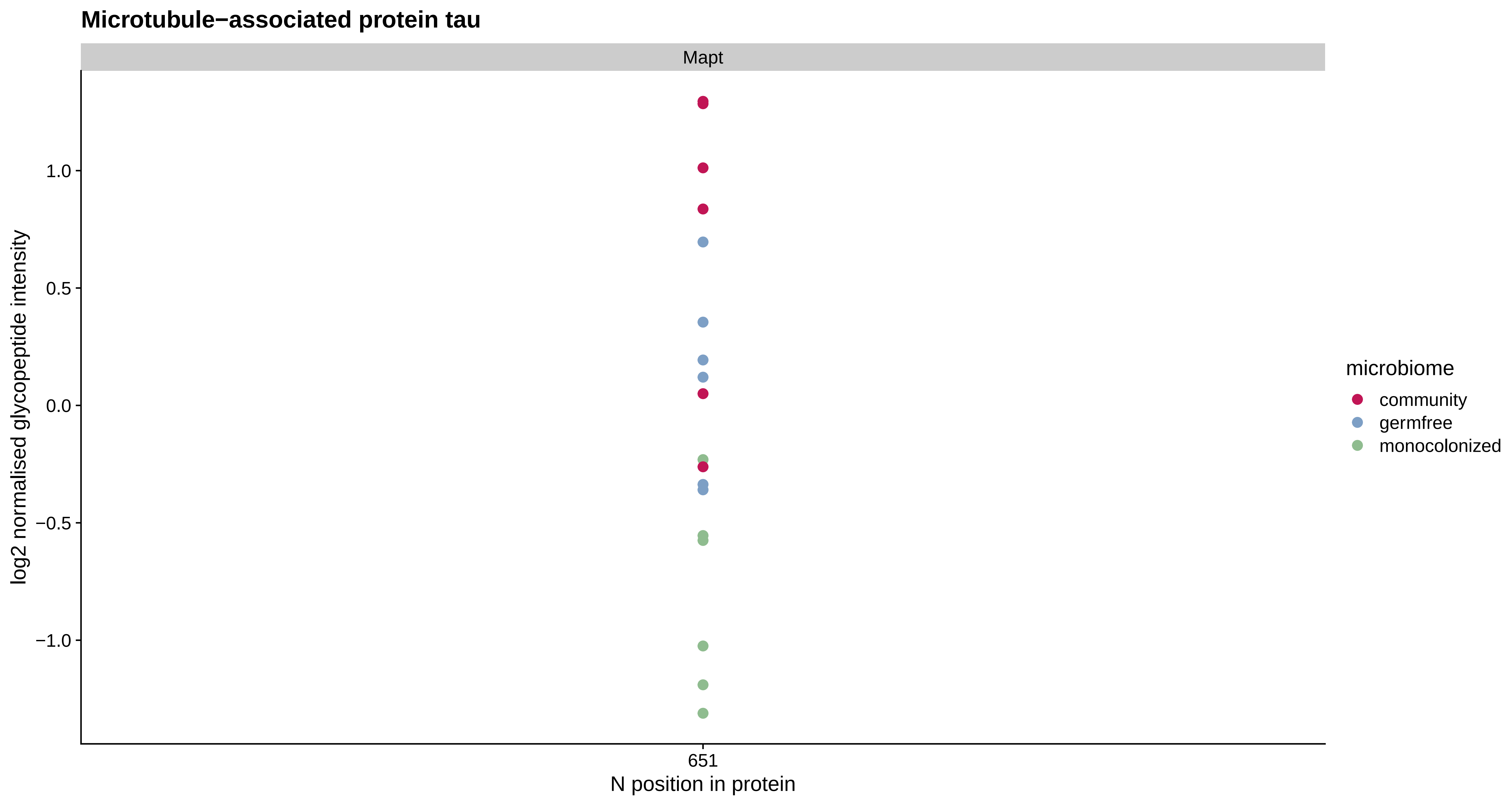
Microtubule−associated protein tau
Mapt
log2 normalised glycopeptide intensity
1.0
0.5
0.0
−0.5
−1.0
651
N position in protein
microbiome
community
germfree
monocolonized

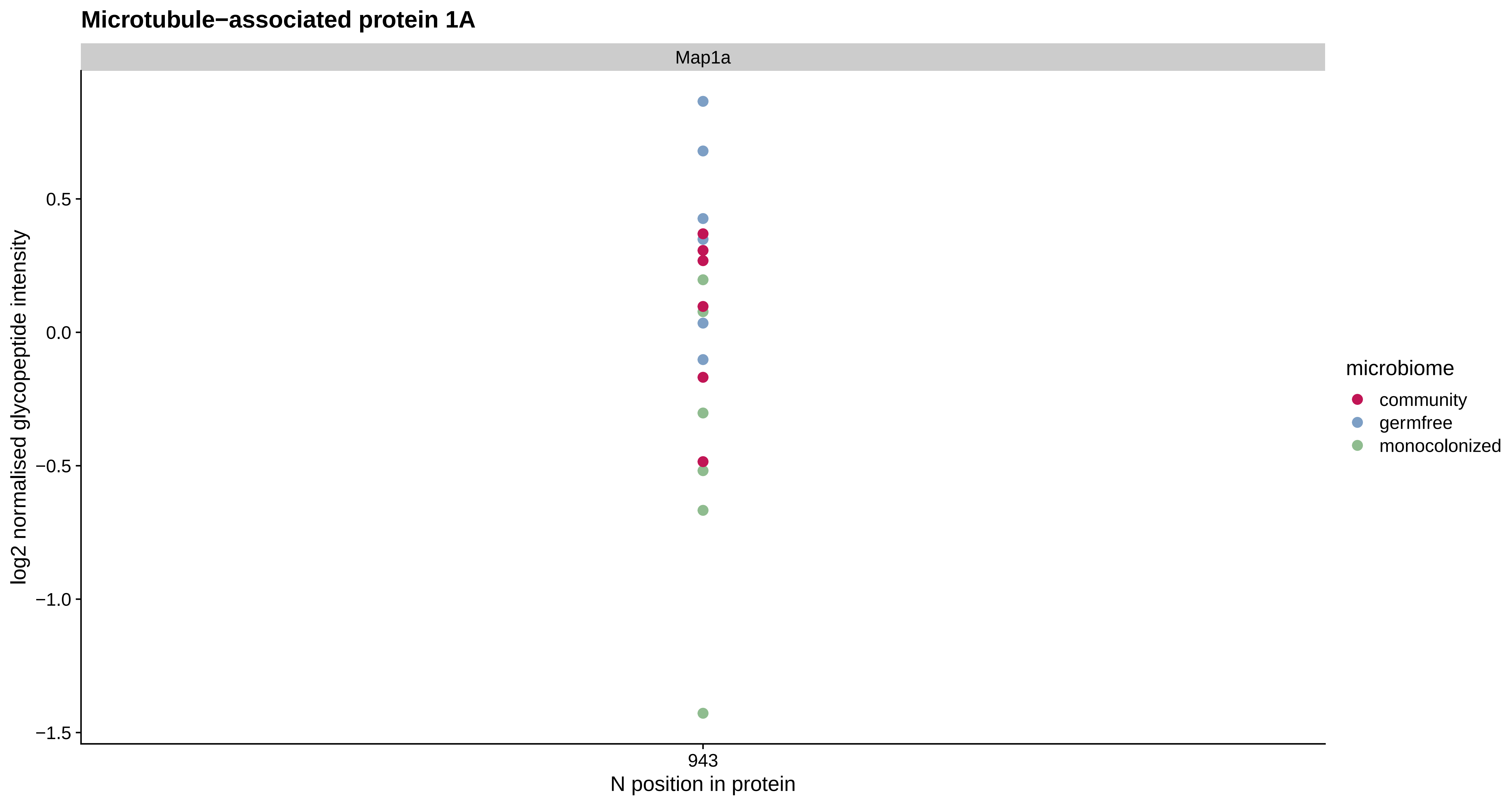
Microtubule−associated protein 1A
Map1a
0.5
0.0
−0.5
−1.0
−1.5
log2 normalised glycopeptide intensity
943
N position in protein
microbiome
community
germfree
monocolonized

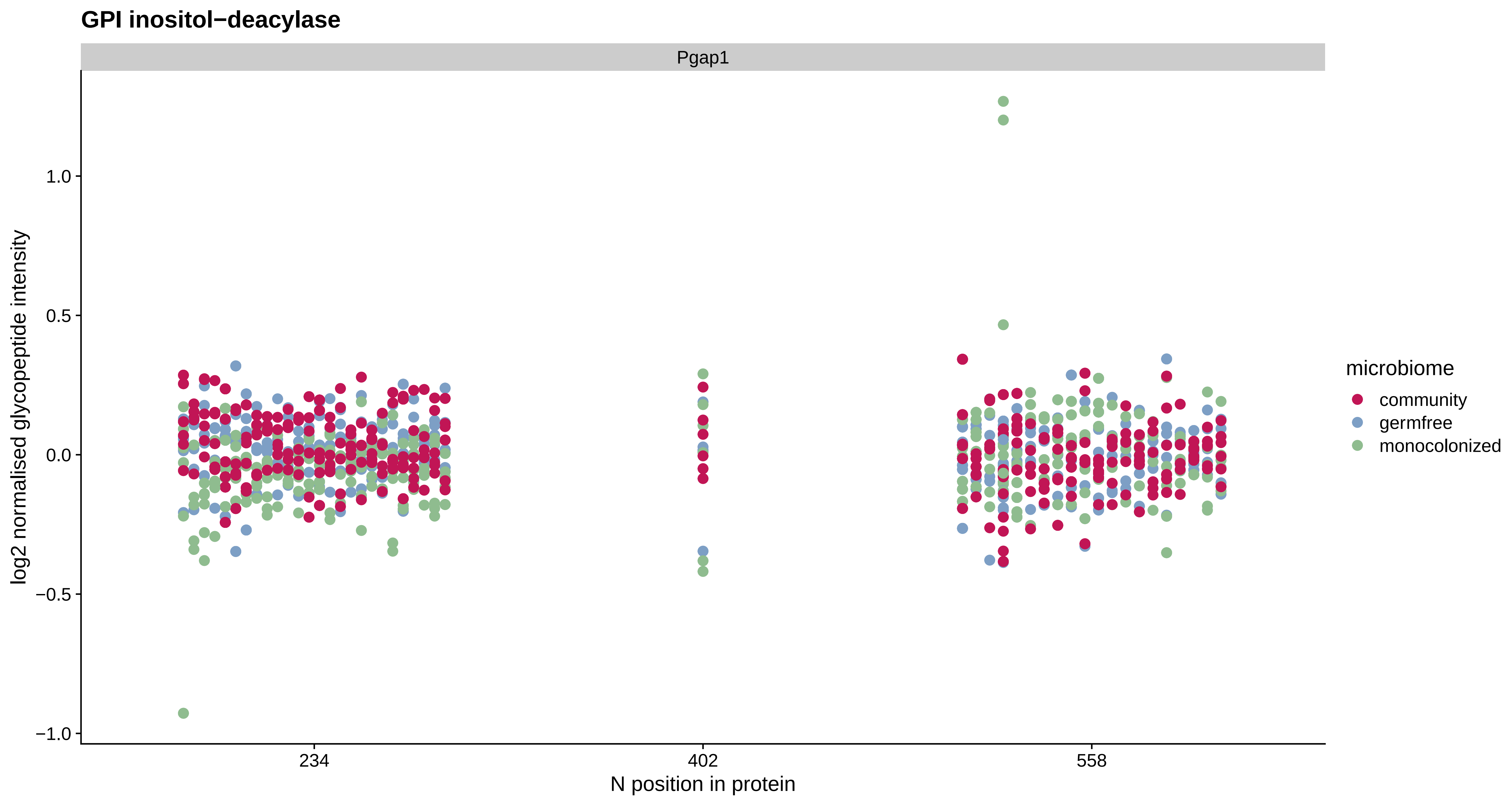
GPI inositol−deacylase
Pgap1
log2 normalised glycopeptide intensity
1.0
0.5
0.0
−0.5
−1.0
234
402
558
N position in protein
microbiome
community
germfree
monocolonized

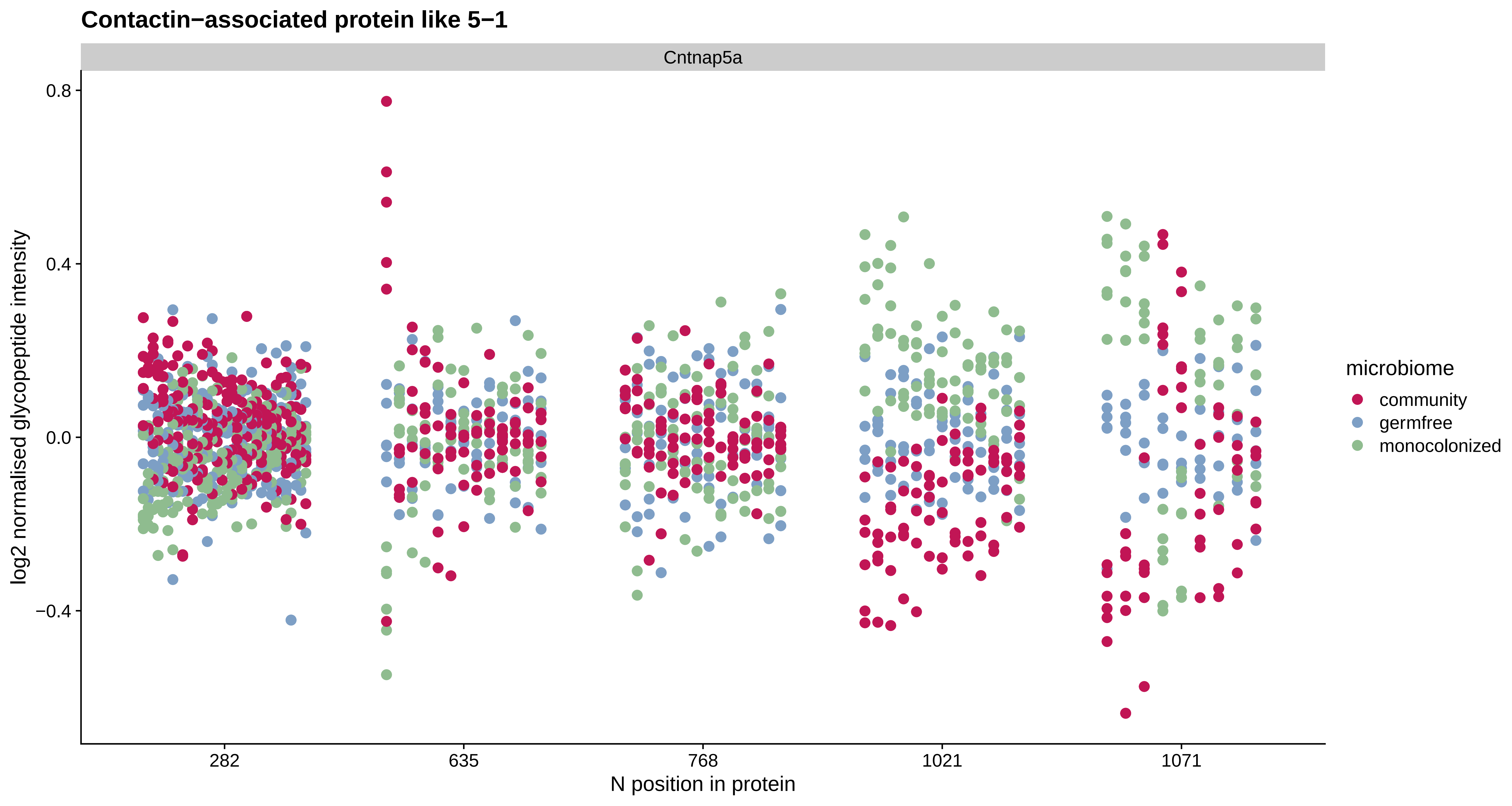

Contactin−associated protein like 5−1
Cntnap5a
0.8
0.4
0.0
−0.4
log2 normalised glycopeptide intensity
282
635
768
1021
1071
N position in protein
microbiome
community
germfree
monocolonized

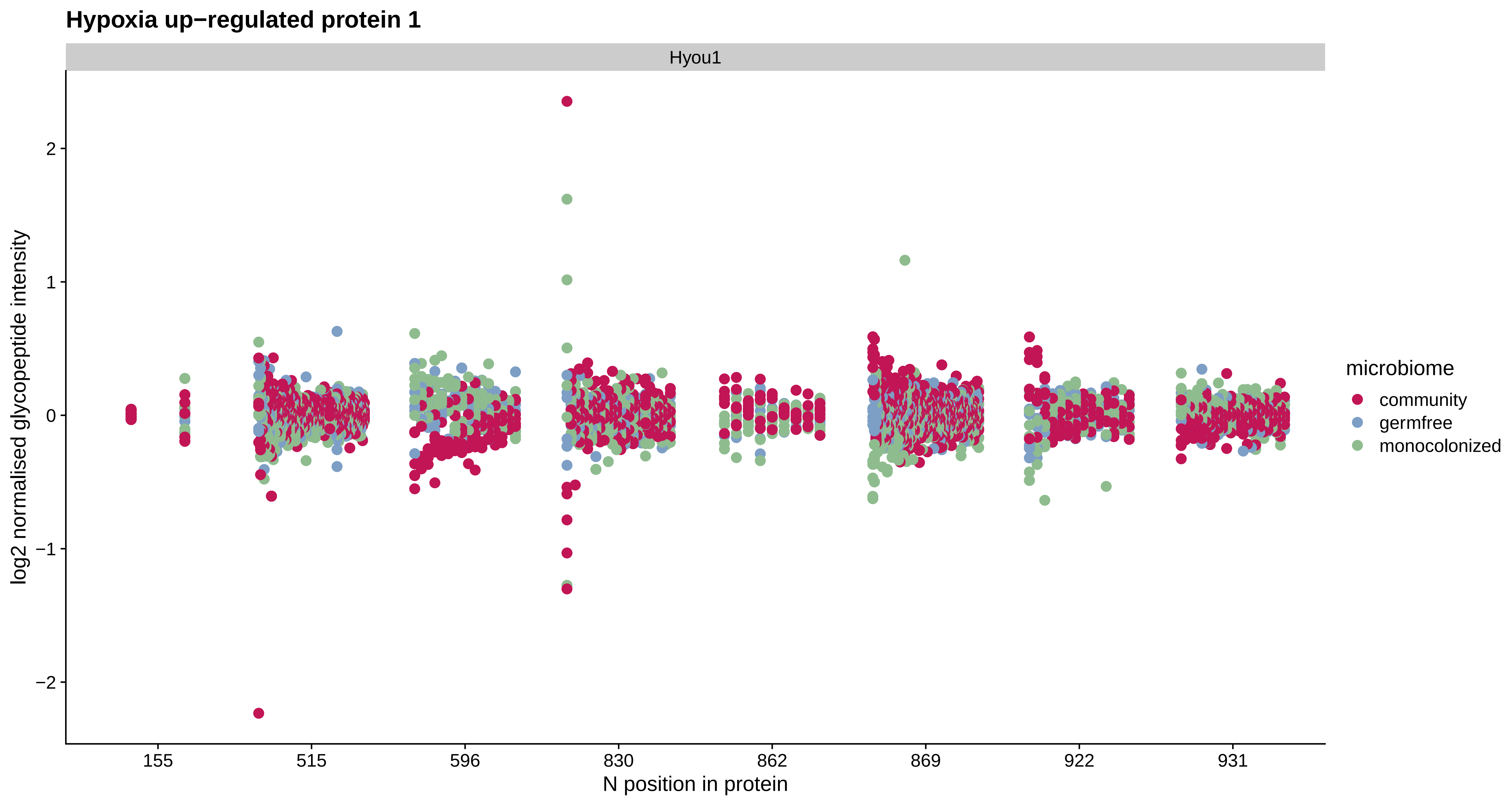
Hypoxia up−regulated protein 1
Hyou1
log2 normalised glycopeptide intensity
2
1
0
−1
−2
155
515
596
830
862
869
922
931
N position in protein
microbiome
community
germfree
monocolonized

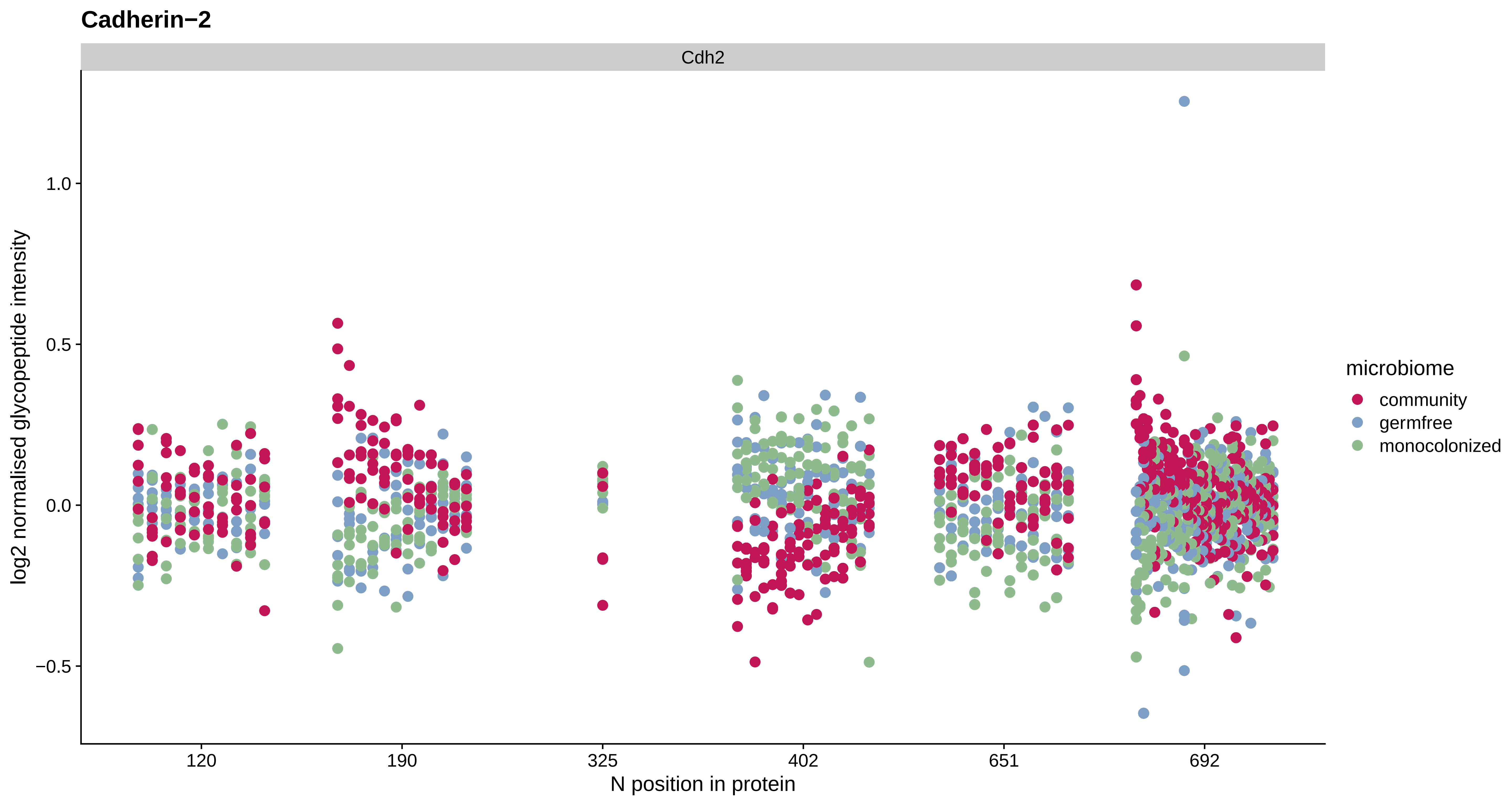

Cadherin−2
Cdh2
log2 normalised glycopeptide intensity
1.0
0.5
0.0
−0.5
120
190
325
402
651
692
N position in protein
microbiome
community
germfree
monocolonized

#### Receptor−type tyrosine−protein phosphatase N2

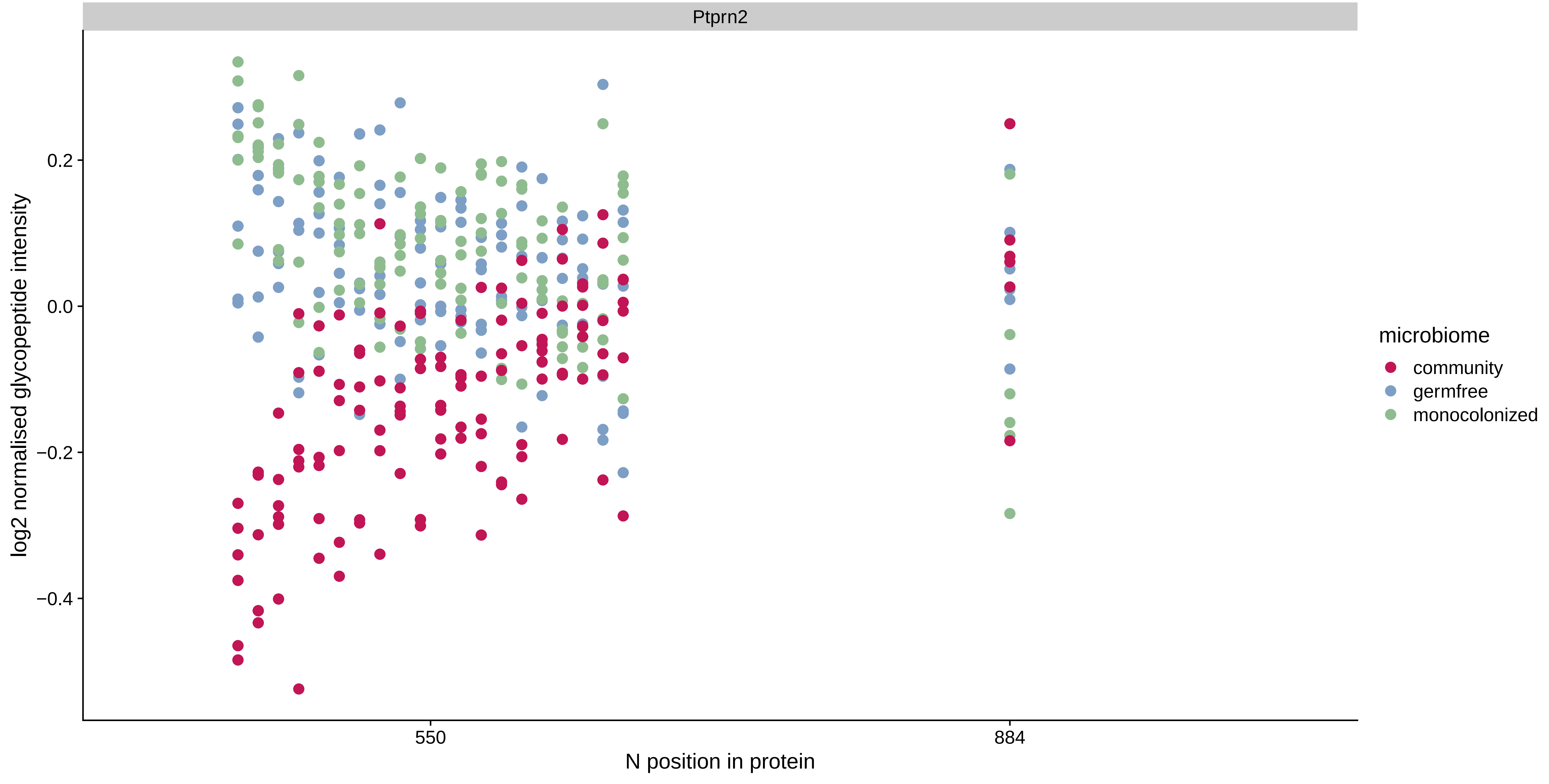

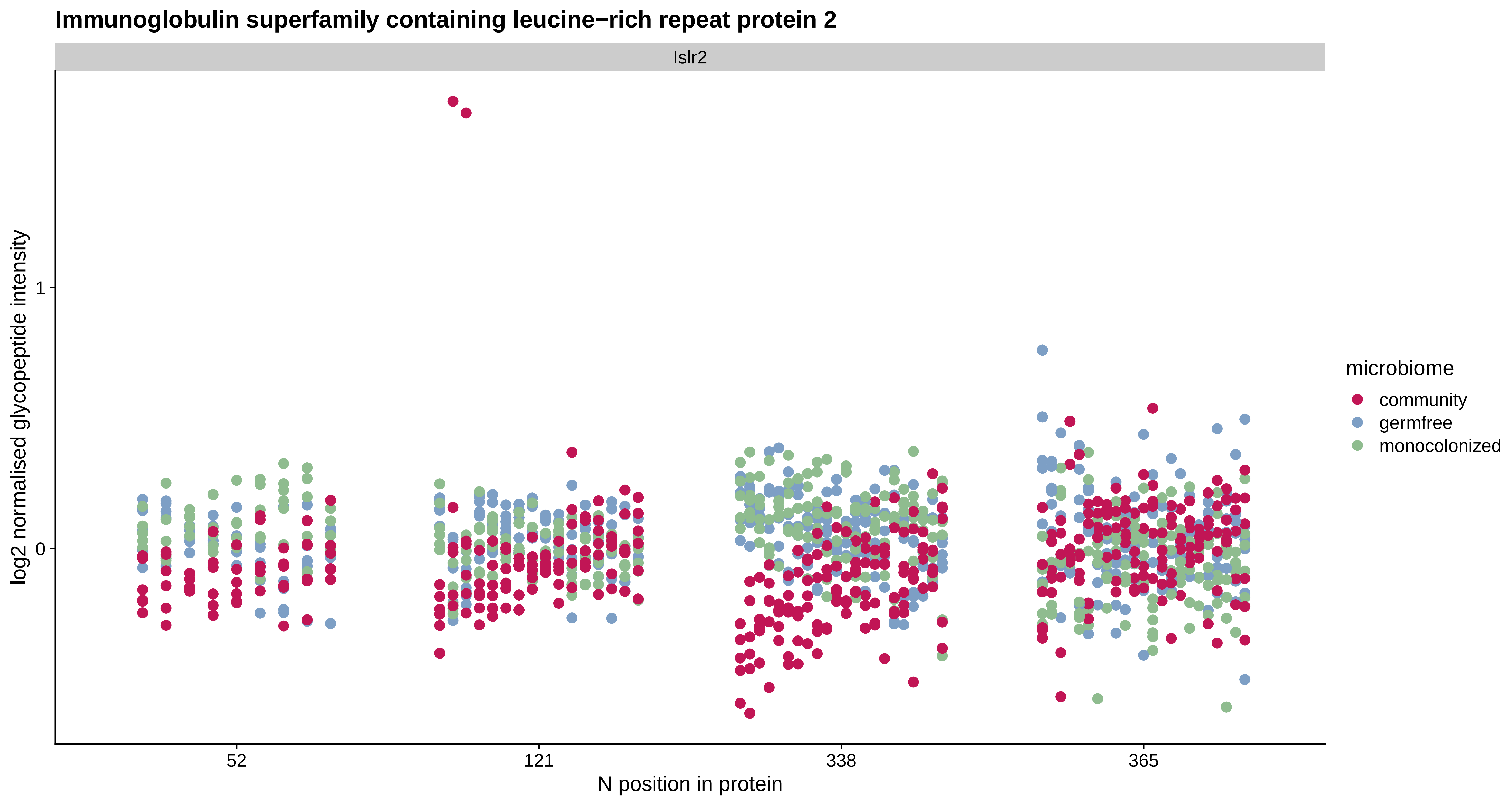

Immunoglobulin superfamily containing leucine−rich repeat protein 2
Islr2
log2 normalised glycopeptide intensity
1
0
52
121
338
365
N position in protein
microbiome
community
germfree
monocolonized

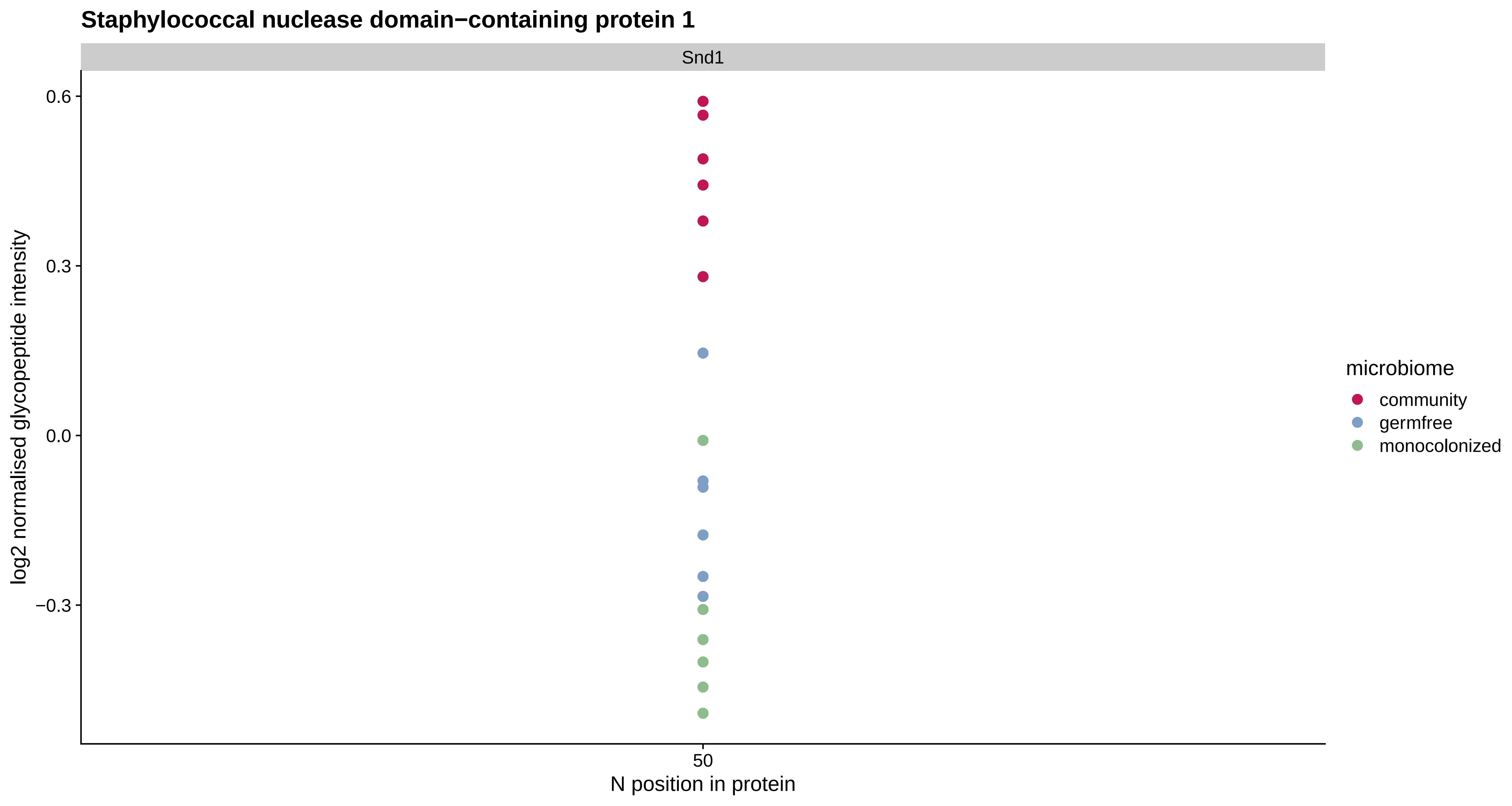

Staphylococcal nuclease domain−containing protein 1
Snd1
0.6
0.3
0.0
−0.3
log2 normalised glycopeptide intensity
50
N position in protein
microbiome
community
germfree
monocolonized

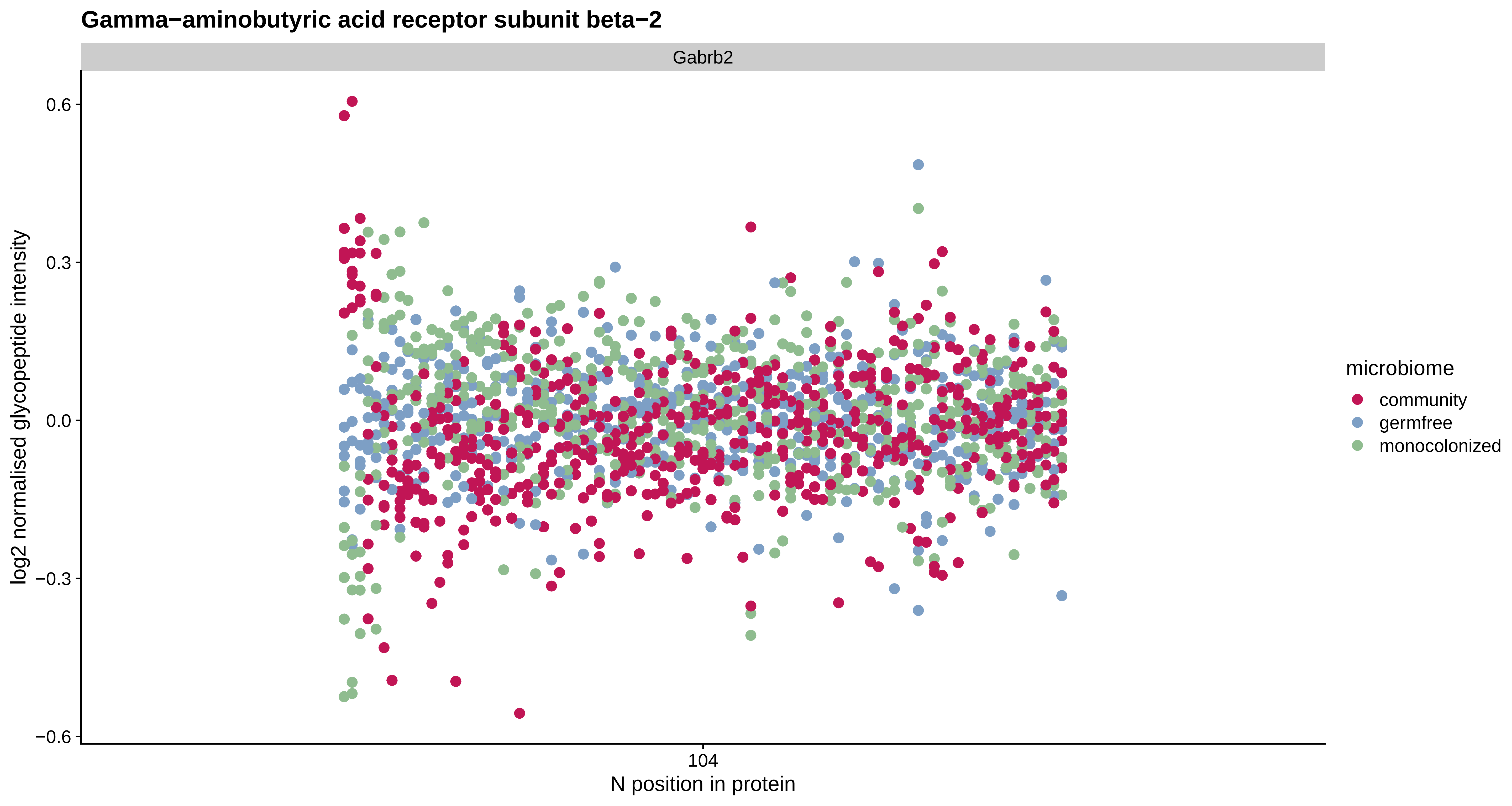
Gamma−aminobutyric acid receptor subunit beta−2
Gabrb2
0.6
0.3
0.0
−0.3
−0.6
log2 normalised glycopeptide intensity
104
N position in protein
microbiome
community
germfree
monocolonized

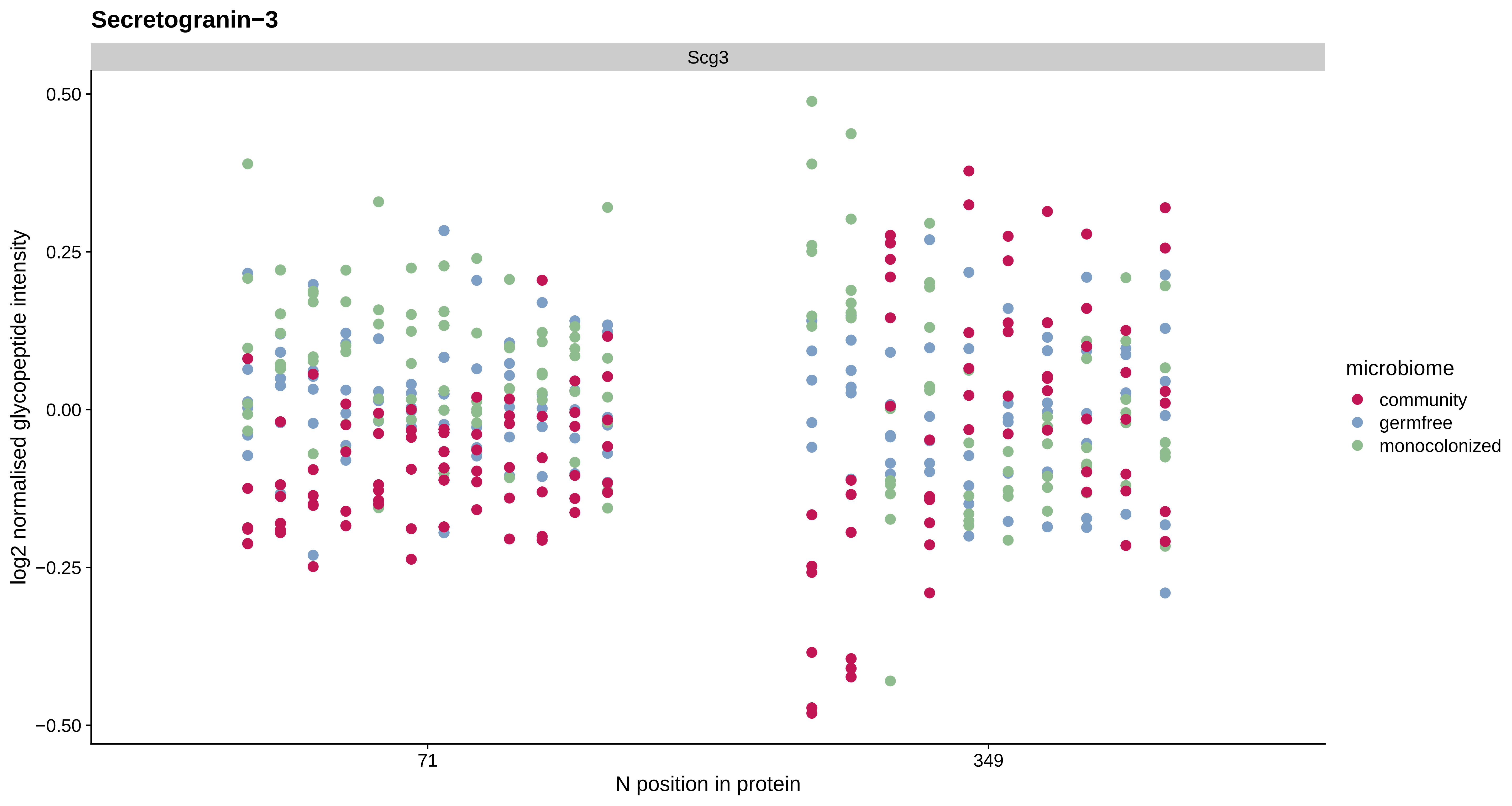

Secretogranin−3
Scg3
0.50
0.25
0.00
−0.25
−0.50
log2 normalised glycopeptide intensity
71
349
N position in protein
microbiome
community
germfree
monocolonized

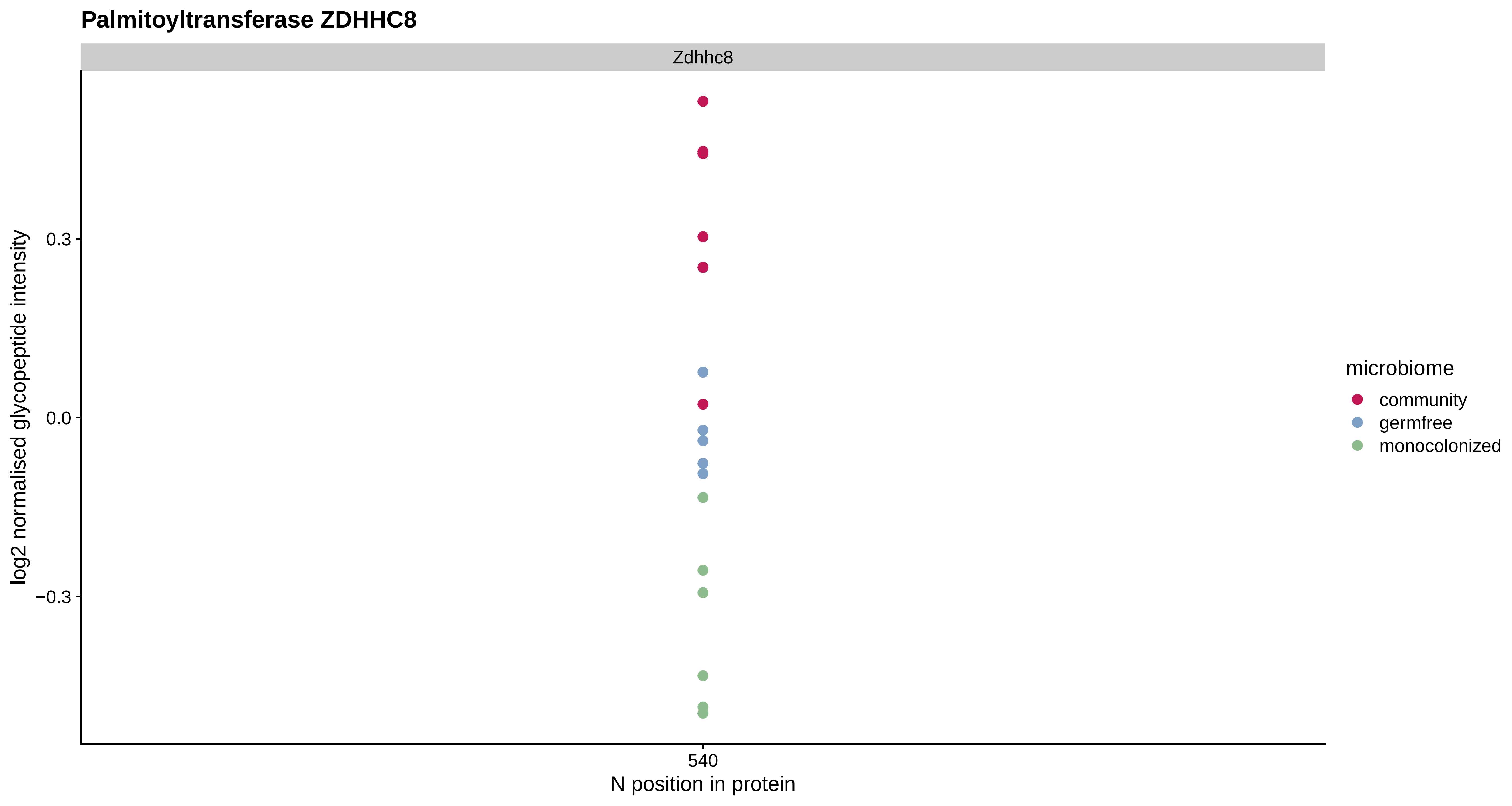

Palmitoyltransferase ZDHHC8
Zdhhc8
0.3
0.0
−0.3
log2 normalised glycopeptide intensity
540
N position in protein
microbiome
community
germfree
monocolonized

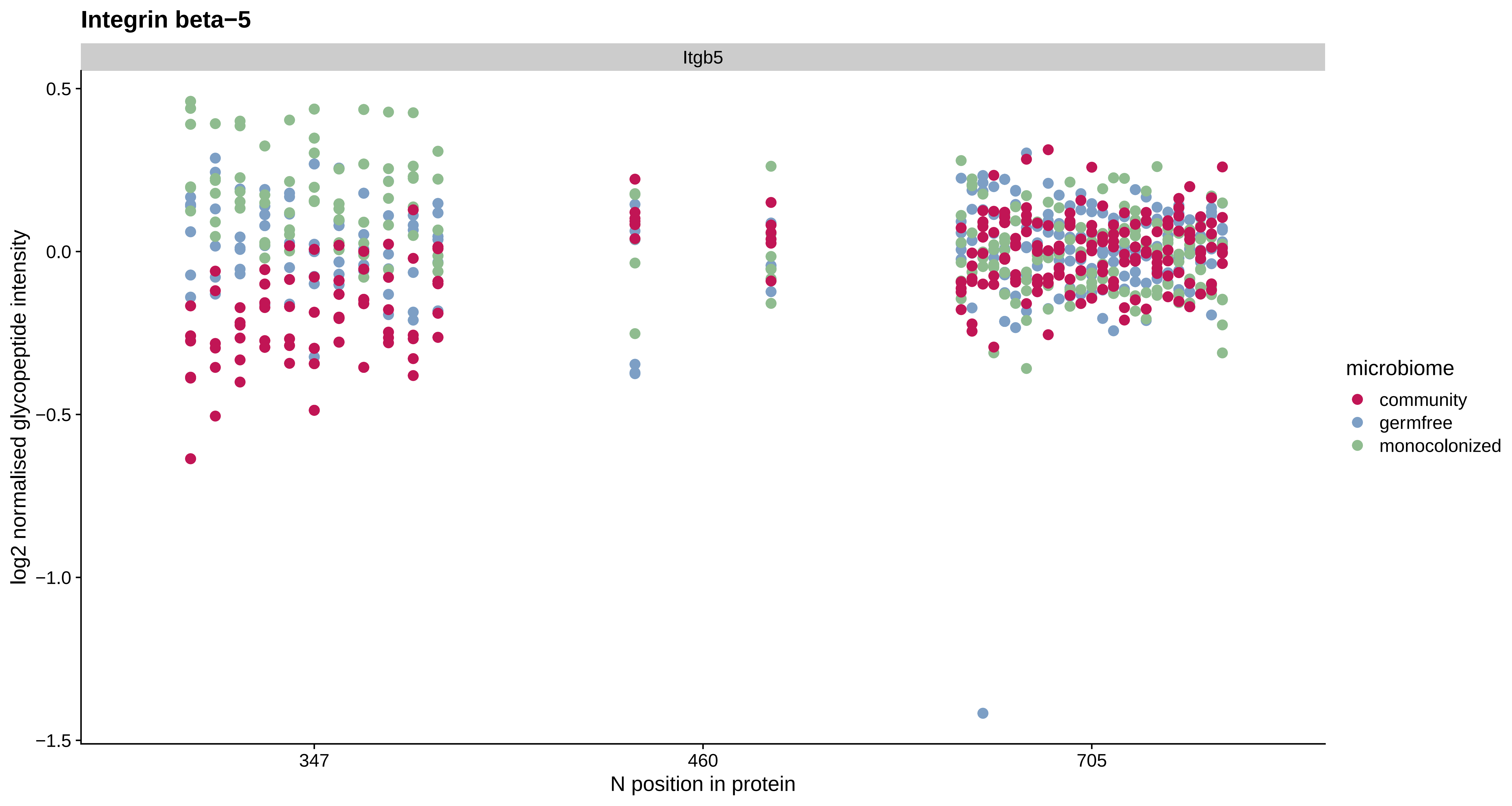
Integrin beta−5
Itgb5
0.5
0.0
−0.5
−1.0
−1.5
log2 normalised glycopeptide intensity
347
460
705
N position in protein
microbiome
community
germfree
monocolonized

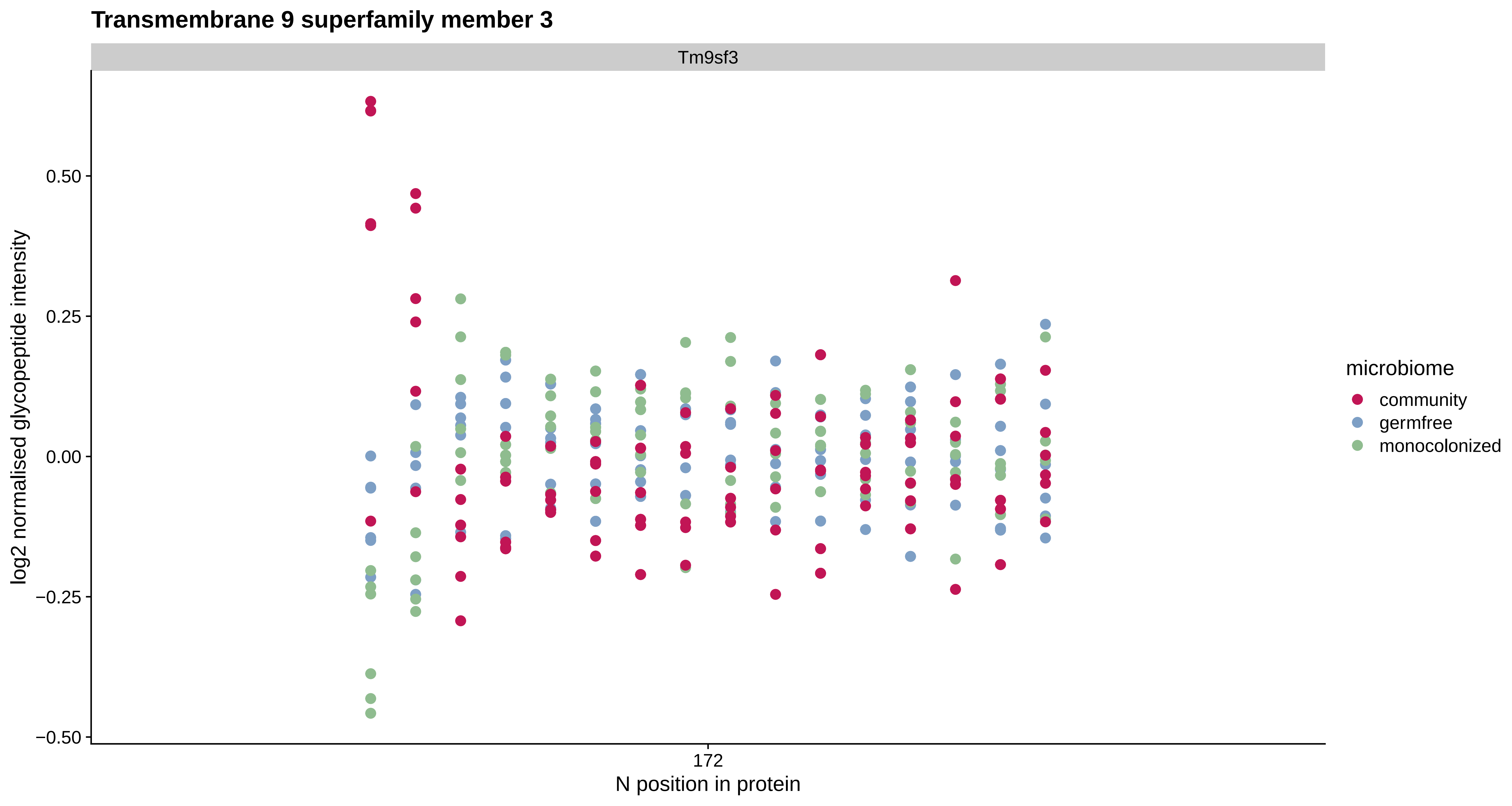
Transmembrane 9 superfamily member 3
Tm9sf3
log2 normalised glycopeptide intensity
0.50
0.25
0.00
−0.25
−0.50
172
N position in protein
microbiome
community
germfree
monocolonized

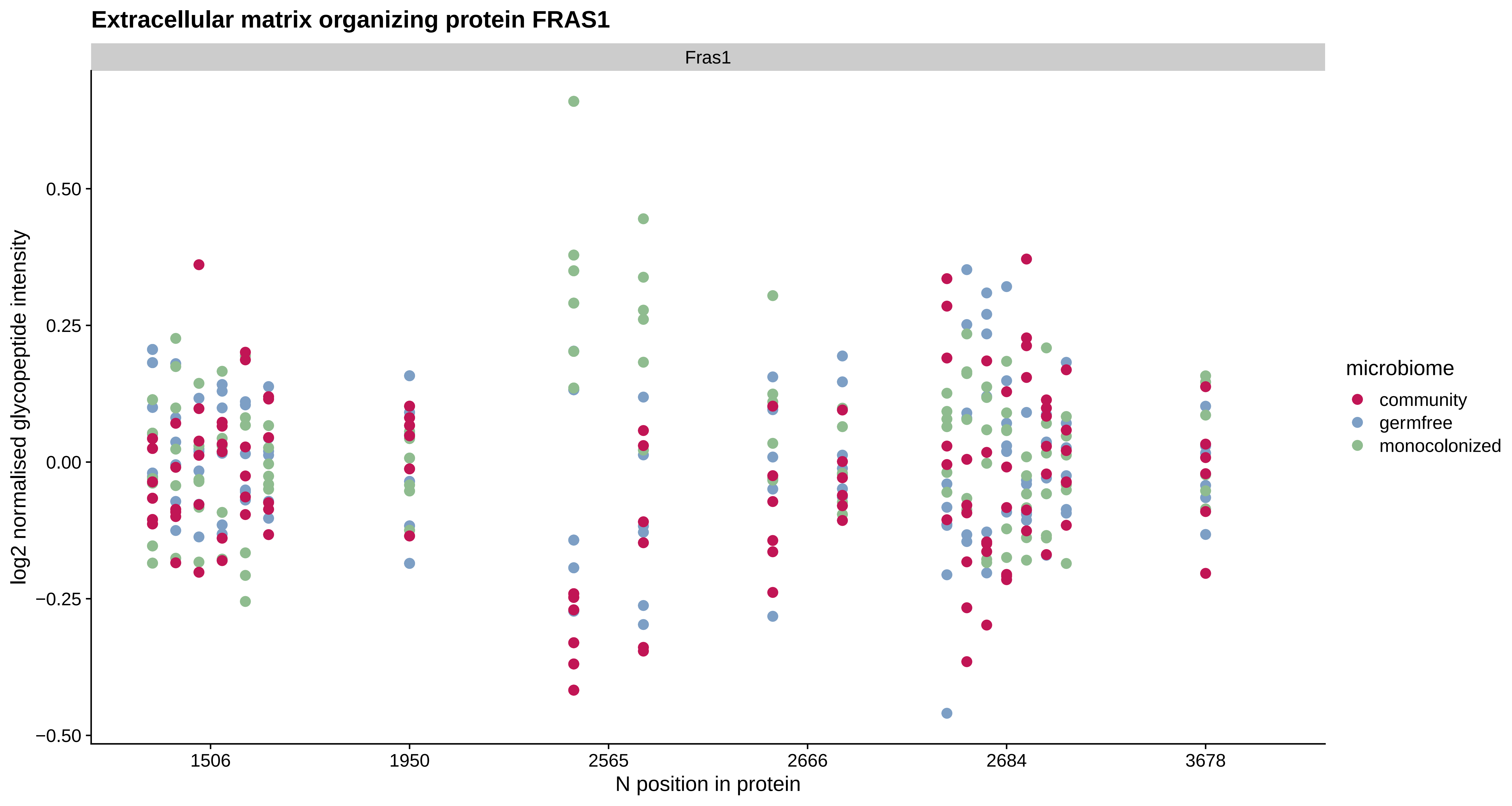
Extracellular matrix organizing protein FRAS1
Fras1
log2 normalised glycopeptide intensity
0.50
0.25
0.00
−0.25
−0.50
1506
1950
2565
2666
2684
3678
N position in protein
microbiome
community
germfree
monocolonized

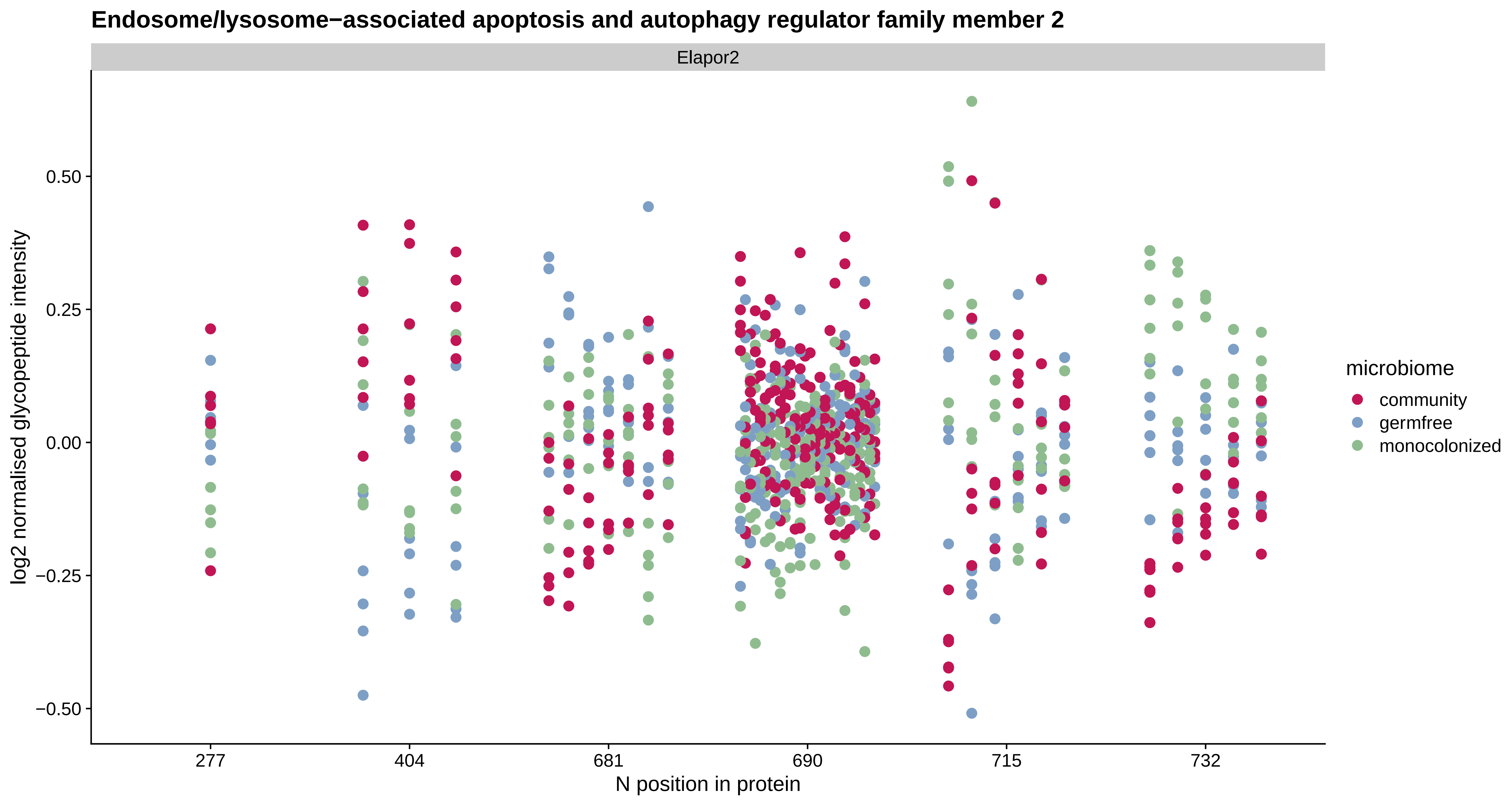

Endosome/lysosome−associated apoptosis and autophagy regulator family member 2
Elapor2
log2 normalised glycopeptide intensity
0.50
0.25
0.00
−0.25
−0.50
277
404
681
690
715
732
N position in protein
microbiome
community
germfree
monocolonized

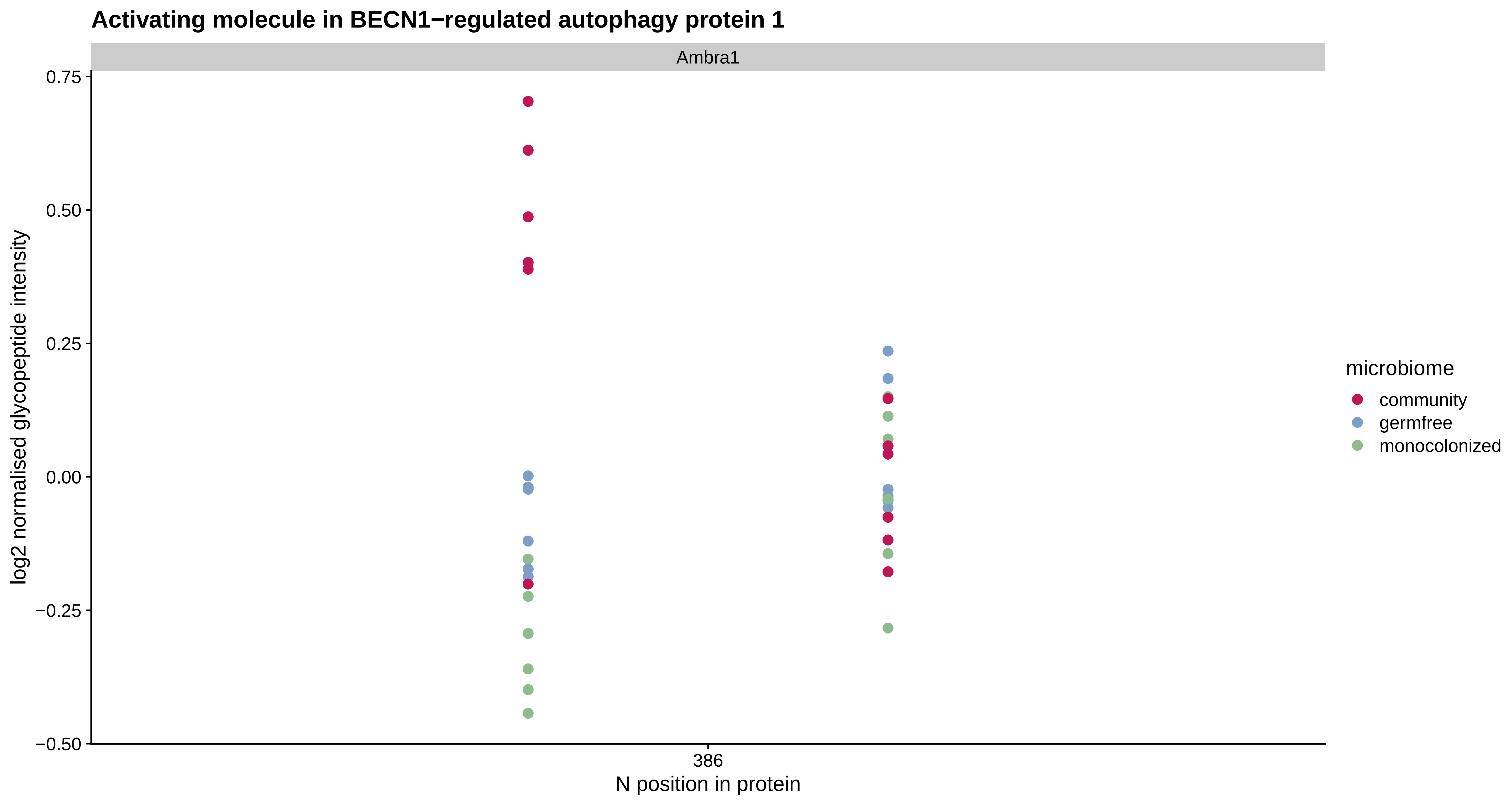
Activating molecule in BECN1−regulated autophagy protein 1
Ambra1
0.75
0.50
0.25
0.00
−0.25
−0.50
log2 normalised glycopeptide intensity
386
N position in protein
microbiome
community
germfree
monocolonized

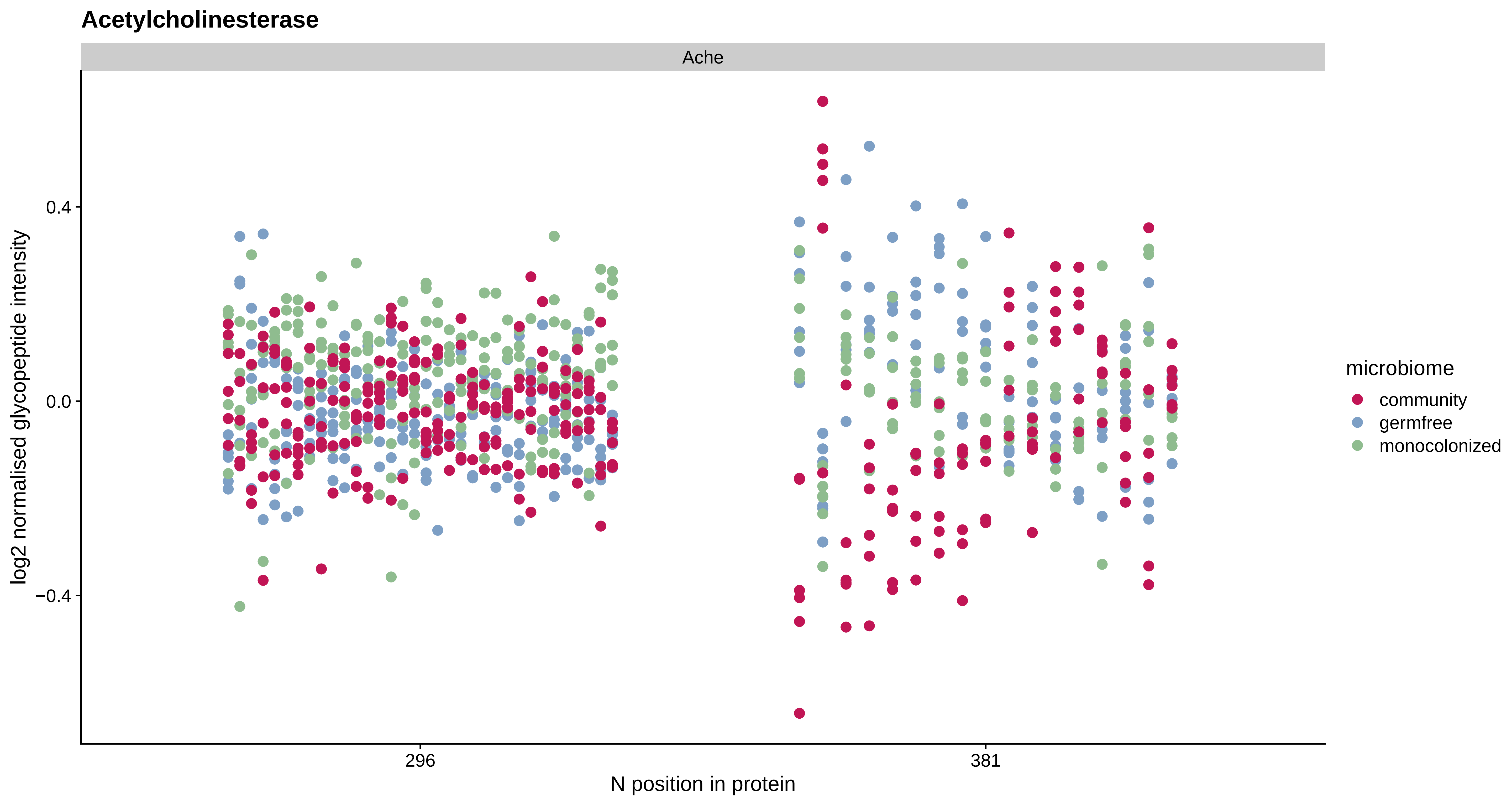
Acetylcholinesterase
Ache
log2 normalised glycopeptide intensity
0.4
0.0
−0.4
296
381
N position in protein
microbiome
community
germfree
monocolonized

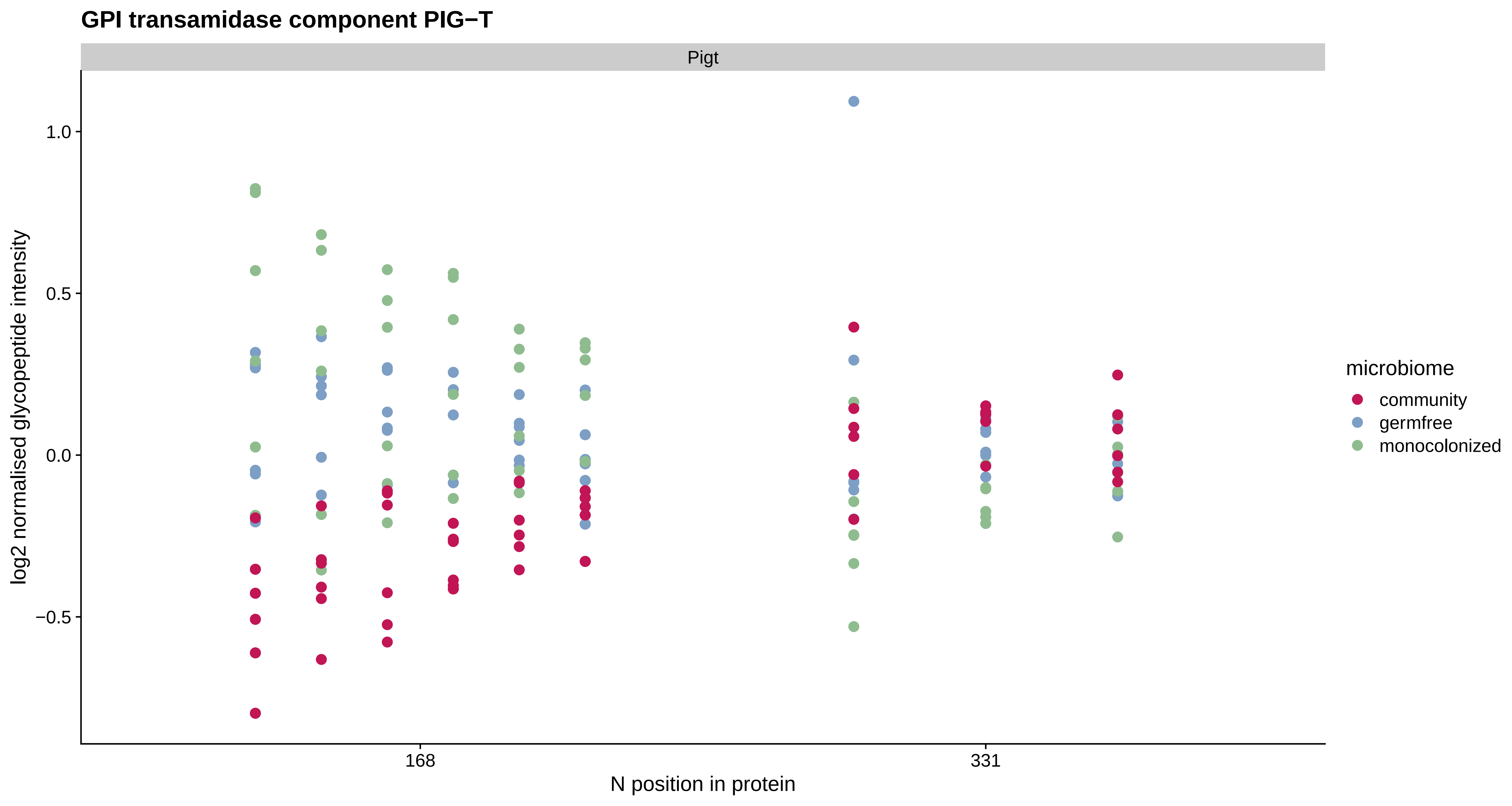

GPI transamidase component PIG−T
Pigt
log2 normalised glycopeptide intensity
1.0
0.5
0.0
−0.5
168
331
N position in protein
microbiome
community
germfree
monocolonized

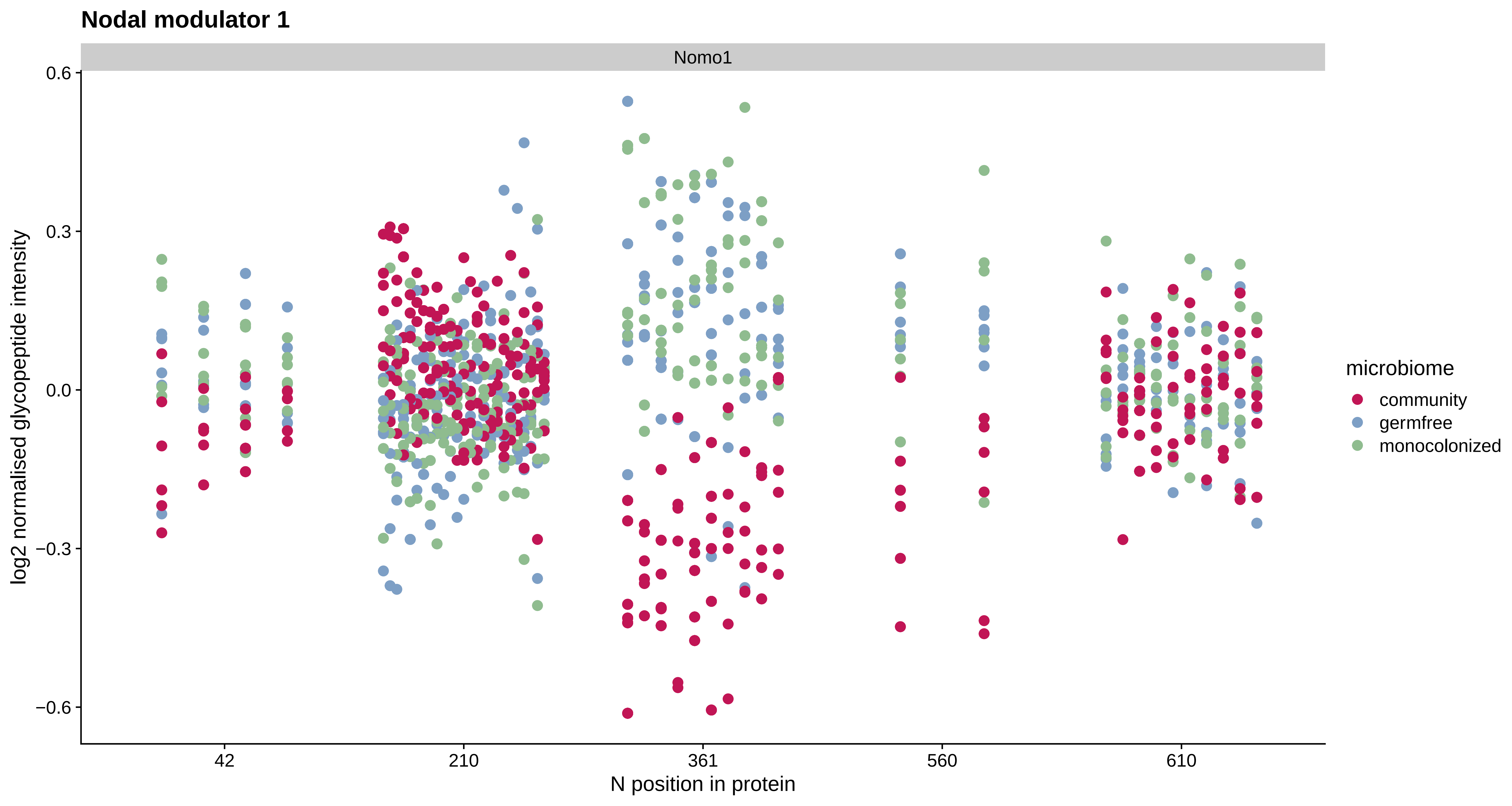
Nodal modulator 1
Nomo1
0.6
0.3
0.0
−0.3
−0.6
log2 normalised glycopeptide intensity
42
210
361
560
610
N position in protein
microbiome
community
germfree
monocolonized

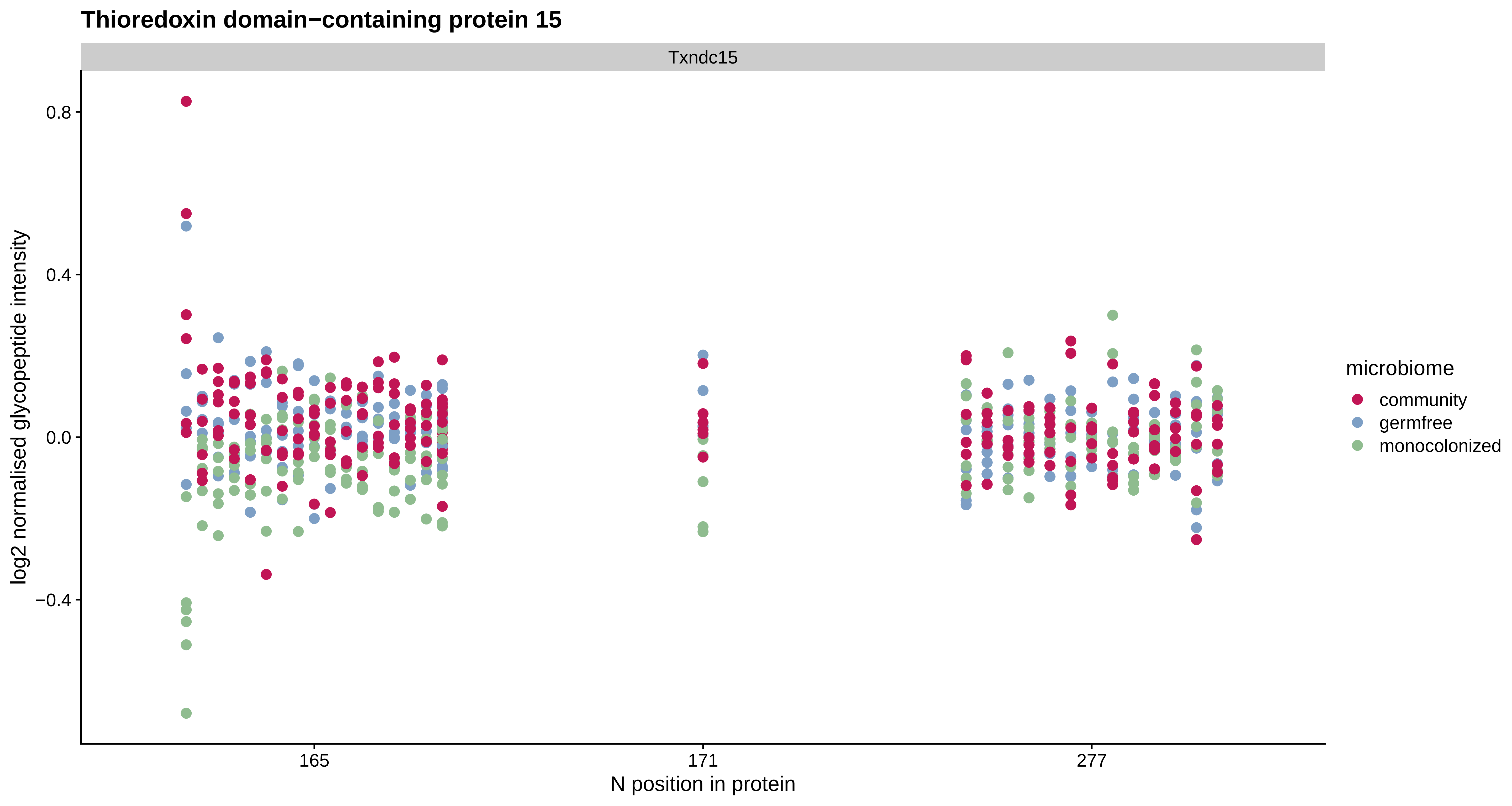
Thioredoxin domain−containing protein 15
Txndc15
0.8
0.4
0.0
−0.4
log2 normalised glycopeptide intensity
165
171
277
N position in protein
microbiome
community
germfree
monocolonized

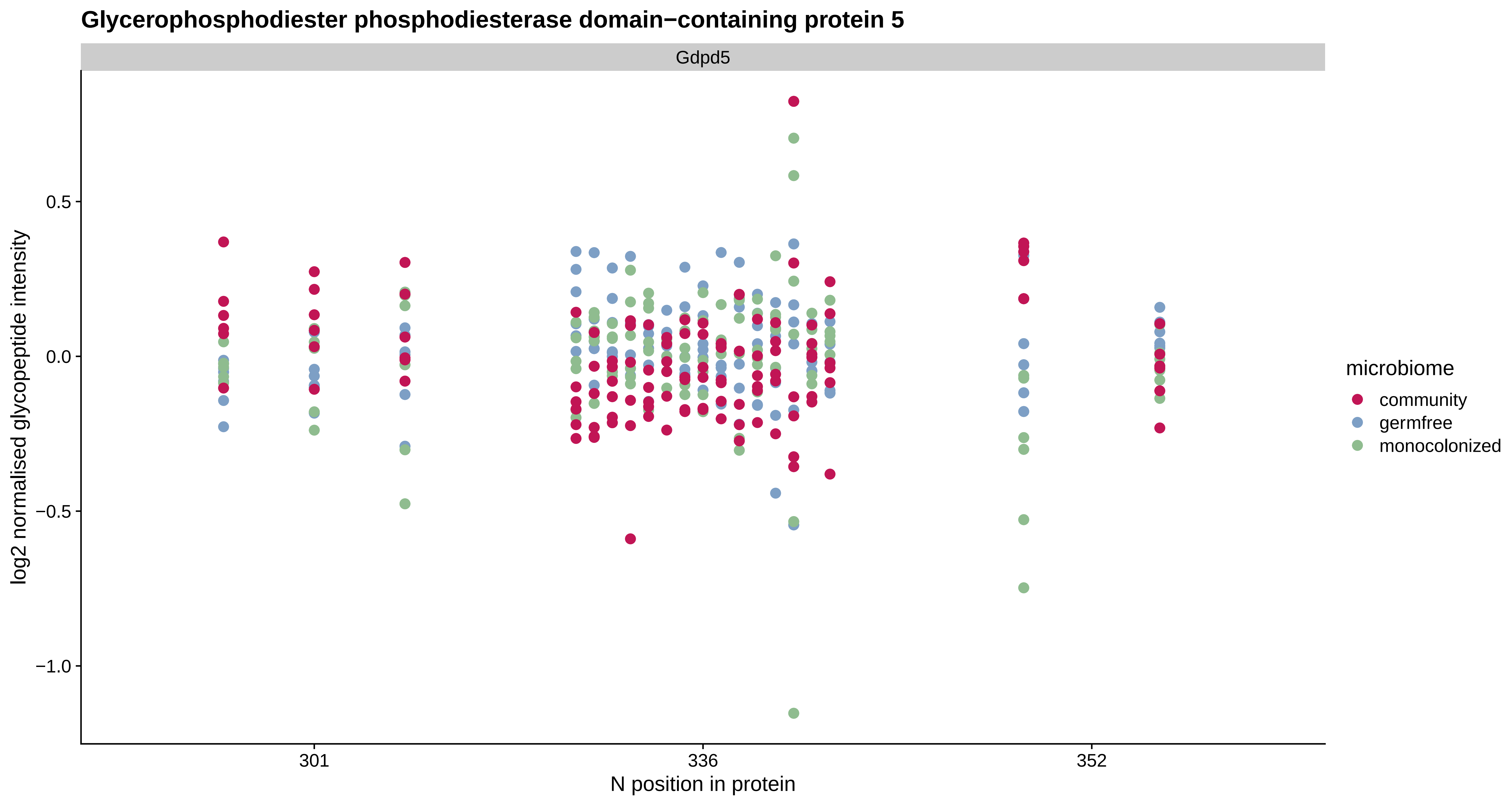

Glycerophosphodiester phosphodiesterase domain−containing protein 5
Gdpd5
log2 normalised glycopeptide intensity
0.5
0.0
−0.5
−1.0
301
336
352
N position in protein
microbiome
community
germfree
monocolonized

#### Receptor−type tyrosine−protein phosphatase F

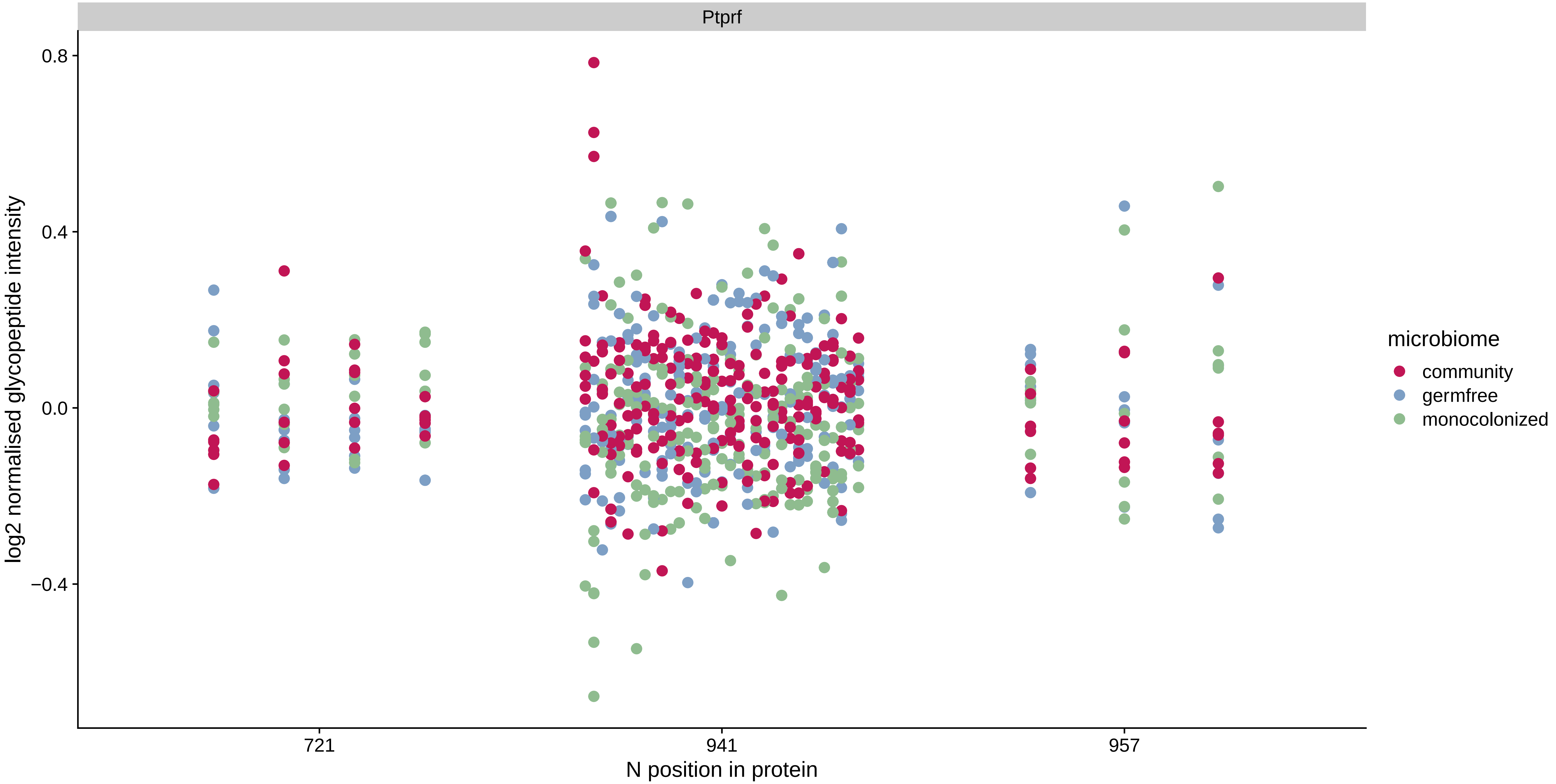

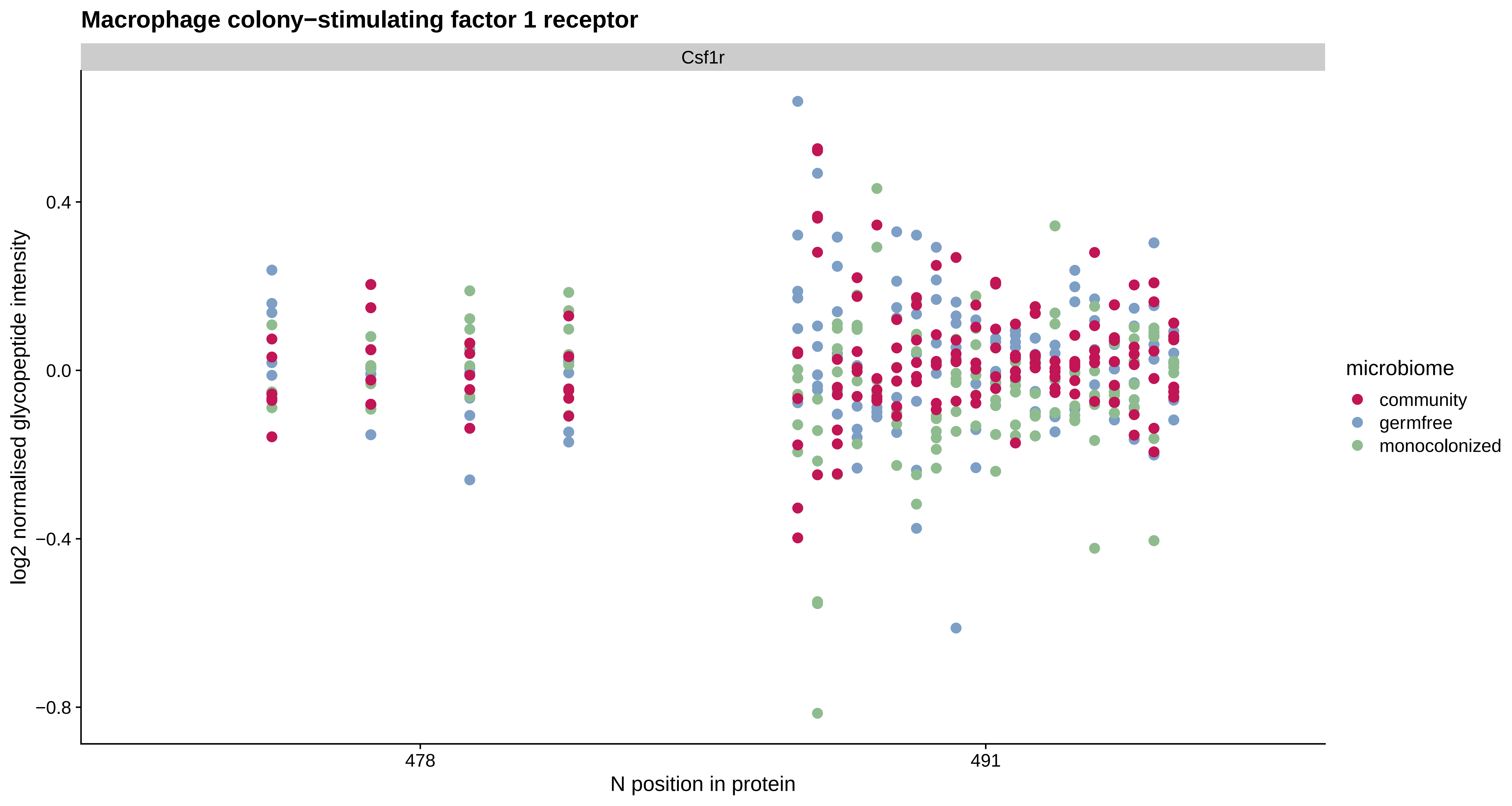
Macrophage colony−stimulating factor 1 receptor
Csf1r
log2 normalised glycopeptide intensity
0.4
0.0
−0.4
−0.8
478
491
N position in protein
microbiome
community
germfree
monocolonized

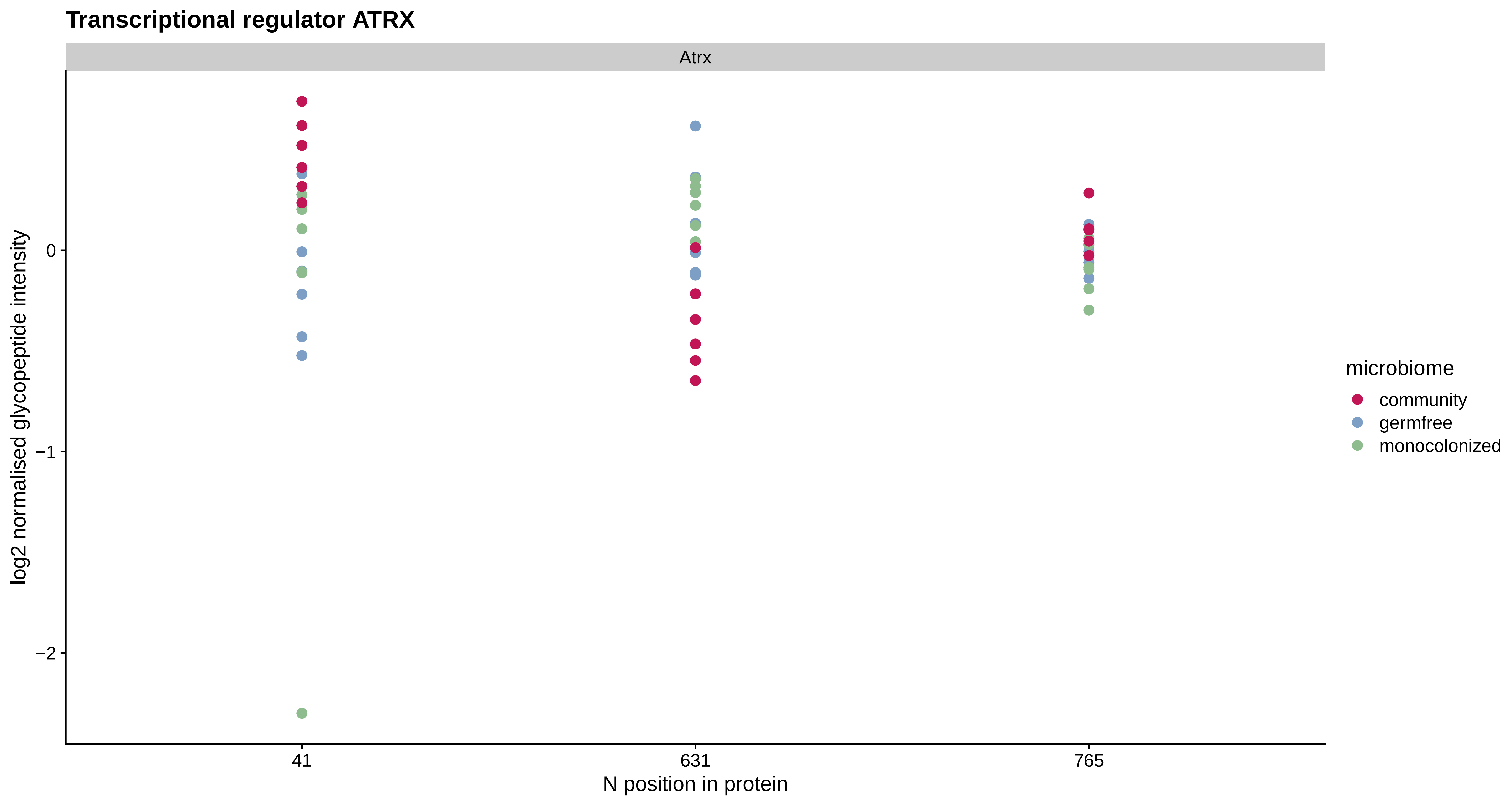

Transcriptional regulator ATRX
Atrx
log2 normalised glycopeptide intensity
0
−1
−2
41
631
765
N position in protein
microbiome
community
germfree
monocolonized

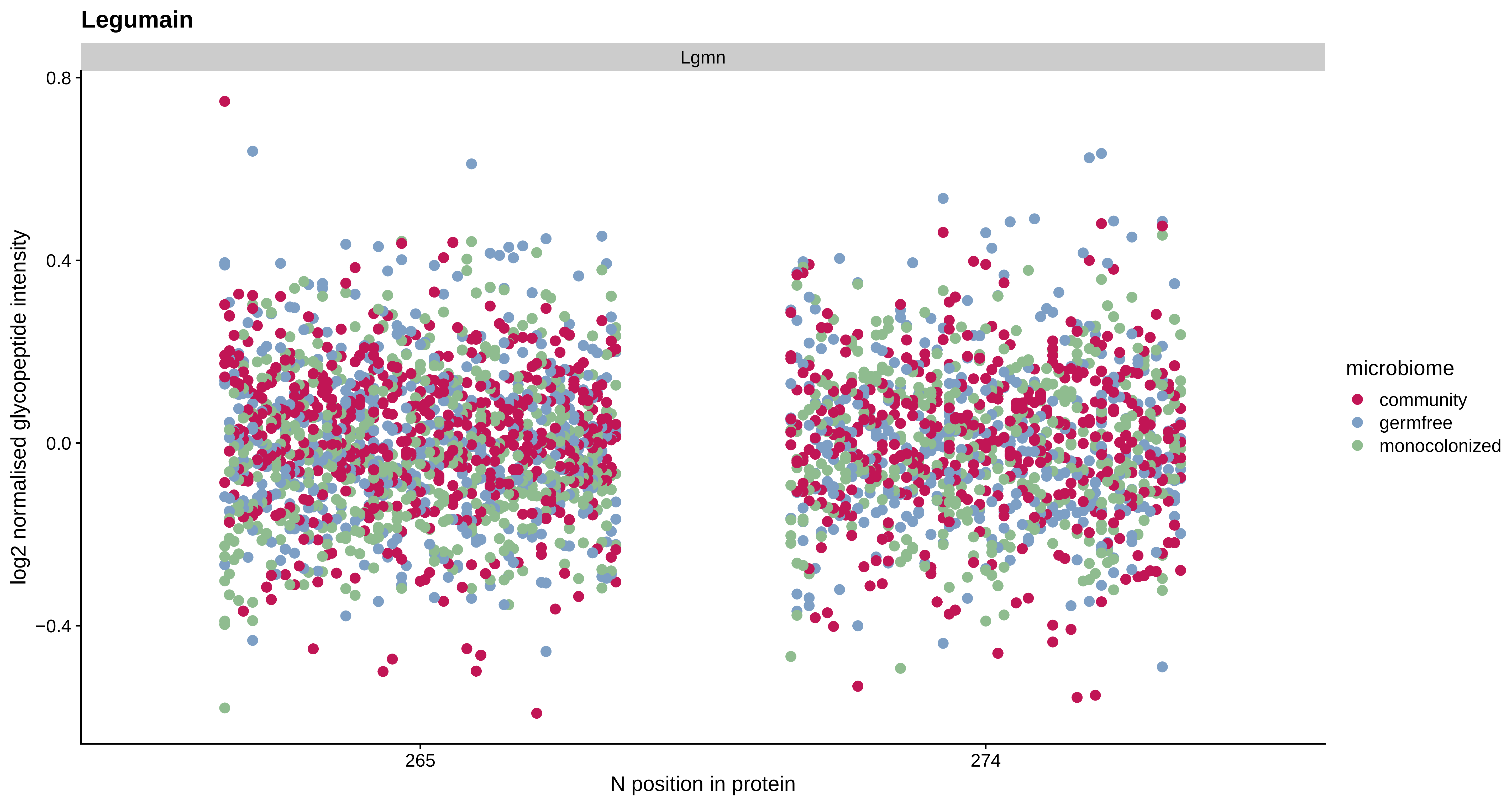

Legumain
Lgmn
0.8
0.4
0.0
−0.4
log2 normalised glycopeptide intensity
265
274
N position in protein
microbiome
community
germfree
monocolonized

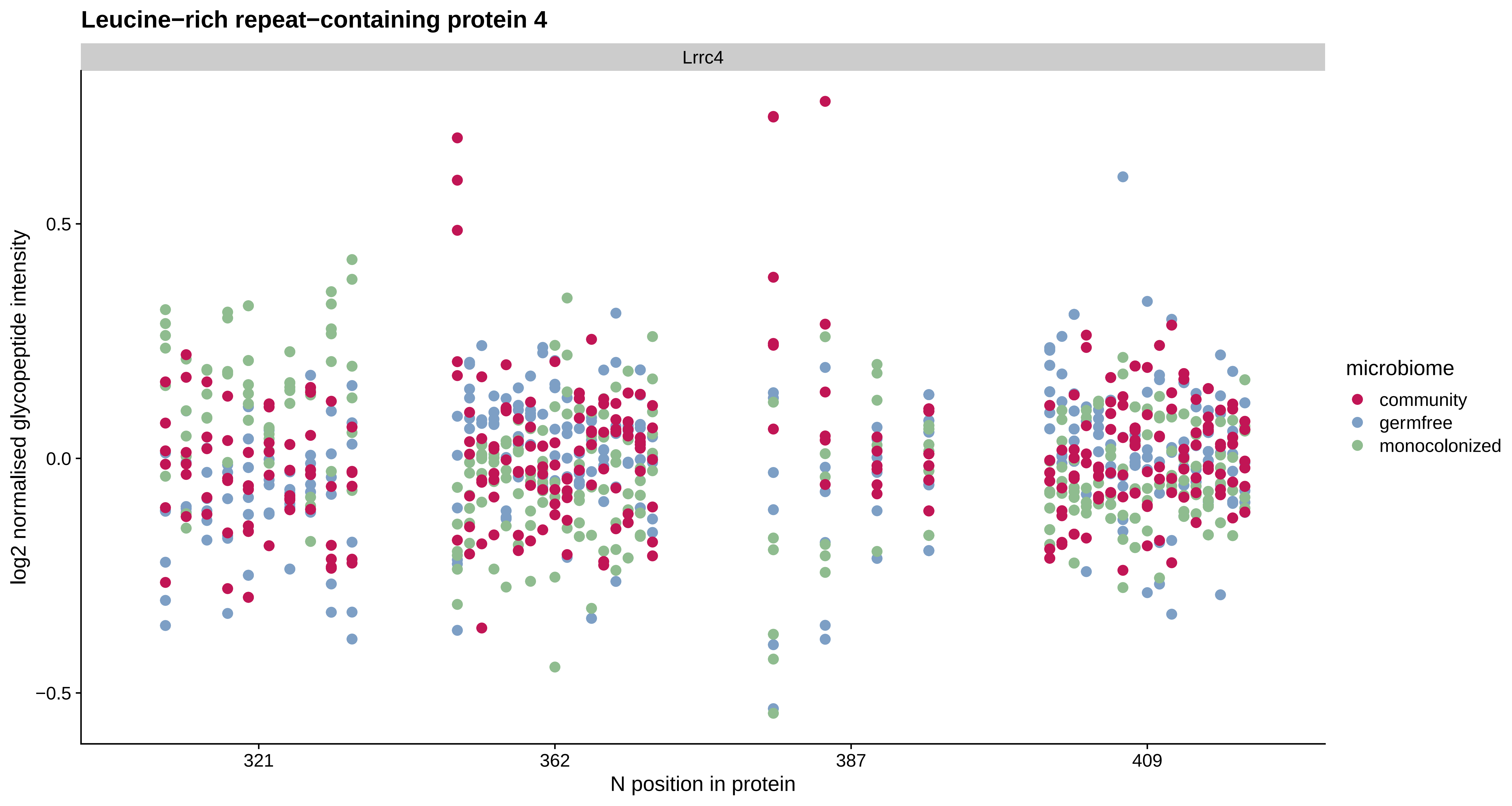

Leucine−rich repeat−containing protein 4
Lrrc4
log2 normalised glycopeptide intensity
0.5
0.0
−0.5
321
362
387
409
N position in protein
microbiome
community
germfree
monocolonized

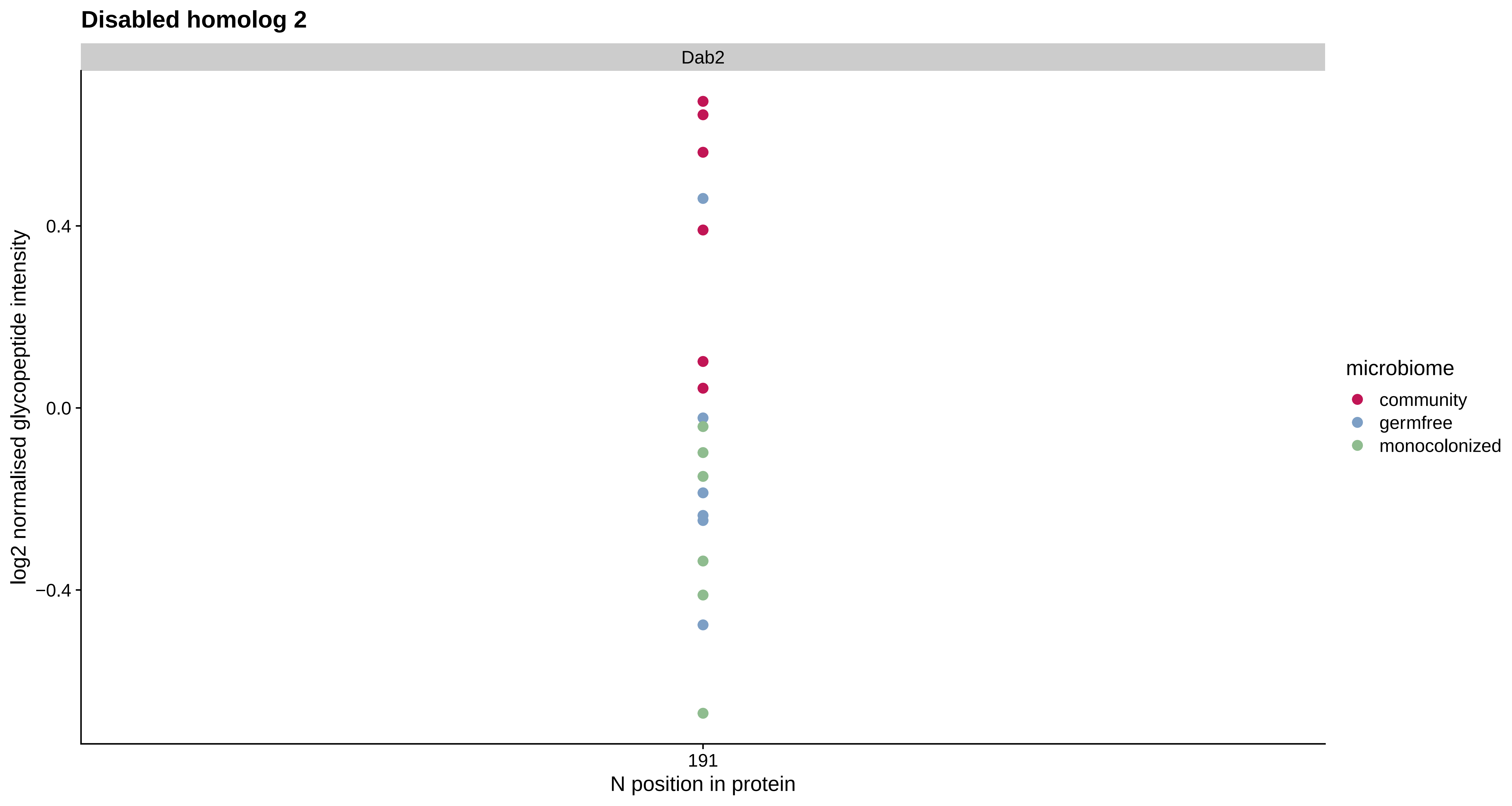

Disabled homolog 2
Dab2
0.4
0.0
−0.4
log2 normalised glycopeptide intensity
191
N position in protein
microbiome
community
germfree
monocolonized

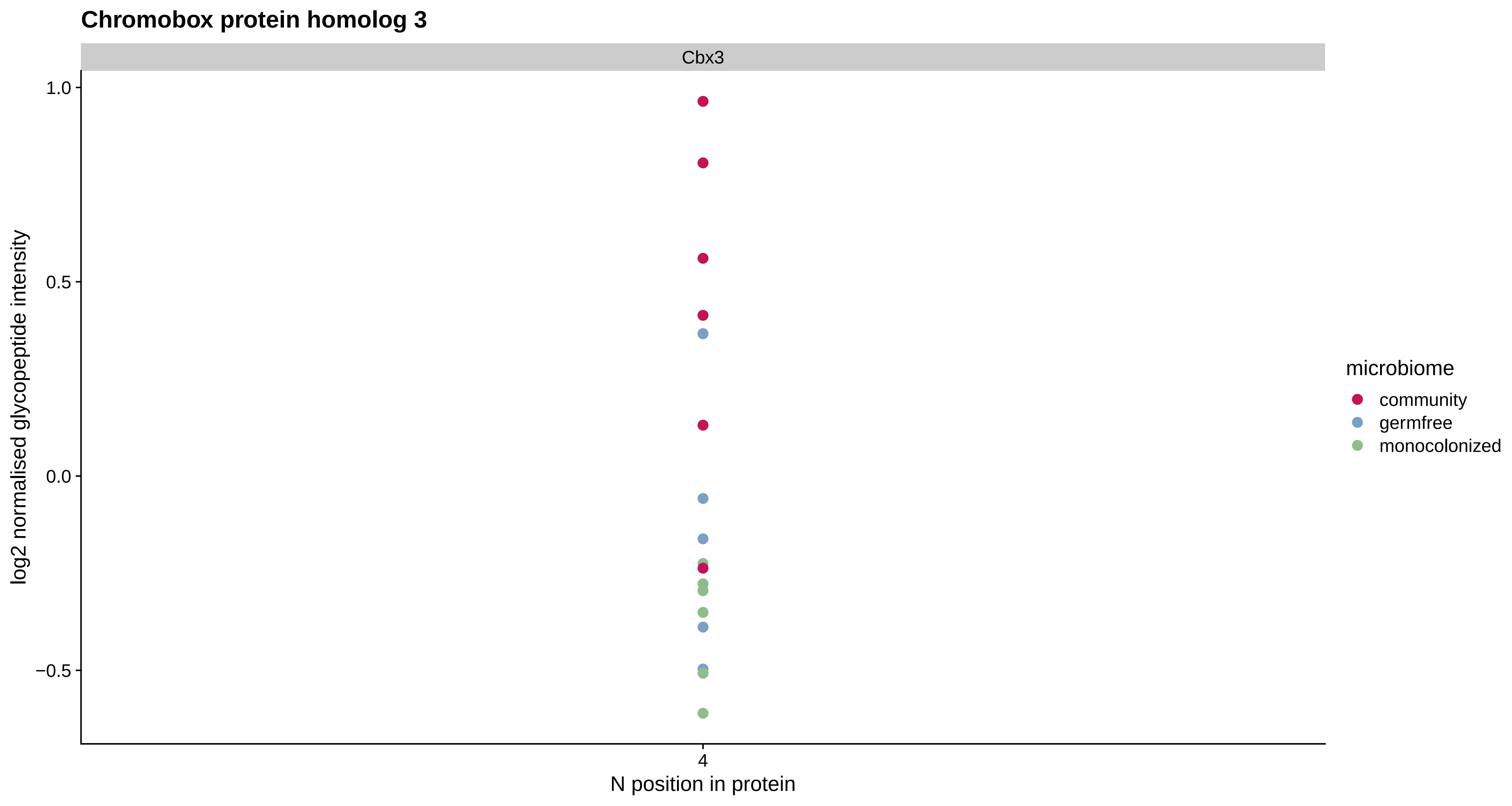
Chromobox protein homolog 3
Cbx3
1.0
0.5
0.0
−0.5
log2 normalised glycopeptide intensity
4
N position in protein
microbiome
community
germfree
monocolonized

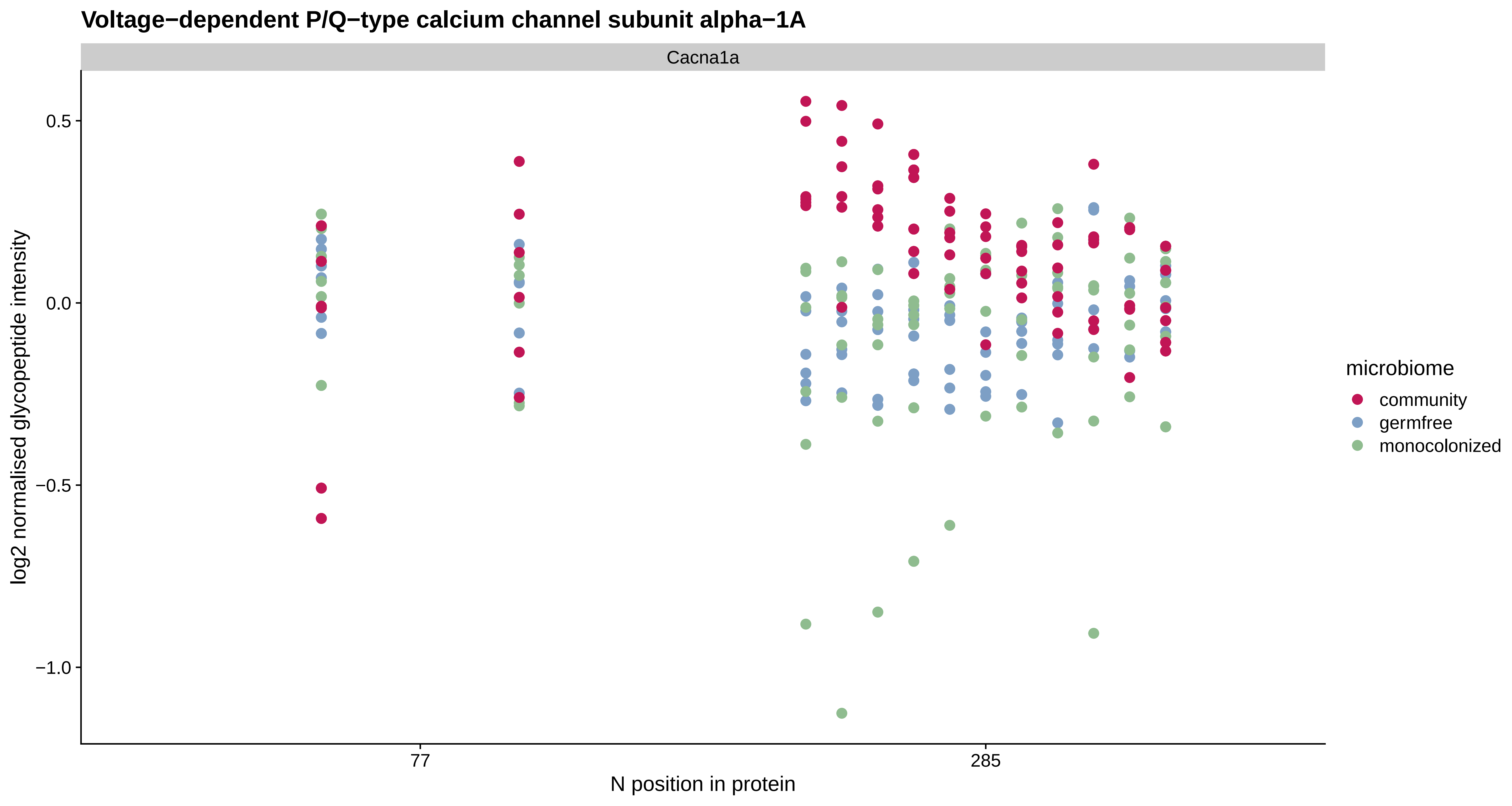

Voltage−dependent P/Q−type calcium channel subunit alpha−1A
Cacna1a
log2 normalised glycopeptide intensity
0.5
0.0
−0.5
−1.0
77
285
N position in protein
microbiome
community
germfree
monocolonized

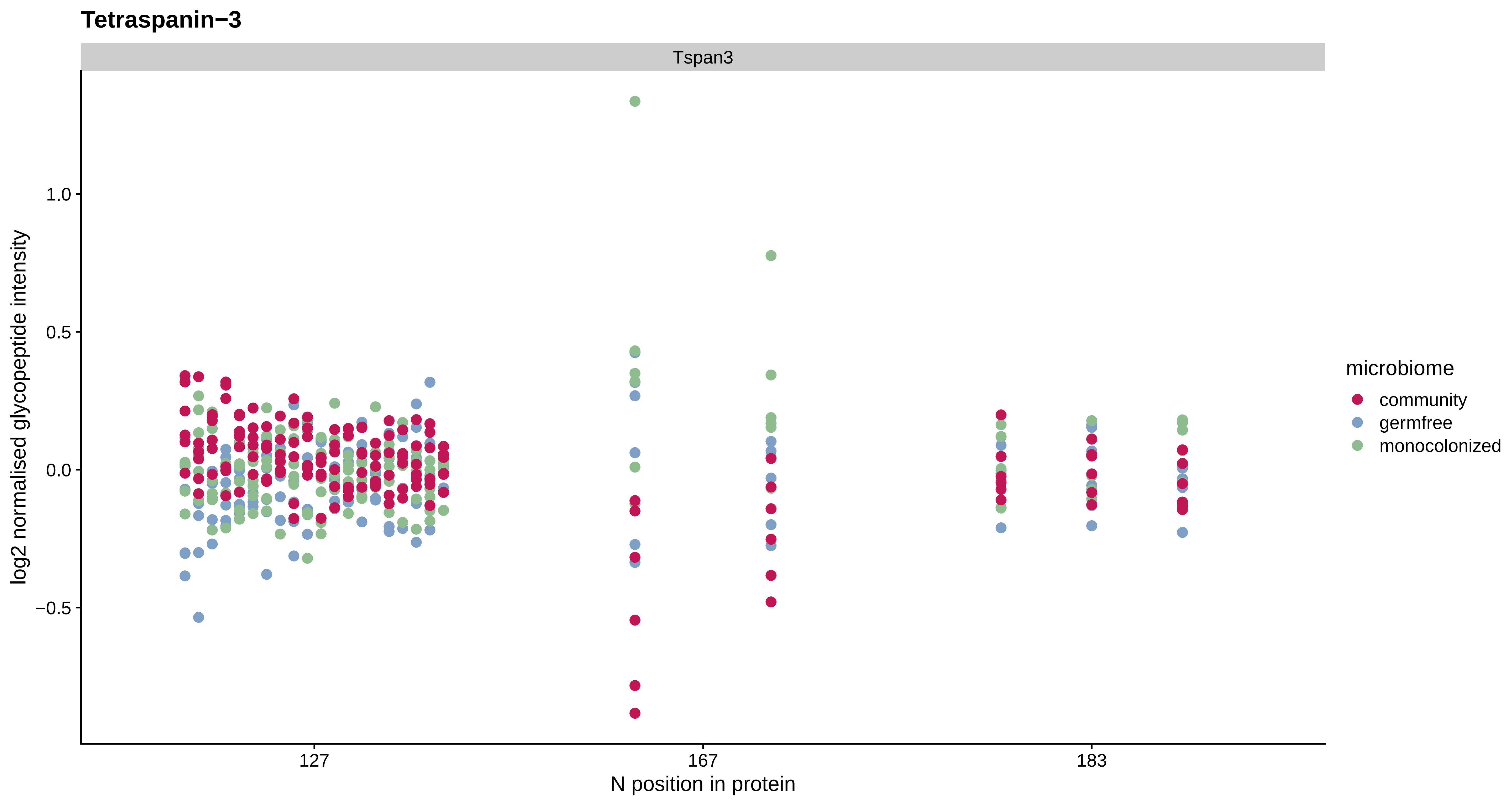

Tetraspanin−3
Tspan3
1.0
0.5
0.0
−0.5
log2 normalised glycopeptide intensity
127
167
183
N position in protein
microbiome
community
germfree
monocolonized

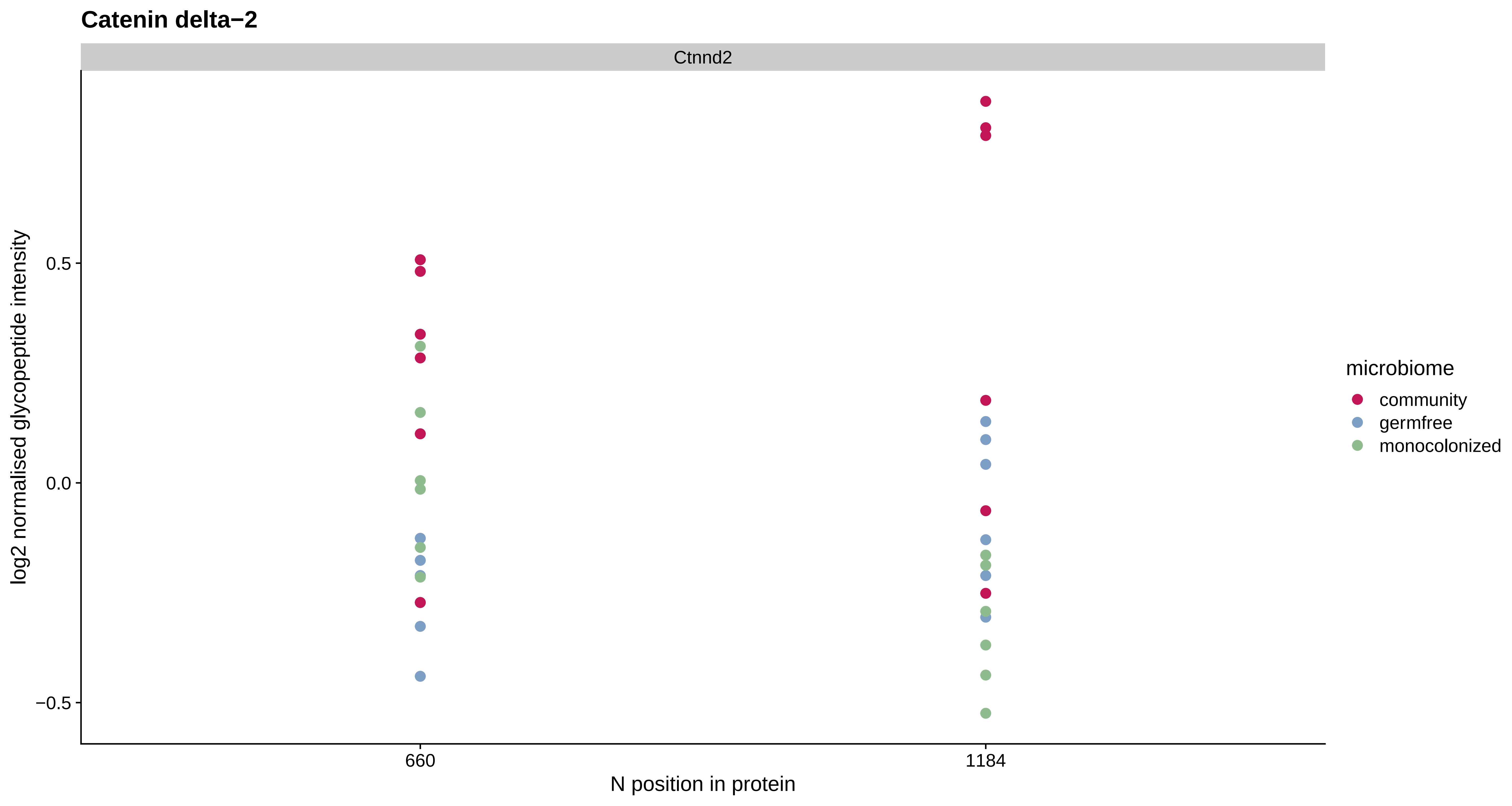
Catenin delta−2
Ctnnd2
log2 normalised glycopeptide intensity
0.5
0.0
−0.5
660
1184
N position in protein
microbiome
community
germfree
monocolonized

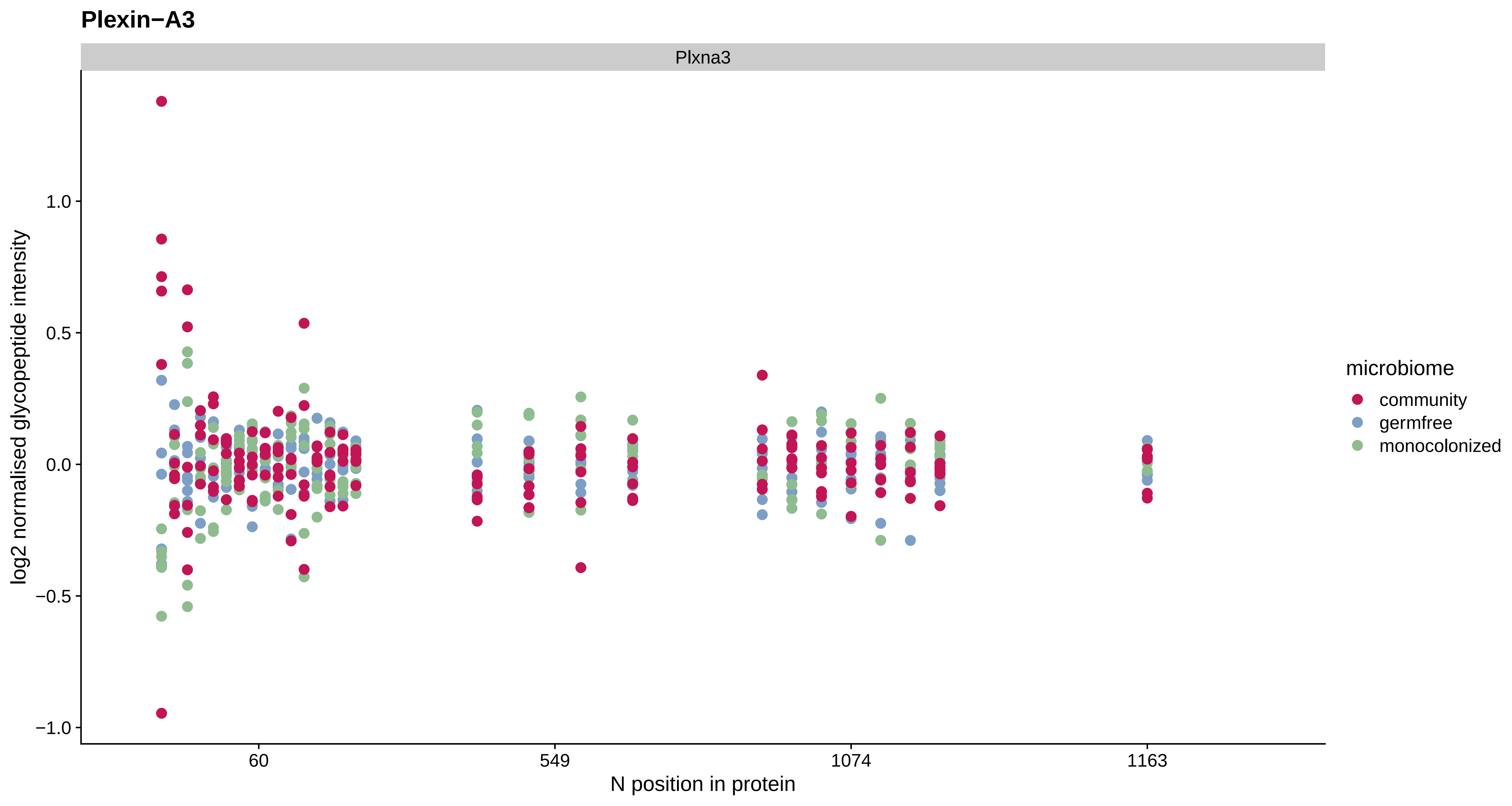

Plexin−A3
Plxna3
log2 normalised glycopeptide intensity
1.0
0.5
0.0
−0.5
−1.0
60
549
1074
1163
N position in protein
microbiome
community
germfree
monocolonized

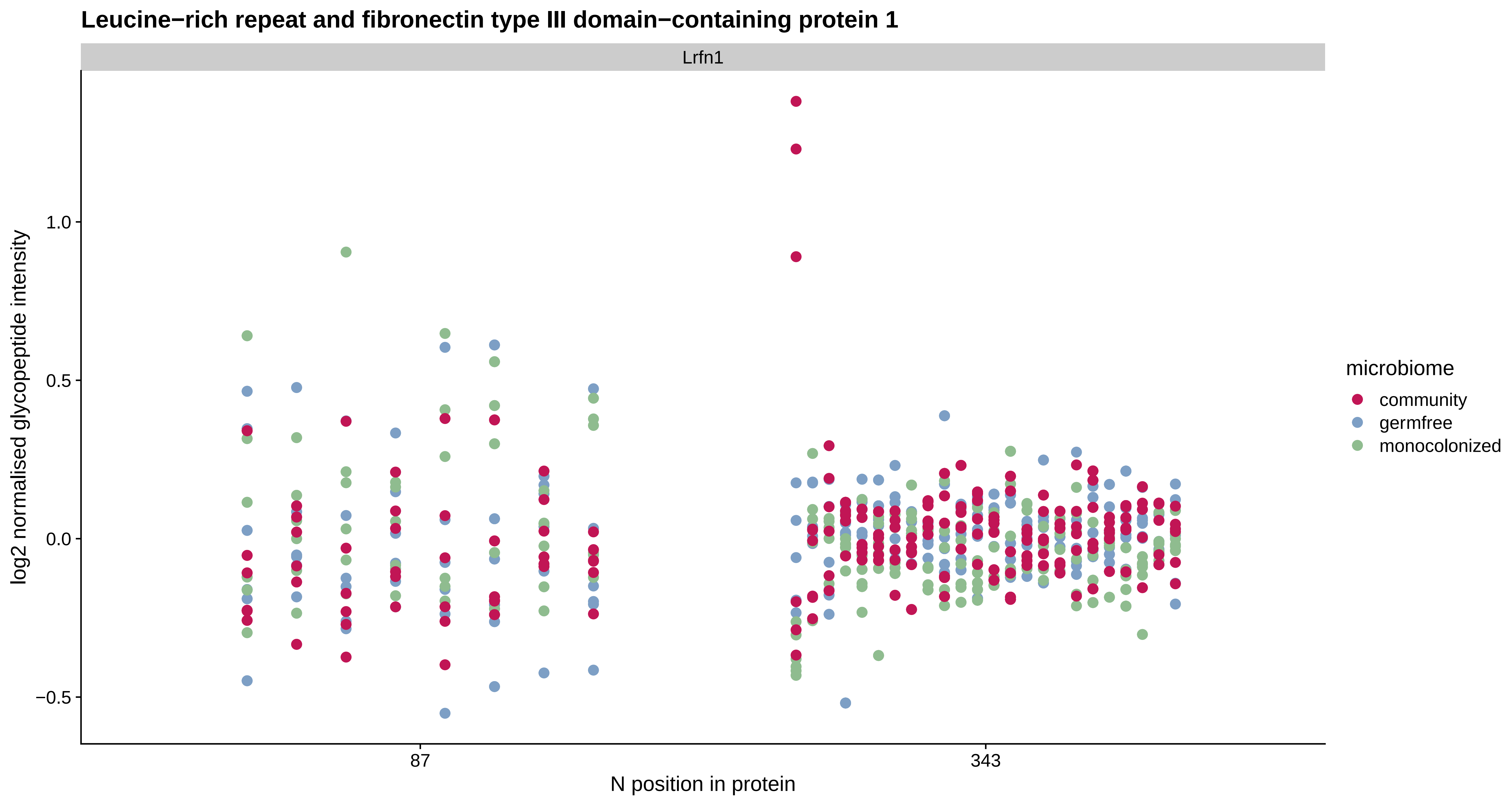

Leucine−rich repeat and fibronectin type III domain−containing protein 1
Lrfn1
log2 normalised glycopeptide intensity
1.0
0.5
0.0
−0.5
87
343
N position in protein
microbiome
community
germfree
monocolonized

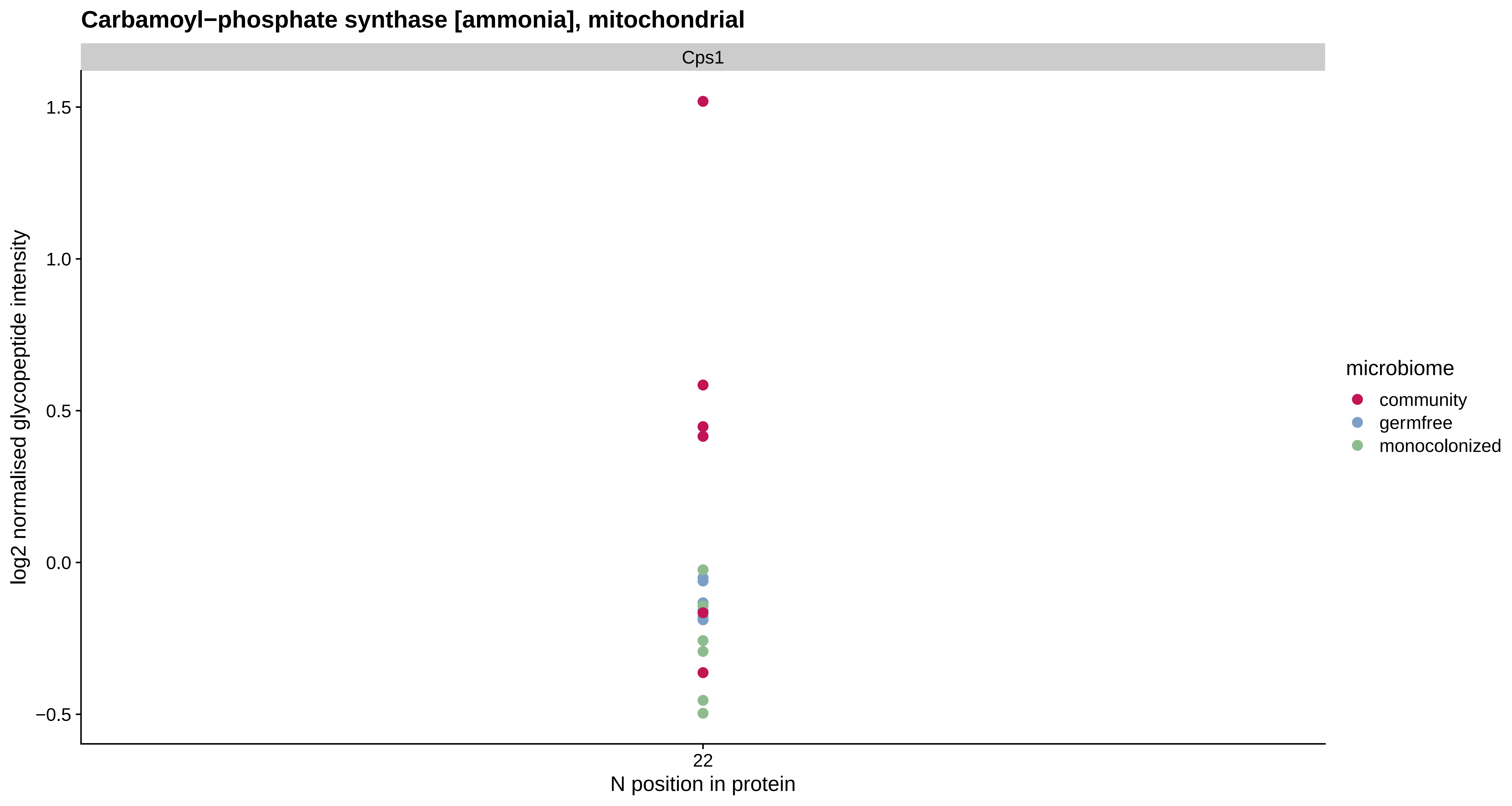
Carbamoyl−phosphate synthase [ammonia], mitochondrial
Cps1
log2 normalised glycopeptide intensity
1.5
1.0
0.5
0.0
−0.5
22
N position in protein
microbiome
community
germfree
monocolonized
